# Whole cryo-FIB lamella acquisition at high magnification with rectangular beam montage cryo-electron tomography

**DOI:** 10.64898/2026.09.27.753993

**Authors:** Hamish G. Brown, Sepideh Valimehr, Denver Styczynski, Manasi Mudaliyar, Srujan Shetty Kundapura, Matthew D. Johnson, Debnath Ghosal, Eric Hanssen

## Abstract

The visualisation of large biological samples in cryo-electron tomography requires both few-nanometre resolution and a wide, ideally >10 µm, field of view. For a single image, increasing magnification trades increased resolution for a smaller field of view. Montaging, where high-magnification images are stitched together to increase field of view, addresses the trade-off but is dose-inefficient with round beams that are standard in transmission electron microscopy (TEM), since this illumination shape does not tessellate. Our workflow greatly simplifies montaging schemes for dose-sensitive specimens, allowing large montages of any shape – from individual cells or other features of interest up to whole lamellae – to be collected at high resolution without compromising electron dose efficiency. We demonstrate the workflow by acquiring montage tomograms of whole yeast (*Saccharomyces cerevisiae)* cells in cryo-FIB lamellae and of human neurons.

## Main

In cryogenic electron tomography (cryo-ET) a three-dimensional volume is reconstructed from a tilt series of images of a biological sample frozen in vitreous ice [1, 2]. The technique allows for the study of biological systems, such as bacteria, eukaryotic cells and even plant and animal tissue, vitrified in a near-native state. Using sub-tomogram averaging, where a 3D volume is reconstructed from multiple instances of a single protein across tilt series, biomolecule structures can be determined at subnanometre resolution. For a single image acquisition in microscopy there is a trade-off between pixel size, which is determined by the image magnification and sets a limit on the maximum attainable resolution, and field of view, which is the product of the number of pixels in the camera and the chosen pixel size [3]. The field of large-area in situ structural and cellular biology is rapidly expanding, driven by the increasing use of carefully prepared cryo-FIB lamellae for high-resolution imaging [4]. Montaging – computationally stitching together a regularly spaced grid of images – enables substantially larger fields of view to be captured at a given pixel size [5]. This approach therefore allows researchers to make more effective use of the limited and valuable real estate available within a cryo-FIB lamella, while retaining the resolution required for detailed structural analysis. A complication of montaging in cryo-electron microscopy, where the specimen is dose-sensitive [6, 7], is that the beam is usually round and electron cameras are square or rectangular. If the round illumination is spread to cover the square camera, then areas adjacent to the field of view will be damaged by the round lobes of the beam but never captured. This deteriorates the quality of the acquisitions in subsequent montage tiles [8]. Previous schemes for montaging in cryo-ET [9, 10] have used standard round beams condensed to be smaller than the size of the electron camera, sometimes with a hexagonal grid for more efficient tiling of the round illumination. To avoid dose hot-spots on the grid the montage array is shifted with each tilt of the sample. If the illumination is made to match the dimensions of the camera – an innovation previously demonstrated with condenser apertures that match our (square) Thermo Fisher Falcon IV and (rectangular) Gatan K3 cameras [11, 12] – then cryo-montage schemes with simpler tiling and with illumination that more efficiently fills the camera become possible [13]. In this manuscript we describe an approach for using these novel electron beam shapes for montage tomography. We discuss the implementation of this approach through scripting for the open-source SerialEM transmission electron microscopy (TEM) automation software [5], and demonstrate the approach by acquiring high-resolution montage tomography datasets of a whole yeast cell and extended human neurons.

The workflow runs on the microscope using just three scripts for acquisition, included both in the supplementary material and on GitHub^1^, developed for the open-source TEM automation software SerialEM [5], which can operate on either the microscope control PC or on the camera control PC. Once the grids are loaded into the TEM column, the microscope is aligned for high-resolution data acquisition. The DetermineOverlapFraction.txt script is then run on an area of empty or broken grid to get a reference beam shape at the magnification that will be used for data acquisition. An interactive utility allows the user to set montage beam overlap. Next, a low magnification atlas is recorded of the entire grid to assist with navigation across the sample and to identify points of interest for tomography acquisition. Areas of interest for montage acquisition are outlined as polygons in the navigator (shown diagrammatically in Fig. 1(a)) and, using the *Acquire at Items* dialogue in SerialEM, images are acquired at an intermediate magnification (5000× magnification, or a pixel size of around 25 Å) at each of the points for stage realignment during the acquisition run. Using *Acquire at Items*, the [setup_polygon_montage.txt](https://github.com/HamishGBrown/Square_beam_montaging/blob/main/SerialEMScripts/SetupPolygonMontage.txt) script is run after each intermediate magnification image acquisition to design the montage tiling that will cover the user-drawn polygon area, shown in Fig. 1(b). A text file is written that lists the number of tilts in the series, tilt angles and image shifts (in camera pixels) for each tile to be acquired, see Fig. 1(c). This output is used both by the acquisition script and during data processing as an initial estimate of image positioning in the montage. A screenshot of the SerialEM interface, with hand-drawn polygon and resulting image acquisition scheme for the first tilt in the series is shown in Fig. 1(d). As overlapping tiles are exposed more than once, the tile positions are shifted laterally at each tilt so that the accumulated dose is spread evenly across the montage (Fig. 1(e)–(f)). In the example shown, this limited the excess exposure from tile overlap to roughly 12% above the intended dose. The full tile geometry, defocus-offset and dose calculations are given in Section 3 of the Materials and Methods.

**Figure 1.**
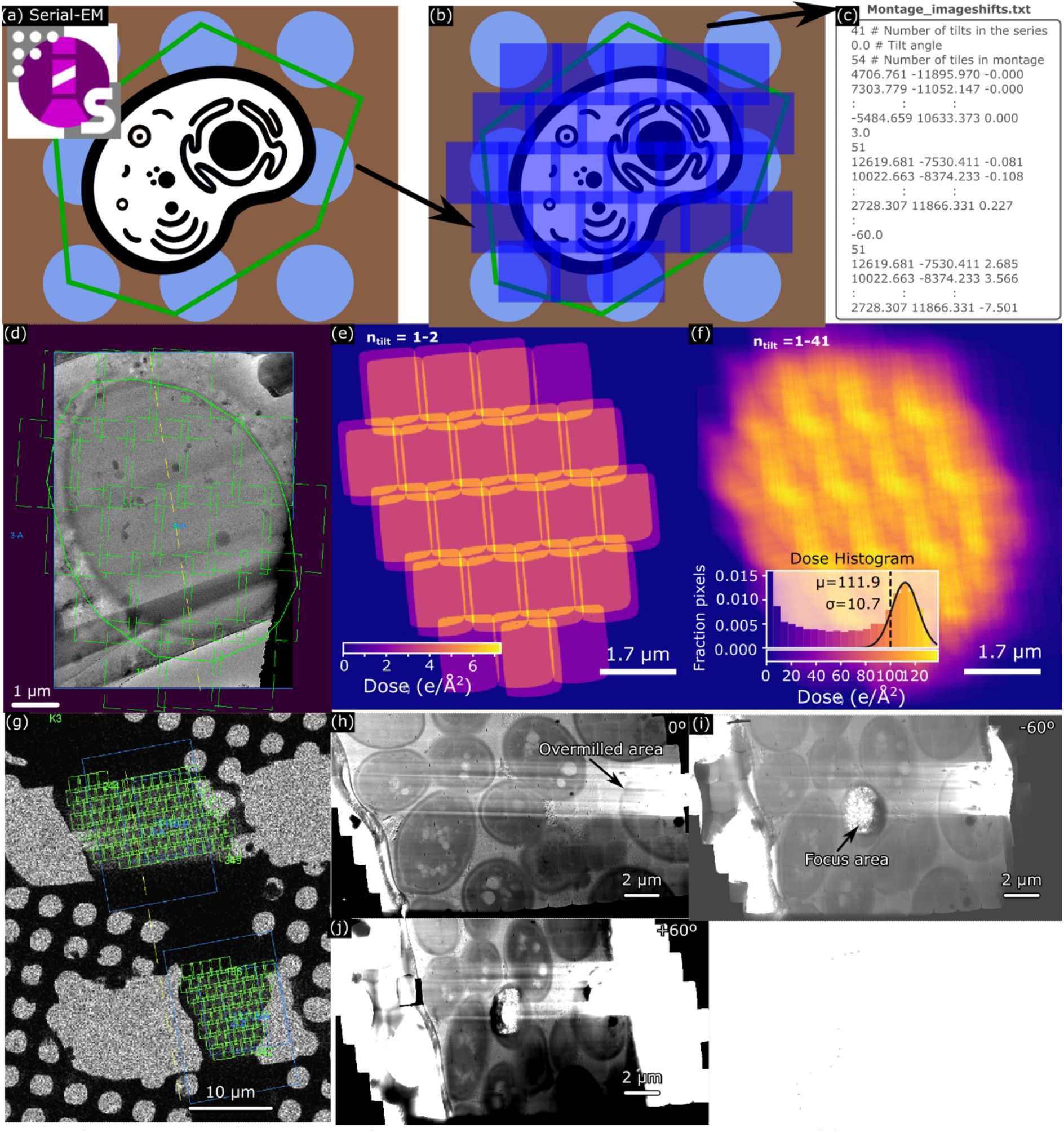
Setup of a montage tilt series in SerialEM and dose accumulation considerations. (a) A region of the grid is selected by a polygon in the SerialEM navigator and (b) a script is run that calculates the montage tiles required to adequately cover the desired region. The tilt series with positions of montage tiles is written to a separate file for the acquisition stage as indicated in (c). (d) The grid of acquisitions, shown plotted atop an overview image in an actual screenshot of the SerialEM interface, is irregular to avoid repeated exposure of overlapping areas and (e) the grid of acquisitions is translated for each tilt step such that the accumulated dose is spread more evenly as shown in (f), resulting in a dose overshoot of about 12% (112 e^−^/Å^2^) above the target dose (100 e^−^/Å^2^). The slight rotation between (d) and (e) is a genuine magnification-dependent image rotation induced by the TEM. (g)–(j) Whole-lamella acquisition: (g) SerialEM screenshot showing the acquisition schemes for two adjacent lamellae, and stitched high-magnification (23 kx, 3.4 Å pixel size) montages at (h) 0°, (i) −60° and (j) +60°.

Tilt series of much larger objects such as FIB lamellae are also achievable as shown in Fig. 1(g)–(j). Fig. 1(g) shows an overview image of two FIB lamellae and their montage acquisition schemes (green squares are camera acquisition positions) in a SerialEM screenshot. The final stitched montages are shown at (h) 0°, (i) −60° and (j) +60° stage tilts. Bubbling is visible on the montage at high tilts because, with the acquisition area extending over the whole lamella, there is no region outside it that can be used for automated focusing in SerialEM.

Montage tilt series are acquired by the Acquire_montage.txt script, which drives the microscope to each tilt angle and positions each tile using coma-free beam–image shift. Coma-free image shifts up to 6 µm are routinely used in single particle workflows and are compatible with protein reconstructions at better than 1.5 Å resolution. This implies object diameters up to 12 µm are possible in montage tomography without noticeable resolution impact, even with sub-tomogram averaging [14]. Focusing and alignment are performed once per tilt step. Full acquisition details are given in Section 4 of the Materials and Methods. A montage of raw tiles is shown in Fig. 2(a).

**Figure 2.**
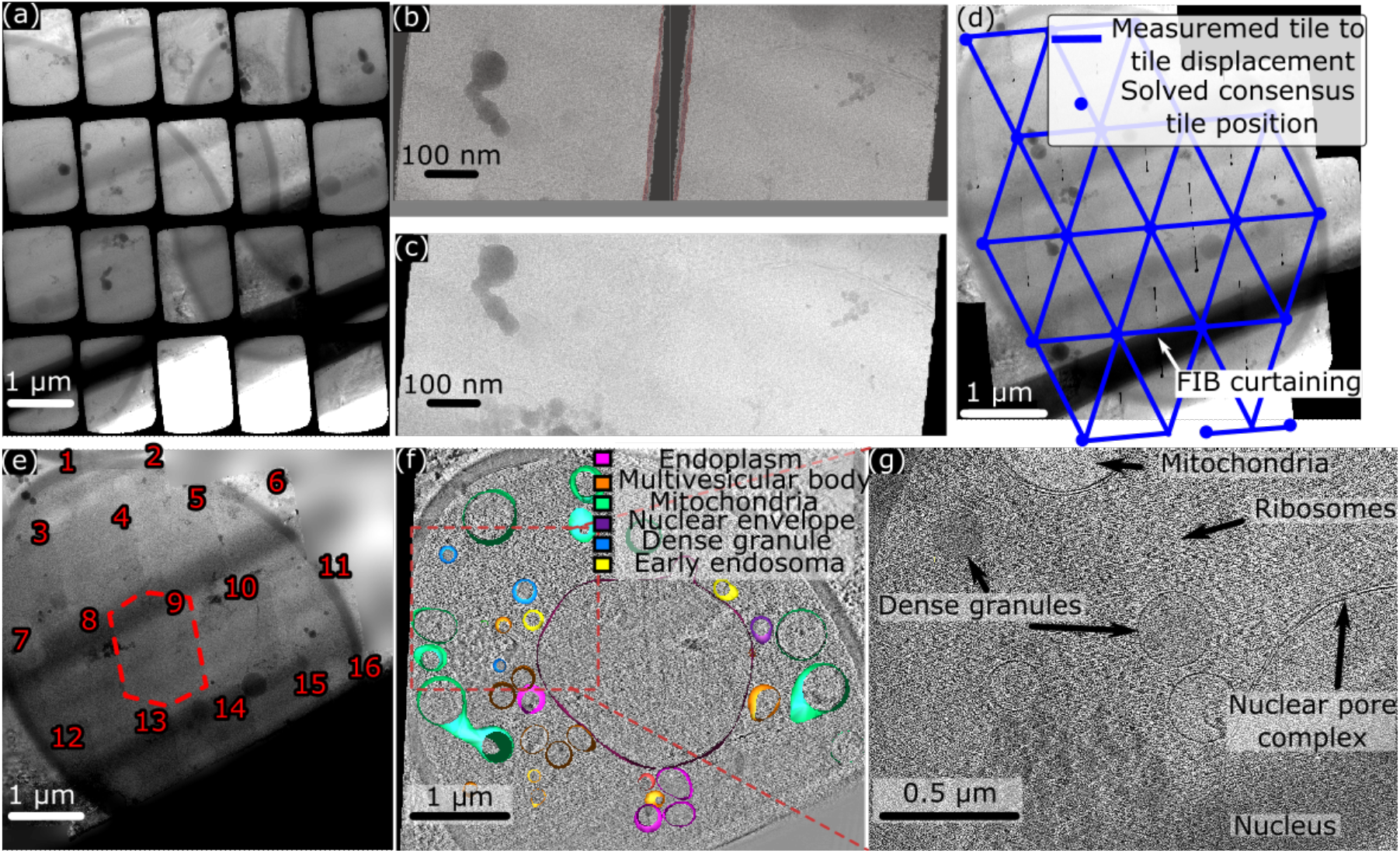
Tomographic reconstruction workflow. (a) A series of images is produced by the microscope and the offsets between images are measured using masked FFTs (masked region overlaid in red) (b), with the stitching of two tiles shown in (c). The consensus stitching is shown in (d), a large region of FIB curtaining is indicated, and regions not covered by the montage are inpainted in (e). A slice of the reconstruction with manual segmentation is shown in (f) and a magnified region showing cellular ultrastructure in (g).

The script recorded dose-fractionated movies which were first aligned using MotionCor2 [15], though single micrographs can also be used. Montages were stitched by the `stitch.py` script, which refined relative positions of overlapping tiles by masked cross-correlation (Fig. 2(b)) – a modification of standard cross-correlation in which only features within the mask are used for alignment – and then solved for a globally consistent set of tile positions from the relative positions by iteratively reweighted least squares (Fig. 2(d)). Sharp tile edges at the extremities of the montage, which create strong artefacts in tomographic reconstruction, were blended into the background using the smoothn [16] algorithm (Fig. 2(e)). Inelastic scattering, which is never fully remediated even when using an energy filter with narrow slit width (typically 10 eV), caused doming of the image tiles as signal was scattered outside the beam. This was corrected by fitting a parametric inelastic scattering model to either all tiles or a representative tile, based on a reference vacuum image, see Section 5 of the Materials and Methods. The doming profile fitted from this procedure was then divided out of the image. The masking, cross-correlation and global-alignment procedures are described in full in Section 6 of the Materials and Methods.

After stitching and blending, the full tilt series of montages can be reconstructed by standard tomography software such as eTomo [17] or AreTomo2 [18]. AreTomo2 was used for the yeast lamella since it uses a fiducial-free alignment method and the lamella lacked gold fiducial markers, whilst eTomo gave excellent results for the human neurons (1.5 nm mean residual for just 10 tracked fiducials across both tomogram regions). We modified the projection-matching algorithm in AreTomo2 to use masked cross-correlations. This was because artefacts such as large contaminants, lamella edges and particularly thick FIB curtaining, which are avoided in standard single-image tomography, are often unavoidable in montage tomography. Since these artefacts often have higher contrast than objects in the lamella, the projection matching algorithm in AreTomo2 tended to have a bias towards aligning to artefacts rather than features in the lamella of interest. The modified version of AreTomo2 is available on its own GitHub^2^ fork and in the supplementary materials. A purpose-built GUI, included in the supplementary information and on GitHub and discussed in detail in Section 7 of the Materials and Methods, was used to design masks for this purpose, either by thresholding or by manually drawing regions to be excluded from 3D alignment and reconstruction.

A three-dimensional rendering of the segmented tomogram and a slice of the full tomogram are shown in Fig. 2(f). A zoomed region of the cell is shown in (g) in which the nucleus and nuclear membrane, early endosome, dense granules, a mitochondrion and ribosomes are visible in the section.

The workflow is efficient for imaging very irregular cells such as human neurons grown on TEM grids. Shown in Fig. 3(a) is a screenshot of SerialEM with a medium-magnification (5000×) overview of a neuron dendrite. The acquisition scheme is plotted in the interface with green lines and a yellow dashed line represents the tilt axis.Tomographic reconstructions from eTomo of two regions (each approximately 9 × 9 tiles in size) cropped out of the full montage are shown in (b) and (c) with an easymode [19] segmentation of membranes (cyan) and ribosomes (yellow) shown in (b). Zoomed regions of these tomograms are shown in (d)–(f) highlighting the dendrite body, a mitochondrion and a microtubule, respectively.

**Figure 3.**
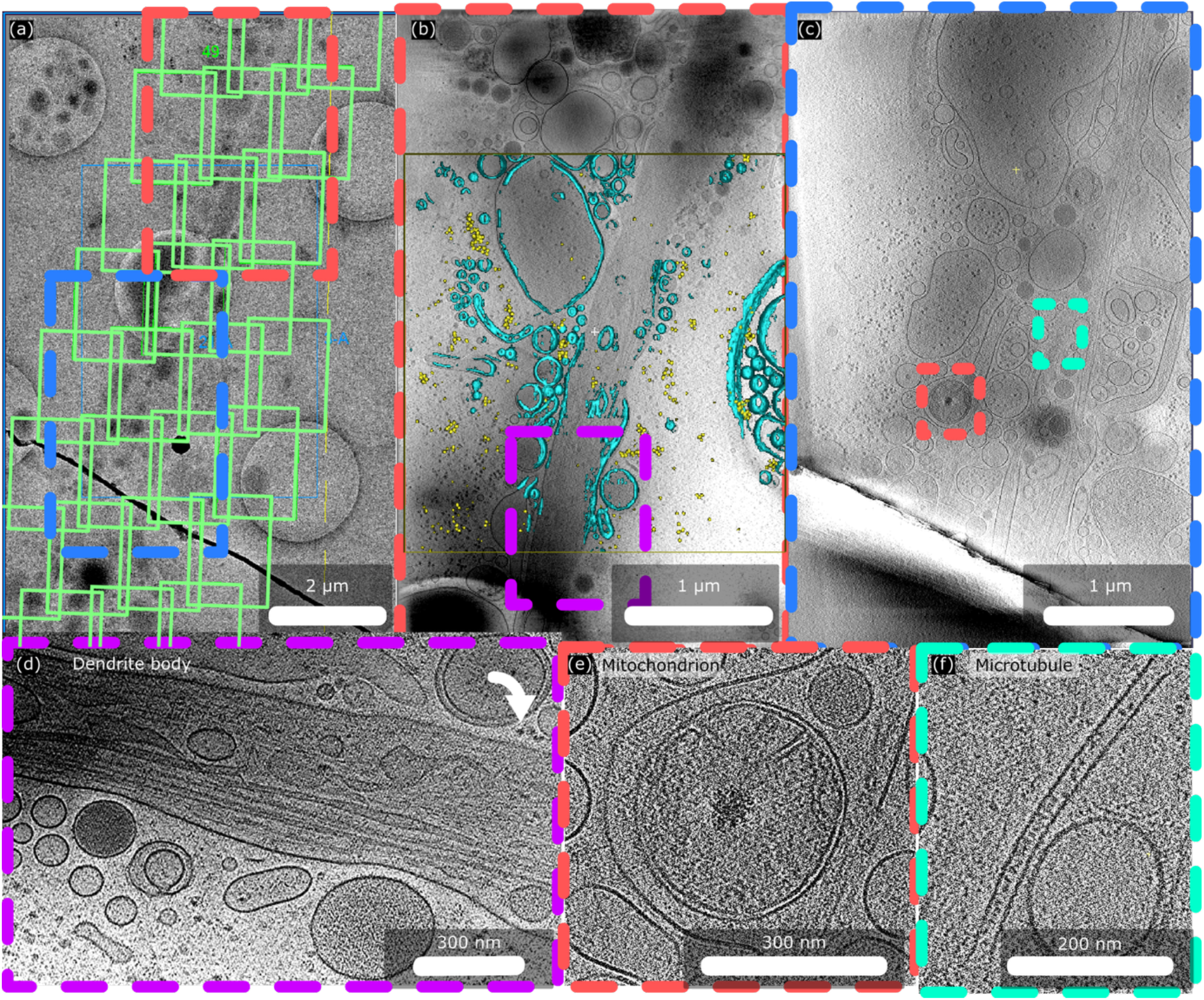
Montage tomography of a human neuron dendrite. (a) Overview image from SerialEM screenshot with montage acquisition scheme plotted. (b) Reconstruction of cropped region 1 with easymode [19] segmentation of membranes (cyan) and ribosomes (yellow) shown. (c) Reconstruction of cropped region 2. Shown in (d)–(f) are zoomed views of the reconstructions showing the dendrite body (rotated 90° due to space considerations), a mitochondrion and a complete microtubule, respectively.

## Conclusion

Cryo-electron tomography allows high-resolution 3D imaging of biological systems in a hydrated near-native state. The inherent dose-sensitivity of hydrated biological specimens means montaging is complicated with standard TEM hardware, since the round beam is mismatched to rectangular electron cameras. In this study we have demonstrated a cryo-tomography montaging scheme using rectangular beams which better match electron cameras and are more sympathetic to montaging. The approach is implemented through the widely adopted SerialEM software package and we have included the scripts in the supplementary material.

## Materials and Methods

This section sets out in full the montage design, dose management, acquisition, stitching and reconstruction procedures summarised in the main text.

## 1 Preparation of milled *Saccharomyces cerevisiae*

Commercial dried yeast (Saccharomyces cerevisiae) was revived in a solution of water with sugar and salt held at 30 °C for 3 h. Quantifoil R1.2/1.3 300-mesh holey carbon copper grids were glow discharged using a GloQube Plus glow discharge system (Quorum Technologies) at 30 mA for 15 s. A 4 μL aliquot of the yeast suspension was applied to each grid, followed by plunge freezing using a Vitrobot Mark IV (Thermo Fisher Scientific). Grids were blotted for 4 s using a blot force of −1 before rapid vitrification in liquid ethane. Frozen grids were stored in liquid nitrogen until further processing.

Cryo-focused ion beam (cryo-FIB) milling was performed using an Aquilos 2 Cryo-FIB/SEM (Thermo Fisher Scientific). Vitrified grids were transferred under cryogenic conditions and maintained below −170 °C throughout the milling procedure. The front side of the grids was sputter coated with platinum for 15 s at 30 mA, followed by a 120 s organometallic platinum coating using the gas injection system (GIS). Suitable regions containing yeast cells were identified using MAPS software (version 3.27, Thermo Fisher Scientific). Lamellae were prepared using AutoTEM Cryo software (version 2.4, Thermo Fisher Scientific). Automated milling was performed at a stage tilt corresponding to a milling angle of 15°. Sequential milling steps were performed using progressively lower ion beam currents, with rough milling at 1 nA, medium milling at 0.3 nA, fine milling at 0.1 nA, and final polishing at 50 pA. This automated milling workflow was used to progressively thin the lamellae while minimising beam-induced damage and producing electron-transparent specimens suitable for cryo-electron tomography.

## 2 Preparation of neurons

Differentiated neurons derived from the immortalized human BE(2)-M17 neuroblastoma cell line were routinely cultured in Dulbecco’s Modified Eagle Medium (DMEM) supplemented with 10% foetal bovine serum (FBS) and 1% penicillin– streptomycin. For cell growth, 200-mesh gold Quantifoil grids with carbon film were incubated for 24 h at 37 °C in complete growth medium consisting of DMEM supplemented with 10% FBS and 1% penicillin–streptomycin.

Following grid pre-conditioning, BE(2)-M17 cells were trypsinised and seeded at approximately 5 × 10^4^ cells per dish containing the pre-conditioned grids. Cells were incubated overnight at 37 °C to allow attachment and growth on the grids. The following day, neuronal differentiation was initiated by replacing the growth medium with DMEM supplemented with 1% FBS and 10 µM retinoic acid (RA). Cells were maintained in RA-containing medium for 3 days, with the medium replaced every 48 h.

Following RA treatment, differentiation was continued in DMEM supplemented with 1% FBS and 5 ng/mL brain-derived neurotrophic factor (BDNF). BDNF-containing medium was replaced every 48 h. After approximately 8–9 days of differentiation, neurons grown on the Quantifoil grids were prepared for cryo-electron tomography. Fiducial markers (10 nm gold nanoparticles) were added before plunge freezing in a Thermo Fisher Vitrobot Mark IV to facilitate tilt-series alignment and tracking during tomographic reconstruction.

## 3 Montage design and dose management

We used the SerialEM microscope automation software for tilt series design and data acquisition. Although the focus of this work is on a workflow for use with square and rectangular beams the workflow is compatible with microscopes that only have round apertures. Three scripts were developed, all of which are included in the supplementary material. The first, `DetermineOverlapFraction.txt,` allows a user to tune montage tile overlap and saves a vacuum reference beam for processing. The second, `setup_polygon_montage.txt`, takes a polygon drawn in the navigator of SerialEM, Fig. 1(a), and calculates the overlapping grid of tiles required for acquiring a montage tilt series that covers the polygon, Fig. 1(b). A full list of tilt angles with the corresponding positions of each tile in the montage is saved to a tilt information file (called Montage_imageshifts.txt in this example) and is reread by SerialEM during batch data acquisition, Fig. 1(c). A defocus offset, given by Δ*f* = *d* tan *θ*, where *d* is the distance of the tile from the tilt axis and θ is the tilt angle, is applied for each montage tile to compensate for the vertical displacement of the planar sample with non-zero tilt angle. The format of the tilt information file is shown in Fig. 1(c). The acquisition scheme for the first tilt is overlaid on the medium magnification image in the SerialEM interface as shown in the screenshot of Fig. 1(d). Overlap of the field of view of the tiles is required to align the images for stitching of the montage and this produces regions of the montage that are doubly and sometimes triply exposed to the beam. To mitigate dose accumulation in specific regions of the grid, the positions of the tiles in the grid are shifted laterally by an amount equal to the size of the overlap region – this is demonstrated in Fig. 1(e). The full dose accumulation for this hypothetical tilt series for a dose-symmetric series between −60° and +60°, with 3° steps (a total of 41 separate tilts), is shown in Fig. 1(f). In this example the dose from a single montage tile is 2.44 e^−^/Å^2^, such that if this were a standard tomographic tilt series the dose accumulated over 41 tilt steps would be 100 e^−^/Å^2^. The extra exposure from montage overlapping in this example is 11.7 e^−^/Å^2^ (i.e. 11.7% more dose than intended), with some variation in the per-pixel dose indicated in the histogram inset to Fig. 1(f), which has an interquartile range of 6.25 e^−^/Å^2^.

## 4 Acquisition

A third script, `Acquire_montage.txt`, reads the text file written by `setup_polygon_montage.txt` and acquires the montage tilt series, tilting the microscope stage to the angles described in the text file, Fig. 1(c), and acquiring images at the coordinates listed for that tilt angle. At each tilt step, a medium-magnification image (approximately 5000×, pixel size roughly 25 Å) is used to align the target with reference to a similar image recorded at the closest tilt in the tilt series (or one acquired when setting up the batch run in SerialEM if this is the first tilt in the series). Focusing is then performed at a region specified by the user when setting up the run. Images are acquired using “image-shift” where beam deflectors in the upper objective lens pole piece shift the position of the beam to the desired coordinate (beam shift) and this shift is balanced by deflection coils in the lower objective (image shift) so that the beam positioning on the camera is maintained. This is a common practice in high-throughput TEM as it avoids mechanical stage movements which are slower, exacerbate mechanical instabilities and are generally less accurate. Focusing and alignment steps are performed in advance of the montage acquisition for each tilt step, as is standard practice in TEM tomography. There are well developed protocols, implemented in SerialEM with the name “coma-free image shift”, for compensating for the lens aberrations that result from the changed optical path of the beam through the objective lens system. This approach was found to be very time efficient – very little time as a proportion of total acquisition time is spent on overheads such as focusing and position alignment relative to data acquisition. A user-specified delay between the application of the image shift and the image acquisition is used to ensure that the beam positioning on the camera and specimen has stabilised. Investigations of this in the context of singleparticle cryo-EM found that the maximum resolution of the images was unaffected by this delay but beam positioning accuracy was affected [20], so users should optimise this parameter for their own microscope based on beam placement stability. The output of a montage tilt-series run is a set of mrc files, one image stack per tilt containing that tilt’s tiles, together with a folder of dose-fractionated movies.

### 5 Tile preprocessing

The standard cryo-EM software MotionCor2 [15] is used for motion correction of raw movies recorded by SerialEM. If applied directly to raw movies, the cross-correlation algorithm will simply measure the displacement of the beam from frame to frame, so the script beam_mask_motioncorr.py first builds an intensity-based mask (using 0.4 times the median value as the cutoff threshold) from the average of the frames of the movie and the largest square that can be inscribed within that mask is calculated. A copy of the movie cropped to that square is then used as input to MotionCor2 and the resulting shifts then applied by beam_mask_motioncorr.py to the whole uncropped movie.

Even with energy-filtered data (a 10 eV slit width was used in all examples here), there are artefacts at the beam edges resulting from inelastic scattering which redistributes intensity from the otherwise sharp beam edges to the outside of the beam. Shown in Fig. 4(a) is a reference beam image, recorded with the stage moved to a hole in the grid. When the beam transits the specimen, inelastic scattering darkens the edges of the beam, and this is particularly noticeable at the seams of the different tiles if the beam images are to be stitched together in a montage. To ameliorate this, a parametrised scattering kernel, assumed to be of the form often used to model plasmon inelastic scattering,

**Figure 4.**
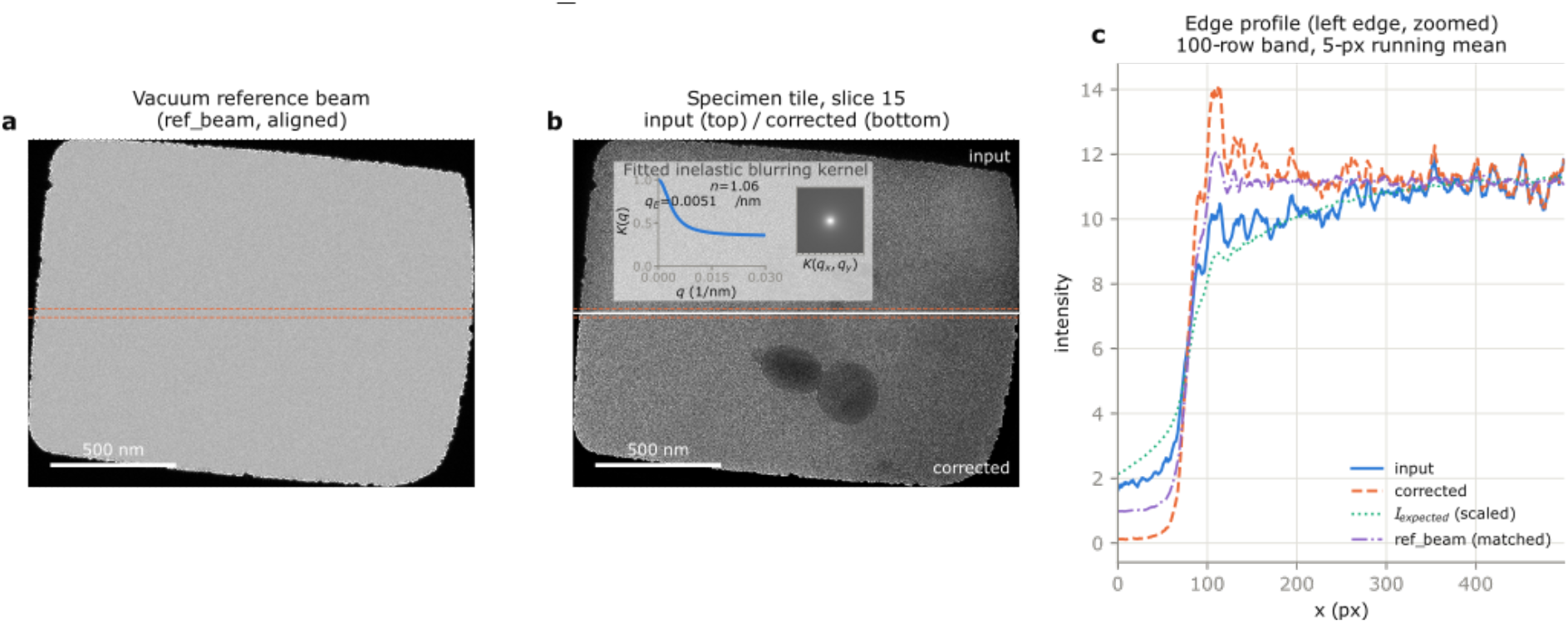
Description of de-doming procedure to compensate for inelastic scattering. (a) A vacuum reference beam, (b) an image from montage acquisition with fitted inelastic blurring kernel inset. The top half of (b) is uncorrected and the bottom half has been corrected by division of the blurred version of the reference beam in (a) that best matches experimental data. Shown in (c) is a line-scan of the edge of the beam.

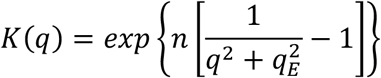

where *q* is the spatial frequency and *q*_*E*_ is a characteristic inelastic scattering width, n is the number of single scattering events, is fitted to the data. The expected image of the beam after inelastic scattering is obtained by applying this kernel to the vacuum reference image, *I*_*exp*_ = *F*^−1^{*F*{*I*_*ref*_} *K*(*q*)} where *F* denotes the Fourier transform, and the ratio *C* = *I*_*ref*_/*I*_*exp*_ then provides a correction map that is divided from the tile within the beam mask (clipped, and applied only where *I*_*exp*_ exceeds 5% of its maximum, so that noise outside the beam is not amplified). The two parameters *n* and *q*_*E*_ are fitted either for every tile or once for a representative tile, by a grid search on a downsampled (64 × 64 pixel) image followed by Nelder–Mead refinement of the match between the measured and expected beam profiles. The result of this correction is shown in Fig. 4(b) with the input tile shown in the top half of the image and the corrected tile in the bottom half. The fitted inelastic blurring kernel is plotted in an inset. A line scan of the edge of the beam is shown in Fig. 4(c); the input intensity forms a smooth dome in profile as the edge of the beam approaches, correction moves this profile up to match the reference beam. In this case there is a slight overcorrection since a single kernel is fitted for each beam and a thickness gradient from left to right is evident in Fig. 4(b).

## 6 Montage stitching

To perform montage stitching, the processing script `stitch.py` takes as input the mrc stacks containing the montage tiles, either output from the motion correction step or straight from the TEM, and the text file containing the tilts and montage image shifts used by SerialEM. The initial image shifts are used as a starting point for refinement. For a montage of a dose-sensitive specimen, the beam size will typically be condensed such that it is smaller than the camera size so that every part of the beam that interacts with the specimen is recorded and included in reconstruction. The first step is to mask out the beam using an intensity threshold that is a fraction of the median – by default this fraction is 0.4 but can be specified by the user. The Krios G4 used in this paper features a fringe-free (Köhler) illumination system where the beam-forming condenser optics are designed so that the condenser aperture and image planes are conjugate image planes. This means that the Fresnel fringes at the beam edge are minimal. These small Fresnel fringes need to be removed from the images in the mrc stack before montage stitching and this is performed by shrinking the mask, removing an outer layer of pixels from the mask. The initial guess of tile displacements and the image masks are used to determine regions of overlap between images. Shown in Fig. 2(b) are two tiles from a montage of the yeast lamella sample with the overlapping regions designated by a red transparent overlay. To refine the alignment of these regions we perform masked cross-correlation [21] on each pair of overlapping tiles. Standard cross-correlation finds the translation that maximises the product of two arrays, so it would identify the correct alignment as the one where the masks, rather than the images contained within them, are aligned. In masked cross-correlation, the cross-correlation product is only calculated in the region of mutual mask overlap, with the overlapping pixels of each tile mean-subtracted and the summed product normalised by the product of their standard deviations over that same overlap [21]. Because this normalisation accounts for the number of overlapping pixels, the score no longer increases simply as the mask overlap grows, so its peak instead better reflects alignment of the image features within the overlap rather than of the masks themselves. The stitched result of the normalised cross-correlation of the two images in Fig. 2(b) is shown in Fig. 2(c). The global alignment of all tiles is then the least-squares solution to a set of linear equations, ***x***_*i*_ − ***x***_*j*_ = Δ***x***_*ij*_, where vector ***x***_*i*_ is the (2D) position of tile *i* and Δ***x***_*ij*_ is the displacement as measured by masked cross-correlation between tiles *i* and *j*. With a standard least-squares solution a spurious cross-correlation alignment between a few pairs of tiles will have a large and noticeable impact on the global alignment so Tukey iteratively reweighted least squares [22] is used to find the global alignment instead. This is an iterative method where outlier measurements, i.e. those that deviate a large amount from the consensus solution, are down-weighted rather than being accepted or rejected once at the outset. Writing the normalised residual of each measured tile pair as u_*ij*_ = |x_*i*_ − x_*j*_ − Δ*x*_*ij*_|/τ where τ is a fixed tolerance (in microns) derived from the spread of the residuals of the initial image-shift positions, each pair is assigned a Tukey bi-weight 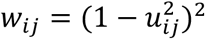 for *u*_*ij*_ < 1 and *w*_*ij*_ = 0 otherwise. The positions are then re-solved as the weighted least-squares problem minimising ∑ *w*_*ij*_ |x_*i*_ − x_*j*_ − Δ*x*_*ij*_|^2^,the residuals and weights are recomputed, and the cycle repeats until the largest change in any tile position falls below a tolerance (typically about ten iterations, with a ceiling of 25). For cases where “islands” of tiles do not have a reliable cross-correlation to the remaining tiles the algorithm defaults to using image shift to connect those sets of tiles to the larger montage. A montage stitched using this approach is shown in Fig. 2(d) with tile displacements plotted atop the image. At the extremities of the montage, the sharp edges of the tiles are readily observable against the background (which is initially set to the mean value inside the tiles of the tomogram). If back projected into the 3D reconstruction volume during tomography reconstruction, these edges create strong artefacts that obscure real features in the tomogram. So as a final step the smoothn algorithm [16] is applied to the stitched montage to blend the background into the montage, results shown in Fig. 2(e).

## 7 Montage masking

Cryo-TEM grids contaminate easily during transfer between sample preparation equipment such as a plunge freezing apparatus and the TEM. Cryo-FIB lamellae are even more prone to this contamination and often feature other artefacts such as curtaining. These artefacts, and other unavoidable features like the edge of a FIB lamella, often produce much more contrast in TEM micrographs than biological features of interest so can bias alignment in tomographic reconstruction, particularly the projection matching approach used by AreTomo2 [18]. Contamination artefacts are usually avoidable by judicious choice of acquisition area in standard tomography workflows but become more difficult to avoid in montage tomography due to the large and contiguous regions acquired. To address some of these challenges we developed a version of AreTomo2 that used masked cross-correlations for projection matching alignment and a simple Python matplotlib-based GUI that allowed the user to design these masks. A screenshot of the utility is shown in Figure 5 GUI for designing tilt-series masks. (a) A binned version of the tilt series is shown top left, hand-drawn masks can be added using the mouse (orange overlay is a previously drawn polygon mask, a new polygon is being added with the indicated blue line) and (b) by selecting a peak in the histogram. The user can move through the tilt series, or select histogram thresholding, using the sliders shown in (c), with buttons allowing different modes of manual masking. The histogram is adjusted to follow the cosine-like scaling of average intensity plotted in (d). A binned representation of the stitched tilt series is shown top left of the interface, Fig. 5(a), and masks can be drawn manually using a paint brush or polygon approach on the image (orange overlay), or by selecting a peak in the histogram, Fig. 5(b). The peaks corresponding to lamella and vacuum are labelled in the figure and the user has selected the lamella intensity peak by double clicking. The Gaussian function fit is plotted in green. The effect on the mask is visible in Fig. 5(a) (current mask overlaid in red). The user can move through the items in the tilt series using the sliders indicated in Fig. 5(c); the other two sliders control how much of the fitted histogram peak is included in the mask (N Sigma) or set a manual intensity threshold instead of peak fitting. Intensity thresholds are adjusted to account for the expected cosine-like variation of mean intensity with stage tilt, plotted in Fig. 5(d).

**Figure 5.**
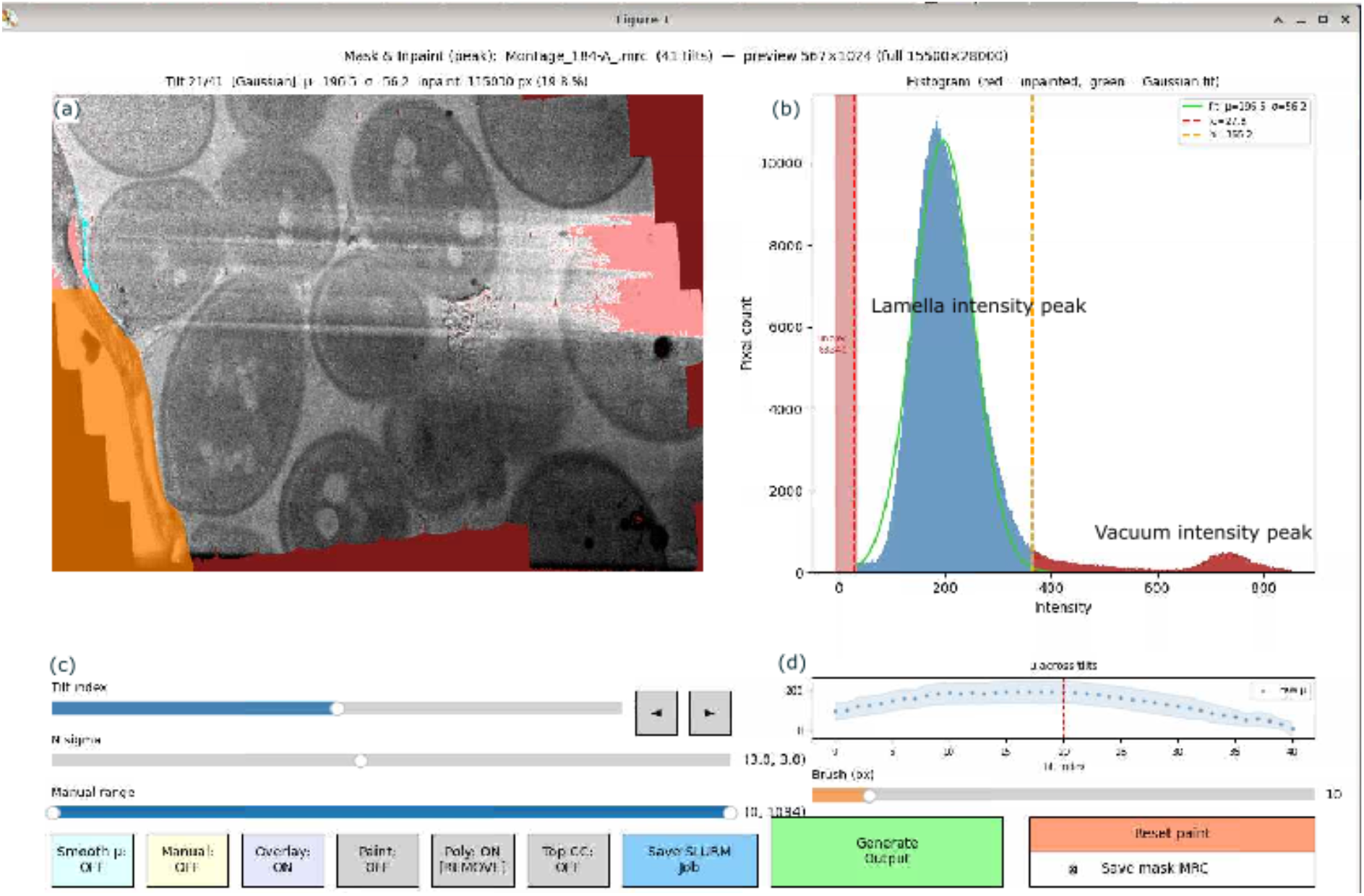
GUI for designing tilt-series masks. (a) A binned version of the tilt series is shown top left, hand-drawn masks can be added using the mouse (orange overlay is a previously drawn polygon mask, a new polygon is being added with the indicated blue line) and (b) by selecting a peak in the histogram. The user can move through the tilt series, or select histogram thresholding, using the sliders shown in (c), with buttons allowing different modes of manual masking. The histogram is adjusted to follow the cosine-like scaling of average intensity plotted in (d).

## 8 Reconstruction

After stitching and blending of montages the full tilt series of montages can be transferred to the AreTomo2 [18] or IMOD [17] tomographic reconstruction software for further processing. A few modifications were made to the AreTomo2 source code. To improve alignment for montage tomograms, which often include unavoidable regions of high contrast such as lamella edges, curtaining or surface contaminants which would dominate projection alignment, we added the option of providing a mask with array dimensions matching the tilt series, which makes AreTomo2 use masked crosscorrelations [21] in the projection matching step. In the modified version, to reduce memory usage for very large montage tomograms the output tomogram is streamed to a memory-mapped mrc file rather than written to RAM and then to disk at the completion of the program as in the standard AreTomo2 version. The modified version of AreTomo2 is available at https://github.com/HamishGBrown/MaskedMemoryEfficientAreTomo.

## 9 Code and data availability

All acquisition and reconstruction code is available on GitHub at https://github.com/HamishGBrown/StitchTomo and https://github.com/HamishGBrown/MaskedMemoryEfficientAreTomo and in the supplementary material. Raw data are available on request.

## 10 Acknowledgements

Authors acknowledge the use of the ThermoFisher Krios G4 cryo-TEM and ThermoFisher Aquilos Cryo-FIB at the Ian Holmes Imaging Centre. Anthropic’s Claude Code (claude.ai) was used for development of Python Serial-EM data acquisition and data processing scripts and for proof reading of the manuscript.

## Footnotes

1 https://github.com/HamishGBrown/StitchTomo

2 https://github.com/HamishGBrown/MaskedMemoryEfficientAreTomo

