## Supplementary material for "Whole cryo-FIB lamella acquisition at high magnification with rectangular beam montage cryo-electron tomography": Source code: metadata_design.md

# A manifest for montage tilt-series metadata

Status: proposal. Nothing here is implemented yet.

## The problem, stated from real failures

Each step of the pipeline is run on its own, writes its outputs, and forgets
what it did. Later steps then re-derive the geometry from the *data*, guess, or
inherit a default that does not match what actually happened. Every one of
these cost real time on this dataset:

| what was lost | how it surfaced |
|---|---|
| `stitch.py --rotate` | 02_stitch.sh has it commented out and the default is 0, but the canvas is rotated 90. Recovered by measuring tile footprints (36.8 px vs 578.6 px) and then correlating the canvas to break the 90/270 tie. |
| which `.aln` made which tomogram | The `.aln` on disk is from `-Patch 5 3`; the tomogram is from `-Patch 10 6`. Reprojection is off by 191 A rms perpendicular to the tilt axis, growing with tilt. |
| whether local patch alignment was used | The reconstruction has it, the reprojection ignores it by design. Neither records the choice, so the two disagreed silently. |
| tile index base | `ctf_results.txt` counts tiles cumulatively across the series; `fix_stack_tile_index.py` exists solely to repair this. |
| `index_map` label meaning | Labels are `0..n_selected-1`, not rows of `Refined_positions`. Costs a `flatnonzero(tile_selection)` that is easy to omit. |
| AreTomo `-TiltCor` offset | The TILT column is corrected (+12 deg here), so joining sections to tilts by angle silently pairs each tilt with the one 4 positions away. |
| canvas pixel size | AreTomo was told 6.87 for a canvas that is 6.852. The header is wrong; the geometry is not. |
| pixel size through inpainting | `inpaint_apply.py` wrote voxel_size 0, so 3dmod rescaled an IMOD model by 6.852 and threw every point off the canvas. |
| phase flipping | A comment in 01_motion_correction.sh says tiles are phase-flipped. They are not. |
| handedness | Never recorded, and no longer measurable from the CTF fits. |

The pattern is the same every time: **a step knew something and did not write it
down**, so a later step had to infer it, and an inference that is 95% reliable
is a bug that appears in 1 dataset in 20.

## What to build

One **manifest per montage tilt-series**, living beside the data, that every
step reads and appends to. Steps stay independent and individually runnable --
the manifest is a record, not a driver.

```
20260713Yeastattempt2/
  Montage_9-A.manifest.json     <- scalars, frames, provenance; human-readable
  Montage_9-A.manifest.h5       <- bulk arrays the JSON points at
```

### 1. Record what the step DID, not what it was asked

This is the single most important rule and it would have caught the rotate=90
case on its own. Parameters go in *after* defaults, `auto` resolution and any
clamping:

```json
"stitch": {
  "requested": {"rotate": null, "binning": 2, "roi": [-5500, 12000, -6000, 12000]},
  "effective": {"rotate": 90, "binning": 2, "roi": [-5500, 12000, -6000, 12000]},
  "auto_resolved": ["rotate"]
}
```

Existing auto-detectors (`rotate="auto"`, the SEC join) do not go away -- they
become *validators* that check the recorded value and fail loudly on a mismatch,
rather than being the source of truth.

### 2. Name every quantity with its frame and units

Most of the failures above are origin/unit confusion. `Refined_positions` says
nothing; the name should:

```
positions_canvas_px_bin2          (21, 2)  rows are ALL tiles, incl. deselected
tile_selection                    (21,)    bool, indexes the above
index_map_labels_are              "selected_subset"   <- not "positions_row"
```

### 3. Frames and transforms as first-class objects

Declare the frames once, then store the affine between *adjacent* ones. Nothing
downstream composes the chain by hand.

```json
"frames": {
  "raw_tile":  {"units": "px", "pixel_size_A": 3.426, "origin": "corner",
                "keyed_by": ["tilt", "tile"]},
  "canvas":    {"units": "px", "pixel_size_A": 6.852, "origin": "corner",
                "keyed_by": ["tilt"]},
  "aligned":   {"units": "px", "pixel_size_A": 13.704, "origin": "corner",
                "keyed_by": ["tilt"]},
  "tomogram":  {"units": "voxel", "pixel_size_A": 13.704, "origin": "centre"}
},
"transforms": [
  {"from": "raw_tile", "to": "canvas",  "h5": "transforms/raw_to_canvas",
   "note": "rot90 k=90, binning 2, tile origin from positions_canvas_px_bin2"},
  {"from": "canvas",   "to": "aligned", "h5": "transforms/canvas_to_aligned",
   "note": "AreTomo global ROT/TX/TY per section, out_bin 2"},
  {"from": "aligned",  "to": "tomogram", "params": {"tilt_axis": "row",
   "z_origin_voxel": 499.5, "handedness": 1, "theta_source": "aln.TILT"}}
]
```

`z_origin_voxel` and `handedness` become recorded, tunable numbers instead of
constants buried in `tomo_to_aligned`. The measured -14.5 voxel z-centre error
has somewhere to live.

### 4. Fingerprint inputs, so mismatches are caught not guessed

Each step records a cheap fingerprint of every input -- `(size, mtime, sha1 of
the first 64 kB + the header)`. A later step verifies that the `.aln` it was
handed is the one that produced the tomogram it was handed. That check alone
turns the `-Patch 5 3` vs `-Patch 10 6` problem from a week of confusion into a
one-line error at startup.

```json
"aretomo": {
  "effective": {"patch": [10, 6], "out_bin": 2, "vol_z": 1000,
                "tilt_cor_offset_deg": 12.0, "local_alignment": true},
  "inputs":  [{"path": ".../Montage_9-A_inpainted.mrc", "fp": "a3f1...", "size": 12902400000}],
  "outputs": [{"path": ".../recon_patch106_bin2.mrc", "fp": "9c02..."},
              {"path": ".../Montage_9-A_patch106.aln", "fp": "77bd..."}]
}
```

Note `"local_alignment": true`. Recording it forces the reprojection code to
either apply the patch table or refuse, instead of silently ignoring it.

### 5. Steps stay one-by-one

The manifest never runs anything. A step:

1. loads the manifest (creating it if absent),
2. verifies its own inputs' fingerprints against what the producing step wrote,
3. does its work,
4. appends/overwrites *its own* section and marks every downstream section
   `"status": "stale"`.

Re-running stitching marks AreTomo and everything after it stale, which shows up
as a warning rather than a wrong answer three steps later. Nothing forces you to
re-run them.

## Format

Hybrid, because the two kinds of metadata want different things:

- **JSON** for scalars, frames, effective parameters, provenance. Diffable,
  greppable, git-friendly, readable in a terminal at 2am.
- **HDF5** for bulk arrays (`positions`, `index_map`, `beam_masks`, per-section
  transforms), which is where they already live. The JSON holds the h5 path and
  dataset name.

STAR was considered and rejected for the manifest itself: it has no good way to
express nesting or heterogeneous per-step records. The RELION export stays STAR,
generated *from* the manifest.

## Migration

Do not rewrite the pipeline. Three steps, each useful alone:

1. **`montage_manifest bootstrap`** -- run the existing auto-detection once
   against a finished dataset and write down what it finds (rotate, SEC join,
   pixel sizes, tilt list). Immediately gives 20260713Yeastattempt2 a manifest
   without re-running anything.
2. **Make writers record.** Add a `manifest.record_step(...)` call at the end of
   `beam_mask_motioncorr`, `stitch`, `inpaint_apply`, and an AreTomo wrapper.
   Each is a few lines and changes no behaviour.
3. **Make readers consume.** `MontageProjector` takes a manifest instead of
   eleven keyword arguments, and its auto-detectors become validators. This is
   where the payoff lands: `--roi`, `--pixel-size`, `--binning`, `--out-bin`,
   `--rotate`, `--extra-shift`, `--handedness` all stop being CLI arguments that
   have to be kept in sync by hand with 02_stitch.sh.

Step 1 is worth doing on its own even if 2 and 3 never happen.
