## Supplementary material for "Whole cryo-FIB lamella acquisition at high magnification with rectangular beam montage cryo-electron tomography": Source code: Montage tomography SOP.docx

### Montage tomography Serial-EM SOP

1. Make folder for acquisition, this will store atlases, gridsquares and the montages theselves when SerialEM prompts you on start up.
2. Make Atlas (remember to make Navigator map) and align to lowest SA mag (
3. Setup low dose presets:
   a. View, nanoprobe 6400 x mag (lowest SA mag where beam covers camera)
   b. Record, get beam as close as possible to size of camera
   c. Focus, same mag and spot size as exposure but make sure beam covers camera
4. Perform microscope alignments
   a. Move to vacuum
   b. C2 aperture alignment and astigmatism
   c. Beam tilt pivot points
   d. Filter alignments (tune gif)
   e. Gain reference (in GMS, also acquire record image with beam size to be used in acquisition)
   f. Dose (measure e/pix/s in GMS, calculate e/A/s and work out exposure time)
   g. Move to carbon
   h. Rotation centre (this is really important!)
   i. Objective astigmatism
   j. Objective coma
   k. Beam tilt pivot points
   l. Beam shift
   m. View – Record alignment in Serial-EM
   n. Use Determine overlap_function.py script to work out your beam overlap. Write these values into the Setup_square_montage.py script. **Run the scripts otherwise these values wont be updated!**
5. Acquire search montages:
   a. Add points to Atlas in middle of Grid squares and tick “acquire” for each of these points in the navigator
   b. File -> Open New, call the new file “searchmontages.mrc”
   c. File -> set up Montage, make sure that stage
   d. Navigator -> Acquire at items, remember to tick “make navigator maps”
6. Add montage positions (or polygons) to searchmontages maps.
   a. Select to acquire each of these in the navigator.
   b. File -> Open New, call the new file “ViewImages.mrc”
   c. Use “Acquire at items” and make sure that the following settings are chosen:
      i. images are being acquired at view preset
      ii. New navigator maps are being made at each point
      iii. perform rough and fine eucentric jobs before acquisition
      iv. Run “Setup_square_montage.py” or “Setup_square_montage_from_polygon.py” after acquisition as appropriate
   d. Check that the label in “xxx” “Montage_imageshifts_xxx.txt” files corresponds to the navigator labels of the view images.
7. Before running the acquisition script check:
   a. Rotation centre
   b. Tilt pivot points
   c. Beam centering
   d. Focus area in low dose (can edit focus area per montage using “Edit Focus” box in the navigator)
   e. Acquisition time is what you intended it to be.
8. Select the new view image maps in the navigator for acquisition and choose acquire at items choose:
   a. To run the “Montagetiltseries” script at each item.
   b. Close column valves at end.
