## Supplementary material for "Whole cryo-FIB lamella acquisition at high magnification with rectangular beam montage cryo-electron tomography": Source code: aretomo_record.py

```
#!/usr/bin/env python3
"""
Record an AreTomo run in a provenance sidecar beside its output volume.

AreTomo is not wrapped. It runs exactly as it always did, keeps its own exit
code and stdout, and this is called immediately afterwards with the same
argument list::

    args=(-InMrC "$stack" -Patch 5 3 -OutMrC "$vol" -VolZ 2000 -OutBin 4 ...)
    $Aretomo "${args[@]}"
    montage_aretomo_record --variant full \\
        --aln "$outdir/series.aln" -- "${args[@]}"

Passing the array to both is the point: everything else about a run survives
it — the .aln is the product, the volume is on disk, the log echoes the
parameters — but the exact command line does not, and reconstructing it from
the log is a hostage to AreTomo's version.

What the record is for
----------------------
Three questions that currently have no answer once a job has scrolled past:

* **Which stack was reconstructed?** The input is fingerprinted, and its own
  sidecar says which step wrote it, so "was this the inpainted one or the raw
  stitch, the full dose or a half set" is a lookup rather than an inference
  from a directory name.
* **Was the alignment solved or supplied?** ``-AlnFile`` reuses an existing
  solution — the odd/even runs do exactly this — and a stale one silently
  reconstructs a half set in a different frame from the full dose. The file
  is fingerprinted and checked against the one the solving run wrote.
* **Where is the tomogram, in canvas terms?** The per-section ROT/TX/TY table,
  the binning, and ``z_sign`` — the direction of the depth axis, which nothing
  in the file itself reveals — are all recorded, which is the chain a
  projector needs and used to get as loose arguments.

The local (patch) alignment table is *summarised*, not copied: its shape is
recorded and the ``.aln`` is named as where to read it. The manifest used to
duplicate the whole table into an HDF5 sibling, but nothing ever read that
copy — ``montage_projection`` parses the ``.aln`` directly — so it was a
second copy of a file that was already on disk and already pointed at.

Whether the field was *applied* is a different question from whether it
exists, and that one is recorded as a fact: see :func:`local_applied`. Getting
it wrong is what let a reconstruction and a reprojection disagree by 191 A
with nothing on record to say why.

The sidecar goes beside the output volume. Under the manifest this step had to
*find* a shared file first, and on Montage_13A_new ``--manifest auto`` found
nothing and recorded nothing, silently — the name it searched for did not match
the one motion correction had created. There is no name to get wrong here.

AreTomo3 support
-----------------
AreTomo3's CLI uses different flag names for the same concepts (``-InPrefix``
instead of ``-InMrc``, ``-OutDir`` instead of ``-OutMrc``, ``-AtPatch``/
``-AtBin`` instead of ``-Patch``/``-OutBin``) and has no ``-AlnFile``, so an
AreTomo3 run's ``alignment_source`` is always ``"solved"``. Because
``-OutDir`` names a directory, not the reconstructed volume, callers running
AreTomo3 must always pass ``--output`` with the actual volume path — there is
no flag to infer it from, unlike ``-OutMrc``. ``--aln`` should likewise
usually be passed explicitly: AreTomo3 writes it into ``-OutDir`` named after
the ``-InPrefix`` stem, which ``default_aln_path`` approximates via
``output_dir`` but has not been checked against a real AreTomo3 ``.aln`` on
disk.
"""

from __future__ import annotations

import argparse
import logging
import os
import sys

from pathlib import Path

try:
    from . import sidecar
    from .refine_montage_projmatch import parse_aln
except ImportError:  # invoked as a plain script
    import sidecar
    from refine_montage_projmatch import parse_aln

logger = logging.getLogger(__name__)

SIDECAR_STEP = "aretomo"

#: AreTomo flags worth naming, and how many values each takes. Anything else
#: on the command line still reaches the record verbatim through ``argv``;
#: this is only about which values become queryable fields.
_FLAGS = {
    "-InMrc": ("input", 1, str),
    "-OutMrc": ("output", 1, str),
    "-AlnFile": ("aln_in", 1, str),
    "-AngFile": ("angle_file", 1, str),
    "-MaskFile": ("mask_file", 1, str),
    "-TmpFile": ("tmp_file", 1, str),
    "-Patch": ("patch", 2, int),
    "-VolZ": ("volz", 1, int),
    "-OutBin": ("outbin", 1, float),
    # AreTomo3: same concepts, different flag names -- see "AreTomo3 support"
    # above. -InPrefix is a full tilt-series path in single-tilt-series mode
    # (the only mode this pipeline uses), same shape as -InMrc. -OutDir is a
    # directory, not the output volume, so it is kept separate from "output"
    # rather than aliased to it.
    "-InPrefix": ("input", 1, str),
    "-OutDir": ("output_dir", 1, str),
    "-AtPatch": ("patch", 2, int),
    "-AtBin": ("outbin", 1, float),
    "-TiltCor": ("tiltcor", -1, float),   # 1, or "1 <offset>"
    "-TiltAxis": ("tilt_axis", -1, float),
    "-PixSize": ("pixel_size_A", 1, float),
    "-Kv": ("voltage_kV", 1, float),
    "-Cs": ("cs_mm", 1, float),
    "-Defoc": ("defocus", 1, float),
    "-FlipVol": ("flip_vol", 1, int),
    "-FlipInt": ("flip_int", 1, int),
    "-Wbp": ("wbp", 1, int),
    "-Align": ("align", 1, int),
    "-DarkTol": ("dark_tol", 1, float),
    "-Gpu": ("gpu", -1, int),
    "-Sart": ("sart", -1, int),
}


def parse_aretomo_argv(argv: list) -> dict:
    """Pull the named flags out of an AreTomo command line.

    Case-insensitive on flag names, because AreTomo accepts ``-InMrC`` and
    ``-InMrc`` alike and shell scripts in the wild use both. Unknown flags are
    not an error: they stay in the recorded ``argv``, which is verbatim.
    """
    lookup = {k.lower(): v for k, v in _FLAGS.items()}
    parsed: dict = {}
    i = 0
    while i < len(argv):
        token = argv[i]
        entry = lookup.get(token.lower()) if token.startswith("-") else None
        if entry is None:
            i += 1
            continue
        name, count, cast = entry
        values = []
        j = i + 1
        # count -1 means "greedy": take values until the next flag. A bare
        # negative number is a value, not a flag (-TiltAxis -8.3).
        while j < len(argv) and (count < 0 or len(values) < count):
            nxt = argv[j]
            if nxt.startswith("-") and not _is_number(nxt):
                break
            values.append(nxt)
            j += 1
        try:
            cast_values = [cast(v) for v in values]
        except ValueError:
            cast_values = values
        if not cast_values:
            parsed[name] = True          # a flag given with no value
        elif len(cast_values) == 1 and count == 1:
            parsed[name] = cast_values[0]
        else:
            parsed[name] = cast_values
        i = j
    return parsed


def _is_number(token: str) -> bool:
    try:
        float(token)
    except ValueError:
        return False
    return True

def default_aln_path(parsed: dict) -> str | None:
    """Where AreTomo puts the .aln when it is not told.

    AreTomo2: the directory comes from ``-OutMrc`` and the *name* from
    ``-InMrc``, which is not the obvious pairing: reconstructing
    ``.../stitched/Montage_9-A_inpainted.mrc`` into ``$scratch/toberotated1.mrc``
    leaves ``$scratch/Montage_9-A_inpainted.aln``. Guessing it from the output
    name instead looks for a file that is never written.

    AreTomo3: there is no ``-OutMrc`` to take a directory from, only
    ``-OutDir`` (parsed as ``output_dir``), which AreTomo3 writes the .aln
    into directly, still named after the input stem. Not verified against a
    real AreTomo3 .aln on disk -- pass ``--aln`` explicitly where possible.
    """
    if not parsed.get("input"):
        return None
    stem = os.path.splitext(os.path.basename(parsed["input"]))[0]
    if parsed.get("output"):
        directory = os.path.dirname(os.path.abspath(parsed["output"]))
    elif parsed.get("output_dir"):
        directory = os.path.abspath(parsed["output_dir"])
    else:
        directory = os.path.dirname(os.path.abspath(parsed["input"]))
    return os.path.join(directory, stem + ".aln")


def local_applied(parsed: dict, aln) -> bool:
    """Did *this* run's output actually get the local (patch) correction?

    Not the same question as "does the .aln have a local table", because of a
    bug in AreTomo2's own reader. ``CLoadAlnFile::mLoadLocal`` scans each patch
    row with::

        sscanf(pcLine, "%d %d %f %f %f %f %f", &t, &p,
           m_pLocalParam->m_pfCoordXs+i, ... m_pLocalParam->m_pfShiftYs+i,
           m_pLocalParam->m_pfGoodShifts);     // <-- no "+i"

    (MrcUtil/CLoadAlnFile.cpp:200-205). Every good flag is written to element
    zero, so after a reload the whole array is the zeros ``Setup`` memset it to,
    except ``[0]``. ``mGCalcLocalShift`` then skips every patch whose flag is
    below 0.9 (GCorrPatchShift.cu:31) and finds none, so ``iCount == 0`` and the
    local shift is identically zero.

    The consequence is worth stating plainly: **a run given ``-AlnFile`` does
    not apply the local alignment, even when the file it was handed contains
    one.** Only the run that *solves* the alignment corrects locally, because
    there the table is still the in-memory one ``CPatchAlignMain`` produced.

    So on this dataset the bin-2 tomogram (solved in-run) is locally corrected
    while the aligned tilt series written beside it with ``-AlnFile`` is not,
    and the two are in different frames by the size of the patch field. That is
    not a montage problem and not something this pipeline introduced; it is
    upstream, and it needs to be in the record because nothing else shows it.
    """
    if not aln.has_local:
        return False
    return not parsed.get("aln_in")


def depth_axis_sign(parsed: dict, clip_rotx: bool) -> int:
    """
    ``z_sign`` for a volume: which way its Z axis runs.

    The reprojection formula in ``montage_projection.tomo_to_aligned`` is the
    textbook reading of AreTomo's own kernels, written for ``z_sign=+1``. A
    volume with *exactly one* depth flip applied needs the opposite sign,
    ``-1``, and that -- AreTomo's ``-FlipVol`` and IMOD's ``clip rotx`` do the
    same thing (CProcessThread.cpp:505-515 vs IMOD's clip, checked directly),
    so it does not matter which of the two did it -- is what every volume this
    pipeline has reconstructed so far actually is, measured on 9-A.

    Both, or neither, is not simply a sign: the axes come out in a different
    order, ``(x, z, y)`` rather than ``(x, y, z)``, so the third column of a
    pick is not depth at all and no value of ``z_sign`` will save it. Those
    cases return +1 (the untransformed reading) and are worth a warning at the
    point of use, not a silent number.
    """
    flips = int(bool(parsed.get("flip_vol"))) + int(bool(clip_rotx))
    return -1 if flips == 1 else 1


def series_base_for_stack(path: str) -> str:
    """Tilt series base name from the stack AreTomo was pointed at."""
    stem = os.path.splitext(os.path.basename(path))[0]
    suffixes = ("_inpainted", "_inpaint", "_odd", "_even", "_mask", "_stitched")
    stripped = True
    while stripped:
        stripped = False
        for suffix in suffixes:
            if stem.endswith(suffix):
                stem = stem[: -len(suffix)]
                stripped = True
    return stem


def classify_input(input_path: str) -> tuple:
    """Say which earlier step produced the stack being reconstructed.

    The question this whole module exists to answer: two plausible inputs sit
    side by side on disk, differing only in whether the beam edges were filled
    in, and nothing until now recorded which one a tomogram came from.

    Under the manifest this searched a shared file's ``outputs`` lists, which
    only worked if that file could be found and had been finalized — two ways
    to get "unknown" that had nothing to do with the input. Now it asks the
    input file itself, which either carries a sidecar or does not.
    """
    doc = sidecar.read_for(input_path)
    if doc is None:
        return "unknown", None
    return doc.get("step", "unknown"), doc


def verify_aretomo_inputs(input_path: str, aln_in: str | None) -> list:
    """Check the input stack and any supplied .aln against what was recorded.

    Returns a list of human-readable warnings, which are also written into the
    sidecar: a mismatch here cannot un-reconstruct the volume that has already
    been written, so it has to be visible in the record rather than only in a
    log that scrolls past.
    """
    warnings = []
    source, doc = classify_input(input_path)

    if doc is not None:
        recorded = doc.get("output_fingerprint") or {}
        if recorded.get("sha1"):
            now = sidecar.fingerprint(input_path)
            if now["sha1"] != recorded["sha1"]:
                warnings.append(
                    f"input stack has changed since {source} recorded writing "
                    f"it ({recorded.get('size')} B then, {now['size']} B now)")
    elif source == "unknown":
        warnings.append(
            f"input stack {os.path.abspath(input_path)} carries no sidecar, "
            "so which step produced it — and with what geometry — is not "
            "recorded anywhere")

    if aln_in:
        doc = sidecar.read_for(aln_in)
        if doc is None:
            warnings.append(
                f"alignment {os.path.abspath(aln_in)} was supplied with "
                "-AlnFile but carries no sidecar, so no recorded AreTomo run "
                "is known to have written it; reusing an alignment of unknown "
                "provenance")

    for message in warnings:
        logger.warning("Sidecar: %s", message)
    return warnings


def local_summary(aln) -> dict:
    """Shape and extent of the patch table, without inlining it.

    The table itself is (sections x patches) numbers — 4100 rows on a 41-tilt
    10x10 run — and the manifest used to copy it into an HDF5 sibling. Nothing
    ever read that copy: ``montage_projection`` parses the ``.aln`` directly
    via ``parse_aln``. So the sidecar records the table's shape and points at
    the file, which is what ``sidecar_metadata_plan.md`` means by bulk arrays
    staying as their own files.
    """
    local = getattr(aln, "local", None)
    if local is None:
        return {"present": False}
    n_sec, n_patch = local.coord_x.shape
    return {
        "present": True,
        "n_sections": int(n_sec),
        "n_patches": int(n_patch),
        "read_from": "the .aln itself, via refine_montage_projmatch.parse_aln",
    }


def aretomo_effective(
    *, argv: list, parsed: dict, input_path: str, output_path: str | None,
    aln_path: str, aln, variant: str, binary: str | None,
    clip_rotx: bool = True, output_pending: bool = False,
) -> dict:
    """Everything about this run a later step could need, in one flat block.

    The manifest split this across a step section, two ``frames`` entries and
    three ``transforms``. Nothing downstream ever walked that graph — the one
    real consumer, ``montage_projection``, reached in for five scalars — so
    the geometry that used to live in transform ``params`` is here, and the
    long-form notes that made those entries worth reading come with it.
    """
    source, _ = classify_input(input_path)
    solved = not parsed.get("aln_in")
    binning = float(parsed.get("outbin") or 1)
    z_sign = depth_axis_sign(parsed, clip_rotx)
    if z_sign == 1:
        logger.warning(
            "flip_vol=%s and clip_rotx=%s means the depth axis is flipped "
            "twice or not at all -- unlike every volume this pipeline has "
            "reconstructed so far. Recording z_sign=+1, but check the volume: "
            "with no flip the axis order is (x, z, y) and picks made in it "
            "are not (X, Y, Z) at all.", parsed.get("flip_vol"), clip_rotx,
        )
    pixel_size = parsed.get("pixel_size_A")

    return {
        "variant": variant,
        "input": os.path.abspath(input_path),
        "input_from": source,
        "output": os.path.abspath(output_path) if output_path else None,
        # True when the volume is still on its way to that path — written to
        # scratch by AreTomo and moved by a queued job.
        "output_pending": bool(output_pending),
        "aln": os.path.abspath(aln_path),
        # Solved here, or handed in from another run. The odd/even runs supply
        # the full-dose solution deliberately; a *stale* supplied file is the
        # failure this distinction exists to make visible.
        "alignment_source": "solved" if solved else "supplied",
        "aln_supplied": (os.path.abspath(parsed["aln_in"])
                         if parsed.get("aln_in") else None),
        "patch": parsed.get("patch"),
        "volz": parsed.get("volz"),
        "outbin": parsed.get("outbin"),
        "tiltcor": parsed.get("tiltcor"),
        "tilt_axis": parsed.get("tilt_axis"),
        "pixel_size_A": pixel_size,
        "tomogram_pixel_size_A": (pixel_size * binning) if pixel_size else None,
        "voltage_kV": parsed.get("voltage_kV"),
        "cs_mm": parsed.get("cs_mm"),
        "flip_vol": parsed.get("flip_vol"),
        "wbp": parsed.get("wbp"),
        "dark_tol": parsed.get("dark_tol"),
        "mask_file": parsed.get("mask_file"),
        "angle_file": parsed.get("angle_file"),
        "binary": binary,
        # Sections are the frames AreTomo *kept*: it drops dark ones from the
        # table rather than flagging them, so this is smaller than the stack
        # and SEC is sparse. Both counts are recorded because their sum should
        # be the stack depth, and a mismatch means the .aln belongs to another
        # stack.
        "n_sections": len(aln.sections),
        "n_dark": len(aln.dark_secs or aln.dark_tilts),
        "dark_secs": [int(s) for s in aln.dark_secs],
        "dark_tilts": [float(t) for t in aln.dark_tilts],
        "sec_range": ([int(aln.sections[0].sec), int(aln.sections[-1].sec)]
                      if aln.sections else None),
        "raw_size": list(aln.raw_size) if aln.raw_size else None,
        "num_patches": aln.num_patches,
        "has_local_alignment": bool(aln.has_local),
        "local_alignment": local_summary(aln),
        # Whether the local field was actually applied to *this* run's output.
        # Not the same as having one in the file: see `local_applied`.
        "local_alignment_applied": local_applied(parsed, aln),

        # -- geometry: what used to be `frames` and `transforms` -------------
        # canvas -> tomogram is really two hops, and collapsing them is what
        # hid the patch field: the local correction lives strictly inside the
        # first one, so a single entry has nowhere to put it and silently
        # describes a global-only chain the reconstruction did not use.
        "clip_rotx": clip_rotx,
        "z_sign": z_sign,
        "canvas_to_aligned": (
            "aligned = R(-ROT) @ (canvas - centre - (TX, TY) - L) + centre, "
            "then divided by outbin. L is the local patch shift, evaluated at "
            "the pre-shift centred coordinate R(ROT) @ (aligned*outbin - "
            "centre); it is zero when there is no patch table. Join on 'sec', "
            "never on the tilt angle."
        ),
        "aligned_to_tomogram": (
            "col = cx + (X - cx) cos(TILT) - z_sign (Z - cz) sin(TILT), "
            "row = Y, with TILT the column as written (any -TiltCor offset "
            "already folded in). z_sign is the direction of the volume's "
            "depth axis: -1 for one depth flip, by 'clip rotx' or by "
            "-FlipVol, which is what every volume this pipeline has "
            "reconstructed so far actually needs (measured on 9-A) even "
            "though it is the opposite of montage_projection's textbook "
            "formula; +1 for both flips or neither. Get it wrong and the "
            "reprojection is exact at 0 deg and out by 2 (Z - cz) sin(TILT) "
            "at the extremes."
        ),
        "sections": aln_section_records(aln),
        "argv": list(argv),
    }


def aln_section_records(aln) -> list:
    """One row per section of the .aln: SEC, ROT, GMAG, TX, TY, TILT.

    SEC is the join key AreTomo itself uses and the only reliable one — the
    TILT column carries any -TiltCor offset, so it does not match the nominal
    stage tilts.

    Dark frames are *absent* from these rows rather than flagged in them:
    AreTomo drops them from the global table, so a 41-tilt stack with 13 dark
    frames yields 28 rows whose SEC starts at 3. Their indices go in
    ``dark_secs`` instead, and the gap in SEC is the record of them.

    Tens of rows of six floats — small enough to inline, which is the whole
    reason the HDF5 sibling is gone.
    """
    return [
        {
            "sec": int(s.sec),
            "rot": float(s.rot),
            "gmag": float(s.gmag),
            "tx": float(s.tx),
            "ty": float(s.ty),
            "tilt": float(s.tilt),
        }
        for s in aln.sections
    ]


def record_aretomo_run(
    *, argv: list, parsed: dict, input_path: str, output_path: str,
    aln_path: str, aln, variant: str, binary: str | None,
    warnings: list, clip_rotx: bool = True, output_pending: bool = False,
) -> Path:
    """Write the sidecar beside the reconstruction and return its path.

    Beside the *output volume*, which is the change that matters. The manifest
    had to be found first — ``--manifest auto`` searched at and above the input
    stack for a file named after the series — and on Montage_13A_new it found
    nothing and recorded nothing, silently, because the name it looked for did
    not match the name motion correction had created. A sidecar is named after
    the file it describes, so there is no name to get wrong and nowhere for it
    to fail to land.
    """
    effective = aretomo_effective(
        argv=argv, parsed=parsed, input_path=input_path,
        output_path=output_path, aln_path=aln_path, aln=aln, variant=variant,
        binary=binary, clip_rotx=clip_rotx, output_pending=output_pending,
    )
    inputs = [sidecar.fingerprint(input_path, role="tilt_series")]
    if parsed.get("aln_in") and os.path.exists(parsed["aln_in"]):
        inputs.append(sidecar.fingerprint(parsed["aln_in"], role="aln_supplied"))

    return sidecar.write(
        output_path,
        SIDECAR_STEP,
        effective,
        inputs=inputs,
        requested={"patch": parsed.get("patch"), "volz": parsed.get("volz"),
                   "outbin": parsed.get("outbin"), "variant": variant},
        extra={"warnings": warnings} if warnings else None,
        # A volume still being moved into place by a queued job has nothing
        # stable to fingerprint yet.
        fingerprint_output=not output_pending,
    )


def main() -> None:
    parser = argparse.ArgumentParser(
        description="Record an AreTomo run in a sidecar beside its output.",
        formatter_class=argparse.RawDescriptionHelpFormatter,
        epilog=__doc__,
    )
    parser.add_argument("--aln", default=None,
                        help="The .aln AreTomo wrote or reused. Defaults to "
                             "-AlnFile when the alignment was supplied, else "
                             "to AreTomo's own naming: the -InMrc basename, in "
                             "the -OutMrc directory")
    parser.add_argument("--variant", default="full",
                        choices=["full", "odd", "even"],
                        help="Which stack this reconstruction is of (default: "
                             "full). Each variant is its own output file with "
                             "its own sidecar, so this is a label rather than "
                             "a record key as it was under the manifest")
    parser.add_argument("--output", default=None, dest="output_override",
                        help="Where the volume really ends up, when -OutMrc is "
                             "not its final home. Reconstructions are commonly "
                             "written to scratch and moved into place by a "
                             "queued 'clip rotx' job. The sidecar is written "
                             "beside THIS path, since that is the file a later "
                             "step will open; the -OutMrc path is still in the "
                             "recorded argv")
    parser.add_argument("--binary", default=None,
                        help="Path to the AreTomo executable, recorded so a "
                             "result can be tied to the build that made it")
    parser.add_argument("--strict", action="store_true", default=False,
                        help="Exit non-zero if the input stack or a supplied "
                             ".aln does not match what was recorded. The "
                             "default warns and records the warning: the "
                             "volume is already written by this point, so the "
                             "job's exit code is about visibility, not rescue")
    parser.add_argument("--no-clip-rotx", action="store_false", default=True,
                        dest="clip_rotx",
                        help="The volume does NOT get IMOD's 'clip rotx' after "
                             "AreTomo. Together with -FlipVol this fixes the "
                             "recorded z_sign, the direction of the volume's "
                             "depth axis: exactly one of the two flips gives "
                             "z_sign -1, both or neither gives +1 and leaves "
                             "the axes in an order picks cannot be read in")
    parser.add_argument("--output-pending", action="store_true", default=False,
                        help="The volume is not at --output yet (a queued job "
                             "will put it there). Records the parameters now "
                             "and skips the output fingerprint")
    parser.add_argument("argv", nargs=argparse.REMAINDER,
                        help="-- followed by the AreTomo command line, "
                             "verbatim")

    args = parser.parse_args()
    logging.basicConfig(level=logging.INFO, format="%(levelname)s: %(message)s")

    argv = list(args.argv)
    if argv and argv[0] == "--":
        argv = argv[1:]
    if not argv:
        parser.error("no AreTomo command line given; pass it after '--'")

    parsed = parse_aretomo_argv(argv)
    if not parsed.get("input"):
        parser.error("could not find -InMrc/-InPrefix in the command line")

    output_path = args.output_override or parsed.get("output")
    if not output_path:
        parser.error("could not determine the output volume path (no "
                     "-OutMrc in the command line, and AreTomo3's -OutDir "
                     "names a directory, not a file) and no --output was "
                     "given; there is nothing to sit beside")

    aln_path = args.aln or default_aln_path(parsed)
    if not aln_path or not os.path.exists(aln_path):
        raise SystemExit(
            f"Alignment file not found: {aln_path}. AreTomo names it after "
            "-InMrc but writes it into the -OutMrc directory; pass --aln if "
            "it is somewhere else.")

    aln = parse_aln(aln_path)
    warnings = verify_aretomo_inputs(parsed["input"], parsed.get("aln_in"))
    if warnings and args.strict:
        raise SystemExit("Input verification failed (--strict): "
                         + "; ".join(warnings))

    path = record_aretomo_run(
        argv=argv, parsed=parsed, input_path=parsed["input"],
        output_path=output_path, aln_path=aln_path, aln=aln,
        variant=args.variant, binary=args.binary, warnings=warnings,
        clip_rotx=args.clip_rotx, output_pending=args.output_pending,
    )
    applied = local_applied(parsed, aln)
    print(f"Sidecar: {path}  [variant={args.variant}, "
          f"{len(aln.sections)} sections, local_applied={applied}]")
    if aln.has_local and not applied:
        print("NOTE: this run was given -AlnFile, so AreTomo did not apply the "
              "patch field to its output. Reprojection out of that volume must "
              "NOT apply it either.")


if __name__ == "__main__":
    main()
```
