## Supplementary material for "Whole cryo-FIB lamella acquisition at high magnification with rectangular beam montage cryo-electron tomography": Source code: aretomo_to_missalign_xml.py

```
"""Build a miss-alignment training-directory entry from an AreTomo2 result.

miss-alignment (github.com/warpem/miss-alignment) reads warpylib-style .xml
sidecars from its `training_directory`, each pointing at a pre-built tilt
stack under `tiltstack/<name>/<name>.st` -- it does not read AreTomo2's
.mrc/.aln output directly. This script writes that .xml + .st pair from:

  --tilt-stack  AreTomo2's *aligned tilt series* output (the -VolZ 0 run
                that reuses a solved -AlnFile, e.g. `tiltout` in
                03_AreTomo_motioncorr.sh) -- each tilt already has its
                solved shift applied, so no further per-tilt offset is
                written into the XML.
  --aln         The .aln from the same AreTomo2 solve as --tilt-stack (same
                run, or one later reused via -AlnFile). Its ROT column
                becomes tilt_axis_angles, its TILT column becomes angles.
                Row order is assumed to match the stack's Z order, which is
                how AreTomo2 writes both.

Usage:
    python -m processing_scripts.aretomo_to_missalign_xml \\
        --tilt-stack AreTomo_motioncorr/recon_patch10x6mask_bin2tiltseries.mrc \\
        --aln AreTomo_motioncorr/Montage_9-A_inpainted.aln \\
        --training-directory missalignment \\
        --volume-thickness-angstrom 13740
"""

import argparse
from pathlib import Path

import mrcfile
import torch
from warpylib import TiltSeries

from miss_alignment.data.io import TiltSeriesData


def parse_aln(aln_path: Path) -> tuple[torch.Tensor, torch.Tensor]:
    """Parse an AreTomo2 .aln file's ROT and TILT columns, in row order."""
    tilt_axis_angles, tilt_angles = [], []
    with open(aln_path) as fh:
        for line in fh:
            line = line.strip()
            if line.startswith("# Local Alignment"):
                # Per-patch local tracking rows follow, in a different
                # (shorter) column format -- not global SEC/ROT/.../TILT rows.
                break
            if not line or line.startswith("#"):
                continue
            fields = line.split()
            # SEC ROT GMAG TX TY SMEAN SFIT SCALE BASE TILT
            tilt_axis_angles.append(float(fields[1]))
            tilt_angles.append(float(fields[9]))
    return (
        torch.tensor(tilt_axis_angles, dtype=torch.float32),
        torch.tensor(tilt_angles, dtype=torch.float32),
    )


def build_xml(
    tilt_stack: Path,
    aln: Path,
    training_directory: Path,
    volume_thickness_angstrom: float,
    name: str | None = None,
) -> Path:
    name = name or tilt_stack.stem
    training_directory.mkdir(parents=True, exist_ok=True)
    xml_path = training_directory / f"{name}.xml"

    tilt_axis_angles, tilt_angles = parse_aln(aln)

    with mrcfile.open(tilt_stack, permissive=True) as mrc:
        images = torch.from_numpy(mrc.data.copy())
        pixel_size = float(mrc.voxel_size.x)

    n_tilts = images.shape[0]
    if len(tilt_angles) != n_tilts:
        raise ValueError(
            f"{aln} has {len(tilt_angles)} data rows but {tilt_stack} has "
            f"{n_tilts} tilts -- pass the .aln from the same AreTomo2 solve "
            "as this stack."
        )

    tilt_series = TiltSeries(path=str(xml_path), n_tilts=n_tilts)
    tilt_series.angles = tilt_angles
    tilt_series.tilt_axis_angles = tilt_axis_angles
    # AreTomo2's -VolZ 0 output already applied each tilt's solved TX/TY, so
    # the per-tilt offset miss-alignment would otherwise apply stays at zero.
    tilt_series.tilt_axis_offset_x = torch.zeros(n_tilts, dtype=torch.float32)
    tilt_series.tilt_axis_offset_y = torch.zeros(n_tilts, dtype=torch.float32)

    ny, nx = images.shape[-2], images.shape[-1]
    tilt_series.image_dimensions_physical = torch.tensor(
        [nx * pixel_size, ny * pixel_size], dtype=torch.float32
    )
    volume_z_px = round(volume_thickness_angstrom / pixel_size)
    tilt_series.volume_dimensions_physical = torch.tensor(
        [nx * pixel_size, ny * pixel_size, volume_z_px * pixel_size],
        dtype=torch.float32,
    )

    tilt_series_data = TiltSeriesData(xml_metadata_path=xml_path)
    stack_path = Path(tilt_series.tilt_stack_path)
    stack_path.parent.mkdir(parents=True, exist_ok=True)
    with mrcfile.new(stack_path, overwrite=True) as mrc:
        mrc.set_data(images.numpy())
        mrc.voxel_size = pixel_size

    tilt_series_data.save_metadata_to_xml(tilt_series)

    print(f"Wrote {xml_path}")
    print(
        f"Wrote {stack_path} ({n_tilts} tilts, {nx}x{ny}px @ {pixel_size:.3f} A/px, "
        f"volume Z {volume_z_px}px)"
    )
    return xml_path


def main() -> None:
    parser = argparse.ArgumentParser(
        description=__doc__, formatter_class=argparse.RawDescriptionHelpFormatter
    )
    parser.add_argument(
        "--tilt-stack",
        required=True,
        type=Path,
        help="AreTomo2 aligned tilt series (-VolZ 0 output .mrc)",
    )
    parser.add_argument(
        "--aln",
        required=True,
        type=Path,
        help="AreTomo2 .aln from the same alignment solve as --tilt-stack",
    )
    parser.add_argument(
        "--training-directory",
        required=True,
        type=Path,
        help="miss-alignment general.training_directory",
    )
    parser.add_argument(
        "--volume-thickness-angstrom",
        required=True,
        type=float,
        help=(
            "Physical Z thickness of the reconstruction volume used for "
            "training, in Angstroms (e.g. AreTomo2's -VolZ in unbinned "
            "pixels x its unbinned pixel size)"
        ),
    )
    parser.add_argument(
        "--name", default=None, help="tilt series name (default: --tilt-stack stem)"
    )
    args = parser.parse_args()

    build_xml(
        tilt_stack=args.tilt_stack,
        aln=args.aln,
        training_directory=args.training_directory,
        volume_thickness_angstrom=args.volume_thickness_angstrom,
        name=args.name,
    )


if __name__ == "__main__":
    main()
```
