## Supplementary material for "Whole cryo-FIB lamella acquisition at high magnification with rectangular beam montage cryo-electron tomography": Source code: beam_mask_motioncorr.py

```
#!/usr/bin/env python3
"""Motion correction for square-beam montage raw frames.

Workflow
--------
1. Load a multi-frame TIFF
2. Auto-detect the square beam mask from the mean frame
3. Find the largest axis-aligned square inscribed in the mask
4. Crop all sub-frames to that square, write a temporary TIFF
5. Run MotionCor2 on the cropped TIFF to estimate per-frame shifts
6. Parse per-frame shifts from the MotionCor2 log
7. Run ctffind4 on the same motion-corrected crop to estimate the CTF
8. Apply the same shifts (Fourier shift) to the original full-size frames
9. Phase-flip the shifted sum using the crop-estimated CTF
10. Sum the shifted frames → write float32 MRC (single 2-D slice)

The crop is used ONLY for motion/CTF estimation; the output MRC covers the
full detector area, so downstream stitching is unaffected.

CTF estimation and phase-flipping
----------------------------------
By default, ctffind4 estimates the defocus/astigmatism of each tile from
its motion-corrected crop and the fit is logged/saved, but the output
image is left untouched — useful for downstream tools, e.g. IMOD's
``ctfphaseflip``. Pass ``--ctf-flip`` to also phase-flip the full-frame
output before it's written, which only happens when the fit's score is
at least ``--ctf-min-score`` (default 0.1); a low-confidence fit is worse
than none, since flipping with the wrong defocus scrambles real signal.
Tiles below the threshold are left un-flipped but still estimated/logged/
saved. Pass ``--no-ctffind`` to skip CTF estimation entirely (also
disables phase-flipping).

Every tile's fit is appended as a row to a shared ``{output_dir}/ctf_results.txt``
(tile name, defocus1/2, azimuth, phase shift, score, fit resolution — safe
to append to concurrently from multiple SLURM array tasks). With
``--save-diagnostic``, a matching ``{stem}_ctf_diagnostic.png`` is also
written next to each tile's mask/crop diagnostic, showing ctffind4's
amplitude-spectrum fit and the same numeric parameters.

``--save-diagnostic`` also writes a ``{stem}_motion_diagnostic.png`` showing
the accumulated MotionCor2 shift trajectory and the FFT power spectrum of
the crop region before vs. after motion correction — a tighter, more
extended spectrum after correction indicates less residual motion blur.

Output modes
------------
By default each tile is written as its own single-slice MRC file, named
``{stem}.mrc`` (or split into ``odd/`` and ``even/`` subdirectories with
``--split-frames``). This is the layout ``stitch.py`` reads via its
"directory of per-tile MRC files" input mode.

Passing ``--stack-output`` instead accumulates every tile belonging to the
same tilt angle into a single memory-mapped MRC stack (one 2-D section per
tile), matching the layout of a raw SerialEM montage stack (e.g.
``Montage_182-A_0.0.mrc``) that ``stitch.py`` reads via its "glob of
per-tilt MRC stacks" input mode. This requires input filenames to follow
the ``{base}_{tile_index}_{tilt_angle}.tif`` convention. It is safe to call
from multiple concurrent SLURM array tasks (one task per tile): the stack
file is created once under an flock, and each task subsequently writes
only its own z-slice.

Provenance sidecars
-------------------
Every tile writes a small JSON sidecar recording what this step actually did,
rather than leaving later steps to infer it (see
``processing_scripts/sidecar.py`` and ``sidecar_metadata_plan.md``). Each
sidecar carries:

* the step's *effective* parameters — after auto-detection of the MotionCor2
  and ctffind4 executables, after ``--ctf-max-res``'s pixel-size-derived
  default, after the beam-mask threshold default;
* that tile's own results: the beam mask and crop actually used, the shifts
  actually applied, the CTF fit, whether the tile was phase-flipped, and —
  under ``--stack-output`` — the tilt angle, the cumulative tile index, the
  base that index was re-referenced from, and the z-slice it landed on. That
  last group is the mapping ``fix_stack_tile_index.py`` exists to reconstruct
  after the fact.

Under ``--stack-output`` many tasks write z-slices of one shared per-tilt
stack, so each writes its own keyed sidecar at
``<stack>.mrc.sidecars/<tile stem>.json``; with one MRC per tile the sidecar
is simply ``<tile>.mrc.json``. Either way no two array tasks ever open the
same path, so there is no lock, no spool and no finalize job — pass
``--no-sidecar`` to skip them entirely.
"""

from __future__ import annotations

import argparse
import fcntl
import glob
import logging
import os
import re
import shutil
import subprocess
import sys
import tempfile
from collections.abc import Sequence
from pathlib import Path

import time

import mrcfile
import numpy as np
from PIL import Image
from tqdm import tqdm

from .Utilities import make_mask
from . import sidecar
from .sidecar import directory_input, fingerprint

log = logging.getLogger(__name__)

SIDECAR_STEP = "motion_correction"

# ---------------------------------------------------------------------------
# TIFF I/O
# ---------------------------------------------------------------------------

def load_multipage_tiff(path: str) -> np.ndarray:
    """Return (n_frames, H, W) uint8/uint16 array from a multi-page TIFF."""

    # Open the TIFF with Pillow, which handles multi-page TIFFs
    with Image.open(path) as img:
        # Get number of frames in tiff
        n = getattr(img, "n_frames", 1)
        # Seek to the first frame and read it to get shape and dtype
        img.seek(0)
        first = np.asarray(img.copy())
        # Initialise output
        out = np.empty((n, *first.shape), dtype=first.dtype)
        # Load first frame to output
        out[0] = first
        # Load remaining frames
        for i in range(1, n):
            img.seek(i)
            out[i] = np.asarray(img.copy())
    return out


def write_multipage_tiff(frames: np.ndarray, path: str) -> None:
    """Write (n_frames, H, W) array as a multi-page TIFF."""
    # Convert each 2-D frame to its own Pillow image
    pil_frames = [Image.fromarray(f) for f in frames]
    # Save the first frame, appending the rest as additional pages
    pil_frames[0].save(
        path,
        save_all=True,
        append_images=pil_frames[1:],
        compression="tiff_lzw",
    )


# ---------------------------------------------------------------------------
# Beam mask + largest inscribed square
# ---------------------------------------------------------------------------

def largest_inscribed_square(mask: np.ndarray) -> tuple[int, int, int]:
    """Find the largest axis-aligned square with all pixels inside ``mask``.

    Uses the classic DP (dynamic programming) histogram approach: build a
    table where each cell holds the side length of the largest all-inside
    square ending at that cell, growing each entry from the three
    neighbours already solved above, to the left, and diagonally above-left.

    Returns
    -------
    (row_top, col_left, side)  — corner of the square and its side length.
    """
    h, w = mask.shape
    # dp[r, c] = side length of the largest all-inside square whose
    # bottom-right corner is (r, c)
    dp = np.zeros((h, w), dtype=np.int32)
    # First row/column can only extend a square of side 0 or 1
    dp[0, :] = mask[0, :].astype(np.int32)
    dp[:, 0] = mask[:, 0].astype(np.int32)

    # Grow each square from its three neighbours (above, left, diagonal)
    for r in range(1, h):
        for c in range(1, w):
            if mask[r, c]:
                dp[r, c] = int(min(dp[r - 1, c], dp[r, c - 1], dp[r - 1, c - 1])) + 1

    # Largest value in dp is the biggest square found; recover its corner
    idx = int(np.argmax(dp))
    r_br, c_br = divmod(idx, w)
    side = int(dp[r_br, c_br])
    row_top = r_br - side + 1
    col_left = c_br - side + 1
    return row_top, col_left, side


def _fft_friendly_size(n: int) -> int:
    """Largest k ≤ n whose prime factors are all in {2, 3, 5, 7} (CUFFT fast path).
    """
    _CUFFT_PRIMES = (2, 3, 5, 7)
    k = n
    # Walk sizes down from n until one factors entirely into 2/3/5/7
    while k > 1:
        m = k
        for p in _CUFFT_PRIMES:
            while m % p == 0:
                m //= p
        if m == 1:
            return k
        k -= 1
    return 1


# ---------------------------------------------------------------------------
# MotionCor2 interaction
# ---------------------------------------------------------------------------

def _find_motioncor2() -> str:
    """Return path to MotionCor2 executable, searching common locations."""
    # Try the bare command first (works if sbgrid/module has already put it
    # on PATH), then fall back to known install locations
    candidates = [
        "MotionCor2",
        "/programs/x86_64-linux/motioncor2/1.6.4/MotionCor2",
        "/programs/x86_64-linux/motioncor2/1.3.2/MotionCor2",
    ]
    for c in candidates:
        if shutil.which(c):
            return c
    raise FileNotFoundError(
        "MotionCor2 not found. Provide --motioncor2 or load it via sbgrid."
    )


def run_motioncor2(
    in_tiff: str,
    out_mrc: str,
    log_dir: str,
    pixel_size: float,
    bfactor: float,
    gpu: int,
    motioncor2_path: str,
    extra_args: list[str] | None = None,
) -> str:
    """Run MotionCor2 on ``in_tiff``, return captured stdout text."""
    # Build the base MotionCor2 CLI invocation; -OutStack 0 means "don't also
    # write the aligned stack", we only need the shift table it prints
    cmd = [
        motioncor2_path,
        "-InTiff", in_tiff,
        "-OutMrc", out_mrc,
        "-PixSize", str(pixel_size),
        "-Bft", str(bfactor),
        "-Gpu", str(gpu),
        "-LogDir", log_dir,
        "-OutStack", "0",
    ]
    if extra_args:
        cmd.extend(extra_args)

    log.debug("Running: %s", " ".join(cmd))
    result = subprocess.run(cmd, capture_output=True, text=True, check=False)
    if result.returncode != 0:
        # Log the tail of stdout/stderr rather than the full text -- MotionCor2
        # output can be long, and the failure reason is always near the end
        log.error("MotionCor2 stdout:\n%s", result.stdout[-2000:])
        log.error("MotionCor2 stderr:\n%s", result.stderr[-2000:])
        raise RuntimeError(
            f"MotionCor2 failed with exit code {result.returncode}."
        )
    return result.stdout


def parse_motioncor2_shifts(text: str, n_frames: int) -> np.ndarray:
    """Parse per-frame shifts from MotionCor2 stdout or log file text.

    MotionCor2 prints a table like::

        Full-frame alignment
          Frame   x Shift   y Shift
            1      0.000      0.000
            2      0.950      0.030

    Indices may be 0- or 1-based depending on version; both are handled.

    Returns
    -------
    shifts : (n_frames, 2) float32 array, columns = (shift_x, shift_y) in pixels.
    """
    # Matches a table row: leading frame index, then two signed floats
    pattern = re.compile(
        r"^\s*(\d+)\s+([+-]?\d+\.\d+)\s+([+-]?\d+\.\d+)\s*$"
    )
    found: dict[int, tuple[float, float]] = {}
    for line in text.splitlines():
        m = pattern.match(line)
        if m:
            found[int(m.group(1))] = (float(m.group(2)), float(m.group(3)))

    if not found:
        raise ValueError(
            "No shift lines found in MotionCor2 output.\n"
            f"Output was:\n{text[-2000:]}"
        )

    # Handle 0-based or 1-based frame indexing
    offset = min(found.keys())
    shifts = np.zeros((n_frames, 2), dtype=np.float32)
    for i in range(n_frames):
        key = i + offset
        if key in found:
            shifts[i, 0] = found[key][0]
            shifts[i, 1] = found[key][1]
        else:
            log.warning("Frame %d shift not found in output; using 0,0.", i)
    return shifts


# ---------------------------------------------------------------------------
# ctffind4 interaction
# ---------------------------------------------------------------------------

def _find_ctffind4() -> str:
    """Return path to the ctffind4 executable, searching common locations."""
    # Some installs name the binary "ctffind4", others just "ctffind"
    candidates = ["ctffind4", "ctffind"]
    for c in candidates:
        if shutil.which(c):
            return c
    raise FileNotFoundError(
        "ctffind4 not found. Provide --ctffind4 or load it via sbgrid."
    )


def run_ctffind4(
    in_mrc: str,
    diag_mrc: str,
    pixel_size: float,
    voltage_kv: float,
    cs_mm: float,
    amp_contrast: float,
    ctffind4_path: str,
    box_size: int = 512,
    min_res: float = 30.0,
    max_res: float = 5.0,
    min_defocus: float = 5000.0,
    max_defocus: float = 200000.0,
    defocus_step: float = 1000.0,
) -> str:
    """Run ctffind4 on ``in_mrc`` (a single 2-D micrograph); return path to
    the summary results ``.txt`` file it writes next to ``diag_mrc``.

    ctffind4 (v4.1.x) takes all parameters as an ordered sequence of
    answers piped to stdin ("scripted mode") rather than CLI flags; this
    reproduces that exact prompt order. Astigmatism/phase-shift search are
    left on ctffind4's own defaults (searched, unrestrained, no phase-plate
    search) and expert options are skipped.
    """
    # Build the answer sequence in the exact order ctffind4 prompts for it
    answers = "\n".join([
        in_mrc,
        diag_mrc,
        str(pixel_size),
        str(voltage_kv),
        str(cs_mm),
        str(amp_contrast),
        str(box_size),
        str(min_res),
        str(max_res),
        str(min_defocus),
        str(max_defocus),
        str(defocus_step),
        "no",   # Do you know what astigmatism is present?
        "no",   # Slower, more exhaustive search?
        "no",   # Use a restraint on astigmatism?
        "no",   # Find additional phase shift?
        "no",   # Do you want to set expert options?
        "",
    ])
    log.debug("Running ctffind4 on %s", in_mrc)
    result = subprocess.run(
        [ctffind4_path], input=answers, capture_output=True, text=True, check=False,
    )
    if result.returncode != 0:
        log.error("ctffind4 stdout:\n%s", result.stdout[-2000:])
        log.error("ctffind4 stderr:\n%s", result.stderr[-2000:])
        raise RuntimeError(f"ctffind4 failed with exit code {result.returncode}.")

    txt_path = os.path.splitext(diag_mrc)[0] + ".txt"
    if not os.path.exists(txt_path):
        raise RuntimeError(f"ctffind4 did not produce expected results file: {txt_path}")
    return txt_path


def parse_ctffind4_result(txt_path: str) -> dict:
    """Parse the data line of a ctffind4 summary ``.txt`` file.

    Returns a dict with keys ``defocus1``, ``defocus2`` (Angstrom),
    ``azimuth`` (degrees), ``phase_shift`` (radians), ``score``, and
    ``fit_resolution`` (Angstrom) — see ctffind4's own column header
    comment for the source ordering.
    """
    # The file is mostly '#'-prefixed comment lines; the data itself is the
    # single trailing non-comment line
    last_line = None
    with open(txt_path) as fh:
        for line in fh:
            line = line.strip()
            if line and not line.startswith("#"):
                last_line = line
    if last_line is None:
        raise ValueError(f"No CTF result line found in {txt_path}")

    # First column is a running micrograph index, which we don't need
    _, df1, df2, azimuth, phase_shift, score, fit_res = (
        float(x) for x in last_line.split()
    )
    return {
        "defocus1": df1,
        "defocus2": df2,
        "azimuth": azimuth,
        "phase_shift": phase_shift,
        "score": score,
        "fit_resolution": fit_res,
    }


_CTF_RESULTS_HEADER = (
    "# tile\tdefocus1[A]\tdefocus2[A]\tazimuth[deg]\t"
    "phase_shift[rad]\tscore\tfit_resolution[A]\n"
)


def append_ctf_result(results_path: str, tile_name: str, ctf_result: dict) -> None:
    """Append one tile's CTF fit as a row to a shared per-run results file.

    Guarded by an flock on the file itself so concurrent SLURM array tasks
    appending to the same shared file (e.g. one task per tile batch) don't
    interleave lines.
    """
    os.makedirs(os.path.dirname(results_path) or ".", exist_ok=True)
    with open(results_path, "a") as fh:
        # Block until we hold the exclusive lock, so no other task's row
        # can interleave with ours between the size check and the write
        fcntl.flock(fh, fcntl.LOCK_EX)
        try:
            # First writer creates the file, so write the header only once
            if os.fstat(fh.fileno()).st_size == 0:
                fh.write(_CTF_RESULTS_HEADER)
            fh.write(
                f"{tile_name}\t{ctf_result['defocus1']:.2f}\t"
                f"{ctf_result['defocus2']:.2f}\t{ctf_result['azimuth']:.2f}\t"
                f"{ctf_result['phase_shift']:.4f}\t{ctf_result['score']:.5f}\t"
                f"{ctf_result['fit_resolution']:.2f}\n"
            )
        finally:
            fcntl.flock(fh, fcntl.LOCK_UN)


# ---------------------------------------------------------------------------
# Per-tilt memory-mapped MRC stack output
# ---------------------------------------------------------------------------

# Matches tile filenames like  Montage_182-A_182-A_003_-12.0  (stem, no ext)
# -> groups (base, tile_index, tilt_angle)
_TILE_FNAME_RE = re.compile(r"^(.+)_(\d+)_(-?\d+\.?\d*)$")


def parse_tile_filename(path: str) -> tuple[str, int, float]:
    """Parse ``{base}_{tile_index}_{tilt_angle}`` from a tile file's stem.

    Returns (base, tile_index, tilt_angle); -0.0 is normalised to 0.0 so
    tiles at zero tilt group together regardless of sign.
    """
    stem = Path(path).stem
    m = _TILE_FNAME_RE.match(stem)
    if not m:
        raise ValueError(
            f"'{path}' does not match the required "
            "'{{base}}_{{tile_index}}_{{tilt_angle}}' naming convention "
            "for --stack-output."
        )
    base, tile_idx, tilt = m.group(1), int(m.group(2)), float(m.group(3))
    # Fold negative-zero tilt onto positive zero so both group together
    if tilt == -0.0:
        tilt = 0.0
    return base, tile_idx, tilt


def stack_path_for(output_dir: str, base: str, tilt: float) -> str:
    """Path of the shared per-tilt MRC stack for a given base name/tilt."""
    return os.path.join(output_dir, f"{base}_{tilt:g}.mrc")


def tile_index_range_for_tilt(input_path: str, base: str, tilt: float) -> tuple[int, int]:
    """Return (min_idx, max_idx) of tile indices sharing ``base``/``tilt``.

    ``tile_index`` in the ``{base}_{tile_index}_{tilt_angle}`` naming
    convention is a cumulative counter across the *entire* tilt series — it
    does not restart at each tilt — so a given tilt's tiles typically occupy
    some contiguous sub-range like 960-1065 rather than 0-105. Callers
    should subtract ``min_idx`` from a tile's raw index to get its 0-based
    z-slice position within that tilt's stack.

    Only inspects ``input_path``'s own directory (no recursion — cheap even
    on a shared filesystem).
    """
    directory = os.path.dirname(os.path.abspath(input_path)) or "."
    ext = os.path.splitext(input_path)[1]
    min_idx, max_idx = None, None
    # Scan siblings in the same directory, keeping only those that parse and
    # share this tile's base/tilt, and track the index range they span
    for sibling in glob.glob(os.path.join(directory, f"*{ext}")):
        try:
            sib_base, sib_idx, sib_tilt = parse_tile_filename(sibling)
        except ValueError:
            continue
        if sib_base == base and sib_tilt == tilt:
            min_idx = sib_idx if min_idx is None else min(min_idx, sib_idx)
            max_idx = sib_idx if max_idx is None else max(max_idx, sib_idx)
    if min_idx is None:
        raise ValueError(
            f"No sibling tiles found matching base='{base}' tilt={tilt} "
            f"next to {input_path}"
        )
    return min_idx, max_idx


def _check_stack_shape(path: str, shape: tuple[int, int, int]) -> None:
    """Refuse to write into a stack left over from a run of a different shape.

    Re-running with a different ``--rotate`` (or a different tile count) over
    an output directory that still holds the previous run's stacks used to
    surface as a numpy broadcast error per tile, several hundred times, while
    the array task itself exited 0 — a whole run that looked COMPLETED and
    wrote nothing. Say what is actually wrong, and what to do about it.
    """
    # Header-only open is cheap even for a huge memory-mapped stack
    with mrcfile.open(path, mode="r", header_only=True, permissive=True) as mrc:
        header = mrc.header
        existing = (int(header.nz), int(header.ny), int(header.nx))
    if existing == tuple(int(n) for n in shape):
        return
    raise ValueError(
        f"{path} already exists with shape (z,y,x)={existing}, but this run "
        f"produces {tuple(int(n) for n in shape)}. It is from an earlier run "
        "with different parameters — a different --rotate or a different "
        "number of tiles in this tilt. Delete the stacks in this output "
        "directory (and their .stack_markers) or write to a new one."
    )


def _check_stack_pixel_size(path: str, pixel_size: float) -> None:
    """Refuse to write into a stack left over from a run with a different pixel size.

    voxel_size is only set when a stack is first created (below); a resumed
    run with a corrected --pixel-size would otherwise silently keep writing
    tiles into a header that still reports the old value, with no error, no
    warning -- just a wrong pixel size baked into some fraction of the
    stack's slices depending on which run created the file first.
    """
    with mrcfile.open(path, mode="r", header_only=True, permissive=True) as mrc:
        existing = float(mrc.voxel_size.x)
    # Small tolerance for float round-tripping through the MRC header
    if abs(existing - pixel_size) <= 1e-3:
        return
    raise ValueError(
        f"{path} already exists with pixel size {existing:.4f} A, but this "
        f"run requests --pixel-size {pixel_size:.4f}. It is from an earlier "
        "run with a different --pixel-size. Delete the stacks in this "
        "output directory (and their .stack_markers) or write to a new one."
    )


def _create_stack_locked(
    path: str, shape: tuple[int, int, int], pixel_size: float, mrc_mode: int = 2
) -> None:
    """Create an empty MRC stack at ``path`` if it doesn't exist yet.

    ``mrc_mode`` follows the mrcfile convention (2 = float32, the default;
    1 = int16; 0 = int8, e.g. for binary mask stacks).

    Guarded by an flock on a sibling ``.lock`` file so concurrent SLURM
    array tasks racing to create the same stack don't step on each other;
    only the task that wins the lock creates the file, everyone else opens
    the resulting file afterwards. Requires a filesystem with working POSIX
    advisory locks (fine on local/GPFS mounts; unreliable on some NFS setups).
    """
    if os.path.exists(path):
        # Someone already created it -- just make sure it matches this run
        _check_stack_shape(path, shape)
        _check_stack_pixel_size(path, pixel_size)
        return
    os.makedirs(os.path.dirname(path) or ".", exist_ok=True)
    lock_path = path + ".lock"
    with open(lock_path, "w") as lockfh:
        fcntl.flock(lockfh, fcntl.LOCK_EX)
        try:
            # Re-check inside the lock: another task may have created the
            # stack between our check above and acquiring the lock
            if not os.path.exists(path):
                with mrcfile.new_mmap(path, shape=shape, mrc_mode=mrc_mode, overwrite=False) as mrc:
                    mrc.voxel_size = pixel_size
                log.info("Created MRC stack %s  shape=%s  mode=%d", path, shape, mrc_mode)
        finally:
            fcntl.flock(lockfh, fcntl.LOCK_UN)


def write_tile_slice(
    stack_path: str,
    tile_idx: int,
    n_tiles: int,
    data: np.ndarray,
    pixel_size: float,
) -> None:
    """Write ``data`` into z-slice ``tile_idx`` of a shared MRC stack.

    Creates the stack — sized ``(n_tiles, *data.shape)`` — on first use.
    Safe to call concurrently from multiple processes as long as each call
    uses a distinct ``tile_idx``: writes land on disjoint byte ranges of
    the memory-mapped file.
    """
    # Creates the stack on first use; a no-op (after its own checks) once it
    # already exists
    _create_stack_locked(stack_path, (n_tiles, *data.shape), pixel_size)
    with mrcfile.mmap(stack_path, mode="r+") as mrc:
        mrc.data[tile_idx] = data.astype(np.float32, copy=False)


# ---------------------------------------------------------------------------
# Shift application
# ---------------------------------------------------------------------------

def _fourier_shift_2d(frame: np.ndarray, dy: float, dx: float) -> np.ndarray:
    """Apply subpixel shift (dy, dx) to a 2-D array via the Fourier shift theorem."""
    h, w = frame.shape
    # Spatial frequency grids for the real FFT
    fy = np.fft.fftfreq(h)
    fx = np.fft.rfftfreq(w)
    # A linear phase ramp in frequency space is a shift in real space
    phase = np.exp(-2j * np.pi * (dy * fy[:, None] + dx * fx[None, :]))
    f_fft = np.fft.rfft2(frame.astype(np.float32))
    shifted = np.fft.irfft2(f_fft * phase, s=frame.shape)
    return shifted.astype(np.float32)


def apply_shifts_and_sum(
    frames: np.ndarray,
    shifts_xy: np.ndarray,
) -> np.ndarray:
    """Shift each frame by (dx, dy) and return their sum.

    Parameters
    ----------
    frames : (n_frames, H, W) array
    shifts_xy : (n_frames, 2) — column 0 = shift_x, column 1 = shift_y

    Returns
    -------
    (H, W) float32 sum of aligned frames.
    """
    n = frames.shape[0]
    result = np.zeros(frames.shape[1:], dtype=np.float32)
    # Shift and accumulate one frame at a time to avoid holding a second
    # full (n_frames, H, W) array in memory
    for i in range(n):
        dx = float(shifts_xy[i, 0])
        dy = float(shifts_xy[i, 1])
        shifted = _fourier_shift_2d(frames[i].astype(np.float32), dy, dx)
        result += shifted
    return result


def apply_shifts_split(
    frames: np.ndarray,
    shifts_xy: np.ndarray,
) -> tuple[np.ndarray, np.ndarray, np.ndarray]:
    """Shift frames and return (full_sum, odd_sum, even_sum) in a single pass.

    odd_sum  accumulates frames at 0-based indices 0, 2, 4, …
    even_sum accumulates frames at 0-based indices 1, 3, 5, …

    All three sums use the same per-frame Fourier shifts, so no extra FFTs
    are computed compared to a plain sum.
    """
    n = frames.shape[0]
    full_sum = np.zeros(frames.shape[1:], dtype=np.float32)
    odd_sum  = np.zeros(frames.shape[1:], dtype=np.float32)
    even_sum = np.zeros(frames.shape[1:], dtype=np.float32)
    for i in range(n):
        dx = float(shifts_xy[i, 0])
        dy = float(shifts_xy[i, 1])
        # One FFT-based shift per frame, reused for all three running sums
        shifted = _fourier_shift_2d(frames[i].astype(np.float32), dy, dx)
        full_sum += shifted
        if i % 2 == 0:
            odd_sum += shifted
        else:
            even_sum += shifted
    return full_sum, odd_sum, even_sum


# ---------------------------------------------------------------------------
# CTF phase-flipping
# ---------------------------------------------------------------------------

def _electron_wavelength_A(voltage_kv: float) -> float:
    """Relativistic electron wavelength in Angstrom for a given accelerating voltage."""
    v = voltage_kv * 1000.0
    return 12.2639 / np.sqrt(v * (1.0 + 0.97845e-6 * v))


def compute_ctf(
    freq_y: np.ndarray,
    freq_x: np.ndarray,
    defocus1: float,
    defocus2: float,
    azimuth_deg: float,
    voltage_kv: float,
    cs_mm: float,
    amp_contrast: float,
    phase_shift_rad: float = 0.0,
) -> np.ndarray:
    """Evaluate the (astigmatic) CTF over a grid of spatial frequencies.

    ``freq_y``/``freq_x`` are in 1/Angstrom (broadcastable to a common
    shape); ``defocus1``/``defocus2`` in Angstrom, ``azimuth_deg`` in
    degrees — matching ctffind4's output convention.
    """
    wavelength = _electron_wavelength_A(voltage_kv)
    cs_A = cs_mm * 1e7  # mm -> Angstrom
    k2 = freq_x ** 2 + freq_y ** 2
    phi = np.arctan2(freq_y, freq_x)
    az = np.deg2rad(azimuth_deg)
    # Defocus at each spatial-frequency direction, interpolating between the
    # two principal defoci according to the astigmatism azimuth
    dz = (
        0.5 * (defocus1 + defocus2)
        + 0.5 * (defocus1 - defocus2) * np.cos(2.0 * (phi - az))
    )
    # Aberration phase (defocus term minus spherical-aberration term), plus
    # any extra phase plate shift
    chi = (
        np.pi * wavelength * k2 * (dz - 0.5 * cs_A * wavelength ** 2 * k2)
        + phase_shift_rad
    )
    ac = float(np.clip(amp_contrast, -1.0, 1.0))
    w2 = float(np.sqrt(max(1.0 - ac ** 2, 0.0)))
    # Mixed amplitude/phase-contrast CTF
    return -(w2 * np.sin(chi) + ac * np.cos(chi))


def ctf_phase_flip(
    image: np.ndarray,
    pixel_size: float,
    defocus1: float,
    defocus2: float,
    azimuth_deg: float,
    voltage_kv: float,
    cs_mm: float,
    amp_contrast: float,
    phase_shift_rad: float = 0.0,
) -> np.ndarray:
    """Flip the sign of Fourier components where the CTF is negative.

    This is a simple global phase-flip (single defocus/astigmatism value
    applied across the whole image) — it corrects contrast reversals but
    does not correct the amplitude falloff itself.
    """
    h, w = image.shape
    # Spatial frequency grids in 1/Angstrom, matching compute_ctf's convention
    fy = np.fft.fftfreq(h, d=pixel_size)
    fx = np.fft.rfftfreq(w, d=pixel_size)
    ctf = compute_ctf(
        fy[:, None], fx[None, :], defocus1, defocus2, azimuth_deg,
        voltage_kv, cs_mm, amp_contrast, phase_shift_rad,
    )
    # +1 where the CTF is positive, -1 where it's negative -- flipping the
    # sign of Fourier components in the negative lobes
    sign = -np.where(ctf < 0, -1.0, 1.0).astype(np.float32)
    f = np.fft.rfft2(image.astype(np.float32))
    flipped = np.fft.irfft2(f * sign, s=image.shape)
    return flipped.astype(np.float32)


# ---------------------------------------------------------------------------
# Diagnostic plot
# ---------------------------------------------------------------------------

def _block_bin(image: np.ndarray, b: int) -> np.ndarray:
    """Downsample by block-averaging rather than strided decimation.

    Strided slicing (``image[::b, ::b]``) just picks every b-th pixel, which
    keeps the same per-pixel noise. Averaging b×b blocks instead reduces
    noise by roughly sqrt(b*b), which is what "binning" is supposed to do.
    """
    if b <= 1:
        return image
    h, w = image.shape
    # Trim to a multiple of b so the reshape below divides evenly
    h2, w2 = h - h % b, w - w % b
    trimmed = image[:h2, :w2]
    # Reshape into b×b blocks and average over each block
    return trimmed.reshape(h2 // b, b, w2 // b, b).mean(axis=(1, 3))


def save_diagnostic_plot(
    raw_frame: np.ndarray,
    corrected_frame: np.ndarray,
    mask: np.ndarray,
    row_top: int,
    col_left: int,
    side: int,
    output_path: str,
    binning: int = 4,
) -> None:
    """Save a PNG showing the beam mask (on the raw frame) and the crop
    region (on the full-frame motion-corrected sum)."""
    import matplotlib
    matplotlib.use("Agg")
    import matplotlib.pyplot as plt
    import matplotlib.patches as mpatches

    b = max(1, binning)
    raw_img = _block_bin(raw_frame, b)
    corr_img = _block_bin(corrected_frame, b)
    h2, w2 = raw_img.shape[0] * b, raw_img.shape[1] * b
    # Subsample the mask/crop coordinates to match the binned images
    msk = mask[:h2:b, :w2:b]
    r, c, s = row_top // b, col_left // b, max(1, side // b)

    # Percentile-based display range so a few hot/cold outlier pixels don't
    # wash out the contrast
    raw_vmin, raw_vmax = np.percentile(raw_img, [1, 99])
    corr_vmin, corr_vmax = np.percentile(corr_img, [1, 99])

    fig, axes = plt.subplots(1, 2, figsize=(14, 5), constrained_layout=True)
    fig.suptitle(os.path.basename(output_path).replace("_diagnostic.png", ""), fontsize=10)

    # Left: mask overlay on the raw (uncorrected) frame
    axes[0].imshow(raw_img, cmap="gray", vmin=raw_vmin, vmax=raw_vmax,
                    interpolation="nearest")
    # NaN inside the mask makes those pixels transparent in the overlay
    outside = np.where(~msk, 1.0, np.nan)
    axes[0].imshow(outside, cmap="Reds", alpha=0.5, vmin=0, vmax=1,
                   interpolation="nearest")
    axes[0].set_title("Beam mask  (red = excluded)", fontsize=9)
    axes[0].axis("off")

    # Right: crop rectangle on the motion-corrected frame
    axes[1].imshow(corr_img, cmap="gray", vmin=corr_vmin, vmax=corr_vmax,
                    interpolation="nearest")
    rect = mpatches.Rectangle(
        (c, r), s, s,
        linewidth=1.5, edgecolor="lime", facecolor="none",
    )
    axes[1].add_patch(rect)
    axes[1].set_title(
        f"Crop region: {side}×{side} px (unbinned), motion-corrected\n"
        f"origin row={row_top} col={col_left}",
        fontsize=9,
    )
    axes[1].axis("off")

    plt.savefig(output_path, dpi=100)
    plt.close(fig)
    log.info("Diagnostic plot: %s", output_path)


def save_ctf_diagnostic_plot(
    diag_mrc_path: str,
    ctf_result: dict,
    output_path: str,
) -> None:
    """Save a PNG of ctffind4's amplitude-spectrum diagnostic plus fit parameters."""
    import matplotlib
    matplotlib.use("Agg")
    import matplotlib.pyplot as plt

    with mrcfile.open(diag_mrc_path, permissive=True) as mrc:
        spectrum = np.asarray(mrc.data)
    # ctffind4 sometimes writes a single-slice stack rather than a 2-D image
    if spectrum.ndim == 3:
        spectrum = spectrum[0]

    vmin = float(np.percentile(spectrum, 1))
    vmax = float(np.percentile(spectrum, 99))

    fig, ax = plt.subplots(1, 1, figsize=(6, 6))
    # Reserve room below the image for the 7-line summary block -- the
    # default subplot margin is too small for it at fontsize 8, and fig.text
    # at y=0.02 ends up overlapping the bottom of the image instead of
    # sitting under it.
    plt.subplots_adjust(bottom=0.22)
    fig.suptitle(
        os.path.basename(output_path).replace("_ctf_diagnostic.png", ""), fontsize=10
    )
    ax.imshow(spectrum, cmap="gray", vmin=vmin, vmax=vmax, interpolation="nearest")
    ax.set_title("ctffind4 amplitude spectrum  (half fit / half data)", fontsize=8)
    ax.axis("off")

    astigmatism = ctf_result["defocus1"] - ctf_result["defocus2"]
    # Fixed-width numeric columns so the block reads as a table in monospace
    summary = (
        f"defocus1      {ctf_result['defocus1']:>10.0f} A\n"
        f"defocus2      {ctf_result['defocus2']:>10.0f} A\n"
        f"astigmatism   {astigmatism:>10.0f} A\n"
        f"azimuth       {ctf_result['azimuth']:>10.1f} deg\n"
        f"phase shift   {ctf_result['phase_shift']:>10.2f} rad\n"
        f"score         {ctf_result['score']:>10.3f}\n"
        f"fit res.      {ctf_result['fit_resolution']:>10.1f} A"
    )
    fig.text(0.02, 0.02, summary, fontsize=8, family="monospace", va="bottom", ha="left")

    plt.savefig(output_path, dpi=100)
    plt.close(fig)
    log.info("CTF diagnostic plot: %s", output_path)


def _log_power_spectrum(image: np.ndarray) -> np.ndarray:
    """Return the log-scaled, zero-centred power spectrum of a 2-D image.

    The image is mean-subtracted and tapered with a 2-D Hann window before
    the FFT; without this, the crop's sharp rectangular edges dominate the
    spectrum with a bright cross-shaped streak through the origin, and the
    huge DC/near-DC term swamps everything else once averaged into a shared
    display range.
    """
    img = image.astype(np.float64)
    img = img - img.mean()
    # 2-D Hann window tapers the edges to zero before the FFT
    window = np.outer(np.hanning(img.shape[0]), np.hanning(img.shape[1]))
    # fftshift puts the DC/zero-frequency term in the centre for display
    f = np.fft.fftshift(np.fft.fft2(img * window))
    return np.log1p(np.abs(f) ** 2)


def _spectrum_display_range(
    *spectra: np.ndarray, center_exclude_radius: int = 6,
) -> tuple[float, float]:
    """Percentile-based (vmin, vmax) for one or more power spectra.

    Excludes a small disk around the origin (still DC-dominated even after
    windowing) so the percentiles reflect the informative mid/high-frequency
    content rather than being skewed by a handful of huge low-frequency
    pixels.
    """
    h, w = spectra[0].shape
    yy, xx = np.ogrid[:h, :w]
    # Boolean mask of every pixel outside the small disk around the centre
    outside = (yy - h // 2) ** 2 + (xx - w // 2) ** 2 > center_exclude_radius ** 2
    values = np.concatenate([spec[outside] for spec in spectra])
    return float(np.percentile(values, 2.0)), float(np.percentile(values, 99.5))


def save_motion_diagnostic_plot(
    shifts_xy: np.ndarray,
    crop_before: np.ndarray,
    crop_after: np.ndarray,
    output_path: str,
    binning: int = 4,
) -> None:
    """Save a PNG with the accumulated MotionCor2 shift trajectory and the
    FFT power spectrum of the crop region before vs. after correction.

    ``crop_before`` is the unaligned sum of the cropped raw frames;
    ``crop_after`` is MotionCor2's aligned sum of the same crop. A tighter,
    more extended spectrum after correction indicates less residual
    motion blur.
    """
    import matplotlib
    matplotlib.use("Agg")
    import matplotlib.pyplot as plt

    b = max(1, binning)
    spec_before = _log_power_spectrum(crop_before[::b, ::b])
    spec_after = _log_power_spectrum(crop_after[::b, ::b])
    vmin, vmax = _spectrum_display_range(spec_before, spec_after)

    fig, axes = plt.subplots(1, 3, figsize=(18, 5.5), constrained_layout=True)
    fig.suptitle(
        os.path.basename(output_path).replace("_motion_diagnostic.png", ""), fontsize=10
    )

    # Panel 1: accumulated motion trajectory
    dx, dy = shifts_xy[:, 0], shifts_xy[:, 1]
    axes[0].plot(dx, dy, "-o", color="steelblue", markersize=4)
    # Label each point with its frame index so the trajectory's direction is readable
    for i, (x, y) in enumerate(zip(dx, dy)):
        axes[0].annotate(str(i), (x, y), fontsize=7, xytext=(3, 3),
                          textcoords="offset points")
    # Straight-line distance between the first and last frame's shift
    total = float(np.hypot(dx[-1] - dx[0], dy[-1] - dy[0]))
    axes[0].set_title(f"Accumulated motion  (total {total:.1f} px)", fontsize=9)
    axes[0].set_xlabel("shift x (px)")
    axes[0].set_ylabel("shift y (px)")
    axes[0].axhline(0, color="gray", lw=0.5)
    axes[0].axvline(0, color="gray", lw=0.5)
    axes[0].set_aspect("equal", adjustable="datalim")
    axes[0].grid(alpha=0.3)

    # Panel 2/3: FFT of the crop before vs. after correction
    axes[1].imshow(spec_before, cmap="gray", vmin=vmin, vmax=vmax, interpolation="nearest")
    axes[1].set_title("FFT of crop  (before correction)", fontsize=9)
    axes[1].axis("off")

    axes[2].imshow(spec_after, cmap="gray", vmin=vmin, vmax=vmax, interpolation="nearest")
    axes[2].set_title("FFT of crop  (after correction)", fontsize=9)
    axes[2].axis("off")

    plt.savefig(output_path, dpi=100)
    plt.close(fig)
    log.info("Motion diagnostic plot: %s", output_path)


# ---------------------------------------------------------------------------
# Per-file orchestration
# ---------------------------------------------------------------------------

def process_one_tiff(
    input_path: str,
    output_dir: str,
    pixel_size: float,
    motioncor2_path: str,
    bfactor: float = 100.0,
    gpu: int = 0,
    mask_shrink: int = 20,
    mask_threshold: float | None = None,
    mask_threshold_fraction: float | None = None,
    template_mask: np.ndarray | None = None,
    keep_temp: bool = False,
    save_diagnostic: bool = False,
    diagnostic_binning: int = 4,
    diagnostic_subdir: str | None = None,
    mc2_extra: list[str] | None = None,
    split_frames: bool = False,
    stack_output: bool = False,
    overwrite: bool = True,
    run_ctffind: bool = True,
    ctf_flip: bool = False,
    ctffind4_path: str | None = None,
    voltage_kv: float = 300.0,
    cs_mm: float = 2.7,
    amp_contrast: float = 0.07,
    ctf_box_size: int = 512,
    ctf_min_res: float = 30.0,
    ctf_max_res: float | None = None,
    ctf_min_defocus: float = 5000.0,
    ctf_max_defocus: float = 200000.0,
    ctf_defocus_step: float = 1000.0,
    ctf_min_score: float = 0.1,
    rotate: int = 0,
    sidecar_step: tuple | None = None,
    frames_dir: str | None = None,
) -> str:
    """Process a single multi-frame TIFF; return path to the output MRC.

    When ``split_frames`` is True, additional outputs are written for the
    odd- and even-indexed frames — sum of frames at indices 0, 2, 4, … and
    1, 3, 5, … respectively. These half-datasets are ready to use directly
    with cryoCARE or similar.

    When ``stack_output`` is False (default), each tile is written as its
    own single-slice MRC: {output_dir}/{stem}.mrc (and
    {output_dir}/{odd,even}/{stem}.mrc for the split frames).

    When ``stack_output`` is True, ``input_path`` must be named
    ``{base}_{tile_index}_{tilt_angle}.tif``. The tile is instead written
    as one z-slice of a shared per-tilt MRC stack:
    {output_dir}/{base}_{tilt}.mrc (and {output_dir}/{odd,even}/{base}_{tilt}.mrc),
    sized to hold every tile sharing that base/tilt. This matches the raw
    per-tilt montage stack layout ``stitch.py`` expects.

    When ``run_ctffind`` is True (default), ctffind4 estimates the CTF on
    the same motion-corrected crop used for shift estimation. If
    ``ctf_flip`` is also True (default: False), that defocus/astigmatism is
    used to phase-flip the full-frame output before it's written, but only
    if the fit's score is at least ``ctf_min_score``. A low-confidence fit
    is worse than no fit: phase-flipping with the wrong defocus scrambles
    real signal rather than just failing to fix contrast reversals, so
    tiles below the threshold are left un-flipped (their CTF is still
    estimated/logged/saved). ``run_ctffind=False`` implies no phase-flip
    regardless of ``ctf_flip``.

    ``overwrite`` defaults to True, so existing outputs are always
    reprocessed. Pass ``overwrite=False`` to skip inputs whose output
    already exists instead, e.g. to resume a partially-completed run.

    ``rotate`` (0/90/180/270, counterclockwise) rotates the final output
    image(s) as the very last step, after motion correction and CTF
    phase-flipping. 90/270 swap the output's width and height.

    If ``sidecar_step`` is given — ``(effective, requested)`` from
    :func:`motion_correction_effective` — this tile writes a sidecar
    describing what was actually done to it, see :func:`tile_sidecar_record`.
    A tile skipped because its output already exists (``overwrite=False``)
    writes nothing: its sidecar is the one the run that produced it wrote.
    """
    in_stem = Path(input_path).stem

    if stack_output:
        base, raw_tile_idx, tilt = parse_tile_filename(input_path)
        # tile_index is a cumulative counter across the whole tilt series
        # (it does not reset per tilt), so re-base it to a 0-based z-slice
        # position within this tilt's own stack.
        min_idx, max_idx = tile_index_range_for_tilt(input_path, base, tilt)
        tile_idx = raw_tile_idx - min_idx
        n_tiles = max_idx - min_idx + 1
        out_mrc  = stack_path_for(output_dir, base, tilt)
        out_odd  = stack_path_for(os.path.join(output_dir, "odd"), base, tilt)
        out_even = stack_path_for(os.path.join(output_dir, "even"), base, tilt)
        # A shared stack file always "exists" once any tile has written to
        # it, so use a per-tile sentinel to track completion instead.
        marker_dir = os.path.join(output_dir, ".stack_markers")
        marker = os.path.join(marker_dir, f"{in_stem}.done")
        skip = (not overwrite) and os.path.exists(marker)
    else:
        # Same identity fields for the sidecar, but one file per tile: no
        # z-slice, and no need to look at the siblings to re-base the index.
        # The naming convention is optional here, so a failure to parse it is
        # recorded as "unknown" rather than raised.
        try:
            base, raw_tile_idx, tilt = parse_tile_filename(input_path)
        except ValueError:
            base, raw_tile_idx, tilt = "", -1, float("nan")
        min_idx, tile_idx, n_tiles = -1, -1, -1
        out_mrc  = os.path.join(output_dir, f"{in_stem}.mrc")
        out_odd  = os.path.join(output_dir, "odd",  f"{in_stem}.mrc")
        out_even = os.path.join(output_dir, "even", f"{in_stem}.mrc")
        skip = (not overwrite) and os.path.exists(out_mrc)
        if split_frames:
            skip = skip and os.path.exists(out_odd) and os.path.exists(out_even)

    if skip:
        log.info("Output already exists, skipping: %s", out_mrc)
        return out_mrc

    # 1. Load the mean of all sub-frames for mask detection -- not just the
    # first. A single frame's median is 0 on sparse single-electron-counting
    # data (measured: 0.2 counts/px mean, 0 median on one frame vs. 0.25 on
    # the 4-frame mean), which makes a median-relative threshold degenerate.
    # This matches choose_mask_params.load_mean_frame exactly, so a threshold
    # tuned there transfers here unchanged.
    t0 = time.perf_counter()
    with Image.open(input_path) as img:
        n_frames = getattr(img, "n_frames", 1)
        img.seek(0)
        mask_frame = np.asarray(img.copy()).astype(np.float32)
        for i in range(1, n_frames):
            img.seek(i)
            mask_frame += np.asarray(img.copy()).astype(np.float32)
    mask_frame /= n_frames
    log.info("%s: loaded mean of %d frames for mask detection in %.1f s",
             in_stem, n_frames, time.perf_counter() - t0)

    # 2. Beam mask from the mean frame. A fraction takes priority over an
    # absolute count when both are given -- the absolute count from a
    # previous dataset does not transfer across detector/dose regimes, which
    # is exactly what made this mask degenerate on this data (see
    # process_one_tiff's docstring).
    t1 = time.perf_counter()
    if mask_threshold_fraction is not None:
        effective_threshold = mask_threshold_fraction * float(np.median(mask_frame))
    else:
        effective_threshold = mask_threshold
    if template_mask is not None:
        mask = make_mask(mask_frame, template_mask=template_mask, shrinkn=mask_shrink,
                         absolutethreshold=effective_threshold)
    else:
        mask = make_mask(mask_frame, shrinkn=mask_shrink, absolutethreshold=effective_threshold)
    log.info("%s: beam mask in %.1f s", in_stem, time.perf_counter() - t1)

    # 3. Largest inscribed square, rounded to an FFT-friendly size
    t2 = time.perf_counter()
    H, W = mask_frame.shape
    _lis_bin = 8  # run DP on downsampled mask, then scale coords back up
    mask_small = mask[::_lis_bin, ::_lis_bin]
    row_top_s, col_left_s, side_s = largest_inscribed_square(mask_small)
    row_top, col_left, side = row_top_s * _lis_bin, col_left_s * _lis_bin, side_s * _lis_bin
    log.info("%s: largest inscribed square in %.1f s", in_stem, time.perf_counter() - t2)
    if side < 64:
        raise ValueError(
            f"Largest inscribed square is only {side}px — beam mask may have failed."
        )
    inscribed_side = side
    fft_side = _fft_friendly_size(side)
    if fft_side < side:
        # Centre the smaller FFT-friendly square within the inscribed square
        pad = (side - fft_side) // 2
        row_top += pad
        col_left += pad
        log.info(
            "%s: crop adjusted %d→%d px (FFT-friendly)  row=%d col=%d  (full frame %dx%d)",
            in_stem, side, fft_side, row_top, col_left, H, W,
        )
        side = fft_side
    else:
        log.info(
            "%s: square crop  row=%d col=%d side=%d  (full frame %dx%d)",
            in_stem, row_top, col_left, side, H, W,
        )

    # 3b. Diagnostic output directory (the mask/crop plot itself is saved
    # later, at step 8d, once the full-frame motion-corrected sum exists)
    diag_dir = os.path.join(output_dir, diagnostic_subdir) if diagnostic_subdir else output_dir

    # 4. Load all frames
    t2 = time.perf_counter()
    frames = load_multipage_tiff(input_path)
    if frames.ndim == 2:
        frames = frames[np.newaxis]
    log.info("%s: loaded all %d frames in %.1f s",
             in_stem, n_frames, time.perf_counter() - t2)

    # 5. Crop frames and write temp TIFF
    ctf_result: dict | None = None
    with tempfile.TemporaryDirectory(prefix="mc2_", suffix=f"_{in_stem}") as tmpdir:
        t3 = time.perf_counter()
        cropped = frames[:, row_top : row_top + side, col_left : col_left + side]
        tmp_tiff = os.path.join(tmpdir, f"{in_stem}_crop.tif")
        write_multipage_tiff(cropped, tmp_tiff)
        log.info("%s: crop + temp TIFF write in %.1f s", in_stem, time.perf_counter() - t3)

        # 6. Run MotionCor2 on cropped TIFF
        t4 = time.perf_counter()
        tmp_mrc = os.path.join(tmpdir, f"{in_stem}_mc2.mrc")
        stdout_text = run_motioncor2(
            in_tiff=tmp_tiff,
            out_mrc=tmp_mrc,
            log_dir=tmpdir,
            pixel_size=pixel_size,
            bfactor=bfactor,
            gpu=gpu,
            motioncor2_path=motioncor2_path,
            extra_args=mc2_extra,
        )
        log.info("%s: MotionCor2 in %.1f s", in_stem, time.perf_counter() - t4)

        if keep_temp:
            # Written to a subdirectory, not output_dir itself — a flat
            # *.mrc glob over output_dir (e.g. stitch.py's per-tilt stack
            # pattern) must never pick these up alongside the real outputs.
            logs_dir = os.path.join(output_dir, "mc2_logs")
            os.makedirs(logs_dir, exist_ok=True)

            dest = os.path.join(logs_dir, f"{in_stem}.mc2.log")
            with open(dest, "w") as fh:
                fh.write(stdout_text)
            log.info("Saved MC2 output: %s", dest)

            mrc_dest = os.path.join(logs_dir, f"{in_stem}.mc2_crop.mrc")
            shutil.copyfile(tmp_mrc, mrc_dest)
            log.info("Saved MC2 cropped output: %s", mrc_dest)

        # 6b. Run ctffind4 on the motion-corrected crop to estimate the CTF
        if run_ctffind:
            t4b = time.perf_counter()
            diag_mrc = os.path.join(tmpdir, f"{in_stem}_ctffind_diag.mrc")
            box = min(ctf_box_size, side)
            box -= box % 2
            max_res = ctf_max_res if ctf_max_res is not None else max(3.0 * pixel_size, 4.0)
            try:
                txt_path = run_ctffind4(
                    in_mrc=tmp_mrc,
                    diag_mrc=diag_mrc,
                    pixel_size=pixel_size,
                    voltage_kv=voltage_kv,
                    cs_mm=cs_mm,
                    amp_contrast=amp_contrast,
                    ctffind4_path=ctffind4_path,
                    box_size=box,
                    min_res=ctf_min_res,
                    max_res=max_res,
                    min_defocus=ctf_min_defocus,
                    max_defocus=ctf_max_defocus,
                    defocus_step=ctf_defocus_step,
                )
                ctf_result = parse_ctffind4_result(txt_path)
                log.info(
                    "%s: CTF  df1=%.0f  df2=%.0f  az=%.1f  phase=%.2f  score=%.3f  "
                    "fit=%.1fA  (%.1f s)",
                    in_stem, ctf_result["defocus1"], ctf_result["defocus2"],
                    ctf_result["azimuth"], ctf_result["phase_shift"],
                    ctf_result["score"], ctf_result["fit_resolution"],
                    time.perf_counter() - t4b,
                )
                if keep_temp:
                    dest = os.path.join(output_dir, f"{in_stem}.ctffind.txt")
                    shutil.copyfile(txt_path, dest)
                    log.info("Saved ctffind4 results: %s", dest)

                # Aggregate per-tile CTF fit parameters into one shared file
                results_path = os.path.join(output_dir, "ctf_results.txt")
                append_ctf_result(results_path, in_stem, ctf_result)

                if save_diagnostic:
                    os.makedirs(diag_dir, exist_ok=True)
                    ctf_diag_path = os.path.join(diag_dir, f"{in_stem}_ctf_diagnostic.png")
                    save_ctf_diagnostic_plot(diag_mrc, ctf_result, ctf_diag_path)
            except Exception as exc:
                log.error("%s: ctffind4 failed: %s", in_stem, exc)
                ctf_result = None

        # 7. Parse shifts from captured stdout
        shifts_xy = parse_motioncor2_shifts(stdout_text, n_frames)
        log.info("Shifts (x,y px):\n%s", shifts_xy)

        # 7b. Optional motion diagnostic: accumulated shift + before/after FFT
        # of the crop, while tmp_mrc (MotionCor2's aligned crop) still exists
        if save_diagnostic:
            os.makedirs(diag_dir, exist_ok=True)
            motion_diag_path = os.path.join(diag_dir, f"{in_stem}_motion_diagnostic.png")
            t_mdiag = time.perf_counter()
            with mrcfile.open(tmp_mrc, permissive=True) as mrc:
                crop_after = np.asarray(mrc.data).astype(np.float32, copy=False)
            crop_before = cropped.astype(np.float32).sum(axis=0)
            save_motion_diagnostic_plot(
                shifts_xy, crop_before, crop_after, motion_diag_path,
                binning=diagnostic_binning,
            )
            log.info("%s: motion diagnostic plot in %.1f s",
                      in_stem, time.perf_counter() - t_mdiag)

    # 8. Apply shifts to full frames and sum
    t5 = time.perf_counter()
    os.makedirs(output_dir, exist_ok=True)
    shifts_xy[:, 1] *= -1.0  # MotionCor2's y-shifts are inverted relative to numpy indexing
    if split_frames:
        aligned_sum, odd_sum, even_sum = apply_shifts_split(frames, shifts_xy)
    else:
        aligned_sum = apply_shifts_and_sum(frames, shifts_xy)
    log.info("%s: shift application in %.1f s", in_stem, time.perf_counter() - t5)

    # 8d. Optional mask/crop diagnostic — mask on the raw frame, crop region
    # on the full-frame motion-corrected sum (before CTF-flip/rotation, so
    # the crop rectangle still lines up with row_top/col_left/side)
    if save_diagnostic:
        os.makedirs(diag_dir, exist_ok=True)
        diag_path = os.path.join(diag_dir, f"{in_stem}_diagnostic.png")
        t_diag = time.perf_counter()
        save_diagnostic_plot(
            mask_frame, aligned_sum, mask, row_top, col_left, side,
            diag_path, binning=diagnostic_binning,
        )
        log.info("%s: diagnostic plot in %.1f s", in_stem, time.perf_counter() - t_diag)

    # 8b. CTF phase-flip using the crop-estimated defocus/astigmatism —
    # skipped for low-confidence fits, where flipping would scramble real
    # signal rather than correct it.
    phase_flipped = False
    if ctf_flip and ctf_result is not None and ctf_result["score"] >= ctf_min_score:
        t5b = time.perf_counter()
        flip_args = (
            pixel_size, ctf_result["defocus1"], ctf_result["defocus2"],
            ctf_result["azimuth"], voltage_kv, cs_mm, amp_contrast,
            ctf_result["phase_shift"],
        )
        aligned_sum = ctf_phase_flip(aligned_sum, *flip_args)
        if split_frames:
            odd_sum = ctf_phase_flip(odd_sum, *flip_args)
            even_sum = ctf_phase_flip(even_sum, *flip_args)
        phase_flipped = True
        log.info("%s: CTF phase-flip in %.1f s", in_stem, time.perf_counter() - t5b)
    elif ctf_flip and ctf_result is not None:
        log.warning(
            "%s: CTF score %.3f below threshold %.3f; output NOT phase-flipped.",
            in_stem, ctf_result["score"], ctf_min_score,
        )
    elif ctf_flip and run_ctffind:
        log.warning("%s: CTF estimation failed; output NOT phase-flipped.", in_stem)

    # 8c. Rotate the final output by a multiple of 90 degrees, if requested
    k = (rotate // 90) % 4
    if k:
        aligned_sum = np.ascontiguousarray(np.rot90(aligned_sum, k))
        if split_frames:
            odd_sum = np.ascontiguousarray(np.rot90(odd_sum, k))
            even_sum = np.ascontiguousarray(np.rot90(even_sum, k))

    # 9. Write output MRC(s) (float32, single 2-D slice each)
    t6 = time.perf_counter()
    if stack_output:
        write_tile_slice(out_mrc, tile_idx, n_tiles, aligned_sum, pixel_size)
        if split_frames:
            for label, data, path in (
                ("odd", odd_sum, out_odd), ("even", even_sum, out_even)
            ):
                write_tile_slice(path, tile_idx, n_tiles, data, pixel_size)
                log.info("Wrote %s frames: %s [tile %d]", label, path, tile_idx)
        os.makedirs(marker_dir, exist_ok=True)
        Path(marker).touch()
    else:
        with mrcfile.new(out_mrc, overwrite=True) as mrc:
            mrc.set_data(aligned_sum)
            mrc.voxel_size = pixel_size

        if split_frames:
            for label, data in (("odd", odd_sum), ("even", even_sum)):
                sub_dir = os.path.join(output_dir, label)
                os.makedirs(sub_dir, exist_ok=True)
                out_path = os.path.join(sub_dir, f"{in_stem}.mrc")
                with mrcfile.new(out_path, overwrite=True) as mrc:
                    mrc.set_data(data)
                    mrc.voxel_size = pixel_size
                log.info("Wrote %s frames: %s", label, out_path)
    log.info("%s: MRC write in %.1f s", in_stem, time.perf_counter() - t6)

    if sidecar_step is not None:
        step_effective, step_requested = sidecar_step
        write_tile_sidecar(
            output_path=out_mrc,
            input_path=input_path,
            frames_dir=frames_dir,
            stack_output=stack_output,
            step_effective=step_effective,
            step_requested=step_requested,
            tile_record=tile_sidecar_record(
                tile=in_stem,
                base=base,
                tilt_deg=tilt,
                tile_index_raw=raw_tile_idx,
                tile_index_base=min_idx,
                z_slice=tile_idx,
                n_tiles_in_tilt=n_tiles,
                output_path=out_mrc,
                frame_shape=(H, W),
                output_shape=aligned_sum.shape,
                n_frames=n_frames,
                mask_fraction=float(mask.mean()),
                mask_shrink=mask_shrink,
                mask_threshold=effective_threshold,
                crop=(row_top, col_left, side, inscribed_side),
                shifts_xy=shifts_xy,
                ctf_result=ctf_result,
                phase_flipped=phase_flipped,
                rotate=rotate,
            ),
        )

    return out_mrc


def tile_sidecar_record(
    *,
    tile: str,
    base: str,
    tilt_deg: float,
    tile_index_raw: int,
    tile_index_base: int,
    z_slice: int,
    n_tiles_in_tilt: int,
    output_path: str,
    frame_shape: tuple[int, int],
    output_shape: tuple[int, int],
    n_frames: int,
    mask_fraction: float,
    mask_shrink: int,
    mask_threshold: float | None,
    crop: tuple[int, int, int, int],
    shifts_xy: np.ndarray,
    ctf_result: dict | None,
    phase_flipped: bool,
    rotate: int,
) -> dict:
    """One tile's own results: what was measured and what was applied.

    Every field is an outcome, not a request. ``crop`` is where the crop
    actually landed after the FFT-friendly shrink, ``shifts_xy`` are the
    shifts in the sense they were applied to the full frames (MotionCor2's
    y-convention already inverted), and ``phase_flipped`` says whether this
    particular tile was flipped — which depends on its own CTF score, so it
    cannot be read off the step's parameters.

    A tile with no usable CTF fit still gets the ``ctf`` keys, filled with
    NaN, so the folded arrays stay dense and rows keep lining up.
    """
    row_top, col_left, side, inscribed_side = crop
    ctf = ctf_result or {}
    nan = float("nan")
    return {
        "tile": tile,
        "base": base,
        "tilt_deg": float(tilt_deg),
        # Cumulative across the series, and the value subtracted from it to
        # get z_slice — the pair fix_stack_tile_index.py has to guess at.
        "tile_index_raw": int(tile_index_raw),
        "tile_index_base": int(tile_index_base),
        "z_slice": int(z_slice),
        "n_tiles_in_tilt": int(n_tiles_in_tilt),
        "output_path": str(output_path),
        "frame_shape": [int(frame_shape[0]), int(frame_shape[1])],
        "output_shape": [int(output_shape[0]), int(output_shape[1])],
        "n_frames": int(n_frames),
        "mask": {
            "fraction": mask_fraction,
            "shrink_px": int(mask_shrink),
            "threshold": nan if mask_threshold is None else float(mask_threshold),
        },
        "crop": {
            "row_top": int(row_top), "col_left": int(col_left), "side": int(side),
            "inscribed_side": int(inscribed_side),
        },
        "shifts_xy_px": np.asarray(shifts_xy, dtype=float).tolist(),
        "total_motion_px": float(np.hypot(
            shifts_xy[-1, 0] - shifts_xy[0, 0], shifts_xy[-1, 1] - shifts_xy[0, 1],
        )),
        "ctf": {
            "defocus1": float(ctf.get("defocus1", nan)),
            "defocus2": float(ctf.get("defocus2", nan)),
            "azimuth": float(ctf.get("azimuth", nan)),
            "phase_shift": float(ctf.get("phase_shift", nan)),
            "score": float(ctf.get("score", nan)),
            "fit_resolution": float(ctf.get("fit_resolution", nan)),
            "fitted": ctf_result is not None,
        },
        "phase_flipped": bool(phase_flipped),
        "rotate_applied_deg": int((rotate // 90) % 4) * 90,
    }


# ---------------------------------------------------------------------------# ---------------------------------------------------------------------------
# Provenance sidecars: this step's effective parameters
# ---------------------------------------------------------------------------


def motion_correction_effective(
    args: argparse.Namespace,
    *,
    mc2_path: str,
    ctffind4_path: str | None,
    ctf_max_res: float,
) -> tuple[dict, dict, list]:
    """This step's effective parameters, its requested ones, and what was auto.

    Computed once per process and written into every tile's sidecar. Values
    that were auto-detected or defaulted are listed in ``auto_resolved``, so
    reading a sidecar later tells you not just what was used but that nobody
    chose it.

    Hundreds of array tasks run this step with identical parameters. Under the
    manifest that was a write race, settled with ``if_absent=True`` and a lock;
    with sidecars each task writes only its own tiles' files, so the same
    values simply appear in each — no coordination, and no chance of a task
    silently disagreeing with the record another task got there first to write.
    """
    auto_resolved = []
    if args.motioncor2 is None:
        auto_resolved.append("motioncor2")
    if args.run_ctffind and args.ctffind4 is None:
        auto_resolved.append("ctffind4")
    if args.ctf_max_res is None:
        auto_resolved.append("ctf_max_res_A")
    if args.mask_threshold is None and args.mask_threshold_fraction is None:
        auto_resolved.append("mask_threshold")

    effective = {
        "pixel_size_A": args.pixel_size,
        "bfactor": args.bfactor,
        "motioncor2": mc2_path,
        "mc2_extra_args": args.mc2_args or [],
        "mask_shrink": args.mask_shrink,
        # None (both) means make_mask's own default: 0.4 x the image median.
        # A fraction varies per-tile by construction, so there is no single
        # absolute number to record here either -- see each tile's own record.
        "mask_threshold": args.mask_threshold,
        "mask_threshold_fraction": args.mask_threshold_fraction,
        "mask_threshold_rule": (
            "fraction of image median" if args.mask_threshold_fraction is not None
            else "absolute" if args.mask_threshold is not None
            else "0.4 * image median (make_mask default)"
        ),
        "template_mask": args.template_mask,
        "run_ctffind": args.run_ctffind,
        "ctffind4": ctffind4_path,
        "voltage_kV": args.voltage,
        "cs_mm": args.cs,
        "amplitude_contrast": args.amp_contrast,
        "ctf_box_size": args.ctf_box_size,
        "ctf_min_res_A": args.ctf_min_res,
        "ctf_max_res_A": ctf_max_res,
        "ctf_defocus_search_A": [args.ctf_min_defocus, args.ctf_max_defocus,
                                 args.ctf_defocus_step],
        # Requested, not achieved: whether a given tile was actually flipped
        # depends on its own CTF score, which is in each tile's own record.
        "ctf_flip_requested": args.ctf_flip,
        "ctf_min_score": args.ctf_min_score,
        "rotate_deg": int((args.rotate // 90) % 4) * 90,
        "split_frames": args.split_frames,
        "output_layout": "per_tilt_stack" if args.stack_output else "per_tile_mrc",
        "output_dir": os.path.abspath(args.output_dir),
        # The frame this step defines: full detector frames, corner origin,
        # keyed by tilt and tile, at the pixel size every later binning is
        # relative to. Under the manifest this was a separate `frames` section
        # that only the first task wrote; it is three scalars, so it rides in
        # `effective` now rather than needing a document of its own.
        "frame": "raw_tile",
        "frame_pixel_size_A": args.pixel_size,
        "frame_note": ("output of motion correction; full detector area, "
                       "not the crop"),
        "auto_resolved": auto_resolved,
    }
    requested = {
        "motioncor2": args.motioncor2,
        "ctffind4": args.ctffind4,
        "mask_threshold": args.mask_threshold,
        "mask_threshold_fraction": args.mask_threshold_fraction,
        "ctf_max_res_A": args.ctf_max_res,
        "rotate_deg": args.rotate,
    }
    return effective, requested, auto_resolved


def write_tile_sidecar(
    *,
    output_path: str,
    input_path: str,
    frames_dir: str | None,
    stack_output: bool,
    step_effective: dict,
    step_requested: dict,
    tile_record: dict,
) -> None:
    """Write one tile's sidecar.

    Under ``--stack-output`` the output is a z-slice of a per-tilt stack that
    other tasks are still writing into, so the sidecar is keyed by the tile's
    input stem — one uncontended path per task — and the stack itself is not
    fingerprinted, because its bytes are not stable until every task is done.
    With one MRC per tile the output is this task's alone, so the sidecar is
    the plain ``<tile>.mrc.json`` and the fingerprint is meaningful.

    The raw frames are recorded as a directory summary rather than hundreds of
    fingerprints: that is the call ``sidecar_metadata_plan.md`` made under
    "Open questions", and it keeps a per-tile sidecar smaller than the thing
    it describes.
    """
    # Fingerprint the raw TIFF itself when it's a real file, plus a cheap
    # directory-level summary of the whole frames directory it came from
    inputs = [fingerprint(input_path, role="frames_stack")] if (
        os.path.isfile(input_path)) else []
    if frames_dir:
        inputs.append(directory_input(frames_dir, role="frames"))

    sidecar.write(
        output_path,
        SIDECAR_STEP,
        {**step_effective, **tile_record},
        inputs=inputs,
        key=tile_record["tile"] if stack_output else None,
        requested=step_requested,
        fingerprint_output=not stack_output,
    )


# ---------------------------------------------------------------------------
# CLI
# ---------------------------------------------------------------------------

DEFAULT_SLURM_TEMPLATE = """\
#!/bin/bash
#SBATCH --job-name=mc2_frames
#SBATCH --output={log_dir}/mc2_%A_%a.out
#SBATCH --error={log_dir}/mc2_%A_%a.err
#SBATCH -p gpu-l40s
#SBATCH --mem=32G
#SBATCH --time=00:30:00
#SBATCH --gres=gpu:1
#SBATCH --array=0-{n_minus_1}%{max_concurrent}  # %N caps simultaneous tasks

frames_dir="{frames_dir}"
output_dir="{output_dir}"
pixel_size={pixel_size}
bfactor={bfactor}

source /programs/sbgrid.shrc
module load CUDA/12.2.0

MOTIONCOR2=/programs/x86_64-linux/motioncor2/1.6.4/MotionCor2_1.6.4_Cuda121_Mar312023

mapfile -t tiffs < <(ls "${{frames_dir}}"/*.tif | sort)
input="${{tiffs[$SLURM_ARRAY_TASK_ID]}}"

beam_mask_motioncorr \\
    --input "${{input}}" \\
    --output-dir "${{output_dir}}" \\
    --pixel-size ${{pixel_size}} \\
    --bfactor ${{bfactor}} \\
    --motioncor2 ${{MOTIONCOR2}} \\
    --gpu 0

# Each task writes its own tiles' provenance sidecars as it goes (they are on
# by default), so there is nothing to finalize afterwards and no dependent
# job to queue. Read one back with:
#   montage_provenance show {output_dir}/<base>_<tilt>.mrc
"""


def main() -> None:
    parser = argparse.ArgumentParser(
        description=__doc__,
        formatter_class=argparse.RawDescriptionHelpFormatter,
    )
    parser.add_argument(
        "--input", "-i", nargs="+", required=True,
        help="Input TIFF file(s) or glob pattern(s). Each TIFF is a multi-frame movie.",
    )
    parser.add_argument(
        "--output-dir", "-o", required=True,
        help="Directory to write motion-corrected MRC files.",
    )
    parser.add_argument(
        "--pixel-size", type=float, required=True,
        help="Pixel size in Ångström (used by MotionCor2 for dose weighting etc.).",
    )
    parser.add_argument(
        "--motioncor2", default=None,
        help="Path to MotionCor2 executable. Auto-detected if not provided.",
    )
    parser.add_argument(
        "--bfactor", type=float, default=100.0,
        help="B-factor for MotionCor2 alignment (default: 100).",
    )
    parser.add_argument(
        "--gpu", type=int, default=0,
        help="GPU index for MotionCor2 (default: 0).",
    )
    parser.add_argument(
        "--mask-shrink", type=int, default=20,
        help="Erosion radius (px) to shrink beam mask away from edge (default: 20).",
    )
    parser.add_argument(
        "--mask-threshold", type=float, default=None,
        help="Absolute intensity threshold for beam detection. "
             "If omitted, uses 0.4 × image median (same default as make_mask). "
             "An absolute count does not transfer across datasets shot at "
             "different dose/detector modes -- prefer --mask-threshold-fraction "
             "unless a fixed count is specifically what was tuned. "
             "Use choose_mask_params to find the right value interactively.",
    )
    parser.add_argument(
        "--mask-threshold-fraction", type=float, default=None,
        help="Beam-detection threshold as a fraction of each tile's own "
             "image median, matching choose_mask_params' live '× median' "
             "readout exactly -- so a value confirmed there transfers here "
             "unchanged, per-tile, regardless of dose/detector mode. Takes "
             "priority over --mask-threshold when both are given.",
    )
    parser.add_argument(
        "--template-mask", default=None,
        help="Path to a .npy binary mask to use as beam template (numpy bool array).",
    )
    parser.add_argument(
        "--keep-logs", action="store_true",
        help="Copy the MotionCor2 log, MotionCor2's cropped aligned MRC "
             "(.mc2_crop.mrc), and ctffind4 results (.ctffind.txt) to the "
             "output directory.",
    )
    parser.add_argument(
        "--voltage", type=float, default=300.0,
        help="Acceleration voltage in kV, used for CTF estimation/phase-flipping "
             "(default: 300).",
    )
    parser.add_argument(
        "--cs", type=float, default=2.7,
        help="Spherical aberration in mm, used for CTF estimation/phase-flipping "
             "(default: 2.7).",
    )
    parser.add_argument(
        "--amp-contrast", type=float, default=0.07,
        help="Amplitude contrast fraction, used for CTF estimation/phase-flipping "
             "(default: 0.07).",
    )
    parser.add_argument(
        "--ctffind4", default=None,
        help="Path to the ctffind4 executable. Auto-detected if not provided.",
    )
    parser.add_argument(
        "--no-ctffind", dest="run_ctffind", action="store_false",
        help="Skip CTF estimation on the motion-corrected crop entirely "
             "(also disables CTF phase-flipping).",
    )
    parser.add_argument(
        "--ctf-flip", dest="ctf_flip", action="store_true", default=False,
        help="Phase-flip the output MRC(s) using the crop-estimated CTF "
             "(default: off; CTF is still estimated/logged/saved when "
             "ctffind4 runs.",
    )
    parser.add_argument(
        "--ctf-box-size", type=int, default=512,
        help="ctffind4 amplitude-spectrum box size in pixels (default: 512; "
             "clamped to the crop size).",
    )
    parser.add_argument(
        "--ctf-min-res", type=float, default=30.0,
        help="ctffind4 minimum resolution to fit, in Angstrom (default: 30).",
    )
    parser.add_argument(
        "--ctf-max-res", type=float, default=None,
        help="ctffind4 maximum resolution to fit, in Angstrom. "
             "Defaults to max(3 x pixel size, 4 A) if omitted.",
    )
    parser.add_argument(
        "--ctf-min-defocus", type=float, default=5000.0,
        help="ctffind4 minimum defocus search bound, in Angstrom (default: 5000).",
    )
    parser.add_argument(
        "--ctf-max-defocus", type=float, default=200000.0,
        help="ctffind4 maximum defocus search bound, in Angstrom (default: "
             "200000 — square-beam montage tiles are often imaged at very "
             "high defocus).",
    )
    parser.add_argument(
        "--ctf-defocus-step", type=float, default=1000.0,
        help="ctffind4 defocus search step, in Angstrom (default: 1000).",
    )
    parser.add_argument(
        "--ctf-min-score", type=float, default=0.1,
        help="Minimum ctffind4 cross-correlation score required to phase-flip "
             "a tile (default: 0.1). Tiles fit below this are left un-flipped "
             "but still estimated/logged/saved to ctf_results.txt — "
             "phase-flipping with a low-confidence fit scrambles real signal "
             "rather than correcting it.",
    )
    parser.add_argument(
        "--rotate", type=int, default=0, choices=[0, 90, 180, 270],
        help="Rotate the final output image(s) counterclockwise by this many "
             "degrees (default: 0). Applied as the last step, after motion "
             "correction and CTF phase-flipping; 90/270 swap width and height.",
    )
    parser.add_argument(
        "--save-diagnostic", nargs="?", const="", default=None, metavar="SUBDIR",
        help="Save PNG diagnostic plots for each input file: "
             "{stem}_diagnostic.png (beam mask on the raw frame; crop "
             "rectangle on the motion-corrected frame), "
             "{stem}_motion_diagnostic.png (accumulated MotionCor2 shift "
             "trajectory and before/after FFT of the crop), and, unless "
             "--no-ctffind, {stem}_ctf_diagnostic.png (ctffind4 fit). "
             "Optionally give a sub-directory name (relative to output_dir) "
             "to place the PNGs in, e.g. --save-diagnostic diagnostics.",
    )
    parser.add_argument(
        "--diagnostic-binning", type=int, default=4,
        help="Downsample factor for diagnostic plots (default: 4).",
    )
    parser.add_argument(
        "--split-frames", action="store_true",
        help="Also write odd- and even-frame sums to {output_dir}/odd/ and "
             "{output_dir}/even/. Useful for cryoCARE and similar denoising "
             "tools that require two independent half-datasets.",
    )
    parser.add_argument(
        "--stack-output", action="store_true",
        help="Write tiles into a shared per-tilt memory-mapped MRC stack "
             "(one z-slice per tile) instead of one MRC file per tile. "
             "Matches the raw-montage-stack layout stitch.py reads, so its "
             "output can be fed straight into stitch.py without going "
             "through the per-tile-directory input mode. Requires input "
             "filenames of the form {base}_{tile_index}_{tilt_angle}.tif "
             "and is safe to call concurrently (e.g. one SLURM array task "
             "per tile) — the stack file is created once under a file "
             "lock and each task then writes only its own slice.",
    )
    parser.add_argument(
        "--no-overwrite", dest="overwrite", action="store_false",
        help="Skip inputs whose output already exists instead of "
             "reprocessing them (resumes a partially-completed run). "
             "By default, existing outputs are always reprocessed.",
    )
    parser.add_argument(
        "--no-sidecar", dest="sidecar", action="store_false", default=True,
        help="Do not write a provenance sidecar per output tile. Sidecars are "
             "ON by default and cost one small JSON write per tile: they "
             "record this step's effective parameters plus that tile's mask, "
             "crop, shifts, CTF fit and — under --stack-output — the tilt, "
             "cumulative tile index and z-slice it landed on, which is the "
             "mapping fix_stack_tile_index.py exists to reconstruct. Each "
             "task writes only its own tiles' files, so there is nothing to "
             "coordinate between array tasks and no finalize job.",
    )
    parser.add_argument(
        "--max-failures", type=int, default=None, metavar="N",
        help="Exit non-zero if more than N inputs fail (default: only when "
             "every input failed). Use --max-failures 0 to treat any tile "
             "failure as a job failure.",
    )
    parser.add_argument(
        "--mc2-args", nargs=argparse.REMAINDER, default=[],
        help="Extra arguments forwarded verbatim to MotionCor2 (after --).",
    )
    parser.add_argument(
        "--print-slurm", action="store_true",
        help="Print a SLURM array job script for the matched input files and exit.",
    )
    parser.add_argument(
        "--max-concurrent", type=int, default=200,
        help="Max simultaneously active array tasks (default: 200). "
             "Sets the %%N throttle in the SLURM array directive.",
    )
    parser.add_argument(
        "--verbose", "-v", action="store_true",
    )
    args = parser.parse_args()

    logging.basicConfig(
        level=logging.DEBUG if args.verbose else logging.INFO,
        format="%(asctime)s %(levelname)s %(message)s",
    )

    # Resolve glob patterns, falling back to a literal path if it isn't
    # actually a glob (e.g. a single filename with no wildcard characters)
    paths: list[str] = []
    for pat in args.input:
        expanded = sorted(glob.glob(pat))
        if expanded:
            paths.extend(expanded)
        elif os.path.exists(pat):
            paths.append(pat)
        else:
            log.warning("Pattern matched no files: %s", pat)
    paths = sorted(set(paths))

    if not paths:
        sys.exit("No input TIFF files found.")

    if args.print_slurm:
        frames_dir = os.path.dirname(os.path.abspath(paths[0]))
        log_dir = os.path.join(os.path.abspath(args.output_dir), "logs")
        script = DEFAULT_SLURM_TEMPLATE.format(
            frames_dir=frames_dir,
            output_dir=os.path.abspath(args.output_dir),
            log_dir=log_dir,
            pixel_size=args.pixel_size,
            bfactor=args.bfactor,
            n_minus_1=len(paths) - 1,
            max_concurrent=args.max_concurrent,
        )
        print(script)
        return

    mc2_path = args.motioncor2 or _find_motioncor2()
    ctffind4_path = (args.ctffind4 or _find_ctffind4()) if args.run_ctffind else None

    template_mask = None
    if args.template_mask:
        template_mask = np.load(args.template_mask)

    mc2_extra = args.mc2_args or []
    # Strip a leading '--' separator if present
    if mc2_extra and mc2_extra[0] == "--":
        mc2_extra = mc2_extra[1:]

    os.makedirs(args.output_dir, exist_ok=True)

    # The step's parameters, computed once and written into every tile's own
    # sidecar. Under the manifest this had to be grouped per tilt series,
    # because one shared file per series was the thing being written; a
    # sidecar is named after its output file, so a chunk straddling two series
    # needs no special handling at all.
    sidecar_step = None
    if args.sidecar:
        effective, requested, auto_resolved = motion_correction_effective(
            args,
            mc2_path=mc2_path,
            ctffind4_path=ctffind4_path,
            ctf_max_res=(
                args.ctf_max_res if args.ctf_max_res is not None
                else max(3.0 * args.pixel_size, 4.0)
            ),
        )
        sidecar_step = (effective, requested)
        if auto_resolved:
            log.info("Sidecar: auto-resolved %s", ", ".join(auto_resolved))

    n_ok = 0
    failures: list[tuple[str, str]] = []
    for p in tqdm(paths, desc="Motion correcting"):
        try:
            out = process_one_tiff(
                input_path=p,
                output_dir=args.output_dir,
                pixel_size=args.pixel_size,
                motioncor2_path=mc2_path,
                bfactor=args.bfactor,
                gpu=args.gpu,
                mask_shrink=args.mask_shrink,
                mask_threshold=args.mask_threshold,
                mask_threshold_fraction=args.mask_threshold_fraction,
                template_mask=template_mask,
                keep_temp=args.keep_logs,
                save_diagnostic=args.save_diagnostic is not None,
                diagnostic_binning=args.diagnostic_binning,
                diagnostic_subdir=args.save_diagnostic or None,
                mc2_extra=mc2_extra,
                split_frames=args.split_frames,
                stack_output=args.stack_output,
                overwrite=args.overwrite,
                run_ctffind=args.run_ctffind,
                ctf_flip=args.ctf_flip,
                ctffind4_path=ctffind4_path,
                voltage_kv=args.voltage,
                cs_mm=args.cs,
                amp_contrast=args.amp_contrast,
                ctf_box_size=args.ctf_box_size,
                ctf_min_res=args.ctf_min_res,
                ctf_max_res=args.ctf_max_res,
                ctf_min_defocus=args.ctf_min_defocus,
                ctf_max_defocus=args.ctf_max_defocus,
                ctf_defocus_step=args.ctf_defocus_step,
                ctf_min_score=args.ctf_min_score,
                rotate=args.rotate,
                sidecar_step=sidecar_step,
                frames_dir=os.path.dirname(os.path.abspath(p)) or ".",
            )
            log.info("Wrote: %s", out)
            n_ok += 1
        except Exception as exc:
            log.error("FAILED %s: %s", p, exc)
            failures.append((p, str(exc)))


    print(f"Done: {n_ok}/{len(paths)} files processed successfully.")
    if failures:
        # Group failures by their error message so one recurring problem
        # shows up as a single line with a count, not hundreds of duplicates
        by_reason: dict[str, int] = {}
        for _, reason in failures:
            by_reason[reason] = by_reason.get(reason, 0) + 1
        print(f"{len(failures)} file(s) failed:")
        for reason, count in sorted(by_reason.items(), key=lambda kv: -kv[1]):
            print(f"  {count:5d}  {reason}")

    # Individual tiles do fail on real data — a beam mask that finds nothing,
    # a MotionCor2 crash — and failing the array task for one of them would
    # break the afterok chain for no reason. A task that wrote *nothing* is a
    # different animal: that is a misconfiguration, and it used to exit 0 and
    # be reported COMPLETED. Fail loudly instead.
    if paths and n_ok == 0:
        sys.exit(
            f"No file was processed successfully out of {len(paths)}; "
            "see the failures above."
        )
    if args.max_failures is not None and len(failures) > args.max_failures:
        sys.exit(
            f"{len(failures)} failures exceeds --max-failures "
            f"{args.max_failures}."
        )


if __name__ == "__main__":
    main()
```
