## Supplementary material for "Whole cryo-FIB lamella acquisition at high magnification with rectangular beam montage cryo-electron tomography": Source code: Choose_fringe_size.py

```
"""
Interactive tool for choosing the fringe-removal size used during square-beam
montage stitching.  Load one frame from an MRC or multi-page TIFF stack, then
drag the sliders to find the fringe size (in unbinned pixels) and beam
threshold at which beam-edge artefacts are cleanly excluded without eroding
too much real sample signal. Pass the chosen values as --fringe and
--maskabsolutethreshold to the main stitching script.
"""

import os

import matplotlib
matplotlib.use('TkAgg')  # X11-forwarding compatible backend
import mrcfile
import numpy as np
from matplotlib import pyplot as plt
from matplotlib.widgets import Slider
from PIL import Image
import argparse
from .Utilities import fourier_interpolate, make_mask
from .mask_tuning_widgets import make_fraction_text, make_nav_buttons, make_threshold_slider


def open_stack(path):
    """
    Open an MRC or multi-page TIFF as a lazily-indexable stack.

    Returns ``(get_slice, n_slices, close)``: ``get_slice(idx)`` reads one 2D
    frame without loading the rest, ``n_slices`` is the stack length (1 for a
    single-image file), and ``close()`` releases the file handle. Only the
    requested frame is read either way -- via mrcfile's memmap for .mrc, via
    PIL's per-frame seek for .tif/.tiff -- so browsing a large raw movie or
    stitched montage stays cheap.
    """
    ext = os.path.splitext(path)[1].lower()
    if ext in (".tif", ".tiff"):
        img = Image.open(path)
        n_slices = getattr(img, "n_frames", 1)

        def get_slice(idx):
            img.seek(idx)
            return np.asarray(img.copy())

        return get_slice, n_slices, img.close

    mrc = mrcfile.mrcmemmap.MrcMemmap(path)
    n_slices = mrc.data.shape[0] if mrc.data.ndim == 3 else 1

    def get_slice(idx):
        return np.asarray(mrc.data[idx] if mrc.data.ndim == 3 else mrc.data)

    return get_slice, n_slices, mrc.close


def parse_commandline():
    parser = argparse.ArgumentParser(
        description=(
            "Interactively choose the fringe-removal size for square-beam montage "
            "stitching.  Opens a side-by-side view of the raw image and the masked "
            "image, with a slider controlling how many pixels are eroded inward from "
            "the detected beam edge.  Drag the slider until beam-edge fringes are "
            "removed without cutting into real sample signal, then note the value and "
            "pass it as --fringe to the main stitching script."
        ),
        formatter_class=argparse.ArgumentDefaultsHelpFormatter,
    )
    parser.add_argument(
        "-i", "--input",
        help=(
            "MRC or multi-page TIFF stack to preview.  The first frame/tilt is "
            "used by default.  Append a colon and a zero-based index to select "
            "a specific slice, e.g. file.mrc:5 or file.tif:5 loads slice 5."
        ),
        required=True,
        type=str,
    )
    parser.add_argument(
        "-b", "--binning",
        help=(
            "Integer downsampling factor applied via Fourier interpolation before "
            "display.  Use a higher value to speed up the preview on large datasets. "
            "Slider values are always reported in unbinned pixels so the chosen fringe "
            "size can be passed directly to the stitching script."
        ),
        required=False,
        type=int,
        default=4,
    )
    parser.add_argument(
        "-mt", "--maskthreshold",
        help=(
            "Beam-detection threshold expressed as a fraction of the image median. "
            "Pixels below this fraction are considered outside the beam.  Increase if "
            "the mask leaks into dark sample regions; decrease if the beam is not fully "
            "detected."
        ),
        type=float,
        default=0.4,
        required=False,
    )
    parser.add_argument(
        "-ma", "--maskabsolutethreshold",
        help=(
            "Absolute beam-detection threshold in image counts.  When set, overrides "
            "--maskthreshold.  Useful when the image median varies across a tilt series "
            "and a consistent count level is known."
        ),
        type=float,
        default=None,
        required=False,
    )
    return vars(parser.parse_args())


def main():
    args = parse_commandline()

    # Parse optional slice index from "file.mrc:N" syntax.
    input_str = args["input"]
    if ":" in input_str:
        input_path, slice_str = input_str.rsplit(":", 1)
        slice_idx = int(slice_str)
    else:
        input_path = input_str
        slice_idx = 0

    get_slice, n_slices, close_stack = open_stack(input_path)

    def get_image(idx):
        raw = get_slice(idx)
        return fourier_interpolate(raw, [x // args["binning"] for x in raw.shape])

    image = get_image(slice_idx)

    # Robust display range: avoid saturation from hot/dead pixels.
    vmin = np.percentile(image, 10)
    vmax = np.percentile(image, 90)

    image_median = float(np.median(image))
    initial_threshold = (
        args["maskabsolutethreshold"]
        if args["maskabsolutethreshold"] is not None
        else image_median * args["maskthreshold"]
    )

    fig, ax = plt.subplots(1, 2, figsize=(12, 6))
    plt.subplots_adjust(bottom=0.44)

    im0 = ax[0].imshow(image, vmin=vmin, vmax=vmax)
    ax[0].set_title(f"Original image  (slice {slice_idx} / {n_slices - 1})")
    ax[0].axis("off")

    initial_fringe_px = 20

    def make_title(fringe_px, threshold, median):
        fraction = threshold / median if median != 0 else 0.0
        return (
            f"Fringes masked  |  fringe = {fringe_px:.0f} px  |  "
            f"threshold = {threshold:.1f} counts = {fraction:.3f} × median"
        )

    mask = make_mask(
        image,
        initial_fringe_px / args["binning"],
        absolutethreshold=initial_threshold,
    )
    masked_display = ax[1].imshow(np.where(mask, image, 0), vmin=vmin, vmax=vmax)
    ax[1].set_title(make_title(initial_fringe_px, initial_threshold, image_median))
    ax[1].axis("off")

    # --- Sliders ---
    ax_fringe = plt.axes([0.2, 0.31, 0.65, 0.03])
    fringe_slider = Slider(
        ax_fringe, "Fringe size (unbinned px)", 1, 200,
        valinit=initial_fringe_px, valstep=1, valfmt="%0.0f",
    )

    threshold_slider = make_threshold_slider(
        fig, [0.2, 0.21, 0.65, 0.03], image, initial_threshold,
        label="Abs. threshold (counts)",
    )

    # Text line below the threshold slider showing the fraction of current median.
    update_frac_text = make_fraction_text(
        fig, 0.2, 0.155, prefix="  →  ", fontsize=8, color="dimgray",
    )
    update_frac_text(initial_threshold, image_median)

    # --- Slice navigation buttons ---
    slice_label = fig.text(
        0.5, 0.08, f"Slice {slice_idx} / {n_slices - 1}",
        ha="center", fontsize=10,
    )

    # Mutable state shared between callbacks.
    state = {"idx": slice_idx, "image": image, "median": image_median}

    def update(val):
        img = state["image"]
        med = state["median"]
        fringe_px = fringe_slider.val
        threshold = threshold_slider.val
        new_mask = make_mask(img, fringe_px / args["binning"], absolutethreshold=threshold)
        masked_display.set_data(np.where(new_mask, img, 0))
        ax[1].set_title(make_title(fringe_px, threshold, med))
        update_frac_text(threshold, med)
        fig.canvas.draw_idle()

    def load_slice(idx):
        idx = max(0, min(n_slices - 1, idx))
        state["idx"] = idx
        img = get_image(idx)
        state["image"] = img
        med = float(np.median(img))
        state["median"] = med
        new_vmin = np.percentile(img, 10)
        new_vmax = np.percentile(img, 90)
        im0.set_data(img)
        im0.set_clim(new_vmin, new_vmax)
        masked_display.set_clim(new_vmin, new_vmax)
        ax[0].set_title(f"Original image  (slice {idx} / {n_slices - 1})")
        slice_label.set_text(f"Slice {idx} / {n_slices - 1}")
        update(None)

    fringe_slider.on_changed(update)
    threshold_slider.on_changed(update)
    make_nav_buttons(
        fig, [0.30, 0.07, 0.13, 0.05], [0.57, 0.07, 0.13, 0.05], n_slices,
        load_slice, prev_label="← Prev slice", next_label="Next slice →",
        start=slice_idx,
    )

    plt.show()
    close_stack()


if __name__ == "__main__":
    main()
```
