## Supplementary material for "Whole cryo-FIB lamella acquisition at high magnification with rectangular beam montage cryo-electron tomography": Source code: imod_aligned_to_aln.py

```
"""Write an AreTomo-format .aln for a stack IMOD has already aligned.

AreTomo only writes a .aln when it solves its own alignment: pointing it at
an already-aligned stack via -AlnFile (which makes it auto-set -Align 0,
CInput.cpp) skips that step, and CSaveAlnFile.cpp's mSaveAlignment returns
early whenever alignment was skipped -- nothing gets written. But this
repo's montage_aretomo_record hard-requires an .aln to exist, and -AlnFile
is also how such a run is honestly recorded as alignment_source="supplied"
rather than "solved" (aretomo_record.py:362) -- so reconstructing an
IMOD-aligned stack through AreTomo still needs a real .aln on disk, just
one this writes instead of AreTomo.

This writes an identity one -- ROT=TX=TY=0, GMAG=1, one row per tilt in
stack order -- carrying only the real tilt angles from IMOD's .tlt. That
mirrors what IMOD's newstack + tiltalign already baked into the stack's
pixels: the shifts and tilt-axis rotation are applied, not left to AreTomo
to redo. NumPatches = 0: IMOD's alignment has no per-patch component to
record. Row format matches AreTomo2's own writer (MrcUtil/CSaveAlnFile.cpp)
so parse_aln (refine_montage_projmatch.py) reads it like any AreTomo file.

Usage:
    python -m processing_scripts.imod_aligned_to_aln \\
        --tlt imod/Montage_21-A.tlt \\
        --stack imod/Montage_21-A_ali.mrc \\
        --output AreTomo_from_imod/Montage_21-A_ali.aln
"""

import argparse

import mrcfile


def read_tilts(tlt_path: str) -> list[float]:
    with open(tlt_path) as fh:
        return [float(line.split()[0]) for line in fh if line.strip()]


def write_aln(output_path: str, raw_size: tuple[int, int, int],
              tilts: list[float]) -> None:
    with open(output_path, "w") as fh:
        fh.write("# AreTomo Alignment / Priims bprmMn \n")
        fh.write(f"# RawSize = {raw_size[0]} {raw_size[1]} {raw_size[2]}\n")
        fh.write("# NumPatches = 0\n")
        fh.write("# SEC     ROT         GMAG       TX          TY      "
                  "SMEAN     SFIT    SCALE     BASE     TILT\n")
        for i, tilt in enumerate(tilts):
            fh.write(f"{i:5d}  {0.0:9.4f}  {1.0:9.5f}  {0.0:9.3f}  "
                      f"{0.0:9.3f}  {1.0:7.2f}  {1.0:7.2f}  {1.0:7.2f}  "
                      f"{0.0:7.2f}  {tilt:8.2f}\n")


def main():
    ap = argparse.ArgumentParser(description=__doc__,
        formatter_class=argparse.RawDescriptionHelpFormatter)
    ap.add_argument("--tlt", required=True,
                     help="IMOD .tlt tilt-angle file, one angle per section, "
                          "in the same order as --stack")
    ap.add_argument("--stack", required=True,
                     help="the IMOD-aligned MRC stack this .aln is for "
                          "(read only for its dimensions, i.e. RawSize)")
    ap.add_argument("--output", required=True, help=".aln file to write")
    args = ap.parse_args()

    tilts = read_tilts(args.tlt)

    with mrcfile.open(args.stack, header_only=True, permissive=True) as mrc:
        raw_size = (int(mrc.header.nx), int(mrc.header.ny), int(mrc.header.nz))

    if len(tilts) != raw_size[2]:
        raise SystemExit(
            f"{args.tlt} has {len(tilts)} angles but {args.stack} has "
            f"{raw_size[2]} sections -- these must match 1:1, in the same "
            "order, or AreTomo will pair the wrong angle with the wrong "
            "section.")

    write_aln(args.output, raw_size, tilts)
    print(f"Wrote {args.output}: RawSize={raw_size}, {len(tilts)} sections, "
          "identity ROT/TX/TY (already applied by IMOD).")


if __name__ == "__main__":
    main()
```
