## Supplementary material for "Whole cryo-FIB lamella acquisition at high magnification with rectangular beam montage cryo-electron tomography": Source code: imod_tiltalign_to_aln.py

```
"""Write an AreTomo-format .aln from a real (non-identity) IMOD alignment.

Sibling of ``imod_aligned_to_aln.py``, which only covers the case where the
stack fed to AreTomo was *already* IMOD-aligned (so the honest .aln is
identity). Here the stack AreTomo/the rest of this pipeline needs to walk
back to is the RAW stitched canvas, and IMOD's alignment between the two --
tiltalign's per-section rotation/shift, combined with the prealignment -- is
a real, non-identity transform that has to be carried through, not discarded.

Where that transform lives, for a `tiltalign`-produced series (see
align.com): `tiltalign`'s OutputTransformFile (the raw .tltxf) only
describes the correction solved on top of the already-prealigned stack
(ImageFile in align.com is the _preali.mrc, not the raw canvas). align.com's
own `xfproduct` step immediately combines it with the prealignment (.prexg)
into one raw-canvas -> aligned-stack transform, `<series>_fid.xf` (then
copied to `<series>.xf`, and that is what `newst.com`'s TransformFile
actually applies). That combined .xf -- not the bare .tltxf -- is what
belongs here as --xf; using the bare .tltxf would silently drop the
prealignment shift.

Derivation of the ROT/GMAG/TX/TY this writes:

IMOD's .xf format (six numbers per line, coordinates centred on the image
middle) is

    q - centre = A @ (p - centre) + (DX, DY)

AreTomo's own .xf writer uses the canonical form (CSaveXF.cpp, also the form
this repo's aretomo_record.py / refine_montage_projmatch.py read out of a
real AreTomo .aln)

    q - centre = R(-ROT) @ (p - centre - (TX, TY)) + centre - centre
                = [GMAG * R(-ROT)] @ (p - centre) - [GMAG * R(-ROT)] @ (TX, TY)

so matching term by term: A = GMAG * R(-ROT), (DX, DY) = -A @ (TX, TY). Both
formats are centred, raw-canvas-pixel, degrees -- so this is an exact
decomposition, not a fit, PROVIDED A is a pure similarity (rotation + one
uniform scale, no shear/stretch). align.com has XStretchOption 0 and
SkewOption 0, so that holds here; ``decompose_row`` checks it against a
second, independent read of ROT off the matrix and refuses to write a row
where they disagree by more than 0.05 deg.

Usage:
    python -m processing_scripts.imod_tiltalign_to_aln \\
        --xf imod/Montage_21-A.xf \\
        --tlt imod/Montage_21-A.tlt \\
        --raw-stack stitched/Montage_21-A.mrc \\
        --output AreTomo_from_imod/Montage_21-A_tiltalign.aln
"""

import argparse
import math

import mrcfile


def read_tilts(tlt_path: str) -> list[float]:
    with open(tlt_path) as fh:
        return [float(line.split()[0]) for line in fh if line.strip()]


def read_xf(xf_path: str) -> list[tuple[float, float, float, float, float, float]]:
    rows = []
    with open(xf_path) as fh:
        for line in fh:
            line = line.strip()
            if not line:
                continue
            nums = [float(x) for x in line.split()]
            if len(nums) != 6:
                raise SystemExit(
                    f"{xf_path}: expected 6 numbers per line (A11 A12 A21 "
                    f"A22 DX DY), got {len(nums)}: {line!r}")
            rows.append(tuple(nums))
    return rows


def decompose_row(
    a11: float, a12: float, a21: float, a22: float, dx: float, dy: float,
) -> tuple[float, float, float, float]:
    """One IMOD .xf row -> AreTomo-style (rot_deg, gmag, tx, ty). See module
    docstring for the derivation."""
    det = a11 * a22 - a12 * a21
    if det <= 0:
        raise SystemExit(
            f"Row [{a11} {a12} {a21} {a22} {dx} {dy}] is not a proper "
            f"rotation+scale (det={det:.6g} <= 0) -- a reflection or "
            "degenerate transform, which this decomposition cannot handle.")
    gmag = math.sqrt(det)

    rot = math.degrees(math.atan2(a12, a11))
    rot_check = math.degrees(math.atan2(-a21, a22))
    diff = (rot - rot_check + 180) % 360 - 180
    if abs(diff) > 0.05:
        raise SystemExit(
            f"Row [{a11} {a12} {a21} {a22} {dx} {dy}] is not a clean "
            f"similarity transform (ROT from A11/A12={rot:.4f} deg, from "
            f"A21/A22={rot_check:.4f} deg, disagree by {diff:.4f} deg) -- "
            "XStretchOption/SkewOption may not have been 0 for this "
            "alignment, and this decomposition does not apply to it.")

    # (TX, TY) = -A^-1 @ (DX, DY); A^-1 = (1/det) * [[A22,-A12],[-A21,A11]]
    inv_dx = (a22 * dx - a12 * dy) / det
    inv_dy = (-a21 * dx + a11 * dy) / det
    return rot, gmag, -inv_dx, -inv_dy


def write_aln(
    output_path: str, raw_size: tuple[int, int, int],
    tilts: list[float], rows: list[tuple[float, float, float, float]],
) -> None:
    with open(output_path, "w") as fh:
        fh.write("# AreTomo Alignment / Priims bprmMn \n")
        fh.write(f"# RawSize = {raw_size[0]} {raw_size[1]} {raw_size[2]}\n")
        fh.write("# NumPatches = 0\n")
        fh.write("# SEC     ROT         GMAG       TX          TY      "
                  "SMEAN     SFIT    SCALE     BASE     TILT\n")
        for i, (tilt, (rot, gmag, tx, ty)) in enumerate(zip(tilts, rows)):
            fh.write(f"{i:5d}  {rot:9.4f}  {gmag:9.5f}  {tx:9.3f}  "
                      f"{ty:9.3f}  {1.0:7.2f}  {1.0:7.2f}  {1.0:7.2f}  "
                      f"{0.0:7.2f}  {tilt:8.2f}\n")


def main():
    ap = argparse.ArgumentParser(description=__doc__,
        formatter_class=argparse.RawDescriptionHelpFormatter)
    ap.add_argument("--xf", required=True,
                     help="IMOD's combined raw-canvas -> aligned-stack "
                          "transform (prealignment x tiltalign -- e.g. "
                          "xfproduct's output/newst.com's TransformFile, "
                          "NOT the bare tiltalign .tltxf -- see module "
                          "docstring), one row per section in stack order")
    ap.add_argument("--tlt", required=True,
                     help="IMOD .tlt tilt-angle file, same order as --xf")
    ap.add_argument("--raw-stack", required=True,
                     help="the RAW (pre-alignment) stack --xf maps from -- "
                          "read only for its dimensions, i.e. RawSize. For a "
                          "stitched montage this is the stitched canvas "
                          "(e.g. stitched/Montage_21-A.mrc), NOT the "
                          "IMOD-aligned stack newstack wrote.")
    ap.add_argument("--output", required=True, help=".aln file to write")
    args = ap.parse_args()

    tilts = read_tilts(args.tlt)
    xf_rows = read_xf(args.xf)
    if len(tilts) != len(xf_rows):
        raise SystemExit(
            f"{args.tlt} has {len(tilts)} angles but {args.xf} has "
            f"{len(xf_rows)} rows -- these must match 1:1, in the same "
            "order, or a tilt will be paired with the wrong section's "
            "transform.")

    with mrcfile.open(args.raw_stack, header_only=True, permissive=True) as mrc:
        raw_size = (int(mrc.header.nx), int(mrc.header.ny), int(mrc.header.nz))
    if raw_size[2] != len(tilts):
        raise SystemExit(
            f"{args.raw_stack} has {raw_size[2]} sections but {args.tlt} "
            f"has {len(tilts)} angles -- --raw-stack must be the actual "
            "stack --xf was solved against (same section count), not a "
            "differently-cropped copy.")

    decomposed = [decompose_row(*row) for row in xf_rows]
    write_aln(args.output, raw_size, tilts, decomposed)
    print(f"Wrote {args.output}: RawSize={raw_size}, {len(tilts)} sections, "
          "ROT/GMAG/TX/TY decomposed from IMOD's .xf.")


if __name__ == "__main__":
    main()
```
