## Supplementary material for "Whole cryo-FIB lamella acquisition at high magnification with rectangular beam montage cryo-electron tomography": Source code: inpaint_apply.py

```
#!/usr/bin/env python3
"""
Parallel inpainter — companion to mask_and_inpaint.py / mask_and_inpaint_peak.py.

Reads a JSON params file generated by the interactive GUI (which stores the
per-tilt (lo, hi) intensity thresholds), then masks and inpaints every tilt in
parallel using ProcessPoolExecutor.  Designed to run on cluster nodes via SLURM.

Usage
-----
    python inpaint_apply.py --input file.mrc --params file_params.json \\
                            --output file_inpainted.mrc --cpus 16

JSON params format (produced by the GUI's "Save SLURM" button)
---------------------------------------------------------------
    {
      "input":        "/abs/path/to/input.mrc",
      "output":       "/abs/path/to/output.mrc",
      "save_mask":    true,                       # optional; defaults to true.
                                                  # --dont-save-mask overrides it.
      "mask_mrc":     "/abs/path/to/mask.mrc",   # optional; present when paint/polygon
                                                  # overlays were drawn in the GUI.
                                                  # When present, the spatial mask
                                                  # overrides per-tilt lo/hi thresholds.
      "tilts": {
        "0":  {"lo": 40.5,  "hi": 253.0},
        "1":  {"lo": 38.2,  "hi": 248.1},
        ...
      }
    }
"""

# Postponed annotation evaluation: the sidecar helpers below use "str | None",
# which is a runtime TypeError on the Python 3.9 in the cluster's conda envs
# unless annotations are strings.
from __future__ import annotations

import argparse
import json
import logging
import multiprocessing
import os
import sys
import time
from concurrent.futures import ProcessPoolExecutor, as_completed

import numpy as np
import numpy.ma as ma
import mrcfile
from scipy.ndimage import label, uniform_filter, zoom
from tqdm import tqdm

try:
    from .smoothn import smoothn
    from . import sidecar
except ImportError:
    from smoothn import smoothn
    import sidecar

logger = logging.getLogger(__name__)

#: Name of this step in its sidecars. The mask that drives it is
#: recorded separately, by mask_and_inpaint.py, as "inpaint_mask".
SIDECAR_STEP = "inpaint_apply"


# ---------------------------------------------------------------------------
# Default SLURM template — imported by the GUI scripts
# ---------------------------------------------------------------------------

DEFAULT_SLURM_TEMPLATE = """\
#!/bin/bash
#SBATCH --job-name={job_name}
#SBATCH --ntasks=1
#SBATCH --cpus-per-task={n_cpus}
#SBATCH --mem={mem_gb}G
#SBATCH --time={time}
#SBATCH --output={log_file}

python {script_path} \\
    --input    {input} \\
    --output   {output} \\
    --params   {params} \\
    --ntotal   {n_cpus} \\
    --npertilt {npertilt} \\
    --bin      {bin_factor} \\
    --variant  full

# ---------------------------------------------------------------------------
# Odd/even half-set inpainting, for Noise2Noise denoising (cryoCARE etc.).
# Uncomment both blocks to also inpaint the half-sets written by
# beam_mask_motioncorr.py.
#
# --mask points both halves at the same mask stack the full-dose run uses, so
# all three stacks are masked identically.  This is essential and not optional:
# without it the half-sets would re-derive a mask from the lo/hi thresholds in
# the params file, which were fitted to full-dose intensities and do not
# transfer to half-dose data.  A pixel that is real data in one half and
# synthetic fill in the other is a signal mismatch that the denoiser treats as
# noise and tries to average away.
#
# The fill itself is still fitted independently to each half's own surrounding
# pixels, so no intensity rescaling is needed.
#
# The mask below is normally the one baked by the GUI at "Save SLURM Job" time,
# so it already exists and these can run in any order.  --dont-save-mask stops
# them writing redundant copies of it.
#
# --variant labels each run. The three write different output stacks, so each
# gets its own sidecar beside its own stack with no keying needed.
# ---------------------------------------------------------------------------

# python {script_path} \\
#     --input    {input_odd} \\
#     --output   {output_odd} \\
#     --params   {params} \\
#     --mask     {mask} \\
#     --ntotal   {n_cpus} \\
#     --npertilt {npertilt} \\
#     --bin      {bin_factor} \\
#     --variant  odd \\
#     --dont-save-mask

# python {script_path} \\
#     --input    {input_even} \\
#     --output   {output_even} \\
#     --params   {params} \\
#     --mask     {mask} \\
#     --ntotal   {n_cpus} \\
#     --npertilt {npertilt} \\
#     --bin      {bin_factor} \\
#     --variant  even \\
#     --dont-save-mask
"""


# ---------------------------------------------------------------------------
# Inpainting
# ---------------------------------------------------------------------------
#
# One implementation, used by both the cluster runner below and the GUI's
# "Generate" button in mask_and_inpaint.py. They used to differ — the GUI let
# smoothn choose s per tilt by GCV and then feathered a 15 px rim of real data
# into the fill, the cluster run used a fixed s and a hard boundary — so the
# preview never showed what the job would write.

#: Smoothing parameter handed to smoothn. Large is stiff: the fill is a smooth
#: continuation of the surrounding data rather than an attempt to invent
#: detail. Not auto-selected per tilt, so every tilt is filled alike.
SMOOTHN_S = 1e7

#: Iteration cap for smoothn's DCT solve.
SMOOTHN_MAX_ITER = 500


def inpaint_masked_tilt(img: np.ndarray, mask: np.ndarray, *,
                        bin_factor: int = 2, workers: int = 1,
                        smoothn_s: float = SMOOTHN_S,
                        max_iter: int = SMOOTHN_MAX_ITER) -> np.ndarray:
    """Fill the pixels where *mask* is False; return float32.

    ``mask``: True = keep, False = inpaint. Only masked-out pixels are ever
    written, so the boundary is followed at full resolution whatever
    ``bin_factor`` is — the binning only makes the smoothn fit cheaper (see
    :func:`_bin_good_mean`), it does not soften the edge.
    """
    bad = ~mask
    if not bad.any():
        return img.astype(np.float32)

    if bin_factor > 1:
        binned_img, binned_good = _bin_good_mean(img, mask, bin_factor)
        binned_field = smoothn(ma.array(binned_img, mask=~binned_good),
                               s=smoothn_s, max_iter=max_iter, workers=workers)
        inpainted = _upsample_to(binned_field, img.shape)
    else:
        inpainted = smoothn(ma.array(img, mask=bad), s=smoothn_s,
                            max_iter=max_iter, workers=workers)

    result = img.copy()
    result[bad] = inpainted[bad]
    return result.astype(np.float32)


# ---------------------------------------------------------------------------
# Binning helpers
# ---------------------------------------------------------------------------

def _bin_good_mean(img: np.ndarray, good: np.ndarray, bin_factor: int) -> tuple:
    """
    Block-downsample *img* by *bin_factor*, ignoring bad pixels (weight 0) so
    they can't leak into good samples near the mask boundary.

    Returns (binned_img, binned_good).  A binned pixel is only marked good if
    a clear majority of the input pixels in its block were good; otherwise it
    is left for smoothn to fill in, same as at full resolution.
    """
    good_f = good.astype(np.float32)
    num = uniform_filter(img.astype(np.float32) * good_f, size=bin_factor, mode="nearest")
    den = uniform_filter(good_f, size=bin_factor, mode="nearest")
    valid = den > 0.5
    filled = np.zeros_like(num)
    filled[valid] = num[valid] / den[valid]
    return filled[::bin_factor, ::bin_factor], valid[::bin_factor, ::bin_factor]


def _upsample_to(field: np.ndarray, shape: tuple) -> np.ndarray:
    """Upsample *field* back to *shape* with bilinear interpolation."""
    zh, zw = shape[0] / field.shape[0], shape[1] / field.shape[1]
    up = zoom(field, (zh, zw), order=1)
    up = up[:shape[0], :shape[1]]
    if up.shape != shape:
        up = np.pad(up, ((0, shape[0] - up.shape[0]), (0, shape[1] - up.shape[1])),
                    mode="edge")
    return up


# ---------------------------------------------------------------------------
# Per-tilt worker  (module-level so ProcessPoolExecutor can pickle it)
# ---------------------------------------------------------------------------

def _process_tilt(args: tuple) -> tuple:
    """
    Load one tilt from the mmap'd MRC, mask it, inpaint with smoothn, and
    return (tilt_idx, result_float32, mask_int8).

    Opening the MRC inside each worker avoids serialising large arrays across
    the process boundary while still allowing concurrent reads (mmap, read-only).

    If mask_path is provided the pre-baked spatial mask is used directly (paint/
    polygon overlays from the GUI are already baked in); otherwise the mask is
    recomputed from the lo/hi intensity thresholds.

    If bin_factor > 1, smoothn is run on a downsampled copy of the tilt (bad
    pixels excluded from the block average) and the resulting smooth field is
    upsampled back to full resolution before being applied to the bad pixels;
    this is much cheaper than running smoothn at full resolution and, since
    only the already-bad pixels are ever overwritten, the mask boundary is
    still followed at full resolution regardless of the bin factor.
    """
    tilt_idx, input_path, lo, hi, smoothn_workers, mask_path, bin_factor = args

    # Concurrent read-only mmap access is safe on all major OS/filesystems
    with mrcfile.mmap(input_path, mode='r', permissive=True) as mrc:
        img = np.array(mrc.data[tilt_idx], dtype=float)

    if mask_path is not None:
        # Use the pre-baked spatial mask (includes paint/polygon overlays)
        with mrcfile.mmap(mask_path, mode='r', permissive=True) as mrc:
            mask = mrc.data[tilt_idx].astype(bool)
    else:
        # --- mask: largest connected component inside [lo, hi] ---
        raw = (img >= lo) & (img <= hi)
        if raw.any():
            labeled, n_feat = label(raw)
            if n_feat > 0:
                sizes = np.bincount(labeled.ravel())
                sizes[0] = 0
                mask = (labeled == sizes.argmax()).astype(bool)
            else:
                mask = raw.astype(bool)
        else:
            mask = raw.astype(bool)

    result = inpaint_masked_tilt(img, mask, bin_factor=bin_factor,
                                 workers=smoothn_workers)

    return tilt_idx, result, mask.astype(np.int8)


# ---------------------------------------------------------------------------
# Main runner
# ---------------------------------------------------------------------------

def run(input_path: str, output_path: str, params: dict,
        ntotal: int = None, npertilt: int = 4,
        bin_factor: int = 2, save_mask: bool = True,
        write_sidecar: bool = True, variant: str = "full") -> None:
    """
    Apply per-tilt masks from *params["tilts"]* to *input_path* and write the
    inpainted stack to *output_path*.

    Tilts are processed concurrently: *ntotal* is the total thread budget and
    *npertilt* is how many of those threads smoothn's DCT/IDCT get per tilt,
    so ``ntotal // npertilt`` tilts run at once, each in its own process.
    Lower *npertilt* trades DCT thread-parallelism (which saturates well
    before 8-16 threads, more so once binning has shrunk the array) for more
    concurrent tilts, which also caps peak memory per tilt.

    If *save_mask* is True (the default), also write the binary mask stack to
    ``<output_stem>_mask.mrc`` (mrc_mode=0, int8).  When a pre-baked
    ``mask_mrc`` is already referenced in *params*, the same mask data is
    re-emitted per-tilt so both outputs sit alongside each other.

Unless *write_sidecar* is False, a provenance sidecar is written beside the
    output stack holding the step's parameters and one row per tilt, each
    carrying the fraction of that tilt which is synthetic — the number that
    otherwise exists nowhere, and the one worth knowing before picking
    particles in a region that was filled in. *variant* distinguishes the
    full-dose run from the odd/even half-set runs, which write different
    stacks from the same mask; each is its own output file with its own
    sidecar.
    """
    tilt_params = params["tilts"]
    n_tilts     = len(tilt_params)
    total_cpus = multiprocessing.cpu_count()
    if ntotal is None:
        ntotal = total_cpus
    npertilt = max(1, npertilt)
    n_concurrent = max(1, ntotal // npertilt)

    mask_path = params.get("mask_mrc")   # None when no paint/polygon overlays were saved

    print(f"Input           : {input_path}")
    print(f"Output          : {output_path}")
    print(f"Tilts           : {n_tilts}")
    print(f"Total threads   : {ntotal}")
    print(f"Threads/tilt    : {npertilt}")
    print(f"Concurrent tilts: {n_concurrent}")
    print(f"Bin factor      : {bin_factor}")
    print(f"smoothn s       : {SMOOTHN_S:g}")
    if mask_path:
        print(f"Spatial mask    : {mask_path}")
    else:
        print("Spatial mask    : none  (using lo/hi intensity thresholds)")

    with mrcfile.mmap(input_path, mode='r', permissive=True) as mrc:
        shape = tuple(mrc.data.shape)
        voxel_size = mrc.voxel_size.copy()

    # A supplied mask is only meaningful if it matches the stack it is applied to.
    # Failing loudly here beats silently inpainting the wrong pixels — particularly
    # for odd/even half-sets, which must share the full-dose run's mask exactly.
    if mask_path:
        if not os.path.exists(mask_path):
            raise FileNotFoundError(
                f"Mask stack not found: {mask_path}\n"
                "If this is an odd/even half-set run, the full-dose run writes the "
                "mask and must complete first."
            )
        with mrcfile.mmap(mask_path, mode='r', permissive=True) as mrc:
            mask_shape = tuple(mrc.data.shape)
        if mask_shape != shape:
            raise ValueError(
                f"Mask shape {mask_shape} does not match input shape {shape} "
                f"({mask_path})"
            )
        # A mask older than the stack it masks usually means the stack was re-stitched
        # after the mask was baked.  Shape alone won't catch this, so warn loudly —
        # but don't refuse, since copies and archive restores can reset mtimes.
        if os.path.getmtime(mask_path) < os.path.getmtime(input_path):
            print(
                "\n"
                "  ****************************  WARNING  ****************************\n"
                "  The mask stack is OLDER than the input stack:\n"
                f"    mask  {time.strftime('%Y-%m-%d %H:%M:%S', time.localtime(os.path.getmtime(mask_path)))}  {mask_path}\n"
                f"    input {time.strftime('%Y-%m-%d %H:%M:%S', time.localtime(os.path.getmtime(input_path)))}  {input_path}\n"
                "  If the input was re-stitched since the mask was made, re-bake the\n"
                "  mask from the GUI before trusting this result.\n"
                "  *******************************************************************\n",
                file=sys.stderr, flush=True,
            )

    # Re-emitting a mask that was handed to us would just duplicate an existing
    # multi-GB file, so skip it and point at the original instead.
    if save_mask and mask_path:
        print("Mask output     : skipped (mask was supplied, not derived)")
        save_mask = False

    mask_out_path = os.path.splitext(output_path)[0] + "_mask.mrc" if save_mask else None

    # Verified before the work, written after it: the sidecar carries each
    # tilt's filled fraction, which does not exist until the tilts are done.
    sidecar_warnings = (
        verify_inpaint_inputs(input_path, mask_path) if write_sidecar else [])

    tasks = [
        (i, input_path, float(tilt_params[str(i)]["lo"]), float(tilt_params[str(i)]["hi"]),
         npertilt, mask_path, bin_factor)
        for i in range(n_tilts)
    ]

    # Filled per tilt as the results land, then spooled below: the mask is in
    # hand here anyway, so the cost of knowing how much of each tilt is
    # synthetic is one mean() per tilt.
    filled_fraction: dict = {}

    def _keep(idx, result, mask, out_mrc, mask_mrc):
        out_mrc.data[idx] = result
        if mask_mrc is not None:
            mask_mrc.data[idx] = mask
        filled_fraction[idx] = float(1.0 - (mask.astype(bool).mean()))

    def _write_tilts(out_mrc, mask_mrc):
        if n_concurrent <= 1:
            for task in tqdm(tasks, desc="Inpainting tilts"):
                idx, result, mask = _process_tilt(task)
                _keep(idx, result, mask, out_mrc, mask_mrc)
            return

        with ProcessPoolExecutor(max_workers=n_concurrent) as executor:
            futures = [executor.submit(_process_tilt, task) for task in tasks]
            for future in tqdm(as_completed(futures), total=len(futures),
                               desc="Inpainting tilts"):
                idx, result, mask = future.result()
                _keep(idx, result, mask, out_mrc, mask_mrc)

    with mrcfile.new_mmap(output_path, shape=shape, mrc_mode=2, overwrite=True) as out_mrc:
        # Carry the pixel size over. Without it the output header says 0, and
        # anything downstream that trusts the header silently uses 1 A/px --
        # 3dmod rescales an IMOD model by the ratio and throws every point clean
        # off the canvas, which looks like the model being empty.
        out_mrc.voxel_size = voxel_size
        if mask_out_path:
            with mrcfile.new_mmap(mask_out_path, shape=shape, mrc_mode=0, overwrite=True) as mask_mrc:
                mask_mrc.voxel_size = voxel_size
                _write_tilts(out_mrc, mask_mrc)
            print(f"Mask  → {mask_out_path}")
        else:
            _write_tilts(out_mrc, None)

    print(f"Saved → {output_path}")

    if write_sidecar:
        effective, requested = inpaint_effective(
            variant=variant, input_path=input_path, output_path=output_path,
            mask_out_path=mask_out_path, mask_path=mask_path, params=params,
            bin_factor=bin_factor, save_mask=save_mask, n_tilts=n_tilts,
            shape=shape, voxel_size=voxel_size, ntotal=ntotal,
            npertilt=npertilt, tilt_params=tilt_params,
            filled_fraction=filled_fraction,
        )
        write_inpaint_sidecar(
            input_path=input_path, output_path=output_path,
            mask_path=mask_path, effective=effective, requested=requested,
            warnings=sidecar_warnings,
        )

# ---------------------------------------------------------------------------
# Provenance sidecar
# ---------------------------------------------------------------------------
#
# Unlike motion correction and stitching this is a single process writing a
# whole stack, so its sidecar is the simple unkeyed `<output>.json` and the
# per-tilt table is inlined in it. What this step does have is a handoff — the
# mask was designed in another process, possibly days earlier — so the
# verification below matters more here than the concurrency did.


def series_base_for_input(path: str) -> str:
    """Tilt series base name from a stack this step reads or writes."""
    stem = os.path.splitext(os.path.basename(path))[0]
    # Repeated because these compose: Series_A_odd_inpainted_mask.mrc is one
    # file this step can be pointed at. Longest first, so "_inpainted" is not
    # left as a stray "ed" by "_inpaint".
    suffixes = ("_inpainted", "_inpaint", "_odd", "_even", "_mask")
    stripped = True
    while stripped:
        stripped = False
        for suffix in suffixes:
            if stem.endswith(suffix):
                stem = stem[: -len(suffix)]
                stripped = True
    return stem


def verify_inpaint_inputs(input_path: str, mask_path: str | None) -> list:
    """Check the stack and mask against what earlier steps recorded.

    The mtime warning this step already prints is a proxy for the real
    question — was this mask made from *this* stack? — and it cannot tell a
    re-stitch from an archive restore. Fingerprints can.

    Returns the warnings so they can go into the sidecar as well as the log:
    the stack is already inpainted by the time anyone reads the log.
    """
    warnings = []

    doc = sidecar.read_for(input_path)
    if doc is None:
        warnings.append(
            f"{os.path.abspath(input_path)} carries no sidecar, so which step "
            "produced it — and with what geometry — is not recorded")
    elif doc.get("step") != "stitch":
        warnings.append(
            f"{os.path.abspath(input_path)} was written by "
            f"{doc.get('step')!r}, not by stitching")
    else:
        recorded = doc.get("output_fingerprint") or {}
        if recorded.get("sha1"):
            now = sidecar.fingerprint(input_path)
            if now["sha1"] != recorded["sha1"]:
                warnings.append(
                    "the input stack has changed since stitching recorded "
                    "writing it")

    if mask_path:
        mask_doc = sidecar.read_for(mask_path)
        if mask_doc is None:
            warnings.append(
                f"mask {os.path.abspath(mask_path)} carries no sidecar; it may "
                "be from a different session than this stack")
        else:
            for problem in sidecar.verify_inputs(mask_doc):
                warnings.append(f"the mask's own inputs: {problem}")

    for message in warnings:
        logger.warning("Sidecar: %s", message)
    if not warnings:
        logger.info("Sidecar: input stack and mask both check out.")
    return warnings


def inpaint_effective(
    *, variant: str, input_path: str, output_path: str,
    mask_out_path: str | None, mask_path: str | None, params: dict,
    bin_factor: int, save_mask: bool, n_tilts: int, shape, voxel_size,
    ntotal, npertilt: int, tilt_params: dict, filled_fraction: dict,
) -> tuple[dict, dict]:
    """This run's effective parameters, including the per-tilt table.

    The per-tilt rows — thresholds and how much of each tilt is now synthetic
    — are inlined rather than spooled. There are as many as there are tilts,
    which is tens; the JSONL spool and the fold job that used to carry them
    existed for motion correction's thousands, not for this.
    """
    source = "baked_mask" if mask_path else "thresholds"
    tilts = []
    for index in sorted(int(k) for k in tilt_params):
        entry = tilt_params[str(index)]
        tilts.append({
            "tilt_index": int(index),
            "lo": float(entry["lo"]),
            "hi": float(entry["hi"]),
            "mask_source": source,
            "filled_fraction": float(
                filled_fraction.get(index, float("nan"))),
        })

    filled = [t["filled_fraction"] for t in tilts
              if t["filled_fraction"] == t["filled_fraction"]]  # drop NaN
    effective = {
        "variant": variant,
        "input": os.path.abspath(input_path),
        "output": os.path.abspath(output_path),
        "mask_output": os.path.abspath(mask_out_path) if mask_out_path else None,
        "params_file": params.get("_params_path"),
        # Which of the two ways the mask was arrived at. "baked_mask" means a
        # spatial mask stack overrode the thresholds, which is the only mode
        # in which the odd/even halves can be masked identically to the full
        # dose — their intensities do not support the same lo/hi.
        "mask_source": source,
        "mask_mrc": os.path.abspath(mask_path) if mask_path else None,
        "smoothn_s": float(SMOOTHN_S),
        "smoothn_max_iter": int(SMOOTHN_MAX_ITER),
        "bin_factor": int(bin_factor),
        "save_mask": bool(save_mask),
        "n_tilts": int(n_tilts),
        "stack_shape": [int(v) for v in shape],
        "pixel_size_A": float(voxel_size.x),
        "filled_fraction_mean": (sum(filled) / len(filled)) if filled else None,
        "filled_fraction_max": max(filled) if filled else None,
        "tilts": tilts,
        # Inpainting replaces pixel values and moves nothing, so the canvas
        # frame stitching declared still describes this stack exactly. Said
        # out loud because the alternative — a reader assuming a new frame and
        # looking for a transform that does not exist — is the failure this
        # record is for.
        "frame": "canvas",
        "geometry_unchanged": True,
    }
    requested = {
        "bin_factor": int(bin_factor),
        "ntotal": ntotal,
        "npertilt": int(npertilt),
        "variant": variant,
    }
    return effective, requested


def write_inpaint_sidecar(
    *, input_path: str, output_path: str, mask_path: str | None,
    effective: dict, requested: dict, warnings: list,
) -> None:
    """One sidecar beside the inpainted stack.

    ``inputs`` names both the stitched stack and the mask, so "which mask was
    this stack inpainted with, and was it made from this stack" is one
    chain-hop rather than a cross-reference between two records.
    """
    inputs = [sidecar.fingerprint(input_path, role="stitched_stack")]
    if mask_path and os.path.exists(mask_path):
        inputs.append(sidecar.fingerprint(mask_path, role="mask"))

    sidecar.write(
        output_path,
        SIDECAR_STEP,
        effective,
        inputs=inputs,
        requested=requested,
        extra={"warnings": warnings} if warnings else None,
    )


def main() -> None:
    parser = argparse.ArgumentParser(
        description="Parallel MRC inpainter driven by a GUI-produced JSON params file.",
        formatter_class=argparse.RawDescriptionHelpFormatter,
        epilog=__doc__,
    )
    parser.add_argument("--input",  required=True,
                        help="Input MRC tilt series")
    parser.add_argument("--output", default=None,
                        help="Output MRC (default: <input>_inpainted.mrc)")
    parser.add_argument("--params", required=True,
                        help="JSON params file from the interactive GUI")
    parser.add_argument("--ntotal", type=int, default=None, dest="ntotal",
                        help="Total thread budget shared across concurrent tilts "
                             "(default: all CPUs)")
    parser.add_argument("--npertilt", type=int, default=4, dest="npertilt",
                        help="DCT/IDCT threads per tilt inside smoothn; "
                             "ntotal // npertilt tilts run concurrently, each in "
                             "its own process (default: 4)")
    parser.add_argument("--bin", type=int, default=2, dest="bin_factor",
                        help="Downsample factor used for the smoothn fit before "
                             "upsampling back to full resolution; 1 disables "
                             "binning (default: 2)")
    parser.add_argument("--mask", default=None, dest="mask_path",
                        help="Pre-baked binary mask stack (int8 MRC) to use instead "
                             "of the per-tilt lo/hi thresholds; overrides the params "
                             "file's mask_mrc entry. Use this to force odd/even "
                             "half-sets through exactly the same mask as the "
                             "full-dose run, whose lo/hi thresholds do not transfer "
                             "to half-intensity data")
    parser.add_argument("--no-sidecar", dest="sidecar", action="store_false",
                        default=True,
                        help="Do not write a provenance sidecar beside the "
                             "inpainted stack. Sidecars are ON by default and "
                             "record this step's effective parameters, the "
                             "mask it used, and one row per tilt carrying the "
                             "fraction of that tilt which is now synthetic — "
                             "the number that otherwise exists nowhere")
    parser.add_argument("--variant", default="full",
                        choices=["full", "odd", "even"],
                        help="Which dose fraction this run inpaints (default: "
                             "full). The half-sets write different stacks from "
                             "the same mask; each is its own output file with "
                             "its own sidecar, so this is a label rather than "
                             "a record key as it was under the manifest")
    parser.add_argument("--dont-save-mask", action="store_true", default=False,
                        dest="dont_save_mask",
                        help="Do not write the binary mask stack to "
                             "<output>_mask.mrc (int8, mrc_mode=0); the mask "
                             "is written by default")
    args = parser.parse_args()
    logging.basicConfig(level=logging.INFO, format="%(levelname)-8s %(message)s")

    for path, label_ in [(args.input, "input"), (args.params, "params")]:
        if not os.path.exists(path):
            sys.exit(f"Error: {label_} file not found: {path}")

    output = args.output or os.path.splitext(args.input)[0] + "_inpainted.mrc"

    with open(args.params) as f:
        params = json.load(f)
    # Recorded rather than re-derived: which params file drove this run is
    # part of the answer to "how was this stack made".
    params["_params_path"] = os.path.abspath(args.params)

    # Mask saving is on by default.  The GUI's "Save mask MRC" checkbox writes
    # save_mask into the JSON (also on by default), and --dont-save-mask is an
    # override that turns it off regardless — used by the odd/even half-set runs,
    # which would otherwise each rewrite the same mask stack.
    save_mask = bool(params.get("save_mask", True)) and not args.dont_save_mask

    # --mask overrides the params file, so half-set runs can point at the mask the
    # full-dose run wrote rather than re-deriving one from thresholds that no
    # longer apply.
    if args.mask_path:
        params["mask_mrc"] = os.path.abspath(args.mask_path)

    run(args.input, output, params,
        ntotal=args.ntotal, npertilt=args.npertilt,
        bin_factor=args.bin_factor, save_mask=save_mask,
        write_sidecar=args.sidecar, variant=args.variant)


if __name__ == "__main__":
    main()
```
