## Supplementary material for "Whole cryo-FIB lamella acquisition at high magnification with rectangular beam montage cryo-electron tomography": Source code: JointoMRC.py

```
"""
Join a tilt-series of stitched montage TIFFs into a single MRC stack for IMOD/AreTomo,
writing companion files: .rawtlt, tiltAngles.txt, and an HDF5 file containing the
refined tile positions and index maps from each tilt.

Deprecated, because the new stitch.py workflow now writes a single MRC and .rawtlt directly.
"""

import argparse
import glob
import os
import re

import mrcfile
import numpy as np
from PIL import Image
from tqdm import tqdm

from .Utilities import load_array_from_hdf5, save_array_to_hdf5

# Disable PIL's safety limit — montage TIFFs are legitimately very large.
Image.MAX_IMAGE_PIXELS = None


def parse_commandline():
    parser = argparse.ArgumentParser(
        description="Join montage TIFF files into a single MRC for IMOD, filling blank areas."
    )
    parser.add_argument(
        "-i", "--input",
        required=True,
        type=str,
        help="Directory containing stitched montage tilt series TIFF files",
    )
    parser.add_argument(
        "-o", "--output",
        required=False,
        type=str,
        help="Output directory (default: same as input)",
    )
    parser.add_argument(
        "-I", "--image_shifts",
        required=False,
        type=str,
        help="Path to text file containing tilts and image shifts at every tilt",
    )
    parser.add_argument(
        "-p", "--pixel_size",
        required=True,
        type=float,
        help="Pixel size in Angstroms (of the binned, stitched montage tiles).",
    )
    parser.add_argument(
        "-O", "--acquisition_order",
        action="store_true",
        help="Stack images in acquisition order (by file mtime) instead of tilt order "
             "(e.g. for Relion dose weighting).",
    )
    return vars(parser.parse_args())


def extract_number(filename):
    """Return the tilt angle embedded in a filename like Montage_-18.0.tif."""
    match = re.search(r"([-]?\d+\.?\d*)\.tif", filename)
    return float(match.group(1)) if match else 0


def sort_files(fnams, acquisition_order=True):
    """Sort TIFFs either by acquisition time (for dose weighting) or by tilt angle.

    Acquisition order uses each file's mtime rather than any embedded
    metadata: stitch.py preserves the source raw montage's mtime on its
    stitched output, so file mtime order matches true acquisition order.
    """
    if acquisition_order:
        return sorted(fnams, key=os.path.getmtime)
    return sorted(fnams, key=extract_number)


def main():
    args = parse_commandline()
    inputdir = args["input"]
    outputdir = args["output"] if args["output"] is not None else inputdir

    fnams = glob.glob(os.path.join(inputdir, "Montage*.tif"))
    fnams = sort_files(fnams, acquisition_order=args["acquisition_order"])

    # Filter to TIFFs that share the canvas size of the first file; warn and skip others.
    with Image.open(fnams[0]) as img0:
        first_size = img0.size
        first_dtype = np.array(img0).dtype
    compliant = [fnams[0]]
    for fnam in fnams[1:]:
        with Image.open(fnam) as img:
            if img.size != first_size:
                print(f"Warning: skipping {fnam} — size {img.size} does not match {first_size}")
            else:
                compliant.append(fnam)
    fnams = compliant

    # PIL returns (width, height); numpy/MRC use (height, width) — hence the reversal.
    # Dimensions are rounded down to even numbers for MRC compatibility.
    shape = (len(fnams), *[2 * (x // 2) for x in first_size[::-1]])

    outname = os.path.basename(os.path.commonprefix(fnams)) or "tilt_series"
    outputfile = os.path.join(outputdir, outname + ".mrc")

    # Match the MRC mode/dtype to the input TIFFs' own dtype rather than forcing int16.
    try:
        mrc_mode = mrcfile.utils.mode_from_dtype(first_dtype)
    except ValueError:
        if first_dtype == np.dtype(np.int32):
            # PIL has no native signed 8/16-bit TIFF mode: stitch.py's int16
            # (or int8) canvases get silently widened to int32 on write.
            # Narrow back down to int16, the only signed MRC-representable
            # type such data can actually have come from.
            first_dtype = np.dtype(np.int16)
            mrc_mode = mrcfile.utils.mode_from_dtype(first_dtype)
        else:
            raise
    mrc_dtype = mrcfile.utils.dtype_from_mode(mrc_mode)
    with mrcfile.new_mmap(outputfile, shape, mrc_mode=mrc_mode, overwrite=True) as mrc:
        for i, f in enumerate(tqdm(fnams, desc="Joining TIFFs into MRC")):
            mrc.data[i] = np.array(Image.open(f), dtype=mrc_dtype)[: shape[1], : shape[2]]

    tilts = [extract_number(fnam) for fnam in fnams]

    # .rawtlt for IMOD: one tilt angle per line.
    tilt_file = os.path.join(outputdir, outname + ".rawtlt")
    with open(tilt_file, "w") as f:
        f.write("\n".join(str(t) for t in tilts) + "\n")

    # tiltAngles.txt for AreTomo: tilt_angle acquisition_index (1-based).
    tilt_angles_file = os.path.join(outputdir, "tiltAngles.txt")
    with open(tilt_angles_file, "w") as f:
        f.write("\n".join(f"{tilt} {i + 1}" for i, tilt in enumerate(tilts)) + "\n")

    psize = [args["pixel_size"]] * len(fnams)

    # Collect per-tilt HDF5 files containing refined tile positions and index maps,
    # pad positions arrays to a common shape, and save everything into one HDF5.
    h5fnams = sorted(
        glob.glob(os.path.join(inputdir, "Montage*refined_positions.h5")),
        key=extract_number,
    )
    h5outname = os.path.commonprefix(fnams) + ".h5"
    positions = [load_array_from_hdf5(fnam, "Refined_positions") for fnam in h5fnams]
    indexmap = np.asarray([load_array_from_hdf5(fnam, "index_map") for fnam in h5fnams])

    if positions:
        # Montages can differ in tile count across tilts; pad to the largest shape.
        max_shape = np.max([list(arr.shape) for arr in positions], axis=0).tolist()
        positions_padded = np.zeros((len(positions), *max_shape), dtype=positions[0].dtype)
        for i, arr in enumerate(positions):
            slices = tuple(slice(0, s) for s in arr.shape)
            positions_padded[(i, *slices)] = arr
        positions = positions_padded

    save_array_to_hdf5(
        [tilts, positions, indexmap, fnams, psize],
        h5outname,
        ["tilts", "Refined_positions", "index_map", "Montage_filenames", "binned_pixel_size"],
    )

    print(
        f"Written {len(fnams)} files to {outputfile}, "
        f"{outname}.rawtlt for IMOD, and {h5outname}"
    )


if __name__ == "__main__":
    main()
```
