## Supplementary material for "Whole cryo-FIB lamella acquisition at high magnification with rectangular beam montage cryo-electron tomography": Source code: mask_and_inpaint.py

```
#!/usr/bin/env python3
"""
Peak-following mask editor and inpainter for montage tilt series MRC files.

Workflow
--------
1. Click on the histogram peak you want to keep.
2. The script fits a Gaussian to that peak and propagates the fit across all
   tilts so that tilt-angle-dependent darkening is tracked automatically.
3. Optionally enforce a smooth µ curve (robust smoothn) to stabilise fits at
   high tilts where the peak drifts close to zero.
4. Optionally fall back to a manual range slider (like mask_and_inpaint.py)
   for full control.
5. Click "Generate Output" to inpaint every tilt with smoothn and save.
"""

import argparse
import json
import logging
import math
import os
import sys

import numpy as np
import mrcfile
import matplotlib.pyplot as plt
import matplotlib.widgets as mwidgets
from matplotlib.font_manager import FontProperties
from matplotlib.path import Path as MplPath
from mpl_toolkits.axes_grid1.anchored_artists import AnchoredSizeBar
from scipy.ndimage import label, zoom
from scipy.optimize import curve_fit
from tqdm import tqdm

from .smoothn import smoothn          # still used to smooth the µ/σ curves
from .Utilities import fourier_interpolate
from . import sidecar
from .sidecar import fingerprint
from .inpaint_apply import (
    DEFAULT_SLURM_TEMPLATE,
    SMOOTHN_S,
    SMOOTHN_MAX_ITER,
    inpaint_masked_tilt,
    series_base_for_input,
)

logger = logging.getLogger(__name__)

#: Name of this step in its sidecars. Applying the mask is a separate step
#: ("inpaint_apply"), usually a separate process on a cluster.
SIDECAR_STEP = "inpaint_mask"


# ---------------------------------------------------------------------------
# Binning
# ---------------------------------------------------------------------------

def bin_tilt_series(mmap_data, max_px: int = 1024, method: str = "zoom"):
    n, ny, nx = mmap_data.shape
    longest = max(ny, nx)
    if longest <= max_px:
        return np.stack([mmap_data[i].astype(np.float32) for i in range(n)]), 1.0
    factor = max_px / longest
    ny_out, nx_out = int(round(ny * factor)), int(round(nx * factor))
    print(
        f"Binning preview:  {ny}×{nx}  →  {ny_out}×{nx_out}  "
        f"(factor {factor:.3f}, {method})"
    )
    if method == "fourier":
        # Real Fourier-space decimation (crop high frequencies, then inverse
        # transform) rather than bilinear resampling — free of the aliasing
        # a plain zoom lets through, at the cost of ringing near sharp edges
        # (beam boundary, hot pixels).
        binned = np.stack([
            fourier_interpolate(mmap_data[i].astype(np.float32), (ny_out, nx_out))
            for i in tqdm(range(n), desc="Fourier-binning preview")
        ]).astype(np.float32)
    else:
        binned = np.stack([
            zoom(mmap_data[i].astype(np.float32), (factor, factor), order=1)
            for i in tqdm(range(n), desc="Binning preview")
        ])
    return binned, factor


def nice_scalebar_length(pixel_size_A: float, image_width_px: int) -> tuple:
    """
    Pick a round-number scalebar length near 18% of the image width.

    Returns (length_px, label), rounding to a 1/2/5 step in whichever of
    Å/nm/µm keeps the displayed number closest to human-friendly.
    """
    pixel_size_nm = pixel_size_A / 10.0
    target_nm = 0.18 * image_width_px * pixel_size_nm
    exponent = math.floor(math.log10(target_nm))
    nice_nm = target_nm
    for mult in (1, 2, 5, 10):
        candidate = mult * 10 ** exponent
        if candidate >= target_nm:
            nice_nm = candidate
            break
    if nice_nm >= 1000:
        label = f"{nice_nm / 1000:g} µm"
    elif nice_nm >= 1:
        label = f"{nice_nm:g} nm"
    else:
        label = f"{nice_nm * 10:g} Å"
    length_px = nice_nm / pixel_size_nm
    return length_px, label


# ---------------------------------------------------------------------------
# Gaussian peak fitting
# ---------------------------------------------------------------------------

def _gauss(x, amplitude, mu, sigma):
    return amplitude * np.exp(-0.5 * ((x - mu) / max(abs(sigma), 1e-9)) ** 2)


def fit_gaussian_peak(image: np.ndarray, center_hint: float,
                      search_window: float = None) -> tuple:
    """
    Fit a Gaussian to the histogram peak nearest to *center_hint*.

    Returns (mu, sigma).  search_window is the half-width of the search
    region; defaults to 30 % of the full intensity range.
    """
    flat = image.ravel().astype(float)
    val_range = float(flat.max() - flat[flat>0].min())
    if val_range == 0:
        return float(center_hint), 1.0

    if search_window is None:
        search_window = 0.30 * val_range

    nbins = 512
    counts, edges = np.histogram(flat[flat>0], bins=nbins)
    centers = 0.5 * (edges[:-1] + edges[1:])

    in_window = np.abs(centers - center_hint) < search_window
    if not in_window.any():
        in_window = np.ones(len(centers), dtype=bool)
    peak_bin = int(np.argmax(np.where(in_window, counts, 0)))
    peak_center = float(centers[peak_bin])

    fit_mask = np.abs(centers - peak_center) < search_window
    x = centers[fit_mask]
    y = counts[fit_mask].astype(float)

    if len(x) < 4:
        return peak_center, search_window / 4

    try:
        p0 = [float(y.max()), peak_center, search_window / 4]
        popt, _ = curve_fit(
            _gauss, x, y, p0=p0,
            bounds=([0.0, float(x.min()), val_range * 5e-4],
                    [np.inf, float(x.max()), search_window]),
            maxfev=4000,
        )
        return float(popt[1]), float(abs(popt[2]))
    except Exception:
        w = y / y.sum() if y.sum() > 0 else np.ones_like(y) / len(y)
        mu = float(np.dot(w, x))
        sigma = float(np.sqrt(np.dot(w, (x - mu) ** 2)))
        return mu, max(sigma, val_range * 0.01)


# ---------------------------------------------------------------------------
# Mask helpers
# ---------------------------------------------------------------------------

def largest_connected_component(binary_mask: np.ndarray) -> np.ndarray:
    labeled, n_features = label(binary_mask)
    if n_features == 0:
        return np.zeros_like(binary_mask, dtype=bool)
    sizes = np.bincount(labeled.ravel())
    sizes[0] = 0
    return (labeled == sizes.argmax()).astype(bool)


def compute_mask_from_range(image: np.ndarray, lo: float,
                             hi: float) -> np.ndarray:
    """True for pixels in [lo, hi] belonging to the largest connected component."""
    raw = (image >= lo) & (image <= hi)
    if not raw.any():
        return raw.astype(bool)
    return largest_connected_component(raw)


# ---------------------------------------------------------------------------
# Inpainting
# ---------------------------------------------------------------------------

def inpaint_tilt(image: np.ndarray, mask: np.ndarray,
                 bin_factor: int = 2, workers: int = 1) -> np.ndarray:
    """
    Inpaint pixels where *mask* is False.  mask: True = good pixel, False =
    inpaint.

    A thin wrapper on :func:`inpaint_masked_tilt`, which is what the SLURM
    runner uses, so what this window writes is what the cluster job writes.
    It used to be a separate implementation: smoothn's s chosen per tilt by
    GCV instead of the fixed 1e7, no binning, and a 15 px feather that
    replaced a rim of real data just inside the mask with the smooth field.
    None of that reached the cluster output, so the preview was a picture of
    a result nobody ever kept.
    """
    return inpaint_masked_tilt(image.astype(float), mask,
                               bin_factor=bin_factor, workers=workers)


# ---------------------------------------------------------------------------
# Interactive editor
# ---------------------------------------------------------------------------

class PeakMaskEditor:
    def __init__(self, mrc_mmap, input_path: str,
                 output_path: str = None, max_px: int = 1024,
                 preview_binning: str = "zoom",
                 n_sigma_default: float = 3.0,
                 slurm_template: str = None, slurm_cpus: int = 8,
                 slurm_npertilt: int = 4, slurm_bin: int = 2,
                 slurm_mem: int = 32, slurm_time: str = "02:00:00",
                 load_mask_path: str = None, write_sidecars: bool = True):
        self._mrc = mrc_mmap
        self._mmap_data = mrc_mmap.data
        if self._mmap_data.ndim == 2:
            self._mmap_data = self._mmap_data[np.newaxis]
        elif self._mmap_data.ndim != 3:
            raise ValueError(f"Expected 2D or 3D MRC data, got shape {self._mmap_data.shape}")
        self._full_shape = self._mmap_data.shape  # (n, h, w) — no data loaded yet
        # Copied, not referenced: the caller's mmap is closed when its `with`
        # block exits, and every MRC this class writes has to carry the input's
        # pixel size. A header saying 0 is read as 1 A/px downstream, which
        # rescales IMOD models by the ratio and throws every point off canvas.
        self._voxel_size = mrc_mmap.voxel_size.copy()
        self.data, self.bin_factor = bin_tilt_series(
            self._mmap_data, max_px, method=preview_binning
        )
        self.n_tilts   = self.data.shape[0]
        self.input_path  = input_path
        self.output_path = output_path or (
            os.path.splitext(input_path)[0] + "_inpainted.mrc"
        )
        self.current_tilt  = self.n_tilts // 2
        self.n_sigma_lo    = n_sigma_default
        self.n_sigma_hi    = n_sigma_default
        self.overlay_visible = True
        self.slurm_template = slurm_template
        self.slurm_cpus     = slurm_cpus       # ntotal: total thread budget for the job
        self.slurm_npertilt = slurm_npertilt   # DCT threads per tilt; ntotal // npertilt run concurrently
        self.slurm_bin      = slurm_bin        # smoothn fit resolution downsample factor
        self.slurm_mem      = slurm_mem
        self.slurm_time     = slurm_time

        # Sidecars go beside whatever this window writes, so unlike the
        # manifest there is nothing to locate up front and nothing to bake
        # into the params file for the cluster run to inherit — inpaint_apply
        # writes its own sidecar beside its own output.
        #
        # The failure this inverts is the silent one: the mask's provenance is
        # only in this window, in someone's head, and losing it silently is the
        # thing this exists to prevent. It is on unless explicitly disabled.
        self.write_sidecars = write_sidecars

        # --- Gaussian fit state ---
        self.fitted_mu    = np.full(self.n_tilts, np.nan)
        self.fitted_sigma = np.full(self.n_tilts, np.nan)
        self.smooth_mu    = None  # filled by _compute_smooth_mu
        self.smooth_sigma = None
        self._peak_selected = False
        self.use_smooth     = False  # toggled by button after fitting

        # --- Manual override state ---
        self.manual_mode     = False
        self.manual_overrides: dict = {}   # {tilt_idx: (lo, hi)}
        self._slider_updating = False      # guard against re-entrant slider updates

        # --- Paint overlay state ---
        self._paint_mode   = False
        self._painting     = False
        self._paint_dir    = "add"    # "add" or "remove" — set on press
        self._paint_add:    dict = {} # {tilt_idx: bool ndarray, binned coords}
        self._paint_remove: dict = {} # {tilt_idx: bool ndarray, binned coords}
        self._brush_size   = 10       # radius in display pixels

        # --- Top connected-component filter ---
        self._keep_top_cc = False   # re-apply LCC to final mask after paint overlay

        # --- Polygon drawing state ---
        self._poly_mode       = False
        self._poly_verts: list = []   # (x, y) in data coords
        self._poly_commit_dir = "remove" # "add" or "remove" — toggled by 'r' key

        # --- Sentinel detection (stitch --mark-uncovered writes -1 for no-tile regions) ---
        self._has_uncovered = bool((self.data < 0).any())
        if self._has_uncovered:
            n_unc = int((self.data < 0).sum())
            pct   = 100.0 * n_unc / self.data.size
            print(
                f"Detected {n_unc} uncovered pixels ({pct:.1f} %) marked with -1 "
                f"by stitch --mark-uncovered.  Setting initial lo=0 to exclude them."
            )
        # Compute intensity bounds from valid (non-sentinel) pixels only
        valid_px = self.data[self.data >= 0] if self._has_uncovered else self.data.ravel()
        self._global_min = -1.0 if self._has_uncovered else float(np.percentile(valid_px, 0.1))
        self._global_max = float(np.percentile(valid_px, 99.9))

        # --- Existing mask, loaded from a previous session ---
        # Kept mmap'd (full resolution) for generate/save; a binned copy is
        # precomputed for interactive display.  Acts as the base mask that
        # paint/polygon edits and the top-CC filter layer on top of, in place
        # of the "everything kept" default, whenever no Gaussian/manual
        # intensity threshold is active for a tilt.
        self._mask_mrc = None
        self.loaded_mask_preview = None
        if load_mask_path:
            self._mask_mrc = mrcfile.mmap(load_mask_path, mode="r", permissive=True)
            mask_data = self._mask_mrc.data
            if mask_data.ndim == 2:
                mask_data = mask_data[np.newaxis]
            if mask_data.shape != self._full_shape:
                self._mask_mrc.close()
                self._mask_mrc = None
                raise ValueError(
                    f"--load-mask shape {mask_data.shape} does not match "
                    f"input shape {self._full_shape}"
                )
            print(f"Loaded existing mask: {load_mask_path}  {mask_data.shape}")
            h_bin, w_bin = self.data.shape[1], self.data.shape[2]
            _, h_full, w_full = self._full_shape
            self.loaded_mask_preview = np.stack([
                zoom(mask_data[i].astype(np.float32), (h_bin / h_full, w_bin / w_full),
                     order=0) > 0.5
                for i in range(self.n_tilts)
            ])

        self._build_ui()
        self._refresh()

    # ------------------------------------------------------------------
    # Fitting
    # ------------------------------------------------------------------

    def _fit_all_tilts(self, ref_tilt: int, ref_mu: float, ref_sigma: float):
        """Propagate Gaussian fit from ref_tilt outward in both directions."""
        self.fitted_mu[ref_tilt]    = ref_mu
        self.fitted_sigma[ref_tilt] = ref_sigma

        mu, sigma = ref_mu, ref_sigma
        for i in range(ref_tilt + 1, self.n_tilts):
            img_i   = self.data[i]
            valid_i = img_i[img_i >= 0] if self._has_uncovered else img_i.ravel()
            rng     = float(valid_i.max() - valid_i.min()) if valid_i.size > 0 else float(img_i.max() - img_i.min())
            window  = max(4.0 * sigma, 0.05 * rng)
            mu, sigma = fit_gaussian_peak(img_i, mu, search_window=window)
            self.fitted_mu[i]    = mu
            self.fitted_sigma[i] = sigma

        mu, sigma = ref_mu, ref_sigma
        for i in range(ref_tilt - 1, -1, -1):
            img_i   = self.data[i]
            valid_i = img_i[img_i >= 0] if self._has_uncovered else img_i.ravel()
            rng     = float(valid_i.max() - valid_i.min()) if valid_i.size > 0 else float(img_i.max() - img_i.min())
            window  = max(4.0 * sigma, 0.05 * rng)
            mu, sigma = fit_gaussian_peak(img_i, mu, search_window=window)
            self.fitted_mu[i]    = mu
            self.fitted_sigma[i] = sigma

        self._peak_selected = True

    def _compute_smooth_mu(self):
        """
        Fit a smooth curve through the per-tilt µ and σ values using robust
        smoothn so that bad fits at high tilts (peak near zero) are down-weighted
        and the smooth curve follows the physically expected drift.
        """
        self.smooth_mu    = smoothn(self.fitted_mu.copy(),    robust=True)
        self.smooth_sigma = smoothn(self.fitted_sigma.copy(), robust=True)
        # Ensure sigma stays positive
        self.smooth_sigma = np.maximum(self.smooth_sigma, 1e-3)

    # ------------------------------------------------------------------
    # Threshold helper — single source of truth for (lo, hi, mu, sigma)
    # ------------------------------------------------------------------

    def _get_lo_hi(self, tilt_idx: int):
        """
        Return (lo, hi, mu, sigma) for the given tilt.

        Priority:  manual per-tilt override  >  smooth µ  >  raw Gaussian fit.
        Returns (None, None, None, None) if no peak has been selected and no
        override exists for this tilt.
        """
        if tilt_idx in self.manual_overrides:
            lo, hi = self.manual_overrides[tilt_idx]
            half   = (hi - lo) / 2.0
            avg_n_sigma = (self.n_sigma_lo + self.n_sigma_hi) / 2.0
            sigma  = half / max(avg_n_sigma, 1e-6)
            return lo, hi, lo + half, sigma

        if not self._peak_selected:
            return None, None, None, None

        if self.use_smooth and self.smooth_mu is not None:
            mu    = float(self.smooth_mu[tilt_idx])
            sigma = float(self.smooth_sigma[tilt_idx])
        else:
            mu    = float(self.fitted_mu[tilt_idx])
            sigma = float(self.fitted_sigma[tilt_idx])

        return max(mu - self.n_sigma_lo * sigma, 0), mu + self.n_sigma_hi * sigma, mu, sigma

    # ------------------------------------------------------------------
    # Mask helper — single source of truth for the auto/loaded base mask
    # ------------------------------------------------------------------

    def _base_mask(self, tilt_idx: int, img: np.ndarray) -> tuple:
        """
        Return (mask, has_threshold) — the Gaussian/manual intensity-threshold
        mask for this tilt, or everything-kept if no threshold is active.

        This is intentionally independent of any mask loaded via --load-mask:
        the loaded mask is combined in afterwards with logical_and (see
        _loaded_mask_at / _effective_mask_full), so it always constrains the
        final mask rather than being overridden by threshold/paint/polygon edits.
        """
        lo, hi, _, _ = self._get_lo_hi(tilt_idx)
        if lo is not None:
            return compute_mask_from_range(img, lo, hi), True
        return np.ones(img.shape, dtype=bool), False

    def _loaded_mask_at(self, tilt_idx: int, full_res: bool):
        """Return the --load-mask mask for this tilt at preview or full resolution, or None."""
        if full_res:
            return self._mask_mrc.data[tilt_idx].astype(bool) if self._mask_mrc is not None else None
        return self.loaded_mask_preview[tilt_idx] if self.loaded_mask_preview is not None else None

    def _effective_mask_full(self, tilt_idx: int, img_full: np.ndarray) -> np.ndarray:
        """
        Full-resolution mask for one tilt: threshold mask, then top-CC (so it
        only prunes the intensity-based selection, not manual edits), then
        paint/polygon overlay, then intersected (logical_and) with the loaded
        mask if any, so the loaded mask is always a constraint, never
        something later edits can override.
        """
        mask, _ = self._base_mask(tilt_idx, img_full)
        if self._keep_top_cc and mask.any():
            mask = largest_connected_component(mask)
        h_bin, w_bin   = self.data.shape[1], self.data.shape[2]
        h_full, w_full = img_full.shape
        p_add = self._paint_add.get(tilt_idx)
        p_rem = self._paint_remove.get(tilt_idx)
        if p_add is not None:
            mask = mask | (zoom(p_add.astype(np.float32),
                                 (h_full / h_bin, w_full / w_bin), order=0) > 0.5)
        if p_rem is not None:
            mask = mask & ~(zoom(p_rem.astype(np.float32),
                                  (h_full / h_bin, w_full / w_bin), order=0) > 0.5)
        loaded = self._loaded_mask_at(tilt_idx, full_res=True)
        if loaded is not None:
            mask = np.logical_and(mask, loaded)
        return mask

    # ------------------------------------------------------------------
    # UI construction
    # ------------------------------------------------------------------

    def _build_ui(self):
        self.fig = plt.figure(figsize=(14, 9))
        full_shape    = self._full_shape
        preview_shape = self.data.shape
        bin_note = (
            f"  —  preview {preview_shape[1]}×{preview_shape[2]}"
            f" (full {full_shape[1]}×{full_shape[2]})"
            if self.bin_factor < 1.0 else ""
        )
        self.fig.suptitle(
            f"Mask & Inpaint (peak):  {os.path.basename(self.input_path)}"
            f"  ({self.n_tilts} tilts){bin_note}",
            fontsize=10,
        )

        # ---- image and histogram panels ----
        # ax_hist and ax_mu need a wider vertical gap than a bare y-delta
        # suggests: ax_hist's xlabel sits below it and ax_mu's title sits
        # above it, both competing for the same strip of figure space.
        # The img/hist horizontal gap is widened too: with add_axes (as
        # opposed to subplots), tick labels and the y-axis label are drawn
        # outside the given rectangle, so ax_hist's own "Pixel count" label
        # and y-tick numbers need clearance from ax_img's right edge.
        self.ax_img  = self.fig.add_axes([0.04, 0.38, 0.48, 0.55])
        self.ax_hist = self.fig.add_axes([0.60, 0.38, 0.37, 0.55])
        self.ax_mu   = self.fig.add_axes([0.60, 0.19, 0.37, 0.07])

        # ---- sliders ----
        ax_tilt   = self.fig.add_axes([0.04, 0.21, 0.42, 0.03])
        ax_prev   = self.fig.add_axes([0.47, 0.195, 0.03, 0.05])
        ax_next   = self.fig.add_axes([0.51, 0.195, 0.03, 0.05])
        ax_nsigma = self.fig.add_axes([0.04, 0.15, 0.50, 0.03])
        ax_manual = self.fig.add_axes([0.04, 0.09, 0.50, 0.03])

        # ---- buttons (bottom row) ----
        ax_btn_smooth   = self.fig.add_axes([0.04, 0.02, 0.06, 0.06])
        ax_btn_manual   = self.fig.add_axes([0.11, 0.02, 0.06, 0.06])
        ax_btn_overlay  = self.fig.add_axes([0.18, 0.02, 0.06, 0.06])
        ax_btn_paint    = self.fig.add_axes([0.25, 0.02, 0.06, 0.06])
        ax_btn_polygon  = self.fig.add_axes([0.32, 0.02, 0.06, 0.06])
        ax_btn_top_cc   = self.fig.add_axes([0.39, 0.02, 0.06, 0.06])
        ax_btn_slurm    = self.fig.add_axes([0.46, 0.02, 0.10, 0.06])
        ax_btn_gen     = self.fig.add_axes([0.57, 0.02, 0.17, 0.08])

        # ---- brush slider + reset/save controls (right column, between mu-plot and buttons) ----
        ax_brush            = self.fig.add_axes([0.60, 0.11, 0.37, 0.03])
        ax_btn_reset_paint  = self.fig.add_axes([0.76, 0.06, 0.21, 0.04])
        ax_chk_save_mask    = self.fig.add_axes([0.76, 0.02, 0.21, 0.04])

        # Tilt slider
        # Labels are set via ax.set_title(loc="left") rather than the Slider's
        # own `label=` argument: matplotlib draws that label to the *left* of
        # the axes, which clips off the figure for sliders anchored near the
        # left edge (x0=0.04).
        self.sl_tilt = mwidgets.Slider(
            ax_tilt, "", 0, max(self.n_tilts - 1, 1),
            valinit=self.current_tilt, valstep=1, color="steelblue",
        )
        ax_tilt.set_title("Tilt index", loc="left", fontsize=9, pad=2)
        self.sl_tilt.on_changed(self._on_tilt)

        self.btn_prev = mwidgets.Button(ax_prev, "◄", color="lightgray", hovercolor="silver")
        self.btn_next = mwidgets.Button(ax_next, "►", color="lightgray", hovercolor="silver")
        self.btn_prev.on_clicked(lambda _: self._step_tilt(-1))
        self.btn_next.on_clicked(lambda _: self._step_tilt(+1))

        # N-sigma range slider (Gaussian mode) — lower/upper multipliers, applied
        # to every tilt without a manual override
        self.sl_nsigma = mwidgets.RangeSlider(
            ax_nsigma, "", 0.5, 6.0,
            valinit=[self.n_sigma_lo, self.n_sigma_hi], color="mediumpurple",
        )
        ax_nsigma.set_title("N sigma", loc="left", fontsize=9, pad=2)
        self.sl_nsigma.on_changed(self._on_nsigma)

        # Manual range slider — lower bound starts at 0 when sentinel pixels present
        _manual_lo_init = max(0.0, self._global_min) if self._has_uncovered else self._global_min
        self.sl_manual = mwidgets.RangeSlider(
            ax_manual, "",
            self._global_min, self._global_max,
            valinit=[_manual_lo_init, self._global_max],
        )
        ax_manual.set_title("Manual range", loc="left", fontsize=9, pad=2)
        self.sl_manual.on_changed(self._on_manual_range)
        ax_manual.set_alpha(0.3)  # visually dim until manual mode is active

        # Smooth µ button
        self.btn_smooth = mwidgets.Button(
            ax_btn_smooth, "Smooth µ:\nOFF",
            color="lightcyan", hovercolor="cyan",
        )
        self.btn_smooth.on_clicked(self._on_toggle_smooth)

        # Manual mode button
        self.btn_manual = mwidgets.Button(
            ax_btn_manual, "Manual:\nOFF",
            color="lightyellow", hovercolor="yellow",
        )
        self.btn_manual.on_clicked(self._on_toggle_manual)

        # Overlay toggle
        self.btn_overlay = mwidgets.Button(
            ax_btn_overlay, "Overlay:\nON",
            color="lavender", hovercolor="mediumpurple",
        )
        self.btn_overlay.on_clicked(self._on_toggle_overlay)

        # Save SLURM job button
        self.btn_slurm = mwidgets.Button(
            ax_btn_slurm, "Save SLURM\nJob",
            color="lightskyblue", hovercolor="deepskyblue",
        )
        self.btn_slurm.on_clicked(self._on_save_slurm)

        # Generate button
        self.btn_gen = mwidgets.Button(
            ax_btn_gen, "Generate\nOutput",
            color="palegreen", hovercolor="lime",
        )
        self.btn_gen.on_clicked(self._on_generate)

        # Paint mode button
        self.btn_paint = mwidgets.Button(
            ax_btn_paint, "Paint:\nOFF",
            color="lightgray", hovercolor="silver",
        )
        self.btn_paint.on_clicked(self._on_toggle_paint)

        # Polygon mode button
        self.btn_polygon = mwidgets.Button(
            ax_btn_polygon, self._poly_btn_label(),
            color="lightgray", hovercolor="silver",
        )
        self.btn_polygon.on_clicked(self._on_toggle_polygon)

        # Top connected-component filter button
        self.btn_top_cc = mwidgets.Button(
            ax_btn_top_cc, "Top CC:\nOFF",
            color="lightgray", hovercolor="silver",
        )
        self.btn_top_cc.on_clicked(self._on_toggle_top_cc)

        # Brush radius slider
        self.sl_brush = mwidgets.Slider(
            ax_brush, "", 1, 80,
            valinit=self._brush_size, valstep=1, valfmt="%0.0f",
            color="sandybrown",
        )
        ax_brush.set_title("Brush (px)", loc="left", fontsize=9, pad=2)
        self.sl_brush.on_changed(self._on_brush_changed)

        # Reset paint button
        self.btn_reset_paint = mwidgets.Button(
            ax_btn_reset_paint, "Reset paint",
            color="lightsalmon", hovercolor="tomato",
        )
        self.btn_reset_paint.on_clicked(self._on_reset_paint)

        # Save mask checkbox
        self.chk_save_mask = mwidgets.CheckButtons(
            ax_chk_save_mask, ["Save mask MRC"], [True]
        )

        # ---- image display ----
        img0 = self.data[0]
        self._im = self.ax_img.imshow(
            img0, cmap="gray", origin="lower",
            vmin=float(np.percentile(img0, 1)),
            vmax=float(np.percentile(img0, 99)),
        )
        self._overlay_im = self.ax_img.imshow(
            np.zeros((*img0.shape, 4), dtype=np.float32),
            origin="lower", aspect="auto", interpolation="nearest",
        )
        self.ax_img.axis("off")

        # Scalebar — display pixel size accounts for the preview binning
        # (bin_tilt_series shrinks the array by self.bin_factor, so each
        # displayed pixel covers 1/bin_factor full-res pixels).
        pixel_size_A = float(self._voxel_size.x)
        if pixel_size_A > 0:
            display_pixel_size_A = pixel_size_A / self.bin_factor
            length_px, label = nice_scalebar_length(
                display_pixel_size_A, img0.shape[1]
            )
            self.ax_img.add_artist(AnchoredSizeBar(
                self.ax_img.transData, length_px, label, loc="lower right",
                pad=0.6, color="yellow", frameon=False, size_vertical=max(img0.shape[0] / 200, 1),
                fontproperties=FontProperties(size=10),
            ))

        self.ax_hist.set_title("Click on the peak to select it", fontsize=9)
        self.ax_mu.set_title("µ across tilts", fontsize=8)
        self.ax_mu.set_xlabel("Tilt index", fontsize=7)
        self.ax_mu.tick_params(labelsize=7)

        # Brush cursor circle drawn on the image axes
        self._brush_cursor = plt.Circle(
            (0, 0), self._brush_size,
            fill=False, color="yellow", linestyle="--", linewidth=1.2,
            visible=False,
        )
        self.ax_img.add_patch(self._brush_cursor)

        # Polygon drawing artists (always present, hidden until used)
        self._poly_line,       = self.ax_img.plot([], [], color="cyan", lw=1.5, ls="--")
        self._poly_close_line, = self.ax_img.plot([], [], color="cyan", lw=1.0, ls=":")
        self._poly_dots,       = self.ax_img.plot([], [], "o", color="cyan", ms=4)

        self.fig.canvas.mpl_connect("button_press_event",   self._on_mouse_press)
        self.fig.canvas.mpl_connect("button_release_event", self._on_mouse_release)
        self.fig.canvas.mpl_connect("motion_notify_event",  self._on_mouse_motion)
        self.fig.canvas.mpl_connect("key_press_event",      self._on_key_press)

    # ------------------------------------------------------------------
    # Display refresh
    # ------------------------------------------------------------------

    def _refresh(self):
        img = self.data[self.current_tilt]
        lo, hi, mu, sigma = self._get_lo_hi(self.current_tilt)

        has_threshold = lo is not None
        ti = self.current_tilt
        auto_mask, _ = self._base_mask(ti, img)
        loaded = self._loaded_mask_at(ti, full_res=False)

        # Apply paint overlay on top of the auto-computed mask
        p_add = self._paint_add.get(ti)
        p_rem = self._paint_remove.get(ti)
        active = has_threshold or loaded is not None or p_add is not None or p_rem is not None
        mask = auto_mask.copy()
        if self._keep_top_cc and mask.any():
            mask = largest_connected_component(mask)
        if p_add is not None:
            mask |= p_add
        if p_rem is not None:
            mask &= ~p_rem
        # The loaded mask is a hard constraint: intersect (never override) so that
        # no threshold/paint/polygon edit can re-include pixels it excludes.
        if loaded is not None:
            mask = np.logical_and(mask, loaded)
        bad = ~mask

        # -- image + overlay --
        # Contrast is stretched over the threshold/loaded-mask selection only, not
        # the paint/polygon overlay, so manually adding back outlier pixels
        # (fiducials, hot pixels, etc.) doesn't blow out the display range on every
        # edit.  Also exclude sentinel pixels so -1 doesn't crush the range.
        clim_mask = np.logical_and(auto_mask, loaded) if loaded is not None else auto_mask
        valid_for_clim = img[clim_mask & (img >= 0)]
        if valid_for_clim.size == 0:
            valid_for_clim = img[img >= 0] if (img >= 0).any() else img.ravel()
        self._im.set_data(img)
        self._im.set_clim(float(np.percentile(valid_for_clim, 1)),
                          float(np.percentile(valid_for_clim, 99)))

        rgba = np.zeros((*img.shape, 4), dtype=np.float32)
        if self.overlay_visible and active:
            # Pixels painted "add" but still excluded (blocked by the loaded-mask
            # constraint) count as bad, not rescued, so they show red like any
            # other excluded pixel rather than a misleading cyan.
            rescued = (p_add & mask) if p_add is not None else np.zeros(img.shape, bool)
            auto_bad_only = bad & ~rescued
            rgba[auto_bad_only] = [1.0, 0.2, 0.2, 0.5]
            # Paint-remove strokes: orange
            if p_rem is not None:
                rgba[p_rem] = [1.0, 0.5, 0.0, 0.75]
            # Paint-add strokes that actually took effect: cyan
            if p_add is not None:
                rgba[p_add & mask] = [0.0, 0.8, 1.0, 0.45]
        self._overlay_im.set_data(rgba)
        self._overlay_im.set_extent(
            [-0.5, img.shape[1] - 0.5, -0.5, img.shape[0] - 0.5]
        )

        n_bad = int(bad.sum())
        pct   = 100.0 * n_bad / bad.size
        if has_threshold:
            mode_tag = (
                "manual override" if self.current_tilt in self.manual_overrides
                else ("smooth µ" if self.use_smooth else "Gaussian")
            )
            if loaded is not None:
                mode_tag += " ∩ loaded mask"
            title = (
                f"Tilt {self.current_tilt + 1}/{self.n_tilts}  [{mode_tag}]  "
                f"µ={mu:.1f}  σ={sigma:.1f}  "
                f"inpaint: {n_bad} px ({pct:.1f} %)"
            )
        elif active:
            title = (
                f"Tilt {self.current_tilt + 1}/{self.n_tilts}  [loaded mask]  "
                f"inpaint: {n_bad} px ({pct:.1f} %)"
            )
        else:
            title = (f"Tilt {self.current_tilt + 1}/{self.n_tilts}  "
                     "—  click histogram to select peak")
        self.ax_img.set_title(title, fontsize=9)

        # -- histogram --
        self.ax_hist.clear()
        flat  = img.ravel()
        # When uncovered-sentinel pixels (-1) are present, show them as a separate
        # bar to the left; compute the main histogram range from valid pixels only.
        uncov_mask = flat < 0
        n_uncov = int(uncov_mask.sum())
        valid_flat = flat[~uncov_mask] if n_uncov > 0 else flat
        h_lo  = float(np.percentile(valid_flat, 0.2)) if valid_flat.size else 0.0
        h_hi  = float(np.percentile(valid_flat, 99.8)) if valid_flat.size else 1.0
        counts, bins, patches = self.ax_hist.hist(
            valid_flat, bins=256, range=(h_lo, h_hi), color="steelblue", alpha=0.8
        )
        if n_uncov > 0:
            self.ax_hist.axvspan(h_lo - (h_hi - h_lo) * 0.05, h_lo,
                                 color="firebrick", alpha=0.4)
            self.ax_hist.text(
                h_lo - (h_hi - h_lo) * 0.02, counts.max() * 0.5,
                f"uncov.\n{n_uncov}", color="firebrick", fontsize=7,
                ha="right", va="center",
            )
        bin_centers = 0.5 * (bins[:-1] + bins[1:])

        if has_threshold:
            for patch, left, right in zip(patches, bins[:-1], bins[1:]):
                if right <= lo or left >= hi:
                    patch.set_facecolor("firebrick")
                    patch.set_alpha(0.85)

            if not self.manual_mode:
                amp = float(counts[np.argmin(np.abs(bin_centers - mu))])
                x_fine = np.linspace(h_lo, h_hi, 500)
                self.ax_hist.plot(
                    x_fine, _gauss(x_fine, amp, mu, sigma),
                    color="limegreen", lw=1.5,
                    label=f"fit  µ={mu:.1f}  σ={sigma:.1f}",
                )
                # If smooth mode is active, also draw raw Gaussian faintly
                if self.use_smooth:
                    raw_mu    = float(self.fitted_mu[self.current_tilt])
                    raw_sigma = float(self.fitted_sigma[self.current_tilt])
                    raw_amp   = float(counts[np.argmin(np.abs(bin_centers - raw_mu))])
                    self.ax_hist.plot(
                        x_fine, _gauss(x_fine, raw_amp, raw_mu, raw_sigma),
                        color="limegreen", lw=0.8, ls=":", alpha=0.5,
                        label=f"raw  µ={raw_mu:.1f}",
                    )

            self.ax_hist.axvline(lo, color="red",    lw=1.5, ls="--",
                                  label=f"lo={lo:.1f}")
            self.ax_hist.axvline(hi, color="orange", lw=1.5, ls="--",
                                  label=f"hi={hi:.1f}")
            self.ax_hist.legend(fontsize=7, loc="upper right")
            self.ax_hist.set_title(
                "Histogram  (red = inpainted,  green = Gaussian fit)", fontsize=9
            )
        elif active:
            self.ax_hist.set_title(
                "Histogram  (loaded mask active — click a peak to also "
                "apply an intensity threshold)", fontsize=9
            )
        else:
            self.ax_hist.set_title("Click on the peak you want to keep",
                                    fontsize=9)

        self.ax_hist.set_xlabel("Intensity")
        self.ax_hist.set_ylabel("Pixel count")

        # -- µ-vs-tilt summary --
        self._redraw_mu_plot()

        self.fig.canvas.draw_idle()

    def _redraw_mu_plot(self):
        ax = self.ax_mu
        ax.clear()
        valid = ~np.isnan(self.fitted_mu)
        if valid.any():
            idx = np.where(valid)[0]
            ax.plot(idx, self.fitted_mu[valid],
                    "o", ms=2, color="steelblue", alpha=0.6, label="raw µ")
            ax.fill_between(
                idx,
                self.fitted_mu[valid] - self.fitted_sigma[valid],
                self.fitted_mu[valid] + self.fitted_sigma[valid],
                alpha=0.15, color="steelblue",
            )
            if self.smooth_mu is not None:
                ax.plot(np.arange(self.n_tilts), self.smooth_mu,
                        color="darkorange", lw=1.5, label="smooth µ")
                ax.fill_between(
                    np.arange(self.n_tilts),
                    self.smooth_mu - self.smooth_sigma,
                    self.smooth_mu + self.smooth_sigma,
                    alpha=0.20, color="darkorange",
                )
        if self.manual_overrides:
            ov_idx = sorted(self.manual_overrides)
            ov_mid = [(self.manual_overrides[i][0] + self.manual_overrides[i][1]) / 2
                      for i in ov_idx]
            ax.scatter(ov_idx, ov_mid, color="red", s=18, zorder=5,
                       label=f"manual ({len(ov_idx)} tilt{'s' if len(ov_idx) != 1 else ''})")
            for i in ov_idx:
                lo_i, hi_i = self.manual_overrides[i]
                ax.plot([i, i], [lo_i, hi_i], color="red", lw=1.5, alpha=0.55)
        ax.axvline(self.current_tilt, color="red", lw=1, ls="--")
        ax.set_title("µ across tilts", fontsize=8)
        ax.set_xlabel("Tilt index", fontsize=7)
        ax.tick_params(labelsize=7)
        if valid.any() or self.manual_mode:
            ax.legend(fontsize=7, loc="upper right")

    # ------------------------------------------------------------------
    # Widget callbacks
    # ------------------------------------------------------------------

    def _update_slider_for_tilt(self, tilt_idx: int):
        """Sync the manual range slider to tilt_idx's current effective lo/hi."""
        lo, hi, _, _ = self._get_lo_hi(tilt_idx)
        if lo is None:
            lo, hi = self._global_min, self._global_max
        self._slider_updating = True
        self.sl_manual.set_val([lo, hi])
        self._slider_updating = False

    def _step_tilt(self, delta: int):
        new_tilt = int(np.clip(self.current_tilt + delta, 0, self.n_tilts - 1))
        if new_tilt != self.current_tilt:
            self.sl_tilt.set_val(new_tilt)  # triggers _on_tilt → _refresh

    def _on_tilt(self, val):
        self.current_tilt = int(round(val))
        if self._poly_verts:
            self._clear_polygon_drawing()
        if self.manual_mode:
            self._update_slider_for_tilt(self.current_tilt)
        self._refresh()

    def _on_nsigma(self, val):
        self.n_sigma_lo, self.n_sigma_hi = float(val[0]), float(val[1])
        self._refresh()

    def _on_manual_range(self, val):
        if not self.manual_mode or self._slider_updating:
            return
        self.manual_overrides[self.current_tilt] = (float(val[0]), float(val[1]))
        self._refresh()

    def _on_toggle_smooth(self, _event):
        if not self._peak_selected:
            print("Select a peak first (click the histogram).")
            return
        self.use_smooth = not self.use_smooth
        if self.use_smooth and self.smooth_mu is None:
            print("Computing smooth µ curve …")
            self._compute_smooth_mu()
            print(f"  smooth µ range: {self.smooth_mu.min():.1f} – "
                  f"{self.smooth_mu.max():.1f}")
        self.btn_smooth.label.set_text(
            "Smooth µ:\nON" if self.use_smooth else "Smooth µ:\nOFF"
        )
        self.btn_smooth.ax.set_facecolor(
            "cyan" if self.use_smooth else "lightcyan"
        )
        self._refresh()

    def _on_toggle_manual(self, _event):
        self.manual_mode = not self.manual_mode
        if self.manual_mode:
            # Seed slider from this tilt's current effective range so the user
            # starts from a sensible position rather than the global extremes
            self._update_slider_for_tilt(self.current_tilt)
        self.btn_manual.label.set_text(
            "Manual:\nON" if self.manual_mode else "Manual:\nOFF"
        )
        self.btn_manual.ax.set_facecolor(
            "yellow" if self.manual_mode else "lightyellow"
        )
        self.sl_manual.ax.set_alpha(1.0 if self.manual_mode else 0.3)
        self._refresh()

    def _on_toggle_overlay(self, _event):
        self.overlay_visible = not self.overlay_visible
        self.btn_overlay.label.set_text(
            "Overlay:\nON" if self.overlay_visible else "Overlay:\nOFF"
        )
        self._refresh()

    def _on_toggle_top_cc(self, _event):
        self._keep_top_cc = not self._keep_top_cc
        if self._keep_top_cc:
            self.btn_top_cc.label.set_text("Top CC:\nON")
            self.btn_top_cc.ax.set_facecolor("lightgreen")
        else:
            self.btn_top_cc.label.set_text("Top CC:\nOFF")
            self.btn_top_cc.ax.set_facecolor("lightgray")
        self._refresh()

    def _on_mouse_press(self, event):
        # Polygon drawing on image panel
        if self._poly_mode and event.inaxes is self.ax_img and event.xdata is not None:
            if event.button == 3:
                # Right-click: commit as REMOVE (bonus shortcut where it works)
                if len(self._poly_verts) >= 3:
                    self._finalize_polygon("remove")
                    self._clear_polygon_drawing()
                    self._refresh()
                else:
                    self._clear_polygon_drawing()
                return
            # Left-click: place vertex, or close and commit if near start or double-click
            if event.dblclick or self._is_near_poly_start(event.xdata, event.ydata):
                if len(self._poly_verts) >= 3:
                    self._finalize_polygon(self._poly_commit_dir)
                    self._refresh()
                self._clear_polygon_drawing()
            else:
                self._poly_verts.append((event.xdata, event.ydata))
                self._update_poly_display(cursor_xy=None)
            return

        # Paint on image panel
        if self._paint_mode and event.inaxes is self.ax_img and event.xdata is not None:
            self._painting = True
            self._paint_dir = "add" if event.button == 1 else "remove"
            self._paint_at(event.xdata, event.ydata)
            return

        # Histogram peak selection (existing behaviour)
        if event.inaxes is not self.ax_hist or event.xdata is None:
            return
        if self.manual_mode:
            return
        x_click = float(event.xdata)
        img = self.data[self.current_tilt]
        print(f"Fitting peak near x={x_click:.2f}  (tilt {self.current_tilt + 1}) …")
        mu, sigma = fit_gaussian_peak(img, x_click)
        print(f"  Fitted:  µ={mu:.2f}  σ={sigma:.2f}")
        print(f"Propagating fit across all {self.n_tilts} tilts …")
        self._fit_all_tilts(self.current_tilt, mu, sigma)
        self.smooth_mu    = None
        self.smooth_sigma = None
        if self.use_smooth:
            print("Recomputing smooth µ curve …")
            self._compute_smooth_mu()
        print("Done.")
        self._refresh()

    def _on_mouse_release(self, _event):
        self._painting = False

    def _on_mouse_motion(self, event):
        # Polygon mode: update the rubber-band close-line following the cursor
        if self._poly_mode and self._poly_verts:
            in_img = event.inaxes is self.ax_img and event.xdata is not None
            if in_img:
                self._update_poly_display(cursor_xy=(event.xdata, event.ydata))
            self.fig.canvas.draw_idle()
            return

        if not self._paint_mode:
            return
        in_img = event.inaxes is self.ax_img and event.xdata is not None
        self._brush_cursor.set_visible(in_img)
        if in_img:
            self._brush_cursor.set_center((event.xdata, event.ydata))
            self._brush_cursor.set_radius(self._brush_size)
            if self._painting:
                self._paint_at(event.xdata, event.ydata)
        self.fig.canvas.draw_idle()

    def _paint_at(self, xdata: float, ydata: float):
        """Apply one brush stamp at (xdata, ydata) in image data coordinates."""
        h, w = self.data.shape[1], self.data.shape[2]
        ti   = self.current_tilt
        r    = self._brush_size
        cx, cy = int(round(xdata)), int(round(ydata))
        ys, xs = np.ogrid[:h, :w]
        circle = (ys - cy) ** 2 + (xs - cx) ** 2 <= r ** 2
        if self._paint_dir == "add":
            if ti not in self._paint_add:
                self._paint_add[ti] = np.zeros((h, w), dtype=bool)
            self._paint_add[ti][circle] = True
            if ti in self._paint_remove:
                self._paint_remove[ti][circle] = False
        else:
            if ti not in self._paint_remove:
                self._paint_remove[ti] = np.zeros((h, w), dtype=bool)
            self._paint_remove[ti][circle] = True
            if ti in self._paint_add:
                self._paint_add[ti][circle] = False
        self._refresh()

    def _on_brush_changed(self, val):
        self._brush_size = int(val)
        self._brush_cursor.set_radius(self._brush_size)
        self.fig.canvas.draw_idle()

    def _on_reset_paint(self, _event):
        ti = self.current_tilt
        self._paint_add.pop(ti, None)
        self._paint_remove.pop(ti, None)
        self._refresh()

    # ------------------------------------------------------------------
    # Polygon drawing
    # ------------------------------------------------------------------

    def _poly_btn_label(self):
        return f"Poly: {'ON' if self._poly_mode else 'OFF'}\n[{self._poly_commit_dir.upper()}]"

    def _on_toggle_polygon(self, _event):
        self._poly_mode = not self._poly_mode
        if self._poly_mode:
            # Deactivate paint mode so they don't interfere
            if self._paint_mode:
                self._paint_mode = False
                self.btn_paint.label.set_text("Paint:\nOFF")
                self.btn_paint.ax.set_facecolor("lightgray")
                self._brush_cursor.set_visible(False)
            self.btn_polygon.ax.set_facecolor("lightcyan")
            print(
                "Polygon mode ON  —  left-click: place vertex  |  "
                "click near start or Enter: commit  |  "
                "c: close via nearest edge points (min-area) then commit  |  "
                "r: toggle add/remove  |  "
                "Backspace: undo last vertex  |  "
                "Esc: cancel"
            )
        else:
            self._clear_polygon_drawing()
            self.btn_polygon.ax.set_facecolor("lightgray")
        self.btn_polygon.label.set_text(self._poly_btn_label())
        self.fig.canvas.draw_idle()

    def _on_toggle_paint(self, _event):
        self._paint_mode = not self._paint_mode
        if self._paint_mode:
            # Deactivate polygon mode so they don't interfere
            if self._poly_mode:
                self._poly_mode = False
                self._clear_polygon_drawing()
                self.btn_polygon.label.set_text("Poly:\nOFF")
                self.btn_polygon.ax.set_facecolor("lightgray")
            self.btn_paint.label.set_text("Paint:\nON")
            self.btn_paint.ax.set_facecolor("lightgreen")
        else:
            self.btn_paint.label.set_text("Paint:\nOFF")
            self.btn_paint.ax.set_facecolor("lightgray")
            self._brush_cursor.set_visible(False)
        self.fig.canvas.draw_idle()

    def _is_near_poly_start(self, xdata: float, ydata: float) -> bool:
        """True if (xdata, ydata) is within ~10 display pixels of the first vertex."""
        if len(self._poly_verts) < 2:
            return False
        x0, y0 = self._poly_verts[0]
        disp_click = self.ax_img.transData.transform((xdata, ydata))
        disp_start = self.ax_img.transData.transform((x0, y0))
        return float(np.hypot(*(disp_click - disp_start))) < 10.0

    def _update_poly_display(self, cursor_xy=None):
        """Redraw the polygon outline artists from current vertices."""
        if not self._poly_verts:
            self._poly_line.set_data([], [])
            self._poly_close_line.set_data([], [])
            self._poly_dots.set_data([], [])
            return
        xs = [v[0] for v in self._poly_verts]
        ys = [v[1] for v in self._poly_verts]
        # Main line connecting placed vertices
        self._poly_line.set_data(xs, ys)
        self._poly_dots.set_data(xs, ys)
        # Rubber-band line: from last vertex to cursor (and dotted back to start)
        if cursor_xy is not None:
            cx, cy = cursor_xy
            self._poly_close_line.set_data(
                [xs[-1], cx, xs[0]], [ys[-1], cy, ys[0]]
            )
        else:
            self._poly_close_line.set_data([], [])

    def _finalize_polygon(self, direction: str):
        """Commit the current polygon interior to the paint overlay."""
        if len(self._poly_verts) < 3:
            return
        h, w = self.data.shape[1], self.data.shape[2]
        verts = self._poly_verts + [self._poly_verts[0]]  # close the path
        path = MplPath(verts)
        yy, xx = np.mgrid[:h, :w]
        pts = np.column_stack([xx.ravel(), yy.ravel()])
        inside = path.contains_points(pts).reshape(h, w)
        ti = self.current_tilt
        if direction == "add":
            if ti not in self._paint_add:
                self._paint_add[ti] = np.zeros((h, w), dtype=bool)
            self._paint_add[ti] |= inside
            if ti in self._paint_remove:
                self._paint_remove[ti] &= ~inside
        else:  # "remove"
            if ti not in self._paint_remove:
                self._paint_remove[ti] = np.zeros((h, w), dtype=bool)
            self._paint_remove[ti] |= inside
            if ti in self._paint_add:
                self._paint_add[ti] &= ~inside

    def _clear_polygon_drawing(self):
        """Discard in-progress polygon vertices and hide artists."""
        self._poly_verts = []
        self._poly_line.set_data([], [])
        self._poly_close_line.set_data([], [])
        self._poly_dots.set_data([], [])

    def _close_via_edge(self):
        """
        Snap each open endpoint perpendicularly to the nearest image boundary
        point (anywhere on the edge, not just corners), then walk the boundary
        between those two snap points in the direction that minimises the
        enclosed polygon area (shoelace), then finalise.
        """
        if not self._poly_verts:
            return
        h, w  = self.data.shape[1], self.data.shape[2]
        W, H  = float(w - 1), float(h - 1)
        P     = 2.0 * (W + H)   # perimeter

        # Clockwise corners: BL, BR, TR, TL
        corners  = [(0.0, 0.0), (W, 0.0), (W, H), (0.0, H)]
        corner_s = [0.0, W, W + H, 2.0*W + H]

        def snap(x, y):
            """Project (x, y) onto the nearest image boundary edge."""
            x, y = float(np.clip(x, 0, W)), float(np.clip(y, 0, H))
            d_l, d_r, d_b, d_t = x, W - x, y, H - y
            best = min(d_l, d_r, d_b, d_t)
            if best == d_l: return (0.0, y)
            if best == d_r: return (W,   y)
            if best == d_b: return (x,   0.0)
            return                  (x,   H)

        def to_s(px, py):
            """Clockwise boundary parameter s ∈ [0, P) for a point on the boundary."""
            if py == 0.0: return px                    # bottom  (left → right)
            if px == W:   return W + py                # right   (bottom → top)
            if py == H:   return W + H + (W - px)     # top     (right → left)
            return                W + H + W + (H - py) # left    (top → bottom)

        def arc(s_from, pt_from, s_to, pt_to, cw: bool):
            """Boundary vertices from pt_from to pt_to via CW (cw=True) or CCW arc."""
            if cw:
                span     = (s_to - s_from) % P
                delta_fn = lambda cs: (cs - s_from) % P
            else:
                span     = (s_from - s_to) % P
                delta_fn = lambda cs: (s_from - cs) % P
            if span == 0:
                return [pt_from, pt_to]
            mid = sorted(
                (delta_fn(cs), corners[i])
                for i, cs in enumerate(corner_s)
                if 0 < delta_fn(cs) < span
            )
            return [pt_from] + [pt for _, pt in mid] + [pt_to]

        def shoelace(verts):
            n  = len(verts)
            xs = [v[0] for v in verts]
            ys = [v[1] for v in verts]
            return abs(sum(xs[i]*ys[(i+1)%n] - xs[(i+1)%n]*ys[i]
                           for i in range(n))) / 2.0

        user     = list(self._poly_verts)
        pt_last  = snap(*user[-1])
        pt_first = snap(*user[0])
        s_last   = to_s(*pt_last)
        s_first  = to_s(*pt_first)

        path_cw  = arc(s_last, pt_last, s_first, pt_first, cw=True)
        path_ccw = arc(s_last, pt_last, s_first, pt_first, cw=False)

        chosen = path_cw if shoelace(user + path_cw) <= shoelace(user + path_ccw) \
                         else path_ccw
        self._poly_verts.extend(chosen)
        self._finalize_polygon(self._poly_commit_dir)
        self._clear_polygon_drawing()
        self._refresh()

    def _on_key_press(self, event):
        if not self._poly_mode:
            return
        if event.key == "escape":
            self._clear_polygon_drawing()
            self.fig.canvas.draw_idle()
        elif event.key == "enter" and len(self._poly_verts) >= 3:
            self._finalize_polygon(self._poly_commit_dir)
            self._clear_polygon_drawing()
            self._refresh()
        elif event.key == "c" and self._poly_verts:
            self._close_via_edge()
        elif event.key == "backspace" and self._poly_verts:
            self._poly_verts.pop()
            self._update_poly_display()
            self.fig.canvas.draw_idle()
        elif event.key == "r":
            self._poly_commit_dir = "remove" if self._poly_commit_dir == "add" else "add"
            self.btn_polygon.label.set_text(self._poly_btn_label())
            print(f"Polygon direction → {self._poly_commit_dir.upper()}")
            self.fig.canvas.draw_idle()

    # ------------------------------------------------------------------
    # Output generation
    # ------------------------------------------------------------------

    def _on_generate(self, _event):
        has_paint = bool(self._paint_add or self._paint_remove)
        if (not self._peak_selected and not self.manual_mode
                and self.loaded_mask_preview is None and not has_paint):
            print("Please click on the histogram to select the peak first "
                  "(or load a mask with --load-mask, or paint/draw a polygon mask).")
            return

        print("\nGenerating output …")
        print(f"  Mode         : "
              f"{'manual' if self.manual_mode else ('smooth µ' if self.use_smooth else 'Gaussian')}")
        print(f"  Output path  : {self.output_path}")
        # The same fit resolution and smoothing strength the SLURM job uses,
        # so this output and the cluster's are the same computation.
        print(f"  Bin factor   : {self.slurm_bin}  (--slurm-bin)")
        print(f"  smoothn s    : {SMOOTHN_S:g}")

        _, h_full, w_full = self._full_shape
        save_mask = bool(self.chk_save_mask.get_status()[0])
        ti = self.current_tilt
        comparison_orig = None
        comparison_proc = None

        mask_path = os.path.splitext(self.output_path)[0] + "_mask.mrc"
        mask_mrc = (
            mrcfile.new_mmap(mask_path, shape=(self.n_tilts, h_full, w_full),
                             mrc_mode=0, overwrite=True)
            if save_mask else None
        )
        if mask_mrc is not None:
            mask_mrc.voxel_size = self._voxel_size

        filled_fraction = {}
        try:
            with mrcfile.new_mmap(self.output_path,
                                  shape=(self.n_tilts, h_full, w_full),
                                  mrc_mode=2, overwrite=True) as out_mrc:
                out_mrc.voxel_size = self._voxel_size
                for i in tqdm(range(self.n_tilts), desc="Inpainting tilts"):
                    img = self._mmap_data[i].astype(float)
                    mask = self._effective_mask_full(i, img)

                    result = inpaint_tilt(img, mask, bin_factor=self.slurm_bin)
                    out_mrc.data[i] = result
                    filled_fraction[i] = float(1.0 - mask.mean())
                    if mask_mrc is not None:
                        mask_mrc.data[i] = mask.astype(np.int8)
                    if i == ti:
                        comparison_orig = img.astype(np.float32)
                        comparison_proc = result
        finally:
            if mask_mrc is not None:
                mask_mrc.close()

        print(f"Saved → {self.output_path}")
        if save_mask:
            print(f"Mask  → {mask_path}")

        outputs = [fingerprint(self.output_path)]
        if save_mask:
            outputs.append({"role": "mask", **fingerprint(mask_path)})
        self._record_mask_step(outputs)
        self._record_gui_inpaint(self.output_path,
                                 mask_path if save_mask else None,
                                 filled_fraction)

        self._show_comparison(comparison_orig, comparison_proc)

    def _show_comparison(self, orig: np.ndarray, proc: np.ndarray):
        ti   = self.current_tilt
        lo, hi, mu, sigma = self._get_lo_hi(ti)
        mask = self._effective_mask_full(ti, orig)
        bad  = ~mask

        valid = orig[mask] if mask.any() else orig.ravel()
        vlo   = float(np.percentile(valid, 1))
        vhi   = float(np.percentile(valid, 99))

        fig, axes = plt.subplots(1, 3, figsize=(15, 5))
        axes[0].imshow(orig, cmap="gray", origin="lower", vmin=vlo, vmax=vhi)
        axes[0].set_title(f"Original  (tilt {ti + 1})")
        axes[0].axis("off")

        rgba = np.zeros((*orig.shape, 4), dtype=np.float32)
        rgba[bad] = [1.0, 0.2, 0.2, 0.5]
        axes[1].imshow(orig, cmap="gray", origin="lower", vmin=vlo, vmax=vhi)
        axes[1].imshow(rgba, origin="lower")
        mask_title = f"Mask  (lo={lo:.1f}, hi={hi:.1f})" if lo is not None else "Mask  (loaded/paint)"
        axes[1].set_title(mask_title)
        axes[1].axis("off")

        axes[2].imshow(proc, cmap="gray", origin="lower", vmin=vlo, vmax=vhi)
        axes[2].set_title("Inpainted output")
        axes[2].axis("off")

        fig.suptitle(f"Output written to:  {self.output_path}", fontsize=10)
        plt.tight_layout()
        plt.show()

    # ------------------------------------------------------------------
    # SLURM / parallel output helpers
    # ------------------------------------------------------------------

    def _save_params_json(self) -> str:
        """Serialise per-tilt thresholds (and paint overlays) to JSON + optional mask MRC."""
        has_paint = bool(self._paint_add or self._paint_remove)
        has_loaded_mask = self.loaded_mask_preview is not None
        if not self._peak_selected and not self.manual_overrides and not has_paint and not has_loaded_mask:
            print("No peak selected, no manual overrides, no loaded mask, and no paint "
                  "overlays — nothing to save.")
            return None

        stem = os.path.splitext(self.input_path)[0]
        params_path = stem + "_inpaint_params.json"

        tilts = {}
        for i in range(self.n_tilts):
            lo, hi, _, _ = self._get_lo_hi(i)
            if lo is None:
                lo, hi = self._global_min, self._global_max
            tilts[str(i)] = {"lo": float(lo), "hi": float(hi)}

        save_mask = bool(self.chk_save_mask.get_status()[0])

        payload = {
            "input":         os.path.abspath(self.input_path),
            "output":        os.path.abspath(self.output_path),
            "save_mask":     save_mask,
            "tilts":         tilts,
        }
        # Bake the final spatial masks into a companion MRC so inpaint_apply.py uses
        # them exactly.  Required when paint/polygon overlays, a loaded mask or Top CC
        # mean the mask can't be reproduced from the lo/hi thresholds alone — and also
        # whenever "Save mask MRC" is ticked, because odd/even half-sets need this file:
        # the thresholds were fitted to full-dose intensities and do not transfer to
        # half-dose data, so the halves cannot re-derive the mask for themselves.
        if save_mask or has_paint or has_loaded_mask or self._keep_top_cc:
            _, h_full, w_full = self._full_shape
            mask_mrc_path  = stem + "_inpaint_mask.mrc"
            with mrcfile.new_mmap(mask_mrc_path,
                                  shape=(self.n_tilts, h_full, w_full),
                                  mrc_mode=0, overwrite=True) as mrc:
                # inpaint_apply.py checks this mask against the stack it masks;
                # a 0 pixel size here is one more way that comparison misleads.
                mrc.voxel_size = self._voxel_size
                for i in tqdm(range(self.n_tilts), desc="Baking masks"):
                    img = self._mmap_data[i].astype(float)
                    m = self._effective_mask_full(i, img)
                    mrc.data[i] = m.astype(np.int8)
            payload["mask_mrc"] = os.path.abspath(mask_mrc_path)
            print(f"Mask MRC  → {mask_mrc_path}")

        with open(params_path, "w") as f:
            json.dump(payload, f, indent=2)
        print(f"Params JSON → {params_path}")

        outputs = [{"role": "params", **fingerprint(params_path)}]
        if "mask_mrc" in payload:
            outputs.append({"role": "mask", **fingerprint(payload["mask_mrc"])})
        self._record_mask_step(outputs)
        return params_path

    # ------------------------------------------------------------------
    # Provenance sidecars
    # ------------------------------------------------------------------
    #
    # This window is the mask's provenance and nothing else records it: the
    # thresholds live in a JSON beside the data, and whether a mask came from
    # a Gaussian fit, a hand-drawn polygon or a previous session's file is
    # visible only to whoever was sitting here at the time.

    def _mask_provenance(self) -> dict:
        """How the mask was arrived at — the part that cannot be re-derived."""
        return {
            "peak_fit": bool(self._peak_selected),
            "smooth_mu": bool(self.use_smooth),
            "manual_mode": bool(self.manual_mode),
            "n_manual_overrides": len(self.manual_overrides),
            "painted_tilts": sorted(
                set(self._paint_add) | set(self._paint_remove)),
            "loaded_mask": (
                os.path.abspath(self._mask_mrc._iostream.name)
                if self._mask_mrc is not None else None),
            "keep_top_cc": bool(self._keep_top_cc),
            "uncovered_sentinel": bool(self._has_uncovered),
            # True when the mask can be rebuilt from the lo/hi thresholds
            # alone. False means the baked mask MRC is the only record of it —
            # and that the odd/even half-sets must be given that file, since
            # they cannot re-derive it from their own intensities.
            "reproducible_from_thresholds": not (
                bool(self._paint_add) or bool(self._paint_remove)
                or self._mask_mrc is not None or self._keep_top_cc),
        }

    def _threshold_source(self, tilt_index: int, lo) -> str:
        if lo is None:
            return "global"
        if tilt_index in self.manual_overrides:
            return "manual"
        return "smooth_fit" if self.use_smooth else "fit"

    def _record_mask_step(self, outputs: list) -> None:
        """Write the mask design's sidecar beside its primary artifact.

        ``outputs`` is fingerprint dicts, most significant first: the mask MRC
        when one was baked, else the params JSON. The sidecar sits beside that
        first entry, because that is the file inpaint_apply is pointed at, and
        the rest are named in ``effective`` as companions.

        Guarded: a sidecar that cannot be written must not cost someone the
        mask they just spent twenty minutes drawing.
        """
        if not self.write_sidecars or not outputs:
            return
        try:
            provenance = self._mask_provenance()
            effective = {
                "input": os.path.abspath(self.input_path),
                "n_tilts": int(self.n_tilts),
                "stack_shape": [int(v) for v in self._full_shape],
                "pixel_size_A": float(self._voxel_size.x),
                "preview_bin_factor": int(self.bin_factor),
                "global_min": float(self._global_min),
                "global_max": float(self._global_max),
                "n_sigma_lo": float(self.n_sigma_lo),
                "n_sigma_hi": float(self.n_sigma_hi),
                # A mask changes no geometry, so the canvas frame stitching
                # declared still describes these pixels.
                "frame": "canvas",
                "geometry_unchanged": True,
                **provenance,
            }
            tilts = []
            for i in range(self.n_tilts):
                lo, hi, mu, sigma = self._get_lo_hi(i)
                source = self._threshold_source(i, lo)
                if lo is None:
                    lo, hi = self._global_min, self._global_max
                tilts.append({
                    "tilt_index": i,
                    "lo": float(lo),
                    "hi": float(hi),
                    "threshold_source": source,
                    "fitted_mu": float(mu) if mu is not None else float("nan"),
                    "fitted_sigma": (float(sigma) if sigma is not None
                                     else float("nan")),
                })
            effective["tilts"] = tilts
            effective["auto_resolved"] = ([] if self.manual_mode
                                          else ["lo", "hi"])
            effective["companion_outputs"] = [o["path"] for o in outputs[1:]]
            path = sidecar.write(
                outputs[0]["path"],
                SIDECAR_STEP,
                effective,
                inputs=[fingerprint(self.input_path, role="stitched_stack")],
                requested={"n_sigma": float(self.n_sigma_lo)},
            )
            print(f"Sidecar  → {path}  [inpaint_mask]")
        except Exception as exc:            # noqa: BLE001 - never lose the mask
            logger.warning("Could not write the sidecar (%s); "
                           "the mask and params files are unaffected.", exc)

    def _record_gui_inpaint(self, output_path: str, mask_out_path: str,
                            filled_fraction: dict) -> None:
        """Record the "Generate Output" button as an apply step in its own right.

        Pressing it does the whole job locally — mask *and* fill — so it is
        recorded as both steps rather than pretending the stack it wrote came
        from the cluster.
        """
        if not self.write_sidecars:
            return
        try:
            tilts = []
            for i in range(self.n_tilts):
                lo, hi, _, _ = self._get_lo_hi(i)
                if lo is None:
                    lo, hi = self._global_min, self._global_max
                tilts.append({
                    "tilt_index": i,
                    "lo": float(lo),
                    "hi": float(hi),
                    "mask_source": "gui_interactive",
                    "filled_fraction": float(
                        filled_fraction.get(i, float("nan"))),
                })
            sidecar.write(
                output_path,
                "inpaint_apply",
                {
                    "variant": "full",
                    "input": os.path.abspath(self.input_path),
                    "output": os.path.abspath(output_path),
                    "mask_output": (os.path.abspath(mask_out_path)
                                    if mask_out_path else None),
                    "mask_source": "gui_interactive",
                    "smoothn_s": float(SMOOTHN_S),
                    "smoothn_max_iter": int(SMOOTHN_MAX_ITER),
                    "bin_factor": int(self.slurm_bin),
                    "n_tilts": int(self.n_tilts),
                    "pixel_size_A": float(self._voxel_size.x),
                    "frame": "canvas",
                    "geometry_unchanged": True,
                    "ran_in": "mask_and_inpaint GUI, not on the cluster",
                    "tilts": tilts,
                },
                inputs=[fingerprint(self.input_path, role="stitched_stack")],
                requested={"bin_factor": int(self.slurm_bin)},
            )
        except Exception as exc:            # noqa: BLE001
            logger.warning("Could not record the inpainting (%s); "
                           "the output stack is unaffected.", exc)

    def _write_slurm_script(self, params_path: str) -> str:
        """Write a SLURM batch script that calls inpaint_apply. Returns file path."""
        stem = os.path.splitext(self.input_path)[0]
        script_path = stem + "_inpaint_slurm.sh"
        apply_script = os.path.join(
            os.path.dirname(os.path.abspath(__file__)), "inpaint_apply.py"
        )

        if self.slurm_template and os.path.exists(self.slurm_template):
            with open(self.slurm_template) as f:
                template = f.read()
        else:
            template = DEFAULT_SLURM_TEMPLATE

        # Half-set paths matching beam_mask_motioncorr.py's odd/even sums, used by
        # the commented-out Noise2Noise blocks in the default template.
        in_stem, in_ext = os.path.splitext(os.path.abspath(self.input_path))

        # Prefer the mask just baked alongside the params file: it already exists, so
        # the half-set jobs can run in any order.  Only when nothing was baked do we
        # fall back to the mask the full-dose run itself emits, which does impose an
        # ordering constraint.
        with open(params_path) as f:
            saved_payload = json.load(f)
        mask_path = saved_payload.get("mask_mrc") or (
            os.path.splitext(os.path.abspath(self.output_path))[0] + "_mask.mrc"
        )

        job_name = os.path.basename(stem)[:32]
        content  = template.format(
            job_name   = job_name,
            n_cpus     = self.slurm_cpus,
            npertilt   = self.slurm_npertilt,
            bin_factor = self.slurm_bin,
            mem_gb     = self.slurm_mem,
            time       = self.slurm_time,
            log_file   = stem + "_inpaint_%j.log",
            script_path= apply_script,
            input      = os.path.abspath(self.input_path),
            output     = os.path.abspath(self.output_path),
            params     = os.path.abspath(params_path),
            input_odd  = f"{in_stem}_odd{in_ext}",
            output_odd = f"{in_stem}_odd_inpainted{in_ext}",
            input_even = f"{in_stem}_even{in_ext}",
            output_even= f"{in_stem}_even_inpainted{in_ext}",
            mask       = mask_path,
        )
        with open(script_path, "w") as f:
            f.write(content)
        print(f"SLURM script → {script_path}")
        print(f"Submit with:    sbatch {script_path}")
        return script_path

    def _on_save_slurm(self, _event):
        params_path = self._save_params_json()
        if params_path is not None:
            self._write_slurm_script(params_path)

    def show(self):
        plt.show()
        if self._mask_mrc is not None:
            self._mask_mrc.close()


# ---------------------------------------------------------------------------
# CLI
# ---------------------------------------------------------------------------

def main():
    parser = argparse.ArgumentParser(
        description=(
            "Peak-following mask editor for montage tilt series MRC files.\n"
            "Click on the histogram peak you want to keep; the Gaussian fit\n"
            "is propagated across all tilts to track intensity drift with tilt.\n"
            "Use 'Smooth µ' to stabilise high-tilt fits; use 'Manual' for full\n"
            "manual control with a range slider."
        ),
        formatter_class=argparse.RawDescriptionHelpFormatter,
    )
    parser.add_argument("input", help="Input MRC tilt series file")
    parser.add_argument(
        "-o", "--output",
        help="Output MRC file (default: <input>_inpainted.mrc)",
        default=None,
    )
    parser.add_argument(
        "-b", "--max-pixels",
        help="Longest dimension of the preview (default: 1024).",
        type=int, default=1024, dest="max_pixels",
    )
    parser.add_argument(
        "--preview-binning",
        choices=["zoom", "fourier"], default="zoom", dest="preview_binning",
        help="Method used to downsample tilts to --max-pixels for display: "
             "'zoom' is bilinear resampling (default, fast, some aliasing); "
             "'fourier' crops in Fourier space before inverse-transforming, "
             "which avoids aliasing at the cost of some ringing near sharp "
             "edges. Preview only — does not affect the full-resolution mask.",
    )
    parser.add_argument(
        "-n", "--n-sigma",
        help="Initial N-sigma threshold, applied to both the lower and upper "
             "bound (default: 3.0).",
        type=float, default=3.0, dest="n_sigma",
    )
    parser.add_argument(
        "--slurm-template",
        help="Path to a custom SLURM script template (uses built-in template if omitted).",
        default=None, dest="slurm_template",
    )
    parser.add_argument(
        "--slurm-cpus", type=int, default=8, dest="slurm_cpus",
        help="Total thread budget (ntotal) for the SLURM task; "
             "ntotal // --slurm-npertilt tilts run concurrently (default: 8).",
    )
    parser.add_argument(
        "--slurm-npertilt", type=int, default=4, dest="slurm_npertilt",
        help="DCT threads per tilt (npertilt) inside smoothn (default: 4).",
    )
    parser.add_argument(
        "--slurm-bin", type=int, default=2, dest="slurm_bin",
        help="Downsample factor for the smoothn fit before upsampling back to "
             "full resolution; 1 disables binning (default: 2).",
    )
    parser.add_argument(
        "--slurm-mem", type=int, default=32, dest="slurm_mem",
        help="Memory in GB for SLURM job (default: 32).",
    )
    parser.add_argument(
        "--slurm-time", default="02:00:00", dest="slurm_time",
        help="Wall-clock time limit for SLURM job (default: 02:00:00).",
    )
    parser.add_argument(
        "--load-mask",
        help="Path to an existing mask MRC (e.g. a previous session's "
             "*_mask.mrc / *_inpaint_mask.mrc) to load as the starting mask. "
             "Paint/polygon edits and Top CC are applied on top of it.",
        default=None, dest="load_mask",
    )
    parser.add_argument(
        "--no-sidecar", action="store_false", default=True, dest="sidecar",
        help="Do not write a provenance sidecar for the mask. Sidecars are ON "
             "by default and record how the mask was arrived at (fit, manual, "
             "painted, loaded), the per-tilt thresholds, and whether it can be "
             "reproduced from those thresholds alone — which is what decides "
             "whether the odd/even half-sets must be handed the baked mask "
             "file. The sidecar sits beside the mask MRC when one is baked, "
             "else beside the params JSON.",
    )
    args = parser.parse_args()
    logging.basicConfig(level=logging.INFO, format="%(levelname)-8s %(message)s")

    if not os.path.exists(args.input):
        sys.exit(f"Error: file not found: {args.input}")
    if args.load_mask and not os.path.exists(args.load_mask):
        sys.exit(f"Error: mask file not found: {args.load_mask}")

    print(f"Opening {args.input} as memory-map …")
    with mrcfile.mmap(args.input, mode="r", permissive=True) as mrc:
        ndim = mrc.data.ndim
        if ndim not in (2, 3):
            sys.exit(f"Error: expected 2-D or 3-D MRC data, got shape {mrc.data.shape}")
        shape = mrc.data.shape
        n = 1 if ndim == 2 else shape[0]
        h, w = shape[-2], shape[-1]
        print(
            f"Opened:  {n} tilt(s),  {h} × {w} px  "
            f"({n * h * w * mrc.data.itemsize / 1e9:.2f} GB on disk)"
        )
        editor = PeakMaskEditor(
            mrc, args.input, args.output,
            max_px=args.max_pixels, preview_binning=args.preview_binning,
            n_sigma_default=args.n_sigma,
            slurm_template=args.slurm_template,
            slurm_cpus=args.slurm_cpus,
            slurm_npertilt=args.slurm_npertilt,
            slurm_bin=args.slurm_bin,
            slurm_mem=args.slurm_mem,
            slurm_time=args.slurm_time,
            load_mask_path=args.load_mask,
            write_sidecars=args.sidecar,
        )
        editor.show()


if __name__ == "__main__":
    main()
```
