## Supplementary material for "Whole cryo-FIB lamella acquisition at high magnification with rectangular beam montage cryo-electron tomography": Source code: mask_tuning_widgets.py

```
"""
Shared matplotlib-widget building blocks for the interactive mask-tuning
tools (Choose_fringe_size.py, Choose_mask_params.py): a percentile-ranged
threshold slider, a "threshold as fraction of median" text readout, and
clamped prev/next navigation buttons.
"""

import numpy as np
from matplotlib.widgets import Button, Slider


def make_threshold_slider(fig, rect, image, initial_threshold,
                           label="Threshold (counts)", pct_lo=1, pct_hi=99):
    """Slider spanning [pct_lo, pct_hi] percentile of image counts.

    The percentile range (rather than min/max) keeps hot/dead outlier
    pixels from collapsing the usable slider range.
    """
    ax = fig.add_axes(rect)
    return Slider(
        ax, label,
        float(np.percentile(image, pct_lo)), float(np.percentile(image, pct_hi)),
        valinit=initial_threshold,
    )


def make_fraction_text(fig, x, y, prefix="", **text_kwargs):
    """Create a text artist reporting threshold as a fraction of the median.

    Returns update(threshold, median) which sets the text.
    """
    text = fig.text(x, y, "", **text_kwargs)

    def update(threshold, median):
        frac = threshold / median if median else 0.0
        text.set_text(f"{prefix}{frac:.4f} × median   (median = {median:.1f} counts)")

    return update


def make_nav_buttons(fig, prev_rect, next_rect, count, on_navigate,
                      prev_label="← Prev", next_label="Next →", start=0):
    """Wire prev/next buttons over a clamped index range [0, count).

    on_navigate(new_idx) is called after each click with the clamped index.
    Returns (btn_prev, btn_next, index) where index is a mutable
    single-element list holding the current index.
    """
    index = [start]

    def step(delta):
        def handler(event):
            index[0] = max(0, min(count - 1, index[0] + delta))
            on_navigate(index[0])
        return handler

    ax_prev = fig.add_axes(prev_rect)
    ax_next = fig.add_axes(next_rect)
    btn_prev = Button(ax_prev, prev_label)
    btn_next = Button(ax_next, next_label)
    btn_prev.on_clicked(step(-1))
    btn_next.on_clicked(step(1))
    return btn_prev, btn_next, index
```
