## Supplementary material for "Whole cryo-FIB lamella acquisition at high magnification with rectangular beam montage cryo-electron tomography": Source code: montage_projection.py

```
#!/usr/bin/env python3
"""
Project 3D tomogram coordinates back onto individual montage tiles.

A square-beam montage tomogram is reconstructed from a stitched tilt series,
so a 3D pick has no direct relationship to the raw data: the pixel it came from
lives somewhere in one of ~20 tiles making up one tilt, and which tile that is
changes from tilt to tilt. This module inverts the whole chain, taking a point
(X, Y, Z) in the reconstruction to (tilt, tile, z-slice, x, y) in the
per-tilt tile stacks written by ``beam_mask_motioncorr --stack-output``.

The chain, in order, and where each link came from
--------------------------------------------------
1. **tomogram -> aligned frame.** Inverting the rotation in
   ``refine_montage_projmatch.reproject_volume``::

       col_al = cx + (X - cx) cos(theta) - s (Z - cz) sin(theta)
       row_al = Y

   ``theta`` is from the AreTomo ``.aln`` TILT column as written, which already 
   includes AreTomo's ``-TiltCor`` offset.

   ``s`` is ``z_sign``, the direction of the volume's depth axis, and defaults
   TO -1, i.e. the formula above with the sign flipped, this assumes standard
   AreTomo reconstruction followed by a clip rotx (-90 degrees about x axis) 
   rotation. 

   ``cx`` and ``cz`` are the volume centre, ``(n-1)/2`` -- which is a claim
   about where AreTomo put the reconstruction, not a measurement of it. This
   link is the one that can be tested on its own, because AreTomo writes the
   aligned stack it reconstructed from: ``measure_reprojection_residual
   --frame aligned`` measures the residual there, with links 2-4 out of the
   loop, and its fit names which of ``cz``, ``cx``, ``theta`` is off. The
   answer goes back in through ``z_centre_offset`` / ``x_centre_offset`` /
   ``theta_offset_deg``.

2. **aligned -> canvas.** The inverse of
   ``refine_montage_projmatch.alignment_matrix``, whose sign convention
   (``rot-1 shift-1``) was settled against AreTomo's own aligned stack, plus
   the **local (patch) shift** when the volume was reconstructed with
   ``-Patch``. The patch field is not a refinement of the rigid alignment but a
   term inside it -- AreTomo adds it between the rotation and the translation
   -- and omitting it left a 191 A position-dependent error. See
   ``docs/reprojection_geometry.md`` for the derivation and the full equation.

3. **canvas -> tile.** ``MontageGeometry`` places tile *t* from its position in
   microns. A point may fall inside several overlapping tiles; ``index_map``
   decides whether it is covered by real data at all, and among the covering
   tiles we take the one that holds the point furthest from any tile edge.

4. **canvas pixel -> raw tile pixel.** Undo ``stitch.py --rotate`` and the
   stitch binning.

5. **tile -> z-slice.** ``index_map`` labels run 0..n_selected-1, *not* over the
   rows of ``Refined_positions``; map through ``flatnonzero(tile_selection)``.
   The result indexes the per-tilt stack directly, because that stack has one
   slice per position row (verified for all 41 tilts of dataset 9-A).

Two traps worth stating explicitly, because both are silent
-----------------------------------------------------------
* ``stitch.py --rotate`` is **not** recorded in the HDF5, and getting it wrong
  displaces every tile by (2880-2046)/2 = 417 canvas pixels. ``rotate="auto"``
  measures it from ``index_map`` rather than trusting an argument, but a tile
  footprint is unchanged by 180 degrees, so auto can only narrow it to k or
  k+2. Prefer :meth:`MontageProjector.from_sidecars`, which reads the argument
  stitch was actually given (``transforms: raw_tile -> canvas, rot90_k``) and
  settles it outright. Note that ``rot90_k`` is stitch's *own* rotation, which
  is not the same as the total from the detector frame: when motion correction
  has already rotated the tiles, stitch applies 0 and the sidecars record both
  numbers separately.
* ``index_map`` labels index the *selected* tiles. When ``tile_selection`` has
  any False entry (it does, on more than half the tilts of 9-A) label != row of
  ``Refined_positions`` and everything downstream lands on the wrong tile.
"""

from __future__ import annotations

import logging
import os
from dataclasses import dataclass, field
from typing import Dict, List, Optional, Sequence, Tuple

import h5py
import numpy as np

from .refine_montage_projmatch import (
    AlnFile,
    Convention,
    LocalAlignment,
    MontageGeometry,
    SectionAlignment,
    alignment_matrix,
    alignment_shift_rc,
    find_position_files,
    map_sections_to_tilts,
    parse_aln,
)
from .Utilities import roll_no_periodic

logger = logging.getLogger(__name__)

#: Cap on the fixed-point iterations that invert the local shift field. The
#: contraction is strong (|dL| << 1), so this is a safety net, not a budget.
_LOCAL_INVERSE_ITERS = 20


# ---------------------------------------------------------------------------
# Per-tilt bookkeeping
# ---------------------------------------------------------------------------


@dataclass
class TiltInfo:
    """Everything needed to place a point onto the tiles of one tilt."""

    tilt: float  # nominal stage tilt, from the filename
    sec: SectionAlignment  # the .aln row joined by SEC
    positions_um: np.ndarray  # (n_tiles, 2), full length, (x, y) microns
    tile_selection: np.ndarray  # (n_tiles,) bool
    h5path: str
    stack_path: Optional[str] = None  # per-tilt tile stack MRC
    _index_map: Optional[np.ndarray] = field(default=None, repr=False)
    _beam_masks: Optional[np.ndarray] = field(default=None, repr=False)

    @property
    def selected(self) -> np.ndarray:
        """Rows of ``positions_um`` that ``index_map`` labels 0, 1, 2, ..."""
        return np.flatnonzero(self.tile_selection)

    def index_map(self) -> np.ndarray:
        """Load (and cache) the canvas partition. ~150 MB per tilt."""
        if self._index_map is None:
            with h5py.File(self.h5path, "r") as h:
                self._index_map = h["index_map"][:]
        return self._index_map

    def beam_masks(self) -> Optional[np.ndarray]:
        """
        Per-tile square-beam footprints, ``(n_selected, *tile_shape)`` bool.

        stitch.py stores a single eroded beam-reference mask
        (``beam_reference_mask``) plus each tile's ``(dy, dx)`` alignment
        shift (``beam_shifts``), both in tile-local canvas coordinates --
        i.e. binned and already rotated -- and ``beam_shifts`` in *selected*
        tile order, like ``Refined_positions`` restricted to the selection.
        A tile's actual mask is never stored; it is reconstructed here by
        rolling the reference mask into place, exactly as stitch.py does
        per tile during the run.

        This matters for extraction, not just for tidiness. The illuminated
        square is smaller than the detector frame and sits off-centre in it, so
        a particle can be comfortably inside the tile rectangle and still have
        the beam edge running through its box. On dataset 9-A that was ~15% of
        boxes, and a beam edge inside a particle box is a far worse artefact
        than a missing observation.
        """
        if self._beam_masks is None:
            with h5py.File(self.h5path, "r") as h:
                if "beam_reference_mask" not in h or "beam_shifts" not in h:
                    return None
                ref_mask = h["beam_reference_mask"][:].astype(bool)
                shifts = h["beam_shifts"][:]
            self._beam_masks = np.stack(
                [
                    roll_no_periodic(ref_mask, (-int(dy), -int(dx)), axis=(0, 1))
                    for dy, dx in shifts
                ]
            )
        return self._beam_masks

    def drop_index_map(self) -> None:
        self._index_map = None
        self._beam_masks = None


# ---------------------------------------------------------------------------
# Assignment result
# ---------------------------------------------------------------------------


@dataclass
class TileHit:
    """One particle seen in one tile of one tilt."""

    tilt: float
    tile: int  # row of Refined_positions == z-slice of the per-tilt stack
    x: float  # raw (unbinned) tile pixel, column
    y: float  # raw (unbinned) tile pixel, row
    canvas_rc: Tuple[float, float]
    depth: float  # along-beam offset from the tomogram centre, unbinned px
    edge_margin: float  # distance to the nearest tile edge, raw px


# ---------------------------------------------------------------------------
# The projector
# ---------------------------------------------------------------------------


class MontageProjector:
    """
    Maps reconstruction coordinates onto the raw tiles of a square-beam montage.

    All 3D input is in *reconstruction voxels*, (X, Y, Z) = (column, row, slice)
    of the AreTomo volume after ``clip rotx`` -- i.e. exactly what
    ``model2point`` prints for a pick made in that volume.
    """

    def __init__(
        self,
        aln_path: str,
        positions_dir: str,
        recon_shape: Sequence[int],  # (nz, ny, nx) of the reconstruction
        roi: Sequence[float] = (-5500, 12000, -6000, 12000),
        pixel_size: float = 3.426,
        binning: int = 2,
        out_bin: int = 2,
        rotate: object = "auto",
        raw_tile_shape: Optional[Tuple[int, int]] = None,
        tile_dir: Optional[str] = None,
        conv: Convention = Convention(),
        extra_shift: Sequence[float] = (0.0, 0.0),
        handedness: int = 1,
        exclude_sections: Sequence[int] = (),
        use_local: bool = True,
        z_centre_offset: float = 0.0,
        x_centre_offset: float = 0.0,
        theta_offset_deg: float = 0.0,
        z_sign: int = -1,
    ):
        # Read AreTomo's .aln file
        self.aln = parse_aln(aln_path)
        self.recon_shape = tuple(int(x) for x in recon_shape)
        self.out_bin = int(out_bin)
        self.binning = int(binning)
        self.pixel_size = float(pixel_size)
        self.conv = conv
        self.extra_shift = np.asarray(extra_shift, dtype=float)
        self.handedness = int(handedness)

        # The three free parameters of link 1, all zero by default because the
        # defaults are *derivations*, not guesses: cz = (nz-1)/2 assumes AreTomo
        # centred the slab on the specimen mid-plane, cx = (nx-1)/2 assumes it
        # centred the volume on the aligned image, and theta is the .aln TILT
        # column. Each is a thing that could be false, and each leaves its own
        # signature in the reprojection residual (see
        # measure_reprojection_residual's fit); these knobs exist so a fitted
        # coefficient can be applied and the measurement repeated, rather than
        # the derivation being argued about. NOT for tuning until the residual
        # looks small -- an offset here is a claim about the reconstruction.
        self.z_centre_offset = float(z_centre_offset)
        self.x_centre_offset = float(x_centre_offset)
        self.theta_offset_deg = float(theta_offset_deg)

        # Which way the volume's Z axis runs, relative to the orientation
        # tomo_to_aligned's formula (below) is written for -- the textbook
        # reading of AreTomo's own kernels, no correction folded in. -1 is
        # that orientation, and -1 is also what every volume this pipeline has
        # made so far actually needs: AreTomo without -FlipVol, then `clip
        # rotx`, measured on 9-A. +1 is the other way round -- both flips
        # applied, or neither -- and unusual enough to warn about.
        #
        # This is a property of the FILE, not a fitted correction, so unlike
        # the three offsets above it comes from the sidecar. Getting it wrong
        # costs 2*dz*sin(theta) of position, which is ZERO at the 0 deg section
        # -- the one an IMOD model is easiest to check on -- and grows to the
        # full 2*dz at the extremes. Check it at +-60 deg or not at all.
        if int(z_sign) not in (1, -1):
            raise ValueError(f"z_sign must be +1 or -1, got {z_sign}")
        self.z_sign = int(z_sign)
        if self.z_sign == 1:
            logger.warning(
                "z_sign=+1: the volume's depth axis is being taken as running "
                "opposite to every volume this pipeline has reconstructed so "
                "far (clip rotx, no -FlipVol). Check flip_vol/clip_rotx.")
        if any((self.z_centre_offset, self.x_centre_offset, self.theta_offset_deg)):
            logger.warning(
                "tomogram -> aligned frame is being shifted from its derived "
                "values: z centre %+.2f, x centre %+.2f recon voxels, theta "
                "%+.4f deg.", self.z_centre_offset, self.x_centre_offset,
                self.theta_offset_deg,
            )

        # The patch field, if the reconstruction was made with one. Applying it
        # is the default because the picks come out of a volume that already
        # has it baked in; `use_local=False` exists to reproduce the old,
        # global-only behaviour for comparison, not as a normal setting.
        self.local: Optional[LocalAlignment] = self.aln.local if use_local else None
        if self.aln.has_local and self.local is None:
            logger.warning(
                "%s has a local (patch) alignment but use_local=False: picks "
                "will be reprojected with the global alignment only. Expect a "
                "position-dependent error the size of the patch field.",
                os.path.basename(aln_path),
            )
        elif self.local is None and not self.aln.has_local:
            logger.info(
                "No local alignment in the .aln; the reconstruction had better "
                "have been made without -Patch, or reprojection will be off by "
                "the patch field.",
            )

        # The tomogram is `binning * out_bin` coarser than a raw tile pixel.
        # Derive it from the binning factors, never from the ratio of recorded
        # pixel sizes: AreTomo was given -PixSize 6.87 for a canvas that is
        # really 6.852 A/px, so the header says 13.74 A when the truth is
        # 13.704 A. The geometry is exact; only the label is wrong.
        self.tomo_bin = self.binning * self.out_bin

        posfiles = find_position_files(positions_dir)
        mapping, offset = map_sections_to_tilts(self.aln, list(posfiles))
        self.tilt_offset = offset
        # Every tilt that has a position file, dark frames included. This is the
        # slice order of the stitched canvas stack and of the .aln SEC index --
        # `self.tilts` below holds only the sections AreTomo kept, so the two
        # must not be confused when indexing into a stack.
        self.all_tilts: List[float] = sorted(posfiles)

        stacks = _find_tile_stacks(tile_dir) if tile_dir else {}
        if raw_tile_shape is None:
            raw_tile_shape = _probe_tile_shape(stacks)
        self.raw_tile_shape = tuple(int(x) for x in raw_tile_shape)

        self.tilts: Dict[float, TiltInfo] = {}
        for tilt, path in sorted(posfiles.items()):
            sec = mapping.get(tilt)
            if sec is None:
                logger.info("tilt %+6.1f: dark frame in the .aln, skipped", tilt)
                continue
            if sec.sec in exclude_sections:
                logger.warning("tilt %+6.1f: SEC %d excluded by request", tilt, sec.sec)
                continue
            with h5py.File(path, "r") as h:
                pos = h["Refined_positions"][:]
                sel = np.asarray(h["tile_selection"][:], dtype=bool)
                canvas_shape = h["index_map"].shape
            self.tilts[tilt] = TiltInfo(
                tilt=tilt, sec=sec, positions_um=pos, tile_selection=sel,
                h5path=path, stack_path=stacks.get(tilt),
            )
            self.canvas_shape = tuple(int(x) for x in canvas_shape)

        if not self.tilts:
            raise ValueError("No tilts survived the .aln join -- check the inputs.")

        if str(rotate) == "auto":
            rotate = self._detect_rotation(roi)
        self.rotate = int(rotate)

        self.geom = MontageGeometry.from_stitch_args(
            roi=roi, pixel_size=pixel_size, binning=binning,
            raw_tile_shape=self.raw_tile_shape, rotate=self.rotate,
        )
        if tuple(self.geom.canvas_shape) != self.canvas_shape:
            raise ValueError(
                f"Canvas mismatch: --roi/--pixel-size/--binning reconstruct "
                f"{self.geom.canvas_shape} but index_map is {self.canvas_shape}. "
                f"Check them against 02_stitch.sh."
            )
        logger.info(
            "canvas %s, tile on canvas %s, rotate %d, tomogram binning %d",
            self.geom.canvas_shape, self.geom.tile_shape, self.rotate, self.tomo_bin,
        )

    # -- construction from the provenance sidecars ---------------------------

    @classmethod
    @classmethod
    def from_sidecars(
        cls,
        tomogram_path: str,
        aln_path: str,
        positions_dir: str,
        recon_shape: Sequence[int],
        *,
        canvas_path: Optional[str] = None,
        **overrides,
    ) -> "MontageProjector":
        """
        Build a projector by walking the tomogram's provenance chain.

        Every geometric constant this class takes -- the ROI, the tile pixel
        size, the stitch binning, the rotation, AreTomo's OutBin -- was chosen
        by an earlier step and then had to be *restated* on the command line
        here, correctly, from memory. That is where the 417-pixel rotation trap
        in this module's docstring came from, and no amount of care at the call
        site can fix it: the arguments are a second copy of facts that already
        exist. This reads the originals.

        Anchored at ``tomogram_path``, not at a shared file for the series.
        That is the substantive change from the manifest version and it closes
        the hazard the submission scripts had to warn about in prose: "TOMOGRAM
        AND ALN MUST BE THE SAME RUN". The AreTomo sidecar sits beside *this*
        volume and names the exact stack and alignment that made it, so a
        ``-Patch 5 3`` volume can no longer be silently paired with a
        ``-Patch 10 6`` run's geometry -- the chain simply leads somewhere else.

        ``rotate`` is the value that matters most. Detecting it from
        ``index_map`` works but cannot separate k from k+2 -- a tile footprint
        is unchanged by 180 degrees -- so it flips a coin on an error that maps
        every particle to the opposite corner of its tile. The stitch sidecar
        records the argument stitch was actually given, which settles it.

        Explicit keyword arguments still win, so a value can be overridden for
        a test without editing a sidecar. Anything the chain does not have
        falls back to the constructor's default, with a warning naming it.
        """
        try:
            from . import sidecar
        except ImportError:
            import sidecar

        aretomo = _sidecar_effective(sidecar, tomogram_path, "aretomo")
        # The canvas the tomogram came from: named by the AreTomo sidecar's
        # own input, so there is nothing to pass and nothing to get wrong.
        # inpaint_apply sits between the two on some series and changes no
        # geometry, so the walk skips through it to the stitch record.
        # The chain wins over the caller's canvas. `--canvas-stack` is the
        # stack the *tools* display, which on an inpainted series is not the
        # stack AreTomo was given; preferring it would walk to the wrong
        # stitch record, or to none. It is only a fallback for a tomogram
        # whose own sidecar is missing.
        stitch: Dict[str, object] = {}
        canvas = (aretomo.get("input") if aretomo else None) or canvas_path
        if canvas:
            stitch = _sidecar_effective(sidecar, canvas, "stitch")
            if not stitch:
                inpaint = _sidecar_effective(sidecar, canvas, "inpaint_apply")
                if inpaint.get("input"):
                    stitch = _sidecar_effective(
                        sidecar, inpaint["input"], "stitch")

        kwargs: Dict[str, object] = {}
        missing = []

        def take(name, value, label):
            if value is None:
                missing.append(label)
            else:
                kwargs[name] = value

        take("roi", stitch.get("roi_px"), "stitch roi_px")
        take("pixel_size", stitch.get("pixel_size_A"), "stitch pixel_size_A")
        take("binning", stitch.get("binning"), "stitch binning")
        take("out_bin", aretomo.get("outbin"), "aretomo outbin")
        # The rotation stitch applied to the tiles it was handed. NOT
        # `rotate_deg_from_raw_frames`, which is the total from the detector
        # frame and already includes whatever motion correction did -- the tile
        # stacks this projector reads are motion correction's output, so the
        # only rotation still to undo is stitch's own.
        rot90_k = stitch.get("rot90_k")
        if rot90_k is None and stitch.get("rotate_deg") is not None:
            rot90_k = (int(stitch["rotate_deg"]) // 90) % 4
        take("rotate", None if rot90_k is None else int(rot90_k) * 90,
             "stitch rotate")

        # Which way the volume's depth axis runs. Recorded by aretomo_record
        # from -FlipVol and whether `clip rotx` was applied, because it is a
        # fact about how the file was made and nothing downstream can see it:
        # a reversed depth axis reprojects perfectly at 0 deg and is wrong by
        # 2*dz*sin(theta) everywhere else.
        z_sign = aretomo.get("z_sign")
        if z_sign is None and "clip_rotx" in aretomo:
            from .aretomo_record import depth_axis_sign

            z_sign = depth_axis_sign({"flip_vol": aretomo.get("flip_vol")},
                                     bool(aretomo.get("clip_rotx")))
            logger.info(
                "Sidecar predates z_sign; derived %+d from flip_vol=%s, "
                "clip_rotx=%s.", z_sign, aretomo.get("flip_vol"),
                aretomo.get("clip_rotx"))
        take("z_sign", z_sign, "aretomo z_sign")

        if missing:
            logger.warning(
                "The provenance chain from %s is missing %s; falling back to "
                "the built-in default(s) for those. Check them against the "
                "submission scripts, and consider `montage_backfill` if this "
                "series predates sidecars.",
                os.path.basename(tomogram_path), ", ".join(missing),
            )
        kwargs.update(overrides)
        logger.info(
            "Geometry from sidecars: roi=%s pixel_size=%s binning=%s "
            "out_bin=%s rotate=%s z_sign=%s",
            kwargs.get("roi"), kwargs.get("pixel_size"), kwargs.get("binning"),
            kwargs.get("out_bin"), kwargs.get("rotate"), kwargs.get("z_sign"),
        )

        # A -Patch tomogram whose local field AreTomo did not actually apply is
        # reconstructed in the global-only frame, so applying it here would
        # introduce the very error this is meant to remove. The record knows
        # which; see aretomo_record.local_applied.
        if "use_local" not in overrides:
            applied = aretomo.get("local_alignment_applied")
            if applied is False and aretomo.get("has_local_alignment"):
                logger.warning(
                    "The sidecar says this reconstruction was made with "
                    "-AlnFile, so AreTomo did not apply the patch field to it. "
                    "Reprojecting WITHOUT the local correction to match.")
                kwargs["use_local"] = False

        proj = cls(aln_path=aln_path, positions_dir=positions_dir,
                   recon_shape=recon_shape, **kwargs)
        proj._check_against_sidecars(aretomo, stitch)
        return proj

    def _check_against_sidecars(self, aretomo: dict, stitch: dict) -> None:
        """Cross-check the reconstructed canvas against what was recorded."""
        shape = stitch.get("canvas_shape_px")
        if shape and tuple(int(x) for x in shape) != tuple(self.canvas_shape):
            logger.error(
                "Canvas shape disagrees with the stitch sidecar: %s here, %s "
                "recorded by stitch.", self.canvas_shape, tuple(shape),
            )
        recorded = stitch.get("canvas_pixel_size_A")
        if recorded and abs(recorded - self.geom.canvas_pixel_size) > 1e-3:
            logger.error(
                "Canvas pixel size disagrees with the stitch sidecar: %.4f A "
                "here, %.4f A recorded.", self.geom.canvas_pixel_size, recorded,
            )
        raw_size = aretomo.get("raw_size")
        if raw_size and tuple(raw_size[:2]) != tuple(self.canvas_shape[::-1]):
            logger.error(
                "The .aln's RawSize %s is not this canvas %s — the alignment "
                "may belong to a different stitch.",
                tuple(raw_size[:2]), tuple(self.canvas_shape[::-1]),
            )


    # -- rotation detection -------------------------------------------------

    def _detect_rotation(self, roi: Sequence[float]) -> int:
        """
        Measure ``stitch.py --rotate`` from index_map rather than trusting it.

        The HDF5 does not record the rotation, and getting it wrong shifts every
        tile by half the difference of the tile's two side lengths -- 417 px for
        9-A, far more than any residual we care about. Each candidate rotation
        predicts a tile footprint; the right one puts the centroid of each
        index_map label near the centre of its predicted footprint. Only tiles
        whose footprint lies wholly on the canvas are scored, because a clipped
        footprint biases the centroid regardless of the rotation.
        """
        tilt = min(self.tilts, key=abs)
        info = self.tilts[tilt]
        imap = info.index_map()
        full = info.selected
        scores = {}
        for rot in (0, 90, 180, 270):
            geom = MontageGeometry.from_stitch_args(
                roi=roi, pixel_size=self.pixel_size, binning=self.binning,
                raw_tile_shape=self.raw_tile_shape, rotate=rot,
            )
            org = geom.tile_origin_rc(info.positions_um)
            h, w = geom.tile_shape
            resid = []
            for label in range(int(imap.max())):
                r0, c0 = org[full[label]]
                if r0 < 0 or c0 < 0 or r0 + h > imap.shape[0] or c0 + w > imap.shape[1]:
                    continue
                m = imap == label
                if not m.any():
                    continue
                rr, cc = np.nonzero(m)
                resid.append(
                    np.hypot(rr.mean() - (r0 + h / 2), cc.mean() - (c0 + w / 2))
                )
            scores[rot] = float(np.mean(resid)) if resid else np.inf
        info.drop_index_map()
        best = min(scores, key=scores.get)
        logger.info(
            "rotate=auto on tilt %+.1f: %s -> %d deg",
            tilt, {k: round(v, 1) for k, v in scores.items()}, best,
        )
        if scores[best] > 200:
            logger.error(
                "Even the best rotation leaves a %.0f px mean residual. The ROI, "
                "pixel size or binning are probably wrong for this dataset.",
                scores[best],
            )
        # A footprint test can only see the tile's *shape*, and k and k+2 give
        # the same shape in the same place. So this settles 0-or-180 vs
        # 90-or-270 and nothing more. Getting the remaining 180 deg wrong maps
        # every particle to the diametrically opposite point of its tile, which
        # the patch average in overlay_picks_on_montage will show as pure noise
        # -- that check is not optional.
        opposite = (best + 180) % 360
        if abs(scores[opposite] - scores[best]) < 0.05 * max(scores[best], 1.0):
            logger.warning(
                "rotate=%d and rotate=%d are indistinguishable from tile "
                "footprints alone (%.1f vs %.1f px). Taking %d; confirm it with "
                "the patch average, or with --canvas-stack.",
                best, opposite, scores[best], scores[opposite], best,
            )
        return best

    # -- which slice of which stack -----------------------------------------

    def canvas_slice_index(self, tilt: float) -> int:
        """
        z index of ``tilt`` in the stitched canvas stack.

        The canvas stack holds *every* tilt in ascending order, dark frames
        included -- the same set the .aln indexes with SEC.
        """
        return self.all_tilts.index(tilt)

    def aligned_slice_index(self, tilt: float) -> int:
        """
        z index of ``tilt`` in AreTomo's aligned tilt series.

        That stack holds only the sections AreTomo *kept*, so its z index is the
        rank of the section among the surviving ones and not SEC, and not the
        canvas index either. Confusing the two silently shifts every comparison
        by one slice past the first dark frame.
        """
        surviving = sorted(s.sec for s in self.aln.sections)
        return surviving.index(self.tilts[tilt].sec.sec)

    # -- the chain ----------------------------------------------------------

    def theta(self, tilt: float) -> float:
        """
        Reprojection angle in radians: the .aln TILT column as written, plus
        ``theta_offset_deg`` (normally zero -- see :meth:`tomo_to_aligned`).
        """
        return np.deg2rad(self.tilts[tilt].sec.tilt + self.theta_offset_deg)

    def volume_centre(self) -> Tuple[float, float]:
        """
        ``(cx, cz)`` of the reconstruction, in recon voxels: the point the
        reprojection shear turns about.

        The derived values are the centre of the volume, ``(n-1)/2`` on each
        axis. That is an assumption about what AreTomo did, not a measurement of
        it -- in particular ``cz`` assumes the ``-VolZ`` slab was centred on the
        specimen mid-plane -- and ``z_centre_offset`` / ``x_centre_offset``
        exist to correct it when the residual fit says otherwise.
        """
        nz, ny, nx = self.recon_shape
        return ((nx - 1) / 2.0 + self.x_centre_offset,
                (nz - 1) / 2.0 + self.z_centre_offset)

    def tomo_to_aligned(self, tilt: float, xyz: np.ndarray) -> np.ndarray:
        """
        (N, 3) reconstruction voxels -> (N, 3) of (row, col, depth) in the
        aligned frame, still at the reconstruction's own sampling.

        ``depth`` is the along-beam coordinate, zero at the tomogram centre, and
        is what sets a particle's defocus relative to the tilt image's nominal
        value. Its sign is the classic tomography handedness ambiguity, so it is
        multiplied by ``handedness`` (+-1) rather than asserted.

        ``col`` and ``depth`` are the two components of ONE orthogonal map of
        the x-z plane through theta (the standard rotation formula). Flipping
        the sign of the ``dz`` term in ``col`` alone destroys that -- the
        determinant becomes cos(2 theta), and at 45 deg ``depth`` comes out
        identically equal to ``col - cx`` -- so the depth-axis direction
        enters once, as ``z_sign`` on ``dz``, and the 2x2 below stays
        length-preserving whatever it is set to.

        This link has exactly three fitted-in-principle parameters -- ``cx``,
        ``cz`` and ``theta`` -- and each is derived rather than fitted. Test
        them by measuring the residual against AreTomo's *aligned* stack
        (``measure_reprojection_residual --frame aligned``), which exercises
        this method and nothing else, and feed a fitted coefficient back
        through ``volume_centre``'s offsets or ``theta_offset_deg``. A depth
        axis pointing the other way shows up in that same fit, as a depth scale
        factor of -1; the answer to that one is ``z_sign``, not an offset.
        """
        xyz = np.atleast_2d(np.asarray(xyz, dtype=float))
        # Retrieve tomography volume center
        cx, cz = self.volume_centre()
        # Retrieve the tilt angles in radians
        th = self.theta(tilt)
        # calculate cosine and sine
        cos_t, sin_t = np.cos(th), np.sin(th)
        # Distance from the centre of the volume. dz is measured along the beam
        # with z_sign fixing which way that is; see __init__.
        dx = xyz[:, 0] - cx
        dz = self.z_sign * (xyz[:, 2] - cz)
        # Project the 3D coordinates to the frame of the aligned tilt image in
        # x (column). Textbook sign -- see the class docstring's link-1 note
        # for why z_sign defaults to -1 rather than the reading below.
        col = cx + dx * cos_t - dz * sin_t
        # aligned tilt series is along y axis so y coo]rdinate goes through unchanged
        row = xyz[:, 1]
        # depth is distance along the beam direction, ie. deviation from mean
        # image defocus. Its overall sign is handedness's job.
        depth = self.handedness * (dx * sin_t + dz * cos_t)
        return np.stack([row, col, depth], axis=1)

    def aligned_to_canvas(self, tilt: float, aligned_rc: np.ndarray) -> np.ndarray:
        """
        (N, 2) aligned (row, col) -> (N, 2) canvas (row, col).

        Two terms, and for a ``-Patch`` reconstruction the second is not small.

        The global term inverts :func:`alignment_matrix` as before. The local
        term is AreTomo's patch field, which its correction kernel applies
        *between* the rotation and the global shift::

            p_in = centre + v + L(v) + G,   v = R(ROT) (p_out - centre_out)

        (Correct/GCorrPatchShift.cu:117-133). Read in that order the inverse is
        explicit rather than iterative, because ``v`` is exactly what the purely
        global inverse already computes: ``v = p_global - centre + shift_rc``,
        so the local shift is simply *added* to the globally-inverted position.
        No fixed point, no ambiguity.

        Skipping it is what put a 191 A position-dependent error into the
        RELION export (job 28886886): the field is a median 12 px and a p90 of
        32 canvas px on 9-A, it varies within a single tilt, and no global
        shift can absorb it -- which is precisely why AreTomo measured it.
        """
        sec = self.tilts[tilt].sec
        M, t = alignment_matrix(sec, self.canvas_shape, self.conv, self.out_bin)
        Minv = np.linalg.inv(M)
        aligned_rc = np.atleast_2d(np.asarray(aligned_rc, dtype=float))
        canvas = (aligned_rc - t) @ Minv.T
        if self.local is not None:
            centre = np.array(self.canvas_shape, dtype=float) / 2.0
            # AreTomo's pre-shift, image-centred coordinate, in (row, col).
            v_rc = canvas - centre + alignment_shift_rc(sec, self.conv)
            # The field is defined in (x, y); the table is in unbinned canvas
            # pixels and so is `canvas`, so no rescaling enters here.
            shift_xy = self.local.shift_at(sec.sec, v_rc[:, ::-1])
            canvas = canvas + shift_xy[:, ::-1]
        return canvas + self.extra_shift

    def canvas_to_aligned(self, tilt: float, canvas_rc: np.ndarray) -> np.ndarray:
        """
        (N, 2) canvas (row, col) -> (N, 2) aligned (row, col), the inverse of
        :meth:`aligned_to_canvas`.

        This direction *is* implicit -- ``v + L(v)`` has to be inverted for
        ``v`` -- so it is solved by fixed-point iteration. ``L`` is a smooth
        Gaussian blend of tens of pixels over a field of thousands, so the
        iteration is a strong contraction and converges to well under a tenth
        of a pixel in a handful of steps; the loop asserts that rather than
        assuming it.
        """
        sec = self.tilts[tilt].sec
        M, t = alignment_matrix(sec, self.canvas_shape, self.conv, self.out_bin)
        canvas_rc = np.atleast_2d(np.asarray(canvas_rc, dtype=float)) - self.extra_shift
        if self.local is not None:
            centre = np.array(self.canvas_shape, dtype=float) / 2.0
            shift_rc = alignment_shift_rc(sec, self.conv)
            q = canvas_rc - centre + shift_rc  # = v + L(v)
            v = q.copy()
            for _ in range(_LOCAL_INVERSE_ITERS):
                shift_xy = self.local.shift_at(sec.sec, v[:, ::-1])
                new = q - shift_xy[:, ::-1]
                delta = float(np.abs(new - v).max()) if new.size else 0.0
                v = new
                if delta < 1e-3:
                    break
            else:
                logger.warning(
                    "tilt %+.1f: local-shift inversion still moving by %.2f px "
                    "after %d iterations.", tilt, delta, _LOCAL_INVERSE_ITERS,
                )
            canvas_rc = v + centre - shift_rc
        return canvas_rc @ M.T + t

    def tomo_to_canvas(self, tilt: float, xyz: np.ndarray) -> np.ndarray:
        """(N, 3) reconstruction voxels -> (N, 2) canvas (row, col)."""
        al = self.tomo_to_aligned(tilt, xyz)
        return self.aligned_to_canvas(tilt, al[:, :2])

    def canvas_to_raw_tile(self, local_rc: np.ndarray) -> np.ndarray:
        """
        Tile-local canvas pixels -> raw (unbinned, unrotated) tile pixels.

        ``stitch.py`` applies ``np.rot90(tile, k)`` to the *binned* tile, and
        ``np.rot90(A, 1)[i, j] == A[j, W-1-i]`` for A of shape (H, W). A binned
        pixel b covers raw pixels [b*bin, b*bin+bin-1], whose centre is at
        b*bin + (bin-1)/2.
        """
        local_rc = np.atleast_2d(np.asarray(local_rc, dtype=float))
        i, j = local_rc[:, 0], local_rc[:, 1]
        h, w = self.geom.tile_shape  # the tile AS PLACED, i.e. after rotation
        k = (self.rotate // 90) % 4
        if k == 0:
            rb, cb = i, j
        elif k == 1:
            rb, cb = j, (h - 1) - i
        elif k == 2:
            rb, cb = (h - 1) - i, (w - 1) - j
        else:
            rb, cb = (w - 1) - j, i
        off = (self.binning - 1) / 2.0
        return np.stack([self.binning * rb + off, self.binning * cb + off], axis=1)

    # -- tile assignment ----------------------------------------------------

    def assign(
        self,
        tilt: float,
        xyz: np.ndarray,
        margin: float = 0.0,
        require_coverage: bool = True,
    ) -> List[Optional[TileHit]]:
        """
        Assign each 3D point to at most one tile of this tilt.

        A candidate tile is only accepted if the *whole extraction box* lies
        inside that tile's illuminated square -- not merely inside the tile
        rectangle. The beam is smaller than the frame and offset within it, so
        the rectangle test lets the beam edge cut through ~15% of boxes, and a
        beam edge inside a particle box is worse than no observation at all.
        Among the tiles that pass, we keep the one holding the point furthest
        from its beam edge.

        ``index_map`` is consulted only as a coverage test: it is a partition,
        so its winner at a pixel is not necessarily the roomiest tile, but a
        fill value there does mean no tile contributed real data.

        Returns one entry per input point, ``None`` where the point is not
        usable in this tilt.
        """
        info = self.tilts[tilt]
        canvas = self.tomo_to_canvas(tilt, xyz)
        al = self.tomo_to_aligned(tilt, xyz)
        org = self.geom.tile_origin_rc(info.positions_um)
        h, w = self.geom.tile_shape
        sel = info.selected
        imap = info.index_map() if require_coverage else None
        fill = int(imap.max()) if imap is not None else None

        beams = info.beam_masks()
        if beams is None:
            logger.warning(
                "tilt %+.1f has no beam_masks in its HDF5 (pre-dating beam-mask "
                "storage); falling back to the tile rectangle, which does not "
                "keep the beam edge out of the box.", tilt,
            )
        half = int(np.ceil(margin / self.binning))  # box half-width, canvas px
        margin_canvas = margin / self.binning
        out: List[Optional[TileHit]] = []
        for n in range(canvas.shape[0]):
            rc = canvas[n]
            if imap is not None:
                ri, ci = int(round(rc[0])), int(round(rc[1]))
                if not (0 <= ri < imap.shape[0] and 0 <= ci < imap.shape[1]):
                    out.append(None)
                    continue
                if int(imap[ri, ci]) >= fill:
                    out.append(None)  # no tile contributed data here
                    continue
            best, best_margin = None, -np.inf
            for label, tile in enumerate(sel):
                local = rc - org[tile]
                if not (0 <= local[0] < h and 0 <= local[1] < w):
                    continue
                m = min(local[0], h - 1 - local[0], local[1], w - 1 - local[1])
                if m < margin_canvas:
                    continue
                if beams is not None:
                    r0, c0 = int(round(local[0])) - half, int(round(local[1])) - half
                    if r0 < 0 or c0 < 0 or r0 + 2 * half > h or c0 + 2 * half > w:
                        continue
                    if not beams[label, r0:r0 + 2 * half, c0:c0 + 2 * half].all():
                        continue  # the beam edge crosses this box
                if m > best_margin:
                    best, best_margin = (tile, local), m
            if best is None:
                out.append(None)
                continue
            tile, local = best
            raw = self.canvas_to_raw_tile(local)[0]
            out.append(
                TileHit(
                    tilt=tilt, tile=int(tile), x=float(raw[1]), y=float(raw[0]),
                    canvas_rc=(float(rc[0]), float(rc[1])),
                    depth=float(al[n, 2]) * self.tomo_bin,
                    edge_margin=float(best_margin) * self.binning,
                )
            )
        return out

    # -- RELION projection matrix ------------------------------------------

    def projection_matrix(self, tilt: float, tile: int) -> np.ndarray:
        """
        The 4x4 RELION tomogram projection matrix for one (tilt, tile) image.

        Maps a 3D position in *unbinned tilt-series pixels* with the origin at
        the tomogram corner -- which is what RELION reconstructs from
        ``rlnCenteredCoordinate*Angst`` plus ``rlnTomoSize*`` -- to
        ``(x, y, depth)`` in raw tile pixels.

        Every link in the chain is affine, so the matrix is recovered exactly by
        evaluating the chain on the origin and the three unit vectors. That is
        deliberate: composing the rotations, the inverse alignment matrix, the
        rot90 and the binning offsets by hand is four more chances to drop a
        sign, and this cannot disagree with the mapping ``assign()`` uses.
        """
        info = self.tilts[tilt]
        org = self.geom.tile_origin_rc(info.positions_um)[tile]

        def f(p: np.ndarray) -> np.ndarray:
            xyz = np.asarray(p, dtype=float)[None, :] / self.tomo_bin
            al = self.tomo_to_aligned(tilt, xyz)
            rc = self.aligned_to_canvas(tilt, al[:, :2])[0] - org
            raw = self.canvas_to_raw_tile(rc)[0]
            return np.array([raw[1], raw[0], al[0, 2] * self.tomo_bin])

        origin = f(np.zeros(3))
        P = np.zeros((4, 4))
        P[3, 3] = 1.0
        for k in range(3):
            e = np.zeros(3)
            e[k] = 1.0
            P[:3, k] = f(e) - origin
        P[:3, 3] = origin
        return P

    def tomo_size_unbinned(self) -> Tuple[int, int, int]:
        """(X, Y, Z) size of the tomogram in unbinned tilt-series pixels."""
        nz, ny, nx = self.recon_shape
        return (nx * self.tomo_bin, ny * self.tomo_bin, nz * self.tomo_bin)

    def centred_angst(self, xyz: np.ndarray) -> np.ndarray:
        """
        Reconstruction voxels -> ``rlnCenteredCoordinate{X,Y,Z}Angst``.

        RELION undoes this as ``coord_px = centred/apix + rlnTomoSize/2``, so
        as long as ``rlnTomoSize`` is ``recon_shape * tomo_bin`` the size/2 vs
        (size-1)/2 choice cancels and the round trip is exact.
        """
        xyz = np.atleast_2d(np.asarray(xyz, dtype=float))
        nz, ny, nx = self.recon_shape
        centre = np.array([nx, ny, nz], dtype=float) / 2.0
        return (xyz - centre) * (self.pixel_size * self.tomo_bin)


# ---------------------------------------------------------------------------
# Helpers
# ---------------------------------------------------------------------------


def add_geometry_args(parser) -> None:
    """
    Add the canvas-geometry arguments every reprojection tool shares.

    Four tools had four private copies of this block, all with the same
    hard-coded ROI and pixel size in their defaults. That is four places for a
    dataset's geometry to be restated and one of them to go stale. The
    arguments stay, because overriding one for a test is useful, but the
    provenance sidecars should be the normal route: they read what the steps
    actually did instead of asking the caller to remember it. They are read
    automatically -- the tomogram names its own chain -- so there is no flag
    to remember either; ``--no-sidecars`` opts out.
    """
    g = parser.add_argument_group("geometry")
    g.add_argument("--no-sidecars", dest="sidecars", action="store_false",
                   default=True,
                   help="do not read geometry from the tomogram's provenance "
                        "sidecars. They supply the ROI, pixel size, binning, "
                        "OutBin, the stitch rotation and z_sign, and say "
                        "whether the tomogram had the patch alignment applied; "
                        "without them every one of those falls back to a "
                        "hard-coded default that belongs to another dataset. "
                        "Explicitly given values below win either way")
    g.add_argument("--roi", nargs=4, type=float,
                   default=[-5500, 12000, -6000, 12000])
    g.add_argument("--pixel-size", type=float, default=3.426)
    g.add_argument("--binning", type=int, default=2)
    g.add_argument("--out-bin", type=int, default=2)
    g.add_argument("--rotate", default="auto",
                   help="stitch.py --rotate; 'auto' measures it from index_map, "
                        "which cannot tell k from k+2 -- prefer the sidecar")
    g.add_argument("--extra-shift", nargs=2, type=float, default=[0.0, 0.0])
    g.add_argument("--handedness", type=int, choices=[1, -1], default=1)
    g.add_argument("--no-local", dest="use_local", action="store_false",
                   default=True,
                   help="ignore the .aln's patch alignment. Only correct when "
                        "the tomogram was reconstructed without one; otherwise "
                        "it reintroduces the error the patch field removes")

    # Link 1 of the chain. The three offsets are NOT in the sidecars and never
    # will be: they are corrections to what the reconstruction actually did,
    # measured by measure_reprojection_residual --frame aligned, which prints
    # the exact flags to paste here. Leave them at zero until a fit says
    # otherwise. --z-sign is the opposite kind of thing -- a fact about the
    # file, which the sidecars do carry -- and is here only to override it.
    a = parser.add_argument_group("tomogram -> aligned frame")
    a.add_argument("--z-sign", type=int, choices=[1, -1], default=-1,
                   help="direction of the volume's Z axis. -1 (default) is an "
                        "AreTomo volume with exactly one depth flip applied, "
                        "by `clip rotx` or by -FlipVol -- what every volume "
                        "this pipeline has made so far needs; +1 is one with "
                        "both or neither. Normally comes from a sidecar; a "
                        "fitted depth scale factor of -1 is what says to "
                        "change it")
    a.add_argument("--z-centre-offset", type=float, default=0.0,
                   help="recon voxels added to cz = (nz-1)/2, the depth the "
                        "shear turns about. Apply the fitted sin(theta) "
                        "coefficient here, sign included")
    a.add_argument("--x-centre-offset", type=float, default=0.0,
                   help="recon voxels added to cx = (nx-1)/2. Apply MINUS the "
                        "fitted cos(theta)-1 coefficient")
    a.add_argument("--theta-offset", type=float, default=0.0,
                   help="degrees added to the .aln TILT column. Apply MINUS the "
                        "fitted dx sin(theta) coefficient, in degrees")


def build_projector(args, recon_shape: Sequence[int], **extra) -> "MontageProjector":
    """Construct a :class:`MontageProjector` from :func:`add_geometry_args`.

    By default the geometry comes from the tomogram's own provenance sidecars
    and only the arguments the caller actually typed override it. With
    ``--no-sidecars`` -- or a tomogram that has none -- this is the old
    behaviour exactly, defaults and all.
    """
    import sys

    rotate = args.rotate if args.rotate == "auto" else int(args.rotate)
    common = dict(
        tile_dir=getattr(args, "tile_dir", None),
        extra_shift=args.extra_shift,
        handedness=args.handedness,
        # Corrections to link 1, so they apply with or without sidecars --
        # sidecars record what the steps did, not where they were wrong.
        z_centre_offset=getattr(args, "z_centre_offset", 0.0),
        x_centre_offset=getattr(args, "x_centre_offset", 0.0),
        theta_offset_deg=getattr(args, "theta_offset", 0.0),
        **extra,
    )
    tomogram = getattr(args, "tomogram", None)
    if not getattr(args, "sidecars", True) or not tomogram:
        return MontageProjector(
            aln_path=args.aln, positions_dir=args.positions_dir,
            recon_shape=recon_shape, roi=args.roi, pixel_size=args.pixel_size,
            binning=args.binning, out_bin=args.out_bin, rotate=rotate,
            use_local=args.use_local, z_sign=getattr(args, "z_sign", -1),
            **common,
        )

    # Only forward what was typed. Passing the parser's defaults through as
    # "overrides" would silently beat the sidecar with the very hard-coded
    # numbers this is meant to replace -- and `use_local` in particular must
    # stay absent unless asked for, because from_sidecars turns it off by
    # itself when the record says the tomogram never had the field applied.
    typed = " ".join(sys.argv)
    overrides = {}
    for flag, name, value in (("--roi", "roi", args.roi),
                              ("--pixel-size", "pixel_size", args.pixel_size),
                              ("--binning", "binning", args.binning),
                              ("--out-bin", "out_bin", args.out_bin)):
        if flag in typed:
            overrides[name] = value
    if "--rotate" in typed:
        overrides["rotate"] = rotate
    # Like --rotate, and unlike the link-1 offsets: the sidecar has an opinion
    # on this one, so passing the parser's default through would beat a
    # recorded -1 with a hard-coded +1.
    if "--z-sign" in typed:
        overrides["z_sign"] = args.z_sign
    if not args.use_local:
        overrides["use_local"] = False
    if overrides:
        logger.warning("Overriding the sidecars with command-line %s",
                       ", ".join(sorted(overrides)))
    return MontageProjector.from_sidecars(
        tomogram, aln_path=args.aln, positions_dir=args.positions_dir,
        recon_shape=recon_shape,
        canvas_path=getattr(args, "canvas_stack", None),
        **overrides, **common,
    )


def _sidecar_effective(sidecar, path: str, step: str) -> dict:
    """``effective`` from ``path``'s sidecar, if it is the step we expected.

    Returns an empty dict rather than raising when there is no sidecar, so a
    series that predates them still runs on the argument defaults -- loudly,
    via the "missing" warning in :meth:`MontageProjector.from_sidecars`,
    rather than by silently substituting another dataset's geometry.
    """
    if not path or not os.path.exists(path):
        return {}
    doc = sidecar.read_for(path)
    if doc is None:
        return {}
    if doc.get("step") != step:
        logger.warning("%s carries a %r sidecar, not %r.",
                       os.path.basename(path), doc.get("step"), step)
        return {}
    if doc.get("reconstructed"):
        logger.info(
            "%s's %s sidecar was RECONSTRUCTED by backfill, not written by "
            "the run itself — its parameters are inferred from surviving "
            "evidence. Treat a disagreement with the data as the sidecar's "
            "fault first.", os.path.basename(path), step)
    return doc.get("effective") or {}


def _find_tile_stacks(directory: str) -> Dict[float, str]:
    """Map tilt angle -> per-tilt tile stack, e.g. Montage_9-A_9-A_-12.mrc."""
    from .refine_montage_projmatch import tilt_from_filename

    out: Dict[float, str] = {}
    for name in sorted(os.listdir(directory)):
        if not name.endswith(".mrc") or "_refined_positions" in name:
            continue
        try:
            out[tilt_from_filename(name)] = os.path.join(directory, name)
        except ValueError:
            continue
    if not out:
        raise FileNotFoundError(f"No per-tilt tile stacks found in {directory}")
    return out


def _probe_tile_shape(stacks: Dict[float, str]) -> Tuple[int, int]:
    import mrcfile

    if not stacks:
        raise ValueError("Pass --tile-dir or --tile-shape so the tile size is known.")
    path = stacks[sorted(stacks)[0]]
    with mrcfile.open(path, header_only=True, permissive=True) as m:
        return (int(m.header.ny), int(m.header.nx))


def load_picks(path: str) -> np.ndarray:
    """
    Read an IMOD ``model2point`` text file into an (N, 3) array of (X, Y, Z).

    Accepts 3-column ``x y z`` and the 4/5-column forms that carry object and
    contour numbers, by taking the last three columns.
    """
    rows = []
    with open(path) as fh:
        for line in fh:
            line = line.strip()
            if not line or line.startswith("#"):
                continue
            parts = line.split()
            if len(parts) < 3:
                continue
            rows.append([float(x) for x in parts[-3:]])
    if not rows:
        raise ValueError(f"No coordinates parsed from {path}")
    return np.asarray(rows, dtype=float)


def stack_slice_index(info: TiltInfo, tile: int) -> int:
    """
    z-slice of the per-tilt tile stack holding tile ``tile``.

    ``Refined_positions`` is full length and the stack has one slice per row, so
    this is the identity -- but it is a real assumption (it fails on stacks
    written before the tile-index re-basing fix, see fix_stack_tile_index.py) and
    is worth naming rather than inlining.
    """
    return int(tile)
```
