## Supplementary material for "Whole cryo-FIB lamella acquisition at high magnification with rectangular beam montage cryo-electron tomography": Source code: montage_to_relion.py

```
#!/usr/bin/env python3
"""
Export a square-beam montage tilt series and a set of 3D picks as a RELION-5
tomogram + particle set, ready for ``relion_tomo_subtomo``.

The idea
--------
RELION assumes one tilt image per tilt. A montage has ~20, one per tile, and a
given particle appears in exactly one of them. Rather than stitch anything or
reimplement the pseudo-subtomogram maths (Zivanov et al. 2022, eqs 8-10), we
declare **one RELION "tilt image" per (tilt, tile)** -- up to ~650 per
tomogram, though only the ones a kept particle actually lands in are written
(see the pruning step in ``main``). Two RELION-5 features make that work
as-is:

* ``rlnTomoProjX/Y/Z/W`` is a general 4x4 affine from tomogram coordinates to
  image coordinates, so a tile's offset within the montage is just part of the
  matrix. Nothing has to know that the images are tiles.
* ``rlnTomoVisibleFrames`` marks, per particle, which of those images actually
  contain it. A particle sits in ~1/20 of the images per tilt and RELION
  already expects to be told this.

Defocus needs one extra step. RELION derives each particle's defocus from its
3D position through the ``rlnTomoProjZ`` row, adding a ramp across the tomogram
that at 45 deg is microns deep. But this acquisition scheme *already removed*
that ramp on the microscope -- ``MontageTiltSeries.txt`` calls ``ChangeFocus``
with a per-tile ``tan(alpha) * y`` offset before every Record -- so every tile
of a tilt was taken at the same defocus at its own field centre. Each image is
therefore given ``rlnDefocusU/V = df0(tilt) - depth(tile centre)``, which
cancels RELION's ramp back out at the tile centre and leaves it doing the right
thing *within* a tile. See ``fit_defocus``.

What it writes
--------------
    <out>/tomograms.star                 data_global, one row per tomogram
    <out>/tilt_series/<name>.star        one row per (tilt, tile)
    <out>/particles.star                 data_optics + data_particles
    <out>/optimisation_set.star          ties the two together
    <out>/defocus_model.tsv              measured vs modelled defocus per tile

Then::

    relion_tomo_subtomo --i <out>/optimisation_set.star --b 256 --crop 128 \\
        --bin 2 --j 8 --o Subtomograms/

Sanity first
------------
Run ``overlay_picks_on_montage`` before trusting any of this. The coordinate
chain has four sign conventions in it and a wrong one produces a perfectly
well-formed star file full of noise.
"""

from __future__ import annotations

import argparse
import logging
import os
import re
from dataclasses import dataclass
from typing import Dict, List, Optional, Sequence, Tuple

import numpy as np

from .montage_projection import (
    TileHit, add_geometry_args, build_projector, load_picks)

logger = logging.getLogger(__name__)

_CTF_STEM_RE = re.compile(r"^(?P<base>.+)_(?P<idx>\d+)_(?P<tilt>-?\d+\.?\d*)$")

STAR_VERSION = "# version 50001"


# ---------------------------------------------------------------------------
# STAR writing
# ---------------------------------------------------------------------------


def write_block(fh, name: str, columns: Sequence[str], rows: Sequence[Sequence]) -> None:
    """Write one ``data_<name>`` loop block."""
    fh.write(f"\ndata_{name}\n\nloop_\n")
    for i, col in enumerate(columns, start=1):
        fh.write(f"_{col} #{i}\n")
    for row in rows:
        fh.write(" ".join(_fmt(v) for v in row) + "\n")
    fh.write("\n")


def _fmt(v) -> str:
    if isinstance(v, float):
        return f"{v:.6f}"
    return str(v)


def _mat_str(v: np.ndarray) -> str:
    """RELION's bracketed vector literal, e.g. ``[1.0,0.0,0.0,0.0]``."""
    return "[" + ",".join(f"{x:.8g}" for x in v) + "]"


# ---------------------------------------------------------------------------
# CTF
# ---------------------------------------------------------------------------


@dataclass
class CtfFit:
    tilt: float
    slice_index: int  # z-slice of the per-tilt stack
    df1: float
    df2: float
    azimuth: float
    score: float
    fit_res: float

    @property
    def mean_defocus(self) -> float:
        return 0.5 * (self.df1 + self.df2)


def parse_ctf_results(path: str, base: Optional[str] = None) -> Dict[float, List[CtfFit]]:
    """
    Read ``beam_mask_motioncorr``'s ctf_results.txt into per-tilt fits.

    The ``{tile_index}`` in the tile names is a cumulative counter across the
    whole tilt series, not a per-tilt index (the same convention that
    fix_stack_tile_index.py exists to repair), so it is re-based per tilt here.
    Rows whose base name does not match ``base`` are ignored -- the file can
    contain more than one series.
    """
    fits: Dict[float, List[CtfFit]] = {}
    with open(path) as fh:
        for line in fh:
            # Ignore comments
            if line.startswith("#") or not line.strip():
                continue
            parts = line.split()
            # lines not containing 7 items are skipped
            if len(parts) < 7:
                continue
            m = _CTF_STEM_RE.match(parts[0])
            if m is None or (base is not None and m.group("base") != base):
                continue
            tilt = float(m.group("tilt"))
            # Read items df1, df2 (defocii in x and y), azimuth, score and fit resolution into object
            fits.setdefault(tilt, []).append(
                CtfFit(
                    tilt=tilt, slice_index=int(m.group("idx")),
                    df1=float(parts[1]), df2=float(parts[2]), azimuth=float(parts[3]),
                    score=float(parts[5]), fit_res=float(parts[6]),
                )
            )
    for tilt, group in fits.items():
        base_idx = min(f.slice_index for f in group)
        for f in group:
            f.slice_index -= base_idx
    return fits


def tile_reference_depth(proj: MontageProjector, tilt: float, tile: int) -> float:
    """
    Along-beam offset, in Angstroms, of the tile's own field centre.

    The point taken is the one on the tomogram mid-plane that projects to the
    centre of this tile's image. That is the position CTFFIND's single defocus
    estimate for the tile actually refers to, so subtracting this from the
    measured defocus puts every tile of a tilt on a common footing.
    """
    P = proj.projection_matrix(tilt, tile)
    ny_t, nx_t = proj.raw_tile_shape
    z = proj.recon_shape[0] / 2.0 * proj.tomo_bin
    A = P[:2, :2]
    b = np.array([nx_t / 2.0, ny_t / 2.0]) - P[:2, 2] * z - P[:2, 3]
    xy = np.linalg.solve(A, b)
    depth_px = P[2, 0] * xy[0] + P[2, 1] * xy[1] + P[2, 2] * z + P[2, 3]
    return float(depth_px) * proj.pixel_size


def _clipped_mean(values: Sequence[float], n_sigma: float = 3.0) -> float:
    """Mean after dropping points more than ``n_sigma`` MAD from the median."""
    v = np.asarray(values, dtype=float)
    if v.size < 3:
        return float(v.mean())
    med = np.median(v)
    mad = float(np.median(np.abs(v - med))) * 1.4826
    if mad <= 0:
        return float(v.mean())
    keep = np.abs(v - med) <= n_sigma * mad
    return float(v[keep].mean()) if keep.any() else float(med)


def fit_defocus(
    proj: MontageProjector,
    fits: Dict[float, List[CtfFit]],
    tilts: Sequence[float],
    min_score: float,
    max_fit_res: float,
    min_tiles: int = 3,
) -> Tuple[Dict[float, Tuple[float, float, float]], List[Sequence]]:
    """
    Collapse the per-tile CTF fits to one defocus per tilt, plus astigmatism.

    The tilt geometry puts a defocus ramp across a montage -- at 45 deg the
    tiles of one tilt span microns -- but the acquisition already takes it out.
    ``Setup_square_montage_from_polygon.calculate_defocii`` writes a per-tile
    ``tan(alpha) * y`` offset as the third column of the image-shift file, and
    ``MontageTiltSeries.txt`` applies it with ``ChangeFocus`` before each
    ``Record``. Every tile of a tilt was therefore exposed at the same defocus
    at its own field centre, and the estimator is simply the mean over the tiles
    whose fit is credible. No depth term.

    That was checked rather than assumed: regressing measured defocus on
    geometric depth within a tilt, over the 116 good tiles of Montage_9-A, gave
    a slope of -0.030 +- 0.110, consistent with 0 and 9 sigma from the +1 an
    uncompensated montage would show. It is not re-checked per export -- a
    montage acquired without the ``ChangeFocus`` compensation is obvious by eye
    and unusable anyway. ``defocus_model.tsv`` carries ``depth_A`` beside each
    tile's fit, so the regression can be repeated on any export that wants it.

    The depth term does not vanish, it moves: RELION reconstructs each
    particle's defocus from ``rlnTomoProjZ`` relative to the tomogram centre, so
    ``main`` writes ``df0 - depth(tile centre)`` per image to cancel that ramp
    back out. Returned here is ``{tilt: (dfU, dfV, angle)}`` *at zero depth*.

    Per-tile CTFFIND on this data is mostly noise -- one tile at ~3 e-/A^2 does
    not give enough Thon-ring signal and only 18% of tiles pass the cut -- so
    tilts with fewer than ``min_tiles`` credible fits are interpolated from
    their neighbours instead of believed. On Montage_9-A that matters: +30 deg
    has exactly one survivor, 1.3 um away from every other tilt's mean.
    """
    per_tilt: Dict[float, Tuple[float, float, float]] = {}
    df0_by_tilt: Dict[float, float] = {}
    good_by_tilt: Dict[float, List[CtfFit]] = {}
    depths: Dict[Tuple[float, int], float] = {}
    status: Dict[Tuple[float, int], str] = {}

    for tilt in tilts:
        info = proj.tilts[tilt]
        group = {f.slice_index: f for f in fits.get(tilt, [])}
        good: List[CtfFit] = []
        for tile in info.selected:
            t = int(tile)
            # Not used by the fit -- the acquisition already flattened the
            # ramp -- but written to defocus_model.tsv, and recomputed in main
            # to cancel the one RELION adds back.
            depths[(tilt, t)] = tile_reference_depth(proj, tilt, t)
            f = group.get(t)
            if f is None:
                status[(tilt, t)] = "nofit"
            elif f.score >= min_score and f.fit_res <= max_fit_res:
                status[(tilt, t)] = "good"
                good.append(f)
            else:
                status[(tilt, t)] = "rejected"
        good_by_tilt[tilt] = good

        if len(good) >= min_tiles:
            df0 = _clipped_mean([f.mean_defocus for f in good])
            a = float(np.median([f.df1 - f.df2 for f in good]))
            angles = np.deg2rad(2.0 * np.array([f.azimuth for f in good]))
            ang = float(np.rad2deg(np.angle(np.mean(np.exp(1j * angles)))) / 2.0)
            df0_by_tilt[tilt] = df0
            per_tilt[tilt] = (df0 + a / 2.0, df0 - a / 2.0, ang)
        else:
            logger.warning(
                "tilt %+6.1f: only %d credible CTF fits (need %d), interpolating",
                tilt, len(good), min_tiles,
            )

    if not df0_by_tilt:
        raise ValueError(
            "No tilt has %d tiles passing --ctf-min-score / --ctf-max-fit-res. "
            "Loosen them, lower --ctf-min-tiles, or supply --defocus directly."
            % min_tiles
        )

    # Fill the gaps by interpolating df0 against tilt angle.
    known = np.array(sorted(df0_by_tilt))
    vals = np.array([df0_by_tilt[t] for t in known])
    for tilt in tilts:
        if tilt not in per_tilt:
            df0 = float(np.interp(tilt, known, vals))
            per_tilt[tilt] = (df0, df0, 0.0)

    logger.info(
        "defocus per tilt: %.0f - %.0f A (%d of %d tilts measured, rest "
        "interpolated)",
        min(v[0] for v in per_tilt.values()), max(v[0] for v in per_tilt.values()),
        len(df0_by_tilt), len(tilts),
    )

    diag: List[Sequence] = []
    for (tilt, tile), st in sorted(status.items()):
        f = {g.slice_index: g for g in good_by_tilt[tilt]}.get(tile)
        if f is None:
            for g in fits.get(tilt, []):
                if g.slice_index == tile:
                    f = g
                    break
        depth_A = depths[(tilt, tile)]
        written = 0.5 * (per_tilt[tilt][0] + per_tilt[tilt][1]) - depth_A
        diag.append((
            f"{tilt:+.1f}", tile,
            f"{f.df1:.1f}" if f else "-", f"{f.df2:.1f}" if f else "-",
            f"{f.score:.4f}" if f else "-", f"{f.fit_res:.2f}" if f else "-",
            f"{depth_A:.1f}", f"{written:.1f}", st,
        ))
    return per_tilt, diag


# ---------------------------------------------------------------------------
# Acquisition order / dose
# ---------------------------------------------------------------------------


def acquisition_order(
    tilts: Sequence[float], mode: str, order_file: Optional[str], mtime_dir: Optional[str]
) -> Dict[float, int]:
    """Map tilt angle -> 0-based position in the acquisition sequence."""
    tilts = list(tilts)
    if mode == "file":
        if not order_file:
            raise ValueError("--acquisition-order file needs --order-file")
        seq = [float(x) for x in open(order_file).read().split()]
    elif mode == "mtime":
        if not mtime_dir:
            raise ValueError("--acquisition-order mtime needs --mtime-dir")
        from .refine_montage_projmatch import tilt_from_filename

        found = []
        for name in os.listdir(mtime_dir):
            if not name.endswith((".mrc", ".tif")):
                continue
            try:
                found.append((os.path.getmtime(os.path.join(mtime_dir, name)),
                              tilt_from_filename(name)))
            except ValueError:
                continue
        seq = [t for _, t in sorted(found)]
    else:
        logger.warning(
            "Acquisition order taken as tilt order. For a dose-symmetric scheme "
            "that is wrong, and every dose weight with it -- pass "
            "--acquisition-order mtime or --order-file if dose matters."
        )
        seq = sorted(tilts)

    order = {}
    for i, t in enumerate(seq):
        order.setdefault(float(t), i)
    missing = [t for t in tilts if t not in order]
    if missing:
        raise ValueError(f"No acquisition order for tilts {missing}")
    return order


# ---------------------------------------------------------------------------
# Main
# ---------------------------------------------------------------------------


def parse_commandline(argv=None):
    p = argparse.ArgumentParser(
        description=__doc__, formatter_class=argparse.RawDescriptionHelpFormatter
    )
    p.add_argument("--picks", required=True, help="model2point text file of 3D picks")
    p.add_argument("--tomogram", required=True, help="the reconstruction the picks were made in")
    p.add_argument("--aln", required=True, help="AreTomo .aln for that reconstruction")
    p.add_argument("--positions-dir", required=True, help="dir of *_refined_positions.h5")
    p.add_argument("--tile-dir", required=True, help="dir of per-tilt tile stacks")
    p.add_argument("-o", "--out", required=True, help="output directory")
    p.add_argument("--tomo-name", default=None, help="rlnTomoName (default: from --tomogram)")

    add_geometry_args(p)
    p.add_argument("--exclude-sections", type=int, nargs="*", default=[],
                   help="SEC indices to drop (e.g. sections with junk shifts)")

    g = p.add_argument_group("particles")
    g.add_argument("--box", type=int, default=256,
                   help="extraction box in unbinned tile px; a particle is only "
                        "declared visible where the box fits inside the tile")
    g.add_argument("--min-frames", type=int, default=1,
                   help="drop particles visible in fewer images than this")

    g = p.add_argument_group("CTF")
    g.add_argument("--ctf-results", default=None,
                   help="ctf_results.txt from beam_mask_motioncorr")
    g.add_argument("--ctf-base", default=None, help="restrict to this base name")
    g.add_argument("--ctf-min-score", type=float, default=0.03)
    g.add_argument("--ctf-max-fit-res", type=float, default=20.0)
    g.add_argument("--ctf-min-tiles", type=int, default=3,
                   help="a tilt needs this many credible fits to be believed; "
                        "below it, its defocus is interpolated from neighbours")
    g.add_argument("--defocus", type=float, default=None,
                   help="use this defocus (A) at the tile centre for every "
                        "tilt instead of fitting")
    g.add_argument("--voltage", type=float, default=300.0)
    g.add_argument("--cs", type=float, default=2.7)
    g.add_argument("--amp-contrast", type=float, default=0.07)

    g = p.add_argument_group("dose")
    g.add_argument("--dose-per-tilt", type=float, default=0.0,
                   help="e-/A^2 per exposure; each area is exposed once per tilt")
    g.add_argument("--acquisition-order", choices=["tilt", "mtime", "file"], default="tilt")
    g.add_argument("--order-file", default=None)
    g.add_argument("--mtime-dir", default=None)

    p.add_argument("--materialise", action="store_true",
                   help="write one flat .mrcs of every tile instead of "
                        "referencing slices of the existing per-tilt stacks")
    p.add_argument("--verbose", "-v", action="store_true")
    return p.parse_args(argv)


def main(argv=None) -> int:
    args = parse_commandline(argv)
    logging.basicConfig(
        level=logging.DEBUG if args.verbose else logging.INFO,
        format="%(levelname)s %(message)s",
    )
    import mrcfile

    # Get tomogram shape from header
    with mrcfile.open(args.tomogram, header_only=True, permissive=True) as m:
        recon_shape = (int(m.header.nz), int(m.header.ny), int(m.header.nx))
    logger.info("reconstruction %s = (nz, ny, nx)", recon_shape)

    # Build projector (projects particle coordinates from tomogram to tilt series)
    # from manifest 
    proj = build_projector(args, recon_shape,
                           exclude_sections=args.exclude_sections)

    picks = load_picks(args.picks)
    logger.info("%d picks from %s", len(picks), args.picks)
    tilts = sorted(proj.tilts)

    # ---- one image row per (tilt, tile), in a fixed order --------------
    images: List[Tuple[float, int]] = []
    for tilt in tilts:
        for tile in proj.tilts[tilt].selected:
            images.append((tilt, int(tile)))
    image_row = {key: i for i, key in enumerate(images)}
    logger.info("%d tilt images (%d tilts x ~%.1f tiles)",
                len(images), len(tilts), len(images) / len(tilts))

    # ---- assign every pick in every tilt -------------------------------
    visible = np.zeros((len(picks), len(images)), dtype=np.int8)
    hits: Dict[Tuple[int, float], TileHit] = {}
    for tilt in tilts:
        assigned = proj.assign(tilt, picks, margin=args.box / 2.0)
        proj.tilts[tilt].drop_index_map()
        for n, hit in enumerate(assigned):
            if hit is None:
                continue
            key = (hit.tilt, hit.tile)
            if key not in image_row:
                continue
            visible[n, image_row[key]] = 1
            hits[(n, tilt)] = hit
        logger.info("tilt %+6.1f: %4d/%d picks land in a tile",
                    tilt, sum(h is not None for h in assigned), len(picks))

    n_seen = visible.sum(axis=1)
    keep = np.flatnonzero(n_seen >= args.min_frames)
    logger.info(
        "particles visible in %d-%d images (median %d); keeping %d/%d",
        n_seen.min(), n_seen.max(), int(np.median(n_seen)), len(keep), len(picks),
    )
    if len(keep) == 0:
        logger.error(
            "No particle is visible anywhere. That is a geometry failure, not a "
            "thin dataset -- run overlay_picks_on_montage before going further."
        )
        return 1

    # ---- prune to images at least one kept particle actually uses -------
    #
    # `images` above is every selected tile of every tilt, regardless of
    # whether a particle lands in it -- on Montage_9-A that is 641 rows for
    # 290 kept particles, of which only 148 are ever visible to one of them.
    # relion_tomo_subtomo loads every image its tilt series star references
    # before extracting a single particle (slurm_templates/
    # 12_subtomo_validate.sh's own memory note is written on exactly that
    # row count), so writing the unused rows costs real memory downstream for
    # nothing. `keep` only, not all picks: a dropped particle's visibility
    # should not keep an otherwise-unused image alive.
    used = np.flatnonzero(visible[keep].any(axis=0))
    n_images_full = len(images)
    images = [images[i] for i in used]
    visible = visible[:, used]
    logger.info(
        "%d/%d tilt images are visible to a kept particle; pruning the rest "
        "from the export", len(images), n_images_full,
    )

    # ---- CTF ------------------------------------------------------------
    if args.defocus is not None:
        per_tilt = {t: (args.defocus, args.defocus, 0.0) for t in tilts}
        diag = []
    elif args.ctf_results:
        fits = parse_ctf_results(args.ctf_results, args.ctf_base)
        per_tilt, diag = fit_defocus(
            proj, fits, tilts, args.ctf_min_score, args.ctf_max_fit_res,
            args.ctf_min_tiles,
        )
    else:
        raise SystemExit("Pass --ctf-results or --defocus.")

    # Order over *every* tilt, not just the ones that survived into `tilts`.
    # Dose accumulates during acquisition whether or not AreTomo later called a
    # frame dark, so counting position among the survivors under-doses every
    # tilt that follows a dropped one.
    order = acquisition_order(
        proj.all_tilts, args.acquisition_order, args.order_file, args.mtime_dir
    )
    if args.dose_per_tilt <= 0:
        logger.warning(
            "--dose-per-tilt is 0, so rlnMicrographPreExposure is 0 everywhere "
            "and RELION will not dose-weight. Supply the real dose per exposure."
        )

    # ---- write ----------------------------------------------------------
    out = os.path.abspath(args.out)
    os.makedirs(os.path.join(out, "tilt_series"), exist_ok=True)
    name = args.tomo_name or os.path.splitext(os.path.basename(args.tomogram))[0]

    micrograph_names = _image_sources(proj, images, out, name, args.materialise)

    ts_path = os.path.join(out, "tilt_series", f"{name}.star")
    rows = []
    mats, ref_depths = [], []
    for i, (tilt, tile) in enumerate(images):
        P = proj.projection_matrix(tilt, tile)
        # RELION adds each particle's own ProjZ depth, measured from the
        # *tomogram* centre, to the value written here. The microscope focused
        # on the *tile* centre (ChangeFocus per tile in MontageTiltSeries.txt),
        # so subtracting that tile's depth turns RELION's ramp into one that is
        # zero at the tile centre -- which is what was actually acquired.
        depth_A = tile_reference_depth(proj, tilt, tile)
        mats.append(P)
        ref_depths.append(depth_A)
        dfu, dfv, dfa = per_tilt[tilt]
        dfu, dfv = dfu - depth_A, dfv - depth_A
        rows.append([
            _mat_str(P[0]), _mat_str(P[1]), _mat_str(P[2]), _mat_str(P[3]),
            dfu, dfv, dfa,
            float(np.cos(np.deg2rad(tilt))),
            float(order[tilt] * args.dose_per_tilt),
            float(tilt),
            micrograph_names[i],
        ])
    with open(ts_path, "w") as fh:
        fh.write(STAR_VERSION + "\n")
        write_block(fh, name, [
            "rlnTomoProjX", "rlnTomoProjY", "rlnTomoProjZ", "rlnTomoProjW",
            "rlnDefocusU", "rlnDefocusV", "rlnDefocusAngle", "rlnCtfScalefactor",
            "rlnMicrographPreExposure", "rlnTomoNominalStageTiltAngle",
            "rlnMicrographName",
        ], rows)

    _report_particle_defocus(proj, per_tilt, images, mats, ref_depths,
                             picks, visible, keep)

    sx, sy, sz = proj.tomo_size_unbinned()
    tomo_path = os.path.join(out, "tomograms.star")
    with open(tomo_path, "w") as fh:
        fh.write(STAR_VERSION + "\n")
        write_block(fh, "global", [
            "rlnTomoName", "rlnVoltage", "rlnSphericalAberration",
            "rlnAmplitudeContrast", "rlnMicrographOriginalPixelSize", "rlnTomoHand",
            "rlnOpticsGroupName", "rlnTomoTiltSeriesPixelSize",
            "rlnTomoTiltSeriesStarFile", "rlnTomoTomogramBinning",
            "rlnTomoSizeX", "rlnTomoSizeY", "rlnTomoSizeZ",
            "rlnTomoReconstructedTomogram",
        ], [[
            name, args.voltage, args.cs, args.amp_contrast, proj.pixel_size,
            args.handedness, "optics1", proj.pixel_size,
            os.path.relpath(ts_path, out), float(proj.tomo_bin),
            sx, sy, sz, os.path.abspath(args.tomogram),
        ]])

    centred = proj.centred_angst(picks)
    prows = []
    for j, n in enumerate(keep, start=1):
        vis = "[" + ",".join(str(int(v)) for v in visible[n]) + "]"
        prows.append([
            name, float(centred[n, 0]), float(centred[n, 1]), float(centred[n, 2]),
            f"{name}/{j}", 1, 0.0, 0.0, 0.0, vis,
        ])
    part_path = os.path.join(out, "particles.star")
    with open(part_path, "w") as fh:
        fh.write(STAR_VERSION + "\n")
        write_block(fh, "optics", [
            "rlnOpticsGroup", "rlnOpticsGroupName", "rlnSphericalAberration",
            "rlnVoltage", "rlnAmplitudeContrast", "rlnTomoTiltSeriesPixelSize",
        ], [[1, "optics1", args.cs, args.voltage, args.amp_contrast, proj.pixel_size]])
        write_block(fh, "particles", [
            "rlnTomoName", "rlnCenteredCoordinateXAngst", "rlnCenteredCoordinateYAngst",
            "rlnCenteredCoordinateZAngst", "rlnTomoParticleName", "rlnOpticsGroup",
            "rlnAngleRot", "rlnAngleTilt", "rlnAnglePsi", "rlnTomoVisibleFrames",
        ], prows)

    with open(os.path.join(out, "optimisation_set.star"), "w") as fh:
        fh.write(STAR_VERSION + "\n\ndata_\n\n")
        fh.write(f"_rlnTomoParticlesFile   {os.path.relpath(part_path, out)}\n")
        fh.write(f"_rlnTomoTomogramsFile   {os.path.relpath(tomo_path, out)}\n\n")

    if diag:
        with open(os.path.join(out, "defocus_model.tsv"), "w") as fh:
            fh.write("#tilt\ttile\tdf1\tdf2\tscore\tfit_res\tdepth_A\t"
                     "df_written\tstatus\n")
            for row in diag:
                fh.write("\t".join(str(x) for x in row) + "\n")

    logger.info("wrote %s", out)
    logger.info(
        "next: relion_tomo_subtomo --i %s --b %d --crop %d --bin 2 --j 8 --o Subtomograms/",
        os.path.join(out, "optimisation_set.star"), args.box, args.box // 2,
    )
    return 0


def _report_particle_defocus(
    proj: MontageProjector,
    per_tilt: Dict[float, Tuple[float, float, float]],
    images: Sequence[Tuple[float, int]],
    mats: Sequence[np.ndarray],
    ref_depths: Sequence[float],
    picks: np.ndarray,
    visible: np.ndarray,
    keep: np.ndarray,
) -> None:
    """
    Reproduce RELION's per-particle defocus and check it is sane.

    Worth doing because the per-image ``rlnDefocusU`` looks alarming on its own:
    at 60 deg a tile's field centre back-projects to a mid-plane point several
    microns outside the reconstructed box, so the written defocus can come out
    negative. That is harmless, and this is the check that shows why -- RELION
    adds the particle's own depth straight back, and the two differ only across
    the width of one tile. Anything outside a couple of microns of the fitted
    defocus means the depth term has the wrong sign or scale.
    """
    pos = np.asarray(picks, dtype=float) * proj.tomo_bin
    vals = []
    for n in keep:
        for i in np.flatnonzero(visible[n]):
            P = mats[i]
            depth = float(P[2, :3] @ pos[n] + P[2, 3]) * proj.pixel_size
            df0 = 0.5 * sum(per_tilt[images[i][0]][:2])
            vals.append(df0 - ref_depths[i] + depth)
    if not vals:
        return
    v = np.array(vals)
    logger.info(
        "per-particle defocus RELION will use: %.2f - %.2f um (median %.2f, "
        "sd %.2f) over %d particle-image pairs",
        v.min() / 1e4, v.max() / 1e4, np.median(v) / 1e4, v.std() / 1e4, v.size,
    )
    if v.min() < 0 or np.ptp(v) > 4e4:
        logger.warning(
            "  that spread is far wider than one tile can explain -- suspect "
            "the sign of --handedness or the tile depth term."
        )


def _image_sources(
    proj: MontageProjector,
    images: Sequence[Tuple[float, int]],
    out: str,
    name: str,
    materialise: bool,
) -> List[str]:
    """
    Produce the ``rlnMicrographName`` for each (tilt, tile) image.

    By default these are ``<slice>@<per-tilt stack>``, which costs nothing: the
    stacks that ``beam_mask_motioncorr --stack-output`` already wrote hold one
    tile per slice in exactly the order of ``Refined_positions``.

    They cannot be pointed at directly, though. RELION rejects ``N@file.mrc``
    with "stacks of images in MRC-format should have extension .mrcs; .mrc
    extensions are reserved for 3D maps" -- it decides by extension alone, and
    the per-tilt stacks are .mrc. The two formats are identical, so a symlink
    per tilt satisfies it for nothing; ``--materialise`` writes the same data
    out flat and costs ~60 GB.
    """
    if not materialise:
        linkdir = os.path.join(out, "tilt_stacks")
        os.makedirs(linkdir, exist_ok=True)
        links: Dict[float, str] = {}
        for tilt in sorted({t for t, _ in images}):
            stack = proj.tilts[tilt].stack_path
            if stack is None:
                raise ValueError(f"No tile stack found for tilt {tilt}")
            link = os.path.join(
                linkdir, os.path.splitext(os.path.basename(stack))[0] + ".mrcs"
            )
            if os.path.lexists(link):
                os.remove(link)
            os.symlink(os.path.abspath(stack), link)
            links[tilt] = link
        logger.info("symlinked %d tile stacks as .mrcs in %s", len(links), linkdir)
        return [f"{tile + 1}@{links[tilt]}" for tilt, tile in images]

    import mrcfile

    ny, nx = proj.raw_tile_shape
    dest = os.path.join(out, f"{name}_tiles.mrcs")
    logger.info("materialising %d tiles into %s (%.1f GB)",
                len(images), dest, len(images) * ny * nx * 4 / 1e9)
    with mrcfile.new_mmap(dest, (len(images), ny, nx), mrc_mode=2, overwrite=True) as dst:
        dst.voxel_size = proj.pixel_size
        for i, (tilt, tile) in enumerate(images):
            with mrcfile.mmap(proj.tilts[tilt].stack_path, mode="r", permissive=True) as src:
                dst.data[i] = src.data[tile]
    return [f"{i + 1}@{os.path.abspath(dest)}" for i in range(len(images))]


if __name__ == "__main__":
    raise SystemExit(main())
```
