## Supplementary material for "Whole cryo-FIB lamella acquisition at high magnification with rectangular beam montage cryo-electron tomography": Source code: mrc_to_gif.py

```
#!/usr/bin/env python3
"""
Convert an MRC tilt series stack to an animated GIF.

Usage:
    python mrc_to_gif.py -i tiltseries.mrc -o tiltseries.gif
    python mrc_to_gif.py -i tiltseries.mrc -o tiltseries.gif -s 0.25 --fps 10
    python mrc_to_gif.py -i tiltseries.mrc -o tiltseries.gif -s 512      # longest edge = 512 px
    python mrc_to_gif.py -i tiltseries.mrc -o tiltseries.gif --rawtlt tiltseries.rawtlt
    python mrc_to_gif.py -i tomogram.mrc -o tomogram.gif --depth-label
    python mrc_to_gif.py -i tiltseries.mrc -o tiltseries.gif --mp4  # also writes tiltseries.mp4
    python mrc_to_gif.py -i tiltseries.mrc -o tiltseries.gif -x 200,1800 -y 100,1500  # crop before scaling
    python mrc_to_gif.py -i tomogram.mrc -o tomogram.gif -z 10,50  # keep only frames 10-49
"""

import argparse
import os
import shutil
import subprocess
import sys
import tempfile
from pathlib import Path
import numpy as np
import mrcfile
from PIL import Image, ImageDraw, ImageFont


def parse_args():
    p = argparse.ArgumentParser(
        description="Read an MRC tilt series stack, downsample, and write an animated GIF."
    )
    p.add_argument("-i", "--input", required=True, help="Input MRC file.")
    p.add_argument("-o", "--output", default=None,
                   help="Output GIF file. Defaults to the input path with extension replaced by .gif.")
    p.add_argument(
        "-s", "--scale", default=0.25,
        help="Downscale factor (float fraction, e.g. 0.25) or target size for the "
             "longest edge in pixels (int > 1). Default: 0.25",
    )
    p.add_argument(
        "-x", "--xrange", default=None,
        help="Crop each frame horizontally to pixels x1,x2 (e.g. '200,1800'), "
             "in the input file's own pixel coordinates (i.e. before --bin, "
             "if given) — applied after slicing along --axis and before "
             "--scale. Default: no crop.",
    )
    p.add_argument(
        "-y", "--yrange", default=None,
        help="Crop each frame vertically to pixels y1,y2 (e.g. '100,1500'), "
             "in the input file's own pixel coordinates (i.e. before --bin, "
             "if given) — applied after slicing along --axis and before "
             "--scale. y1,y2 use an IMOD-style origin at the BOTTOM-left of "
             "the image (y=0 = bottom row, increasing upward), matching "
             "3dmod; the crop itself still applies y1,y2 as top-origin array "
             "row indices, unaffected by the vertical flip applied to the "
             "rendered output (see frame_to_pil). Default: no crop.",
    )

    p.add_argument(
        "-z", "--zrange", default=None,
        help="Crop the movie depth-wise to frames z1,z2 (e.g. '10,50'), i.e. "
             "keep only frames [z1, z2) along --axis, in the input file's own "
             "pixel/frame coordinates (i.e. before --bin, if given). Applied "
             "before --scale; --rawtlt/--depth-label labels stay anchored to "
             "the uncropped volume. Default: no crop.",
    )
    p.add_argument(
        "--fps", type=float, default=8,
        help="Frames per second in the output GIF. Default: 8",
    )
    p.add_argument(
        "--loop", type=int, default=0,
        help="Number of GIF loops (0 = loop forever). Default: 0",
    )
    p.add_argument(
        "--contrast", type=float, default=3.0,
        help="Clip data at +/-N standard deviations from the mean before scaling "
             "to 8-bit. Higher = more contrast, fewer saturated pixels. Default: 3",
    )
    p.add_argument(
        "--cmap", choices=["gray", "hot", "plasma"], default="gray",
        help="Colourmap to apply. Default: gray",
    )
    p.add_argument(
        "--stat-frames", type=int, default=5,
        help="Number of evenly-spaced frames used to estimate global mean/std "
             "for contrast normalisation (avoids loading the full stack). Default: 5",
    )
    p.add_argument(
        "--rawtlt", default=None,
        help="Path to a .rawtlt file (one tilt angle per line). When provided, "
             "each frame is labelled with its tilt angle in degrees.",
    )
    p.add_argument(
        "--depth-label", action="store_true",
        help="Label each frame with its position along the sliced axis in nm, "
             "for tomograms. The spacing comes from the MRC header (or "
             "--aperpix). Mutually exclusive with --rawtlt, which is the "
             "tilt-series equivalent.",
    )
    p.add_argument(
        "--depth-origin", choices=["center", "start"], default="center",
        help="Where zero sits for --depth-label: center=mid-plane of the "
             "volume, so labels run -N nm … +N nm; start=first frame. "
             "Default: center",
    )
    p.add_argument(
        "--label-size", type=int, default=None,
        help="Font size for the frame label in output pixels. "
             "Defaults to ~4%% of the output image height.",
    )
    p.add_argument(
        "--label-color", default="white",
        help="Label text colour (any PIL colour name or hex, e.g. 'white', '#ffff00'). "
             "Default: white",
    )
    p.add_argument(
        "--label-pos", choices=["tl", "tr", "bl", "br"], default="bl",
        help="Label position: tl=top-left, tr=top-right, bl=bottom-left, br=bottom-right. "
             "Default: bl",
    )
    p.add_argument(
        "--axis", choices=["z", "y", "x"], default="z",
        help="Axis to slice along. z=tilt series (default), y=tomogram Y slices, x=tomogram X slices.",
    )
    p.add_argument(
        "--bin", type=int, default=None, dest="bin_factor",
        help="Bin the volume by this integer factor using IMOD's binvol before processing. "
             "For --axis z (tilt series) bins XY only; for y/x bins all 3 dimensions. "
             "When set, --scale is still applied afterwards for any additional resize.",
    )
    p.add_argument(
        "--rotate", type=int, choices=[0, 90, 180, 270], default=0,
        help="Rotate each frame by this many degrees counter-clockwise. Default: 0",
    )
    p.add_argument(
        "--bounce", action="store_true",
        help="Play frames forward then backward (ping-pong) instead of looping from the start.",
    )
    p.add_argument(
        "--aperpix", type=float, default=None,
        help="Input pixel size in Angstroms/pixel, for the scale bar. Read "
             "from the MRC header when omitted; give it explicitly when the "
             "header says 0 (unset) or is known to be wrong.",
    )
    p.add_argument(
        "--scalebar", type=float, default=None,
        help="Scale bar length in nm. If --aperpix is given but --scalebar is omitted, "
             "a round value ~15%% of the image width is chosen automatically.",
    )
    p.add_argument(
        "--scalebar-pos", choices=["tl", "tr", "bl", "br"], default="br",
        help="Scale bar position. Default: br",
    )
    p.add_argument(
        "--scalebar-color", default="white",
        help="Scale bar and label colour. Default: white",
    )
    p.add_argument(
        "--mp4", action="store_true",
        help="Also convert the output GIF to an MP4 (H.264, via ffmpeg), "
             "written alongside it with the same basename.",
    )
    p.add_argument(
        "--tmpdir", default=None,
        help="Where to write the binned copy made by --bin. Defaults to "
             "$TMPDIR if set, otherwise the output file's directory — not "
             "/tmp, which is node-local, often small, and sometimes a RAM "
             "disk, while these volumes run to tens of GB.",
    )
    return p.parse_args()


def parse_range(value, name, limit):
    """Parse a '-x'/'-y' 'a,b' crop argument, or None if not given."""
    if value is None:
        return None
    parts = value.split(",")
    if len(parts) != 2:
        sys.exit(f"--{name} expects 'a,b' (e.g. '200,1800'), got {value!r}")
    try:
        a, b = (int(p) for p in parts)
    except ValueError:
        sys.exit(f"--{name} expects integer pixel indices, got {value!r}")
    if not (0 <= a < b <= limit):
        sys.exit(f"--{name} range {a},{b} out of bounds for frame size {limit}")
    return a, b


def compute_output_size(h, w, scale_arg):
    try:
        scale = float(scale_arg)
    except ValueError:
        sys.exit("Cannot parse --scale value: {0}".format(scale_arg))

    if 0 < scale <= 1.0:
        new_h = max(1, int(round(h * scale)))
        new_w = max(1, int(round(w * scale)))
    else:
        target = int(scale)
        ratio = target / max(h, w)
        new_h = max(1, int(round(h * ratio)))
        new_w = max(1, int(round(w * ratio)))

    return new_h, new_w


def load_tilt_angles(rawtlt_path, nframes):
    """Read tilt angles from a .rawtlt file.
    Accepts single-value lines (angle) or two-value lines (angle acquisition_order).
    """
    with open(rawtlt_path) as f:
        angles = [float(line.split()[0]) for line in f if line.strip()]
    if len(angles) != nframes:
        print(
            f"  Warning: {rawtlt_path} has {len(angles)} angles but MRC has "
            f"{nframes} frames. Angles will be truncated or padded with None."
        )
        angles = (angles + [None] * nframes)[:nframes]
    return angles


def _load_font(font_size):
    for font_path in [
        "/usr/share/fonts/liberation/LiberationSans-Bold.ttf",
        "/usr/share/fonts/truetype/liberation/LiberationSans-Bold.ttf",
        "/usr/share/fonts/truetype/dejavu/DejaVuSans-Bold.ttf",
        "/usr/share/fonts/dejavu/DejaVuSans-Bold.ttf",
    ]:
        try:
            return ImageFont.truetype(font_path, size=font_size)
        except (IOError, OSError):
            continue
    return ImageFont.load_default()


def _text_size(draw, text, font):
    try:
        bbox = draw.textbbox((0, 0), text, font=font)
        return bbox[2] - bbox[0], bbox[3] - bbox[1]
    except AttributeError:
        return draw.textsize(text, font=font)


def _color_for_mode(color, mode):
    """Convert a PIL color name/hex to the right fill type for the image mode."""
    from PIL import ImageColor
    if mode == "L":
        return ImageColor.getcolor(color, "L")
    return color  # RGB accepts strings directly


def _draw_shadowed_text(draw, xy, text, font, fg, shadow):
    x, y = xy
    for dx, dy in ((1, 1), (-1, -1), (1, -1), (-1, 1)):
        draw.text((x + dx, y + dy), text, font=font, fill=shadow)
    draw.text((x, y), text, font=font, fill=fg)


def nice_scalebar_nm(image_width_px, aperpix_output):
    """Pick a round scale bar length (nm) that is ~15% of the image width."""
    target_px = image_width_px * 0.15
    target_nm = target_px * aperpix_output / 10.0  # Å → nm
    # Round to a nice value: 1, 2, 5, 10, 20, 50, 100, 200, 500, …
    magnitude = 10 ** np.floor(np.log10(target_nm))
    for nice in [1, 2, 5, 10]:
        candidate = nice * magnitude
        if candidate >= target_nm * 0.5:
            return candidate
    return magnitude * 10


def add_scalebar(img, bar_nm, aperpix_output, font_size, color, position):
    """Draw a scale bar and label directly onto the image (L or RGB, in-place)."""
    bar_px = int(round(bar_nm * 10 / aperpix_output))  # nm→Å, then ÷ Å/px
    if bar_px < 2:
        return img

    label = f"{bar_nm:g} nm" if bar_nm < 1000 else f"{bar_nm/1000:g} µm"
    fg = _color_for_mode(color, img.mode)
    bg = _color_for_mode("black", img.mode)

    draw = ImageDraw.Draw(img)
    font = _load_font(font_size)
    tw, th = _text_size(draw, label, font)

    W, H = img.size
    margin = max(6, font_size // 3)
    bar_h = max(2, font_size // 5)

    block_w = max(bar_px, tw)
    x0 = margin if position in ("tl", "bl") else W - block_w - margin
    y0 = margin if position in ("tl", "tr") else H - (bar_h + 2 + th) - margin

    bar_x0 = x0 + (block_w - bar_px) // 2
    bar_x1 = bar_x0 + bar_px
    bar_y1 = y0 + bar_h

    for dx, dy in ((1, 1), (-1, -1), (1, -1), (-1, 1)):
        draw.rectangle([bar_x0 + dx, y0 + dy, bar_x1 + dx, bar_y1 + dy], fill=bg)
    draw.rectangle([bar_x0, y0, bar_x1, bar_y1], fill=fg)

    _draw_shadowed_text(draw, (x0 + (block_w - tw) // 2, y0 + bar_h + 2), label, font, fg, bg)
    return img


def add_label(img, text, font_size, color, position):
    """Draw a frame label directly onto the image (L or RGB, in-place)."""
    fg = _color_for_mode(color, img.mode)
    bg = _color_for_mode("black", img.mode)

    draw = ImageDraw.Draw(img)
    font = _load_font(font_size)
    tw, th = _text_size(draw, text, font)

    W, H = img.size
    margin = max(4, font_size // 4)
    x = margin if position in ("tl", "bl") else W - tw - margin
    y = margin if position in ("tl", "tr") else H - th - margin

    _draw_shadowed_text(draw, (x, y), text, font, fg, bg)
    return img


def to_gif_frame(img):
    """Convert L or RGB image to GIF palette mode without dithering."""
    if img.mode == "L":
        # Direct copy: pixel value n → palette index n → grey(n, n, n). Lossless.
        img_p = Image.frombytes("P", img.size, img.tobytes())
        img_p.putpalette([i for i in range(256) for _ in range(3)])
        return img_p
    if img.mode == "RGB":
        return img.quantize(colors=256, dither=0)
    return img  # already P


def pixel_size_from_header(mrc, axis):
    """Å/px along the *horizontal* axis of the displayed frame, or None.

    The scale bar is drawn horizontally, so the relevant number depends on
    which way the stack is being sliced: frames are (y, x) along z, (z, x)
    along y, and (z, y) along x. Anisotropic volumes are rare but a scale bar
    off by the z sampling is worse than no scale bar.

    An MRC whose header was never filled in reads back as 0, which every
    downstream tool then treats as 1 Å/px. That is precisely the failure that
    puts a scale bar off by a factor of a thousand, so 0 is reported as
    "absent" rather than believed.
    """
    voxel = mrc.voxel_size
    horizontal = {"z": voxel.x, "y": voxel.x, "x": voxel.y}[axis]
    value = float(horizontal)
    return value if value > 1e-6 else None


def depth_pixel_size_from_header(mrc, axis):
    """Å between successive frames along the sliced axis, or None.

    The complement of pixel_size_from_header: that one measures across a
    frame, for the scale bar, this one measures the step from one frame to
    the next, which is what a depth label counts. Slicing along y makes each
    step a step in y, and so on.

    As there, a header that was never filled in reads back as 0, so 0 is
    reported as absent rather than believed — a depth axis off by a factor of
    a thousand is worse than no depth label.
    """
    voxel = mrc.voxel_size
    step = {"z": voxel.z, "y": voxel.y, "x": voxel.x}[axis]
    value = float(step)
    return value if value > 1e-6 else None


def depth_labels(nframes, step_a, axis, origin):
    """Per-frame position labels, e.g. 'z = +12.3 nm'.

    Positions describe the input volume, not the output image, so they are
    unaffected by --scale: frame i is where it is regardless of how many
    pixels it gets drawn in.
    """
    step_nm = step_a / 10.0  # Å → nm
    zero = (nframes - 1) / 2.0 if origin == "center" else 0.0
    fmt = "{0} = {1:+.1f} nm" if origin == "center" else "{0} = {1:.1f} nm"
    return [fmt.format(axis, (i - zero) * step_nm) for i in range(nframes)]


def dims_for_axis(nz, ny, nx, axis):
    """(nframes, height, width) for the chosen slice axis, given raw z/y/x sizes."""
    if axis == "z":
        return nz, ny, nx
    if axis == "y":
        return ny, nz, nx
    return nx, nz, ny  # axis == "x"


def make_frame_getter(data, axis):
    """Return (nframes, height, width, get_frame_fn) for the chosen slice axis."""
    nz, ny, nx = data.shape
    nframes, h, w = dims_for_axis(nz, ny, nx, axis)
    if axis == "z":
        return nframes, h, w, lambda i: data[i]
    if axis == "y":
        return nframes, h, w, lambda i: data[:, i, :]
    # axis == "x"
    return nframes, h, w, lambda i: data[:, :, i]


def estimate_stats(nframes, get_frame, n_sample, n_std):
    """Sample a few evenly-spaced frames to estimate global clipping limits."""
    indices = np.linspace(0, nframes - 1, min(n_sample, nframes), dtype=int)
    sample = np.concatenate([get_frame(i).ravel() for i in indices]).astype(np.float32)
    mu, sigma = sample.mean(), sample.std()
    return float(mu - n_std * sigma), float(mu + n_std * sigma)


def frame_to_pil(frame, lo, hi, new_w, new_h, cmap_name):
    # Flipped vertically so row 0 of the array renders at the bottom,
    # matching IMOD/3dmod's bottom-left origin convention rather than the
    # top-left origin numpy arrays default to.
    frame = np.flipud(frame)
    f = np.clip(frame.astype(np.float32), lo, hi)
    u8 = ((f - lo) / (hi - lo) * 255).astype(np.uint8)
    img = Image.fromarray(u8, mode="L").resize((new_w, new_h), Image.LANCZOS)

    if cmap_name == "gray":
        return img  # stay in L mode — lossless, no palette quantization

    import matplotlib.cm as cm
    cmap = cm.get_cmap(cmap_name)
    rgba = cmap(np.array(img) / 255.0)
    rgb = (rgba[..., :3] * 255).astype(np.uint8)
    return Image.fromarray(rgb, mode="RGB")


def resolve_tmpdir(explicit, output_path):
    """Where to put the binned copy, which is a sizeable file.

    Not /tmp by default, which is what ``tempfile`` would choose. /tmp is
    node-local and often small — 64 GB on this cluster's login node, and under
    Slurm it is a per-job private mount — while these volumes run to tens or
    hundreds of GB. On plenty of other systems /tmp is a tmpfs, i.e. RAM, and
    filling it takes the node down rather than just the job.

    So: an explicit ``--tmpdir`` wins, then ``TMPDIR`` if the site or the
    scheduler set one, and failing both the output's own directory, which is
    on the same filesystem as the data and therefore sized for it.
    """
    if explicit:
        return explicit
    if os.environ.get("TMPDIR"):
        return os.environ["TMPDIR"]
    return str(Path(output_path).resolve().parent)


def run_binvol(input_path, bin_factor, axis, tmpdir=None):
    """Bin an MRC file using IMOD binvol; returns path to a temporary binned file."""
    if not shutil.which("binvol"):
        sys.exit("binvol not found in PATH — is IMOD installed and on PATH?")

    os.makedirs(tmpdir, exist_ok=True)
    # A dedicated directory with a name that does not exist yet, rather than
    # NamedTemporaryFile: binvol renames any existing output file to "<name>~"
    # before writing, so pre-creating the file leaves a stray backup behind
    # that nothing cleans up. Anything binvol does write lands inside this
    # directory, which is removed whole.
    workdir = tempfile.mkdtemp(dir=tmpdir, prefix="mrc_to_gif_")
    out_path = str(Path(workdir) / "binned.mrc")

    # Tilt series: bin XY only to preserve frame count; tomogram: full 3-D bin.
    #
    # There is no -binXY option in binvol and never has been in any IMOD
    # version. -binning sets the factor for all three axes and -zbinning
    # overrides Z, so "1" there leaves the frames alone.
    if axis == "z":
        cmd = ["binvol", "-binning", str(bin_factor), "-zbinning", "1",
               input_path, out_path]
    else:
        cmd = ["binvol", "-binning", str(bin_factor), input_path, out_path]

    print(f"  Running: {' '.join(cmd)}")
    result = subprocess.run(cmd, capture_output=True, text=True)
    if result.returncode != 0:
        shutil.rmtree(workdir, ignore_errors=True)
        sys.exit(f"binvol failed:\n{result.stderr or result.stdout}")

    return out_path


def convert_gif_to_mp4(gif_path, fps):
    """Convert a GIF to an MP4 (H.264/yuv420p) via ffmpeg, same basename.

    GIF frame delays are quantised to 1/100s, so anything requesting faster
    than 100 fps collapses to a delay of 0-1 centiseconds; ffmpeg's GIF
    demuxer treats that as invalid and silently substitutes a 10 fps default
    (the same fallback browsers use), turning e.g. a 1000-fps request into a
    video 100x longer than intended. -r before -i overrides the demuxer's
    per-frame timing with the real requested fps instead of trusting the
    (possibly-clamped) delays baked into the GIF.
    """
    if not shutil.which("ffmpeg"):
        sys.exit("ffmpeg not found in PATH — install it or drop --mp4.")

    mp4_path = str(Path(gif_path).with_suffix(".mp4"))
    cmd = [
        "ffmpeg", "-y", "-r", str(fps), "-i", gif_path,
        "-movflags", "faststart",
        "-pix_fmt", "yuv420p",
        # H.264 requires even width/height; GIF frame sizes are not guaranteed to be.
        "-vf", "scale=trunc(iw/2)*2:trunc(ih/2)*2",
        mp4_path,
    ]
    print(f"Converting to MP4: {' '.join(cmd)}")
    result = subprocess.run(cmd, capture_output=True, text=True)
    if result.returncode != 0:
        sys.exit(f"ffmpeg failed:\n{result.stderr}")
    print(f"  Wrote {mp4_path}")


def main():
    args = parse_args()
    if args.rawtlt and args.depth_label:
        sys.exit(
            "--rawtlt and --depth-label both write a label into the same "
            "corner; pass one. --rawtlt is for a tilt series, --depth-label "
            "for a tomogram."
        )
    if args.output is None:
        args.output = str(Path(args.input).with_suffix(".gif"))
        print(f"Output: {args.output}")
    duration_ms = int(round(1000 / args.fps))

    # -x/-y/-z are specified in the *input* file's own pixel coordinates, not
    # whatever --bin produces, so parse and bounds-check them against the
    # input's own header before any binning happens. Header-only: no data
    # read yet.
    with mrcfile.open(args.input, header_only=True, permissive=True) as m0:
        nz0, ny0, nx0 = int(m0.header.nz), int(m0.header.ny), int(m0.header.nx)
    full_nframes0, h0, w0 = dims_for_axis(nz0, ny0, nx0, args.axis)
    zrange = parse_range(args.zrange, "zrange", full_nframes0)
    xrange = parse_range(args.xrange, "xrange", w0)
    yrange = parse_range(args.yrange, "yrange", h0)
    if yrange is not None:
        # --yrange is documented as an IMOD-style bottom-left origin, but
        # y1,y2 are applied below as plain top-origin array row indices —
        # this crop is independent of the vertical flip frame_to_pil now
        # applies to the rendered output.
        ya, yb = yrange
        yrange = (ya, yb)

    binned_tmp = None
    mrc_path = args.input
    if args.bin_factor is not None:
        tmpdir = resolve_tmpdir(args.tmpdir, args.output)
        print(f"Binning with IMOD binvol (factor {args.bin_factor}) ...")
        print(f"  Working directory for the binned copy: {tmpdir}")
        binned_tmp = run_binvol(args.input, args.bin_factor, args.axis, tmpdir)
        mrc_path = binned_tmp

    print(f"Reading {mrc_path} ...")
    mrc = mrcfile.mmap(mrc_path, mode="r", permissive=True)
    data = mrc.data

    if data.ndim == 2:
        data = data[np.newaxis]

    nframes, h, w, get_frame = make_frame_getter(data, args.axis)
    axis_label = {"z": "Z (tilt series)", "y": "Y slices", "x": "X slices"}[args.axis]
    print(f"  {data.shape}, dtype {data.dtype}  →  {nframes} frames along {axis_label}, {h} × {w} px each")

    # -x/-y/-z were parsed against the *input's* dimensions above; binvol's
    # own output (`data`) is smaller by --bin, so convert them into that
    # array's coordinates now. binvol bins XY only for --axis z (frame count,
    # i.e. nz, is untouched by -zbinning 1), but bins all three dimensions
    # for --axis y/x (the slicing axis is binned along with the frame), so
    # zrange only needs converting in the latter case.
    if args.bin_factor is not None:
        bf = args.bin_factor

        def _to_binned(r, limit):
            if r is None:
                return None
            a, b = r
            a2 = a // bf
            b2 = max(a2 + 1, b // bf)
            return (min(a2, limit - 1), min(b2, limit))

        xrange = _to_binned(xrange, w)
        yrange = _to_binned(yrange, h)
        if args.axis != "z":
            zrange = _to_binned(zrange, nframes)
        if xrange is not None or yrange is not None or zrange is not None:
            print(f"  --bin {bf}: crop ranges given in pre-bin pixels, "
                  f"converted to binned coordinates")

    full_nframes = nframes
    if zrange is not None:
        z0, z1 = zrange
        # A closure captures the *variable*, not its value at definition
        # time, and both this wrapping and the x/y one below live in the
        # same function scope. Naming this depth_get_frame rather than
        # reusing a shared name (e.g. sliced_get_frame for both) matters:
        # reusing one would let the x/y block's reassignment retarget the
        # slot this lambda already closed over, making it call itself.
        depth_get_frame = get_frame
        get_frame = lambda i: depth_get_frame(i + z0)
        nframes = z1 - z0
        print(f"  Cropped depth to z[{z0}:{z1}]  →  {nframes} frames")

    if xrange is not None or yrange is not None:
        x0, x1 = xrange if xrange is not None else (0, w)
        y0, y1 = yrange if yrange is not None else (0, h)
        uncropped_get_frame = get_frame
        get_frame = lambda i: uncropped_get_frame(i)[y0:y1, x0:x1]
        h, w = y1 - y0, x1 - x0
        print(f"  Cropped to x[{x0}:{x1}] y[{y0}:{y1}]  →  {h} × {w} px")

    new_h, new_w = compute_output_size(h, w, args.scale)
    print(f"  Output size: {new_h} × {new_w} px  ({new_h/h:.1%} of original)")

    # One label per frame, or None for no labelling at all. A tilt series
    # takes its labels from the .rawtlt file, a tomogram derives them from the
    # voxel spacing; downstream neither the drawing nor --bounce cares which.
    frame_labels = None
    if args.rawtlt:
        print(f"Loading tilt angles from {args.rawtlt} ...")
        # Loaded against full_nframes (the file has one angle per uncropped
        # frame), then sliced to match --zrange so angle i still lines up
        # with frame i of the cropped movie.
        tilt_angles = load_tilt_angles(args.rawtlt, full_nframes)
        if zrange is not None:
            tilt_angles = tilt_angles[z0:z1]
        print(f"  {len(tilt_angles)} angles: {tilt_angles[0]:.1f}° … {tilt_angles[-1]:.1f}°")
        frame_labels = [None if a is None else f"{a:+.1f}°" for a in tilt_angles]
    elif args.depth_label:
        # Read from whichever file is being drawn, so a --bin run picks up the
        # spacing binvol wrote rather than the original's. Falling back to
        # --aperpix assumes isotropic voxels, which is why it says so.
        step_a = depth_pixel_size_from_header(mrc, args.axis)
        step_source = f"{Path(mrc_path).name} header"
        if step_a is None and args.aperpix is not None:
            step_a, step_source = args.aperpix, "--aperpix (assuming cubic voxels)"
        if step_a is None:
            sys.exit(
                f"--depth-label needs the {args.axis} spacing, and "
                f"{Path(mrc_path).name} has none in its header (0 means "
                "unset). Pass --aperpix."
            )
        # Labelled against full_nframes so e.g. --depth-origin center still
        # marks the mid-plane of the uncropped volume, then sliced to match
        # --zrange — a crop should not shift what "z = 0" means.
        frame_labels = depth_labels(full_nframes, step_a, args.axis, args.depth_origin)
        if zrange is not None:
            frame_labels = frame_labels[z0:z1]
        span_nm = (nframes - 1) * step_a / 10.0
        print(f"  Depth labels: {frame_labels[0]} … {frame_labels[-1]} "
              f"({step_a:.4g} Å/frame from {step_source}, {span_nm:.1f} nm total)")

    # Scale bar setup — compute effective pixel size of the output image.
    #
    # Falling back to the header reads it from the file actually being drawn,
    # which is the binned copy when --bin-factor was used: binvol rescales the
    # pixel size, so this stays correct without knowing the bin factor here.
    aperpix = args.aperpix
    aperpix_source = "--aperpix"
    if aperpix is None:
        aperpix = pixel_size_from_header(mrc, args.axis)
        aperpix_source = f"{Path(mrc_path).name} header"

    scalebar_nm = None
    aperpix_output = None
    if aperpix is not None:
        scale_factor = new_w / w  # output/input pixel ratio
        aperpix_output = aperpix / scale_factor  # Å/px in the output image
        scalebar_nm = args.scalebar if args.scalebar is not None else nice_scalebar_nm(new_w, aperpix_output)
        bar_px = int(round(scalebar_nm * 10 / aperpix_output))
        print(f"  Pixel size: {aperpix:.4g} Å/px (from {aperpix_source})")
        print(f"  Scale bar: {scalebar_nm:g} nm = {bar_px} output pixels  "
              f"(effective pixel size {aperpix_output:.2f} Å/px)")
    elif args.scalebar is not None:
        sys.exit(
            f"--scalebar needs a pixel size, and {Path(mrc_path).name} has "
            "none in its header (0 means unset). Pass --aperpix."
        )

    font_size = args.label_size or max(12, int(new_h * 0.04))

    print(f"Estimating contrast from {min(args.stat_frames, nframes)} sample frames ...")
    lo, hi = estimate_stats(nframes, get_frame, args.stat_frames, args.contrast)
    print(f"  Clipping range: [{lo:.1f}, {hi:.1f}]")

    print("Converting frames ...")
    pil_frames = []
    for i in range(nframes):
        img = frame_to_pil(get_frame(i), lo, hi, new_w, new_h, args.cmap)
        if args.rotate:
            img = img.rotate(args.rotate, expand=True)  # works for L and RGB
        if frame_labels is not None and frame_labels[i] is not None:
            img = add_label(img, frame_labels[i], font_size,
                            args.label_color, args.label_pos)
        if scalebar_nm is not None:
            img = add_scalebar(img, scalebar_nm, aperpix_output, font_size,
                               args.scalebar_color, args.scalebar_pos)
        pil_frames.append(to_gif_frame(img))
        print(f"  frame {i+1}/{nframes}", end="\r")
    print()

    mrc.close()
    if binned_tmp is not None:
        # The whole working directory, so any backup file binvol left beside
        # the binned copy goes with it.
        shutil.rmtree(Path(binned_tmp).parent, ignore_errors=True)

    if args.bounce:
        # forward + reversed, excluding duplicated end-points
        pil_frames = pil_frames + pil_frames[-2:0:-1]
        print(f"  Bounce mode: {len(pil_frames)} frames total (forward + reverse)")

    print(f"Writing {args.output}  ({len(pil_frames)} frames @ {args.fps} fps) ...")
    pil_frames[0].save(
        args.output,
        format="GIF",
        save_all=True,
        append_images=pil_frames[1:],
        duration=duration_ms,
        loop=args.loop,
        optimize=False,
    )
    if args.mp4:
        convert_gif_to_mp4(args.output, args.fps)

    print("Done.")


if __name__ == "__main__":
    main()
```
