## Supplementary material for "Whole cryo-FIB lamella acquisition at high magnification with rectangular beam montage cryo-electron tomography": Source code: overlay_picks_on_montage.py

```
#!/usr/bin/env python3
"""
Check the pick -> tile coordinate chain by looking at it, one link at a time.

``montage_to_relion`` composes four coordinate transforms, each with its own
sign convention. A wrong one produces a perfectly well-formed star file whose
particles are noise, and the failure is invisible until a refinement has burned
a day of GPU. This script projects the picks forward and draws them, in three
stages chosen so that each isolates one link:

``aligned``
    Predicted positions on AreTomo's own aligned tilt series. Exercises only
    the tomogram -> aligned-frame shear. Note that this stage still needs the
    ``.aln``, and not merely for bookkeeping: with ``-TiltCor 1 12`` the section
    at nominal 0 deg reprojects at TILT = +12 deg, so sin(theta) = 0.21 and a
    particle 500 px off the mid-plane moves ~100 px in the image. There is no
    tilt at which the geometry reduces to "drop the Z coordinate". The nearest
    thing to a free lunch is the section whose *TILT column* is nearest zero --
    nominal -12 deg here -- where the Z term genuinely vanishes and a
    discrepancy can only come from the 2D part.

``canvas``
    Predicted positions on the stitched montage. Adds the aligned -> canvas
    inverse alignment. A systematic rotation here means the ROT convention;
    a systematic translation means TX/TY.

``patches``
    Cuts a box out of the *raw tile* at each predicted position and contact-
    sheets them, plus their average. This is the one that matters: it exercises
    the tile assignment, the rot90 and the binning offset, and it is the only
    stage that can tell you the picks landed on density rather than merely in
    a plausible place. The average patch also *measures* the residual offset,
    which is reported as the ``--extra-shift`` to feed back to
    ``montage_to_relion``.

Examples
--------
    overlay_picks_on_montage --picks Ribosomes.txt --tomogram recon_bin2.mrc \\
        --aln Montage_9-A_inpainted.aln --positions-dir stitched_motioncorr \\
        --tile-dir motion_corrected --mode all --tilt -12 --tilt 0 --tilt 30 \\
        --aligned-stack recon_patch53mask_bin2tiltseries.mrc \\
        --canvas-stack stitched_motioncorr/Montage_9-A.mrc -o overlays/
"""

from __future__ import annotations

import argparse
import logging
import os
from typing import List, Optional, Sequence

import matplotlib

matplotlib.use("Agg")

import matplotlib.pyplot as plt  # noqa: E402
import mrcfile  # noqa: E402
import numpy as np  # noqa: E402

from .montage_projection import (  # noqa: E402
    add_geometry_args, build_projector, load_picks)

logger = logging.getLogger(__name__)


def _show(ax, img: np.ndarray, sigma: float = 3.0) -> None:
    """Display with a robust contrast range; montage data has huge outliers."""
    finite = img[np.isfinite(img)]
    if finite.size == 0:
        return
    med = np.median(finite)
    mad = np.median(np.abs(finite - med)) * 1.4826 or finite.std() or 1.0
    ax.imshow(img, cmap="gray", vmin=med - sigma * mad, vmax=med + sigma * mad,
              origin="upper", interpolation="nearest")


def _downsample(img: np.ndarray, target: int = 2000) -> tuple:
    step = max(1, int(max(img.shape) // target))
    return img[::step, ::step], step


# ---------------------------------------------------------------------------
# Stages
# ---------------------------------------------------------------------------


def overlay_aligned(proj, picks, tilt, stack, outdir) -> Optional[str]:
    """Predicted positions on AreTomo's aligned tilt series."""
    sec = proj.tilts[tilt].sec
    zi = proj.aligned_slice_index(tilt)
    with mrcfile.mmap(stack, mode="r", permissive=True) as m:
        if zi >= m.data.shape[0]:
            logger.error("aligned stack has %d slices, need %d", m.data.shape[0], zi)
            return None
        img = np.asarray(m.data[zi], dtype=np.float32)

    al = proj.tomo_to_aligned(tilt, picks)
    small, step = _downsample(img)
    fig, ax = plt.subplots(figsize=(10, 10))
    _show(ax, small)
    ax.scatter(al[:, 1] / step, al[:, 0] / step, s=28, facecolors="none",
               edgecolors="#ff3b30", linewidths=0.8)
    ax.set_title(f"aligned frame, nominal tilt {tilt:+.1f} deg "
                 f"(.aln SEC {sec.sec}, TILT {sec.tilt:+.2f})")
    ax.set_axis_off()
    path = os.path.join(outdir, f"aligned_{tilt:+.1f}.png")
    fig.savefig(path, dpi=140, bbox_inches="tight")
    plt.close(fig)
    return path


def overlay_canvas(proj, picks, tilt, stack, outdir) -> Optional[str]:
    """Predicted positions on the stitched montage canvas."""
    zi = proj.canvas_slice_index(tilt)
    with mrcfile.mmap(stack, mode="r", permissive=True) as m:
        if zi >= m.data.shape[0]:
            logger.error("canvas stack has %d slices, need %d", m.data.shape[0], zi)
            return None
        img = np.asarray(m.data[zi], dtype=np.float32)

    rc = proj.tomo_to_canvas(tilt, picks)
    small, step = _downsample(img)
    fig, ax = plt.subplots(figsize=(10, 10))
    _show(ax, small)
    ax.scatter(rc[:, 1] / step, rc[:, 0] / step, s=28, facecolors="none",
               edgecolors="#ff3b30", linewidths=0.8)
    ax.set_title(f"stitched canvas, tilt {tilt:+.1f} deg")
    ax.set_axis_off()
    path = os.path.join(outdir, f"canvas_{tilt:+.1f}.png")
    fig.savefig(path, dpi=140, bbox_inches="tight")
    plt.close(fig)
    return path


def verify_rotation(proj, tilt: float, canvas_stack: str) -> None:
    """
    Settle the 180 deg that tile footprints cannot.

    ``MontageProjector`` detects ``stitch.py --rotate`` from the shape and
    placement of the tile footprints, which is blind to a half turn. Here the
    raw tile is binned and rotated both ways and correlated against the piece of
    canvas it was supposed to have produced, masked to the pixels ``index_map``
    attributes to it. The stitched canvas is literally made of these pixels, so
    the correct rotation correlates near 1 and the wrong one near 0. If the two
    scores are close, the tile has too little contrast -- try another tilt.
    """
    info = proj.tilts[tilt]
    imap = info.index_map()
    zi = proj.all_tilts.index(tilt)
    with mrcfile.mmap(canvas_stack, mode="r", permissive=True) as m:
        canvas = np.asarray(m.data[zi], dtype=np.float32)

    # Use the label with the most canvas pixels, i.e. the least-clipped tile.
    counts = np.bincount(imap.ravel(), minlength=int(imap.max()) + 1)
    label = int(np.argmax(counts[: int(imap.max())]))
    tile = int(info.selected[label])
    r0, c0 = np.round(proj.geom.tile_origin_rc(info.positions_um)[tile]).astype(int)
    h, w = proj.geom.tile_shape
    if r0 < 0 or c0 < 0 or r0 + h > imap.shape[0] or c0 + w > imap.shape[1]:
        logger.warning("rotation check: tile %d is clipped by the canvas, skipping", tile)
        info.drop_index_map()
        return

    ref = canvas[r0:r0 + h, c0:c0 + w]
    mask = (imap[r0:r0 + h, c0:c0 + w] == label)
    info.drop_index_map()
    if mask.sum() < 10000:
        logger.warning("rotation check: tile %d owns too little canvas, skipping", tile)
        return

    with mrcfile.mmap(info.stack_path, mode="r", permissive=True) as m:
        raw = np.asarray(m.data[tile], dtype=np.float32)
    b = proj.binning
    ny, nx = raw.shape[0] // b * b, raw.shape[1] // b * b
    binned = raw[:ny, :nx].reshape(ny // b, b, nx // b, b).mean(axis=(1, 3))

    scores = {}
    for k in (proj.rotate // 90 % 4, (proj.rotate // 90 + 2) % 4):
        cand = np.rot90(binned, k)
        if cand.shape != ref.shape:
            continue
        a, bb = cand[mask], ref[mask]
        a = a - a.mean()
        bb = bb - bb.mean()
        denom = np.sqrt((a * a).sum() * (bb * bb).sum())
        scores[k * 90] = float((a * bb).sum() / denom) if denom else 0.0

    logger.info("rotation check on tilt %+.1f, tile %d: %s",
                tilt, tile, {k: round(v, 4) for k, v in scores.items()})
    if not scores:
        return
    best = max(scores, key=scores.get)
    if best != proj.rotate:
        logger.error(
            "ROTATION IS WRONG: the canvas prefers rotate=%d (r=%.3f) over the "
            "assumed %d (r=%.3f). Re-run with --rotate %d.",
            best, scores[best], proj.rotate, scores.get(proj.rotate, float("nan")), best,
        )
    elif scores[best] < 0.3:
        logger.warning("rotation check inconclusive (best r=%.3f)", scores[best])
    else:
        logger.info("rotation confirmed: rotate=%d (r=%.3f)", best, scores[best])


def _cut(proj, picks, tilts, box, jitter_A=0.0, rng=None):
    """Cut a box from the raw tile at each predicted position."""
    half = box // 2
    patches, labels = [], []
    for tilt in tilts:
        pts = picks
        if jitter_A:
            # Displace within the specimen plane only, so the decoys sit in the
            # same tiles, at the same dose and the same defocus, and differ from
            # the real positions in one respect alone: they are not on a particle.
            d = jitter_A / (proj.pixel_size * proj.tomo_bin)
            ang = rng.uniform(0, 2 * np.pi, len(picks))
            pts = picks + np.stack(
                [d * np.cos(ang), d * np.sin(ang), np.zeros(len(picks))], axis=1
            )
        hits = proj.assign(tilt, pts, margin=half)
        proj.tilts[tilt].drop_index_map()
        stack = proj.tilts[tilt].stack_path
        if stack is None:
            continue
        with mrcfile.mmap(stack, mode="r", permissive=True) as m:
            for n, hit in enumerate(hits):
                if hit is None:
                    continue
                xi, yi = int(round(hit.x)), int(round(hit.y))
                if not (half <= xi < m.data.shape[2] - half
                        and half <= yi < m.data.shape[1] - half):
                    continue
                patch = np.asarray(
                    m.data[hit.tile, yi - half:yi + half, xi - half:xi + half],
                    dtype=np.float32,
                )
                if patch.shape != (box, box):
                    continue
                patches.append(patch)
                labels.append(f"p{n} t{tilt:+.0f} tile{hit.tile}")
    return patches, labels


def overlay_patches(proj, picks, tilts, box, outdir, max_show=64,
                    particle_diameter=300.0, decoy=True):
    """
    Cut a box from the raw tile at each predicted position; sheet them, average
    them, and compare that average against a decoy average.

    Two things make the naive version of this test useless, and both were
    learned the hard way on 9-A:

    * A single tile at ~3 e-/A^2 is nearly pure noise at full sampling, so the
      mean has to be judged at the particle's own spatial scale. A ribosome is
      ~88 px across at 3.426 A/px; looking at the unfiltered mean is looking at
      the wrong frequency band and finds nothing however good the geometry is.
    * "There is a blob" is not evidence on its own. So the same average is also
      built from decoy positions, displaced sideways within the same tiles, and
      the two are compared. If the real average is not clearly stronger, the
      geometry is wrong or the picks are not particles -- and the test says so
      instead of leaving it to the eye.
    """
    rng = np.random.default_rng(0)
    patches, labels = _cut(proj, picks, tilts, box)
    if not patches:
        logger.error("No patches could be cut -- every pick fell outside a tile.")
        return None
    logger.info("cut %d patches of %d px from %d tilts", len(patches), box, len(tilts))
    arr = np.stack(patches)
    # Normalise each patch before averaging: tiles differ in mean and in dose,
    # and an unnormalised mean is dominated by whichever tile is brightest.
    arr = (arr - arr.mean(axis=(1, 2), keepdims=True)) / (
        arr.std(axis=(1, 2), keepdims=True) + 1e-6
    )
    mean = arr.mean(axis=0)

    n_show = min(max_show, len(patches))
    ncol = int(np.ceil(np.sqrt(n_show)))
    nrow = int(np.ceil(n_show / ncol))
    fig, axes = plt.subplots(nrow, ncol, figsize=(1.5 * ncol, 1.5 * nrow))
    for k, ax in enumerate(np.atleast_1d(axes).ravel()):
        ax.set_axis_off()
        if k < n_show:
            _show(ax, arr[k])
            ax.set_title(labels[k], fontsize=5)
    fig.suptitle(f"raw-tile patches at predicted positions (box {box} px)")
    sheet = os.path.join(outdir, "patches_sheet.png")
    fig.savefig(sheet, dpi=150, bbox_inches="tight")
    plt.close(fig)

    # -- the residual measurement, at the particle's own spatial scale --
    sigma_px = particle_diameter / proj.pixel_size / 6.0
    dr, dc, contrast, sm = _peak_offset(mean, sigma_px)

    decoy_sm, decoy_contrast = None, None
    if decoy:
        dpatches, _ = _cut(proj, picks, tilts, box,
                           jitter_A=2.5 * particle_diameter, rng=rng)
        if dpatches:
            darr = np.stack(dpatches)
            darr = (darr - darr.mean(axis=(1, 2), keepdims=True)) / (
                darr.std(axis=(1, 2), keepdims=True) + 1e-6
            )
            _, _, decoy_contrast, decoy_sm = _peak_offset(darr.mean(axis=0), sigma_px)

    panels = [("picks", sm, contrast)]
    if decoy_sm is not None:
        panels.append((f"decoy (+{2.5 * particle_diameter:.0f} A)", decoy_sm,
                       decoy_contrast))
    fig, axes = plt.subplots(1, len(panels), figsize=(5 * len(panels), 5.4))
    for ax, (title, img, con) in zip(np.atleast_1d(axes), panels):
        _show(ax, img, sigma=4)
        ax.axhline(box / 2, color="#ff3b30", lw=0.5)
        ax.axvline(box / 2, color="#ff3b30", lw=0.5)
        ax.set_title(f"{title}\n|peak|/rms = {con:.2f}")
        ax.set_axis_off()
    np.atleast_1d(axes)[0].plot(box / 2 + dc, box / 2 + dr, "+", color="#34c759", ms=12)
    fig.suptitle(f"mean of {len(patches)} patches, low-passed to "
                 f"~{particle_diameter:.0f} A")
    avg = os.path.join(outdir, "patches_mean.png")
    fig.savefig(avg, dpi=150, bbox_inches="tight")
    plt.close(fig)

    logger.info("mean patch: offset (%+0.1f, %+0.1f) raw px, |peak|/rms = %.2f",
                dr, dc, contrast)
    if decoy_contrast is not None:
        logger.info("decoy mean: |peak|/rms = %.2f", decoy_contrast)
    ok = contrast >= 2.0 and (decoy_contrast is None or contrast > 1.5 * decoy_contrast)
    if not ok:
        logger.warning(
            "The picks' average is not clearly stronger than the decoy's "
            "(%.2f vs %s). Either the geometry is wrong or these picks are not "
            "particles -- do not proceed to extraction on this evidence.",
            contrast, "n/a" if decoy_contrast is None else f"{decoy_contrast:.2f}",
        )
    else:
        # Map the residual back through the rot90 into a canvas-pixel shift.
        k = (proj.rotate // 90) % 4
        v = np.array([dr, dc]) / proj.binning
        for _ in range(k):
            v = np.array([-v[1], v[0]])
        logger.info("  -> try  --extra-shift %+.2f %+.2f  (canvas px)", -v[0], -v[1])
    return avg


def _peak_offset(mean: np.ndarray, sigma_px: float, search: int = 24):
    """
    Offset of the mean patch's central feature from the box centre.

    Low-passed to the particle's own scale first -- at full sampling a low-dose
    tile average is dominated by frequencies far above anything a 300 A particle
    puts signal into, and the peak search just finds noise. The peak is signed
    either way, since particles may be dark or bright depending on whether
    contrast was inverted upstream, and the rms it is measured against comes
    from the same filtered image so the two are comparable.
    """
    from scipy.ndimage import gaussian_filter

    sm = gaussian_filter(mean, max(sigma_px, 1.0))
    n = sm.shape[0]
    c = n // 2
    win = sm[c - search:c + search, c - search:c + search]
    rms = sm.std() or 1.0
    flat = win - sm.mean()
    idx = np.unravel_index(np.argmax(np.abs(flat)), flat.shape)
    return float(idx[0] - search), float(idx[1] - search), float(abs(flat[idx]) / rms), sm


# ---------------------------------------------------------------------------


def parse_commandline(argv=None):
    p = argparse.ArgumentParser(
        description=__doc__, formatter_class=argparse.RawDescriptionHelpFormatter
    )
    p.add_argument("--picks", required=True)
    p.add_argument("--tomogram", required=True)
    p.add_argument("--aln", required=True)
    p.add_argument("--positions-dir", required=True)
    p.add_argument("--tile-dir", required=True)
    p.add_argument("-o", "--out", required=True)
    p.add_argument("--mode", choices=["aligned", "canvas", "patches", "all"],
                   default="all")
    p.add_argument("--tilt", type=float, action="append", default=None,
                   help="tilt(s) to draw; repeatable. Default: the section whose "
                        ".aln TILT is nearest zero, plus the two extremes.")
    p.add_argument("--aligned-stack", default=None)
    p.add_argument("--canvas-stack", default=None)
    p.add_argument("--box", type=int, default=256)
    p.add_argument("--particle-diameter", type=float, default=300.0,
                   help="A; sets the low-pass the patch average is judged at")
    p.add_argument("--no-decoy", action="store_true")
    p.add_argument("--patch-tilts", type=int, default=9,
                   help="how many of the lowest-|TILT| sections to average over")

    add_geometry_args(p)
    p.add_argument("--verbose", "-v", action="store_true")
    return p.parse_args(argv)


def main(argv=None) -> int:
    args = parse_commandline(argv)
    logging.basicConfig(
        level=logging.DEBUG if args.verbose else logging.INFO,
        format="%(levelname)s %(message)s",
    )
    os.makedirs(args.out, exist_ok=True)

    with mrcfile.open(args.tomogram, header_only=True, permissive=True) as m:
        recon_shape = (int(m.header.nz), int(m.header.ny), int(m.header.nx))

    proj = build_projector(args, recon_shape)
    picks = load_picks(args.picks)
    logger.info("%d picks", len(picks))

    available = sorted(proj.tilts)
    if args.tilt:
        tilts = [min(available, key=lambda t: abs(t - x)) for x in args.tilt]
    else:
        # The .aln TILT nearest zero is where the Z term drops out entirely, so
        # it isolates the 2D part of the chain; the extremes are where a wrong
        # Z sign is most obvious.
        flat = min(available, key=lambda t: abs(proj.tilts[t].sec.tilt))
        tilts = sorted({available[0], flat, available[-1]})
    logger.info("drawing tilts %s (.aln TILT %s)",
                [f"{t:+.1f}" for t in tilts],
                [f"{proj.tilts[t].sec.tilt:+.1f}" for t in tilts])

    # A table is often enough to spot a gross error before any image is drawn.
    for tilt in tilts:
        hits = proj.assign(tilt, picks, margin=args.box / 2.0)
        proj.tilts[tilt].drop_index_map()
        got = [h for h in hits if h is not None]
        logger.info("tilt %+6.1f: %d/%d picks placed", tilt, len(got), len(picks))
        for h in got[:5]:
            logger.info(
                "   tile %2d (z-slice %2d)  x=%7.1f y=%7.1f  canvas=(%.0f, %.0f)  "
                "edge margin %5.0f px", h.tile, h.tile, h.x, h.y,
                h.canvas_rc[0], h.canvas_rc[1], h.edge_margin,
            )

    if args.mode in ("aligned", "all"):
        if args.aligned_stack:
            for tilt in tilts:
                p = overlay_aligned(proj, picks, tilt, args.aligned_stack, args.out)
                if p:
                    logger.info("wrote %s", p)
        elif args.mode == "aligned":
            logger.error("--aligned-stack is required for mode 'aligned'")

    if args.mode in ("canvas", "all"):
        if args.canvas_stack:
            verify_rotation(proj, tilts[len(tilts) // 2], args.canvas_stack)
            for tilt in tilts:
                p = overlay_canvas(proj, picks, tilt, args.canvas_stack, args.out)
                if p:
                    logger.info("wrote %s", p)
        elif args.mode == "canvas":
            logger.error("--canvas-stack is required for mode 'canvas'")

    if args.mode in ("patches", "all"):
        low = sorted(available, key=lambda t: abs(proj.tilts[t].sec.tilt))
        overlay_patches(proj, picks, low[: args.patch_tilts], args.box, args.out,
                        particle_diameter=args.particle_diameter,
                        decoy=not args.no_decoy)

    return 0


if __name__ == "__main__":
    raise SystemExit(main())
```
