## Supplementary material for "Whole cryo-FIB lamella acquisition at high magnification with rectangular beam montage cryo-electron tomography": Source code: picks_to_imod_model.py

```
#!/usr/bin/env python3
"""
Write IMOD models of reprojected 3D picks, so the coordinate chain can be
checked in 3dmod against the actual images.

A PNG overlay shows you where the points went; a model lets you scrub through
sections with the images underneath, which is what actually catches a slowly
accumulating error. Two models, matching the two things worth checking:

``tiltseries``
    One contour per particle, one point per tilt, on the tilt series. Exactly
    the shape of an IMOD fiducial model, and readable the same way: step
    through Z with the contour selected and the point should stay nailed to the
    same feature. A residual that grows towards high tilt is the Z/shear term;
    a constant offset is TX/TY; a rotation that opens up across the field is
    ROT. Available in two frames:

      --frame canvas   the stitched montage (6.852 A/px here), 41 sections,
                       including the ones AreTomo later called dark
      --frame aligned  AreTomo's own aligned stack (13.704 A/px), 35 sections,
                       in surviving-SEC order

``tiles``
    For one tilt, the raw per-tilt stack -- where each Z section is a *tile*,
    not a view -- with each particle plotted on the tile it was assigned to.
    This is the end of the chain: it exercises the tile assignment, the rot90
    and the binning offset, and it is where you find out whether a particle
    really is where the projection says it is in the unstitched data.

    A second object holds the picks that fall inside a tile's rectangle but
    were rejected because the extraction box would have crossed the square-beam
    edge. Those should sit in a band around each tile's illuminated square; if
    they are scattered through the middle of tiles instead, the beam masks are
    not being read correctly.

Section numbering is 0-based in the model, matching the image Z index, so
3dmod's slider (which counts from 1) will read one higher.
"""

from __future__ import annotations

import argparse
import logging
import os
import shutil
import subprocess
from typing import List, Optional, Sequence, Tuple

import mrcfile
import numpy as np

from .montage_projection import (
    add_geometry_args, build_projector, load_picks)

logger = logging.getLogger(__name__)

# object, contour, x, y, z -- verified to round-trip through
# point2model / model2point -zero without -zcoord (a -0.5 shift otherwise).
Point = Tuple[int, int, float, float, float]


def write_model(
    points: Sequence[Point],
    image: str,
    out_mod: str,
    names: Sequence[str],
    colours: Sequence[Tuple[int, int, int]],
    symbol_size: int,
    scattered: bool,
) -> Optional[str]:
    """
    Write a point list and convert it to a .mod with IMOD's point2model.

    Marks the points with a 2D circle (``-circle``), never a sphere
    (``-sphere``). A sphere radius is a radius in *pixels applied in Z as well*,
    so 3dmod draws the point on every section the sphere intersects. That is
    right for a tomogram, where Z is a spatial axis, and wrong for both stacks
    here, where Z is a tilt number or a tile number: a 300 A particle is a
    22-44 pixel radius, so every point would be drawn on every section and the
    model would be unreadable. ``-circle`` sets ``pointsize 0`` and a 2D symbol
    instead, which 3dmod draws only on the point's own section.

    The text file is kept alongside the model: if point2model is not on PATH the
    conversion is the only part that fails, and the command to finish it by hand
    is printed rather than the work being lost.
    """
    out_txt = os.path.splitext(out_mod)[0] + ".txt"
    with open(out_txt, "w") as fh:
        for ob, co, x, y, z in points:
            fh.write(f"{ob} {co} {x:.2f} {y:.2f} {z:.1f}\n")

    cmd = ["point2model", "-image", image, "-circle", str(int(symbol_size))]
    cmd.append("-scat" if scattered else "-open")
    for n in names:
        cmd += ["-name", n]
    for c in colours:
        # IMOD's PIP parser takes a multi-value option as one comma-separated
        # argument. Passing "0 255 0" leaves 255 and 0 as positional arguments
        # and point2model then tries to open "255" as the input file.
        cmd += ["-color", f"{c[0]},{c[1]},{c[2]}"]
    cmd += [out_txt, out_mod]

    if shutil.which("point2model") is None:
        logger.warning(
            "point2model not on PATH (source /programs/sbgrid.shrc); wrote %s. "
            "Convert it with:\n  %s", out_txt, " ".join(cmd),
        )
        return None
    res = subprocess.run(cmd, capture_output=True, text=True)
    if res.returncode != 0:
        logger.error("point2model failed: %s", res.stderr.strip() or res.stdout.strip())
        return None
    logger.info("wrote %s  (%d points)", out_mod, len(points))
    logger.info("    3dmod %s %s", image, out_mod)
    return out_mod


# ---------------------------------------------------------------------------


def model_tiltseries(proj, picks, frame, image, out_mod, symbol_size, margin):
    """
    One contour per particle, one point per tilt, in the tilt series frame.

    Object 1 carries the tracks. Object 2 marks the (particle, tilt) pairs that
    have no usable tile -- the particle is outside the montage, or too close to
    a beam edge for a box. Those are not errors, but a particle that vanishes
    from object 1 and appears in object 2 at low tilt is worth a look.
    """
    if frame == "aligned":
        z_of = {t: proj.aligned_slice_index(t) for t in proj.tilts}
        project = proj.tomo_to_aligned
    else:
        z_of = {t: proj.canvas_slice_index(t) for t in proj.tilts}
        project = proj.tomo_to_canvas

    with mrcfile.open(image, header_only=True, permissive=True) as m:
        nx, ny, nz = int(m.header.nx), int(m.header.ny), int(m.header.nz)

    tracks: List[Point] = []
    dropped: List[Point] = []
    n_out = 0
    for tilt in sorted(proj.tilts):
        rc = project(tilt, picks)[:, :2]
        hits = proj.assign(tilt, picks, margin=margin)
        proj.tilts[tilt].drop_index_map()
        z = z_of[tilt]
        if z >= nz:
            logger.warning("tilt %+.1f maps to section %d but %s has %d",
                           tilt, z, os.path.basename(image), nz)
            continue
        for n in range(len(picks)):
            # IMOD y is the numpy row index: MRC stores rows bottom-up and
            # 3dmod displays them the same way, so no flip is needed here even
            # though the PNG overlays (origin="upper") look vertically mirrored.
            y, x = float(rc[n, 0]), float(rc[n, 1])
            if not (0 <= x < nx and 0 <= y < ny):
                n_out += 1
                continue
            if hits[n] is None:
                dropped.append((2, n + 1, x, y, z))
            else:
                tracks.append((1, n + 1, x, y, z))

    points = sorted(tracks + dropped, key=lambda p: (p[0], p[1], p[4]))
    logger.info(
        "%s frame: %d tracked points, %d with no usable tile, %d outside the image",
        frame, len(tracks), len(dropped), n_out,
    )
    return write_model(
        points, image, out_mod,
        names=["reprojected picks", "no usable tile"],
        colours=[(0, 255, 0), (255, 0, 0)],
        symbol_size=symbol_size, scattered=False,
    )


def model_tiles(proj, picks, tilt, out_mod, symbol_size, margin):
    """
    One tilt's raw tile stack, with each particle on the tile it was assigned.

    Z here is the tile index, not a view. Object 1 is the assignment actually
    used; object 2 is every (particle, tile) pair that lies inside the tile but
    was rejected for crossing the beam edge, which is what makes the beam mask
    visible as a band rather than something you have to take on trust.
    """
    info = proj.tilts[tilt]
    if info.stack_path is None:
        logger.error("no tile stack for tilt %+.1f", tilt)
        return None

    hits = proj.assign(tilt, picks, margin=margin)
    canvas = proj.tomo_to_canvas(tilt, picks)
    org = proj.geom.tile_origin_rc(info.positions_um)
    beams = info.beam_masks()
    h, w = proj.geom.tile_shape
    half = int(np.ceil(margin / proj.binning))

    used: List[Point] = []
    rejected: List[Point] = []
    for n, hit in enumerate(hits):
        if hit is not None:
            used.append((1, hit.tile + 1, hit.x, hit.y, float(hit.tile)))
        # Everything geometrically inside a tile but not chosen, so the
        # rejection band around each beam is visible.
        for label, tile in enumerate(info.selected):
            if hit is not None and tile == hit.tile:
                continue
            local = canvas[n] - org[tile]
            if not (0 <= local[0] < h and 0 <= local[1] < w):
                continue
            if beams is not None:
                r0, c0 = int(round(local[0])) - half, int(round(local[1])) - half
                inside = (r0 >= 0 and c0 >= 0 and r0 + 2 * half <= h
                          and c0 + 2 * half <= w
                          and beams[label, r0:r0 + 2 * half, c0:c0 + 2 * half].all())
                if inside:
                    continue  # usable, just not the roomiest -- not a rejection
            raw = proj.canvas_to_raw_tile(local[None, :])[0]
            rejected.append((2, int(tile) + 1, float(raw[1]), float(raw[0]), float(tile)))
    info.drop_index_map()

    points = sorted(used + rejected, key=lambda p: (p[0], p[1]))
    logger.info(
        "tilt %+.1f: %d particles placed on %d tiles, %d beam-edge rejections",
        tilt, len(used), len({p[1] for p in used}), len(rejected),
    )
    return write_model(
        points, info.stack_path, out_mod,
        names=[f"picks on tiles, tilt {tilt:+.1f}", "rejected: box crosses beam edge"],
        colours=[(0, 255, 0), (255, 0, 0)],
        symbol_size=symbol_size, scattered=True,
    )


# ---------------------------------------------------------------------------


def parse_commandline(argv=None):
    p = argparse.ArgumentParser(
        description=__doc__, formatter_class=argparse.RawDescriptionHelpFormatter
    )
    p.add_argument("--picks", required=True)
    p.add_argument("--tomogram", required=True)
    p.add_argument("--aln", required=True)
    p.add_argument("--positions-dir", required=True)
    p.add_argument("--tile-dir", required=True)
    p.add_argument("-o", "--out", required=True)
    p.add_argument("--mode", choices=["tiltseries", "tiles", "both"], default="both")
    p.add_argument("--frame", choices=["canvas", "aligned"], default="canvas")
    p.add_argument("--tiltseries-image", default=None,
                   help="the stack the tiltseries model goes with: the stitched "
                        "montage for --frame canvas, AreTomo's aligned stack for "
                        "--frame aligned")
    p.add_argument("--tilt", action="append", default=None, metavar="ANGLE",
                   help="tilt(s) for the per-tile model; repeatable, or 'all' "
                        "for every tilt. The default is the section whose .aln "
                        "TILT is nearest zero -- which is precisely the tilt "
                        "where a reprojection error perpendicular to the tilt "
                        "axis is multiplied by sin(theta) = 0 and so cannot be "
                        "seen. Pass the extremes as well.")
    p.add_argument("--box", type=int, default=256,
                   help="extraction box, unbinned px; sets the beam-edge margin")
    p.add_argument("--particle-diameter", type=float, default=300.0,
                   help="A; sets the displayed circle radius")

    add_geometry_args(p)
    p.add_argument("--verbose", "-v", action="store_true")
    return p.parse_args(argv)


def main(argv=None) -> int:
    args = parse_commandline(argv)
    logging.basicConfig(
        level=logging.DEBUG if args.verbose else logging.INFO,
        format="%(levelname)s %(message)s",
    )
    os.makedirs(args.out, exist_ok=True)

    with mrcfile.open(args.tomogram, header_only=True, permissive=True) as m:
        recon_shape = (int(m.header.nz), int(m.header.ny), int(m.header.nx))

    proj = build_projector(args, recon_shape)
    picks = load_picks(args.picks)
    logger.info("%d picks", len(picks))
    margin = args.box / 2.0
    radius = args.particle_diameter / 2.0

    if args.mode in ("tiltseries", "both"):
        if not args.tiltseries_image:
            logger.error("--tiltseries-image is required for the tiltseries model")
        else:
            # Sphere radius in the pixels of whichever frame we are drawing in.
            apix = (proj.pixel_size * proj.binning if args.frame == "canvas"
                    else proj.pixel_size * proj.tomo_bin)
            model_tiltseries(
                proj, picks, args.frame, args.tiltseries_image,
                os.path.join(args.out, f"picks_tiltseries_{args.frame}.mod"),
                symbol_size=max(int(round(radius / apix)), 2), margin=margin,
            )

    if args.mode in ("tiles", "both"):
        available = sorted(proj.tilts)
        if args.tilt and any(str(x).strip().lower() == "all" for x in args.tilt):
            tilts = available
        elif args.tilt:
            tilts = [min(available, key=lambda t: abs(t - float(x)))
                     for x in args.tilt]
        else:
            tilts = [min(available, key=lambda t: abs(proj.tilts[t].sec.tilt))]
        logger.info("per-tile models for tilt(s): %s",
                    ", ".join(f"{t:+.1f}" for t in tilts))
        for tilt in tilts:
            model_tiles(
                proj, picks, tilt,
                os.path.join(args.out, f"picks_tiles_{tilt:+.1f}.mod"),
                symbol_size=max(int(round(radius / proj.pixel_size)), 2), margin=margin,
            )
    return 0


if __name__ == "__main__":
    raise SystemExit(main())
```
