## Supplementary material for "Whole cryo-FIB lamella acquisition at high magnification with rectangular beam montage cryo-electron tomography": Source code: resize_reference_beam.py

```
#!/usr/bin/env python3
"""
Rescale a reference beam image to match the beam size actually recorded in a
real tile.

A --referencebeam image (see stitch.py) is only useful for locating and
masking the beam if its illuminated disc is the same physical size, in
pixels, as the beam in the data it is used on. If the reference was imaged at
a different beam/aperture setting than the data -- an easy mistake to make
between sessions -- a mask built from it at native size sits at the wrong
radius no matter how well it is translated into position.

This searches over candidate scipy.ndimage.zoom factors, and for each one:
resizes the reference, aligns it (translation only, via the same edge
cross-correlation stitch.py uses) to a real tile, and scores the match by the
IoU between the two beam masks. The zoom factor with the best IoU is applied
to the reference at full resolution and written out.

The search runs in two passes -- a coarse sweep over the full range, then a
finer sweep around the coarse winner -- and, for speed, both passes work on a
downsampled copy of the images (--downsample). Only the final resize, applied
once the best scale is known, runs at full resolution.
"""

import argparse
import logging
import os

import mrcfile
import numpy as np
from scipy.ndimage import zoom

from .Utilities import (
    align_beam_reference,
    crop,
    fourier_interpolate,
    make_mask,
    roll_no_periodic,
)

logger = logging.getLogger(__name__)


def _load_slice(path, index):
    """Read one 2D frame from an MRC (or TIFF) stack, plus its voxel_size."""
    ext = os.path.splitext(path)[1].lower()
    if ext in (".mrc", ".mrcs", ".rec"):
        with mrcfile.open(path, permissive=True) as m:
            data = np.asarray(m.data)
            voxel_size = m.voxel_size.copy()
    else:
        from PIL import Image
        img = Image.open(path)
        img.seek(index)
        data = np.asarray(img)
        voxel_size = None
    if data.ndim == 3:
        data = data[index]
    return data.astype(np.float32), voxel_size


def _iou_at_scale(reference, tile, factor, maskthreshold, maskabsolutethreshold):
    """IoU between tile's own beam mask and the reference's, resized by
    `factor` and translation-aligned to the tile."""
    resized = zoom(reference, factor, order=1)
    resized = crop(resized, tile.shape)

    # Same edge cross-correlation, and the same sign convention rolling the
    # result into place, that make_mask's template path and stitch.py use.
    dy, dx = align_beam_reference(tile, resized)
    resized_mask = make_mask(
        resized, shrinkn=0,
        medianthreshold=maskthreshold, absolutethreshold=maskabsolutethreshold,
    )
    aligned_mask = roll_no_periodic(resized_mask, (-dy, -dx), axis=(0, 1))
    tile_mask = make_mask(
        tile, shrinkn=0,
        medianthreshold=maskthreshold, absolutethreshold=maskabsolutethreshold,
    )

    intersection = np.logical_and(aligned_mask, tile_mask).sum()
    union = np.logical_or(aligned_mask, tile_mask).sum()
    return float(intersection) / float(union) if union else 0.0


def search_scales(reference, tile, factors, maskthreshold, maskabsolutethreshold):
    """Score every candidate scale factor; return (best_factor, best_iou, all_scores)."""
    scores = [
        _iou_at_scale(reference, tile, f, maskthreshold, maskabsolutethreshold)
        for f in factors
    ]
    for f, s in zip(factors, scores):
        logger.info("  scale %.4f  ->  IoU %.4f", f, s)
    best = int(np.argmax(scores))
    return factors[best], scores[best], scores


def parse_commandline():
    parser = argparse.ArgumentParser(
        description=__doc__,
        formatter_class=argparse.ArgumentDefaultsHelpFormatter,
    )
    parser.add_argument(
        "-r", "--reference", required=True,
        help="Reference beam MRC/TIFF to rescale. For multi-frame files, "
             "append :INDEX to select a frame (e.g. Reference.mrc:0).",
    )
    parser.add_argument(
        "-t", "--tile", default=None,
        help="MRC/TIFF stack holding a real tile at the correct beam size. "
             "For multi-frame files, append :INDEX to select a frame "
             "(e.g. Montage_21-A_0.0.mrc:0). Required unless --scale is given.",
    )
    parser.add_argument(
        "--scale", type=float, default=None,
        help="Skip the search entirely and resize --reference by this zoom "
             "factor directly -- e.g. reusing the scale a previous run "
             "already found. --tile is not needed when this is given.",
    )
    parser.add_argument(
        "-o", "--output", default=None,
        help="Output path. Defaults to reference_resized.mrc alongside --reference.",
    )
    parser.add_argument(
        "--min-scale", type=float, default=0.7,
        help="Lower bound of the coarse zoom-factor search.",
    )
    parser.add_argument(
        "--max-scale", type=float, default=1.3,
        help="Upper bound of the coarse zoom-factor search.",
    )
    parser.add_argument(
        "--coarse-steps", type=int, default=61,
        help="Number of zoom factors sampled across [--min-scale, --max-scale].",
    )
    parser.add_argument(
        "--fine-steps", type=int, default=41,
        help="Number of zoom factors sampled in the refinement pass.",
    )
    parser.add_argument(
        "--fine-window", type=float, default=0.03,
        help="Half-width (in zoom factor) of the refinement pass around the "
             "coarse best.",
    )
    parser.add_argument(
        "--downsample", type=int, default=4,
        help="Downsampling factor applied only while searching for the best "
             "scale, for speed. The final resize is always done at full "
             "resolution.",
    )
    parser.add_argument(
        "-mt", "--maskthreshold", type=float, default=0.4,
        help="Beam-detection threshold as a fraction of the image median, "
             "used only for scoring candidate scales.",
    )
    parser.add_argument(
        "-ma", "--maskabsolutethreshold", type=float, default=None,
        help="Absolute beam-detection threshold; overrides --maskthreshold "
             "if given.",
    )
    return vars(parser.parse_args())


def _split_index(path_arg):
    if ":" in os.path.basename(path_arg):
        path, idx_str = path_arg.rsplit(":", 1)
        return path, int(idx_str)
    return path_arg, 0


def main():
    logging.basicConfig(level=logging.INFO, format="%(message)s")
    args = parse_commandline()

    ref_path, ref_index = _split_index(args["reference"])
    reference, voxel_size = _load_slice(ref_path, ref_index)

    if args["scale"] is not None:
        best_scale = args["scale"]
        logger.info("Using given scale: %.4f (skipping search)", best_scale)
    else:
        if args["tile"] is None:
            raise ValueError("--tile is required unless --scale is given.")
        tile_path, tile_index = _split_index(args["tile"])
        tile, _ = _load_slice(tile_path, tile_index)

        if args["downsample"] > 1:
            d = args["downsample"]
            reference_small = fourier_interpolate(reference, [s // d for s in reference.shape])
            tile_small = fourier_interpolate(tile, [s // d for s in tile.shape])
        else:
            reference_small, tile_small = reference, tile

        logger.info(
            "Coarse search: %d scales in [%.3f, %.3f]",
            args["coarse_steps"], args["min_scale"], args["max_scale"],
        )
        coarse_factors = np.linspace(args["min_scale"], args["max_scale"], args["coarse_steps"])
        best_coarse, coarse_iou, _ = search_scales(
            reference_small, tile_small, coarse_factors,
            args["maskthreshold"], args["maskabsolutethreshold"],
        )
        logger.info("Coarse best: scale=%.4f  IoU=%.4f", best_coarse, coarse_iou)

        fine_lo = max(args["min_scale"], best_coarse - args["fine_window"])
        fine_hi = min(args["max_scale"], best_coarse + args["fine_window"])
        logger.info("Refining: %d scales in [%.4f, %.4f]", args["fine_steps"], fine_lo, fine_hi)
        fine_factors = np.linspace(fine_lo, fine_hi, args["fine_steps"])
        best_scale, best_iou, _ = search_scales(
            reference_small, tile_small, fine_factors,
            args["maskthreshold"], args["maskabsolutethreshold"],
        )
        logger.info("Best scale: %.4f  (IoU=%.4f)", best_scale, best_iou)

    # Apply the found scale to the full-resolution reference.
    resized_full = zoom(reference, best_scale, order=1).astype(np.float32)

    output = args["output"] or os.path.join(
        os.path.dirname(os.path.abspath(ref_path)), "reference_resized.mrc"
    )
    with mrcfile.new(output, overwrite=True) as m:
        m.set_data(resized_full)
        if voxel_size is not None:
            m.voxel_size = voxel_size
    logger.info("Wrote %s  (scale=%.4f, shape=%s)", output, best_scale, resized_full.shape)


if __name__ == "__main__":
    main()
```
