## Supplementary material for "Whole cryo-FIB lamella acquisition at high magnification with rectangular beam montage cryo-electron tomography": Source code: roi_editor.py

```
#!/usr/bin/env python3
"""
AreTomo2 ROI file editor
========================
Interactive matplotlib GUI for creating and editing the patch-centre
ROI file used by AreTomo2's -RoiFile option.

Usage
-----
    python3 roi_editor.py stack.mrc [--ang angles.txt] [--roi existing.txt] [-o out.txt] [--bin 4]

Controls
--------
    Left-click  (empty area)   Add a patch centre
    Left-drag   (on point)     Move it
    Right-click (on point)     Delete it
    G / "Set grid" button      Replace all points with a regular grid
    S / Ctrl-S / Save button   Save ROI file
    ◀ / ▶ buttons or ←/→       Browse slices

ROI file format (AreTomo2 convention)
--------------------------------------
    Two whitespace-separated integer columns: X  Y
    One patch centre per row, no header.
    Coordinates are in full-resolution pixel space.  If the loaded MRC is
    itself binned (e.g. a bin4 preview stack), pass --bin to match — points
    are picked in the displayed (binned) image and scaled up by that factor
    when saved, and any --roi loaded is scaled back down for display.
    (default --bin: 4)
    Origin is BOTTOM-LEFT (MRC/IMOD convention: y=0 is the first row
    stored in the MRC file, which is displayed at the bottom).
    Points must be picked on the zero-tilt image — the GUI auto-selects
    that slice and marks it "[ZERO-TILT]" in the title.

Tile-grid overlay
-----------------
    Red dashed rectangles show the tile boundaries AreTomo2 would use
    when auto-generating the same NX×NY grid.  Tile size = imgW/NX × imgH/NY
    (matches CPatchTargets::DetectTargets).  The overlay updates whenever
    you change the grid spec.

Dependencies
------------
    pip install mrcfile matplotlib numpy
"""

import sys
import os
import argparse
from typing import Optional
import numpy as np
import matplotlib
matplotlib.use('TkAgg')   # change to Qt5Agg / GTK3Agg if TkAgg unavailable
import matplotlib.pyplot as plt
from matplotlib.patches import Rectangle
from matplotlib.widgets import Button, TextBox
import mrcfile


# ---------------------------------------------------------------------------
# File I/O helpers
# ---------------------------------------------------------------------------

def load_tilt_angles(angfile: str) -> np.ndarray:
    """Return 1-D array of tilt angles (degrees) from a text file."""
    angles = []
    with open(angfile) as fh:
        for line in fh:
            line = line.strip()
            if line and not line.startswith('#'):
                try:
                    angles.append(float(line.split()[0]))
                except ValueError:
                    pass
    return np.array(angles, dtype=float)


def zero_tilt_index(angles: np.ndarray) -> int:
    """Index of the tilt angle closest to 0°."""
    return int(np.argmin(np.abs(angles)))


def load_roi(path: str) -> list:
    """Load an existing ROI file → list of [x, y] int pairs."""
    pts = []
    with open(path) as fh:
        for line in fh:
            line = line.strip()
            if line and not line.startswith('#'):
                cols = line.split()
                if len(cols) >= 2:
                    pts.append([int(float(cols[0])), int(float(cols[1]))])
    return pts


def save_roi(points: list, path: str) -> None:
    """Write ROI file: two integer columns X Y, one patch per row."""
    with open(path, 'w') as fh:
        for x, y in points:
            fh.write(f"{int(x)}\t{int(y)}\n")
    print(f"[roi_editor] Saved {len(points)} patches → {path}")
    for i, (x, y) in enumerate(points):
        print(f"  {i+1:3d}   x={x:6d}   y={y:6d}")


def make_grid(nx: int, ny: int, img_w: int, img_h: int) -> list:
    """
    Regular patch-centre grid matching AreTomo2's CPatchTargets::DetectTargets
    (integer arithmetic throughout, as in the C++ source):
        iPatX  = imgW / nx          (integer division)
        iCentX = ix * iPatX + iPatX / 2
        iPatY  = imgH / ny
        iCentY = iy * iPatY + iPatY / 2
    iy=0 gives the BOTTOM row of tiles (y=0 at bottom-left / MRC origin).
    """
    pat_w = img_w // nx
    pat_h = img_h // ny
    pts = []
    for iy in range(ny):
        for ix in range(nx):
            cx = ix * pat_w + pat_w // 2
            cy = iy * pat_h + pat_h // 2
            pts.append([cx, cy])
    return pts


# ---------------------------------------------------------------------------
# Main editor
# ---------------------------------------------------------------------------

class RoiEditor:
    """Interactive patch-centre editor backed by matplotlib."""

    # Radius in screen pixels within which a click "hits" an existing point.
    PICK_RADIUS_PX = 14
    # Maximum display dimension in pixels (image downsampled for speed).
    MAX_DISPLAY_PX = 2048

    def __init__(
        self,
        mrc_path: str,
        ang_path: Optional[str] = None,
        roi_path: Optional[str] = None,
        out_path: Optional[str] = None,
        grid: tuple = (5, 3),
        bin: int = 4,
    ):
        self.mrc_path = mrc_path
        self.out_path = out_path or os.path.splitext(mrc_path)[0] + "_roi.txt"
        self.bin = bin

        # ---- open MRC (memory-mapped so large stacks stay on disk) ----------
        self._mrc = mrcfile.mmap(mrc_path, mode='r', permissive=True)
        raw = self._mrc.data
        if raw.ndim == 2:
            raw = raw[np.newaxis]           # single image → (1, ny, nx)
        self._raw = raw                     # shape (nz, ny, nx), NOT loaded yet
        self.nz, self.img_h, self.img_w = self._raw.shape

        # ---- display downsampling (coordinates are always full-resolution) --
        self._ds = max(1, max(self.img_w, self.img_h) // self.MAX_DISPLAY_PX)
        print(f"[roi_editor] Stack {self.img_w}×{self.img_h}×{self.nz}  "
              f"display scale 1/{self._ds}")
        print(f"[roi_editor] Input treated as bin{self.bin} — points are picked "
              f"in bin{self.bin} pixels and scaled ×{self.bin} to full-resolution "
              "on save (use --bin 1 if the input stack is already full-resolution)")

        # ---- tilt angles / zero-tilt slice ----------------------------------
        self.angles = None
        if ang_path and os.path.exists(ang_path):
            self.angles = load_tilt_angles(ang_path)
            self.slice_idx = zero_tilt_index(self.angles)
            print(f"[roi_editor] Zero-tilt slice: {self.slice_idx+1} "
                  f"({self.angles[self.slice_idx]:.1f}°)")
        else:
            self.slice_idx = self.nz // 2
            print(f"[roi_editor] No angle file — using middle slice "
                  f"{self.slice_idx+1}/{self.nz}")

        # ---- initial patch points -------------------------------------------
        self.grid_nx, self.grid_ny = grid
        if roi_path and os.path.exists(roi_path):
            full_res_pts = load_roi(roi_path)
            self.points = [[x / self.bin, y / self.bin] for x, y in full_res_pts]
            print(f"[roi_editor] Loaded {len(self.points)} points from {roi_path} "
                  f"(scaled to bin{self.bin} display)")
        else:
            self.points = make_grid(self.grid_nx, self.grid_ny,
                                    self.img_w, self.img_h)

        # ---- interaction state ----------------------------------------------
        self._drag_idx   = None   # index of point being dragged
        self._drag_moved = False  # distinguish drag from click

        self._build_figure()

    # ------------------------------------------------------------------ build

    def _build_figure(self):
        self.fig, self.ax = plt.subplots(figsize=(13, 8))
        self.fig.subplots_adjust(bottom=0.13)

        # Image — origin='lower' so y=0 is at the BOTTOM, matching MRC/AreTomo
        # convention.  extent maps data coordinates directly to full-res pixels.
        img_ds = self._get_slice_ds()
        self.im = self.ax.imshow(
            img_ds, cmap='gray', origin='lower',
            extent=[0, self.img_w, 0, self.img_h],
            aspect='equal', interpolation='nearest')
        self._auto_contrast(img_ds)

        # Tile-grid overlay (AreTomo2 auto-grid reference)
        self._tile_rects = []
        self._draw_tiles()

        # Patch-centre scatter
        xs, ys = self._xys()
        self.scat = self.ax.scatter(
            xs, ys, s=90, c='red', marker='o', zorder=5,
            linewidths=1.2, edgecolors='white')

        # Point-index labels
        self._pt_labels = []
        self._redraw_labels()

        self.ax.set_xlim(0, self.img_w)
        self.ax.set_ylim(0, self.img_h)
        self.ax.set_xlabel('X  (full-res pixels, origin bottom-left)', fontsize=9)
        self.ax.set_ylabel('Y  (full-res pixels, origin bottom-left)', fontsize=9)
        self._set_title()

        # ---- widget row -----------------------------------------------------
        ax_gval  = self.fig.add_axes([0.09, 0.025, 0.09, 0.05])
        ax_grid  = self.fig.add_axes([0.19, 0.025, 0.11, 0.05])
        ax_save  = self.fig.add_axes([0.45, 0.025, 0.11, 0.05])
        ax_prev  = self.fig.add_axes([0.65, 0.025, 0.05, 0.05])
        ax_next  = self.fig.add_axes([0.71, 0.025, 0.05, 0.05])
        ax_sinfo = self.fig.add_axes([0.77, 0.025, 0.20, 0.05])

        self.txt_grid  = TextBox(ax_gval, 'Grid ',
                                 initial=f'{self.grid_nx} {self.grid_ny}')
        self.btn_grid  = Button(ax_grid,  'Set grid')
        self.btn_save  = Button(ax_save,  'Save ROI')
        self.btn_prev  = Button(ax_prev,  '◀')
        self.btn_next  = Button(ax_next,  '▶')
        self.lbl_slice = Button(ax_sinfo, self._slice_label())
        self.lbl_slice.label.set_fontsize(8)

        self.btn_grid.on_clicked(self._cb_set_grid)
        self.btn_save.on_clicked(self._cb_save)
        self.btn_prev.on_clicked(self._cb_prev)
        self.btn_next.on_clicked(self._cb_next)

        # ---- canvas events --------------------------------------------------
        self.fig.canvas.mpl_connect('button_press_event',   self._on_press)
        self.fig.canvas.mpl_connect('motion_notify_event',  self._on_motion)
        self.fig.canvas.mpl_connect('button_release_event', self._on_release)
        self.fig.canvas.mpl_connect('key_press_event',      self._on_key)

        plt.show()
        self._mrc.close()

    # ---------------------------------------------------------------- drawing

    def _get_slice_ds(self) -> np.ndarray:
        """Return the current slice, downsampled for display."""
        slc = np.array(self._raw[self.slice_idx], dtype=np.float32)
        if self._ds > 1:
            slc = slc[::self._ds, ::self._ds]
        return slc

    def _auto_contrast(self, img: np.ndarray) -> None:
        lo, hi = np.percentile(img, (2, 98))
        self.im.set_clim(lo, hi)

    def _xys(self):
        if not self.points:
            return np.array([]), np.array([])
        a = np.array(self.points, dtype=float)
        return a[:, 0], a[:, 1]

    def _draw_tiles(self) -> None:
        """
        Redraw the auto-grid reference overlay.

        These rectangles show the tile boundaries AreTomo2 uses when it
        generates patch centres automatically (i.e. when NO -RoiFile is
        given).  When a ROI file IS provided, AreTomo ignores these tiles
        entirely — the points are just bare centre coordinates with no
        fixed region attached.  The overlay is a placement guide only.
        """
        for r in self._tile_rects:
            r.remove()
        self._tile_rects = []
        pw = self.img_w // self.grid_nx
        ph = self.img_h // self.grid_ny
        for iy in range(self.grid_ny):
            for ix in range(self.grid_nx):
                r = Rectangle(
                    (ix * pw, iy * ph), pw, ph,
                    linewidth=0.8, edgecolor='red', facecolor='red',
                    linestyle='--', alpha=0.06, zorder=2)
                self.ax.add_patch(r)
                rb = Rectangle(
                    (ix * pw, iy * ph), pw, ph,
                    linewidth=0.8, edgecolor='red', facecolor='none',
                    linestyle='--', alpha=0.55, zorder=3)
                self.ax.add_patch(rb)
                self._tile_rects.extend([r, rb])
        # Label the overlay so it's clear this is a reference, not owned regions
        if self._tile_rects:
            pw_f = self.img_w / self.grid_nx
            ph_f = self.img_h / self.grid_ny
            t = self.ax.text(
                pw_f * 0.02, ph_f * 0.02,
                f'auto-grid ref ({self.grid_nx}×{self.grid_ny})\n'
                f'tile {pw}×{ph} px (bin{self.bin})  '
                f'/  {pw * self.bin}×{ph * self.bin} px full-res\n'
                '(not used when ROI file given)',
                color='red', fontsize=6, alpha=0.8, zorder=8,
                verticalalignment='bottom')
            self._tile_rects.append(t)

    def _redraw_labels(self) -> None:
        for t in self._pt_labels:
            t.remove()
        self._pt_labels = []
        for i, (x, y) in enumerate(self.points):
            t = self.ax.text(x + self.img_w * 0.005, y + self.img_h * 0.005,
                             str(i + 1), color='magenta', fontsize=7,
                             zorder=7, clip_on=True)
            self._pt_labels.append(t)

    def _refresh(self) -> None:
        xs, ys = self._xys()
        if len(xs):
            self.scat.set_offsets(np.column_stack([xs, ys]))
        else:
            self.scat.set_offsets(np.empty((0, 2)))
        self._redraw_labels()
        self._set_title()
        self.fig.canvas.draw_idle()

    def _set_title(self) -> None:
        ang_str = ''
        if self.angles is not None and self.slice_idx < len(self.angles):
            ang_str = f'  {self.angles[self.slice_idx]:+.1f}°'
        zt = '  [ZERO-TILT ✓]' if self._is_zero_tilt() else ''
        tile_w = self.img_w // self.grid_nx
        tile_h = self.img_h // self.grid_ny
        self.ax.set_title(
            f'{os.path.basename(self.mrc_path)}  |  '
            f'slice {self.slice_idx+1}/{self.nz}{ang_str}{zt}  |  '
            f'{len(self.points)} patches  |  '
            f'tile size {tile_w}×{tile_h} px  |  '
            'right-click=delete · drag=move · click=add',
            fontsize=8.5)

    def _slice_label(self) -> str:
        if self.angles is not None and self.slice_idx < len(self.angles):
            return f'slice {self.slice_idx+1}/{self.nz}  ({self.angles[self.slice_idx]:.1f}°)'
        return f'slice {self.slice_idx+1}/{self.nz}'

    def _update_slice(self) -> None:
        img_ds = self._get_slice_ds()
        self.im.set_data(img_ds)
        self._auto_contrast(img_ds)
        self.lbl_slice.label.set_text(self._slice_label())
        self._set_title()
        self.fig.canvas.draw_idle()

    def _is_zero_tilt(self) -> bool:
        if self.angles is not None:
            return self.slice_idx == zero_tilt_index(self.angles)
        return self.slice_idx == self.nz // 2

    # --------------------------------------------------------- hit testing

    def _find_point(self, xdata: float, ydata: float) -> Optional[int]:
        """Return index of the nearest point within PICK_RADIUS_PX, else None."""
        if not self.points:
            return None
        # Convert screen-pixel radius to data-space units
        xlim = self.ax.get_xlim()
        ylim = self.ax.get_ylim()
        bbox = self.ax.get_window_extent()
        rx = self.PICK_RADIUS_PX * (xlim[1] - xlim[0]) / max(bbox.width,  1)
        ry = self.PICK_RADIUS_PX * (ylim[1] - ylim[0]) / max(bbox.height, 1)
        pts = np.array(self.points, dtype=float)
        dist = np.sqrt(((pts[:, 0] - xdata) / rx) ** 2 +
                       ((pts[:, 1] - ydata) / ry) ** 2)
        idx = int(np.argmin(dist))
        return idx if dist[idx] <= 1.0 else None

    # --------------------------------------------------------- event handlers

    def _on_press(self, event):
        if event.inaxes != self.ax or event.xdata is None:
            return

        hit = self._find_point(event.xdata, event.ydata)

        if event.button == 3:           # right-click → delete
            if hit is not None:
                del self.points[hit]
                self._refresh()
            return

        if event.button == 1:
            if hit is not None:         # start drag
                self._drag_idx   = hit
                self._drag_moved = False
            else:                       # add new point
                x = int(round(np.clip(event.xdata, 0, self.img_w - 1)))
                y = int(round(np.clip(event.ydata, 0, self.img_h - 1)))
                self.points.append([x, y])
                self._refresh()

    def _on_motion(self, event):
        if self._drag_idx is None or event.inaxes != self.ax:
            return
        if event.xdata is None:
            return
        x = int(round(np.clip(event.xdata, 0, self.img_w - 1)))
        y = int(round(np.clip(event.ydata, 0, self.img_h - 1)))
        self.points[self._drag_idx] = [x, y]
        self._drag_moved = True
        self._refresh()

    def _on_release(self, event):
        self._drag_idx   = None
        self._drag_moved = False

    def _on_key(self, event):
        k = event.key
        if k in ('s', 'ctrl+s'):
            self._cb_save(None)
        elif k in ('left', 'pageup'):
            self._cb_prev(None)
        elif k in ('right', 'pagedown'):
            self._cb_next(None)
        elif k == 'g':
            self._cb_set_grid(None)

    # --------------------------------------------------------- button callbacks

    def _cb_set_grid(self, _):
        raw = self.txt_grid.text.strip()
        try:
            parts = raw.split()
            nx, ny = int(parts[0]), int(parts[1])
            assert nx > 0 and ny > 0
        except Exception:
            print(f"[roi_editor] Invalid grid '{raw}' — use 'NX NY' e.g. '5 3'")
            return
        self.grid_nx, self.grid_ny = nx, ny
        self.points = make_grid(nx, ny, self.img_w, self.img_h)
        self._draw_tiles()
        self._refresh()

    def _cb_save(self, _):
        full_res_pts = [[x * self.bin, y * self.bin] for x, y in self.points]
        save_roi(full_res_pts, self.out_path)

    def _cb_prev(self, _):
        if self.slice_idx > 0:
            self.slice_idx -= 1
            self._update_slice()

    def _cb_next(self, _):
        if self.slice_idx < self.nz - 1:
            self.slice_idx += 1
            self._update_slice()


# ---------------------------------------------------------------------------
# Entry point
# ---------------------------------------------------------------------------

def main():
    ap = argparse.ArgumentParser(
        description='Interactive ROI file editor for AreTomo2 -RoiFile',
        formatter_class=argparse.RawDescriptionHelpFormatter,
        epilog=__doc__)
    ap.add_argument('mrc',
                    help='Input MRC tilt-series stack')
    ap.add_argument('--ang', '--angfile', dest='ang', default=None,
                    help='Tilt-angle file (one angle per line) — used to '
                         'auto-select the zero-tilt slice')
    ap.add_argument('--roi', dest='roi', default=None,
                    help='Existing ROI file to load as starting points')
    ap.add_argument('-o', '--out', dest='out', default=None,
                    help='Output ROI file path '
                         '(default: <mrc_stem>_roi.txt)')
    ap.add_argument('--grid', nargs=2, type=int, default=[5, 3],
                    metavar=('NX', 'NY'),
                    help='Initial patch grid when no --roi is given '
                         '(default: 5 3)')
    ap.add_argument('--bin', dest='bin', type=int, default=4,
                    help='Binning factor of the input MRC relative to the '
                         'full-resolution image AreTomo expects for -RoiFile. '
                         'Points are picked in the (binned) displayed image '
                         'and scaled up by this factor when saved; pass '
                         '--bin 1 if the input stack is already full-resolution '
                         '(default: 4)')
    args = ap.parse_args()

    RoiEditor(
        mrc_path=args.mrc,
        ang_path=args.ang,
        roi_path=args.roi,
        out_path=args.out,
        grid=tuple(args.grid),
        bin=args.bin,
    )


if __name__ == '__main__':
    main()
```
