## Supplementary material for "Whole cryo-FIB lamella acquisition at high magnification with rectangular beam montage cryo-electron tomography": Source code: sidecar.py

```
#!/usr/bin/env python3
"""Per-output provenance sidecars: one small JSON file beside each output.

See ``sidecar_metadata_plan.md``. Supersedes ``manifest.py``, which was 1037
lines almost entirely devoted to making one shared JSON+HDF5 file safe for
many processes to write — ``flock``'d transactions, a JSONL spool, a separate
``finalize`` sbatch pass, ``stale`` propagation across a fixed step graph. All
of that solved concurrent writes to one file. Sidecars remove the shared file,
so none of it is needed.

The rule is one writer per file. A step writes ``<output>.json`` next to what
it produced; when many processes contribute to one output — motion correction's
array tasks all writing z-slices of a shared per-tilt stack, stitch's array
tasks all writing z-slices of one canvas — each writes its own *keyed* sidecar
under ``<output>.sidecars/<key>.json`` instead. No two processes ever open the
same path, so there is nothing to lock and nothing to fold afterwards.

Step-level parameters (ROI, binning, rotation) are repeated in every keyed
sidecar rather than hoisted into a shared file. They are a couple of dozen
scalars, and the redundancy is the point: it is what keeps every sidecar
independently readable and every write uncontended.

Provenance is a chain, not a registry. Each sidecar's ``inputs`` name the files
it was derived from, fingerprinted, so :func:`chain` can walk backwards from a
tomogram to the raw frames by following pointers — and :func:`verify_inputs`
can say "this is not the file that made this". Nothing has to be kept current
by a writer that is not itself producing something.

Rule number one is inherited unchanged from the manifest: **record what the
step DID, not what it was asked.** ``effective`` holds the values in force
after defaults, ``auto`` resolution and clamping; ``requested`` (optional)
holds what was passed in. The two differing is the whole point.
"""

from __future__ import annotations

import hashlib
import json
import logging
import os
import socket
import sys
import tempfile
from datetime import datetime, timezone
from pathlib import Path
from typing import Any, Iterable, Iterator, Sequence

import numpy as np

log = logging.getLogger(__name__)

#: Bumped only for a change that an old reader could misread. Adding keys to
#: ``effective`` does not qualify; renaming or re-meaning one does.
SCHEMA = "montage-sidecar/1"

#: Pipeline step names. Unlike the manifest's PIPELINE_STEPS this is *not* an
#: order with stale-propagation attached — it is just the vocabulary, so a
#: typo'd step name is catchable. Order is documentation only.
STEPS = (
    "motion_correction",
    "stitch",
    "inpaint_mask",
    "inpaint_apply",
    "aretomo",
    "relion_export",
)

#: How many bytes of a file's head go into its fingerprint. Enough to cover an
#: MRC/TIFF header plus the start of the data — cheap on a shared filesystem,
#: while still catching a file regenerated with different parameters.
FINGERPRINT_BYTES = 65536

#: Consecutive value arguments beyond this many are collapsed to a count when
#: a command line is recorded (see :func:`summarise_argv`).
_ARGV_RUN_LIMIT = 6


# ---------------------------------------------------------------------------
# Fingerprints and paths
# ---------------------------------------------------------------------------

def fingerprint(path: str | os.PathLike, *, role: str | None = None) -> dict:
    """Cheap content fingerprint of one file: ``(size, mtime, sha1 of head)``.

    Deliberately not a full hash — these files run to tens of gigabytes and
    this is called once per input per step. Size plus the first
    :data:`FINGERPRINT_BYTES` catches a file that was rewritten with different
    parameters (different header, different shape, different first slice),
    which is the failure mode this is here to catch. It will not catch a
    change confined to the tail of a file of identical size.
    """
    p = Path(path)
    st = p.stat()
    h = hashlib.sha1()
    h.update(str(st.st_size).encode())
    with open(p, "rb") as fh:
        h.update(fh.read(FINGERPRINT_BYTES))
    out = {
        "path": str(p.resolve()),
        "size": st.st_size,
        "mtime": st.st_mtime,
        "sha1": h.hexdigest(),
    }
    if role:
        out["role"] = role
    return out


def canonical_path(path: str | os.PathLike) -> str:
    """One name per file, for comparing against a recorded ``path``.

    :func:`fingerprint` stores ``Path.resolve()``, so recorded paths have their
    symlinks resolved. Anything asking "is this the file that step recorded?"
    has to resolve too, or the same file reached by two names does not match —
    ``/home/user/project`` symlinked to ``/data/gpfs/.../project`` is the case
    that prompted this, and it silently made every input look unrecognised.
    """
    return str(Path(path).resolve())


def fingerprint_many(paths: Iterable[str | os.PathLike], *,
                     role: str | None = None) -> list[dict]:
    """Fingerprint several files, skipping (with a warning) any that are missing."""
    out = []
    for p in paths:
        try:
            out.append(fingerprint(p, role=role))
        except OSError as exc:
            log.warning("Cannot fingerprint %s: %s", p, exc)
    return out


def directory_input(directory: str | os.PathLike, *, pattern: str = "*",
                    role: str | None = None) -> dict:
    """Summarise a whole input directory instead of fingerprinting every file.

    Motion correction reads hundreds of raw frames per tile. Inlining hundreds
    of fingerprints into a per-tile JSON would make the sidecar bigger than
    anything it describes, for provenance nobody walks at that granularity —
    so a directory input is recorded as a path plus a count, which is what
    ``sidecar_metadata_plan.md`` settled on under "Open questions".
    """
    d = Path(directory)
    try:
        n = sum(1 for _ in d.glob(pattern))
    except OSError:
        n = -1
    out = {"path": str(d.resolve()), "kind": "directory",
           "pattern": pattern, "n_files": n}
    if role:
        out["role"] = role
    return out


# ---------------------------------------------------------------------------
# Sidecar locations
# ---------------------------------------------------------------------------

def sidecar_path(output: str | os.PathLike, key: str | None = None) -> Path:
    """Where the sidecar for ``output`` lives.

    ``key=None`` — one producer, one output — is ``<output>.json``. A key is
    for the many-writers-one-file case and gives
    ``<output>.sidecars/<key>.json``, one uncontended path per writer.

    The key is slugified, because tilt angles (``-7.5``) and tile stems are
    what get passed and neither is guaranteed to be a safe filename.
    """
    out = Path(output)
    if key is None:
        return out.with_name(out.name + ".json")
    return out.with_name(out.name + ".sidecars") / f"{_slug(key)}.json"


def _slug(key: str) -> str:
    """Filename-safe form of a sidecar key, preserving readability."""
    safe = "".join(c if (c.isalnum() or c in "+-._") else "_" for c in str(key))
    return safe or "_"


def sidecar_dir(output: str | os.PathLike) -> Path:
    """The ``<output>.sidecars/`` directory (may not exist)."""
    out = Path(output)
    return out.with_name(out.name + ".sidecars")


def find(output: str | os.PathLike, key: str | None = None) -> Path | None:
    """An existing sidecar for ``output``, or None.

    With no key this prefers the unkeyed ``<output>.json`` but falls back to
    *any* keyed sidecar, because step-level parameters are repeated in every
    one — so a reader that only wants "what binning made this canvas" does not
    have to know whether the step ran keyed or not.
    """
    direct = sidecar_path(output)
    if direct.exists():
        return direct
    if key is not None:
        keyed = sidecar_path(output, key)
        return keyed if keyed.exists() else None
    others = sorted(sidecar_dir(output).glob("*.json"))
    return others[0] if others else None


def find_all(output: str | os.PathLike) -> list[Path]:
    """Every sidecar for ``output``, unkeyed first then keyed, sorted."""
    out = []
    direct = sidecar_path(output)
    if direct.exists():
        out.append(direct)
    out.extend(sorted(sidecar_dir(output).glob("*.json")))
    return out


# ---------------------------------------------------------------------------
# Write / read
# ---------------------------------------------------------------------------

def write(
    output: str | os.PathLike,
    step: str,
    effective: dict,
    *,
    inputs: Sequence[dict] | None = None,
    key: str | None = None,
    requested: dict | None = None,
    extra: dict | None = None,
    reconstructed: bool = False,
    fingerprint_output: bool = True,
) -> Path:
    """Write ``output``'s sidecar and return where it went.

    ``inputs`` are fingerprint dicts from :func:`fingerprint`,
    :func:`fingerprint_many` or :func:`directory_input` — give them a ``role``
    when a step has more than one kind (inpainting takes a stack *and* a mask).

    ``reconstructed=True`` marks a sidecar assembled after the fact from
    surviving evidence rather than written by the run itself, e.g. by
    ``montage_backfill``. It is deliberately part of the document and not a
    footnote: a reconstructed record is an inference, and a reader weighing
    "is this the geometry that made this file" needs to know which it is
    holding.
    """
    if step not in STEPS:
        log.warning("Sidecar step %r is not one of %s — writing it anyway, but "
                    "check for a typo.", step, ", ".join(STEPS))

    path = sidecar_path(output, key)
    path.parent.mkdir(parents=True, exist_ok=True)

    doc: dict[str, Any] = {
        "schema": SCHEMA,
        "step": step,
        "output": canonical_path(output) if os.path.exists(output)
        else str(Path(output).absolute()),
        "key": key,
        "written_at": _now(),
        "provenance": _provenance(),
        "inputs": list(inputs or []),
        "effective": effective,
    }
    if requested is not None:
        doc["requested"] = requested
    if reconstructed:
        doc["reconstructed"] = True
    if extra:
        doc.update(extra)
    if fingerprint_output:
        try:
            doc["output_fingerprint"] = fingerprint(output)
        except OSError as exc:
            # Normal for a volume still on its way to that path (AreTomo
            # writing to scratch, rotation queued behind it) — record the
            # parameters now, fingerprint on a later pass.
            log.debug("No output fingerprint for %s yet: %s", output, exc)

    _atomic_write_json(path, doc)
    log.info("Sidecar: %s", path)
    return path


def read(path: str | os.PathLike) -> dict:
    """Read one sidecar by its own path."""
    with open(path) as fh:
        return json.load(fh)


def read_for(output: str | os.PathLike, key: str | None = None) -> dict | None:
    """The sidecar describing ``output``, or None if it has none."""
    found = find(output, key)
    return read(found) if found else None


def read_all_for(output: str | os.PathLike) -> list[dict]:
    """Every sidecar describing ``output`` — the keyed set, in key order."""
    return [read(p) for p in find_all(output)]


def _atomic_write_json(path: Path, doc: dict) -> None:
    """Write via a temp file and ``os.replace``, so a crash cannot truncate.

    There is no locking anywhere in this module — one writer per path is the
    whole design — but a job killed mid-write would still leave an unreadable
    sidecar, and that is cheap to rule out.
    """
    fd, tmp = tempfile.mkstemp(dir=str(path.parent), suffix=".tmp")
    try:
        with os.fdopen(fd, "w") as fh:
            json.dump(doc, fh, indent=2, sort_keys=False, default=_jsonable)
            fh.write("\n")
        os.replace(tmp, path)
    except BaseException:
        try:
            os.unlink(tmp)
        except OSError:
            pass
        raise


# ---------------------------------------------------------------------------
# Reading a chain
# ---------------------------------------------------------------------------

def input_paths(doc: dict, *, role: str | None = None) -> list[str]:
    """Paths this sidecar's step read, optionally just those in one role."""
    out = []
    for item in doc.get("inputs") or []:
        if role is not None and item.get("role") != role:
            continue
        if item.get("kind") == "directory":
            continue  # a directory summary is not a file to chain into
        if item.get("path"):
            out.append(item["path"])
    return out


def chain(output: str | os.PathLike, *, max_depth: int = 12) -> list[dict]:
    """Walk ``input`` pointers back from ``output``, nearest step first.

    Returns the sidecars encountered, deduplicated by path. Breadth-first, so
    a many-to-one step (stitch reading dozens of tiles) contributes its inputs
    at one level rather than recursing down the first one only.

    ``max_depth`` is a loop guard: nothing in this pipeline is more than about
    six steps deep, and a cycle would otherwise hang a diagnostic tool.
    """
    seen: set[str] = set()
    out: list[dict] = []
    frontier = [str(output)]
    for _ in range(max_depth):
        if not frontier:
            break
        nxt: list[str] = []
        for target in frontier:
            for path in find_all(target):
                if str(path) in seen:
                    continue
                seen.add(str(path))
                doc = read(path)
                out.append(doc)
                nxt.extend(input_paths(doc))
        # Distinct upstream files only: 109 tile sidecars naming the same
        # frame directory should not be walked 109 times.
        frontier = list(dict.fromkeys(nxt))
    return out


def verify_inputs(doc: dict) -> list[str]:
    """Complaints about ``doc``'s inputs as they stand on disk *now*.

    This is the check the ``-Patch 5 3`` vs ``-Patch 10 6`` mix-up needed and
    never had: it says "the file that made this is not the file at that path
    any more", which no amount of care at the call site can notice.

    A fingerprint that matches proves the file is unchanged since the step ran.
    A mismatch does not prove which of the two is wanted — only that they
    differ, which is exactly when a human should look.
    """
    problems = []
    for item in doc.get("inputs") or []:
        if item.get("kind") == "directory" or not item.get("path"):
            continue
        path = item["path"]
        if not os.path.exists(path):
            problems.append(f"missing: {path}")
            continue
        if item.get("sha1") is None:
            continue
        now = fingerprint(path)
        if now["sha1"] != item["sha1"] or now["size"] != item.get("size"):
            problems.append(
                f"changed since the step ran: {path} "
                f"(recorded {item.get('size')} B, now {now['size']} B)")
    return problems


# ---------------------------------------------------------------------------
# Small shared helpers (moved here from manifest.py, which this replaces)
# ---------------------------------------------------------------------------

def _now() -> str:
    return datetime.now(timezone.utc).isoformat(timespec="seconds")


def _jsonable(obj: Any) -> Any:
    """Fallback encoder: numpy scalars/arrays and Paths appear in records."""
    if isinstance(obj, (np.integer,)):
        return int(obj)
    if isinstance(obj, (np.floating,)):
        return float(obj)
    if isinstance(obj, (np.bool_,)):
        return bool(obj)
    if isinstance(obj, np.ndarray):
        return obj.tolist()
    if isinstance(obj, Path):
        return str(obj)
    raise TypeError(f"Not JSON-serialisable: {type(obj)}")


def summarise_argv(argv: Sequence[str]) -> list[str]:
    """Collapse long runs of value arguments, e.g. a chunk of input paths.

    An array task's argv is *not* the step's input list — it is one chunk of
    it — and spelling out fifty paths invites exactly that misreading. Runs
    longer than ``_ARGV_RUN_LIMIT`` keep their first and last entry with a
    count in between; the authoritative list is ``inputs``.
    """
    summarised: list[str] = []
    run: list[str] = []

    def flush() -> None:
        if len(run) > _ARGV_RUN_LIMIT:
            summarised.extend([run[0], f"... {len(run) - 2} more ...", run[-1]])
        else:
            summarised.extend(run)
        run.clear()

    for arg in argv:
        if arg.startswith("-"):
            flush()
            summarised.append(arg)
        else:
            run.append(arg)
    flush()
    return summarised


def _provenance() -> dict:
    """Who wrote this sidecar, from where, with what version of the code."""
    try:
        from importlib.metadata import version
        pkg_version = version("SquareBeamMontaging")
    except Exception:
        pkg_version = "unknown"
    return {
        "host": socket.gethostname(),
        "user": os.environ.get("USER", "unknown"),
        "package_version": pkg_version,
        "slurm_job": os.environ.get("SLURM_ARRAY_JOB_ID")
        or os.environ.get("SLURM_JOB_ID"),
        "command": summarise_argv(sys.argv),
        "cwd": os.getcwd(),
    }


# ---------------------------------------------------------------------------
# montage_provenance CLI — read-only, nothing in the pipeline depends on it
# ---------------------------------------------------------------------------

def _cmd_show(args) -> int:
    docs = chain(args.output) if args.chain else [
        read(p) for p in find_all(args.output)]
    if not docs:
        print(f"No sidecar for {args.output}")
        return 1
    if args.json:
        json.dump(docs, sys.stdout, indent=2, default=_jsonable)
        print()
        return 0

    for doc in docs:
        recon = "  [RECONSTRUCTED]" if doc.get("reconstructed") else ""
        print(f"\n=== {doc.get('step', '?')}{recon} "
              f"key={doc.get('key')!r} ===")
        print(f"  output   {doc.get('output')}")
        prov = doc.get("provenance") or {}
        print(f"  written  {doc.get('written_at')}  "
              f"job={prov.get('slurm_job')}  by {prov.get('user')}")
        for item in doc.get("inputs") or []:
            role = f"[{item['role']}] " if item.get("role") else ""
            if item.get("kind") == "directory":
                print(f"  input    {role}{item['path']}  "
                      f"({item.get('n_files')} files matching "
                      f"{item.get('pattern')})")
            else:
                print(f"  input    {role}{item.get('path')}")
        for k, v in (doc.get("effective") or {}).items():
            text = repr(v)
            if len(text) > 88:
                text = text[:85] + "..."
            print(f"    {k:32} {text}")
    return 0


def _cmd_verify(args) -> int:
    docs = [read(p) for p in find_all(args.output)]
    if not docs:
        print(f"No sidecar for {args.output}")
        return 1
    bad = 0
    for doc in docs:
        problems = verify_inputs(doc)
        label = f"{doc.get('step')} key={doc.get('key')!r}"
        if problems:
            bad += 1
            print(f"{label}: {len(problems)} problem(s)")
            for p in problems:
                print(f"    {p}")
        elif args.verbose:
            print(f"{label}: inputs match")
    if not bad:
        print(f"All {len(docs)} sidecar(s) for {os.path.basename(args.output)} "
              f"have inputs matching what is on disk.")
    return 1 if bad else 0


def main() -> None:
    import argparse

    p = argparse.ArgumentParser(
        prog="montage_provenance",
        description="Read provenance sidecars written beside pipeline outputs. "
                    "Read-only: nothing in the pipeline depends on this.")
    sub = p.add_subparsers(dest="cmd", required=True)

    s = sub.add_parser("show", help="print the sidecar(s) for a file")
    s.add_argument("output", help="the OUTPUT file, not its .json")
    s.add_argument("--chain", action="store_true",
                   help="follow inputs back through every upstream step")
    s.add_argument("--json", action="store_true")
    s.set_defaults(func=_cmd_show)

    v = sub.add_parser("verify", help="check inputs still match their fingerprints")
    v.add_argument("output", help="the OUTPUT file, not its .json")
    v.add_argument("-v", "--verbose", action="store_true")
    v.set_defaults(func=_cmd_verify)

    args = p.parse_args()
    logging.basicConfig(level=logging.WARNING, format="%(message)s")
    raise SystemExit(args.func(args))


if __name__ == "__main__":
    main()
```
