## Supplementary material for "Whole cryo-FIB lamella acquisition at high magnification with rectangular beam montage cryo-electron tomography": Source code: Simulate_dose.py

```
import argparse
from Utilities import *
from mpl_toolkits.axes_grid1.anchored_artists import AnchoredSizeBar
from scipy.optimize import curve_fit


def _gaussian(x, amplitude, mean, sigma):
    return amplitude * np.exp(-0.5 * ((x - mean) / sigma) ** 2)


def read_tilts_and_image_shifts_from_file(filename):
    """
    Reads the tilt angles and image shifts from a file and returns them as numpy arrays.

    Parameters
    ----------
    filename : str
        The path to the file containing the tilt angles and image shifts.

    Returns
    -------
    tilts : ndarray
        An array of tilt angles.
    imageshifts : list of ndarray
        A list of arrays, each containing the image shifts for a particular tilt.
    zs : list of ndarray
        A list of arrays, each containing the z values for a particular tilt.
    """
    # Read lines into a list
    with open(filename, "r", encoding="utf-8") as f:
        content = f.read().strip().split("\n")

    # Number of tilts is the first item in file
    num_tilts = int(content[0])
    tilts = []
    imageshifts = []
    zs = []

    # index tracks our place in the file as we consume it tilt-block by tilt-block
    index = 1
    for _ in range(num_tilts):
        # First line of a tilt block is the tilt angle itself
        tilt = float(content[index])
        tilts.append(tilt)
        # Second line is how many image-shift rows follow for this tilt
        num_shifts = int(content[index + 1])
        shifts = []
        z_values = []
        for i in range(num_shifts):
            # Each row is a shift coordinate with a trailing z value in its last column
            shift_data = list(map(float, content[index + 2 + i].split()))
            shifts.append(shift_data[:-1])
            z_values.append(shift_data[-1])
        imageshifts.append(np.array(shifts))
        zs.append(np.array(z_values))
        # Advance past the tilt line, the count line, and all of its shift rows
        index += 2 + num_shifts

    return np.array(tilts), imageshifts, zs

def compute_canvas_geometry(imageshifts,beam,beampixelsize,maxgridshape=[2048,2048]):
    """
    Determines the canvas pixel size and shape needed to fit every tile in
    imageshifts (across all of its tilts) without clipping.

    Kept separate from simulate_dose so that callers building up a dose map
    incrementally (e.g. one additional tilt at a time) can compute this once
    from the full set of tilts and reuse the same canvas for every call -
    otherwise each call would size its own canvas from whatever subset of
    tiles it was given, so the pixel size (and anything measured in canvas
    pixels, like a scale bar) would drift slightly from call to call.

    Parameters
    ----------
    imageshifts : list of ndarray
        A list of arrays, each containing the image shifts for a particular tilt in camera pixels.
    beam : ndarray
        An image of the beam
    beampixelsize : float
        The pixel size for the beam profile.
    maxgridshape : list
        Upper bound on the number of pixels along each grid dimension. The
        canvas pixel size is chosen so that the larger physical extent of the
        tiles maps to no more than this many pixels, and the grid itself is
        then sized exactly to the data (per axis) so every tile fits without
        clipping.

    Returns
    -------
    min_shift : ndarray
        The minimum image shift (in Angstrom), used as the canvas origin.
    pixel_size : float
        Physical size of one canvas pixel, in Angstrom.
    gridshape : list
        Shape of the canvas, in pixels.
    """
    # Determine real space sizing of beam shift ipixels
    beamshape = np.asarray(beam.shape)
    min_shift = np.min([np.min(shift,axis=0) for shift in imageshifts],axis=0) #- beamshape//2
    max_shift = np.max([np.max(shift,axis=0) for shift in imageshifts],axis=0) + beamshape
    # Convert to Angstrom
    min_shift = min_shift * beampixelsize
    max_shift = max_shift * beampixelsize
    ptp_shift = max_shift - min_shift

    # Pixel size is set by whichever axis is most constrained by maxgridshape,
    # then the grid is sized exactly to the data so no axis is clipped or
    # padded out to a fixed square. ptp_shift is reversed here because its
    # components are (y, x) (see y0, x0 = position.astype(int) below) while
    # gridshape/dose axes are ordered (x, y), matching how dose is indexed
    # further down as dose[cx0:cx1, cy0:cy1].
    pixel_size = max(ptp_shift[::-1] / np.asarray(maxgridshape))
    gridshape = (np.ceil(ptp_shift[::-1] / pixel_size).astype(int) + 1).tolist()

    return min_shift, pixel_size, gridshape

def simulate_dose(tilts,imageshifts,beam,beampixelsize,eperA2,maxgridshape=[2048,2048],geometry=None):
    """
    Simulates the dose for a set of tilt angles and image shifts.

    Parameters
    ----------
    tilts : ndarray
        An array of tilt angles.
    imageshifts : list of ndarray
        A list of arrays, each containing the image shifts for a particular tilt in camera pixels.
    beam : ndarray
        An image of the beam
    beampixelsize : float
        The pixel size for the beam profile.
    eperA2 : float
        The number of electrons per Angstrom squared in the beam.
    maxgridshape : list
        Upper bound on the number of pixels along each grid dimension, used
        only when geometry is not supplied (see compute_canvas_geometry).
    geometry : tuple, optional
        (min_shift, pixel_size, gridshape) as returned by
        compute_canvas_geometry. Pass this in when calling simulate_dose
        repeatedly for growing subsets of the same tilt series, so every call
        shares one fixed canvas instead of each one auto-fitting its own
        (which would make the pixel size, and the scale bar, drift between
        calls). If omitted, the canvas is auto-fit to just this call's tiles.

    Returns
    -------
    dose : ndarray
        The dose for each pixel in the grid.
    """
    beamshape = np.asarray(beam.shape)
    if geometry is None:
        geometry = compute_canvas_geometry(imageshifts,beam,beampixelsize,maxgridshape)
    min_shift, pixel_size, gridshape = geometry

    # Canvas that will accumulate dose contributions from every tile
    dose = np.zeros(gridshape)

    # Canvas-origin offset in output pixels, for reference (not used further below)
    origin = min_shift/pixel_size

    # Resize beam to match the shift range
    newsize = (beamshape * beampixelsize / pixel_size).astype(int)

    # Resample the beam profile onto the canvas pixel grid...
    beam = fourier_interpolate(beam,newsize)
    # ...then rebuild the mask so ringing introduced by the resampling doesn't leak into the dose map
    beam = make_mask(beam,shrinkn = False)

    for i,tilt in enumerate(tilts):
        # Dilate beam to match tilted grid size
        tiltedbeamshape = (newsize*np.asarray([1,1/np.cos(np.deg2rad(tilt))])).astype(int)
        # Dose per pixel drops by cos(tilt) since the same beam current is spread over a larger, foreshortened area
        dilatedbeam = fourier_interpolate(beam,tiltedbeamshape)*eperA2*np.cos(np.deg2rad(tilt))


        for j in range(imageshifts[i].shape[0]):
            
            position = (imageshifts[i][j]*beampixelsize - min_shift)/pixel_size
            # Desired coordinate of upper left of tile
            y0, x0 = position.astype(int)
        

            # Skip tiles that have fallen off canvas
            if x0 > gridshape[0] or y0 > gridshape[1]:
                continue
            if x0 +tiltedbeamshape[0] <0 or y0 +tiltedbeamshape[1] <0:
                continue

            # Truncate canvas coordinate beginnings to be >= 0
            cy0, cx0 = [max(coord, 0) for coord in [y0, x0]]
            # Truncate canvas coordinate maximum to be <= canvas array limits
            cx1, cy1  = [
                min(coord, limit) for coord, limit in zip([x0 + tiltedbeamshape[0],y0 + tiltedbeamshape[1] ], gridshape)
            ]

            # Size of tile that will make it onto the montage canvas
            X = cx1 - cx0
            Y = cy1 - cy0

            # Coordinates of tile, if x0 (or y0) < 0 this implies some of the tile
            # falls off the left (or upper) edge of canvas so is not included
            tx0 = -min(x0,0)
            tx1 = tx0 + X
            ty0 = -min(y0,0)
            ty1 = ty0 + Y
            # Add this tile's contribution onto the shared canvas, accumulating dose where tiles overlap
            dose[cx0:cx1, cy0:cy1] += dilatedbeam[tx0:tx1,ty0:ty1]
            # overlap[cx0:cx1, cy0:cy1] += np.where(msk, np.uint8(1), np.uint8(0))[tx0:tx1,ty0:ty1]

    # Single panel: dose map with the histogram and colorbar inset on top of it
    fig, ax = plt.subplots(figsize=(6, 6))
    # Cap the colour scale at the highest dose actually reached
    vmax = np.max(dose)
    # Target dose if every pixel received uniform coverage across all tilts
    desired_dose = eperA2*len(tilts)

    # Sequential colormap since we're showing absolute dose, not over/under exposure
    # Flip of y-axis since mrc convention is opposite to numpy/matplotlib
    im = ax.imshow(dose[::-1], cmap='plasma', vmin=0, vmax=vmax)
    # Axis ticks are just pixel indices, not physically meaningful, so hide them and rely on the scale bar instead
    ax.axis('off')

    # Scale bar sized to ~20% of the canvas width, labelled in the nearest
    # convenient physical unit (nm below 1 um, otherwise um)
    bar_pixels = 0.2 * dose.shape[1]
    bar_angstrom = bar_pixels * pixel_size
    if bar_angstrom < 1e4:
        bar_label = f'{bar_angstrom / 10:.0f} nm'
    else:
        bar_label = f'{bar_angstrom / 1e4:.1f} µm'
    # Anchored to a point in data (pixel) coordinates rather than the default
    # axes-fraction position, so the bar sits inside the actual image content
    # instead of the whitespace matplotlib adds to letterbox a non-square image
    margin_x = 0.05 * dose.shape[1]
    margin_y = 0.05 * dose.shape[0]
    scalebar = AnchoredSizeBar(
        ax.transData, bar_pixels, bar_label, loc='lower right',
        bbox_to_anchor=(dose.shape[1] - margin_x, dose.shape[0] - margin_y),
        bbox_transform=ax.transData,
        pad=0.3, borderpad=0, color='white', frameon=False, size_vertical=0.01 * dose.shape[0],
    )
    ax.add_artist(scalebar)

    # Histogram inset: bars area is width*height of the axes, kept at <=15%
    # (here ~9%), sat in the top-right corner away from the scale bar, with a
    # semi-transparent panel so the dose map stays visible behind it
    hist_ax = ax.inset_axes([0.50, 0.72, 0.46, 0.20], transform=ax.transAxes)
    hist_ax.set_facecolor((1, 1, 1, 0.55))
    # A thin colorbar sits immediately below the histogram, sharing its x-axis
    # extent, so it doubles as the histogram's dose axis instead of a separate
    # one taking up more space
    cbar_ax = ax.inset_axes([0.50, 0.685, 0.46, 0.03], transform=ax.transAxes)

    # 'sturges' picks the bin count from the sample size alone, so it stays
    # sensible for both a handful of tilts (where 100 fixed bins leaves mostly
    # empty gaps between narrow overlap-count spikes) and many tilts (where
    # 100 bins is noisier than the underlying distribution warrants).
    # density=True normalises the bars so they integrate (sum(height*width)) to 1
    counts, bin_edges, patches = hist_ax.hist(dose.flatten(), bins='sturges', density=True)
    bin_centers = 0.5 * (bin_edges[:-1] + bin_edges[1:])
    bin_width = bin_edges[1] - bin_edges[0]
    # Colour each bar by its dose value, using the same colormap/scale as the dose map
    for patch, center in zip(patches, bin_centers):
        patch.set_facecolor(im.cmap(im.norm(center)))

    hist_ax.axvline(desired_dose, color='k', linestyle='dashed', linewidth=1)
    hist_ax.set_xlim(0, vmax)
    # The dose axis is carried by the colorbar underneath, so hide the histogram's own
    hist_ax.tick_params(axis='x', bottom=False, labelbottom=False)
    hist_ax.set_ylabel('Fraction of pixels', fontsize=7)
    hist_ax.tick_params(labelsize=6)

    cbar = fig.colorbar(im, cax=cbar_ax, orientation='horizontal')
    cbar.set_label('Dose (e/Å$^2$)', fontsize=7)
    cbar.ax.tick_params(labelsize=6)

    # The background spike at dose ~ 0 dwarfs the rest of the distribution, so
    # scale the y-axis off the next largest bin instead of that spike
    near_zero = bin_centers < 0.1 * desired_dose
    visible_counts = counts[~near_zero]
    if visible_counts.size:
        hist_ax.set_ylim(0, visible_counts.max() * 1.2)

    # Fit a Gaussian to the well-exposed peak just to the right of the target
    # dose line, to quantify the mean/spread of dose where tiles fully overlap
    peak_mask = bin_centers >= desired_dose
    if peak_mask.sum() >= 3:
        peak_centers = bin_centers[peak_mask]
        peak_counts = counts[peak_mask]
        p0 = [peak_counts.max(), peak_centers[np.argmax(peak_counts)], np.ptp(peak_centers) / 6]
        try:
            # Lower-bound sigma at half a bin width; otherwise, when the peak is
            # only 1-2 bins wide (few tilts), curve_fit can "succeed" by collapsing
            # onto a spike-thin, meaningless width
            popt, _ = curve_fit(
                _gaussian, peak_centers, peak_counts, p0=p0,
                bounds=([0, peak_centers.min(), bin_width / 2], [np.inf, peak_centers.max(), np.ptp(peak_centers)]),
            )
            fit_mean, fit_sigma = popt[1], abs(popt[2])
            # Plot the fit curve across its own +/-5 sigma range rather than just the
            # (possibly narrower) bin range it was fitted to, clipped to non-negative dose
            fit_x = np.linspace(max(fit_mean - 5 * fit_sigma, 0), fit_mean + 5 * fit_sigma, 200)
            hist_ax.plot(fit_x, _gaussian(fit_x, *popt), 'r-', linewidth=1)
            hist_ax.text(
                0.97, 0.92, f'μ={fit_mean:.1f}\nσ={fit_sigma:.1f}', transform=hist_ax.transAxes,
                ha='right', va='top', fontsize=6, color='r',
            )
            # print(f'Peak dose (Gaussian fit): mean={fit_mean:.2f}, std={fit_sigma:.2f} e/Angstrom^2')
        except RuntimeError:
            print('Gaussian fit to dose peak did not converge')

    # plt.show()
    return dose,pixel_size,fig

if __name__=='__main__':

    parser = argparse.ArgumentParser(description='Simulate accumulated dose across a tilt series montage.')
    parser.add_argument('imageshifts', help='Path to the image shifts file (see read_tilts_and_image_shifts_from_file)')
    parser.add_argument('beamfilename', help='Path to the reference beam profile .mrc file')
    parser.add_argument('--dose', type=float, default=100,
                         help='Total dose in electrons per Angstrom^2 across the whole tilt series (default: 100)')
    args = parser.parse_args()

    filename = args.imageshifts
    beamfilename = args.beamfilename

    # Read tilt angles and image shifts
    tilts,imageshifts, _ = read_tilts_and_image_shifts_from_file(filename)
    # imageshifts /=2
    # Load beam profile
    beam = np.asarray(mrcfile.open(beamfilename,'r').data)
    # Native beam pixel size, kept fixed across iterations below (simulate_dose returns
    # a different, per-call canvas pixel_size that must not be fed back in as this)
    beampixelsize = mrcfile.open(beamfilename,'r').voxel_size.x
    # Split the total dose evenly across every tilt in the series
    eperA2 = args.dose/len(tilts)

    # Fix the canvas geometry once from the full tilt series, so every frame below
    # shares the same pixel size and scale bar reference as tiles are progressively
    # added, instead of each frame auto-fitting (and drifting) its own canvas
    geometry = compute_canvas_geometry(imageshifts,beam,beampixelsize)

    # Re-run for each successive subset of tilts so we can watch dose accumulate frame by frame
    for i in tqdm(np.arange(1,len(tilts)+1),desc='tilt'):
        dose,pixel_size,fig = simulate_dose(tilts[:i],imageshifts[:i],beam,beampixelsize,eperA2,geometry=geometry)
        # plt.show()
        # Keep one frame per cumulative tilt count for a dose_1.png ... dose_N.png sequence
        fig.savefig('dose/dose_{0}.pdf'.format(i))
        plt.close(fig)
    # Only the final (all-tilts) dose map from the loop above is kept for the saved outputs below
    np.save('dose.npy',dose)
    # Save dose
    m = mrcfile.new('dose.mrc',data=dose.astype(np.float32),overwrite=True)
    # m._set_voxel_size(pixel_size,pixel_size,1)
    m.close()

    
```
