## Supplementary material for "Whole cryo-FIB lamella acquisition at high magnification with rectangular beam montage cryo-electron tomography": Source code: smoothn.py

```
# -*- coding: utf-8 -*-
"""
Implementation of smoothn(y)
"""
__all__ = [
    'smoothn'
]
# ====== IMPORT LIBRARIES ======

import numpy as np
from scipy.fft import dctn, idctn
from scipy.optimize import minimize
from scipy.ndimage.morphology import distance_transform_edt


# ================================================================================================
def smoothn(y, s=None, tolz=1e-3, z0=None, w=None, di=None,
            robust=False, max_iter=1000, robust_iter=3, workers=1,
            return_s=False):
    """
    Fast smooths an array of data based on the cosine discrete transform. Allows to choose the smoothing parameter,
    or calculate it using a GCVscore.
    Also, it implements a robust iteration (False by default) to ignore outlier points.

    Parameters
    ----------
    y : np.ma.MaskedArray or np.ndarray
        Data to be smoothed and filled.
    s : None or float
        Smoothing parameter, calculated automatically if not provided to minimize gcv score.
    tolz : float
        Tolerance of iteration over z (prediction of data).
    z0 : np.ndarray
        Initial guess for z, calculated with nearest neighbor if not provided
    w : np.ndarray
        Weights array, if not provided assumes all data has the same confidence
    di : tuple
        Grid dimensions, assumed regular if not given
    robust : bool
        Indicates if the robust iteration is executed
    workers : int
        Number of workers assigned to Scipy functions `fft.dctn` and `fft.idctn`
    max_iter : int
        Maximum number of iterations. Default: 1,000
    robust_iter : int
        Number of robust iterations to perform. Notice that the max_iter value works inside the number of robust
        iterations, leading to a maximum of `robust_iter * max_iter` iterations. Default: 3
    return_s : bool
        Indicates if s value should be returned. Useful for fine-tuning after calculation.

    Returns
    -------
    np.ndarray
        Gives the smoothed gridded data, with missing values filled with the prediction done

    References
    ----------
    [1] Garcia D. (2010) Robust smoothing of gridded data in one and higher dimensions with missing values.
    Computational Statistics & Data Analysis, 2010; 54:1167-1178.
    """
    # Create MaskedArray in case it contains (NaN, Inf)
    y = to_masked(y)

    # Define dimensions
    n = y.size
    d = y.ndim
    shape = y.shape
    nf = y.count()

    # Create spacing vector
    if di is None:
        di = np.ones(d)
    di /= np.max(di)

    # Create weights matrix
    if w is None:
        w = np.ones(shape)
    w = w/np.max(w) * ~y.mask
    if np.any(w < 0):
        raise ValueError('All weights must be >=0')

    # Define if it has a weight matrix and if the smoothing factor needs to be calculated
    isweight = np.any(w < 1)
    isauto = s is None

    # Create Lambda tensor (see reference) and calculate Gamma if S is given
    lamb = create_lambda(y, di)
    if not isauto:
        gamm = 1 / (1 + s * lamb ** 2)
        s_lim = None
    else:
        # Set limit values for S to be calculated
        gamm = None
        h_lim = np.array([1-1e-5, 1e-5])
        s_lim = (((1 + np.sqrt(1 + 8 * h_lim ** (2 / d))) / (4 * h_lim ** (2 / d))) ** 2 - 1) / 16

    # ===== Initial values for iteration =====
    w_tot = w.copy()
    # Initial condition for z if is weighted
    if isweight:
        # Initialize z
        if z0:
            z = z0.copy()
        else:
            z = initz(y)
    else:
        z = np.zeros(shape)

    z0 = z

    y_fill = y.filled(0.0)  # Assign arbitrary value to missing data

    # ===== START ITERATIONS =====

    for i in range(robust_iter):

        aow = np.sum(w_tot)/n
        miss = np.inf

        for k in range(max_iter):

            if miss < tolz:
                break

            # Calculate discrete cosine transform
            xdc = w_tot * (y_fill - z) + z
            dcty = dctn(xdc, norm='ortho', workers=workers)

            # Estimate value of s
            if isauto:
                s = minimize(gcv_score, x0=np.array(1.0),
                             args=(y, w_tot, n, nf, dcty, lamb, aow), bounds=[s_lim]).x
                gamm = 1 / (1 + s * lamb ** 2)

            # Calculate inverse discrete cosine transform
            z = idctn(gamm * dcty, norm='ortho', workers=workers)

            # Check jump, if it's less than tolerance iterations are terminated
            res = z0-z
            miss = isweight * np.sqrt(np.sum(res ** 2)) / np.sqrt(np.sum(z ** 2))
            z0 = z

        if robust:
            u = studentized_residuals(y_fill, z, ~y.mask, s)
            w_tot = w * biweight(u)
            isweight = True
        else:
            break

    return z if not return_s else (z, s)


# ================================================================================================
def gcv_score(s, y, w_tot, n, nf, dcty, lamb, aow=None):
    """
    Calculates the Generalised Cross-Validation (GCV) score based on the weights assigned to the data.

    Parameters
    ----------
    s : float
        The smoothing parameter used in the algorithm.
    y : np.ma.MaskedArray
        The input data as a masked array.
    w_tot : np.ndarray
        The array of weights assigned to the data.
    n : int
        The size of the input data array, not calculated to save resources.
    nf : int
        The number of finite elements in the input data array, not calculated to save resources.
    dcty : np.ndarray
        The discrete cosine transform of the input data, considering the prediction and weights.
    lamb : np.ndarray
        The lambda tensor of the input data array.
    aow : Optional[float], default=None
        The amount of weight in the input data, which is the mean of the weights.

    Returns
    -------
    float
        The calculated GCV score for the input data based on the assigned weights.

    Notes
    -----
    The GCV score is a statistical measure used for selecting the optimal smoothing parameter in
    non-parametric regression problems, such as smoothing splines. The method is based on the principle
    of minimizing the expected mean squared error (MSE) of the model, and it is widely used in signal
    processing, computer vision, and machine learning applications.
    """
    gamm = 1 / (1 + s * lamb ** 2)

    if aow > 0.9:
        a = dcty * (gamm - 1)
        rss = np.sum(a ** 2)
    else:
        y_hat = idctn(dcty * gamm, norm='ortho')
        dy = y.compressed() - y_hat[~y.mask]
        rss = np.sum((np.sqrt(w_tot[~y.mask]) * dy) ** 2)

    trh = gamm.sum()
    gcv = rss / nf / (1 - trh / n) ** 2
    return gcv


# ================================================================================================
def to_masked(y):
    """
    Converts an input array into a masked array with NaN values masked.

    Parameters
    ----------
    y : ndarray
        The input array to convert.

    Returns
    -------
    ma.MaskedArray
        The masked array with NaN values masked.
    """
    if not np.ma.isMaskedArray(y):
        y = np.ma.masked_invalid(y)
    return y


# ================================================================================================
def create_lambda(y, dx):
    """
    Creates a Lambda tensor for the `smoothn` function. It contains the eigenvalues of the difference matrix used in
    the penalized least-squares process.

    Parameters
    ----------
    y : np.ndarray
        The data to process.
    dx : tuple of float
        The spatial difference between steps in each dimension.

    Returns
    -------
    np.ndarray
        The Lambda tensor

    References
    ----------
    * Per Christian Hansen, "Rank-Deficient and Discrete Ill-Posed Problems: Numerical Aspects of Linear Inversion",
    SIAM, 1998.
    """
    d = y.ndim
    lamb = np.zeros(y.shape)
    for i in range(d):
        siz0 = np.ones(d, dtype=int)
        siz0[i] = y.shape[i]
        lamb += (2 - 2 * np.cos(np.pi * np.arange(y.shape[i]).reshape(siz0) / y.shape[i])) / dx[i] ** 2
    return lamb


# ================================================================================================
def initz(y):
    """
    Initialize the estimation values of y using nearest interpolation. Implemented for 1D, 2D and 3D.

    Parameters
    ----------
    y : np.ma.MaskedArray
        Input masked array of shape (n,) for 1D, (m, n) for 2D, or (l, m, n) for 3D.

    Returns
    -------
    np.ndarray
        Array with the first guess of z, of the same shape as y.
    """
    if np.any(y.mask):
        _, li = distance_transform_edt(y.mask, return_indices=True)
        z0 = np.zeros(y.shape)
        z0[~y.mask] = y[~y.mask]
        if y.ndim == 1:
            z0[y.mask] = y[li[0][y.mask]]
        elif y.ndim == 2:
            z0[y.mask] = y[li[0][y.mask], li[1][y.mask]]
        elif y.ndim == 3:
            z0[y.mask] = y[li[0][y.mask], li[1][y.mask], li[2][y.mask]]
        else:
            raise NotImplementedError(f"Algorithm not implemented for {y.ndim}-dimensional data")
    else:
        z0 = y
    return z0


# ================================================================================================
def studentized_residuals(y, z, mask, s):
    """
    Calculate the Studentized residuals of the approximation using the median absolute deviation and diagonal
    elements of the hat matrix H.

    Parameters
    ----------
    y : np.ndarray
        Input data.
    z : np.ndarray
        Data estimation.
    mask : np.ndarray
        Original mask of the data (y.mask).
    s : float
        Smoothing factor.

    Returns
    -------
    np.ndarray
        Array with studentized residuals of the approximation.
    """
    r = y - z

    # Calculate the median absolute deviation
    mad = np.median(np.abs(r[mask] - np.median(r[mask])))

    # Calculate the hat matrix
    n = r.ndim
    a = np.sqrt(1 + 16 * s)
    h = (np.sqrt(1 + a) / (np.sqrt(2) * a)) ** n

    # Calculate the Studentized residuals
    u = r / (1.4826 * mad * np.sqrt(1 - h))

    return u


# ================================================================================================
def biweight(u, c=4.685):
    """
    Calculate the bisquare weighting of the residuals.

    Parameters
    ----------
    u : np.ndarray
        Studentized residual adjusted for standard deviation.
    c : float, optional
        Residual limit. Default is 4.685.

    Returns
    -------
    np.ndarray
        Array with biweights of the residuals.
    """
    subset = np.less(np.abs(u), c)
    bw = (1 - (u / c) ** 2) ** 2 * subset

    return bw
```
