## Supplementary material for "Whole cryo-FIB lamella acquisition at high magnification with rectangular beam montage cryo-electron tomography": Source code: stitch.py

```
# Postponed annotation evaluation: the sidecar helpers below use "str | None",
# which Python 3.9 cannot evaluate at definition time.
from __future__ import annotations

import re
import logging
import networkx as nx
from tqdm import tqdm
from PIL import Image
import os

from skimage.registration import phase_cross_correlation

from scipy.spatial import KDTree
import matplotlib

# SLURM exports the submitting shell's environment, so a job submitted from an
# X11-forwarded session inherits DISPLAY and matplotlib picks an interactive
# backend.  plt.show(block=True) then blocks until the X connection dies, which
# kills the run before it writes its tilt.  Force Agg in batch, unless the
# caller has asked for a specific backend via MPLBACKEND.
if os.environ.get("MPLBACKEND") is None and (
    os.environ.get("SLURM_JOB_ID") or not os.environ.get("DISPLAY")
):
    matplotlib.use("Agg")
import matplotlib.pyplot as plt
import copy
import glob
import mrcfile
import argparse
import numpy as np
from typing import List, Tuple, Dict, Any
from concurrent.futures import ThreadPoolExecutor
import threading
from .Utilities import *
from .smoothn import smoothn

try:
    from .beam_mask_motioncorr import _create_stack_locked
except ImportError:
    from beam_mask_motioncorr import _create_stack_locked

try:
    from . import sidecar
    from .sidecar import canonical_path, fingerprint, fingerprint_many
except ImportError:
    import sidecar
    from sidecar import canonical_path, fingerprint, fingerprint_many

logger = logging.getLogger(__name__)

#: Name of this step in its sidecars.
SIDECAR_STEP = "stitch"


# ---------------------------------------------------------------------------
# Per-tile MRC directory support
# ---------------------------------------------------------------------------

# Matches filenames like  Name_003_-12.0.mrc  →  groups (tile_idx, tilt_angle)
_TILE_FNAME_RE = re.compile(r"^.+_(\d+)_(-?\d+\.?\d*)\.mrc$")


class TileStack:
    """Read-only adapter that presents a list of per-tile MRC files as a
    virtual (N, H, W) stack, duck-typing the ``tiles`` interface
    expected by :func:`montage`.

    Supports integer indexing, boolean-mask indexing (returns a list), and
    the ``.dtype`` / ``.shape`` attributes used by :func:`montage`.
    """

    def __init__(self, paths: list) -> None:
        self._paths = paths
        # Peek at the first tile only, to report dtype/shape without opening
        # every file up front
        with mrcfile.mmap(paths[0], mode="r", permissive=True) as m:
            self.dtype = m.data.dtype
            self.shape = (len(paths),) + tuple(m.data.shape[-2:])

    def __len__(self) -> int:
        return len(self._paths)

    def __getitem__(self, idx):
        # A single integer index loads just that one tile off disk
        if isinstance(idx, (int, np.integer)):
            with mrcfile.mmap(self._paths[int(idx)], mode="r", permissive=True) as m:
                return np.asarray(m.data)
        # bool mask or index array → list, compatible with executor.map
        if isinstance(idx, np.ndarray) and idx.dtype == bool:
            indices = np.where(idx)[0]
        else:
            indices = np.asarray(idx, dtype=int)
        return [self[i] for i in indices]

    @property
    def name(self) -> str:
        """Representative ``{stem}_{tilt}.mrc`` basename for this tilt.

        Per-tile inputs are named ``{stem}_{tile_index}_{tilt_angle}.mrc``;
        dropping the tile index gives every tile of a tilt the same name, so
        this stack's output files (TIFF, plot, position file) are named the
        same way as they would be for a single per-tilt MRC stack.
        """
        base = os.path.basename(self._paths[0])
        m = _TILE_FNAME_RE.match(base)
        if not m:
            return base
        # Remove "_{tile_index}", keeping the leading stem and trailing tilt.
        return base[: m.start(1) - 1] + base[m.end(1) :]


def _group_per_tile_mrcs(directory: str):
    """Group per-tile MRC files from a motioncorr output directory by tilt angle.

    Expects filenames of the form ``{name}_{tile_index}_{tilt_angle}.mrc``
    (the naming convention produced by ``beam_mask_motioncorr``).

    Returns
    -------
    sorted_tilts : list of float
    tile_paths   : dict mapping tilt_angle → list of paths sorted by tile index
    """
    groups: dict = {}
    # Bucket every matching file by its tilt angle, skipping anything that
    # doesn't follow the naming convention
    for path in glob.glob(os.path.join(directory, "*.mrc")):
        m = _TILE_FNAME_RE.match(os.path.basename(path))
        if not m:
            continue
        tile_idx = int(m.group(1))
        tilt = float(m.group(2))
        # Fold negative-zero tilt onto positive zero so both group together
        if tilt == -0.0:
            tilt = 0.0
        groups.setdefault(tilt, []).append((tile_idx, path))

    # Sort tilts ascending, and each tilt's own files by tile index
    sorted_tilts = sorted(groups)
    tile_paths = {t: [p for _, p in sorted(groups[t])] for t in sorted_tilts}
    return sorted_tilts, tile_paths


def _first_tile_shape(file_or_stack):
    """Return (H, W) of the first tile from either an MRC stack path or a TileStack."""
    if isinstance(file_or_stack, str):
        return mrcfile.mmap(file_or_stack).data.shape[-2:]
    return file_or_stack.shape[-2:]


def _image_basename(file_or_stack):
    """Basename used to name this tilt's outputs, for an MRC path or TileStack."""
    if isinstance(file_or_stack, str):
        return os.path.split(file_or_stack)[1]
    return file_or_stack.name


def parse_commandline() -> Dict[str, Any]:
    """
    Parse commandline input.
    """
    parser = argparse.ArgumentParser(
        description="Stitch square beam montage tomography data."
    )
    parser.add_argument(
        "-i",
        "--input",
        help="Either a glob pattern matching per-tilt MRC stacks (e.g. '*.mrc'), "
        "or a directory of per-tile MRC files produced by beam_mask_motioncorr "
        "(files named {stem}_{tile_index}_{tilt_angle}.mrc).",
        required=True,
        type=str,
    )
    parser.add_argument(
        "--pixel-size",
        dest="pixel_size",
        type=float,
        default=None,
        help="Pixel size in Ångström. If omitted, read from the input MRC's "
        "voxel_size header.",
    )
    parser.add_argument(
        "-o",
        "--output",
        help="Where to write stitched output. Three forms are accepted: "
        "(1) omitted, or a directory — the current per-tilt TIFF (lzw) "
        "behaviour, placed in ./[input_filename]_output if omitted; "
        "(2) 'path/to/stack.mrc', optionally with a ':N' suffix — write "
        "directly into a shared memory-mapped MRC stack at that path "
        "(created on first use, flock-guarded, safe to call concurrently), "
        "skipping the separate JointoMRC.py join step entirely. With no "
        "':N' suffix and no --tilt-index, every discovered tilt is stitched "
        "into its own z-slice of the stack in one run; ':N' (or "
        "--tilt-index) restricts this invocation to writing only slice N, "
        "for one SLURM array task per tilt; (3) 'path/to/file.tiff' — "
        "write a single stitched TIFF instead of treating the path as a "
        "directory. Requires exactly one discovered tilt (use "
        "--tilt-index to select it).",
        required=False,
        type=str,
    )
    parser.add_argument(
        "--tilt-index",
        dest="tilt_index",
        help="Restrict this invocation to stitching only the tilt at this "
        "0-based rank (tilt angles sorted ascending) instead of the "
        "whole series — pairs with --output pointing at an MRC stack to "
        "run one SLURM array task per tilt, each writing only its own "
        "slice. If omitted, every discovered tilt is stitched, as "
        "before. Overrides any ':N' suffix on --output.",
        type=int,
        default=None,
        required=False,
    )
    parser.add_argument(
        "-I",
        "--image_shifts",
        help="Path to text file containing the tilts and a list of image shifts at every tilt (requried).",
        required=True,
        type=str,
    )
    parser.add_argument(
        "-b",
        "--binning",
        help="Binning of input data, defaults to 1",
        required=False,
        type=int,
        default=1,
    )
    parser.add_argument(
        "--rotate",
        help="Rotate every input tile (and template mask) counter-clockwise "
        "by this many degrees before masking/alignment/stitching. Must "
        "be a multiple of 90. Useful when the camera's row/col axes are "
        "rotated relative to the stage x/y axes assumed by the image "
        "shift file. Default 0 (no rotation).",
        required=False,
        type=int,
        choices=[0, 90, 180, 270],
        default=0,
    )
    parser.add_argument(
        "-f",
        "--fringe_size",
        help="Size of Fresnel fringes at edge of beam, this will be removed from the gain reference (default 20).",
        required=False,
        type=int,
        default=20,
    )
    parser.add_argument(
        "-s",
        "--skipcrosscorrelation",
        help="Reuse a previous run's alignment instead of refining tile "
        "positions by cross-correlation. The alignment is read from the HDF5 "
        "position file (see --positionfile), which must already exist — this "
        "run will NOT fall back to raw image shifts and will NOT overwrite "
        "the file. As well as tile positions, the saved tile selection, beam "
        "masks and beam-edge-correction parameters are reused, so the stitch "
        "is pixel-for-pixel reproducible from the same tiles. This is the "
        "mode to use for re-stitching even/odd (denoising) half-stacks from "
        "the full-frame alignment: run once without -s on the full frames, "
        "then once with -s on each half.",
        action="store_true",
    )
    parser.add_argument(
        "-m",
        "--max_allowed_imshift_correction",
        help="Maximum allowed correction (in µm) to tile alignments by "
        "cross-correlation, 0.05 µm by default. This is the outlier test used "
        "by '--consensus lstsq'; the robust consensus methods (including "
        "'irls', the default) ignore it and derive their own tolerance from "
        "the data.",
        type=float,
        default=0.05,
        required=False,
    )

    parser.add_argument(
        "--consensus",
        help="How to reconcile the measured pairwise tile shifts into one set "
        "of tile positions. 'irls' (default) keeps every measurement but "
        "iteratively down-weights whichever disagree with the current fit, "
        "which avoids an all-or-nothing decision when the bad measurements are "
        "not cleanly separated from the good ones. 'ransac' instead keeps the "
        "largest set of measurements that are mutually consistent with a "
        "single set of tile positions and fits only those. Both robust methods "
        "size their own tolerance from the spread of the residuals unless "
        "--consensus-threshold is given. 'lstsq' is the original behaviour: "
        "throw away any measurement differing from the microscope's own image "
        "shift by more than --max_allowed_imshift_correction, then fit the "
        "rest by unweighted least squares — a fixed threshold applied to each "
        "overlap in isolation, which cannot catch a measurement that is wrong "
        "but small and discards a correct one that happens to be large.",
        choices=["lstsq", "ransac", "irls"],
        default="irls",
        required=False,
    )

    parser.add_argument(
        "--consensus-threshold",
        dest="consensus_threshold",
        help="Fix the tolerance (in µm) within which a measurement counts as "
        "agreeing with the fitted tile positions, instead of deriving it from "
        "the spread of the corrections cross-correlation asked for. Used by "
        "--consensus ransac and irls; no effect with --consensus lstsq, which "
        "uses --max_allowed_imshift_correction. The derived value is reported "
        "per tilt in the log, so run once without this and look there for a "
        "sensible starting point.",
        type=float,
        default=None,
        required=False,
    )

    parser.add_argument(
        "--ransac-iterations",
        dest="ransac_iterations",
        help="Number of random spanning-tree samples drawn by --consensus "
        "ransac, on top of the deterministic image-shift hypothesis it always "
        "tries first. Default 200.",
        type=int,
        default=200,
        required=False,
    )

    parser.add_argument(
        "--ransac-seed",
        dest="ransac_seed",
        help="Seed for the --consensus ransac sampling, so that a run is "
        "reproducible. Default 0.",
        type=int,
        default=0,
        required=False,
    )

    parser.add_argument(
        "--irls-loss",
        dest="irls_loss",
        help="Weight function for --consensus irls. 'tukey' (default) falls to "
        "exactly zero beyond the tolerance, so gross outliers are removed "
        "outright; 'huber' only tapers, so a bad measurement retains a little "
        "pull but the overlap graph can never be disconnected by the fit.",
        choices=["tukey", "huber"],
        default="tukey",
        required=False,
    )

    parser.add_argument(
        "-S",
        "--correctimageshiftfilefactor",
        help="Sometimes imageshift file needs to be divided by a factor of 2 since serial-EM uses super-resolution pixels.",
        type=float,
        default=1,
        required=False,
    )
    parser.add_argument(
        "-mt",
        "--maskthreshold",
        help="threshold (as fraction of raw image median) for masking of beam.",
        type=float,
        default=0.4,
        required=False,
    )

    parser.add_argument(
        "-ma",
        "--maskabsolutethreshold",
        help="Absolute threshold (in number of image counts) for masking of beam.",
        type=float,
        default=None,
        required=False,
    )

    parser.add_argument(
        "-R",
        "--ROI",
        help="Region of interest in the montage to stitch in format x0:x1,y0:y1 (in pixels).",
        type=int,
        default=None,
        required=False,
        nargs=4,
    )

    parser.add_argument(
        "-nt",
        "--nthreads",
        help="Number of threads for parralel implementation.",
        type=int,
        default=1,
        required=False,
    )

    parser.add_argument(
        "--nosmooth",
        help="Disable smoothing of background regions in the stitched montage.",
        action="store_true",
        default=False,
    )

    parser.add_argument(
        "--mark-uncovered",
        help=(
            "Instead of inpainting uncovered (no-tile) regions, mark them with "
            "pixel value -1. Use this when the output will be processed by "
            "mask_and_inpaint, which detects the sentinel automatically and "
            "includes those regions in its initial mask."
        ),
        action="store_true",
        default=False,
        dest="mark_uncovered",
    )

    parser.add_argument(
        "-pf",
        "--positionfile",
        help="Path of the HDF5 alignment file for this tilt, overriding the "
        "default of <outdir>/<input basename with .mrc replaced>"
        "_refined_positions.h5. Without -s this is the file the refined tile "
        "positions, tile selection, beam masks and beam-edge parameters are "
        "WRITTEN to (overwriting it if it exists). With -s it is the file "
        "they are READ from, and it must exist. Because the default is "
        "derived from the input tile filename, even/odd re-stitches that "
        "share a basename with the full-frame run and write to the same "
        "--output directory pick up the right file with no -pf at all; only "
        "pass -pf when that is not true. Never point several tilts of one "
        "array job at a single -pf path — they will overwrite each other.",
        type=str,
        default=None,
        required=False,
    )

    parser.add_argument(
        "-T",
        "--referencebeam",
        help="Path to a reference beam image (TIFF or MRC) defining the beam shape. When provided, edge-based cross-correlation is used to align a mask built from it to each tile instead of threshold-based masking. For multi-frame MRC or TIFF files, append :INDEX to select a specific frame (e.g. mask.mrc:7).",
        type=str,
        default=None,
        required=False,
    )

    parser.add_argument(
        "--min_mean_intensity",
        help="Tiles with a mean pixel value at or below this threshold are excluded from processing. Default is 5.",
        type=float,
        default=0,
        required=False,
    )

    parser.add_argument(
        "--correct_beam_edges",
        help="Enable correction of beam edge darkening caused by inelastic (plasmon) "
        "scattering.  The image provided via --referencebeam is used as the "
        "vacuum reference.  With no argument, plasmon parameters are fitted "
        "independently for every tile.  With an integer argument N "
        "(e.g. --correct_beam_edges 42), tile N is used to fit the parameters "
        "once and those values are applied to all tiles — useful when a known "
        "interior tile gives a better fit than edge tiles.  "
        "Pass -1 to automatically use the tile closest to the centre of the montage.",
        nargs="?",
        const=True,
        default=False,
        type=int,
    )

    parser.add_argument(
        "--plasmon_energy",
        help="Plasmon energy in eV used for beam-edge correction (--correct_beam_edges). "
        "Default 21 eV, appropriate for vitreous ice / water.",
        type=float,
        default=21.0,
        required=False,
    )

    parser.add_argument(
        "--voltage",
        help="Accelerating voltage in kV used for beam-edge correction "
        "(--correct_beam_edges).  Default 300 kV.",
        type=float,
        default=300.0,
        required=False,
    )

    parser.add_argument(
        "--no-sidecar",
        dest="sidecar",
        action="store_false",
        default=True,
        help="Do not write a provenance sidecar beside each stitched tilt. "
        "Sidecars are ON by default and cost one small JSON write per tilt: "
        "they record this step's effective geometry — rotation, binning, "
        "canvas origin and size — plus that tilt's tile positions and "
        "selection, so nothing downstream has to restate 02_stitch.sh from "
        "memory. There is no shared file and no finalize pass; each array "
        "task writes only its own tilt's sidecar under "
        "<output>.sidecars/<tilt>.json, so no two tasks ever touch the same "
        "path. Turning them off is for throwaway runs only — a stitched "
        "canvas with no sidecar is one whose ROI has to be guessed later, "
        "which is exactly the failure this replaced.",
    )

    parser.add_argument(
        "--input-frame",
        dest="input_frame",
        choices=["auto", "raw", "motion-corrected"],
        default="auto",
        help="What the input tiles are, for the sidecar's benefit. Stitching "
        "can be the first step of the pipeline or the second, and only in the "
        "second case is this step's --rotate applied on top of another one. "
        "'raw' says these tiles come straight off the detector, so the total "
        "rotation from the raw frames is this step's own and these files are "
        "the provenance root, so they get fingerprinted individually; "
        "'motion-corrected' says they do not. 'auto' (default) infers it from "
        "whether the input tiles carry motion_correction sidecars and, "
        "finding none, records the total rotation as unknown rather than "
        "guessing.",
    )

    verbosity = parser.add_mutually_exclusive_group()
    verbosity.add_argument(
        "-v",
        "--verbose",
        help="Print detailed per-tile progress and diagnostic information.",
        action="store_true",
        default=False,
    )
    verbosity.add_argument(
        "-q",
        "--quiet",
        help="Suppress all output except warnings and errors.",
        action="store_true",
        default=False,
    )

    return vars(parser.parse_args())


def stitch(
    ims,
    positions,
    msks,
    pixel_size,
    binning=1,
    montagewidth=None,
    montageorigin=None,
    smooth=True,
    nthreads=1,
    mark_uncovered=False,
):
    """
    Stitch a set of tiles into a single montage canvas.

    Parameters
    ----------
    ims : sequence of numpy.ndarray
        2D image tiles to place into the montage (shape: N, rows, cols).
    positions : numpy.ndarray
        Array of shape (N, 2) with tile positions in microns (x, y order).
    msks : sequence of numpy.ndarray
        Boolean/0-1 masks matching each tile; masked-out pixels are ignored.
    pixel_size : float
        Pixel size in Angstroms for the unbinned images.
    binning : int, optional
        Binning factor applied to the montage. Default is 1.
    montagewidth : None or (2,) array_like, optional
        Full montage size in Angstroms. If None, inferred from positions.
    montageorigin : None or (2,) array_like, optional
        Montage origin in Angstroms (most negative image shift). If None, inferred.
    smooth : bool, optional
        If True, smooth background (uncovered) regions; otherwise fill with median.

    Returns
    -------
    canvas : numpy.ndarray
        Stitched montage image (overlaps averaged).
    indxmap : np.ndarray
        Map of indicating index of tiles
    """
    # Determine the global range of tiles in Angstroms
    pixels = ims[0].shape
    if montagewidth is None:
        width = (
            np.ptp(positions, axis=0) * 1e4
            + np.asarray(pixels[::-1]) * pixel_size * binning
        )
    else:
        width = montagewidth
    if montageorigin is None:
        origin = np.amin(positions, axis=0) * 1e4
    else:
        origin = montageorigin

    # Calculate the size of the global montage canvas in pixels
    size = (
        np.asarray(width / pixel_size / binning, dtype=int)[::-1]
        # + pixels
        # + np.asarray([1, 1])
    )

    logger.debug("Stitch canvas: %d × %d px  |  %d tile(s)", size[0], size[1], len(ims))
    # Initialize the montage canvas and overlap map
    canvas = np.zeros(size)
    overlap = np.zeros(size, dtype=np.uint8)
    # Initialise the index map with n+1, where n is the number of tiles to identify
    # indices that correspond to no tile, indices that correspond to a tile will be
    # overwritten
    indxmap = len(ims) * np.ones(size, dtype=np.uint16)

    # Place each image onto the canvas, applying the masks
    for i, (im, position, msk) in enumerate(zip(ims, positions, msks)):
        # s is shape of tile, size is shape of canvas
        s = im.shape
        # Desired coordinate of upper left of tile
        y0, x0 = [int(x) for x in (position * 1e4 - origin) / pixel_size / binning]

        # Skip tiles that have fallen off canvas
        if x0 > size[0] or y0 > size[1]:
            continue
        if x0 + s[0] < 0 or y0 + s[1] < 0:
            continue

        # Truncate canvas coordinate beginnings to be >= 0
        cy0, cx0 = [max(coord, 0) for coord in [y0, x0]]
        # Truncate canvas coordinate maximum to be <= canvas array limits
        cx1, cy1 = [
            min(coord, limit) for coord, limit in zip([x0 + s[0], y0 + s[1]], size)
        ]

        # Size of tile that will make it onto the montage canvas
        X = cx1 - cx0
        Y = cy1 - cy0

        # Coordinates of tile, if x0 (or y0) < 0 this implies some of the tile
        # falls off the left (or upper) edge of canvas so is not included
        tx0 = -min(x0, 0)
        tx1 = tx0 + X
        ty0 = -min(y0, 0)
        ty1 = ty0 + Y

        # Add this tile's masked pixels into the canvas, count how many tiles
        # covered each pixel, and record which tile last claimed each pixel
        canvas[cx0:cx1, cy0:cy1] += np.where(msk, im, 0)[tx0:tx1, ty0:ty1]
        overlap[cx0:cx1, cy0:cy1] += np.where(msk, np.uint8(1), np.uint8(0))[
            tx0:tx1, ty0:ty1
        ]
        indxmap[cx0:cx1, cy0:cy1][msk[tx0:tx1, ty0:ty1]] = np.uint16(i)

    # Regions no tile covers at all (overlap == 0) have no real data to
    # average -- fill them with the caller's chosen strategy instead
    if mark_uncovered:
        canvas[overlap < 1] = -1  # sentinel: picked up by mask_and_inpaint as uncovered
    elif smooth:
        logger.info("Smoothing background regions")
        smoothed = smoothn(
            np.ma.masked_array(canvas, overlap < 1),
            s=1e7,
            max_iter=100,
            workers=nthreads,
        )
        canvas = np.where(overlap < 1, smoothed, canvas)
    else:
        canvas[overlap < 1] = 0  # np.median(canvas[overlap == 1])

    # Treat uncovered pixels as if covered once, so the divide below is a
    # no-op there instead of a divide-by-zero
    overlap = np.where(overlap > 1, overlap, 1)
    # Average overlapping tiles' contributions at each pixel
    canvas /= overlap

    return canvas, indxmap


def default_positionfile(image, outdir):
    """Path of the HDF5 alignment file stitch.py reads/writes for one tilt.

    Derived from the *input* tile filename, not the output filename, so that
    re-stitching a different frame subset of the same tilt (e.g. the even/odd
    half-stacks used for denoising) from the same input basename resolves to
    the same file, while different tilts of one series never collide.
    """
    return os.path.join(
        outdir, _image_basename(image).replace(".mrc", "_refined_positions.h5")
    )


def save_alignment_state(
    positionfile,
    original_positions,
    positions,
    xcorr,
    overlaps,
    deltas,
    indxmap,
    binned_pixel_size,
    tile_selection,
    beam_reference_mask,
    plasmon_parameters,
    beam_shifts,
    source_image,
    edge_inliers=None,
):
    """Write everything a ``-s`` run needs to reproduce this stitch.

    ``Original_positions``/``Refined_positions`` are full-length (n_tiles, 2)
    arrays in microns; ``tile_selection`` is the length-n_tiles boolean mask of
    tiles that actually made it into the montage, and ``plasmon_parameters`` /
    ``beam_shifts`` are ordered to match ``positions[tile_selection]``. Storing
    the selection alongside the positions is what makes reuse safe: without it
    a reuse run re-derives the selection from its own (noisier) data and can
    admit a tile whose stored position was never refined.

    ``beam_reference_mask`` is the single eroded beam-reference mask (in
    tile-local canvas coordinates); every tile's actual mask is
    ``roll_no_periodic(beam_reference_mask, beam_shifts[k])`` and is not
    itself stored — ``montage_projection.py`` reconstructs it the same way.
    Both ``beam_reference_mask`` and ``beam_shifts`` are None when the run
    had no ``--referencebeam`` and so masked each tile independently; neither
    dataset is written in that case.

    ``edge_inliers`` is the boolean mask, one entry per row of ``overlaps``, of
    which cross-correlation measurements the consensus method accepted. It is
    diagnostic only — nothing reads it back — but it is the record of which
    overlaps the fitted positions actually rest on.
    """
    # Datasets that are always present, paired with their HDF5 names
    arrays = [
        original_positions,
        positions,
        xcorr,
        overlaps,
        np.asarray(deltas),
        indxmap.astype(np.uint16),
        binned_pixel_size,
        np.asarray(tile_selection, dtype=bool),
        np.asarray(plasmon_parameters, dtype=float),
        source_image,
    ]
    names = [
        "Original_positions",
        "Refined_positions",
        "cross_correlations",
        "overlaps",
        "relative_shifts",
        "index_map",
        "binned_pixel_size",
        "tile_selection",
        "plasmon_parameters",
        "source_image",
    ]
    # Optional datasets: only written when the caller actually has them
    # (e.g. no --referencebeam means no beam_reference_mask/beam_shifts)
    if beam_reference_mask is not None:
        arrays.append(np.asarray(beam_reference_mask, dtype=bool))
        names.append("beam_reference_mask")
    if beam_shifts is not None:
        arrays.append(np.asarray(beam_shifts, dtype=np.int32))
        names.append("beam_shifts")
    if edge_inliers is not None:
        arrays.append(np.asarray(edge_inliers, dtype=bool))
        names.append("edge_inliers")
    save_array_to_hdf5(arrays, positionfile, names)


def load_alignment_state(positionfile, n_tiles, mask_shape, source_image=None):
    """Read a previous run's alignment back for reuse under ``-s``.

    Returns a dict with keys ``positions`` (full-length, microns),
    ``tile_selection`` (bool mask or None), ``beam_shifts`` ((n_selected, 2)
    array or None) and ``plasmon_parameters`` ((n_selected, 2) array or None).
    The optional entries are None for files written before those datasets
    existed, in which case the caller recomputes them. Per-tile masks are
    never stored or returned here — the caller rebuilds
    ``beam_reference_mask`` fresh from this run's own ``--referencebeam``
    and rolls it by each ``beam_shifts`` entry.

    Raises ValueError if the stored arrays do not match the data being
    stitched — a mismatch means the file belongs to a different tilt or a
    different binning, and silently proceeding would misplace tiles.
    """

    # Optional datasets simply weren't written by an older run; treat a
    # missing key as "not available" rather than an error
    def _get(name):
        try:
            return load_array_from_hdf5(positionfile, name)
        except KeyError:
            return None

    # Refined_positions is the one dataset every valid alignment file must
    # have; everything else below is optional/backward-compatible
    positions = _get("Refined_positions")
    if positions is None:
        raise ValueError(
            f"{positionfile} has no 'Refined_positions' dataset; it was not "
            "written by a stitch.py refinement run."
        )
    positions = np.asarray(positions)
    if positions.shape != (n_tiles, 2):
        raise ValueError(
            f"{positionfile} holds positions for {positions.shape[0]} tile(s) "
            f"but this tilt has {n_tiles}; the position file belongs to a "
            "different tilt or a different image-shift file."
        )

    # Each optional dataset below is validated against this run's own data
    # shape before being trusted -- a mismatch means the file was written
    # for a different tilt/binning/rotation and reuse would misplace tiles
    tile_selection = _get("tile_selection")
    if tile_selection is not None:
        tile_selection = np.asarray(tile_selection, dtype=bool)
        if tile_selection.shape != (n_tiles,):
            raise ValueError(
                f"{positionfile} holds a tile selection of length "
                f"{tile_selection.shape[0]} but this tilt has {n_tiles} tiles."
            )

    beam_reference_mask = _get("beam_reference_mask")
    if beam_reference_mask is not None:
        beam_reference_mask = np.asarray(beam_reference_mask, dtype=bool)
        if beam_reference_mask.shape != tuple(mask_shape):
            raise ValueError(
                f"{positionfile} holds a beam reference mask of shape "
                f"{beam_reference_mask.shape} but tiles bin to "
                f"{tuple(mask_shape)}; the position file was written with a "
                "different --binning or --rotate."
            )

    plasmon_parameters = _get("plasmon_parameters")
    if plasmon_parameters is not None:
        plasmon_parameters = np.asarray(plasmon_parameters, dtype=float)
        # An empty array means a run that saved the dataset but had no
        # --correct_beam_edges; treat it the same as never having been saved
        if plasmon_parameters.size == 0:
            plasmon_parameters = None

    stored_source = _get("source_image")
    if stored_source is not None and source_image is not None:
        if stored_source != source_image:
            # Not fatal — the even/odd half-stacks share a basename with the
            # full-frame run, so a legitimate reuse always matches, but a
            # deliberate cross-run reuse may not. It is fatal often enough to
            # be worth saying out loud: a --positionfile shared by every tilt
            # of an array job lands here.
            logger.warning(
                "%s was written from %s but is being reused for %s; check "
                "that this is really the same tilt.",
                positionfile,
                stored_source,
                source_image,
            )

    beam_shifts = _get("beam_shifts")
    if beam_shifts is not None:
        beam_shifts = np.asarray(beam_shifts, dtype=int)
        if beam_shifts.size == 0:
            beam_shifts = None
        else:
            # Must have exactly one shift per selected tile
            n_expected = (
                int(tile_selection.sum()) if tile_selection is not None else None
            )
            if n_expected is not None and beam_shifts.shape[0] != n_expected:
                raise ValueError(
                    f"{positionfile} holds {beam_shifts.shape[0]} beam shift(s) "
                    f"but {n_expected} selected tile(s); the file is inconsistent."
                )

    return {
        "positions": positions,
        "tile_selection": tile_selection,
        "plasmon_parameters": plasmon_parameters,
        "beam_shifts": beam_shifts,
    }


def plot_cross_correlation_map(
    image,
    outdir,
    original_positions,
    positions,
    M,
    overlaps,
    deltas,
    canvas,
    ims,
    indxmap,
    xcorr,
    pixel_size,
    binning,
    maxshift,
    montageorigin=None,
    montagewidth=None,
    inliers=None,
):
    """
    Plot initial vs. cross-correlation-refined tile positions (with shift
    vectors between overlapping tiles) to <image>_Plot.pdf. Only meaningful
    after cross-correlation refinement has actually run (i.e. when montage()
    calls this with skipcrosscorrelation=False).

    Shift vectors the consensus rejected are drawn dashed. ``inliers`` is the
    accept/reject mask the consensus method actually returned, one entry per
    row of ``overlaps``; without it the drawing falls back to re-deriving the
    fixed ``maxshift`` test, which is only the right answer for the "lstsq"
    method.
    """
    # One square-ish figure sized to comfortably fit ntiles annotated points
    ntiles = len(original_positions[M])
    figsize = 2 * (int(np.ceil(np.sqrt(ntiles))),)
    xcorfig, xcorax = plt.subplots(figsize=figsize)
    # Contrast-stretch the canvas display around its own mean/std, rather
    # than the full data range, so a few bright/dark outlier pixels don't
    # wash out the tile boundaries
    cmean = np.mean(canvas)
    cstd = np.std(canvas)
    if montageorigin is None or montagewidth is None:
        extent = None
    else:
        # Convert the montage's Angstrom-space origin/width into the pixel
        # extents imshow needs, so tile positions (also in pixels) line up
        extent = [
            montageorigin[0],
            (montageorigin[0] + montagewidth[0]),
            montageorigin[1],
            (montageorigin[1] + montagewidth[1]),
        ]
        extent = np.asarray(extent) / pixel_size
    xcorax.imshow(
        canvas,
        cmap=plt.get_cmap("gist_gray"),
        origin="lower",
        vmin=cmean - cstd,
        vmax=cmean + cstd,
        extent=extent,
    )
    # s in unbinned pixels so all coordinates share the same unit
    s = np.asarray(ims[0].shape)[::-1] * binning
    # Initial (pre-refinement) tile centres: convert microns to pixels and
    # offset from each tile's corner to its centre
    posi = (original_positions[M] * 1e4) / pixel_size
    posi += s / 2
    xcorax.plot(*posi.T, "ko", label="Initial tile positions")
    for i, pos in enumerate(posi):
        xcorax.annotate(str(i), pos)
    # Refined tile centres, same conversion
    posi = (positions[M] * 1e4) / pixel_size
    posi += s / 2
    xcorax.plot(*posi.T, "bo", label="Refined tile positions")
    # xcorrmax = np.amax(xcorr)
    cmap = plt.get_cmap("viridis")
    # Draw each overlap's measured cross-correlation shift as a vector from
    # tile i's centre, dashed if the consensus fit rejected it as an outlier
    for ind, (i, j) in enumerate(overlaps):
        # Retrieve image shifts for the overlapping tiles
        x1 = (original_positions[M][i] * 1e4) / pixel_size

        x2 = (original_positions[M][j] * 1e4) / pixel_size
        dx = [int(x) for x in (x2 - x1)][::-1]
        x1 += s / 2
        delta = (deltas[ind] * 1e4) / pixel_size

        if inliers is not None:
            # Use the consensus method's own accept/reject decision when we
            # have it -- correct for every method, not just "lstsq"
            rejected = not bool(inliers[ind])
        else:
            # Fallback: re-derive the fixed-threshold test "lstsq" itself
            # uses, only correct when the caller ran that method
            rejected = (
                np.linalg.norm(np.asarray(dx) - delta) > maxshift / pixel_size * 1e4
            )
        linestyle = "--" if rejected else "-"

        xcorax.plot(
            [x1[0], x1[0] + delta[1]],
            [x1[1], x1[1] + delta[0]],
            linestyle=linestyle,
            color="b",
        )
    xcorax.set_xlabel("x (unbinned pixels)")
    xcorax.set_ylabel("y (unbinned pixels)")
    plotfile = os.path.join(
        outdir, _image_basename(image).replace(".mrc", "_Plot.pdf")
    )
    xcorfig.savefig(plotfile)
    plt.close(xcorfig)


def montage(
    image,
    outdir,
    positions,
    pixel_size,
    binning=8,
    skipcrosscorrelation=False,
    montagewidth=None,
    montageorigin=None,
    tiles=None,
    maxshift=0.05,
    consensus="irls",
    consensus_threshold=None,
    ransac_iterations=200,
    ransac_seed=0,
    irls_loss="tukey",
    fringe_size=20,
    maskthreshold=0.4,
    maskabsolutethreshold=None,
    nthreads=1,
    positionfile=None,
    smooth=True,
    mark_uncovered=False,
    reference_beam=None,
    min_mean_intensity=20,
    correct_beam_edges=False,
    E_plasmon_eV=21.0,
    voltage_kV=300.0,
    rotate=0,
    output_mrc=None,
    tilt_z_index=None,
    n_tilts_total=None,
    fileout_path=None,
    sidecar_step=None,
    sidecar_input_frame=None,
    tilt_deg=None,
    series_z_index=None,
):
    """
    Creates a montage image from a series of input images by aligning and stitching them together.

    This function takes a set of images, positions them according to provided coordinates,
    and aligns them based on mask images or generated masks. The images are then combined
    into a single montage image.

    Parameters:
    -----------
    image : str
        mrc file containing montage tiles
    outdir : str
        directory for output
    positions : numpy.ndarray
        Array of shape (N, 2) containing the (x, y) coordinates for positioning each image in units of pixels.
        Note that the x,y have opposite convention to standard y,x Python convention
    pixel_size : float
        The pixel size in Angstroms.
    binning : int, optional
        Binning factor to reduce the image size. Default is 8.
    skipcrosscorrelation : bool, optional
        Reuse the alignment saved in positionfile instead of refining tile
        positions by masked cross correlation. The file must exist (a missing
        one raises FileNotFoundError rather than silently falling back to raw
        image shifts), and the saved tile selection, beam masks and beam-edge
        parameters are reused along with the positions so that the stitch is
        reproducible from the same tiles.
    montagewidth : None or (2,) array_like
        size in Angstrom of the full montage in both dimensions, useful for consistency
        with other tilts in the tilt series
    montageorigin : None or (2,) array_like
        Origin point (most negative image shift) in Angstrom of the full montage
        in both dimensions, useful for consistency with other tilts in the tilt
        series
    tiles : None or sliceobject, optional
        Slice object indicating tiles that will be stitched (mainly for testing purposes)
    consensus : str, optional
        How the measured pairwise shifts are reconciled into one set of tile
        positions: "irls" (default, robustly reweighted least squares),
        "ransac", or "lstsq" (the original fixed-threshold rejection followed
        by plain least squares). See the module section "Global consensus of
        tile positions". ``maxshift`` is the rejection threshold for "lstsq"
        only; the robust methods derive their own from the data unless
        ``consensus_threshold`` is given.
    consensus_threshold : float or None, optional
        Tolerance in microns within which a measurement counts as agreeing with
        the fitted positions, for "ransac" and "irls". None (default) derives
        it from the spread of the corrections; see _auto_threshold.
    ransac_iterations : int, optional
        Random spanning-tree samples drawn by "ransac". Default 200.
    ransac_seed : int, optional
        Seed for that sampling, so a run is reproducible. Default 0.
    irls_loss : str, optional
        "tukey" (default) or "huber".
    fringe_size : float, optional
        Size of the Fresnel fringe region excluded from the beam mask. Default is 20.
    medianthreshold : float, optional
        Threshold for (as a fraction of the median) for masking the image. Default is 0.4.
    positionfile : string, optional
        Path of the hdf5 alignment file for this tilt, written when
        skipcrosscorrelation is False and read when it is True. Defaults to
        default_positionfile(image, outdir). Unlike earlier versions, an
        existing file does not by itself suppress refinement — pass
        skipcrosscorrelation to reuse it.
    correct_beam_edges : bool, optional
        When True, beam edge darkening due to plasmon scattering is corrected
        per-tile using plasmon_beam_correction().  The reference_beam image is
        used as the vacuum reference and is aligned to each tile via
        align_beam_reference. Requires reference_beam to be given -- there is
        no tile-0 fallback, since plasmon correction needs a real reference
        beam to correct against.
    E_plasmon_eV : float, optional
        Plasmon energy in eV for beam-edge correction. Default 21 eV.
    voltage_kV : float, optional
        Accelerating voltage in kV for beam-edge correction. Default 300 kV.
    rotate : int, optional
        Rotate every tile (and the template mask) counter-clockwise by this
        many degrees before any further processing. Must be a multiple of
        90. Default 0 (no rotation).
    output_mrc : str or None, optional
        When given, write the stitched canvas into z-slice ``tilt_z_index``
        of a shared memory-mapped MRC stack at this path (created under a
        file lock on first use), instead of only writing the per-tilt TIFF.
        Requires ``tilt_z_index`` and ``n_tilts_total`` to also be given.
    tilt_z_index : int or None, optional
        0-based z-slice this tilt occupies in ``output_mrc``.
    n_tilts_total : int or None, optional
        Total number of tilts in the series — sizes ``output_mrc`` on creation.
    fileout_path : str or None, optional
        Explicit path for the stitched TIFF, overriding the default
        ``outdir/<image>.tif`` naming. Ignored when ``output_mrc`` is given
        (the TIFF is skipped entirely in that case).
    sidecar_step : tuple or None, optional
        ``(effective, requested)`` from :func:`stitch_effective`. When given,
        this tilt's sidecar is written beside the canvas: the step's geometry
        plus tile positions in both microns and canvas pixels, the tile
        selection, and the consensus fit they rest on. Requires
        ``montageorigin`` so the canvas frame is well defined; a run without it
        (positions inferred per tilt) has no series-wide canvas to record
        against, and the sidecar is skipped with a warning.
    sidecar_input_frame : str or None, optional
        Resolved value of ``--input-frame`` (see :func:`resolve_input_frame`).
        ``"raw"`` means stitching is the first step, so this tilt's sidecar
        carries a fingerprint of every input tile file — nothing upstream has
        recorded them.
    tilt_deg : float or None, optional
        Tilt angle of this montage, used as the sidecar's key.
    series_z_index : int or None, optional
        This tilt's 0-based rank among all discovered tilt angles sorted
        ascending — the slice it occupies in the joined stack. Recorded
        whatever the output mode, because it is the number JointoMRC.py and
        everything downstream order sections by; ``tilt_z_index`` only exists
        when this run writes an MRC stack itself.
    Returns:
    --------
    numpy.ndarray
        The resulting stitched montage image.
    """
    if rotate % 90 != 0:
        raise ValueError("rotate must be a multiple of 90 degrees")
    rot_k = (rotate // 90) % 4

    # Load tile data: accept either an MRC stack path or a TileStack adapter.
    tile_selection = tiles  # save before overwriting with actual image data
    tiles = mrcfile.mrcmemmap.MrcMemmap(image).data if isinstance(image, str) else image

    # Output dtype matches the input tiles' own dtype (e.g. float32 for
    # motion-corrected data) rather than forcing int16, which loses precision
    # for values clustered near zero.
    tile_dtype = np.dtype(tiles.dtype)
    if mark_uncovered and np.issubdtype(tile_dtype, np.unsignedinteger):
        raise ValueError(
            f"mark_uncovered requires a signed or floating tile dtype to "
            f"represent the -1 sentinel; got unsigned dtype {tile_dtype}."
        )

    pixels = np.asarray([x // binning for x in tiles.shape[-2:]], dtype=int)
    if rot_k % 2:
        # A 90°/270° rotation swaps height and width for every tile.
        pixels = pixels[::-1]

    # M is the boolean mask describing which tiles to include.
    # None → all tiles; slice or int-index array → convert to bool; bool array → use directly.
    if tile_selection is None:
        M = np.ones(len(tiles), dtype=bool)
    elif isinstance(tile_selection, slice):
        M = np.zeros(len(tiles), dtype=bool)
        M[tile_selection] = True
    else:
        M = np.zeros(len(tiles), dtype=bool)
        M[tile_selection] = True

    # Convert positions from pixels to microns, remove 3rd dimension
    positions = positions[:, :2] * pixel_size * 1e-4

    # ── Alignment reuse ───────────────────────────────────────────────────────
    # skipcrosscorrelation means "reuse the alignment a previous run saved",
    # so the position file must exist. Falling back to raw image shifts here
    # would produce an unrefined montage that looks plausible but is silently
    # misaligned relative to the run it is supposed to match — for even/odd
    # denoising half-stacks that is exactly the failure that matters, so it is
    # an error instead.
    if positionfile is None:
        positionfile = default_positionfile(image, outdir)
    saved = None
    if skipcrosscorrelation:
        if not os.path.exists(positionfile):
            raise FileNotFoundError(
                f"--skipcrosscorrelation was given but no alignment file exists "
                f"at {positionfile}. Run stitch.py once without "
                "--skipcrosscorrelation to create it, or point --positionfile "
                "at the file written by that run."
            )
        logger.info("Reusing tile alignment from %s", positionfile)
        saved = load_alignment_state(
            positionfile, len(tiles), pixels, _image_basename(image)
        )
        positions = saved["positions"]
        if saved["tile_selection"] is not None:
            M = saved["tile_selection"]
        else:
            logger.warning(
                "%s predates tile-selection storage; re-deriving which tiles to "
                "include from this run's own data. Rewrite it with a "
                "refinement run to guarantee the same tiles are used.",
                positionfile,
            )
        if saved["beam_shifts"] is None:
            logger.warning(
                "%s holds no saved beam shifts; they will be recomputed from "
                "this run's data and will differ slightly from the run that "
                "wrote the file. Rewrite it with a refinement run to make the "
                "masking identical.",
                positionfile,
            )
        if correct_beam_edges and (
            saved["plasmon_parameters"] is None
            or np.isnan(saved["plasmon_parameters"]).all()
        ):
            # The refinement run either predates parameter storage or ran
            # without --correct_beam_edges. Refitting per tile here would make
            # the inelastic correction depend on this stack's own noise (and on
            # thread completion order), so the even and odd halves would get
            # measurably different corrections — the whole point of reuse.
            logger.warning(
                "--correct_beam_edges is enabled but %s holds no inelastic "
                "scattering parameters, so they will be refitted from this "
                "run's data and will not match the run that wrote the file. "
                "Re-run the refinement step with --correct_beam_edges to store "
                "them.",
                positionfile,
            )

    # Initialize an empty mask list
    msks = []

    ims = []  # List to store the processed images
    beam_posns = []  # List to store the beam positions

    # Identify template mask. A --referencebeam is a real reference beam image
    # (or the caller's substitute for one) and switches on the fast path:
    # build one mask from it, erode once, and roll it into place per tile
    # (see _process_tile). Without one, each tile is masked independently by
    # thresholding, exactly as before --referencebeam existed -- there is no
    # shared reference to align to, so this is where erosion cost (and any
    # dose-dependent cross-correlation noise) is paid per tile.
    use_template_alignment = reference_beam is not None
    if correct_beam_edges and not use_template_alignment:
        raise ValueError(
            "--correct_beam_edges requires --referencebeam: plasmon correction "
            "corrects each tile against a reference beam image, and that "
            "reference can no longer be silently substituted with tile 0. "
            "Pass --referencebeam, or drop --correct_beam_edges to mask each "
            "tile independently."
        )

    raw_template = None
    template_mask = None
    template_edges = None
    if use_template_alignment:
        if rot_k:
            reference_beam = np.ascontiguousarray(np.rot90(reference_beam, k=rot_k))

        # Keep the raw (non-binary) beam image as the reference for plasmon correction.
        raw_template = np.asarray(reference_beam, dtype=float)

        smooth_template_mask = convolve(reference_beam, Gaussian(3, reference_beam.shape))
        template_edges = np.hypot(
            sobel(smooth_template_mask, axis=0), sobel(smooth_template_mask, axis=1)
        )
        # From here on the reference beam has become a binary mask, so it
        # gets a name that reflects that: template_mask.
        template_mask = make_mask(
            reference_beam,
            shrinkn=0,
            medianthreshold=maskthreshold,
            absolutethreshold=maskabsolutethreshold,
        )
        

        # Erode once here, rather than per tile: the mask's shape is fixed once
        # thresholded from the reference beam, only its position moves tile to
        # tile, and erosion by a disk commutes with translation. Every tile then
        # just rolls this single eroded mask into place (see _process_tile).
        # radius=beam_shrinkn (not beam_shrinkn/2) to match make_mask's
        # fallback threshold path, so --fringe_size shrinks the mask by the
        # same amount whether or not --referencebeam is given.
        beam_shrinkn = fringe_size / binning
        if beam_shrinkn > 0:
            struct_elem = circular_mask(
                [int(beam_shrinkn * 2) + 1] * 2, radius=beam_shrinkn
            )
            template_mask = binary_erosion(template_mask, structure=struct_elem)
    else:
        logger.debug("No --referencebeam provided — masking each tile independently")

    # Plasmon correction parameter state.
    # _plasmon_fixed: when the user passes --correct_beam_edges N, tile N is
    #   fitted once before the thread pool and its (n, q_E) are locked in for
    #   all other tiles, bypassing per-tile optimisation entirely.
    # _plasmon_running: when no reference tile is given, each tile fits its own
    #   parameters but uses the running mean of previous fits as a warm start,
    #   skipping the 25×25 grid and going straight to Nelder-Mead.
    _plasmon_fixed_n = [None]
    _plasmon_fixed_q_E = [None]

    ref_tile_idx = (
        correct_beam_edges
        if (
            isinstance(correct_beam_edges, int)
            and not isinstance(correct_beam_edges, bool)
        )
        else None
    )
    correct_beam_edges = bool(correct_beam_edges)

    if ref_tile_idx == -1:
        # Auto-select the tile whose stage position is closest to the montage centre.
        active_indices = np.where(M)[0]
        active_positions = positions[M]
        centre = active_positions.mean(axis=0)
        ref_tile_idx = int(
            active_indices[np.argmin(np.linalg.norm(active_positions - centre, axis=1))]
        )
        logger.info(f"Auto-selected centre tile {ref_tile_idx} as plasmon reference")

    if ref_tile_idx is not None and correct_beam_edges:
        logger.info(f"Fitting plasmon parameters on reference tile {ref_tile_idx} ...")
        _ref_img = np.asarray(tiles[ref_tile_idx]).copy()
        if rot_k:
            _ref_img = np.ascontiguousarray(np.rot90(_ref_img, k=rot_k))
        if binning > 1:
            _ref_img = fourier_interpolate(
                _ref_img, [x // binning for x in _ref_img.shape]
            )
        _ref_shift = align_beam_reference(_ref_img, raw_template, template_edges)
        _ref_mask = roll_no_periodic(
            template_mask, (-_ref_shift[0], -_ref_shift[1]), axis=(0, 1)
        )
        _, _plasmon_fixed_n[0], _plasmon_fixed_q_E[0] = plasmon_beam_correction(
            _ref_img,
            raw_template,
            _ref_mask,
            pixel_size_nm=pixel_size * binning / 10.0,
            template_edges=template_edges,
            E_plasmon_eV=E_plasmon_eV,
            voltage_kV=voltage_kV,
        )
        logger.info(
            f"Reference tile fit: n={_plasmon_fixed_n[0]:.3f}  "
            f"q_E={_plasmon_fixed_q_E[0]:.5f} cyc/nm"
        )

    _plasmon_lock = threading.Lock()
    _plasmon_n_sum = [0.0]
    _plasmon_qE_sum = [0.0]
    _plasmon_count = [0]

    # Per-tile plasmon parameters saved by a previous refinement run, and the
    # per-tile beam-reference shift, indexed by position within tiles[M].
    # Reusing them (rather than re-deriving them from this run's data) is what
    # makes an even/odd re-stitch differ from the full-frame stitch by noise
    # alone: the beam is located by cross-correlation, which lands a pixel or
    # two differently on half-dose data, and the per-tile plasmon fit
    # warm-starts from a running mean whose value depends on thread completion
    # order. The mask itself is never stored per tile — it is always
    # `roll_no_periodic(template_mask, shift)`, so the shift is the only
    # thing that needs to survive a reuse run.
    _saved_plasmon = saved["plasmon_parameters"] if saved is not None else None
    _saved_shifts = saved["beam_shifts"] if saved is not None else None
    # Per-tile (n_avg, q_E) actually used, for saving. NaN marks "not fitted".
    _plasmon_fits = np.full((int(M.sum()), 2), np.nan)
    # Per-tile (dy, dx) alignment of the beam reference, likewise for saving.
    _beam_shifts = np.zeros((int(M.sum()), 2), dtype=int)

    def _process_tile(indexed_img):
        k, img = indexed_img
        im = np.asarray(img).copy()
        if rot_k:
            im = np.ascontiguousarray(np.rot90(im, k=rot_k))
        if binning > 1:
            im = fourier_interpolate(im, [x // binning for x in im.shape])
        if _saved_shifts is None and np.mean(im) <= min_mean_intensity:
            # Only applied when deriving the tile selection from this run's own
            # data; under reuse the saved selection is authoritative, so that
            # half-dose tiles cannot drop out and desynchronise the saved shifts.
            return None
        if use_template_alignment:
            # The beam reference's alignment to this tile is measured by
            # cross-correlation and so is dose-dependent; reuse the saved shift
            # when there is one, otherwise measure it and record it. This one
            # measurement serves both the mask (rolled into place below) and,
            # when requested, the plasmon correction.
            if _saved_shifts is not None:
                shift = _saved_shifts[k]
            else:
                shift = align_beam_reference(im, raw_template, template_edges)
            _beam_shifts[k] = shift
            mask = roll_no_periodic(template_mask, (-shift[0], -shift[1]), axis=(0, 1))

            # DEBUG: visually sanity-check the rolled mask against the beam.
            # Only the first tile, since plt.show() blocks and this runs in a
            # ThreadPoolExecutor -- remove the `k == 0` guard (or change it to
            # e.g. `k < 5`) to look at more tiles, but do so with --nthreads 1,
            # since matplotlib isn't safe to drive from multiple threads at once.
            # fig, axes = plt.subplots(1, 4, figsize=(15, 5))
            # vmin = np.percentile(np.where(mask, im, 0),1)
            # vmax = np.percentile(np.where(mask, im, 0),99)
            # axes[0].imshow(im, cmap="gray",vmin=vmin,vmax=vmax)
            # axes[0].set_title(f"beam (tile {k})")
            # axes[1].imshow(mask, cmap="gray")
            # axes[1].set_title(f"rolled mask (shift={tuple(shift)})")
            # axes[2].imshow(np.where(mask, im, 0), cmap="gray",vmin=vmin,vmax=vmax)
            # axes[2].set_title("masked beam")
            # axes[3].imshow(reference_beam, cmap="gray",vmin=vmin,vmax=vmax)
            # axes[3].set_title("Reference beam")
            # for ax in axes:
            #     ax.axis("off")
            # fig.tight_layout()
            # plt.show()
        else:
            # No shared reference to align to -- mask this tile independently
            # by thresholding, same as before --referencebeam existed.
            mask = make_mask(
                im,
                shrinkn=fringe_size / binning,
                medianthreshold=maskthreshold,
                absolutethreshold=maskabsolutethreshold,
            )
        if correct_beam_edges:
            if _saved_plasmon is not None:
                n_use, q_E_use = _saved_plasmon[k]
            else:
                n_use, q_E_use = _plasmon_fixed_n[0], _plasmon_fixed_q_E[0]
            if n_use is not None and not np.isnan(n_use):
                # Fixed-parameter mode: reuse a stored or reference-tile fit.
                im, _, _ = plasmon_beam_correction(
                    im,
                    raw_template,
                    mask,
                    pixel_size_nm=pixel_size * binning / 10.0,
                    template_edges=template_edges,
                    E_plasmon_eV=E_plasmon_eV,
                    voltage_kV=voltage_kV,
                    n_fixed=n_use,
                    q_E_fixed=q_E_use,
                    beam_shift=shift,
                )
                n_fit, q_E_fit = n_use, q_E_use
            else:
                # Per-tile mode: use running mean as warm start for the grid search.
                with _plasmon_lock:
                    count = _plasmon_count[0]
                    n_hint = _plasmon_n_sum[0] / count if count else None
                    q_E_hint = _plasmon_qE_sum[0] / count if count else None
                im, n_fit, q_E_fit = plasmon_beam_correction(
                    im,
                    raw_template,
                    mask,
                    pixel_size_nm=pixel_size * binning / 10.0,
                    template_edges=template_edges,
                    E_plasmon_eV=E_plasmon_eV,
                    voltage_kV=voltage_kV,
                    n_hint=n_hint,
                    q_E_hint=q_E_hint,
                    beam_shift=shift,
                )
                if n_fit > 0.0:
                    with _plasmon_lock:
                        _plasmon_n_sum[0] += n_fit
                        _plasmon_qE_sum[0] += q_E_fit
                        _plasmon_count[0] += 1
            _plasmon_fits[k] = (n_fit, q_E_fit)
        return im, mask

    _quiet = not logger.isEnabledFor(logging.INFO)
    max_workers = min(os.cpu_count() or 1, nthreads)
    with ThreadPoolExecutor(max_workers=max_workers) as executor:
        results = list(
            tqdm(
                executor.map(_process_tile, enumerate(tiles[M])),
                total=len(positions[M]),
                desc="Making masks",
                disable=_quiet,
            )
        )
    # Single threaded implementation for testing/debugging
    # results = []
    # for tile in enumerate(tiles[M]):
    #     result = _process_tile(tile)
    #     results.append(result)

    M_indices = np.where(M)[0]
    excluded = [M_indices[i] for i, r in enumerate(results) if r is None]
    if excluded:
        logger.warning(
            "Excluded %d low-intensity tile(s): indices %s", len(excluded), excluded
        )
        M[excluded] = False
        kept = [r is not None for r in results]
        _plasmon_fits = _plasmon_fits[kept]
        _beam_shifts = _beam_shifts[kept]
    logger.debug("%d tile(s) retained after intensity filtering", sum(M))

    ims = [r[0] for r in results if r is not None]
    msks = [r[1] for r in results if r is not None]

    # Calculate the overlaps between adjacent tiles
    overlaps = find_overlaps(
        positions[M], pixels, pixel_size * binning, msks, plot=False
    )

    consensus_stats = {}
    if not skipcrosscorrelation:
        # Refine the tile positions using cross-correlation between overlapping
        # tiles
        original_positions = copy.deepcopy(positions)
        positions[M], xcorr, deltas, edge_inliers = cross_correlate_tiles(
            positions[M],
            ims,
            msks,
            overlaps,
            pixel_size * binning,
            max_correction=maxshift,
            max_workers=nthreads,
            consensus=consensus,
            consensus_threshold=consensus_threshold,
            ransac_iterations=ransac_iterations,
            ransac_seed=ransac_seed,
            irls_loss=irls_loss,
            stats=consensus_stats,
        )
    else:
        # positions/M/masks were already loaded from positionfile above.
        original_positions = positions
        consensus_stats = {"method": "reused", "n_edges": len(overlaps)}

    # Recalculate overlaps before mask clipping since some tiles might have migrated
    # since the last time we did this.

    postrefineoverlaps = find_overlaps(
        positions[M],
        pixels,
        pixel_size * binning,
        msks,
        plot=False,
        minoverlapfrac=1 / np.prod(pixels),
    )

    # Clip masks to ensure overlapping regions are only taken from closest tile.
    # Keep the unclipped beam masks for input to clip_masks_to_overlaps below;
    # what actually gets saved for reuse is the single beam-reference mask
    # (template_mask) plus each tile's shift, since clipping — like the
    # per-tile mask itself — is a deterministic function of those plus the
    # tile positions.
    beam_msks = msks
    msks = clip_masks_to_overlaps(
        beam_msks, positions[M], postrefineoverlaps, pixels, pixel_size * binning
    )

    # Stitch the images together using the refined tile positions
    canvas, indxmap = stitch(
        ims,
        positions[M],
        msks,
        pixel_size,
        binning=binning,
        montagewidth=montagewidth,
        montageorigin=montageorigin,
        smooth=smooth,
        mark_uncovered=mark_uncovered,
        nthreads=nthreads,
    )
    if output_mrc is None:
        fileout = (
            fileout_path
            if fileout_path is not None
            else os.path.join(
                outdir, os.path.splitext(_image_basename(image))[0] + ".tif"
            )
        )
    if not skipcrosscorrelation:
        save_alignment_state(
            positionfile,
            original_positions,
            positions,
            xcorr,
            overlaps,
            deltas,
            indxmap,
            pixel_size / binning,
            M,
            template_mask,
            _plasmon_fits,
            _beam_shifts if use_template_alignment else None,
            _image_basename(image),
            edge_inliers=edge_inliers,
        )
        logger.info("Saved tile alignment to %s", positionfile)
        plot_cross_correlation_map(
            image,
            outdir,
            original_positions,
            positions,
            M,
            overlaps,
            deltas,
            canvas,
            ims,
            indxmap,
            xcorr,
            pixel_size,
            binning,
            maxshift,
            montageorigin=montageorigin,
            montagewidth=montagewidth,
            inliers=edge_inliers,
        )

    # Note that Python Image library does not support write 16-bit integer and writes
    # 32-bit real instead ¯\_(ツ)_/¯
    # Image.fromarray(canvas).save(fileout.replace('.tif','float.tiff'),compression="tiff_lzw")
    # Image.fromarray(canvas.astype(np.uint16)).save(fileout.replace('.tif','nolzw.tiff'))
    # Clipping saves underflow errors upon conversion to uint16, sometimes images values can go
    # negative from smoothing algorithm

    # Clip to the input tiles' own dtype range.  When mark_uncovered is set,
    # uncovered pixels are exactly -1 (no smoothn runs in that path so no
    # other negatives exist). Otherwise clip negatives to 0 so smoothn
    # artefacts don't become false sentinels.
    lo_clip = -1 if mark_uncovered else 0
    hi_clip = (
        None if np.issubdtype(tile_dtype, np.floating) else np.iinfo(tile_dtype).max
    )
    clipped = np.clip(canvas, lo_clip, hi_clip).astype(tile_dtype)
    if output_mrc is None:
        Image.fromarray(clipped).save(fileout, compression="tiff_lzw")
        if isinstance(image, str):
            # Stamp the source raw montage's mtime onto the stitched output so
            # downstream tools (JointoMRC.py's --acquisition_order) can recover
            # true acquisition order from file mtime, with no mdoc needed.
            src_mtime = os.path.getmtime(image)
            os.utime(fileout, (src_mtime, src_mtime))
        save_array_as_png(
            canvas, fileout.replace(".tiff", ".png"), cmap=plt.get_cmap("Greys")
        )

    if output_mrc is not None:
        if tilt_z_index is None or n_tilts_total is None:
            raise ValueError("output_mrc requires both tilt_z_index and n_tilts_total")
        # MRC mode matching the input tiles' own dtype, consistent with
        # JointoMRC.py's stack convention.
        mrc_mode = mrcfile.utils.mode_from_dtype(tile_dtype)
        mrc_dtype = mrcfile.utils.dtype_from_mode(mrc_mode)
        _create_stack_locked(
            output_mrc,
            (n_tilts_total, *clipped.shape),
            pixel_size * binning,
            mrc_mode=mrc_mode,
        )
        with mrcfile.mmap(output_mrc, mode="r+") as out_mrc:
            out_mrc.data[tilt_z_index] = clipped.astype(mrc_dtype, copy=False)
        logger.info(
            "Wrote tilt directly into %s  [slice %d/%d]",
            output_mrc,
            tilt_z_index,
            n_tilts_total,
        )

    if sidecar_step is not None:
        if montageorigin is None:
            logger.warning(
                "Sidecar: no montage origin for this run, so tile positions "
                "have no series-wide canvas to be expressed in; skipping the "
                "sidecar for tilt %s.", tilt_deg,
            )
        else:
            step_effective, step_requested = sidecar_step
            write_tilt_sidecar(
                output_path=output_mrc if output_mrc is not None else fileout,
                image=image,
                tilt_deg=tilt_deg if tilt_deg is not None else float("nan"),
                step_effective=step_effective,
                step_requested=step_requested,
                tilt_record=tilt_sidecar_record(
                    tilt_deg=tilt_deg if tilt_deg is not None else float("nan"),
                    z_slice=(series_z_index if series_z_index is not None
                             else tilt_z_index),
                    positionfile=positionfile,
                    pixel_size=pixel_size,
                    binning=binning,
                    montage_origin_A=montageorigin,
                    canvas_shape_px=canvas.shape,
                    nominal_positions_um=original_positions,
                    refined_positions_um=positions,
                    tile_selection=M,
                    excluded_tiles=excluded,
                    consensus_stats=consensus_stats,
                    rotate=rotate,
                ),
                # First step in the pipeline: nothing upstream has recorded
                # these files, so this sidecar is their only fingerprint.
                fingerprint_inputs=sidecar_input_frame == "raw",
            )

    return canvas


def plot_overlaps(positions, overlaps, show=True):
    # Plot every tile position as a point, connecting overlapping pairs with
    # a line -- overlaps is a condensed (pdist-style) boolean array, one
    # entry per unordered pair, so it has to be expanded back to (i, j)
    m = len(positions)
    fig, ax = plt.subplots()
    ax.plot(*positions.T, "ko")
    for n, val in enumerate(overlaps):
        if val:
            i, j = condensed_to_square(n, m)
            x1 = positions[i]
            x2 = positions[j]
            ax.plot([x1[0], x2[0]], [x1[1], x2[1]], "r-")
    plt.show(block=show)
    return fig


def array_overlap(dx, n):
    """Overlapping indices for 1d array shifted dx relative to each other"""
    # A positive shift moves the second array's slice earlier and the
    # first's later (or vice versa for a negative shift), so only the
    # overlapping tail/head of each is compared
    if dx > 0:
        return [dx, n], [0, n - dx]
    else:
        return [0, n + dx], [-dx, n]


def find_overlaps(positions, pixels, pixel_size, masks, minoverlapfrac=0.01, plot=True):
    """
    Identify pairs of images that overlap in a 2D montage, based on their positions
    and field of view, and further refine the overlap based on binary masks.

    Parameters:
    ----------
    positions : ndarray of shape (m, 2)
        The (x, y) coordinates of `m` images in a 2D montage. Note that
        the positions array is ordered (x,y) opposite to python standard (y,x)
    pixels : tuple or list of length 2
        Dimensions of each image in pixels, as (height, width).
    pixel_size : float
        The size of each pixel in microns.
    masks : list of ndarrays
        Binary masks for each image (same shape as image), used to check if
        a specific region of the images overlaps. If None, the function only
        considers the geometric overlap.
    minoverlapfrac : float, optional
        Minimum fraction of overlap, as a fraction of the average illuminated
        (mask-True) tile area, required for two images to be considered
        overlapping. Default is 0.01 (i.e., 1%).
    plot : float, optional
    Returns:
    -------
    overlapping_inds : list of lists
        List of pairs of indices representing images that overlap.
        Each element is a list of two indices `[i, j]`, where the i-th and j-th
        images overlap.
    """
    m = len(positions)  # Number of images

    from scipy.spatial.distance import pdist

    # Calculate the size of each field of view in microns (height, width)
    # Order of pixels array has to be reversed from python standard to match
    # positions array
    criterion = pixel_size * np.asarray(pixels)[::-1] * 1e-4

    # Calculate (unique) distances between montage pieces
    # with scipy's pdist (pair-wise distance) function and
    # calculate which are within a "field of view" of each other
    # result is stored in a condensed distance matrix

    overlaps = np.logical_and(
        *[pdist(positions[..., i : i + 1]) / criterion[i] < 1.0 for i in range(2)]
    )

    # Visualize the positions and overlap status
    if plot:
        overlapfig = plot_overlaps(positions, overlaps, show=False)
        overlapax = overlapfig.get_axes()[0]

    # List to store the indices of overlapping image pairs
    overlapping_inds = []

    # Update criterion to be the minimum required overlap area in pixels, as a
    # fraction of the average illuminated (mask-True) area rather than the
    # full rectangular tile (which includes non-illuminated corners/edges).
    criterion = minoverlapfrac * np.mean([np.sum(msk) for msk in masks])

    # Iterate over the condensed distance matrix
    for n, val in enumerate(overlaps):
        if val:  # If overlap is detected
            # Convert the index in the condensed matrix back to the square form
            i, j = condensed_to_square(n, m)
            x1 = positions[i]
            x2 = positions[j]

            # Calculate the relative shift in position between the two images, in pixels
            dx = [int(x) for x in (x2 - x1) / pixel_size * 1e4]

            # Determine the overlapping pixel regions in both images
            i1, i2 = array_overlap(dx[1], pixels[0])
            j1, j2 = array_overlap(dx[0], pixels[1])

            # Check if the overlap area in the masks exceeds the minimum criterion
            overlap = (
                np.sum(
                    np.logical_and(
                        masks[i][i1[0] : i1[1], j1[0] : j1[1]],
                        masks[j][i2[0] : i2[1], j2[0] : j2[1]],
                    )
                )
                >= criterion
            )
            if overlap:
                overlapping_inds.append([i, j])
                if plot:
                    x1 = positions[i]
                    x2 = positions[j]
                    overlapax.plot([x1[0], x2[0]], [x1[1], x2[1]], "b--")
    if plot:
        plt.show(block=True)
    return overlapping_inds


def clip_masks_to_overlaps(masks, positions, overlaps, pixels, pixel_size):
    """
    Clip binary masks so when there is overlap, pixels are taken from the closest
    tile to the pixel in the overlapping region.

    Parameters:
    -----------
    masks : list of ndarrays
        List of binary masks for each image tile.
    positions : ndarray of shape (N, 2)
        Array of (x, y) coordinates for each image tile in microns.
    overlaps : list of tuples
        List of pairs of indices representing which tiles overlap.
    pixels : tuple or list of length 2
        Dimensions of each image tile in pixels, as (height, width).
    pixel_size : float
        The size of each pixel in microns.

    Returns:
    --------
    clipped_masks : list of ndarrays
        List of modified binary masks with overlapping regions clipped to
        closest tile.
    """
    clipped_masks = [copy.deepcopy(msk) for msk in masks]

    x = np.arange(masks[0].shape[-1])
    y = np.arange(masks[0].shape[-2])
    distancefromcenter = np.zeros((len(masks), *masks[0].shape[-2:]))
    for i, mask in enumerate(masks):
        y0, x0 = center_of_mass_2d(mask)
        distancefromcenter[i] = (y[:, None] - y0) ** 2 + (x[None, :] - x0) ** 2

    # Pre-compute per-tile canvas offsets using the same integer truncation as stitch,
    # so that dx below matches the actual pixel-level placement of each tile.
    # int(A - B) != int(A) - int(B) in general; computing the difference of rounded
    # individual offsets avoids a systematic 1-pixel gap at seams.
    origin = np.amin(positions, axis=0) * 1e4
    canvas_offsets = np.array(
        [[int(v) for v in (pos * 1e4 - origin) / pixel_size] for pos in positions]
    )  # shape (N, 2): [x_offset, y_offset] per tile

    for i, j in overlaps:
        # Calculate relative shift in pixels, consistent with stitch's canvas placement
        dx_raw = canvas_offsets[j] - canvas_offsets[i]  # [x_shift, y_shift]
        dx = list(dx_raw[::-1])  # [row_shift, col_shift]

        # Get the indices for array overlap
        i1, i2 = array_overlap(dx[0], pixels[0])
        j1, j2 = array_overlap(dx[1], pixels[1])

        # Make a mask where pixels closer to center of tile i are True
        closer = np.less_equal(
            distancefromcenter[i][i1[0] : i1[1], j1[0] : j1[1]],
            distancefromcenter[j][i2[0] : i2[1], j2[0] : j2[1]],
        )
        # Only consider pixels that are in the mask of tile j
        closer = np.where(clipped_masks[j][i2[0] : i2[1], j2[0] : j2[1]], closer, True)
        clipped_masks[i][i1[0] : i1[1], j1[0] : j1[1]] = np.logical_and(
            closer, clipped_masks[i][i1[0] : i1[1], j1[0] : j1[1]]
        )

        closer = np.greater_equal(
            distancefromcenter[i][i1[0] : i1[1], j1[0] : j1[1]],
            distancefromcenter[j][i2[0] : i2[1], j2[0] : j2[1]],
        )
        closer = np.where(clipped_masks[i][i1[0] : i1[1], j1[0] : j1[1]], closer, True)
        clipped_masks[j][i2[0] : i2[1], j2[0] : j2[1]] = np.logical_and(
            closer, clipped_masks[j][i2[0] : i2[1], j2[0] : j2[1]]
        )

    return clipped_masks


# ---------------------------------------------------------------------------
# Global consensus of tile positions from measured pairwise displacements
# ---------------------------------------------------------------------------
#
# Cross-correlation gives one relative displacement per overlapping tile pair.
# Turning those into a single set of tile positions is an over-determined
# linear problem — one equation per measurement, two unknowns per tile — and a
# few of the measurements are simply wrong: a pair whose overlap region holds
# little signal correlates on noise, or the correlation peak hops to a
# neighbouring feature of a repetitive structure.  Ordinary least squares has
# no way to tell those apart from the good measurements and smears their error
# across the whole montage, so the outliers have to be identified either before
# or during the fit.
#
# Three strategies are available, selected by ``--consensus``:
#
#   irls    (the default) keep every measurement but weight it by how well it
#           agrees with the current fit (Tukey or Huber), iterating to
#           convergence.  No hard accept/reject decision, which makes it stable
#           when the outliers are not cleanly separated from the good
#           measurements.
#   ransac  hypothesise a set of tile positions, count the measurements
#           consistent with it, keep the largest consistent set, and fit only
#           those.  The tolerance is derived from the spread of the data unless
#           it is given explicitly.
#   lstsq   the original.  Reject any measurement that differs from the
#           microscope's own image shift by more than
#           ``--max_allowed_imshift_correction``, then fit the rest by
#           unweighted least squares.  A fixed threshold, applied to each edge
#           in isolation: it knows nothing about what the rest of the montage
#           says, so it cannot catch a measurement that is wrong but small, and
#           it throws away a correct measurement that happens to be large.
#
# Throughout this section, tile positions are (n, 2) arrays in microns ordered
# (x, y) — the convention of the image shift file — while a measured
# displacement ``d`` for the edge (i, j) is the position of tile j minus that
# of tile i in microns ordered (y, x), the convention that comes out of
# ``phase_cross_correlation``.


def _bridge_components(nominal_positions, graph):
    """Extra edges tying each disconnected component of ``graph`` to the largest.

    Where the retained cross-correlation measurements leave a group of tiles
    with no reliable link to the rest of the montage, the microscope's own
    image shifts are the only information available about where that group
    sits, so the closest pair of tiles across the gap is constrained to its
    nominal offset.  Returns a list of ``(i, j, d)`` in the same convention as
    the measured edges.
    """
    # Sort so the largest component is last; nothing to bridge if there's
    # only one component (the whole graph is already connected)
    components = sorted(nx.connected_components(graph), key=len)
    if len(components) < 2:
        return []
    largest = list(components.pop())
    # Nearest-neighbour search over the largest component's nominal
    # positions, so each smaller component bridges via its closest tile
    tree = KDTree([nominal_positions[x] for x in largest])
    bridges = []
    for component in components:
        members = list(component)
        distances, indices = tree.query([nominal_positions[i] for i in members])
        closest = int(np.argmin(distances))
        i = members[closest]
        j = largest[int(indices[closest])]
        # Displacement in the (y, x) convention the measured edges use
        bridges.append((i, j, (nominal_positions[j] - nominal_positions[i])[::-1]))
    return bridges


def _least_squares_positions(
    nominal_positions,
    edge_ij,
    edge_d,
    weights=None,
    anchor_idx=None,
    anchor_pos=None,
    anchor_weights=None,
):
    """Weighted least-squares tile positions from pairwise displacements.

    Parameters
    ----------
    nominal_positions : (n, 2) ndarray
        Tile positions in microns, (x, y), from the microscope image shifts.
        Used to bridge components the retained edges leave unconnected, and to
        fix the gauge (least squares determines positions only up to a global
        translation) when no anchors are given.
    edge_ij : (m, 2) int ndarray
        Tile index pair (i, j) for each measurement.
    edge_d : (m, 2) ndarray
        Measured position of tile j minus that of tile i, in microns, (y, x).
    weights : (m,) ndarray, optional
        Per-edge weight.  Edges weighted zero are dropped from the graph
        entirely (and so may trigger bridging); the rest enter the normal
        equations scaled by sqrt(weight).  Defaults to all ones, which
        reproduces the original unweighted solve exactly.
    anchor_idx : (k,) int ndarray, optional
        Tile indices carrying an *absolute* position constraint from an
        independent measurement — e.g. a per-tile correction derived from
        IMOD fiducial-track residuals. Unlike every row above, which
        constrains a difference between two tiles, an anchor constrains one
        tile's position directly. Default: no anchors (identical behaviour
        to before this parameter existed).
    anchor_pos : (k, 2) ndarray, optional
        Target position in microns, (x, y), for each entry of ``anchor_idx``.
    anchor_weights : (k,) ndarray, optional
        Per-anchor weight, same convention as ``weights``. Defaults to all
        ones.

    Returns
    -------
    positions : (n, 2) ndarray
        Fitted positions in microns, (x, y).
    n_bridges : int
        How many nominal-shift bridges had to be added.
    """
    nominal_positions = np.asarray(nominal_positions, dtype=float)
    n = len(nominal_positions)
    edge_ij = np.asarray(edge_ij, dtype=int).reshape(-1, 2)
    edge_d = np.asarray(edge_d, dtype=float).reshape(-1, 2)
    if weights is None:
        weights = np.ones(len(edge_ij))
    weights = np.asarray(weights, dtype=float)
    used = np.where(weights > 0)[0]

    if anchor_idx is None:
        anchor_idx = np.zeros(0, dtype=int)
        anchor_pos = np.zeros((0, 2))
    else:
        anchor_idx = np.asarray(anchor_idx, dtype=int).reshape(-1)
        anchor_pos = np.asarray(anchor_pos, dtype=float).reshape(-1, 2)
    anchor_weights = (
        np.ones(len(anchor_idx))
        if anchor_weights is None
        else np.asarray(anchor_weights, dtype=float).reshape(-1)
    )
    anchor_used = np.where(anchor_weights > 0)[0]

    graph = nx.Graph()
    graph.add_nodes_from(range(n))
    graph.add_edges_from(edge_ij[used].tolist())
    bridges = _bridge_components(nominal_positions, graph)

    n_edge_rows = len(used)
    n_bridge_rows = len(bridges)
    nrows = 2 * (n_edge_rows + n_bridge_rows + len(anchor_used))
    A = np.zeros((nrows, 2 * n))
    b = np.zeros(nrows)
    # Each retained measurement contributes two rows (y and x): "position of
    # j minus position of i equals the measured displacement", scaled by
    # sqrt(weight) so squaring in the least-squares solve gives it its
    # intended weight
    for row, e in enumerate(used):
        i, j = edge_ij[e]
        w = np.sqrt(weights[e])
        for c in range(2):
            A[2 * row + c, 2 * j + c] = w
            A[2 * row + c, 2 * i + c] = -w
            b[2 * row + c] = w * edge_d[e, c]
    # Bridge rows use the same difference constraint but at full weight --
    # they only exist to keep the system connected, not to be down-weighted
    for k, (i, j, d) in enumerate(bridges):
        row = n_edge_rows + k
        for c in range(2):
            A[2 * row + c, 2 * j + c] = 1.0
            A[2 * row + c, 2 * i + c] = -1.0
            b[2 * row + c] = d[c]
    # Absolute anchors constrain one tile's position directly, so — unlike
    # every row above — they can fix the global-translation gauge with real
    # data instead of the arbitrary pin below. anchor_pos is (x, y); the
    # unknowns below are unpacked (y, x) per tile (see `positions = np.stack`),
    # hence the axis flip.
    for row_offset, k in enumerate(anchor_used):
        row = n_edge_rows + n_bridge_rows + row_offset
        i = anchor_idx[k]
        w = np.sqrt(anchor_weights[k])
        for c in range(2):
            A[2 * row + c, 2 * i + c] = w
            b[2 * row + c] = w * anchor_pos[k, 1 - c]

    # Only differences of positions are constrained by the edges/bridges, so
    # without anchors A is rank-deficient by two (a global translation in
    # each axis) and the gauge has to be fixed somewhere.  Fix it *before*
    # the solve, by dropping one tile's two unknowns, rather than solving the
    # singular system and re-centring afterwards.  The latter looks
    # equivalent and is not: np.linalg.lstsq with rcond=-1 does not truncate
    # the two ~1e-16 singular values, so the raw solution picks up a
    # component of order 1e12 µm along the null space and re-centring cancels
    # it only to floating-point precision.  That leaves tens of nm of pure
    # numerical noise in the positions, and makes the solution jump by
    # hundreds of nm in response to a 1e-12 change in the weights.  A single
    # solve merely absorbs that as error; IRLS feeds it back through the
    # weights and never converges, wandering between iterates hundreds of nm
    # apart and stopping wherever the iteration count runs out.
    #
    # Anchors do not remove the need for this pin: dropping a column is
    # harmless even when anchors are present (it just means that if the
    # pinned tile itself has an anchor row, that one row is ignored — every
    # other tile's anchor still counts), and it keeps this function
    # well-posed on every IRLS iteration, including ones where Tukey has
    # zero-weighted every anchor.
    gauge_tile = int(np.argmin(np.linalg.norm(nominal_positions, axis=1)))
    free = np.ones(2 * n, dtype=bool)
    free[2 * gauge_tile : 2 * gauge_tile + 2] = False
    x = np.zeros(2 * n)
    x[free] = np.linalg.lstsq(A[:, free], b, rcond=None)[0]
    # Unknowns are interleaved (y_0, x_0, y_1, x_1, ...); positions are (x, y)
    positions = np.stack([x[1::2], x[::2]], axis=-1)

    # Pin the tile closest to the origin to its nominal absolute position
    return (
        positions - positions[gauge_tile] + nominal_positions[gauge_tile],
        len(bridges),
    )


def _edge_residuals(positions, edge_ij, edge_d):
    """Disagreement in microns between each measurement and a set of positions."""
    predicted = (positions[edge_ij[:, 1]] - positions[edge_ij[:, 0]])[:, ::-1]
    return np.linalg.norm(predicted - edge_d, axis=1)


def _robust_scale(residuals):
    """MAD-based Gaussian-equivalent sigma of a set of residual magnitudes.
    
    MAD is median absolute deviation from the median, and 1.4826 is the 
    factor that makes it equivalent to the standard deviation for a Gaussian 
    distribution. Ie. median - 1.4826 * MAD and median + 1.4826 * MAD is the 
    68% confidence interval for a Gaussian distribution.  
    See https://en.wikipedia.org/wiki/Median_absolute_deviation
    """
    if residuals.size == 0:
        return 0.0
    return 1.4826 * np.median(np.abs(residuals - np.median(residuals)))


def _auto_threshold(residuals, k=3.0, floor=0.0):
    """Auto inlier detection: k robust sigma above the typical residual.

    If no threshhold for inlier is given for the iterative re-weighted least 
    squares, derive one from the spread of the residuals of the input positions.
    this is median + k * robust scale, with a floor to avoid over-fitting to a 
    single outlier.  The robust scale is the MAD-based Gaussian-equivalent 
    sigma of the distribution (where 50% of points fall about median). By default
    k=3.0, which is the 99.7% confidence interval for a Gaussian distribution.
    """
    if residuals.size == 0:
        return max(floor, 1e-12)
    return max(np.median(residuals) + k * _robust_scale(residuals), floor, 1e-12)


def ransac_positions(
    nominal_positions,
    edge_ij,
    edge_d,
    threshold=None,
    n_iterations=200,
    lo_iterations=5,
    floor=0.0,
    seed=0,
):
    """RANSAC consensus over the tile-position graph.

    A hypothesis here is a full set of tile positions, and the minimal sample
    that generates one is a spanning tree of the overlap graph: fix the root
    and every other tile follows from the measured displacements along the
    tree.  Each hypothesis is scored by how many of *all* the measurements it
    explains to within ``threshold``, and the largest consensus set wins.

    Two things make this practical for a montage of a few hundred tiles, where
    a textbook RANSAC would need an astronomical number of samples for all
    n - 1 edges of a tree to be drawn clean:

    * The microscope's own image shifts are used as the first hypothesis.  A
      genuine correction is small by construction, so the nominal positions are
      already close to the answer and usually produce a near-complete consensus
      set on their own; the random spanning trees exist to escape the case
      where they do not (a stage jump, a mis-set image shift file).
    * Every hypothesis is locally optimised (LO-RANSAC): its consensus set is
      re-fitted by least squares and the inliers re-selected, repeatedly, so a
      roughly-right hypothesis is pulled to the actual optimum rather than
      being scored where it landed.

    Random spanning trees are drawn by giving every edge a uniform random
    weight and taking the minimum spanning tree.

    Parameters
    ----------
    threshold : float or None
        Inlier tolerance in microns.  ``None`` derives it from the spread of
        the residuals of the nominal hypothesis.
    floor : float
        Lower bound on any derived threshold, in microns.  Correlation peaks
        are located to the (binned) pixel, so there is no point in a tolerance
        finer than that.
    seed : int
        Seed for the spanning-tree sampling, so a run is reproducible.

    Returns
    -------
    positions : (n, 2) ndarray in microns, (x, y)
    inliers : (m,) bool ndarray
    threshold : float
    n_bridges : int
    """
    nominal_positions = np.asarray(nominal_positions, dtype=float)
    edge_ij = np.asarray(edge_ij, dtype=int).reshape(-1, 2)
    edge_d = np.asarray(edge_d, dtype=float).reshape(-1, 2)
    n_edges = len(edge_ij)
    rng = np.random.default_rng(seed)

    # Nothing to reconcile with no measurements at all
    if n_edges == 0:
        return nominal_positions.copy(), np.zeros(0, dtype=bool), float(floor), 0

    # Derive a tolerance from how far the raw nominal positions already are
    # from being self-consistent, unless the caller fixed one
    if threshold is None:
        threshold = _auto_threshold(
            _edge_residuals(nominal_positions, edge_ij, edge_d), floor=floor
        )
    threshold = float(threshold)

    # Build the overlap graph once; each RANSAC trial below only changes the
    # edge weights used to draw a random spanning tree from it
    graph = nx.Graph()
    graph.add_nodes_from(range(len(nominal_positions)))
    for e, (i, j) in enumerate(edge_ij):
        graph.add_edge(int(i), int(j), index=e)

    def _propagate(tree):
        """Positions implied by treating the displacements on ``tree`` as exact."""
        positions = nominal_positions.copy()
        for component in nx.connected_components(tree):
            # Each component is anchored at its own root's nominal position, so
            # components the tree does not link keep their nominal relationship.
            for parent, child in nx.bfs_edges(tree, min(component)):
                e = tree[parent][child]["index"]
                d_xy = edge_d[e][::-1]
                if int(edge_ij[e, 0]) == parent:
                    positions[child] = positions[parent] + d_xy
                else:
                    positions[child] = positions[parent] - d_xy
        return positions

    def _score(positions):
        # How many measurements a set of positions explains within tolerance,
        # and their total residual (for tie-breaking equally-sized consensus sets)
        residuals = _edge_residuals(positions, edge_ij, edge_d)
        inliers = residuals <= threshold
        return inliers, int(inliers.sum()), float(residuals[inliers].sum())

    # Winning hypothesis so far, by inlier count then by lower total residual
    best = {"inliers": None, "count": -1, "cost": np.inf}

    def _consider(positions):
        # Local optimisation (LO-RANSAC): score a hypothesis, then repeatedly
        # re-fit by least squares on its current inlier set and re-score,
        # until the inlier set stops changing -- pulls a roughly-right
        # hypothesis to the actual optimum instead of scoring it as-is
        inliers, count, cost = _score(positions)
        for _ in range(lo_iterations):
            if count == 0:
                break
            refined, _ = _least_squares_positions(
                nominal_positions, edge_ij, edge_d, inliers.astype(float)
            )
            new = _score(refined)
            settled = np.array_equal(new[0], inliers)
            inliers, count, cost = new
            if settled:
                break
        if count > best["count"] or (count == best["count"] and cost < best["cost"]):
            best.update(inliers=inliers, count=count, cost=cost)

    # Always try the microscope's own image shifts first -- for a well
    # behaved montage this alone already gives a near-complete consensus
    _consider(nominal_positions)
    for _ in range(int(n_iterations)):
        if best["count"] == n_edges:
            break
        # A minimum spanning tree over uniformly random edge weights is a
        # uniformly random spanning tree -- this is the random hypothesis draw
        random_weights = rng.random(n_edges)
        for e, (i, j) in enumerate(edge_ij):
            graph[int(i)][int(j)]["weight"] = random_weights[e]
        _consider(_propagate(nx.minimum_spanning_tree(graph)))

    inliers = best["inliers"]
    if inliers is None or not inliers.any():
        logger.warning(
            "RANSAC found no self-consistent set of overlap measurements at a "
            "tolerance of %.4f µm; falling back to using all %d of them.",
            threshold,
            n_edges,
        )
        inliers = np.ones(n_edges, dtype=bool)
    # Final answer: refit on the winning hypothesis's inlier set
    positions, n_bridges = _least_squares_positions(
        nominal_positions, edge_ij, edge_d, inliers.astype(float)
    )
    return positions, inliers, threshold, n_bridges


def irls_positions(
    nominal_positions,
    edge_ij,
    edge_d,
    loss="tukey",
    threshold=None,
    n_iterations=25,
    floor=0.0,
    tol=1e-9,
    anchor_idx=None,
    anchor_pos=None,
    anchor_threshold=None,
    anchor_loss="huber",
):
    """Iteratively reweighted least squares consensus over the tile-position graph.

    Rather than deciding once which measurements to keep, this fits all of
    them, down-weights whichever disagree with the fit, and repeats.  The
    weight function is a Tukey biweight (weight falls to exactly zero at
    ``threshold``, so gross outliers are genuinely removed) or a Huber loss
    (weight falls off as 1/residual beyond ``threshold`` but never reaches
    zero, so the graph can never be disconnected by the fit — safer, but a wild
    measurement still exerts a little pull).

    Tukey is the default because of what happens to a tile all of whose
    overlaps were measured badly, which in practice means a corner tile with
    only two neighbours.  Tukey zeroes both, the tile drops out of the graph
    and is placed by its nominal image shift — the right answer.  Huber leaves
    it attached to two wrong measurements and drags it wherever they point.

    The iteration is started from the microscope's image shifts rather than
    from an ordinary least-squares fit; Tukey is not convex and needs a
    sensible starting point, and the nominal positions are one by construction.

    ``threshold`` is fixed for the whole iteration, and defaults to the same
    fit-independent tolerance :func:`ransac_positions` derives.  Re-estimating
    it from the current fit's residuals — the textbook IRLS recipe — does not
    work here: down-weighting a measurement shrinks the residuals of the ones
    that remain, which shrinks the next scale estimate, which down-weights
    more.  On real montages that ratchet runs away until the fit is
    interpolating a spanning tree and most of the measurements have been thrown
    out, and it does so at a different point on every tilt.
    
    Inputs
    ------
    nominal_positions : (n,2) nd_array in microns, (x, y) with the initial guess of positions for the tiles
    edge_ij : The indices of each tile cross correlation
    edge_d : Measured tile displacement for each tile cross-correlation
    anchor_idx : (k,) int ndarray, optional
        Tile indices carrying an absolute-position constraint from an
        independent measurement — e.g. a per-tile correction derived by
        averaging IMOD fiducial-track residuals (tiltalign's global 3D
        projection-model fit) over the fiducials landing in that tile.
        Unlike the cross-correlation edges, which only ever compare two
        overlapping tiles within one image, this ties the fit to a signal
        that is consistent across the whole tilt series. Default: no
        anchors, i.e. identical behaviour to before this parameter existed.
    anchor_pos : (k, 2) ndarray, optional
        Target position in microns, (x, y), for each entry of ``anchor_idx``
        — typically this tilt's own stitched position for that tile plus its
        fiducial-derived correction.
    anchor_threshold : float or None, optional
        Tolerance in microns for the anchor residuals' Tukey/Huber weight,
        analogous to ``threshold`` for the edges but kept on its own scale:
        a cross-correlation peak on real image content and an averaged
        fiducial-track residual have different noise characteristics, so
        they should not be forced to share one tolerance. None (default)
        derives it from the spread of the anchor residuals at the starting
        positions, the same way ``threshold`` is derived — this is
        unreliable with only a handful of anchors, so pass an explicit
        value when there are only a few tiles with usable anchor data.
    anchor_loss : str, optional
        "huber" (default) or "tukey", independent of ``loss``. Default
        differs from ``loss`` deliberately: with hundreds of cross-correlation
        edges Tukey's hard cutoff is fine, but with only a handful of anchors
        (typical for fiducial-derived corrections, at most one or two tiles
        in a hundred), a measurement sitting near the threshold flips
        between full weight and exactly zero every iteration as the fit
        shifts back and forth in response — a stable period-2 limit cycle,
        not slow convergence, confirmed on real data. Huber's weight never
        reaches zero, so it cannot disconnect and reconnect an anchor like
        that; it converges instead of oscillating, at the cost of a wild
        anchor retaining a little pull rather than being fully rejected.

    Returns
    -------
    positions : (n, 2) ndarray in microns, (x, y)
    weights : (m,) ndarray
        Final per-edge weight in [0, 1].
    inliers : (m,) bool ndarray
        Measurements within the tolerance, for reporting and plotting.
    threshold : float
        That tolerance, in microns.
    n_bridges : int
    anchor_weights : (k,) ndarray
        Final per-anchor weight in [0, 1]. Empty if no anchors were given.
    anchor_threshold : float
        The anchor tolerance actually used. 0.0 if no anchors were given.
    """
    # Force numpy arrays
    nominal_positions = np.asarray(nominal_positions, dtype=float)
    edge_ij = np.asarray(edge_ij, dtype=int).reshape(-1, 2)
    edge_d = np.asarray(edge_d, dtype=float).reshape(-1, 2)

    # Check that loss is one of the expected values
    if loss not in ("tukey", "huber"):
        raise ValueError(f"unknown IRLS loss {loss!r}; expected 'tukey' or 'huber'")
    if anchor_loss not in ("tukey", "huber"):
        raise ValueError(
            f"unknown IRLS anchor_loss {anchor_loss!r}; expected 'tukey' or 'huber'"
        )

    if anchor_idx is None:
        anchor_idx = np.zeros(0, dtype=int)
        anchor_pos = np.zeros((0, 2))
    else:
        anchor_idx = np.asarray(anchor_idx, dtype=int).reshape(-1)
        anchor_pos = np.asarray(anchor_pos, dtype=float).reshape(-1, 2)
    has_anchors = len(anchor_idx) > 0

    def _tukey_or_huber(u, which):
        if which == "tukey":
            # Tukey biweight: weight falls to zero at u=1, and is zero beyond that.
            return np.where(u < 1.0, (1.0 - u**2) ** 2, 0.0)
        return np.where(u <= 1.0, 1.0, 1.0 / np.maximum(u, 1e-12))

    def _anchor_residuals(pos):
        return np.linalg.norm(pos[anchor_idx] - anchor_pos, axis=1)

    # Make a copy of nominal positions to avoid modifying the input array
    positions = nominal_positions.copy()

    # Initialize weights to ones for all edges (and anchors, if any)
    weights = np.ones(len(edge_ij))
    anchor_weights = np.ones(len(anchor_idx))
    n_bridges = 0

    # Determine the threshold for inlier detection. If not provided,
    # compute it based on the residuals of the nominal positions so that
    # 50% of residuals are within the threshold.
    if threshold is None:
        threshold = _auto_threshold(
            _edge_residuals(positions, edge_ij, edge_d), floor=floor
        )
    threshold = float(max(threshold, 1e-12))

    # Same derivation as above, but on the anchor residuals' own scale — see
    # the anchor_threshold docstring for why this is not shared with threshold.
    if has_anchors and anchor_threshold is None:
        anchor_threshold = _auto_threshold(_anchor_residuals(positions), floor=floor)
    anchor_threshold = float(max(anchor_threshold, 1e-12)) if has_anchors else 0.0

    moved = 0.0
    converged = False
    # Classic IRLS loop: re-weight each measurement (and anchor) by how well
    # it agrees with the current fit, re-fit with those weights, repeat until
    # the positions stop moving by more than tol
    for _ in range(int(n_iterations)):
        weights = _tukey_or_huber(
            _edge_residuals(positions, edge_ij, edge_d) / threshold, loss
        )
        if has_anchors:
            anchor_weights = _tukey_or_huber(
                _anchor_residuals(positions) / anchor_threshold, anchor_loss
            )
        new_positions, n_bridges = _least_squares_positions(
            nominal_positions,
            edge_ij,
            edge_d,
            weights,
            anchor_idx=anchor_idx,
            anchor_pos=anchor_pos,
            anchor_weights=anchor_weights,
        )
        moved = (
            np.max(np.linalg.norm(new_positions - positions, axis=1))
            if len(positions)
            else 0.0
        )
        positions = new_positions
        if moved < tol:
            converged = True
            break
    if not converged:
        # The iteration normally settles in about ten passes.  Stopping on the
        # count instead means the answer is whichever iterate the loop happened
        # to end on, so say so rather than reporting it as a fit.
        logger.warning(
            "IRLS did not converge in %d iterations (last step %.1f nm); "
            "the tile positions are the last iterate, not a converged fit",
            int(n_iterations),
            moved * 1e3,
        )

    # Reported inlier set is a plain threshold test on the final fit, distinct
    # from the continuous weights the fit itself actually used
    inliers = _edge_residuals(positions, edge_ij, edge_d) <= threshold
    return positions, weights, inliers, threshold, n_bridges, anchor_weights, anchor_threshold


def cross_correlate_tiles(
    positions,
    tiles,
    masks,
    overlaps,
    pixel_size,
    max_correction=0.3,
    generate_plot=False,
    cross_corr_param_file=None,
    parallelize=True,
    max_workers=None,
    consensus="irls",
    consensus_threshold=None,
    ransac_iterations=200,
    ransac_seed=0,
    irls_loss="tukey",
    stats=None,
    anchor_idx=None,
    anchor_pos=None,
    anchor_threshold=None,
):
    """
    Perform cross-correlation-based alignment of image tiles and adjust their positions.

    This function aligns overlapping image tiles by calculating the relative shifts between them
    using masked phase cross-correlation. The shifts are then used to adjust the global positions
    of the tiles through least squares minimization. The function supports visualizing the tile
    positions and shift vectors if desired.

    Args:
        positions (ndarray): Nx2 array of the initial x, y coordinates of the tiles in micrometers.
        tiles (list of ndarrays): List of 2D arrays representing the image tiles.
        masks (list of ndarrays): List of binary masks representing the valid regions of the tiles.
        overlaps (list of tuples): Pairs of indices representing which tiles overlap and should be aligned.
        pixel_size (float): The pixel size in microns.
        max_correction (float, optional): Maximum allowed correction (in microns) for shifts between tiles. Defaults to 0.1.
                                          Only used by the "lstsq" consensus method, which rejects measurements exceeding it.
        generate_plot (bool or string, optional): Whether to generate a plot visualizing the tile positions and shift vectors. Defaults to False.
                                                  If a string this will be the filename that the plot will be saved as.
        parallelize (bool, optional): Whether to parallelize cross-correlation work. Defaults to True.
        max_workers (int, optional): Maximum worker threads to use when parallelizing. Defaults to a tuned value.
        consensus (str, optional): How to reconcile the measured pairwise shifts into one set of positions:
                                   "irls" (default, robustly reweighted least squares over all measurements),
                                   "ransac" (largest self-consistent set of measurements, then least squares on it) or
                                   "lstsq" (the original fixed-threshold reject then plain least squares).
                                   See the module section "Global consensus of tile positions".
        consensus_threshold (float, optional): Tolerance in microns within which a measurement counts as agreeing with
                                               the fitted positions, for "ransac" and "irls". None (default) derives it
                                               from the spread of the corrections; see _auto_threshold.
        ransac_iterations (int, optional): Random spanning-tree samples drawn by "ransac". Defaults to 200.
        ransac_seed (int, optional): Seed for that sampling, so a run is reproducible. Defaults to 0.
        irls_loss (str, optional): "tukey" (default; zero weight beyond the tolerance) or "huber".
        stats (dict, optional): If given, filled in with the consensus fit's own numbers —
                                method, the tolerance actually used (which the robust methods
                                derive from the data, so it is not knowable from the arguments),
                                edge and inlier counts, and how many tile groups had to be
                                bridged by image shift alone. An out-parameter rather than an
                                extra return value so existing callers keep working.
        anchor_idx ((k,) int ndarray, optional): Tile indices carrying an absolute-position
                                constraint from an independent measurement — e.g. a per-tile
                                correction derived from IMOD fiducial-track residuals. Only
                                used by "irls"; ignored by "ransac"/"lstsq". See
                                irls_positions for the rationale.
        anchor_pos ((k, 2) ndarray, optional): Target position in microns, (x, y), for each
                                entry of anchor_idx.
        anchor_threshold (float, optional): Tolerance in microns for the anchor residuals'
                                robust weighting, kept separate from consensus_threshold. None
                                (default) derives it from the anchor residuals themselves.

    Returns:
        tuple: (positions, xcorr, deltas, inliers) — the updated Nx2 array of x, y tile coordinates,
        the cross-correlation peak values, the measured pairwise shifts (one per entry of `overlaps`,
        in microns and (y, x) order) and a boolean mask of which of those the consensus accepted.

    Notes:
        - The alignment is solved as a least squares problem (Ax = b) to minimize the relative shifts between overlapping tiles.
        - If some tiles are not connected to others through reliable shift determinations, their positions are adjusted
        using the initial relative positions inferred from the microscope.

    References:
        - Masked phase cross-correlation: https://scikit-image.org/docs/stable/auto_examples/registration/plot_masked_register_translation.html
        - Dirk Padfield, "Masked object registration in the Fourier domain", IEEE Transactions on Image Processing, 2011.
    """
    pixels = tiles[0].shape
    cmap = plt.get_cmap("viridis")

    # The global alignment of the tiles is solved as a least squares Ax = b
    # matrix problem (https://en.wikipedia.org/wiki/Linear_least_squares), with
    # two unknowns per tile; see _least_squares_positions and the consensus
    # methods that call it.

    genplot = False
    if type(generate_plot) is bool:
        genplot = generate_plot
        show = True
        savefig = False
    elif type(generate_plot) is str:
        genplot = True
        show = False
        savefig = True

    if genplot:
        xcorfig, xcorax = plt.subplots(figsize=(8, 8))
        xcorax.plot(*positions.T, "ko", label="Initial tile positions")
        for i, pos in enumerate(positions):
            xcorax.annotate(str(i), pos)

    xcorr = []
    deltas = []

    # Debug/testing path: reuse a previous run's measured shifts instead of
    # re-running cross-correlation, so the consensus fit can be re-run alone
    if cross_corr_param_file is not None:
        xcorr, deltas = [
            load_array_from_hdf5(cross_corr_param_file, x)
            for x in ("cross_correlations", "relative_shifts")
        ]
    # Debug only: re-runs the cross correlation and a full two-tile stitch for
    # every overlapping pair and dumps a PDF each, into the CWD under names that
    # collide between concurrent array tasks.  Costs ~14 min per tilt.
    # plot_individual_cross_correlation(tiles,masks,positions,overlaps,pixel_size,filename_template='cross_corr_{0}_{1}.pdf')
    import concurrent.futures
    import os

    def _compute_overlap(args):
        ind, (i, j) = args
        # Retrieve image shifts for the overlapping tiles
        x1 = positions[i]
        x2 = positions[j]

        # Calculate relative shift in pixels
        dx = [int(x) for x in (x2 - x1) / pixel_size * 1e4][::-1]
        if cross_corr_param_file is None:
            # Align tiles by masked phase cross correlation, see:
            # https://scikit-image.org/docs/stable/auto_examples/registration/plot_masked_register_translation.html
            # and Padfield, Dirk. "Masked object registration in the Fourier domain." IEEE Transactions on image processing 21.5 (2011): 2706-2718.

            # Get the indices for array overlap
            i1, i2 = array_overlap(dx[0], pixels[0])
            j1, j2 = array_overlap(dx[1], pixels[1])

            # Crop to the overlap region before cross-correlating.
            # phase_cross_correlation internally FFTs at (2H × 2W) in
            # complex128; using the full tiles wastes memory proportional to
            # (full_tile / overlap)^2. Both crops already show the same
            # physical region, so the algorithm returns only the fine
            # correction; dx is added back to recover the total shift in
            # full-tile coordinates.
            overlap_mask = np.logical_and(
                masks[i][i1[0] : i1[1], j1[0] : j1[1]],
                masks[j][i2[0] : i2[1], j2[0] : j2[1]],
            )
            crop_ref = tiles[i][i1[0] : i1[1], j1[0] : j1[1]]
            crop_mov = tiles[j][i2[0] : i2[1], j2[0] : j2[1]]

            # Calculate shift by masked cross correlation with ordering (Y,X)
            fine_shift = phase_cross_correlation(
                crop_ref,
                crop_mov,
                reference_mask=overlap_mask,
                moving_mask=overlap_mask,
            )[0]
            detected_shift = fine_shift + np.array(dx)
            # delta is measured shift in microns
            delta = np.asarray(detected_shift) * pixel_size * 1e-4
        else:
            delta = deltas[ind]
            # xcorrmax = xcorr[ind]
            detected_shift = delta / pixel_size * 1e-4

        return ind, i, j, dx, detected_shift, delta

    if cross_corr_param_file is None:
        # Run every overlap's cross-correlation, in a thread pool unless
        # parallelism is disabled or there's only one overlap to measure
        deltas = [None] * len(overlaps)
        use_parallel = (
            parallelize
            and len(overlaps) > 1
            and (max_workers is None or max_workers != 1)
        )
        if use_parallel:
            worker_count = max_workers
            if worker_count is None:
                worker_count = min(32, (os.cpu_count() or 1) + 4)
            with concurrent.futures.ThreadPoolExecutor(
                max_workers=worker_count
            ) as executor:
                results = list(
                    tqdm(
                        executor.map(_compute_overlap, enumerate(overlaps)),
                        total=len(overlaps),
                        desc="Cross-correlation alignment",
                    )
                )
        else:
            results = [
                _compute_overlap(args)
                for args in enumerate(
                    tqdm(overlaps, desc="Cross-correlation alignment")
                )
            ]
        for ind, _, _, _, _, delta in results:
            deltas[ind] = delta
    else:
        results = [
            _compute_overlap(args)
            for args in enumerate(tqdm(overlaps, desc="Cross-correlation alignment"))
        ]

    # Gather every measurement as an edge of the tile-position graph.  Which of
    # them to trust is the consensus method's decision, taken below rather than
    # here, so that a method can weigh a measurement against the rest of the
    # montage instead of against a fixed threshold alone.  Indexing by `ind`
    # rather than appending keeps the edge order equal to `overlaps` order.
    n_edges = len(results)
    edge_ij = np.zeros((n_edges, 2), dtype=int) # Tile indices i,j
    edge_d = np.zeros((n_edges, 2))  # measured displacements in microns, (y, x)
    nominal_d = np.zeros((n_edges, 2))  # from the microscope image shifts
    for ind, i, j, dx, detected_shift, delta in results:
        edge_ij[ind] = (i, j)
        edge_d[ind] = delta
        nominal_d[ind] = np.asarray(dx, dtype=float) * pixel_size * 1e-4

    # Peak positions are integers, so a tolerance finer than a couple of binned
    # pixels is meaningless; it is the floor under any data-derived threshold.
    threshold_floor = 2 * pixel_size * 1e-4

    if consensus == "lstsq":
        # Original behaviour: a fixed per-edge cap on the correction to the
        # microscope's own image shift, then an unweighted fit of the survivors.
        threshold = max_correction
        inliers = np.linalg.norm(edge_d - nominal_d, axis=1) <= max_correction
        newpositions, n_bridges = _least_squares_positions(
            positions, edge_ij, edge_d, inliers.astype(float)
        )
    elif consensus == "ransac":
        newpositions, inliers, threshold, n_bridges = ransac_positions(
            positions,
            edge_ij,
            edge_d,
            threshold=consensus_threshold,
            n_iterations=ransac_iterations,
            floor=threshold_floor,
            seed=ransac_seed,
        )
    elif consensus == "irls":
        (
            newpositions,
            _weights,
            inliers,
            threshold,
            n_bridges,
            _anchor_weights,
            anchor_threshold,
        ) = irls_positions(
            positions,
            edge_ij,
            edge_d,
            loss=irls_loss,
            threshold=consensus_threshold,
            floor=threshold_floor,
            anchor_idx=anchor_idx,
            anchor_pos=anchor_pos,
            anchor_threshold=anchor_threshold,
        )
    else:
        raise ValueError(
            f"unknown consensus method {consensus!r}; "
            "expected 'lstsq', 'ransac' or 'irls'"
        )

    logger.info(
        "%s consensus: %d/%d overlap measurements retained (tolerance %.4f µm)%s",
        consensus,
        int(np.sum(inliers)),
        n_edges,
        threshold,
        f"; {n_bridges} tile group(s) placed by image shift alone" if n_bridges else "",
    )
    if stats is not None:
        stats.update({
            "method": consensus,
            "threshold": float(threshold),
            "n_edges": int(n_edges),
            "n_inliers": int(np.sum(inliers)),
            "n_bridges": int(n_bridges),
        })
        if anchor_idx is not None and consensus == "irls":
            stats.update({
                "n_anchors": int(len(anchor_idx)),
                "anchor_threshold": float(anchor_threshold),
                "n_anchor_inliers": int(np.sum(_anchor_weights > 0)),
            })
    if n_edges and not np.all(inliers):
        logger.debug(
            "Rejected overlap measurements: %s",
            [tuple(edge_ij[e]) for e in np.where(~inliers)[0]],
        )

    if show:
        plt.show(block=True)
    return newpositions, xcorr, deltas, inliers


def plot_individual_cross_correlation(
    images, masks, positions, overlaps, pixel_size, filename_template=None
):
    """
    Generate a plot showing the two masks and the final aligned images for each pair of overlapping tiles.

    Parameters:
    -----------
    images : list of numpy.ndarray
        List of image tiles.
    masks : list of numpy.ndarray
        List of binary masks corresponding to the image tiles.
    positions : numpy.ndarray
        Array of shape (N, 2) containing the (x, y) coordinates for positioning each image in units of microns.
    overlaps : list of tuples
        List of pairs of indices representing which tiles overlap and should be aligned.
    pixel_size : float
        The pixel size in microns.
    binning : int, optional
        Binning factor to reduce the image size. Default is 1.

    Returns:
    --------
    None
    """
    pixels = images[0].shape

    # One figure per overlapping pair: both tiles' masks, then a two-tile
    # stitch using the freshly measured shift, as a visual sanity check
    for i, j in overlaps:
        x1 = positions[i]
        x2 = positions[j]

        # Same overlap-region geometry as find_overlaps/cross_correlate_tiles
        dx = [int(x) for x in (x2 - x1) / pixel_size * 1e4][::-1]
        i1, i2 = array_overlap(dx[0], pixels[0])
        j1, j2 = array_overlap(dx[1], pixels[1])
        reference_mask = np.zeros_like(masks[i], dtype=bool)
        reference_mask[i1[0] : i1[1], j1[0] : j1[1]] = np.logical_and(
            masks[i][i1[0] : i1[1], j1[0] : j1[1]],
            masks[j][i2[0] : i2[1], j2[0] : j2[1]],
        )
        moving_mask = np.zeros_like(masks[j], dtype=bool)
        moving_mask[i2[0] : i2[1], j2[0] : j2[1]] = np.logical_and(
            masks[i][i1[0] : i1[1], j1[0] : j1[1]],
            masks[j][i2[0] : i2[1], j2[0] : j2[1]],
        )

        # Full-tile (not cropped-to-overlap) cross-correlation, unlike the
        # production path in cross_correlate_tiles -- fine for a one-pair-at-
        # a-time debug plot, too slow to use for every overlap in a montage
        detected_shift = np.asarray(
            phase_cross_correlation(
                images[i],
                images[j],
                reference_mask=reference_mask,
                moving_mask=moving_mask,
            )[0]
        )

        fig = plt.figure(figsize=(8, 12))
        axes = fig.subplot_mosaic([["Image1", "Image2"], ["Stitched", "Stitched"]])
        gs = fig.add_gridspec(3, 1, height_ratios=[1, 1, 2], hspace=0.3)

        axes["Image1"].imshow(
            images[i],
            cmap="gray",
            vmin=np.percentile(images[i][masks[i]], 1),
            vmax=np.percentile(images[i][masks[i]], 99),
        )
        axes["Image1"].imshow(reference_mask, alpha=0.5, cmap="Reds")
        axes["Image1"].set_title(f"Image {i} with Mask")

        axes["Image2"].imshow(
            images[j],
            cmap="gray",
            vmin=np.percentile(images[j][masks[j]], 1),
            vmax=np.percentile(images[j][masks[j]], 99),
        )
        axes["Image2"].imshow(moving_mask, alpha=0.5, cmap="Reds")
        axes["Image2"].set_title(f"Image {j} with Mask")
        # Place tile j relative to tile i using the shift just measured
        # (not the microscope's nominal position), then stitch just this pair
        stitched_positions = np.array(
            [
                positions[i],
                positions[i] + detected_shift[::-1] * pixel_size * 1e-4,
            ]
        )
        clipped_masks = clip_masks_to_overlaps(
            [masks[i], masks[j]],
            stitched_positions,
            [(0, 1)],
            pixels,
            pixel_size,
        )
        aligned_image, _ = stitch(
            [images[i], images[j]],
            stitched_positions,
            clipped_masks,
            pixel_size,
            smooth=False,
        )
        # aligned_image = np.roll(images[j], shift=(-int(detected_shift[0]), -int(detected_shift[1])), axis=(0, 1))
        # axes[1, 0].imshow(images[i], cmap='gray')
        # axes[1, 0].set_title(f"Image {i}")

        axes["Stitched"].imshow(
            aligned_image,
            vmin=np.percentile(images[j][masks[j]], 1),
            vmax=np.percentile(images[j][masks[j]], 99),
            cmap="gray",
        )
        axes["Stitched"].set_title(f"Aligned Image {j}")

        for a in axes.values():
            a.axis("off")

        if filename_template is not None:
            fig.savefig(filename_template.format(i, j))
        else:
            plt.tight_layout()
            plt.show()
        # clear() empties the figure but leaves it registered with pyplot, so a
        # run over many pairs accumulates every figure it ever made.
        plt.close(fig)


def parse_image_shifts(file_path, superres=1):
    """Parse an image shift file and return image shifts in units of pixels"""
    with open(file_path, "r") as file:
        lines = file.readlines()
    ntilts = int(lines[0].strip())
    result = []
    index = 1
    # while index < len(lines):
    for _ in range(ntilts):
        # Parse the tilt angle
        tilt_angle = float(lines[index].strip())
        index += 1

        # Parse the number of rows
        n = int(lines[index].strip())
        index += 1

        # Parse the n x 3 array
        array = []
        for _ in range(n):
            array.append([float(x) for x in lines[index].strip().split()])
            index += 1
        array = np.asarray(array)
        array[:, :2] /= superres
        # Add the parsed data to the result
        result.append((tilt_angle, array))

    # Sort the list by tilt angle and return
    return sorted(result, key=lambda x: x[0])


def generate_image_file_names_from_template(name, datadir, tilt, positions):
    # One path per tile index at this tilt, following the {name}_{tilt}_{tile
    # index}.mrc convention
    imagefiles = [
        os.path.join(datadir, "{0}_{1}_{2}.mrc".format(name, tilt, x))
        for x in range(positions.shape[0])
    ]

    return imagefiles


def _resolve_output_target(output_arg):
    """Classify the ``-o/--output`` argument into an output mode.

    Returns
    -------
    mode : {"dir", "mrc", "tiff"}
    path : str or None
        Directory path (mode "dir", or None to derive from --input),
        MRC stack path (mode "mrc"), or single TIFF path (mode "tiff").
    slice_index : int or None
        Z-slice parsed from a ``:N`` suffix in mode "mrc"; otherwise None.
    """
    if output_arg is None:
        return "dir", None, None
    base = os.path.basename(output_arg)
    # A ":N" suffix only counts if what precedes it is actually an .mrc path
    # and N is a plain integer -- otherwise a colon is just part of the path
    if ":" in base:
        head, idx_str = output_arg.rsplit(":", 1)
        if (
            os.path.basename(head).lower().endswith(".mrc")
            and idx_str.lstrip("-").isdigit()
        ):
            return "mrc", head, int(idx_str)
    if base.lower().endswith(".mrc"):
        return "mrc", output_arg, None
    if base.lower().endswith((".tif", ".tiff")):
        return "tiff", output_arg, None
    return "dir", output_arg, None


def setup_outputdir(args):
    """Create and return the directory the per-tilt images are written to.

    Only used in ``"dir"`` output mode (see :func:`_resolve_output_target`);
    the ``.mrc``/``.tiff`` modes take the dirname of the target instead.

    With no ``-o``, the name is derived from ``--input`` by swapping the
    ``*.mrc`` of the glob for ``_output``, so ``/data/Series_A_*.mrc`` gives a
    relative ``Series_A__output`` beside the *current working directory* — not
    beside the inputs. An input without that literal suffix (a directory of
    per-tile MRCs, say) keeps its own basename, which would then be created in
    the CWD under the same name. Pass ``-o`` and none of that applies.

    ``os.mkdir``, not ``makedirs``: a missing parent is an error worth seeing
    rather than a tree quietly conjured from a typo.
    """
    if args["output"] is None:
        outputdir = os.path.split(args["input"])[1].replace("*.mrc", "_output")
    else:
        outputdir = args["output"]
    logger.info("Results will be written to: %s", outputdir)
    if not os.path.exists(outputdir):
        os.mkdir(outputdir)
    return outputdir


def plot_positions(coordinates, fnam=None, fig=None, color="blue"):
    # Extract X and Y coordinates
    x_coords = [coord[0] for coord in coordinates]
    y_coords = [coord[1] for coord in coordinates]

    # Create a scatter plot
    if fig is None:
        fig, ax = plt.subplots(figsize=(10, 8))
    else:
        ax = fig.get_axes()[0]
        ax.title("Coordinate Points with Index Annotations")
        ax.xlabel("X Coordinates")
        ax.ylabel("Y Coordinates")
        ax.axhline(0, color="black", linewidth=0.5)
        ax.axvline(0, color="black", linewidth=0.5)
        ax.grid(color="gray", linestyle="--", linewidth=0.5)
    ax.scatter(x_coords, y_coords, color=color, label="Coordinates")

    # Annotate each point with its index
    for idx, (x, y) in enumerate(coordinates):
        ax.annotate(
            str(idx),
            (x, y),
            textcoords="offset points",
            xytext=(5, 5),
            ha="center",
            fontsize=8,
        )

    # Set plot title and labels

    ax.legend()

    # Save the plot to a PDF file
    fig.tight_layout()
    if fnam is None:
        plt.show()
    else:
        fig.savefig(fnam)


def plot_positions(coordinates, fnam=None, fig=None, color="blue"):

    # Extract X and Y coordinates
    x_coords = [coord[0] for coord in coordinates]
    y_coords = [coord[1] for coord in coordinates]

    # Create a scatter plot
    if fig is None:
        fig, ax = plt.subplots(figsize=(10, 8))
    else:
        ax = fig.get_axes()[0]
        ax.title("Coordinate Points with Index Annotations")
        ax.xlabel("X Coordinates")
        ax.ylabel("Y Coordinates")
        ax.axhline(0, color="black", linewidth=0.5)
        ax.axvline(0, color="black", linewidth=0.5)
        ax.grid(color="gray", linestyle="--", linewidth=0.5)
    ax.scatter(x_coords, y_coords, color=color, label="Coordinates")

    # Annotate each point with its index
    for idx, (x, y) in enumerate(coordinates):
        ax.annotate(
            str(idx),
            (x, y),
            textcoords="offset points",
            xytext=(5, 5),
            ha="center",
            fontsize=8,
        )

    # Set plot title and labels

    ax.legend()

    # Save the plot to a PDF file
    fig.tight_layout()
    if fnam is None:
        plt.show()
    else:
        fig.savefig(fnam)

# ---------------------------------------------------------------------------
# Provenance sidecars
# ---------------------------------------------------------------------------
#
# This runs one SLURM array task per tilt (--tilt-index with --output
# stack.mrc:N), so every task would be writing to the same canvas. Each writes
# only its own <canvas>.sidecars/<tilt>.json, which no other task touches —
# that is the whole reason there is no locking, no spool and no finalize pass
# here any more. The step-level parameters are identical across tasks and are
# repeated in each tilt's sidecar rather than hoisted into a shared file.


def series_base_for_stack(path: str) -> str:
    """Tilt series base name from a per-tilt stack name, ``{base}_{tilt}.mrc``.

    Nothing in this module needs it now that sidecars are named after the
    output file rather than after the series — it is kept because
    ``montage_backfill`` groups per-tilt stacks by series with it.
    """
    stem = os.path.splitext(os.path.basename(path))[0]
    return re.sub(r"_-?\d+\.?\d*$", "", stem)


def resolve_input_frame(files: list, spec: str) -> str:
    """Decide whether the input tiles are raw or motion-corrected.

    ``auto`` asks the files themselves: a tile stack that motion correction
    produced carries a ``motion_correction`` sidecar, and nothing else does.
    That is a stronger test than the manifest's was — it looked for a
    motion_correction *section* in a shared file, which could be present
    because some other run put it there, whereas this is a property of the
    exact files being read.

    Returning ``unknown`` is deliberate. It makes ``rotate_deg_from_raw_frames``
    null in the record rather than guessing zero, and a null that a reader has
    to handle is much cheaper than a plausible wrong number.
    """
    if spec != "auto":
        return spec
    # Checking a handful of files is enough -- they all come from the same
    # motion-correction run (or none did), no need to read every tile's sidecar
    for f in files[:8]:
        path = f if isinstance(f, str) else getattr(f, "name", None)
        if not path:
            continue
        doc = sidecar.read_for(path)
        if doc and doc.get("step") == "motion_correction":
            logger.info(
                "--input-frame auto: %s has a motion_correction sidecar, so "
                "these tiles are motion-corrected.", os.path.basename(path))
            return "motion-corrected"
    logger.warning(
        "--input-frame auto: no motion_correction sidecar on the input tiles, "
        "so it is not known whether they were motion-corrected. Recording the "
        "total rotation from raw frames as unknown. Pass --input-frame "
        "explicitly to settle it: 'raw' if these came straight off the "
        "detector, 'motion-corrected' if motion correction ran without "
        "writing sidecars.")
    return "unknown"


def verify_stitch_inputs(files: list, input_frame: str) -> None:
    """Warn if the input tiles are not the ones motion correction wrote.

    Only ever warns. A tile legitimately re-made between the two steps is a
    normal thing to do; a tile *silently* re-made with different parameters is
    the failure, and the difference between the two is a human decision.
    """
    if input_frame == "raw":
        return
    checked = changed = missing = 0
    # Re-fingerprint every tile and compare against what motion correction's
    # own sidecar recorded writing, tallying rather than warning per-file
    for f in files:
        path = f if isinstance(f, str) else getattr(f, "name", None)
        if not path or not os.path.exists(path):
            continue
        doc = sidecar.read_for(path)
        if doc is None:
            missing += 1
            continue
        checked += 1
        recorded = doc.get("output_fingerprint") or {}
        if recorded.get("sha1"):
            now = fingerprint(path)
            if now["sha1"] != recorded["sha1"]:
                changed += 1
    if missing:
        logger.info("Sidecars: %d input stack(s) have no motion_correction "
                    "sidecar; their provenance stops here.", missing)
    if changed:
        logger.warning(
            "Sidecars: %d of %d input stack(s) differ from what motion "
            "correction recorded writing. They were re-made after that step "
            "ran; make sure it was with the parameters you want.",
            changed, checked)
    elif checked:
        logger.info("Sidecars: all %d input stack(s) match motion "
                    "correction's record.", checked)


def stitch_effective(
    args: dict,
    *,
    pixel_size: float,
    binning: int,
    montage_origin_A,
    montage_width_A,
    canvas_shape_px,
    n_tilts_total: int,
    template_mask_path: str | None,
    output_mode: str,
    output_target: str | None,
    output_dir: str,
    input_frame: str,
    upstream_rotate_deg: int | None,
) -> tuple[dict, dict, list]:
    """This step's effective parameters, its requested ones, and what was auto.

    Identical for every tilt, and repeated into every tilt's sidecar: these are
    a couple of dozen scalars and the repetition is what lets any single
    sidecar answer "what geometry made this canvas" without opening a second
    file.

    ``auto_resolved`` names the values nobody chose — the pixel size read out
    of an MRC header, the ROI derived from the tile positions, the consensus
    tolerance derived from the data rather than given by
    ``--consensus-threshold``.
    """
    auto_resolved = []
    if args.get("pixel_size") is None:
        auto_resolved.append("pixel_size_A")
    if args.get("consensus_threshold") is None and args["consensus"] != "lstsq":
        # Derived per tilt from the spread of the corrections; the value that
        # was actually used is in each tilt's own record, not here.
        auto_resolved.append("consensus_threshold_um")
    if args.get("ROI") is None:
        auto_resolved.append("roi_px")

    # After motion correction, --rotate here is an *extra* rotation on top of
    # whatever that step already baked into the tiles. Neither step's own
    # value tells you how the canvas sits relative to the detector; the sum
    # does, and that is the number that had to be recovered by measuring tile
    # footprints. Run first, on raw tiles, this step's own value *is* the sum.
    rotate_deg = int((args["rotate"] // 90) % 4) * 90
    if input_frame == "raw":
        cumulative_rotate = rotate_deg
    elif upstream_rotate_deg is None:
        cumulative_rotate = None
        auto_resolved.append("rotate_deg_from_raw_frames")
    else:
        cumulative_rotate = (int(upstream_rotate_deg) + rotate_deg) % 360

    effective = {
        "pixel_size_A": float(pixel_size),
        "binning": int(binning),
        # Which of the two positions in the pipeline this run occupies, and
        # therefore how rotate_deg_from_raw_frames below was arrived at.
        "input_frame": input_frame,
        "canvas_pixel_size_A": float(pixel_size) * int(binning),
        # The value that was commented out of 02_stitch.sh and cost a day —
        # and, next to it, the total that nobody was recording at all. None
        # when motion correction left no record to compose with.
        "rotate_deg": rotate_deg,
        "rot90_k": int((args["rotate"] // 90) % 4),
        "rotate_deg_from_raw_frames": cumulative_rotate,
        "roi_px": list(args["ROI"]) if args.get("ROI") else None,
        "montage_origin_A": [float(v) for v in montage_origin_A],
        "montage_width_A": [float(v) for v in montage_width_A],
        "canvas_shape_px": [int(v) for v in canvas_shape_px],
        "image_shifts_file": os.path.abspath(args["image_shifts"]),
        "image_shift_superres_factor": float(args["correctimageshiftfilefactor"]),
        "skipcrosscorrelation": bool(args["skipcrosscorrelation"]),
        "consensus": args["consensus"],
        "consensus_threshold_um": args["consensus_threshold"],
        "max_allowed_imshift_correction_um": args["max_allowed_imshift_correction"],
        "ransac_iterations": int(args["ransac_iterations"]),
        "ransac_seed": int(args["ransac_seed"]),
        "irls_loss": args["irls_loss"],
        "fringe_size": int(args["fringe_size"]),
        "mask_threshold_fraction_of_median": args["maskthreshold"],
        "mask_absolute_threshold": args["maskabsolutethreshold"],
        "template_mask": template_mask_path,
        "min_mean_intensity": args["min_mean_intensity"],
        "correct_beam_edges": args["correct_beam_edges"],
        "plasmon_energy_eV": args["plasmon_energy"],
        "voltage_kV": args["voltage"],
        "smooth": not args["nosmooth"] and not args["mark_uncovered"],
        "mark_uncovered": bool(args["mark_uncovered"]),
        "output_mode": output_mode,
        # In "dir" mode the target may be unset and derived from --input, so
        # record where the tilts were actually written either way.
        "output_target": (
            os.path.abspath(output_target) if output_target else None),
        "output_dir": os.path.abspath(output_dir),
        "n_tilts_total": int(n_tilts_total),
        "auto_resolved": auto_resolved,
    }
    requested = {
        "pixel_size_A": args.get("pixel_size"),
        "input_frame": args["input_frame"],
        "rotate_deg": args["rotate"],
        "binning": args["binning"],
        "consensus_threshold_um": args.get("consensus_threshold"),
        "roi_px": list(args["ROI"]) if args.get("ROI") else None,
    }
    return effective, requested, auto_resolved


def upstream_rotate_from_sidecars(files: list) -> int | None:
    """``rotate_deg`` motion correction applied to these tiles, if recorded."""
    for f in files[:8]:
        path = f if isinstance(f, str) else getattr(f, "name", None)
        if not path:
            continue
        doc = sidecar.read_for(path)
        if doc and doc.get("step") == "motion_correction":
            value = (doc.get("effective") or {}).get("rotate_deg")
            if value is not None:
                return int(value)
    return None


def _tile_stack_input(stack, *, fingerprint_files: bool = False) -> list:
    """Describe a per-tile MRC directory input for one tilt.

    One tilt is tens of files here rather than the single stack the glob input
    mode gives, so this summarises them the way motion correction summarises
    its raw frames: a directory entry, which :func:`sidecar.verify_inputs`
    skips instead of trying to fingerprint a folder.

    ``fingerprint_files`` adds one fingerprint per tile. That is off for
    motion-corrected input — those files have their own sidecars, so the chain
    already covers them — but when stitching is the *first* step these files
    are the provenance root and no other record covers them at all.
    """
    paths = list(getattr(stack, "_paths", []))
    if not paths:
        return []
    directory = os.path.dirname(os.path.abspath(paths[0]))
    entry = sidecar.directory_input(directory, pattern="*.mrc", role="tiles")
    entry["n_files"] = len(paths)
    entry["note"] = "per-tile MRC input; one file per tile of this tilt"
    out = [entry]
    if fingerprint_files:
        out.extend(fingerprint_many(sorted(paths), role="tile"))
    return out


def tilt_sidecar_record(
    *,
    tilt_deg: float,
    z_slice,
    positionfile: str,
    pixel_size: float,
    binning: int,
    montage_origin_A,
    canvas_shape_px,
    nominal_positions_um,
    refined_positions_um,
    tile_selection,
    excluded_tiles,
    consensus_stats: dict,
    rotate: int,
) -> dict:
    """One tilt's own results: where its tiles ended up and what moved them.

    Positions are stored for *every* tile, selected or not, with the boolean
    selection alongside — the same contract as the position file, and the
    reason ``index_map`` needs ``flatnonzero(tile_selection)`` to be read.
    That indirection is named here rather than left to be rediscovered.
    """
    origin = np.asarray(montage_origin_A, dtype=float)
    scale = float(pixel_size) * int(binning)
    nominal = np.asarray(nominal_positions_um, dtype=float)[:, :2]
    refined = np.asarray(refined_positions_um, dtype=float)[:, :2]
    selection = np.asarray(tile_selection, dtype=bool)

    def to_canvas_px(positions_um):
        return (positions_um * 1e4 - origin) / scale

    # Corrections are only meaningful for tiles that were actually refined.
    correction_um = np.linalg.norm(refined - nominal, axis=1)
    refined_only = correction_um[selection] if selection.any() else correction_um

    return {
        "tilt": f"{tilt_deg:g}",
        "tilt_deg": float(tilt_deg),
        "z_slice": -1 if z_slice is None else int(z_slice),
        "position_file": os.path.abspath(positionfile),
        "n_tiles": int(selection.size),
        "n_tiles_used": int(selection.sum()),
        "excluded_tiles": [int(i) for i in excluded_tiles],
        "tile_selection": selection,
        # Both frames, both named. Microns is what the position file and the
        # image shift file speak; canvas pixels is what the montage is in.
        "positions_nominal_um": nominal,
        "positions_canvas_px": to_canvas_px(refined),
        "positions_nominal_canvas_px": to_canvas_px(nominal),
        "correction_um_max": float(refined_only.max()) if refined_only.size else 0.0,
        "correction_um_median": (
            float(np.median(refined_only)) if refined_only.size else 0.0),
        "canvas_shape_px": [int(v) for v in canvas_shape_px],
        "rotate_applied_deg": int((rotate // 90) % 4) * 90,
        # The consensus fit this tilt's positions actually rest on. threshold
        # is the derived one when --consensus-threshold was not given.
        "consensus": {
            "method": consensus_stats.get("method"),
            "threshold_um": consensus_stats.get("threshold", float("nan")),
            "n_edges": consensus_stats.get("n_edges", 0),
            "n_inliers": consensus_stats.get("n_inliers", 0),
            "n_bridges": consensus_stats.get("n_bridges", 0),
        },
        "index_map_labels_are": "selected_subset",
        "index_map_note": (
            "index_map in the position file labels tiles 0..n_selected-1, "
            "i.e. rows of positions_canvas_px[tile_selection]. Map back with "
            "np.flatnonzero(tile_selection)[label]."
        ),
    }


def write_tilt_sidecar(
    *,
    output_path: str,
    image,
    tilt_deg: float,
    step_effective: dict,
    step_requested: dict,
    tilt_record: dict,
    fingerprint_inputs: bool,
) -> None:
    """Write one tilt's sidecar beside the canvas it was stitched into.

    Keyed by tilt, so an array task writes a path no sibling task shares. The
    step's parameters ride along in ``effective`` next to this tilt's own
    results, which is what makes the file answer both "what was this run's
    geometry" and "where did this tilt's tiles land".
    """
    if isinstance(image, str):
        inputs = [fingerprint(image, role="tile_stack")]
    else:
        inputs = _tile_stack_input(image, fingerprint_files=fingerprint_inputs)

    sidecar.write(
        output_path,
        SIDECAR_STEP,
        {**step_effective, **tilt_record},
        inputs=inputs,
        key=tilt_record["tilt"],
        requested=step_requested,
        # The canvas is a shared stack that other tasks are still writing
        # slices into, so its size and head bytes are not stable while this
        # runs. Fingerprinting it here would record a value that is wrong by
        # the time the run finishes.
        fingerprint_output=False,
    )
def main():
    args = parse_commandline()

    if args["verbose"]:
        log_level = logging.DEBUG
    elif args["quiet"]:
        log_level = logging.WARNING
    else:
        log_level = logging.INFO
    logging.basicConfig(
        level=log_level,
        format="%(asctime)s  %(levelname)-8s  %(message)s",
        datefmt="%H:%M:%S",
    )

    output_mode, output_target, output_slice = _resolve_output_target(args["output"])

    if output_mode == "dir":
        outdir = setup_outputdir(args)
        mrc_path = None
        tiff_path = None
    else:
        outdir = os.path.dirname(output_target) or "."
        os.makedirs(outdir, exist_ok=True)
        logger.info("Results will be written to: %s", output_target)
        mrc_path = output_target if output_mode == "mrc" else None
        tiff_path = output_target if output_mode == "tiff" else None

    # Binning constant
    binning = int(args["binning"])

    imageshifts = parse_image_shifts(
        args["image_shifts"], superres=args["correctimageshiftfilefactor"]
    )

    if os.path.isdir(args["input"]):
        # ── Directory of per-tile MRC files from beam_mask_motioncorr ──────────
        # Files named  {stem}_{tile_index}_{tilt_angle}.mrc  are grouped by tilt.
        tilts, _tile_paths = _group_per_tile_mrcs(args["input"])
        if not tilts:
            raise FileNotFoundError(
                "No per-tile MRC files matching the expected naming pattern "
                f"found in directory: {args['input']}"
            )
        files = [TileStack(_tile_paths[t]) for t in tilts]
        logger.debug(
            "Directory input: %d tilt(s), %d tile(s) each",
            len(tilts),
            len(files[0]),
        )

        # Pixel size: prefer explicit --pixel-size, then MRC voxel_size header
        if args["pixel_size"] is not None:
            pixelsize = args["pixel_size"]
        else:
            with mrcfile.mmap(
                _tile_paths[tilts[0]][0], mode="r", permissive=True
            ) as _m:
                pixelsize = float(_m.voxel_size.x)
            if pixelsize == 0.0:
                raise ValueError(
                    "MRC voxel_size is not set; provide --pixel-size explicitly."
                )
        logger.debug(
            "Pixel size: %.4f Å  |  binning: %d  |  effective: %.4f Å",
            pixelsize,
            binning,
            pixelsize * binning,
        )

    else:
        # ── Glob of per-tilt MRC stacks (original path) ─────────────────────────
        files = sorted(glob.glob(args["input"]))
        if len(files) < 1:
            raise FileNotFoundError("No files matching {0}".format(args["input"]))
        logger.debug("Found %d MRC file(s) matching '%s'", len(files), args["input"])

        # Pixel size: prefer explicit --pixel-size, then MRC voxel_size header
        if args["pixel_size"] is not None:
            pixelsize = args["pixel_size"]
        else:
            with mrcfile.mmap(files[0], mode="r", permissive=True) as _m:
                pixelsize = float(_m.voxel_size.x)
            if pixelsize == 0.0:
                raise ValueError(
                    "MRC voxel_size is not set; provide --pixel-size explicitly."
                )
        logger.debug(
            "Pixel size: %.4f Å  |  binning: %d  |  effective pixel size: %.4f Å",
            pixelsize,
            binning,
            pixelsize * binning,
        )

        # Tilt angle is parsed straight from each filename, e.g.
        # Montage_182-A_-12.0.mrc -> -12.0. -0.0 normalises to 0.0 so tiles
        # at zero tilt aren't treated as a distinct angle.
        tilts = []
        for f in files:
            m = re.search(r"_(-?\d+\.?\d*)\.mrc$", os.path.basename(f))
            if not m:
                raise ValueError(
                    f"Cannot parse tilt angle from filename: {f} "
                    "(expected it to end in '_{tilt_angle}.mrc')"
                )
            t = float(m.group(1))
            tilts.append(0.0 if t == -0.0 else t)

    # Get image shifts from file
    # TODO make SerialEM put this in the mdoc file in microns (not
    # weird TFS units) to obviate this step

    tiltsfromfile = [imageshifts[x][0] for x in range(len(imageshifts))]
    imageshifts = [imageshifts[x][1] for x in range(len(imageshifts))]
    reshapedimshifts = np.concatenate(imageshifts)[:, :2] * pixelsize

    if args["ROI"] is None:
        tile_shape = np.asarray(_first_tile_shape(files[0]))
        if (args["rotate"] // 90) % 2:
            # A 90°/270° rotation swaps each tile's height and width.
            tile_shape = tile_shape[::-1]
        tileFOV = pixelsize * tile_shape
        # Global range of tiles in Angstroms
        globalwidth = np.ptp(reshapedimshifts, axis=0) + tileFOV
        globalorigin = np.amin(reshapedimshifts, axis=0)  # +tileFOV
    else:
        roi = [float(x) for x in args["ROI"]]
        # Convert ROI from pixels (similar to image shifts file) to Angstroms)
        globalorigin = np.array([roi[0], roi[2]]) * pixelsize
        globalwidth = np.array([roi[1] - roi[0], roi[3] - roi[2]]) * pixelsize

    canvas_px = (globalwidth / pixelsize / binning).astype(int)
    logger.debug(
        "Montage canvas: %d × %d px (binned)  |  %.1f × %.1f µm",
        canvas_px[0],
        canvas_px[1],
        globalwidth[0] * 1e-4,
        globalwidth[1] * 1e-4,
    )
    logger.info("Processing %d tilt(s)", len(files))

    # --output pointing at an MRC stack (mode "mrc") writes directly into a
    # shared per-series MRC stack instead of the per-tilt TIFFs, skipping
    # JointoMRC.py. With no --tilt-index, every discovered tilt is stitched
    # into its own z-slice in this one run; --tilt-index (or a ':N' suffix)
    # restricts the run to a single slice, for one SLURM array task per
    # tilt. mode "tiff" writes a single named TIFF instead of a per-tilt
    # directory and always requires exactly one tilt to be selected.
    # z-slice assignment is independent of files/tilts iteration order — it's
    # always each tilt's 0-based rank when every discovered tilt angle is
    # sorted ascending, matching JointoMRC.py's default tilt-angle ordering.
    tilt_index = args["tilt_index"]
    if output_mode == "mrc" and output_slice is not None and tilt_index is None:
        tilt_index = output_slice

    n_tilts_total = len(tilts)
    z_index_map = {t: z for z, t in enumerate(sorted(tilts))}

    if tilt_index is None and output_mode == "tiff":
        if n_tilts_total != 1:
            raise ValueError(
                f"--output {args['output']!r} names a single TIFF output file "
                f"but {n_tilts_total} tilt(s) were discovered; pass "
                "--tilt-index to select one."
            )
        tilt_index = 0

    if tilt_index is not None:
        if not (0 <= tilt_index < n_tilts_total):
            raise ValueError(
                f"--tilt-index {tilt_index} out of range for "
                f"{n_tilts_total} discovered tilt(s)."
            )
        target_tilt = sorted(tilts)[tilt_index]
        keep = [k for k, t in enumerate(tilts) if t == target_tilt]
        files = [files[k] for k in keep]
        tilts = [tilts[k] for k in keep]
        logger.info(
            "Restricting to tilt index %d/%d  (tilt angle %.2f°)",
            tilt_index,
            n_tilts_total,
            target_tilt,
        )

    fringe_size = args["fringe_size"]

    # Load reference beam image if provided
    reference_beam = None
    if args["referencebeam"] is not None:
        tmask_path = args["referencebeam"]
        # Parse optional ":index" suffix, e.g. "file.mrc:7"
        tmask_index = None
        if ":" in os.path.basename(tmask_path):
            tmask_path, idx_str = tmask_path.rsplit(":", 1)
            tmask_index = int(idx_str)
        if tmask_path.lower().endswith(".mrc"):
            with mrcfile.open(tmask_path, "r") as m:
                data = np.asarray(m.data)
                reference_beam = data[tmask_index] if tmask_index is not None else data
        else:
            # Need to flip y of tiff files to match mrc conventions
            img = Image.open(tmask_path)
            if tmask_index is not None:
                img.seek(tmask_index)
            reference_beam = np.asarray(img)[::-1]

        input_tile_shape = tuple(int(x) for x in _first_tile_shape(files[0]))
        if reference_beam.shape[-2:] != input_tile_shape:
            logger.warning(
                "Reference beam %s has shape %s, which does not match the "
                "input tile shape %s; cropping/padding to match.",
                tmask_path,
                reference_beam.shape[-2:],
                input_tile_shape,
            )
            reference_beam = crop(reference_beam, input_tile_shape)
        if binning > 1:
            reference_beam = fourier_interpolate(
                reference_beam, [x // binning for x in reference_beam.shape]
            )
        logger.info("Loaded template mask from %s", tmask_path)

    # Resolved after the tilt selection, so a task restricted to one tilt still
    # records the whole series' parameters — which is why n_tilts_total is used
    # here rather than len(files). The step's parameters are computed once and
    # handed to every tilt, each of which writes them into its own sidecar.
    sidecar_step = None
    input_frame = None
    if args["sidecar"]:
        input_frame = resolve_input_frame(files, args["input_frame"])
        verify_stitch_inputs(files, input_frame)
        effective, requested, auto_resolved = stitch_effective(
            args,
            pixel_size=pixelsize,
            binning=binning,
            montage_origin_A=globalorigin,
            montage_width_A=globalwidth,
            canvas_shape_px=canvas_px[::-1],
            n_tilts_total=n_tilts_total,
            template_mask_path=args["referencebeam"],
            output_mode=output_mode,
            output_target=output_target,
            output_dir=outdir,
            input_frame=input_frame,
            upstream_rotate_deg=upstream_rotate_from_sidecars(files),
        )
        sidecar_step = (effective, requested)
        if auto_resolved:
            logger.info("Sidecar: auto-resolved %s", ", ".join(auto_resolved))

    _quiet = not logger.isEnabledFor(logging.INFO)
    for i, (file, tilt) in enumerate(
        tqdm(
            zip(files, tilts),
            total=len(files),
            desc="Stitching montages",
            disable=_quiet,
        )
    ):
        # _image_basename, not os.path.basename: with a directory of per-tile
        # MRCs `file` is a TileStack, and logging arguments are evaluated
        # whatever the log level, so basename() crashed the whole run.
        logger.debug(
            "[%d/%d] %s  (tilt %.1f°)", i + 1, len(files), _image_basename(file), tilt
        )
        indx = find_closest_index(tiltsfromfile, tilt)
        positions = imageshifts[indx]

        tiles = None
        logger.debug("  %d tile(s) selected", len(positions))
        # plot_positions(positions[tiles][:,:2],color='k')

        mont = montage(
            file,
            outdir,
            positions,
            pixelsize,
            binning=binning,
            skipcrosscorrelation=args["skipcrosscorrelation"],
            montagewidth=globalwidth,
            montageorigin=globalorigin,
            tiles=tiles,
            fringe_size=fringe_size,
            maxshift=args["max_allowed_imshift_correction"],
            consensus=args["consensus"],
            consensus_threshold=args["consensus_threshold"],
            ransac_iterations=args["ransac_iterations"],
            ransac_seed=args["ransac_seed"],
            irls_loss=args["irls_loss"],
            maskthreshold=args["maskthreshold"],
            maskabsolutethreshold=args["maskabsolutethreshold"],
            nthreads=args["nthreads"],
            positionfile=args["positionfile"],
            smooth=not args["nosmooth"] and not args["mark_uncovered"],
            mark_uncovered=args["mark_uncovered"],
            reference_beam=reference_beam,
            min_mean_intensity=args["min_mean_intensity"],
            correct_beam_edges=args["correct_beam_edges"],
            E_plasmon_eV=args["plasmon_energy"],
            voltage_kV=args["voltage"],
            rotate=args["rotate"],
            output_mrc=mrc_path,
            tilt_z_index=z_index_map[tilt] if mrc_path is not None else None,
            n_tilts_total=n_tilts_total if mrc_path is not None else None,
            fileout_path=tiff_path,
            sidecar_step=sidecar_step,
            sidecar_input_frame=input_frame,
            tilt_deg=tilt,
            series_z_index=z_index_map[tilt],
        )


if __name__ == "__main__":
    main()
```
