## Supplementary material for "Whole cryo-FIB lamella acquisition at high magnification with rectangular beam montage cryo-electron tomography": Source code: stitchbyadjacenttilt.py

```
import re
import networkx as nx
from scipy.ndimage import affine_transform
from tqdm import tqdm
from PIL import Image
import os
import sys
from skimage.registration import phase_cross_correlation

from scipy.spatial import KDTree
import matplotlib.pyplot as plt
import copy
import glob
import mrcfile
import argparse
import numpy as np
from typing import List, Tuple, Dict, Any
from .Utilities import *
from .smoothn import smoothn


def parse_commandline() -> Dict[str, Any]:
    """
    Parse commandline input.
    """
    parser = argparse.ArgumentParser(
        description="Stitch square beam montage tomography data."
    )
    parser.add_argument(
        "-i", "--input", help="*.mrc wildcard for raw data", required=True, type=str
    )
    parser.add_argument(
        "-i2", "--input2", help=".mrc file for that will be used as alignment reference", required=False, type=str,default='None',
    )
    parser.add_argument(
        "-o",
        "--output",
        help="Output directory, if left blank output will be placed in a folder named ./[input_filename]_output",
        required=False,
        type=str,
    )
    parser.add_argument(
        "-I",
        "--image_shifts",
        help="Path to text file containing the tilts and a list of image shifts at every tilt (requried).",
        required=True,
        type=str,
    )
    parser.add_argument(
        "-b",
        "--binning",
        help="Binning of input data, defaults to 1",
        required=False,
        type=int,
        default=1,
    )
    parser.add_argument(
        "-f",
        "--fringe_size",
        help="Size of Fresnel fringes at edge of beam, this will be removed from the gain reference (default 20).",
        required=False,
        type=int,
        default=20,
    )
    parser.add_argument(
        "-s",
        "--skipcrosscorrelation",
        help="Skip cross-correlation alignment of montage tiles and default to using imageshifts to stitch montage.",
        action="store_true",
    )
    parser.add_argument(
        "-t",
        "--tiltaxisrotation",
        help="Rotation of tilt axis relative to image, if not provided will take from .mdoc file.",
        type=float,
        required=False,
    )
    parser.add_argument(
        "-M",
        "--maximageshift",
        help="Limit the size of the total montage by excluding images in montage with image shift in microns greater than this value.",
        type=float,
        required=False,
    )
    parser.add_argument(
        "-m",
        "--max_allowed_imshift_correction",
        help="Maximum allowed correction to tile alignments by cross-correlation.",
        type=float,
        default=0.05,
        required=False,
    )

    parser.add_argument(
        "-S",
        "--correctimageshiftfilefactor",
        help="Sometimes imageshift file needs to be divided by a factor of 2 since serial-EM uses super-resolution pixels.",
        type=int,
        default=1,
        required=False,
    )
    parser.add_argument(
        "-mt",
        "--maskthreshold",
        help="threshold (as fraction of raw image median) for masking of beam.",
        type=float,
        default=0.4,
        required=False,
    )

    parser.add_argument(
        "-ma",
        "--maskabsolutethreshold",
        help="Absolute threshold (in number of image counts) for masking of beam.",
        type=float,
        default=None,
        required=False,
    )

    return vars(parser.parse_args())


def stitch(ims,positions,msks,pixel_size,binning=1,montagewidth=None,montageorigin=None,smooth=True,renormalize=False):
    # Determine the global range of tiles in Angstroms
    if montagewidth is None:
        width = np.ptp(positions, axis=0) * 1e4
    else:
        width = montagewidth
    if montageorigin is None:
        origin = np.amin(positions, axis=0) * 1e4
    else:
        origin = montageorigin

    pixels = ims[0].shape

    # Calculate the size of the global montage canvas in pixels
    size = (
        np.asarray(width / pixel_size / binning, dtype=int)[::-1]
        + pixels
        + np.asarray([1, 1])
    )

    # Initialize the montage canvas and overlap map
    canvas = np.zeros(size)
    overlap = np.zeros(size, dtype=np.uint8)

    # Place each image onto the canvas, applying the masks
    for i, (im, position, msk) in enumerate(
        zip(tqdm(ims, desc="Stitching"), positions, msks)
    ): 
        # s is shape of tile, size is shape of canvas
        s = im.shape
        # Desired coordinate of upper left of tile
        y0, x0 = [int(x) for x in (position * 1e4 - origin) / pixel_size / binning]
    

        # Skip tiles that have fallen off canvas
        if x0 > size[0] or y0 > size[1]:
            continue
        if x0 +s[0] <0 or y0 +s[1] <0:
            continue

        # Truncate canvas coordinate beginnings to be >= 0
        cy0, cx0 = [max(coord, 0) for coord in [y0, x0]]
        # Truncate canvas coordinate maximum to be <= canvas array limits
        cx1, cy1 = [
            min(coord, limit) for coord, limit in zip([x0 + s[0], y0 + s[1]], size)
        ]

        # Size of tile that will make it onto the montage canvas
        X = cx1 - cx0
        Y = cy1 - cy0

        # Coordinates of tile, if x0 (or y0) < 0 this implies some of the tile
        # falls off the left (or upper) edge of canvas so is not included
        tx0 = -min(x0,0)
        tx1 = tx0 + X
        ty0 = -min(y0,0)
        ty1 = ty0 + Y

        canvas[cx0:cx1, cy0:cy1] += np.where(msk, im, 0)[tx0:tx1,ty0:ty1]
        overlap[cx0:cx1, cy0:cy1] += np.where(msk, np.uint8(1), np.uint8(0))[tx0:tx1,ty0:ty1]

    # # Normalize the canvas by the overlap map to account for overlapping regions
    # median = np.median(canvas[overlap == 1])

    # positions_pixel = (positions * 1e4) / pixel_size / binning
    # for coords in np.nonzero(canvas[overlap>1]):
    #     # Find closest montage tile
    #     y_idx, x_idx = coords
    #     pt = np.array([x_idx, y_idx])
    #     closest_index = int(np.argmin(np.sum((positions_pixel - pt) ** 2, axis=1)))

    #     # closest_position = positions_pixel[closest_index]
    #     # closest_index is the index of the nearest tile; closest_position is its pixel coords

    if renormalize:
        canvas[overlap>0] /= overlap[overlap>0]
    if smooth:
        print("Smoothing background to montage")
        smoothed = smoothn(np.ma.masked_array(canvas,overlap <1),s=1e7,robust=True)
        canvas = np.where(overlap < 1, smoothed, canvas)
    else:
        canvas[overlap < 1] = np.median(canvas[overlap == 1])
           # fig,ax = plt.subplots()
    # 
    # plt.show(block=True)
        # fig,ax=plt.subplots()
        # p = positions * 1e4 / pixel_size / binning
        # ax.imshow(canvas,cmap='gray',origin='lower')
        # ax.plot(*p.T,'ro',linestyle='-',label='Montage tiles')
        # for i,pos in enumerate(p):
        #     ax.text(pos[0],pos[1],str(i))
        # plt.show(block=True)
    overlap = np.where(overlap > 1, overlap, 1)
    
    return canvas


def montage(
    image,
    alignfile,
    outdir,
    positions,
    refpositions,
    pixel_size,
    compression_factor,
    binning=8,
    skipcrosscorrelation=False,
    montagewidth=None,
    montageorigin=None,
    tiles=None,
    maxshift=0.05,
    fringe_size=20,
    maskthreshold=0.4,
    maskabsolutethreshold=None,
    tilt_axis=0.0
):
    """
    Creates a montage image from a series of input images by aligning and stitching them together.

    This function takes a set of images, positions them according to provided coordinates,
    and aligns them based on mask images or generated masks. The images are then combined
    into a single montage image, with optional gain reference correction.

    Parameters:
    -----------
    image : str
        mrc file containing montage tiles
    outdir : str
        directory for output
    positions : numpy.ndarray
        Array of shape (N, 2) containing the (x, y) coordinates for positioning each image in units of pixels.
        Note that the x,y have opposite convention to standard y,x Python convention
    pixel_size : float
        The pixel size in Angstroms.
    binning : int, optional
        Binning factor to reduce the image size. Default is 8.
    tiltaxisrotation : float, optional
        Rotation angle of the tilt axis, used to correct the positions. Default is 0.
    gainref : numpy.ndarray, optional
        Gain reference image used to normalize the input images. Default is None.
    montagewidth : None or (2,) array_like
        size in Angstrom of the full montage in both dimensions, useful for consistency
        with other tilts in the tilt series
    montageorigin : None or (2,) array_like
        Origin point (most negative image shift) in Angstrom of the full montage
        in both dimensions, useful for consistency with other tilts in the tilt
        series
    tiles : None or sliceobject, optional
        Slice object indicating tiles that will be stitched (mainly for testing purposes)
    gaincorrectedimagefile : None or string,optional
        Path to already gain corrected images, if not provided the program will generate
        the gain corrected images and write them to a mrc file in the output directory.
    fringe_size : float, optional
        Size of the fringe region in pixels for removal from gain reference. Default is 20.
    Matchgainrefmask : bool, optional
        If False, a mask will be generated for each tile. If True, masks will be generated
        by cross correlation alignment of the gain reference mask to each tile. Default is None.
    medianthreshold : float, optional
        Threshold for (as a fraction of the median) for masking the image. Default is 0.4.
    Returns:
    --------
    numpy.ndarray
        The resulting stitched montage image.
    """

    if Matchgainrefmask:
        assert (
            gainref is not None
        ), "Gain reference must be provided if Matchgainrefmask is True"

    # Load images as mrc memory map
    mrcmemmap = mrcfile.mrcmemmap.MrcMemmap(image)
    refmrcmemmap = mrcfile.mrcmemmap.MrcMemmap(alignfile)

    # Get the shape and data type of the montage tiles
    dtype = mrcmemmap.data.dtype
    pixels = np.asarray([x // binning for x in mrcmemmap.data.shape[-2:]], dtype=int)

    # M is the slice object describing the indices of the tiles to include
    # in the montage, if None is passed for tiles then all tiles are included
    # by default.
    if tiles is None:
        M = slice(0, len(mrcmemmap.data))
    else:
        M = tiles

    # Convert positions from pixels to microns, remove 3rd dimension
    positions = positions[:, :2] * pixel_size * 1e-4
    refpositions = refpositions[:, :2] * pixel_size * 1e-4

    # fig,ax = plt.subplots()
    # ax.plot(*positions[M].T,'ro',linestyle='-',label='Montage tiles')
    # for i,pos in enumerate(positions[M]):
    #     ax.text(pos[0],pos[1],str(i))
    # plt.show(block=True)

    # Compress positions of reference images to account for relative tilt
    img = copy.deepcopy(mrcmemmap.data[0])
    # print(tilt_axis,relative_tilt)
    # fig,ax = plt.subplots(ncols=2,nrows=2)
    # ax= ax.ravel()
    # ax[0].imshow(img,vmin=np.percentile(img,1),vmax=np.percentile(img,99),cmap='gist_gray')
    
    T = np.eye(2)
    T[1,1] = compression_factor 
    # if relative_tilt >= 0:
    #      np.cos(np.deg2rad(relative_tilt))
    # else:
    #     T[1,1] = 1/np.cos(np.deg2rad(relative_tilt))
    # o = np.asarray(img.shape)
    # M = np.linalg.inv(rotation_matrix(tilt_axis))
    # ax[1].imshow(affine_transform(img,M,offset =o),vmin=np.percentile(img,1),vmax=np.percentile(img,99),cmap='gist_gray')
    # M = np.linalg.inv(T @ rotation_matrix(tilt_axis))
    # ax[2].imshow(affine_transform(img,M,offset =o),vmin=np.percentile(img,1),vmax=np.percentile(img,99),cmap='gist_gray')
    # M = np.linalg.inv(rotation_matrix(-tilt_axis) @ T @ rotation_matrix(tilt_axis))
    # ax[3].imshow(affine_transform(img,M),vmin=np.percentile(img,1),vmax=np.percentile(img,99),cmap='gist_gray')
    # plt.show(block=True)
    T =  rotation_matrix(-tilt_axis) @ T @ rotation_matrix(tilt_axis)
    refpositions = refpositions @  T

    # Initialize an empty mask list
    msks = []

    ims = []  # List to store the processed images
    for position, img in tqdm(
        zip(positions[M], mrcmemmap.data[M]),
        total=len(positions[M]),
        desc="Generating masks",
    ):
        im = copy.deepcopy(np.asarray(img))
        if binning > 1:
            im = fourier_interpolate(im, [x // binning for x in im.shape])
        msks.append(make_mask(im, shrinkn=fringe_size / binning, medianthreshold=maskthreshold, absolutethreshold=maskabsolutethreshold))
        ims.append(im)

    positionfile = os.path.join(
        outdir, os.path.split(image)[1].replace(".mrc", "_refined_positions.h5")
    )

    refmsks = []
    # if binning <2:
    #     refims = refmrcmemmap.data
    # else:
    refims = []
    # Load image from memory map and convert to numpy array
    for  img in tqdm(refmrcmemmap.data,desc='Masking reference images'):
        im = copy.deepcopy(np.asarray(img))
        # Fourier interpolate if reqeusted
        if binning > 1:
            im = fourier_interpolate(im, [x // binning for x in im.shape])
        # Shrink image to account for extra tilt
        im = affine_transform(im,T,output_shape=pixels)
        refims.append(im)
        refmsks.append(make_mask(im, shrinkn=fringe_size / binning,medianthreshold=maskthreshold,absolutethreshold=maskabsolutethreshold))
        # fig,ax = plt.subplots(ncols=2)
        # ax[0].imshow(refims[-1])
        # ax[1].imshow(np.where(refmsks[-1],1,0))
        # plt.show(block=True)
    
    # Calculate the overlaps between adjacent tiles
    overlaps = find_overlaps(
        positions[M], pixels, pixel_size * binning, msks, plot=False
    )
    from scipy.spatial.distance import cdist
    distmatrix = cdist(positions[M],refpositions)
    reftile = []
    for overlap in overlaps:
        i, j = overlap
        # See which tile in the reference set minimizes the mutual distance
        k = np.argmin(distmatrix[i]**2+distmatrix[j]**2)
        reftile.append(k)

    
    if not skipcrosscorrelation:
        # Refine the tile positions using cross-correlation between overlapping
        # tiles
        original_positions = copy.deepcopy(positions)
        fileout = os.path.join(
            outdir, os.path.splitext(os.path.split(image)[1])[0] + "_cross_correlation_plots.pdf"
        )
        positions[M], xcorr, deltas = cross_correlate_tiles(
            positions[M],
            refpositions,
            ims,
            refims,
            msks,
            refmsks,
            overlaps,
            reftile,
            pixel_size * binning,
            max_correction=maxshift,
            generate_plot=fileout,
            # showplot=True,
        )

        save_array_to_hdf5(
            [original_positions, positions, xcorr, overlaps, np.asarray(deltas)],
            positionfile,
            [
                "Original_positions",
                "Refined_positions",
                "cross_correlations",
                "overlaps",
                "relative_shifts",
            ],
        )
        
    elif os.path.exists(positionfile):
        # If requested to skip cross correlation and an older set of tile
        # alignments exist then load these.
        positions = load_array_from_hdf5(positionfile, "Refined_positions")
        xcorr = load_array_from_hdf5(positionfile, "cross_correlations")
        original_positions = positions
    else:
        original_positions = positions
    # fig,ax = plt.subplots()
    # ax.plot(*positions[M].T,'ro',linestyle='-',label='Montage tiles')
    # for i,pos in enumerate(positions[M]):
    #     ax.text(pos[0],pos[1],str(i))
    # plt.show(block=True)
    # Clip masks to ensure overlapping regions are only taken from closest tile
    msks = clip_masks_to_overlaps(msks,positions[M],overlaps,pixels,pixel_size * binning,ims)

    # Stitche the images together using the refined tile positions
    canvas = stitch(
        ims,
        positions[M],
        msks,
        pixel_size,
        binning=binning,
        montagewidth=montagewidth,
        montageorigin=montageorigin,
    )
    fileout = os.path.join(
        outdir, os.path.splitext(os.path.split(image)[1])[0] + ".tif"
    )
    if not skipcrosscorrelation:
        ntiles = len(original_positions[M])
        figsize = 2 * (int(np.ceil(np.sqrt(ntiles))),)
        xcorfig, xcorax = plt.subplots(figsize=figsize)
        cmean = np.mean(canvas)
        cstd = np.std(canvas)
        xcorax.imshow(
            canvas,
            cmap=plt.get_cmap("gist_gray"),
            origin="lower",
            vmin=cmean - cstd,
            vmax=cmean + cstd,
        )
        s = np.asarray(ims[0].shape)[::-1]
        posi = (original_positions[M] * 1e4 - montageorigin) / pixel_size / binning
        posi += s / 2
        xcorax.plot(*posi.T, "ko", label="Initial tile positions")
        for i, pos in enumerate(posi):
            xcorax.annotate(str(i), pos)
        posi = (positions[M] * 1e4 - montageorigin) / pixel_size / binning
        posi += s / 2
        xcorax.plot(*posi.T, "bo", label="Refined tile positions")
        # xcorrmax = np.amax(xcorr)
        cmap = plt.get_cmap("viridis")
        for ind, (i, j) in enumerate(overlaps):
            # Retrieve image shifts for the overlapping tiles
            x1 = (original_positions[M][i] * 1e4 - montageorigin) / pixel_size / binning

            x2 = (original_positions[M][j] * 1e4 - montageorigin) / pixel_size / binning
            dx = [int(x) for x in (x2 - x1)][::-1]
            x1 += s / 2
            delta = (deltas[ind] * 1e4) / pixel_size / binning

            shifttoolarge = (
                np.linalg.norm(np.asarray(dx) - delta)
                > maxshift / pixel_size / binning * 1e4
            )
            if shifttoolarge:
                linestyle = "--"
            else:
                linestyle = "-"
            xcorax.plot(
                [x1[0], x1[0] + delta[1]],
                [x1[1], x1[1] + delta[0]],
                linestyle=linestyle,
                color="b"
            )
        plotfile = os.path.join(
            outdir, os.path.split(image)[1].replace(".mrc", "_Plot.pdf")
        )
        xcorfig.savefig(plotfile)
    # Note that Python Image library does not support write 16-bit integer and writes
    # 32-bit real instead ¯\_(ツ)_/¯
    Image.fromarray(canvas.astype(np.int16)).save(fileout)
    save_array_as_png(canvas, fileout.replace('.tiff','.png'), cmap=plt.get_cmap('Greys'))
    # return canvas


def plot_overlaps(positions, overlaps, show=True):
    m = len(positions)
    fig, ax = plt.subplots()
    ax.plot(*positions.T, "ko")
    for n, val in enumerate(overlaps):
        if val:
            i, j = condensed_to_square(n, m)
            x1 = positions[i]
            x2 = positions[j]
            ax.plot([x1[0], x2[0]], [x1[1], x2[1]], "r-")
    plt.show(block=show)
    return fig


def array_overlap(dx, n):
    """Overlapping indices for 1d array shifted dx relative to each other"""
    if dx > 0:
        return [dx, n], [0, n - dx]
    else:
        return [0, n + dx], [-dx, n]


def find_overlaps(positions, pixels, pixel_size, masks, minoverlapfrac=0.01, plot=True):
    """
    Identify pairs of images that overlap in a 2D montage, based on their positions
    and field of view, and further refine the overlap based on binary masks.

    Parameters:
    ----------
    positions : ndarray of shape (m, 2)
        The (x, y) coordinates of `m` images in a 2D montage. Note that
        the positions array is ordered (x,y) opposite to python standard (y,x)
    pixels : tuple or list of length 2
        Dimensions of each image in pixels, as (height, width).
    pixel_size : float
        The size of each pixel in microns.
    masks : list of ndarrays
        Binary masks for each image (same shape as image), used to check if
        a specific region of the images overlaps. If None, the function only
        considers the geometric overlap.
    minoverlapfrac : float, optional
        Minimum fraction of overlap (based on pixel area) required for two images
        to be considered overlapping. Default is 0.03 (i.e., 3%).
    plot : float, optional
    Returns:
    -------
    overlapping_inds : list of lists
        List of pairs of indices representing images that overlap.
        Each element is a list of two indices `[i, j]`, where the i-th and j-th
        images overlap.
    """
    m = len(positions)  # Number of images

    from scipy.spatial.distance import pdist

    # Calculate the size of each field of view in microns (height, width)
    # Order of pixels array has to be reversed from python standard to match
    # positions array
    criterion = pixel_size * np.asarray(pixels)[::-1] * 1e-4

    # Calculate (unique) distances between montage pieces
    # with scipy's pdist (pair-wise distance) function and
    # calculate which are within a "field of view" of each other
    # result is stored in a condensed distance matrix

    overlaps = np.logical_and(
        *[pdist(positions[..., i : i + 1]) / criterion[i] < 1.0 for i in range(2)]
    )

    # Visualize the positions and overlap status
    if plot:
        overlapfig = plot_overlaps(positions, overlaps, show=False)
        overlapax = overlapfig.get_axes()[0]

    # List to store the indices of overlapping image pairs
    overlapping_inds = []

    # Update criterion to be the minimum required overlap area in pixels
    criterion = minoverlapfrac * np.prod(pixels)

    # Iterate over the condensed distance matrix
    for n, val in enumerate(overlaps):
        if val:  # If overlap is detected
            # Convert the index in the condensed matrix back to the square form
            i, j = condensed_to_square(n, m)
            x1 = positions[i]
            x2 = positions[j]

            # Calculate the relative shift in position between the two images, in pixels
            dx = [int(x) for x in (x2 - x1) / pixel_size * 1e4]

            # Determine the overlapping pixel regions in both images
            i1, i2 = array_overlap(dx[1], pixels[0])
            j1, j2 = array_overlap(dx[0], pixels[1])

            # Check if the overlap area in the masks exceeds the minimum criterion
            overlap = (
                np.sum(
                    np.logical_and(
                        masks[i][i1[0] : i1[1], j1[0] : j1[1]],
                        masks[j][i2[0] : i2[1], j2[0] : j2[1]],
                    )
                )
                >= criterion
            )
            if overlap:
                overlapping_inds.append([i, j])
                if plot:
                    x1 = positions[i]
                    x2 = positions[j]
                    overlapax.plot([x1[0], x2[0]], [x1[1], x2[1]], "b--")
    if plot:
        plt.show(block=True)
    return overlapping_inds

def clip_masks_to_overlaps(masks, positions, overlaps, pixels, pixel_size,ims):
    """
    Clip binary masks so when there is overlap, pixels are taken from the closest
    tile to the pixel in the overlapping region.

    Parameters:
    -----------
    masks : list of ndarrays
        List of binary masks for each image tile.
    positions : ndarray of shape (N, 2)
        Array of (x, y) coordinates for each image tile in microns.
    overlaps : list of tuples
        List of pairs of indices representing which tiles overlap.
    pixels : tuple or list of length 2
        Dimensions of each image tile in pixels, as (height, width).
    pixel_size : float
        The size of each pixel in microns.

    Returns:
    --------
    clipped_masks : list of ndarrays
        List of modified binary masks with overlapping regions clipped to 
        closest tile.
    """
    clipped_masks = [copy.deepcopy(msk) for msk in masks]
    
    x = np.arange(masks[0].shape[-1])
    y = np.arange(masks[0].shape[-2])
    distancefromcenter = np.zeros((len(masks[0]),*masks[0].shape[-2:]))
    for i,mask in enumerate(masks):
        y0,x0 = center_of_mass_2d(mask)
        distancefromcenter[i] = (y[:,None]-y0)**2 + (x[None,:]-x0)**2

    for i, j in overlaps:
        x1 = positions[i]
        x2 = positions[j]

        # Calculate relative shift in pixels
        dx = [int(x) for x in (x2 - x1) / pixel_size * 1e4][::-1]

        # Get the indices for array overlap
        i1, i2 = array_overlap(dx[0], pixels[0])
        j1, j2 = array_overlap(dx[1], pixels[1])

        # Make a mask where pixels closer to center of tile i are True
        closer = np.less_equal(
            distancefromcenter[i][i1[0] : i1[1], j1[0] : j1[1]],
            distancefromcenter[j][i2[0] : i2[1], j2[0] : j2[1]],
        )
        # Only consider pixels that are in the mask of tile j
        closer = np.where(clipped_masks[j][i2[0] : i2[1], j2[0] : j2[1]],closer,True)
        # Update the mask of tile i to only include pixels closer to its center
        clipped_masks[i][i1[0] : i1[1], j1[0] : j1[1]] = np.logical_and(closer,clipped_masks[i][i1[0] : i1[1], j1[0] : j1[1]])
        # fig,ax = plt.subplots(ncols=2,nrows=2)
        # ax[0,0].imshow(np.where(clipped_masks[i],distancefromcenter[i],0),cmap='gray')
        # ax[0,1].imshow(np.where(clipped_masks[j],distancefromcenter[j],0),cmap='gray')
        # ax[1,0].imshow(closer,cmap='gray')
        # ax[1,1].imshow(clipped_masks[i][i1[0] : i1[1], j1[0] : j1[1]],cmap='gray')
        # plt.show(block=True)
        closer = np.greater(
            distancefromcenter[i][i1[0] : i1[1], j1[0] : j1[1]],
            distancefromcenter[j][i2[0] : i2[1], j2[0] : j2[1]],
        )
        closer = np.where(clipped_masks[i][i1[0] : i1[1], j1[0] : j1[1]],closer,True)
        clipped_masks[j][i2[0] : i2[1], j2[0] : j2[1]] = np.logical_and(closer,clipped_masks[j][i2[0] : i2[1], j2[0] : j2[1]])

        # fig,ax = plt.subplots(ncols=2,nrows=2)
        # ax[0,0].imshow(np.where(clipped_masks[i],distancefromcenter[i],0),cmap='gray')
        # ax[0,1].imshow(np.where(clipped_masks[j],distancefromcenter[j],0),cmap='gray')
        # ax[1,0].imshow(closer,cmap='gray')
        # ax[1,1].imshow(clipped_masks[i][i1[0] : i1[1], j1[0] : j1[1]],cmap='gray')
        # plt.show(block=True)
        # aligned_image = stitch(
        #     ims,
        #     positions,
        #     clipped_masks,
        #     pixel_size,
        #     smooth=False
        # )

    return clipped_masks


def cross_correlate_tiles(
    positions,
    refpositions,
    tiles,
    reftiles,
    masks,
    refmasks,
    overlaps,
    references,
    pixel_size,
    max_correction=0.3,
    generate_plot=False,
    cross_corr_param_file=None,
):
    """
    Perform cross-correlation-based alignment of image tiles and adjust their positions.

    This function aligns overlapping image tiles by calculating the relative shifts between them
    using masked phase cross-correlation. The shifts are then used to adjust the global positions
    of the tiles through least squares minimization. The function supports visualizing the tile
    positions and shift vectors if desired.

    Args:
        positions (ndarray): Nx2 array of the initial x, y coordinates of the tiles in micrometers.
        tiles (list of ndarrays): List of 2D arrays representing the image tiles.
        masks (list of ndarrays): List of binary masks representing the valid regions of the tiles.
        overlaps (list of tuples): Pairs of indices representing which tiles overlap and should be aligned.
        pixel_size (float): The pixel size in microns.
        max_correction (float, optional): Maximum allowed correction (in microns) for shifts between tiles. Defaults to 0.1.
        generate_plot (bool or string, optional): Whether to generate a plot visualizing the tile positions and shift vectors. Defaults to False.
                                                  If a string this will be the filename that the plot will be saved as.

    Returns:
        ndarray: Updated Nx2 array of the adjusted x, y coordinates of the tiles.

    Notes:
        - The alignment is solved as a least squares problem (Ax = b) to minimize the relative shifts between overlapping tiles.
        - If some tiles are not connected to others through reliable shift determinations, their positions are adjusted
        using the initial relative positions inferred from the microscope.

    References:
        - Masked phase cross-correlation: https://scikit-image.org/docs/stable/auto_examples/registration/plot_masked_register_translation.html
        - Dirk Padfield, "Masked object registration in the Fourier domain", IEEE Transactions on Image Processing, 2011.
    """
    pixels = tiles[0].shape
    cmap = plt.get_cmap("viridis")

    # Convert maximum allowed shift in microns (max_correction) to pixels (max_shift)
    max_shift = max_correction / pixel_size * 1e4

    # We will solve global alignment of tiles by least squares Ax = b
    # matrix problem (https://en.wikipedia.org/wiki/Linear_least_squares).
    # N is the number of tiles times two (for x and y coordinates)
    N = 2 * len(positions)
    # Initialize lists which we will append rows for the A and b matrices
    A = []
    b = []

    genplot = False
    if type(generate_plot) is bool:
        genplot = generate_plot
        show = True
        savefig = False
    elif type(generate_plot) is str:
        genplot = True
        show = False
        savefig = True

    if genplot:
        xcorfig, xcorax = plt.subplots(figsize=(8, 8))
        xcorax.plot(*positions.T, "ko", label="Initial tile positions")
        for i, pos in enumerate(positions):
            xcorax.annotate(str(i), pos)

    G = nx.Graph()
    # Add nodes to graph
    G.add_nodes_from(list(range(len(positions))))

    xcorr = []
    deltas = []

    # TODO make this threaded
    if cross_corr_param_file is not None:
        xcorr, deltas = [
            load_array_from_hdf5(cross_corr_param_file, x)
            for x in ("cross_correlations", "relative_shifts")
        ]
    # plot_individual_cross_correlation(tiles,masks,positions,overlaps,pixel_size,filename_template='cross_corr_{0}_{1}.pdf')
    for ind, (i, j) in enumerate(tqdm(overlaps, desc="Cross-correlation alignment")):
        k = references[ind]
        # Retrieve image shifts for the overlapping tiles
        x1 = positions[i]
        x2 = positions[j]
        x3 = refpositions[k]

        # Calculate relative shift in pixels
        dx1 = [int(x) for x in (x3 - x1) / pixel_size * 1e4][::-1]
        dx2 = [int(x) for x in (x3 - x2) / pixel_size * 1e4][::-1]
        if cross_corr_param_file is None:

            # Perform masked phase cross-correlation to find the shift
            # between both images 1 and 2 and the reference image
            # The shift between images 1 and 2 is then the difference
            # between these two shifts
            detected_shifts = []
            # fig,ax = plt.subplots(ncols=2,nrows=2, figsize=(8,4))
            # ax= ax.ravel()
            m=0
            # fig,ax = plt.subplots(ncols=3,nrows=2)
            # ax = ax.ravel()
            for ii,dx in zip([i,j],[dx1, dx2]):
                # Align tiles by masked phase cross correlation, see:
                # https://scikit-image.org/docs/stable/auto_examples/registration/plot_masked_register_translation.html
                # and Padfield, Dirk. "Masked object registration in the Fourier domain." IEEE Transactions on image processing 21.5 (2011): 2706-2718.

                # Get the indices for array overlap
                i1, i2 = array_overlap(dx[0], pixels[0])
                j1, j2 = array_overlap(dx[1], pixels[1])

                # Make masks for (estimated) overlapping areas of each array
                reference_mask = np.zeros_like(masks[ii], dtype=bool)
                reference_mask[i1[0] : i1[1], j1[0] : j1[1]] = np.logical_and(masks[ii][
                    i1[0] : i1[1], j1[0] : j1[1]
                ],refmasks[k][
                    i2[0] : i2[1], j2[0] : j2[1]
                ])
                moving_mask = np.zeros_like(refmasks[k], dtype=bool)
                moving_mask[i2[0] : i2[1], j2[0] : j2[1]] = np.logical_and(masks[ii][
                    i1[0] : i1[1], j1[0] : j1[1]
                ],refmasks[k][
                    i2[0] : i2[1], j2[0] : j2[1]
                ])
                
                # Calculate shift by masked cross correlation with ordering (Y,X)
                shift,error,_ = phase_cross_correlation(
                    tiles[ii],
                    reftiles[k],
                    reference_mask=reference_mask,
                    moving_mask=moving_mask,
                    overlap_ratio=0.5,
                )
                detected_shifts.append(shift)
                
                # aligned_image = stitch(
                #     [tiles[ii], reftiles[k]],
                #     [positions[i], positions[i] + detected_shifts[-1][::-1] * pixel_size * 1e-4],
                #     [masks[ii], refmasks[k]],
                #     pixel_size,
                #     smooth=False,
                #     renormalize=True,
                # )
                # guess_alignment = stitch(
                #     [tiles[ii], reftiles[k]],
                #     [positions[i], positions[i] + np.asarray(dx)[::-1] * pixel_size * 1e-4],
                #     [masks[ii], refmasks[k]],
                #     pixel_size,
                #     smooth=False
                # )
                
                # ax[2+3*m].imshow(aligned_image,cmap='gist_gray')
                # ax[2+3*m].set_title(f'detected shift: {detected_shifts[-1][::-1]} px, Error {error:.4f}')
                # # # ax[1+3*m].imshow(guess_alignment,cmap='gist_gray')
                
                # # # Image.fromarray(aligned_image.astype(np.float32)).save(f'aligned_tile_{ii}_to_ref_{k}.tiff')

                # ax[0+3*m].imshow(tiles[ii],vmin = np.percentile(tiles[ii],5),vmax = np.percentile(tiles[ii],95),cmap='gist_gray')
                # ax[0+3*m].set_title(f'Tile {ii}')
                # ax[1+3*m].imshow(reftiles[k],vmin = np.percentile(reftiles[k],5),vmax = np.percentile(reftiles[k],95),cmap='gist_gray')
                # ax[0+3*m].imshow(reference_mask, alpha=0.1, cmap="Reds")
                # ax[1+3*m].imshow(moving_mask, alpha=0.1, cmap="Reds")
                m+=1
            # plt.show(block=True)
                
            # plt.show(block=True)


            detected_shift = np.asarray(detected_shifts[0]) - np.asarray(detected_shifts[1])
            
            # delta is measured shift in microns
            delta = np.asarray(detected_shift) * pixel_size * 1e-4

            
            # xcorr.append(xcorrmax)
            deltas.append(delta)
        else:
            delta = deltas[ind]
            # xcorrmax = xcorr[ind]
            detected_shift = delta / pixel_size * 1e-4

        # shifttoolarge=True
        if generate_plot:
            if ind == 0:
                label = "Shift vector from cross-correlation"
            else:
                label = None

        shifttoolarge = (
            np.linalg.norm((np.asarray(dx1)-np.asarray(dx2)) * pixel_size * 1e-4 - delta) > max_correction
        )
        if genplot:
            x1 = positions[i]
            # delta = (detected_shift * 1e4) * pixel_size
            if shifttoolarge:
                linestyle = "--"
            else:
                linestyle = "-"
            xcorax.plot(
                [x1[0], x1[0] + delta[1]],
                [x1[1], x1[1] + delta[0]],
                color=cmap(ind / len(overlaps)),
                label=label,
                linestyle=linestyle,
            )
        
        # detected_shift is position j (x2) - position i (x1)
        if not shifttoolarge:
            Arow = np.zeros((2, N))
            Arow[0, 2 * j] = 1
            Arow[0, 2 * i] = -1
            Arow[1, 2 * j + 1] = 1
            Arow[1, 2 * i + 1] = -1
            A.append( Arow)
            b[len(b) :] = ( delta).tolist()
            # Add valid connection to graph
            G.add_edge(i, j)

        # Weight by the maximum value of cross-correlation
        # weights.append(xcorrmax)

    # Join rows of A into numpy matrix
    if len(A) > 0:
        A = np.concatenate(A, axis=0)
    else:
        A = np.asarray(A)
    # Normalize by mean weight
    # meanweight = np.mean(weights)
    # A /= meanweight

    # Function to find the closest points using KDTree
    def find_closest_points_with_kdtree(list1, list2):
        # Create KDTree for the second list
        tree = KDTree(list2)

        # Query the tree with all points in the first list
        distances, indices = tree.query(list1)

        # Find the closest pair
        min_distance_index = distances.argmin()
        closest_point_list1 = list1[min_distance_index]
        closest_point_list2 = list2[indices[min_distance_index]]
        return min_distance_index, indices[min_distance_index]

    # For some tiles there will be no reliable shift determinations
    # from cross-correlation linking them to the other tiles,
    # in this case default to using the original
    # shifts implied by the microscope image shifts if groups of tiles
    # TODO use RANSAC algorithm to find the best fit
    print(nx.is_connected(G))
    if not nx.is_connected(G):
        # if False:
        extraA = []
        islands = sorted(nx.connected_components(G), key=len)
        largest_island = islands.pop()
        largest_island_points = [positions[x] for x in largest_island]
        # anchor = largest_island.pop()
        # for i in np.where(np.all(A == 0, axis=0))[0]:
        for island in islands:
            island_points = [positions[i] for i in island]

            i, j = find_closest_points_with_kdtree(island_points, largest_island_points)

            i = list(island)[i]
            j = list(largest_island)[j]
            x1 = positions[j]
            x2 = positions[i]
            delta = x1 - x2

            # Loop over x and y components
            for jj in range(2):
                # Make new row to add to A matrix
                Arow = np.zeros((N))
                Arow[2 * j + jj] = 1
                Arow[2 * i + jj] = -1
                extraA.append(Arow)

            b[len(b) :] = delta[::-1].tolist()
        A = np.stack(A.tolist() + extraA, axis=0)
    def objective_function(x,A,b,positions,lam):
        obj = np.linalg.norm(A @ x - b)
        obj += lam*np.linalg.norm(positions - x)
        return obj

    def objective_function(x, A, b, positions, lam):
        obj = np.linalg.norm(A @ x - b)
        obj += lam * np.linalg.norm(positions - x)
        return obj

    from scipy.optimize import minimize

    xvec = positions[:, ::-1].ravel()

    x, residuals, rank, s = np.linalg.lstsq(A, np.asarray(b), rcond=-1)
    newy = x[::2]
    newx = x[1::2]

    # The absolute position is unconstrained in least squares minimization
    # Pin the position closest to the origin to its original absolute position
    i = np.argmin(np.linalg.norm(positions, axis=1))
    newx -= newx[i] - positions[i][0]
    newy -= newy[i] - positions[i][1]

    newpositions = np.zeros_like(positions)
    newpositions[:, 0] = newx
    newpositions[:, 1] = newy

    

    if show:
        plt.show(block=True)
    if savefig:
        xcorfig.savefig(generate_plot)
    
    # fig,ax = plt.subplots()
    # ax.plot(*positions.T,'ro',linestyle='-',label='Montage tiles')
    # ax.plot(*newpositions.T,'bo',linestyle='--',label='Montage tiles')
    # for i,pos in enumerate(positions):
    #     ax.text(pos[0],pos[1],str(i))
    # plt.show(block=True)
    return newpositions, xcorr, deltas


def plot_individual_cross_correlation(
    images, masks, positions, overlaps, pixel_size, filename_template=None
):
    """
    Generate a plot showing the two masks and the final aligned images for each pair of overlapping tiles.

    Parameters:
    -----------
    images : list of numpy.ndarray
        List of image tiles.
    masks : list of numpy.ndarray
        List of binary masks corresponding to the image tiles.
    positions : numpy.ndarray
        Array of shape (N, 2) containing the (x, y) coordinates for positioning each image in units of microns.
    overlaps : list of tuples
        List of pairs of indices representing which tiles overlap and should be aligned.
    pixel_size : float
        The pixel size in microns.
    binning : int, optional
        Binning factor to reduce the image size. Default is 1.

    Returns:
    --------
    None
    """
    pixels = images[0].shape

    for i, j in overlaps:
        x1 = positions[i]
        x2 = positions[j]

        dx = [int(x) for x in (x2 - x1) / pixel_size * 1e4][::-1]
        print(dx)
        i1, i2 = array_overlap(dx[0], pixels[0])
        j1, j2 = array_overlap(dx[1], pixels[1])
        print(i1, i2, j1, j2)
        reference_mask = np.zeros_like(masks[i], dtype=bool)
        reference_mask[i1[0] : i1[1], j1[0] : j1[1]] = np.logical_and(masks[i][
            i1[0] : i1[1], j1[0] : j1[1]
        ],masks[j][
            i2[0] : i2[1], j2[0] : j2[1]
        ])
        moving_mask = np.zeros_like(masks[j], dtype=bool)
        moving_mask[i2[0] : i2[1], j2[0] : j2[1]] = np.logical_and(masks[i][
            i1[0] : i1[1], j1[0] : j1[1]
        ],masks[j][
            i2[0] : i2[1], j2[0] : j2[1]
        ])

        detected_shift = np.asarray(phase_cross_correlation(
            images[i], images[j], reference_mask=reference_mask, moving_mask=moving_mask
        )[0])

        fig = plt.figure(figsize=(8, 12))
        axes = fig.subplot_mosaic([["Image1", "Image2"], ["Stitched", "Stitched"]])
        gs = fig.add_gridspec(3, 1, height_ratios=[1, 1, 2], hspace=0.3)

        axes["Image1"].imshow(images[i], cmap="gray",vmin=np.percentile(images[i][masks[i]],1),vmax=np.percentile(images[i][masks[i]],99))
        axes["Image1"].imshow(reference_mask, alpha=0.5, cmap="Reds")
        axes["Image1"].set_title(f"Image {i} with Mask")

        axes["Image2"].imshow(images[j], cmap="gray",vmin=np.percentile(images[j][masks[j]],1),vmax=np.percentile(images[j][masks[j]],99))
        axes["Image2"].imshow(moving_mask, alpha=0.5, cmap="Reds")
        axes["Image2"].set_title(f"Image {j} with Mask")
        print(
            [positions[i], positions[i] + detected_shift * pixel_size * 1e-4],
            detected_shift,
        )
        aligned_image = stitch(
            [images[i], images[j]],
            [positions[i], positions[i] + detected_shift[::-1] * pixel_size * 1e-4],
            [masks[i], masks[j]],
            pixel_size,
        )
        # aligned_image = np.roll(images[j], shift=(-int(detected_shift[0]), -int(detected_shift[1])), axis=(0, 1))
        # axes[1, 0].imshow(images[i], cmap='gray')
        # axes[1, 0].set_title(f"Image {i}")

        axes["Stitched"].imshow(
            aligned_image,
            vmin=np.percentile(aligned_image, 10),
            vmax=np.percentile(aligned_image, 90),
            cmap="gray",
        )
        axes["Stitched"].set_title(f"Aligned Image {j}")

        for a in axes.values():
            a.axis("off")

        if filename_template is not None:
            fig.savefig(filename_template.format(i, j))
        else:
            plt.tight_layout()
            plt.show()


def parse_image_shifts(file_path):
    """Parse an image shift file and return image shifts in units of pixels"""
    with open(file_path, "r") as file:
        lines = file.readlines()
    ntilts = int(lines[0].strip())
    result = []
    index = 1
    # while index < len(lines):
    for _ in range(ntilts):
        # Parse the tilt angle
        tilt_angle = float(lines[index].strip())
        index += 1

        # Parse the number of rows
        n = int(lines[index].strip())
        index += 1

        # Parse the n x 3 array
        array = []
        for _ in range(n):
            array.append([float(x) for x in lines[index].strip().split()])
            index += 1
        array = np.asarray(array)
        # Add the parsed data to the result
        result.append((tilt_angle, array))

    # Sort the list by tilt angle and return
    return sorted(result, key=lambda x: x[0])


def generate_image_file_names_from_template(name, datadir, tilt, positions):
    imagefiles = [
        os.path.join(datadir, "{0}_{1}_{2}.mrc".format(name, tilt, x))
        for x in range(positions.shape[0])
    ]

    return imagefiles


def parse_mdoc(file_path):
    parsed_data = {}
    current_section = None

    # Regular expressions to match key-value pairs and sections
    key_value_regex = re.compile(r"(\S+)\s*=\s*(.+)")
    section_regex = re.compile(r"\[(.+)\]")

    with open(file_path, "r") as file:
        for line in file:
            line = line.strip()

            # Skip empty lines
            if not line:
                continue

            # Check if the line matches a section header in square brackets
            section_match = section_regex.match(line)
            if section_match:
                current_section = section_match.group(1)
                parsed_data[current_section] = {}
                continue

            # Match key-value pairs
            key_value_match = key_value_regex.match(line)
            if key_value_match:
                key, value = key_value_match.groups()
                if current_section:
                    parsed_data[current_section][key] = value
                else:
                    parsed_data[key] = value
    return parsed_data


def extract_tilt_axis_angle(file_path):
    tilt_axis_angle = None
    section_regex = re.compile(r"\[T\s*=\s*.*Tilt\s+axis\s+angle\s*=\s*([-\d.]+)")

    with open(file_path, "r") as file:
        for line in file:
            line = line.strip()

            # Check if the line contains the "Tilt axis angle"
            section_match = section_regex.search(line)
            if section_match:
                tilt_axis_angle = float(section_match.group(1))
                break

    return tilt_axis_angle


def setup_outputdir(args):
    if args["output"] is None:
        outputdir = os.path.split(args["input"])[1].replace("*.mrc", "_output")
    else:
        outputdir = args["output"]
    print("Results will be outputted to {0}".format(outputdir))
    if not os.path.exists(outputdir):
        os.mkdir(outputdir)
    return outputdir


def plot_positions(coordinates, fnam=None, fig=None, color="blue"):
    # Extract X and Y coordinates
    x_coords = [coord[0] for coord in coordinates]
    y_coords = [coord[1] for coord in coordinates]

    # Create a scatter plot
    if fig is None:
        fig, ax = plt.subplots(figsize=(10, 8))
    else:
        ax = fig.get_axes()[0]
        ax.title("Coordinate Points with Index Annotations")
        ax.xlabel("X Coordinates")
        ax.ylabel("Y Coordinates")
        ax.axhline(0, color="black", linewidth=0.5)
        ax.axvline(0, color="black", linewidth=0.5)
        ax.grid(color="gray", linestyle="--", linewidth=0.5)
    ax.scatter(x_coords, y_coords, color=color, label="Coordinates")

    # Annotate each point with its index
    for idx, (x, y) in enumerate(coordinates):
        ax.annotate(
            str(idx),
            (x, y),
            textcoords="offset points",
            xytext=(5, 5),
            ha="center",
            fontsize=8,
        )

    # Set plot title and labels

    ax.legend()

    # Save the plot to a PDF file
    fig.tight_layout()
    if fnam is None:
        plt.show()
    else:
        fig.savefig(fnam)


def plot_positions(coordinates,fnam=None,fig = None,color='blue'):


    # Extract X and Y coordinates
    x_coords = [coord[0] for coord in coordinates]
    y_coords = [coord[1] for coord in coordinates]

    # Create a scatter plot
    if fig is None:
        fig,ax = plt.subplots(figsize=(10, 8))
    else:
        ax = fig.get_axes()[0]
        ax.title('Coordinate Points with Index Annotations')
        ax.xlabel('X Coordinates')
        ax.ylabel('Y Coordinates')
        ax.axhline(0, color='black',linewidth=0.5)
        ax.axvline(0, color='black',linewidth=0.5)
        ax.grid(color = 'gray', linestyle = '--', linewidth = 0.5)
    ax.scatter(x_coords, y_coords, color=color, label='Coordinates')

    # Annotate each point with its index
    for idx, (x, y) in enumerate(coordinates):
        ax.annotate(str(idx), (x, y), textcoords="offset points", xytext=(5, 5), ha='center', fontsize=8)

    # Set plot title and labels

    
    ax.legend()

    # Save the plot to a PDF file
    fig.tight_layout()
    if fnam is None:
        plt.show()
    else:
        fig.savefig(fnam)


def main():
    args = parse_commandline()
    outdir = setup_outputdir(args)

    # Binning constant
    binning = int(args["binning"])

    files = glob.glob(args["input"])
    alignfile = args["input2"]
    if alignfile == 'None':
        match = re.search(r'_(-?[0-9]+(?:\.[0-9]+)?)\.mrc', files[0])
        print(match)
        if match:
            number = float(match.group(1))
        else:
            print('Could not determine alignment tilt from filename, please provide with --input2')
            sys.exit(1)
        
        nexttilt = np.sign(number)*(np.abs(number) +3)
        alignfile = files[0].replace(f'_{number}.mrc', f'_{nexttilt}.mrc')
        print(alignfile)
        if os.path.exists(alignfile):
            print(f'Using alignment tilt file: {alignfile}')
        else:
            prevtilt = np.sign(number)*(np.abs(number) -3)
            alignfile = files[0].replace(f'_{number}.mrc', f'_{prevtilt}.mrc')
            print(alignfile)
            if os.path.exists(alignfile):
                print(f'Using alignment tilt file: {alignfile}')
            else:
                print('Could not determine alignment tilt from filename, please provide with --input2')
                sys.exit(1)
        
    alignmdocfile = alignfile.replace(".mrc", ".mrc.mdoc")
    m = parse_mdoc(alignmdocfile)
    aligntilt = float(m["ZValue = 0"]["TiltAngle"])

    if len(files) < 1:
        raise FileNotFoundError("No files matching {0}".format(args["input"]))
    mdoc_files = [x.replace(".mrc", ".mrc.mdoc") for x in files]

    # Get pixel size, tilt axis rotation from first mdoc file
    mdoc = parse_mdoc(mdoc_files[0])
    pixelsize = float(mdoc["PixelSpacing"])

    # get tilts and image shifts from all mdoc files
    tilts, imageshifts = [[], []]
    indices = []
    # sys.exit()
    for mdoc in mdoc_files:
        m = parse_mdoc(mdoc)
        tilts.append(float(m["ZValue = 0"]["TiltAngle"]))

        # Split the string at '.mrc' and then find the integer before 'mrc'
        split_filename = mdoc.split(".mrc")[0]
        index = split_filename.split("_")[-1]
        indices.append(index)

    # Get image shifts from file
    # TODO make SerialEM put this in the mdoc file in microns (not
    # weird TFS units) to obviate this step
    imageshifts = parse_image_shifts(
        args["image_shifts"]
    )
    tiltsfromfile = [imageshifts[x][0] for x in range(len(imageshifts))]
    imageshifts = [imageshifts[x][1] for x in range(len(imageshifts))]

    indx = find_closest_index(tiltsfromfile, aligntilt)
    refpositions = imageshifts[indx]
    
    reshapedimshifts = np.concatenate(imageshifts)[:, :2] * pixelsize
    if args["maximageshift"] is not None:
        # Convert image shift criterion from microns to Angstroms
        maxim = args["maximageshift"] * 1e4
        mask = np.logical_and(*(np.abs(reshapedimshifts) < maxim).T)
    else:
        mask = np.ones(reshapedimshifts.shape[0], dtype=bool)
    # Global range of tiles in Angstroms
    globalwidth = np.ptp(reshapedimshifts[mask], axis=0)
    globalorigin = np.amin(reshapedimshifts[mask], axis=0)

    fringe_size = args["fringe_size"]

    for i, (file, tilt) in enumerate(
        tqdm(zip(files, tilts), total=len(files), desc="Stitching montages")
    ):
        # positions = imageshifts[i][1]
        indx = find_closest_index(tiltsfromfile, tilt)
        positions = imageshifts[indx]
        
        # plot_positions(positions[:,:2])
        if args['maximageshift'] is not None:
            tiles = np.where(np.logical_and(*(np.abs(positions) < maxim).T))[0]
        else:
            tiles = None
        # plot_positions(positions[tiles][:,:2],color='k')

        # plot_positions(positions[:,:2])
        if args["maximageshift"] is not None:
            tiles = np.where(np.logical_and(*(np.abs(positions) < maxim).T))[0]
        else:
            tiles = None
        # plot_positions(positions[tiles][:,:2],color='k')
        relative_tilt = aligntilt - tilt
        if np.abs(aligntilt) > np.abs(tilt):
            compression_factor = np.cos(np.deg2rad(relative_tilt))
        else:
            compression_factor = 1/np.cos(np.deg2rad(relative_tilt))
        print(compression_factor)
        mont = montage(
            file,
            alignfile,
            outdir,
            positions,
            refpositions,
            pixelsize,
            compression_factor,
            binning=binning,
            gainref=gainref,
            gainrefmask=gainrefmask,
            skipcrosscorrelation=args["skipcrosscorrelation"],
            montagewidth=globalwidth,
            montageorigin=globalorigin,
            tiles=tiles,
            fringe_size=fringe_size,
            # tiles = [24,25,36,37,38,39,54,55,56,57],
            gaincorrectedfile=args["gain_corrected_files"],
            maxshift=args["max_allowed_imshift_correction"],
            Matchgainrefmask=args["Matchgainrefmask"],
            maskthreshold=args["maskthreshold"],
            maskabsolutethreshold=args["maskabsolutethreshold"],
            flattenbeam=args["flattenbeam"],
            tilt_axis=extract_tilt_axis_angle(file.replace(".mrc", ".mrc.mdoc" )),
        )


if __name__ == "__main__":
    main()
```
