## Supplementary material for "Whole cryo-FIB lamella acquisition at high magnification with rectangular beam montage cryo-electron tomography": Source code: subvolume_segment.py

```
"""Split a montaged tomogram into overlapping XY sub-volumes, segment each, merge back.

Why this exists
---------------
easymode resamples a tomogram to the model's pixel size before inference. For the
10 A/px models (membrane, ribosome) and a 13.704 A/px montage that is an *upsample*,
so the working volume grows by 1.3704**3 = 2.6x. On a 1000x4500x4374 montage the
pipeline peaks around 780 GB, which is what killed job 28389962 at --mem=400G.

The peak scales with the XY area of whatever you hand easymode, so cutting the montage
into overlapping XY sub-volumes brings it down quadratically: a 3x3 split peaks near
120 GB. Z is never split -- the models need the full thickness, and Z is the cheap axis.

Note that pre-resampling the montage to 10 A/px does *not* help. Binning cannot reach a
smaller pixel size at all, and feeding easymode a volume already at 10 A/px makes the
tile loop worse (625 GB vs 502 GB), because the output accumulator is then allocated at
the larger size.

Sub-volumes overlap by --overlap pixels and are merged with a linear feather across the
overlap, so seams are blended rather than butt-joined. The overlap must comfortably
exceed the model's receptive field (160 px at 10 A/px = 219 px at 13.704 A/px); the
default of 256 does.

Usage
-----
    subvolume_segment.py plan  <tomogram.mrc> <workdir> [--grid 3 3] [--overlap 256]
    subvolume_segment.py split <workdir>                 # writes sub-volumes (IMOD trimvol)
    subvolume_segment.py merge <workdir> <feature>       # reassembles one feature

`plan` writes workdir/manifest.json, which the other two read, so the tiling arithmetic
lives in exactly one place.
"""

import argparse
import json
import os
import subprocess
import sys

import mrcfile
import numpy as np


def _manifest_path(workdir):
    return os.path.join(workdir, 'manifest.json')


def plan(tomogram, workdir, grid, overlap):
    with mrcfile.open(tomogram, permissive=True, header_only=True) as m:
        nz, ny, nx = int(m.header.nz), int(m.header.ny), int(m.header.nx)
        apix = float(m.voxel_size.x)

    ny_split, nx_split = grid
    tiles = []
    for iy in range(ny_split):
        for ix in range(nx_split):
            # Core (non-overlapping) extent this tile is responsible for, then grown by
            # `overlap` on every interior side. Growing only interior sides keeps the
            # union exactly equal to the original volume.
            y0c, y1c = iy * ny // ny_split, (iy + 1) * ny // ny_split
            x0c, x1c = ix * nx // nx_split, (ix + 1) * nx // nx_split
            y0, y1 = max(0, y0c - overlap), min(ny, y1c + overlap)
            x0, x1 = max(0, x0c - overlap), min(nx, x1c + overlap)
            tiles.append({
                'name': f'sub_y{iy}_x{ix}',
                # inclusive 0-based bounds of the extracted sub-volume
                'y0': y0, 'y1': y1 - 1, 'x0': x0, 'x1': x1 - 1,
                # the part of it this tile actually owns
                'y0_core': y0c, 'y1_core': y1c - 1, 'x0_core': x0c, 'x1_core': x1c - 1,
            })

    manifest = {
        'tomogram': os.path.abspath(tomogram),
        'shape': [nz, ny, nx],
        'apix': apix,
        'grid': [ny_split, nx_split],
        'overlap': overlap,
        'subvolume_dir': os.path.join(os.path.abspath(workdir), 'subvolumes'),
        'segmented_dir': os.path.join(os.path.abspath(workdir), 'segmented'),
        'tiles': tiles,
    }
    os.makedirs(workdir, exist_ok=True)
    with open(_manifest_path(workdir), 'w') as f:
        json.dump(manifest, f, indent=2)

    biggest = max((t['y1'] - t['y0'] + 1) * (t['x1'] - t['x0'] + 1) for t in tiles)
    frac = biggest / float(ny * nx)
    print(f'{tomogram}: {nz}x{ny}x{nx} at {apix:.3f} A/px')
    print(f'{len(tiles)} sub-volumes on a {ny_split}x{nx_split} grid, overlap {overlap} px')
    print(f'largest sub-volume is {frac:.1%} of the montage area')
    print(f'  -> estimated easymode peak ~{783 * frac:.0f} GB (vs ~783 GB for the whole montage)')
    print(f'wrote {_manifest_path(workdir)}')
    return manifest


def load(workdir):
    with open(_manifest_path(workdir)) as f:
        return json.load(f)


def split(workdir):
    man = load(workdir)
    os.makedirs(man['subvolume_dir'], exist_ok=True)
    nz = man['shape'][0]
    for t in man['tiles']:
        out = os.path.join(man['subvolume_dir'], t['name'] + '.mrc')
        if os.path.exists(out):
            print(f'{t["name"]}: exists, skipping')
            continue
        # trimvol takes 1-based inclusive ranges. Z is never trimmed.
        cmd = ['trimvol',
               '-x', f'{t["x0"] + 1},{t["x1"] + 1}',
               '-y', f'{t["y0"] + 1},{t["y1"] + 1}',
               '-z', f'1,{nz}',
               man['tomogram'], out]
        print(' '.join(cmd), flush=True)
        subprocess.run(cmd, check=True)
        # trimvol preserves the pixel size, but be explicit: easymode reads it from
        # the header when --apix is not passed, and a wrong value silently changes
        # the resampling factor.
        subprocess.run(['alterheader', '-del', f'{man["apix"]},{man["apix"]},{man["apix"]}',
                        out], check=True)
    print(f'{len(man["tiles"])} sub-volumes in {man["subvolume_dir"]}')


def _feather(n, lead, trail):
    """Ramp 0->1 over `lead` samples and 1->0 over the last `trail`, flat in between."""
    w = np.ones(n, dtype=np.float32)
    if lead > 0:
        w[:lead] = np.linspace(0.0, 1.0, lead + 2, dtype=np.float32)[1:-1]
    if trail > 0:
        w[n - trail:] = np.linspace(1.0, 0.0, trail + 2, dtype=np.float32)[1:-1]
    return w


def merge(workdir, feature, output=None):
    man = load(workdir)
    nz, ny, nx = man['shape']
    ny_split, nx_split = man['grid']
    seg_dir = man['segmented_dir']

    if output is None:
        stem = os.path.splitext(os.path.basename(man['tomogram']))[0]
        output = os.path.join(workdir, f'{stem}__{feature}.mrc')

    missing = []
    for t in man['tiles']:
        p = os.path.join(seg_dir, f'{t["name"]}__{feature}.mrc')
        # A 10x10x10 stub is easymode's placeholder for a run that never finished.
        if not os.path.exists(p) or os.path.getsize(p) < 10_000:
            missing.append(t['name'])
    if missing:
        sys.exit(f'error: {len(missing)} sub-volume(s) have no usable {feature} segmentation: '
                 + ', '.join(missing))

    # Accumulate in float32 one Z-chunk at a time rather than holding two full-size
    # float32 volumes: at 1000x4500x4374 that would be 158 GB for what is ultimately
    # a 20 GB int8 result.
    acc_chunk = 512
    with mrcfile.new_mmap(output, shape=(nz, ny, nx), mrc_mode=0, overwrite=True) as out:
        for z0 in range(0, nz, acc_chunk):
            z1 = min(nz, z0 + acc_chunk)
            num = np.zeros((z1 - z0, ny, nx), dtype=np.float32)
            den = np.zeros((z1 - z0, ny, nx), dtype=np.float32)
            for t in man['tiles']:
                p = os.path.join(seg_dir, f'{t["name"]}__{feature}.mrc')
                with mrcfile.mmap(p, permissive=True, mode='r') as m:
                    sub = np.asarray(m.data[z0:z1], dtype=np.float32)
                sy, sx = t['y1'] - t['y0'] + 1, t['x1'] - t['x0'] + 1
                if sub.shape[1:] != (sy, sx):
                    sys.exit(f'error: {os.path.basename(p)} is {sub.shape[1:]}, '
                             f'expected {(sy, sx)} -- manifest and segmentations disagree')
                wy = _feather(sy, t['y0_core'] - t['y0'], t['y1'] - t['y1_core'])
                wx = _feather(sx, t['x0_core'] - t['x0'], t['x1'] - t['x1_core'])
                w = wy[:, None] * wx[None, :]
                num[:, t['y0']:t['y1'] + 1, t['x0']:t['x1'] + 1] += sub * w
                den[:, t['y0']:t['y1'] + 1, t['x0']:t['x1'] + 1] += w
            np.divide(num, den, out=num, where=den > 0)
            np.rint(num, out=num)
            np.clip(num, -128, 127, out=num)
            out.data[z0:z1] = num.astype(np.int8)
            print(f'  z {z0}-{z1} of {nz}', flush=True)
        out.voxel_size = man['apix']
    print(f'wrote {output}')


def main():
    ap = argparse.ArgumentParser(description=__doc__,
                                 formatter_class=argparse.RawDescriptionHelpFormatter)
    sub = ap.add_subparsers(dest='command', required=True)

    p = sub.add_parser('plan', help='compute the tiling and write manifest.json')
    p.add_argument('tomogram')
    p.add_argument('workdir')
    p.add_argument('--grid', nargs=2, type=int, default=[3, 3], metavar=('NY', 'NX'))
    p.add_argument('--overlap', type=int, default=256,
                   help='overlap in original px (default 256; must exceed the 219 px '
                        'receptive field of a 160 px 10 A/px tile at 13.704 A/px)')

    p = sub.add_parser('split', help='cut the sub-volumes with IMOD trimvol')
    p.add_argument('workdir')

    p = sub.add_parser('merge', help='feather-blend the segmented sub-volumes back together')
    p.add_argument('workdir')
    p.add_argument('feature')
    p.add_argument('--output', default=None)

    args = ap.parse_args()
    if args.command == 'plan':
        plan(args.tomogram, args.workdir, args.grid, args.overlap)
    elif args.command == 'split':
        split(args.workdir)
    elif args.command == 'merge':
        merge(args.workdir, args.feature, args.output)


if __name__ == '__main__':
    main()
```
