## Supplementary material for "Whole cryo-FIB lamella acquisition at high magnification with rectangular beam montage cryo-electron tomography": Source code: Utilities.py

```
import h5py
import numpy as np
import math
import warnings
from typing import List, Tuple
from scipy.ndimage import correlate, sobel, gaussian_filter, distance_transform_edt
from scipy.ndimage.morphology import binary_fill_holes, binary_erosion, binary_dilation
from tqdm import tqdm
import copy
import mrcfile
import matplotlib.pyplot as plt
import os
import png
import re
from typing import List, Tuple, Dict, Any

def renormalize(array, oldmin=None, oldmax=None, newmax=1.0, newmin=0.0):
    """Rescales the array such that its maximum is newmax and its minimum is newmin."""
    if oldmin is not None:
        min_ = oldmin
    else:
        min_ = array.min()

    if oldmax is not None:
        max_ = oldmax
    else:
        max_ = array.max()

    return (
        np.clip((array - min_) / (max_ - min_), 0.0, 1.0) * (newmax - newmin) + newmin
    )


def parse_mdoc(file_path):
    parsed_data = {}
    current_section = None

    # Regular expressions to match key-value pairs and sections
    key_value_regex = re.compile(r"(\S+)\s*=\s*(.+)")
    section_regex = re.compile(r"\[(.+)\]")

    with open(file_path, "r") as file:
        for line in file:
            line = line.strip()

            # Skip empty lines
            if not line:
                continue

            # Check if the line matches a section header in square brackets
            section_match = section_regex.match(line)
            if section_match:
                current_section = section_match.group(1)
                parsed_data[current_section] = {}
                continue

            # Match key-value pairs
            key_value_match = key_value_regex.match(line)
            if key_value_match:
                key, value = key_value_match.groups()
                if current_section:
                    parsed_data[current_section][key] = value
                else:
                    parsed_data[key] = value
    return parsed_data


def write_mdoc(parsed_data, file_path):
    """
    Write a dictionary (like that produced by parse_mdoc) back to an mdoc file.

    Parameters:
    parsed_data (dict): The dictionary to write.
    file_path (str): The path to the output mdoc file.
    """
    with open(file_path, 'w') as f:
        for key, value in parsed_data.items():
            if isinstance(value, dict):
                f.write(f'[{key}]\n')
                for subkey, subval in value.items():
                    f.write(f'{subkey} = {subval}\n')
            else:
                f.write(f'{key} = {value}\n')


def combine_dicts(dict_list: List[Dict]) -> Dict:
    """
    Combines a set of dictionaries, only keeping key-value pairs that are identical across all dictionaries.

    Parameters:
    -----------
    dict_list : list of dict
        A list of dictionaries to combine.

    Returns:
    --------
    dict
        A dictionary containing only the key-value pairs that are identical in all input dictionaries.
    """
    if not dict_list:
        return {}
    
    if len(dict_list) == 1:
        return dict_list[0].copy()
    
    # Start with the first dictionary
    common_dict = {}
    
    # Get all keys from the first dictionary
    for key, value in dict_list[0].items():
        # Check if this key-value pair is identical in all other dictionaries
        if all(d.get(key) == value for d in dict_list[1:]):
            common_dict[key] = value
    
    return common_dict


def array_to_RGB(arrayin, cmap=plt.get_cmap("viridis"), vmin=None, vmax=None):
    """Convert an array to RGB using a supplied colormap."""

    kwargs = {"oldmin": vmin, "oldmax": vmax}
    return (cmap(renormalize(arrayin, **kwargs))[..., :3] * 256).astype(np.uint8)


def RGB_to_PNG(RGB_array, fnam):
    """Output an RGB array [shape (n,m,3)] as a .png file."""
    # Get array shape
    n, m = RGB_array.shape[:2]

    # Replace filename ending with .png
    fnam_out = os.path.splitext(fnam)[0] + ".png"
    png.fromarray(RGB_array.reshape((n, m * 3)), mode="RGB").save(fnam_out)


def save_array_as_png(array, fnam, cmap=plt.get_cmap("viridis"), vmin=None, vmax=None):
    """Output a numpy array as a .png file."""
    if vmin is None or vmax is None:
        populated = array[array != 0]
        data = populated if populated.size > 0 else array.ravel()
        if vmin is None:
            vmin = np.percentile(data, 2)
        if vmax is None:
            vmax = np.percentile(data, 98)
    RGB_to_PNG(array_to_RGB(array, cmap, vmin=vmin, vmax=vmax), fnam)

def savetomrc(array, fnam, overwrite=True):
    """
    Saves a NumPy array to an MRC file format.

    Parameters
    ----------
    array : numpy.ndarray
        The data array to be saved in MRC format. This is typically a 2D or 3D array.

    fnam : str
        The filename for the output MRC file, including the path if necessary.

    overwrite : bool, optional
        If True (default), overwrites the file if it already exists. If False, raises
        an error if the file exists.

    Returns
    -------
    None
        This function writes the data to disk and does not return a value.

    Notes
    -----
    The MRC (Medical Research Council) file format is commonly used for storing electron microscopy and
    other volumetric data. This function uses the `mrcfile` library to write the NumPy array as MRC data.
    """
    with mrcfile.new(fnam, overwrite=overwrite) as mrc:
        mrc.set_data(array)


def find_closest_index(arr, target):
    """
    Finds the index of the closest value in the array to the target value.

    Parameters:
    arr (list or np.array): A list or numpy array of numbers
    target (float or int): The target value to find the closest match for

    Returns:
    int: The index of the closest value in the array
    """
    # Convert the input array to a numpy array if it's not already
    arr = np.array(arr)

    # Compute the absolute difference between each element and the target
    diff = np.abs(arr - target)

    # Find the index of the minimum difference
    closest_index = np.argmin(diff)

    return closest_index


def correlate(array1, array2, axes=None):
    """
    Fourier correlation of two arrays over specified axes.

    array2 is broadcast to match array1 so axes refers to the dimensions of
    array1
    """
    # input and output shape
    s = array1.shape
    # Broadcast array2 to match array1
    a2 = np.broadcast_to(array2, s)
    # Axes of transformation
    a = axes
    if a is not None:
        s = [s[i] for i in a]
    if np.iscomplexobj(array1) or np.iscomplexobj(a2):
        return np.fft.ifftn(
            np.fft.fftn(array1, s, a) * np.conj(np.fft.fftn(a2, s, a)), s, a
        )
    else:
        return np.fft.irfftn(
            np.fft.rfftn(array1, s, a) * np.conj(np.fft.rfftn(a2, s, a)), s, a
        )


def save_array_to_hdf5(
    arrays: List[np.ndarray],
    filename: str,
    dataset_names: List[str],
    filemode: str = "w",
    compression ="gzip",
    compression_opts=9
) -> None:
    """
    Saves a numpy array to an HDF5 file.

    Parameters:
    arrays (List[np.ndarray]): The numpy arrays to save.
    filename (str): The name of the file where the arrays will be saved.
    dataset_names (List[str]): The names of the datasets in the HDF5 file.
    filemode (str): The file mode for opening the HDF5 file.
    """
    with h5py.File(filename, filemode) as hdf:
        for name, data in zip(dataset_names, arrays):
            if isinstance(data, str):
                hdf.create_dataset(name, data=data)
            elif np.isscalar(data):
                hdf.create_dataset(name, data=data)
            else:
                hdf.create_dataset(name, data=data, compression=compression, compression_opts=compression_opts)


def load_array_from_hdf5(filename: str, dataset_name: str):
    """
    Loads a dataset from an HDF5 file.

    Parameters:
    filename (str): The name of the file from which to load the dataset.
    dataset_name (str): The name of the dataset in the HDF5 file.

    Returns:
    The dataset as a numpy array, string, or scalar depending on what was stored.
    """
    with h5py.File(filename, "r") as hdf:
        dataset = hdf[dataset_name]
        if h5py.check_string_dtype(dataset.dtype):
            value = dataset[()]
            return value.decode("utf-8") if isinstance(value, bytes) else value
        value = dataset[()]
        if value.ndim == 0:
            return value.item()
        return np.array(value)


def calc_row_idx(k: int, n: int) -> int:
    """
    Calculate the row index in a condensed distance matrix.

    Parameters:
    k (int): Index in the condensed distance matrix.
    n (int): Number of original data points.

    Returns:
    int: The row index corresponding to the given condensed index.
    """
    return int(
        math.ceil(
            (1 / 2.0) * (-((-8 * k + 4 * n**2 - 4 * n - 7) ** 0.5) + 2 * n - 1) - 1
        )
    )


def elem_in_i_rows(i: int, n: int) -> int:
    """
    Calculate the number of elements in the first i rows of a condensed distance matrix.

    Parameters:
    i (int): Row index.
    n (int): Number of original data points.

    Returns:
    int: Number of elements in the first i rows.
    """
    return i * (n - 1 - i) + (i * (i + 1)) // 2


def calc_col_idx(k: int, i: int, n: int) -> int:
    """
    Calculate the column index in a condensed distance matrix.

    Parameters:
    k (int): Index in the condensed distance matrix.
    i (int): Row index in the square matrix.
    n (int): Number of original data points.

    Returns:
    int: The column index corresponding to the given condensed index and row index.
    """
    return int(n - elem_in_i_rows(i + 1, n) + k)


def condensed_to_square(k: int, n: int) -> Tuple[int, int]:
    """
    Convert a condensed distance matrix index to a square matrix index.

    Parameters:
    k (int): Index in the condensed distance matrix.
    n (int): Number of original data points.

    Returns:
    tuple: The row and column indices in the square matrix.
    """
    i = calc_row_idx(k, n)
    j = calc_col_idx(k, i, n)
    return i, j


def crop(arrayin: np.ndarray, shapeout: Tuple[int, ...]) -> np.ndarray:
    """
    Crop or zero-pad the last `n` dimensions of `arrayin` to match the specified `shapeout`.

    If the dimensions in `shapeout` are smaller than those of `arrayin`, the function crops
    the array to match the specified size. If the dimensions in `shapeout` are larger, the
    function zero-pads the array to achieve the desired output shape.

    Parameters
    ----------
    arrayin : ndarray
        Input array of shape (..., N1, N2, ..., Nn). The array to be cropped or zero-padded
        along its last `n` dimensions.
    shapeout : tuple of int
        Desired shape for the final `n` dimensions of `arrayin`. The function preserves the
        leading dimensions (those not specified in `shapeout`) and modifies only the trailing
        dimensions.

    Returns
    -------
    arrayout : ndarray
        Output array with shape (..., N1', N2', ..., Nn'), where N1', N2', ..., Nn' match
        `shapeout`. Cropping or zero-padding is applied as necessary.

    Notes
    -----
    - For each dimension, if the desired size is smaller than the input size, the array is
      centered and cropped accordingly.
    - If the desired size is larger than the input size, the array is centered and zero-padded.

    Example
    -------
    >>> arrayin = np.array([[1, 2, 3], [4, 5, 6]])
    >>> shapeout = (4, 2)
    >>> crop(arrayin, shapeout)
    array([[0, 1],
           [4, 5],
           [0, 0],
           [0, 0]])
    """
    # Total number of dimensions in the input array
    ndim = arrayin.ndim

    # Number of trailing dimensions to crop or pad
    n = len(shapeout)

    # Number of leading dimensions not affected by cropping/padding
    nUntouched = ndim - n

    # Calculate the shape of the output array
    shapeout_ = arrayin.shape[:nUntouched] + tuple(shapeout)

    # Initialize the output array with zeros
    arrayout = np.zeros(shapeout_, dtype=arrayin.dtype)

    # Get the shapes of the trailing dimensions for input and output arrays
    oldshape = arrayin.shape[-n:]
    newshape = shapeout[-n:]

    def indices(y: int, y_: int) -> Tuple[slice, slice]:
        """
        Determine the slicing indices for cropping or zero-padding a single dimension.

        Parameters
        ----------
        y : int
            Size of the dimension in the input array.
        y_ : int
            Desired size of the dimension in the output array.

        Returns
        -------
        in_slice : slice
            Slicing indices for the input array.
        out_slice : slice
            Slicing indices for the output array.
        """
        if y > y_:
            # Crop: Center the cropping region within the input dimension
            y1, y2 = (y - y_) // 2, (y + y_) // 2
            in_slice = slice(y1, y2)
            out_slice = slice(0, y_)
        else:
            # Zero-pad: Center the input dimension within the padded output region
            y1_, y2_ = (y_ - y) // 2, (y_ + y) // 2
            in_slice = slice(0, y)
            out_slice = slice(y1_, y2_)
        return in_slice, out_slice

    # Compute the slicing indices for each trailing dimension
    ind = [indices(x, x_) for x, x_ in zip(oldshape, newshape)]
    inind, outind = map(tuple, zip(*ind))

    # Assign the cropped or padded data to the output array
    arrayout[nUntouched * (slice(None),) + outind] = arrayin[
        nUntouched * (slice(None),) + inind
    ]

    return arrayout


def rotation_matrix(theta: float) -> np.ndarray:
    """
    Returns a 2D rotation matrix for a given angle in degrees

    Parameters:
    theta, float :: The rotation angle in degrees

    returns:
    numpy.ndarray: the 2x2 rotation matrix."""

    rad = np.deg2rad(theta)
    ct = np.cos(rad)
    st = np.sin(rad)

    return np.array([[ct, -st], [st, ct]])


def fourier_crop(ain: np.ndarray, shapeout: Tuple[int, ...]) -> np.ndarray:
    """
    Crop or pad a Fourier-transformed array to match the desired output shape.

    Parameters:
    -----------
    ain : numpy.ndarray
        Input array to be cropped or padded in Fourier space.
    shapeout : tuple
        Desired shape of the output array.

    Returns:
    --------
    numpy.ndarray : Cropped or padded array in Fourier space.
    """

    def crop1d(array: np.ndarray, s: int, d: int) -> np.ndarray:
        # Number of dimensions of array
        N = len(array.shape)
        # Size of array that will be transferred to new grid
        s_ = min(array.shape[d], s)
        # Indices of grid region to transfer to new grid
        ind1 = (
            (np.s_[:],) * ((N + d) % N)
            + (np.s_[: s_ // 2 + s_ % 2],)
            + (np.s_[:],) * (N - (N + d) % N - 1)
        )
        ind2 = (
            (np.s_[:],) * ((N + d) % N)
            + (np.s_[-s_ // 2 + s_ % 2 :],)
            + (np.s_[:],) * (N - (N + d) % N - 1)
        )
        if s > array.shape[d]:
            xtra = list(array.shape)
            xtra[d] = s - array.shape[d]
            return np.concatenate(
                [array[ind1], np.zeros(xtra, dtype=array.dtype), array[ind2]], axis=d
            )
        else:
            return np.concatenate([array[ind1], array[ind2]], axis=d)

    array = copy.deepcopy(ain)
    for i, s in enumerate(shapeout):
        array = crop1d(array, s, i - len(shapeout))
    return array


def fourier_interpolate(
    ain: np.ndarray,
    shapeout: Tuple[int, ...],
    norm: str = "conserve_val",
    N: int = None,
    qspace_in: bool = False,
    qspace_out: bool = False,
) -> np.ndarray:
    """
    Perfom fourier interpolation on array ain so that its shape matches shapeout.

    Arguments
    ---------
    ain : (...,Ni,..,Ny,Nx) array_like
        Input numpy array, interpolation will be applied to the n trailing
        dimensions where n is the length of shapeout.
    shapeout : int (n,) , array_like
        Desired shape of output array
    norm : str, optional  {'conserve_val','conserve_norm','conserve_L1'}
        Normalization of output. If 'conserve_val' then array values are preserved
        if 'conserve_norm' L2 norm is conserved under interpolation and if
        'conserve_L1' L1 norm is conserved under interpolation
    N : int, optional
        Number of (trailing) dimensions to Fourier interpolate. By default take
        the length of shapeout
    qspace_in : bool, optional
        Set to True if the input array is in reciprocal space, False if not (default).
    ---------
    ain : (...,Ni,..,Ny,Nx) array_like
        Input numpy array, interpolation will be applied to the n trailing
        dimensions where n is the length of shapeout.
    shapeout : int (n,) , array_like
        Desired shape of output array
    norm : str, optional  {'conserve_val','conserve_norm','conserve_L1'}
        Normalization of output. If 'conserve_val' then array values are preserved
        if 'conserve_norm' L2 norm is conserved under interpolation and if
        'conserve_L1' L1 norm is conserved under interpolation
    N : int, optional
        Number of (trailing) dimensions to Fourier interpolate. By default take
        the length of shapeout
    qspace_in : bool, optional
        Set to True if the input array is in reciprocal space, False if not (default).
        Be careful with setting this to True for a non-complex array.
    qspace_out : bool, optional
        Set to True for reciprocal space output, False for real-space output (default).
    """
    # Import required FFT functions
    from numpy.fft import fftn

    if N is None:
        N = len(shapeout)

    # Make input complex
    aout = np.zeros(np.shape(ain)[:-N] + tuple(shapeout), dtype=complex)

    # Get input dimensions
    shapein = np.shape(ain)[-N:]

    # axes to Fourier transform
    axes = np.arange(-N, 0)

    if qspace_in:
        a = np.asarray(ain, dtype=complex)
    else:
        a = fftn(np.asarray(ain, dtype=complex), axes=axes)

    aout = fourier_crop(a, shapeout)
    # aout = np.fft.fftshift(crop(np.fft.fftshift(a,axes=axes),shapeout),axes=axes)

    # Fourier transform result with appropriate normalization
    if norm == "conserve_val":
        aout *= np.prod(shapeout) / np.prod(np.shape(ain)[-N:])
    elif norm == "conserve_norm":
        aout *= np.sqrt(np.prod(shapeout) / np.prod(np.shape(ain)[-N:]))

    if not qspace_out:
        aout = np.fft.ifftn(aout, axes=axes)

    # Return correct array data type
    if not np.iscomplexobj(ain):
        return np.real(aout)
    else:
        return aout


def get_imageshifts_for_tilt_angle(file_path: str, tilt: float) -> np.ndarray:
    """
    Retrieve image shifts for a specific tilt angle from an mdoc file.

    Parameters:
    -----------
    file_path : str
        Path to the file containing image shifts.
    tilt : float
        The tilt angle for which to retrieve the image shifts.

    Returns:
    --------
    np.ndarray
        A 2D numpy array containing the image shifts for the specified tilt angle.
    """
    with open(file_path, "r") as file:
        lines = file.readlines()

    result = []
    index = 0
    while index < len(lines):
        # Parse the tilt angle
        tilt_angle = float(lines[index].strip())
        index += 1

        # Parse the number of rows
        n = int(lines[index].strip())
        index += 1

        # Parse the n x 3 array
        array = []
        for _ in range(n):
            array.append([float(x) for x in lines[index].strip().split()])
            index += 1

        # Add the parsed data to the result
        if float(tilt) == tilt_angle:
            return np.asarray(array)[:, :2]


def convolve(array1, array2, axes=None):
    """
    Fourier convolution of two arrays over specified axes.

    array2 is broadcast to match array1 so axes refers to the dimensions of
    array1
    """
    # input and output shape
    s = array1.shape
    # Broadcast array2 to match array1
    a2 = np.broadcast_to(array2, s)
    # Axes of transformation
    a = axes
    if a is not None:
        s = [s[i] for i in a]
    if np.iscomplexobj(array1) or np.iscomplexobj(a2):
        return np.fft.ifftn(np.fft.fftn(array1, s, a) * np.fft.fftn(a2, s, a), s, a)
    else:
        return np.fft.irfftn(np.fft.rfftn(array1, s, a) * np.fft.rfftn(a2, s, a), s, a)


def Gaussian(sigma, gridshape):
    r"""
    Calculate a 2D Gaussian function.

    Notes
    -----
    Functional form
    .. math:: 1 / \sqrt { 2 \pi \sigma }  e^{ - ( x^2 + y^2 ) / 2 / \sigma^2 }

    Parameters
    ----------
    sigma : float or (2,) array_like
        The standard deviation of the Gaussian function in pixels
    gridshape : (2,) array_like
        Number of pixels in the grid.
    """
    ysqr = np.fft.fftfreq(gridshape[0], d=1 / gridshape[0]) ** 2
    xsqr = np.fft.fftfreq(gridshape[1], d=1 / gridshape[1]) ** 2

    gaussian = np.exp(-(ysqr[:, None] + xsqr[None, :]) / sigma**2 / 2)
    return gaussian / np.sum(gaussian)

def center_of_mass_2d(array):
    """
    Calculate the center of mass of a 2D array.

    Parameters
    ----------
    array : 2D numpy array
        Input array for which to calculate the center of mass.

    Returns
    -------
    tuple
        Coordinates of the center of mass (row, col).
    """
    total = np.sum(array)
    if total == 0:
        return array.shape[0] / 2, array.shape[1] / 2
        # raise ValueError("The sum of the array elements is zero, cannot determine center of mass.")
    
    rows, cols = np.indices(array.shape)
    center_row = np.sum(rows * array) / total
    center_col = np.sum(cols * array) / total
    
    return center_row, center_col


def broadcast_from_unmeshed(coords):
    """
    For an unmeshed set of coordinates broadcast to a meshed ND array.

    Examples
    --------
    >>> broadcast_from_unmeshed([np.arange(5),np.arange(6)])
    [array([[0, 0, 0, 0, 0, 0],
       [1, 1, 1, 1, 1, 1],
       [2, 2, 2, 2, 2, 2],
       [3, 3, 3, 3, 3, 3],
       [4, 4, 4, 4, 4, 4]]), array([[0, 1, 2, 3, 4, 5],
       [0, 1, 2, 3, 4, 5],
       [0, 1, 2, 3, 4, 5],
       [0, 1, 2, 3, 4, 5],
       [0, 1, 2, 3, 4, 5]])]
    """

    N = len(coords)
    pixels = [a.shape[0] for a in coords]

    # Broadcasting patterns
    R = np.ones((N, N), dtype=np.int16) + np.diag(pixels) - np.eye(N, dtype=np.int16)

    # Broadcast unmeshed grids
    return [np.broadcast_to(a.reshape(rr), pixels) for a, rr in zip(coords, R)]


def r_space_array(pixels, gridsize, meshed=True):
    """
    Return the appropriately scaled ND real space coordinates.

    Parameters
    -----------
    pixels : (N,) array_like
        Pixels in each dimension of a ND array
    gridsize : (N,) array_like
        Dimensions of the array in real space units
    meshed : bool, optional
        Option to output dense meshed grid (True) or output unbroadcasted
        arrays (False)
    """
    # N is the dimensionality of grid
    N = len(pixels)

    # Calculate unmeshed grids
    rspace = [np.fft.fftshift(np.fft.fftfreq(pixels[i], d=1 / gridsize[i])) for i in range(N)]

    # At this point we can return the arrays without broadcasting
    if meshed:
        return broadcast_from_unmeshed(rspace)
    else:
        return rspace


def plot_plasmon_sweep(tile, ref_beam, mask, pixel_size_nm,
                       E_plasmon_eV=21.0, voltage_kV=300.0,
                       n_values=(1, 5, 10, 15, 20)):
    """
    Diagnostic plot: reference beam blurred by the plasmon kernel at a range of
    n values alongside the actual tile.  All images are normalised to the tile's
    mean within the mask so only profile *shape* differences are visible.
    Use this to judge visually whether any n gives a good match to the tile and
    whether q_E (the Lorentzian half-width) is physically reasonable.

    Parameters
    ----------
    tile : (ny, nx) ndarray
    ref_beam : (ny, nx) ndarray
    mask : (ny, nx) bool ndarray
    pixel_size_nm : float
    E_plasmon_eV : float
    voltage_kV : float
    n_values : sequence of float
        n values to sweep (default 1, 5, 10, 15, 20).
    """
    m0c2 = 511e3
    beta2 = 1.0 - (m0c2 / (m0c2 + voltage_kV * 1e3)) ** 2
    q_E = E_plasmon_eV / (1239.8 * np.sqrt(beta2))

    ny, nx = ref_beam.shape
    pad = max(16, int(np.ceil(3.0 / (q_E * pixel_size_nm))))
    ref_padded = np.pad(ref_beam.astype(float), pad, mode='constant', constant_values=0.0)
    ny_p, nx_p = ref_padded.shape
    qy_p = np.fft.fftfreq(ny_p, d=pixel_size_nm)
    qx_p = np.fft.fftfreq(nx_p, d=pixel_size_nm)
    P_single_p = q_E ** 2 / (qy_p[:, None] ** 2 + qx_p[None, :] ** 2 + q_E ** 2)
    ref_fft_p = np.fft.fft2(ref_padded)

    n_list = [0] + list(n_values)
    blurred = []
    for n in n_list:
        K_p = np.exp(n * (P_single_p - 1.0))
        blurred.append(np.real(np.fft.ifft2(ref_fft_p * K_p))[pad:pad + ny, pad:pad + nx])

    # Normalise every image to tile's mean within mask so shape is comparable
    tile_mean = float(np.mean(tile[mask])) if mask.any() else float(np.mean(tile))
    def _norm(img):
        m = float(np.mean(img[mask])) if mask.any() else float(np.mean(img))
        return img * (tile_mean / m) if m != 0 else img

    panels      = [_norm(b) for b in blurred] + [tile.astype(float)]
    panel_titles = [f"ref  n={n}" for n in n_list] + ["tile"]

    n_panels = len(panels)
    half = 50
    row0, row1 = max(ny // 2 - half, 0), min(ny // 2 + half, ny)

    fig = plt.figure(figsize=(3.0 * n_panels, 8))
    gs  = fig.add_gridspec(2, n_panels, height_ratios=[3, 1], hspace=0.35, wspace=0.05)
    img_axes = [fig.add_subplot(gs[0, i]) for i in range(n_panels)]
    ax_ls    = fig.add_subplot(gs[1, :])   # linescan spans all columns

    vmin, vmax = (np.percentile(tile[mask], [1, 99]) if mask.any()
                  else (tile.min(), tile.max()))
    for ax, img, title in zip(img_axes, panels, panel_titles):
        ax.imshow(img, cmap="gray", vmin=vmin, vmax=vmax, origin="lower")
        ax.axhline(row0, color="r", linewidth=0.5, linestyle="--")
        ax.axhline(row1, color="r", linewidth=0.5, linestyle="--")
        ax.set_title(title, fontsize=9)
        ax.axis("off")

    colors = plt.cm.viridis(np.linspace(0.0, 0.85, len(n_list)))
    for img, n, color in zip(panels[:-1], n_list, colors):
        ls = img[row0:row1, :].mean(axis=0)
        ax_ls.plot(ls, color=color, linewidth=1.0, label=f"n={n}")
    ax_ls.plot(tile[row0:row1, :].mean(axis=0), color="r", linewidth=1.5, label="tile")
    ax_ls.set_xlim(0, nx - 1)
    ax_ls.set_xlabel("x (px)")
    ax_ls.set_ylabel("intensity (normalised)")
    ax_ls.legend(fontsize=8, ncol=len(n_list) + 1, loc="upper center",
                 bbox_to_anchor=(0.5, 1.18))

    fig.suptitle(
        f"Plasmon sweep  |  q_E = {q_E:.4f} cyc/nm  "
        f"|  FWHM ≈ {0.5 / q_E / pixel_size_nm:.1f} px  "
        f"|  pixel = {pixel_size_nm:.3f} nm",
        fontsize=10,
    )
    plt.show()


def _estimate_n_avg_lsq(tile, ref_beam, mask, q_E, pixel_size_nm, downsample=64):
    """
    Estimate n_avg by least-squares shape-matching on a downsampled image.

    Finds the n_avg that minimises ||tile - a * I_exp(n_avg)||² within the
    beam mask, where a is an analytically-optimal amplitude scale factor.
    Fitting only the profile *shape* (not its amplitude) makes this robust to
    overall intensity differences between reference and tile (e.g. from dose,
    energy filtering, or thickness).
    """
    from scipy.optimize import minimize_scalar

    ny, nx = tile.shape
    ds_y = min(downsample, ny)
    ds_x = min(downsample, nx)

    tile_ds = fourier_interpolate(tile.astype(float), [ds_y, ds_x])
    ref_ds  = fourier_interpolate(ref_beam.astype(float), [ds_y, ds_x])
    mask_ds = fourier_interpolate(mask.astype(float), [ds_y, ds_x]) > 0.5
    if not mask_ds.any():
        mask_ds = np.ones((ds_y, ds_x), dtype=bool)

    # Pixel size scales with downsampling factor
    py = pixel_size_nm * ny / ds_y
    px = pixel_size_nm * nx / ds_x
    qy = np.fft.fftfreq(ds_y, d=py)
    qx = np.fft.fftfreq(ds_x, d=px)
    P_single_ds = q_E ** 2 / (qy[:, None] ** 2 + qx[None, :] ** 2 + q_E ** 2)
    ref_fft_ds  = np.fft.fft2(ref_ds)

    def residual(n):
        K = np.exp(n * (P_single_ds - 1.0))
        I_exp = np.real(np.fft.ifft2(ref_fft_ds * K))
        t = tile_ds[mask_ds]
        r = I_exp[mask_ds]
        denom = float(np.dot(r, r))
        if denom == 0.0:
            return np.inf
        amp = float(np.dot(t, r)) / denom  # optimal amplitude (closed-form)
        return float(np.sum((t - amp * r) ** 2))

    result = minimize_scalar(residual, bounds=(0.0, 20.0), method='bounded')
    return max(0.0, float(result.x))


def _estimate_plasmon_params_grid(tile, ref_beam, mask, pixel_size_nm,
                                   n_range=(0.0, 25.0),
                                   q_E_range=(0.003, 0.08),
                                   n_points=25, q_E_points=25,
                                   downsample=64, refine=True,
                                   edge_width=10,
                                   n_hint=None, q_E_hint=None):
    """
    2D grid search over (n, q_E) to find the plasmon parameters that best match
    the tile beam-edge profile, followed by an optional local refinement.

    Key design decisions
    --------------------
    Edge-only residual
        The plasmon effect is only visible at the beam boundary.  Fitting the
        flat interior (large area, noisy) dominates the residual and pulls the
        optimiser toward unrealistic (n, q_E) values.  The residual is therefore
        evaluated only on a ring of pixels near the beam boundary.

    Zero-padding on the downsampled convolution
        Without padding the FFT assumes periodic boundary conditions, which
        wraps the beam signal across the frame and biases the fit.

    Parameters
    ----------
    n_range : (float, float)    Search range for mean plasmon event count.
    q_E_range : (float, float)  Search range for q_E in cycles nm⁻¹.
                                Default covers E_plasmon ≈ 3–80 eV at 300 kV.
    n_points, q_E_points : int  Grid resolution along each axis.
    refine : bool               If True, run a local minimiser from the best
                                grid point to sub-grid precision.
    edge_width : int
        Width of the beam-edge ring (in downsampled pixels) used for the
        residual.  Default 10.

    Returns
    -------
    n_best, q_E_best : float
    residuals : (n_points, q_E_points) ndarray   full residual landscape
    n_vals, q_E_vals : 1-D ndarrays              grid axes
    """
    from scipy.optimize import minimize

    ny, nx = tile.shape
    ds_y, ds_x = min(downsample, ny), min(downsample, nx)

    tile_ds = fourier_interpolate(tile.astype(float), [ds_y, ds_x])
    ref_ds  = fourier_interpolate(ref_beam.astype(float), [ds_y, ds_x])
    mask_ds = fourier_interpolate(mask.astype(float), [ds_y, ds_x]) > 0.5
    if not mask_ds.any():
        mask_ds = np.ones((ds_y, ds_x), dtype=bool)

    # Edge ring: pixels within edge_width of the beam boundary, on both sides.
    # Inside ring: beam pixels excluded by erosion — captures the darkened rim.
    # Outside ring: non-beam pixels included by dilation — captures the
    #   plasmon-scattered halo, which is visible in this dataset.
    struct = np.ones((edge_width, edge_width), dtype=bool)
    eroded   = binary_erosion(mask_ds,  structure=struct)
    dilated  = binary_dilation(mask_ds, structure=struct)
    edge_mask_ds = (mask_ds & ~eroded) | (dilated & ~mask_ds)
    if not edge_mask_ds.any():
        edge_mask_ds = mask_ds  # beam too small to erode; use full mask

    py = pixel_size_nm * ny / ds_y
    px = pixel_size_nm * nx / ds_x

    # Zero-pad so the FFT convolution does not wrap the beam across the frame.
    pad_ds = max(4, int(np.ceil(3.0 / (q_E_range[0] * py))))
    ref_ds_pad = np.pad(ref_ds, pad_ds, mode='constant', constant_values=0.0)
    ds_yp, ds_xp = ref_ds_pad.shape
    qy_p = np.fft.fftfreq(ds_yp, d=py)
    qx_p = np.fft.fftfreq(ds_xp, d=px)
    q2_p = qy_p[:, None] ** 2 + qx_p[None, :] ** 2
    ref_fft_p = np.fft.fft2(ref_ds_pad)

    t = tile_ds[edge_mask_ds]

    def residual(n, q_E):
        n   = max(n,   0.0)
        q_E = max(q_E, 1e-6)
        K = np.exp(n * (q_E ** 2 / (q2_p + q_E ** 2) - 1.0))
        I_exp = np.real(np.fft.ifft2(ref_fft_p * K))[pad_ds:pad_ds + ds_y,
                                                       pad_ds:pad_ds + ds_x]
        r = I_exp[edge_mask_ds]
        denom = float(np.dot(r, r))
        if denom == 0.0:
            return np.inf
        amp = float(np.dot(t, r)) / denom
        return float(np.sum((t - amp * r) ** 2))

    if n_hint is not None and q_E_hint is not None:
        # Warm start: skip the grid and go straight to local refinement.
        n_best, q_E_best = float(n_hint), float(q_E_hint)
        residuals = np.empty((0, 0))
        n_vals = q_E_vals = np.empty(0)
    else:
        n_vals   = np.linspace(n_range[0],   n_range[1],   n_points)
        q_E_vals = np.linspace(q_E_range[0], q_E_range[1], q_E_points)

        residuals = np.array([[residual(n, q_E) for q_E in q_E_vals]
                               for n in n_vals])

        best = np.unravel_index(np.argmin(residuals), residuals.shape)
        n_best, q_E_best = float(n_vals[best[0]]), float(q_E_vals[best[1]])

    if refine:
        result = minimize(
            lambda x: residual(x[0], x[1]),
            x0=[n_best, q_E_best],
            method='Nelder-Mead',
            options={'xatol': 1e-3, 'fatol': 1e-6, 'maxiter': 500},
        )
        n_best   = max(0.0,              float(result.x[0]))
        q_E_best = max(float(q_E_range[0]), float(result.x[1]))

    return n_best, q_E_best, residuals, n_vals, q_E_vals


def _plot_plasmon_correction(tile, I_expected, corrected, mask,
                              n_avg, n_avg_method, pixel_size_nm):
    """Diagnostic plot for plasmon_beam_correction.  Call or comment out as needed."""
    ny, nx = tile.shape
    corrected_masked = np.where(mask, corrected, np.nan)

    half = 50
    row0, row1 = max(ny // 2 - half, 0), min(ny // 2 + half, ny)

    ls_tile        = tile[row0:row1, :].mean(axis=0)
    ls_corr_masked = np.nanmean(corrected_masked[row0:row1, :], axis=0)

    fig, axes = plt.subplots(2, 4, figsize=(20, 8),
                             gridspec_kw={"height_ratios": [3, 1]})

    def _clim(img, m):
        valid = img[m] if m.any() else img[np.isfinite(img)]
        valid = valid[np.isfinite(valid)]
        return tuple(np.percentile(valid, [1, 99])) if len(valid) else (img.min(), img.max())

    shared_clim = _clim(tile, mask)
    clims = [shared_clim, _clim(I_expected, mask), _clim(corrected, mask), shared_clim]

    for ax, img, title, (lo, hi) in zip(axes[0],
                               [tile, I_expected, corrected, corrected_masked],
                               ["tile (input)", "I_expected", "tile × correction",
                                "tile × correction (masked)"],
                               clims):
        ax.imshow(img, cmap="gray", vmin=lo, vmax=hi, origin="lower")
        ax.axhline(row0, color="r", linewidth=0.5, linestyle="--")
        ax.axhline(row1, color="r", linewidth=0.5, linestyle="--")
        ax.set_title(title)
        ax.axis("off")

    ax_ls = axes[1, 1]
    ax_ls.plot(ls_tile,        color="C0", label="input")
    ax_ls.plot(ls_corr_masked, color="C3", linestyle="--", label="masked corrected")
    ax_ls.set_xlim(0, nx - 1)
    ax_ls.set_xlabel("x (px)")
    ax_ls.set_ylabel("intensity")
    ax_ls.legend(fontsize=8, loc="upper center", bbox_to_anchor=(0.5, -0.35), ncol=2)

    for ax in (axes[1, 0], axes[1, 2], axes[1, 3]):
        ax.axis("off")

    fig.suptitle(f"Plasmon correction  (n_avg = {n_avg:.3f},  method = {n_avg_method},  "
                 f"pixel = {pixel_size_nm:.3f} nm)")
    plt.tight_layout()
    plt.show()


def align_beam_reference(tile, ref_beam, template_edges=None):
    """Integer (dy, dx) shift that aligns ref_beam to the beam in ``tile``.

    Uses the same edge cross-correlation as make_mask, so a tile's mask and its
    plasmon correction share one registration. The result is data-dependent:
    the same tile at a different dose (an even/odd half-stack, say) can land a
    pixel or two away, so callers that need a bit-for-bit reproducible
    correction should compute this once and feed it back to
    :func:`plasmon_beam_correction` via ``beam_shift``.
    """
    smoothed = convolve(tile.astype(float), Gaussian(3, tile.shape))
    image_edges = np.hypot(sobel(renormalize(smoothed), axis=0),
                           sobel(renormalize(smoothed), axis=1))
    if template_edges is None:
        smooth_ref = convolve(ref_beam.astype(float), Gaussian(3, ref_beam.shape))
        template_edges = np.hypot(sobel(smooth_ref, axis=0), sobel(smooth_ref, axis=1))
    return cross_correlate_alignment(image_edges, template_edges, returncoords=True)


def plasmon_beam_correction(tile, ref_beam, mask, pixel_size_nm,
                             template_edges=None, E_plasmon_eV=21.0, voltage_kV=300.0,
                             n_avg_method='grid', downsample=64,
                             n_range=(0.0, 25.0), q_E_range=(0.003, 0.08),
                             n_hint=None, q_E_hint=None,
                             n_fixed=None, q_E_fixed=None, beam_shift=None):
    """
    Correct beam edge darkening due to inelastic (plasmon) scattering.

    Plasmon scattering blurs the diffraction-plane intensity via a Lorentzian
    kernel (Mkhoyan et al. 2008, Eq. 11 in Brown et al. 2017).  In image space
    this appears as an attenuation of high spatial frequencies, making beam
    edges darker than the centre.

    The correction is:
        1. Align ref_beam to the beam position in tile via edge cross-correlation
           (same approach as make_mask).
        2. Build the single-plasmon Lorentzian in image k-space:
               P(q) = q_E^2 / (q^2 + q_E^2)   [normalised: P(0) = 1]
        3. Estimate n_avg — either by least-squares profile shape-matching
           ('lsq', default) or by the log-ratio of mean intensities
           ('log_ratio').
        4. Form the multi-plasmon (Poisson) kernel:
               K(q) = exp(n_avg * (P(q) - 1))
        5. Compute the expected blurred reference: I_exp = IFFT(FFT(I_ref) * K)
        6. Correction map: C = I_ref / I_exp  (applied within the beam mask)

    Parameters
    ----------
    tile : (ny, nx) ndarray
        Tile image to correct.
    ref_beam : (ny, nx) ndarray
        Raw (non-binary) reference beam image, typically the template image
        acquired through vacuum at the same settings as the data.
    mask : (ny, nx) bool ndarray
        Valid pixel mask (beam interior); used to estimate n_avg and to
        restrict where the correction is applied.
    pixel_size_nm : float
        Effective pixel size in nm (after any binning).
    template_edges : (ny, nx) ndarray, optional
        Pre-computed Sobel edge map of the reference beam, used for alignment.
        If None it is computed from ref_beam.  Pass the same array used by
        make_mask to avoid repeating the edge computation.
    E_plasmon_eV : float
        Plasmon energy in eV.  Default 21 eV (vitreous ice / water).
    voltage_kV : float
        Accelerating voltage in kV.  Default 300.
    n_avg_method : {'grid', 'lsq', 'log_ratio'}
        How to estimate plasmon parameters.
        'grid' (default): 2-D grid search over both n and q_E on a downsampled
        image, followed by Nelder-Mead refinement.  Both parameters are free,
        so the fit does not rely on the physical E_plasmon_eV / voltage_kV
        estimates.  The fitted q_E overrides the physics-based value.
        'lsq': 1-D bounded search over n only, with q_E fixed from physics.
        'log_ratio': fastest but coarsest — log(ref_mean / tile_mean).
    downsample : int
        Side length (pixels) of the image used for 'grid' / 'lsq' fitting.
        Default 64.
    n_range : (float, float)
        Search bounds for n used by the 'grid' method.  Default (0, 25).
    q_E_range : (float, float)
        Search bounds for q_E in cycles nm⁻¹ used by the 'grid' method.
        Default (0.003, 0.08), covering E_plasmon ≈ 3–80 eV at 300 kV.
    beam_shift : (int, int) or None
        Pre-computed (dy, dx) alignment of ref_beam to this tile, as returned
        by :func:`align_beam_reference`.  When None it is measured from the
        tile, which makes the correction depend on the tile's noise; pass a
        stored value to reproduce an earlier run's correction exactly.

    Returns
    -------
    corrected : (ny, nx) ndarray
        Tile with beam-edge darkening corrected.
    """
    # Align ref_beam to the beam position in this tile via edge cross-correlation,
    # matching the approach used in make_mask so both share the same registration.
    if beam_shift is None:
        beam_shift = align_beam_reference(tile, ref_beam, template_edges)
    dy, dx = (int(beam_shift[0]), int(beam_shift[1]))
    ref_beam = roll_no_periodic(ref_beam.astype(float), (-dy, -dx), axis=(0, 1))

    #plot_plasmon_sweep(tile, ref_beam, mask, pixel_size_nm,
                    #    E_plasmon_eV=E_plasmon_eV, voltage_kV=voltage_kV)

    # Relativistic speed factor β
    m0c2 = 511e3  # eV
    beta2 = 1.0 - (m0c2 / (m0c2 + voltage_kV * 1e3)) ** 2

    # Physics-based q_E in cycles nm^{-1} (used as fallback / starting point).
    # hc = 2π·ħc ≈ 1239.8 eV·nm; fftfreq returns cycles/unit not rad/unit.
    hc = 1239.8  # eV·nm
    q_E_phys = E_plasmon_eV / (hc * np.sqrt(beta2))
    q_E = q_E_phys

    if n_fixed is not None and q_E_fixed is not None:
        # Parameters locked in from a reference tile — skip all fitting.
        n_avg, q_E = float(n_fixed), float(q_E_fixed)
    elif n_avg_method == 'grid':
        n_avg, q_E, _res, _nv, _qv = _estimate_plasmon_params_grid(
            tile, ref_beam, mask, pixel_size_nm,
            n_range=n_range, q_E_range=q_E_range, downsample=downsample,
            n_hint=n_hint, q_E_hint=q_E_hint,
        )
        # print(f"Grid fit: n_avg={n_avg:.3f}  q_E={q_E:.5f} cyc/nm  "
        #       f"(E_eff≈{q_E * hc * np.sqrt(beta2):.1f} eV)")
        if q_E < q_E_phys / 5:
            warnings.warn(
                f"Fitted q_E ({q_E:.5f} cyc/nm, E_eff≈{q_E * hc * np.sqrt(beta2):.2f} eV) "
                f"is more than 5× below the physics-based value "
                f"({q_E_phys:.5f} cyc/nm, E_plasmon={E_plasmon_eV:.1f} eV at {voltage_kV:.0f} kV). "
                f"The reference tile is likely unsuitable (thick sample, no visible beam, or "
                f"beam edge not in frame). Use --correct_beam_edges N to specify a different tile.",
                RuntimeWarning, stacklevel=2,
            )
    elif n_avg_method == 'lsq':
        n_avg = _estimate_n_avg_lsq(tile, ref_beam, mask, q_E, pixel_size_nm,
                                    downsample=downsample)
    else:
        # Log-ratio: n_avg ≈ -ln(tile_mean / ref_mean); underestimates when
        # images are energy-filtered because intensity loss is not purely from
        # electrons leaving the beam mask.
        valid = mask if mask.any() else np.ones_like(mask)  # fall back to full image if mask is empty to avoid mean([]) → nan
        ref_mean  = float(np.mean(ref_beam[valid]))
        tile_mean = float(np.mean(tile[valid]))
        if ref_mean <= 0.0:
            return tile.copy(), 0.0, q_E
        n_avg = max(0.0, -np.log(np.clip(tile_mean / ref_mean, 1e-6, 1.0)))

    if n_avg == 0.0:
        return tile.copy(), 0.0, q_E

    ny, nx = ref_beam.shape

    # Zero-pad ref_beam before convolving to avoid wrap-around artefacts when
    # the beam edge is near the image boundary.  Padding = 3 × kernel half-width
    # in pixels; this is the minimum needed so the Lorentzian tail has decayed to
    # a negligible level before it would wrap.  Direct spatial convolution is
    # O(N²·R²) vs O((N+2P)²·log(N+2P)²) here — FFT wins for any realistic R.
    pad = max(16, int(np.ceil(3.0 / (q_E * pixel_size_nm))))
    pad = min(pad, max(ny, nx))  # guard: spurious small q_E can make pad >> tile size
    ref_padded = np.pad(ref_beam, pad, mode='constant', constant_values=0.0)
    ny_p, nx_p = ref_padded.shape
    qy_p = np.fft.fftfreq(ny_p, d=pixel_size_nm)
    qx_p = np.fft.fftfreq(nx_p, d=pixel_size_nm)
    K_p = np.exp(n_avg * (q_E ** 2 / (qy_p[:, None] ** 2 + qx_p[None, :] ** 2 + q_E ** 2) - 1.0))

    # Expected beam profile after plasmon blurring; crop back to original size
    I_expected = np.real(np.fft.ifft2(np.fft.fft2(ref_padded) * K_p))[pad:pad + ny, pad:pad + nx]

    # Correction map: ratio of reference to expected, clipped to avoid
    # amplifying noise outside the beam or in low-signal regions
    threshold = 0.05 * float(np.max(I_expected))
    correction = np.where(
        mask & (I_expected > threshold),
        np.clip(ref_beam / np.where(I_expected > threshold, I_expected, 1.0), 0.0, 10.0),
        1.0,
    )

    corrected = tile * correction

    # _plot_plasmon_correction(tile, I_expected, corrected, mask,
    #                           n_avg, n_avg_method, pixel_size_nm)

    return corrected, n_avg, q_E


def flatten_beam(image,mask,rotation=0):
    """
    Flattens the beam profile in an image by fitting and subtracting a 4th order polynomial surface.

    Parameters:
    image (numpy.ndarray): The input 2D image array.
    mask (numpy.ndarray): A boolean mask array where True values indicate the region of interest.
    rotation (float, optional): The rotation angle in degrees to apply to the coordinates. Default is 0.
++
    Returns:
    numpy.ndarray: The flattened image with the beam profile subtracted.
    """
    im  = fourier_interpolate(image,[256,256])
    msk = make_mask(im)
    y,x = r_space_array(im.shape,gridsize=im.shape,meshed=True)
    y0,x0= center_of_mass_2d(np.where(msk,1,0))
    y = y - (y0 - image.shape[0]//2)
    x = x - (x0 - image.shape[1]//2)
    M = rotation_matrix(-rotation)
    y,x = np.einsum('ij,jkl->ikl',M,np.array([y,x]))
    
    x_masked = x[msk].flatten()
    y_masked = y[msk].flatten()
    m_masked = im[msk].flatten()
    # A = np.column_stack([x_masked.ravel()**4,y_masked.ravel()**4,np.ones_like(x_masked.ravel())])
    A = np.column_stack([x_masked.ravel()**4,y_masked.ravel()**4,x_masked.ravel()**2,y_masked.ravel()**2,np.ones_like(x_masked.ravel())])
    coeffs, _, _, _ = np.linalg.lstsq(A, m_masked, rcond=None)
    fitted_2d = np.zeros_like(image)
    fitted_2d = coeffs[0]*x**4+coeffs[1]*y**4+coeffs[2]*x**2+coeffs[3]*y**2
    # fig,ax = plt.subplots(ncols=3)
    # vmin=np.percentile(image[mask],1)
    # vmax=np.percentile(image[mask],99)
    # ax[0].imshow(image,vmin=vmin,vmax=vmax)
    # ax[1].imshow(fitted_2d)
    mean = np.mean(image[mask])
    image[mask]= image[mask]-fourier_interpolate(fitted_2d,image.shape)[mask]+mean
    # vmin=np.percentile(image[mask],1)
    # vmax=np.percentile(image[mask],99)
    # ax[1].imshow(image,vmin=vmin,vmax=vmax)
    # ax[2].imshow(fourier_interpolate(fitted_2d,image.shape))
    # plt.show(block=True)
    # image[mask] -= fitted_2d[mask]
    return image

def determine_square_beam_angle(m):
    """
    Determines the orientation angle of a square beam in an image.
    Parameters:
    m (numpy.ndarray): The input 2D array representing the image of the square beam.
    Returns:
    float: The orientation angle of the square beam in degrees, adjusted to be within the range [-45, 45] degrees.
    Notes:
    - The function first resizes the image such that the longest axis is 128 pixels to speed up the calculation.
    - It then pads the resized image with zeros.
    - The Radon transform is applied to the padded image to compute the sinogram.
    - The angle corresponding to the minimum value in the central row of the sinogram is determined.
    - This angle is adjusted to be within the range [-45, 45] degrees.
    """
    
    # Reshape so that longest axis is 128 pixels to speed up the calculation
    if m.shape[0]>m.shape[1]:
        news = [128,128*m.shape[1]//m.shape[0]]
    else:    
        news = [128*m.shape[0]//m.shape[1],128]
    m_ = fourier_interpolate(m,news)

    # Pad the image with zeros
    m_ = np.pad(m_,((64,64),(64,64)))
    
    N = 180
    theta = np.linspace(0, 180, N)

    from skimage.transform import radon
    sinogram = fourier_interpolate(radon(m_, theta=theta),[N,N])
    rot = theta[np.argmin(sinogram[sinogram.shape[0]//2])]
    rotmod45 = (rot+45)%90-45
    return rotmod45

def circular_mask(size, radius=None, center=None):
    """
    Create a binary circular mask in a 2D array.

    Parameters:
    - size (int, int): Size of the array (rows, cols).
    - radius (int, optional): Radius of the circle. Defaults to min(size)/2.
    - center (int, int, optional): Center of the circle (row, col). Defaults to the center of the array.

    Returns:
    - mask (2D numpy array): Binary mask with a circle.
    """
    rows, cols = size

    if center is None:
        center = (rows // 2, cols // 2)
    if radius is None:
        radius = min(rows, cols) // 2

    Y, X = np.ogrid[:rows, :cols]
    dist_from_center = np.sqrt((X - center[1]) ** 2 + (Y - center[0]) ** 2)

    return dist_from_center <= radius


def iterative_edge_smoothing(array, mask, niterations=5, pow=4, initial_radius=None):
    """
    Smooths the edges of a given array iteratively using a circular convolution kernel.
    Parameters:
    array (numpy.ndarray): The input array to be smoothed.
    mask (numpy.ndarray): A boolean mask array where True values indicate the regions to be preserved.
    niterations (int, optional): The number of iterations for the smoothing process. Default is 5.
    pow (int, optional): The power to which the iteration index is raised to control the radius of the smoothing kernel. Default is 4.
    Returns:
    numpy.ndarray: The smoothed array with edges processed according to the mask.
    """

    if initial_radius is None:
        R = np.mean(array.shape) / 10
    else:
        R = initial_radius
    for i in tqdm(range(niterations), desc="smoothing"):
        radius = (R - 1) / niterations**pow * np.abs(i - niterations) ** pow + 1
        kernel = np.where(circular_mask(array.shape, radius=radius, center=None), 1, 0)
        kernel = np.fft.fftshift(kernel)
        kernel[0, 0] = 0
        kernel = kernel / np.sum(kernel)
        array = np.where(mask, array, convolve(array, kernel))

    return array


def make_mask(im, shrinkn=20, smoothing_kernel=3, medianthreshold=0.4,
              absolutethreshold=None, template_mask=None, template_edges=None,
              bin_factor=1, erosion_method="binary_erosion"):
    """
    Generate a binary mask from an image by applying a Gaussian filter, filling holes,
    and then shrinking the mask with morphological erosion.

    When a template_mask is provided, edge-based cross-correlation is used to align
    the template to the beam position in the image. This is more robust than
    thresholding when thick sample regions are as bright as (or brighter than) the beam.

    Parameters:
    -----------
    im : numpy.ndarray
        Input 2D image array.
    shrinkn : int, optional
        Number of pixels to shrink the mask boundary inward by. Default is 20.
    smoothing_kernel : float, optional
        Sigma of the Gaussian smoothing kernel before applying the mask (3 by default).
    medianthreshold : float, optional
        Threshold as a fraction of the image median used in the fallback method. Default 0.4.
    absolutethreshold : float, optional
        If provided, used as an absolute threshold instead of the median-based one.
    template_mask : numpy.ndarray, optional
        A binary template of the expected beam shape. When provided, Sobel edge maps of
        the (smoothed) image and of the template boundary are cross-correlated to locate
        the beam, and the template is shifted to that position. The result is insensitive
        to bright/dark sample regions inside the beam.
    erosion_method : {"binary_erosion", "distance_transform"}, optional
        Only used by the fallback (no ``template_mask``) path -- the template
        path always uses ``binary_erosion``. ``"binary_erosion"`` (default)
        erodes with a disk structuring element of radius ``shrinkn``; this is
        the historical approach and removes exactly ``shrinkn`` pixels of
        boundary. ``"distance_transform"`` approximates the same disk erosion
        via ``distance_transform_edt(mask) >= shrinkn`` instead, which is O(N)
        rather than O(N * shrinkn**2) and so is faster for large ``shrinkn``
        on full-resolution images; it removes ``shrinkn - 1`` pixels (an
        off-by-one at the exact boundary from the ``>=`` vs strict-disk
        comparison), which is immaterial at the scale ``shrinkn`` is used at
        here.

    Returns:
    --------
    numpy.ndarray : Binary mask of the same shape as the input image.
    """
    if template_mask is not None:
        # Resize template to match image shape if necessary
        if template_mask.shape != im.shape:
            template_mask = fourier_interpolate(
                template_mask.astype(float), im.shape
            ) > 0.5

        # Compute Sobel edge magnitude on the smoothed image
        smoothed_im = convolve(im, Gaussian(smoothing_kernel, im.shape))
        norm_im = renormalize(smoothed_im)
        image_edges = np.hypot(sobel(norm_im, axis=0), sobel(norm_im, axis=1))

        # Compute Sobel edge magnitude on the template boundary
        if template_edges is None:
            tmpl_float = template_mask.astype(float)
            template_edges = np.hypot(sobel(tmpl_float, axis=0), sobel(tmpl_float, axis=1))

        

        # Find the shift that best aligns template edges to image edges
        dy, dx = cross_correlate_alignment(image_edges, template_edges, returncoords=True)

        # Shift template to align with the beam
        mask = roll_no_periodic(template_mask, (-dy, -dx), axis=(0, 1))


        # Optionally erode to keep away from the beam edge. radius=shrinkn
        # (not shrinkn/2) to match the fallback threshold path below --
        # otherwise the same shrinkn value shrinks half as much here as it
        # does there, which is exactly the mismatch that made Choose_fringe_
        # size.py (always the fallback path) and stitch.py --referencebeam
        # (always this path) disagree on what a given fringe size does.
        if shrinkn > 0:
            struct_elem = circular_mask([shrinkn * 2 + 1] * 2, radius=shrinkn)
            mask = binary_erosion(mask, structure=struct_elem)

        return mask

    # --- Fallback: fast threshold-based approach ---
    # Step 1: Downsample — only the Gaussian smoothing runs at 1/bin_factor
    # resolution, since that's the expensive step. Everything downstream
    # (threshold, fill, erode) runs at full resolution so the mask boundary
    # lands at full-image accuracy rather than being quantized to
    # bin_factor-sized blocks.
    im_work = im[::bin_factor, ::bin_factor] if bin_factor > 1 else im

    # Step 2: Separable Gaussian smooth (replaces full-image FFT convolution)
    smoothed_im = gaussian_filter(im_work.astype(np.float32), sigma=smoothing_kernel)

    # Step 2b: Upsample the smoothed intensity field (not yet a binary mask)
    # back to full resolution. Interpolating the continuous field preserves
    # the true edge position; interpolating an already-binarized mask cannot
    # recover position lost to the downsample.
    if bin_factor > 1:
        smoothed_im = fourier_interpolate(smoothed_im, im.shape)

    # Step 3: Threshold
    if absolutethreshold is not None:
        mask = smoothed_im > absolutethreshold
    else:
        mask = smoothed_im > medianthreshold * np.median(im_work)

    # Step 4: Fill holes
    mask = binary_fill_holes(mask)

    # Step 5: Shrink the mask boundary inward by shrinkn pixels.
    if shrinkn > 0:
        if erosion_method == "binary_erosion":
            struct_elem = circular_mask(
                [shrinkn * 2 + 1] * 2, radius=shrinkn
            )
            mask = binary_erosion(mask, structure=struct_elem)
        elif erosion_method == "distance_transform":
            # O(N) regardless of radius, vs O(N * shrinkn**2) for
            # binary_erosion with a large structuring element.
            mask = distance_transform_edt(mask) >= shrinkn
        else:
            raise ValueError(f"Unknown erosion_method: {erosion_method!r}")

    return mask.astype(bool)


def cross_correlate_alignment(im, template, returncoords=True):
    """
    Align an image to a template using cross-correlation.

    This function calculates the cross-correlation between a template and an image,
    identifies the point of maximum correlation (which indicates the best alignment),
    and either returns the coordinates of this point or returns the image aligned to the template.

    Parameters:
    -----------
    im : numpy.ndarray
        The input image to be aligned. This is the image that will be adjusted.
    template : numpy.ndarray
        The template image that is used as a reference for alignment.
    returncoords : bool, optional
        If True, the function returns the coordinates of the maximum correlation.
        If False, the function returns the aligned image. Default is True.

    Returns:
    --------
    tuple of int or numpy.ndarray :
        If `returncoords` is True, returns a tuple (y, x) representing the coordinates of the
        maximum correlation.
        If `returncoords` is False, returns the aligned image as a numpy array.
    """

    # Step 1: Calculate the cross-correlation between the template and the image
    corr = correlate(template, im)

    # Step 2: Find the coordinates (y, x) of the maximum value in the correlation matrix
    y, x = np.unravel_index(np.argmax(corr), im.shape)

    y, x = [(i + N // 2) % N - N // 2 for i, N in zip((y, x), im.shape)]

    # Step 3: Depending on the value of returncoords, return the coordinates or the aligned image
    if returncoords:
        return y, x  # Return the coordinates of the maximum correlation
    else:
        # Align the image by rolling it to place the max correlation point at the origin
        return np.roll(im, (-y, -x), axis=(-2, -1))


def roll_no_periodic(arr, shift, fill_value=0, axis=None):
    """
    Rolls an array along the given axis or axes but without periodic boundary conditions.
    Vacated positions will be filled with fill_value.

    Parameters:
    - arr: numpy array to be shifted.
    - shift: int or tuple of ints, amount to shift. Positive values shift right/down, negative values shift left/up.
    - fill_value: value to place in the vacated positions.
    - axis: int or tuple of ints, the axis or axes to roll along. If None, roll along all axes.

    Returns:
    - A new numpy array with the same shape as arr, but shifted.
    """
    # If axis is None, shift across all axes
    if axis is None:
        axis = tuple(range(arr.ndim))
        shift = (shift,) * arr.ndim
    elif isinstance(axis, int):
        axis = (axis,)
        shift = (shift,) if isinstance(shift, int) else shift

    result = np.full_like(arr, fill_value)  # Create an array filled with the fill_value

    if len(axis) != len(shift):
        raise ValueError("The number of shifts must match the number of axes.")
    src_slice = [slice(None)] * arr.ndim
    dst_slice = [slice(None)] * arr.ndim

    for ax, s in zip(axis, shift):
        if s == 0:
            continue  # No shift for this axis

        # Determine the slices that will remain after the shift
        if s > 0:
            src_slice[ax] = slice(0, -s)
            dst_slice[ax] = slice(s, None)
        elif s < 0:
            src_slice[ax] = slice(-s, None)
            dst_slice[ax] = slice(0, s)

    result[tuple(dst_slice)] = arr[tuple(src_slice)]

    return result


def _masked_phase_cross_correlation(
    reference_image, moving_image, reference_mask, moving_mask=None, overlap_ratio=0.3
):
    """Masked image translation registration by masked normalized
    cross-correlation.

    Parameters
    ----------
    reference_image : ndarray
        Reference image.
    moving_image : ndarray
        Image to register. Must be same dimensionality as ``reference_image``,
        but not necessarily the same size.
    reference_mask : ndarray
        Boolean mask for ``reference_image``. The mask should evaluate
        to ``True`` (or 1) on valid pixels. ``reference_mask`` should
        have the same shape as ``reference_image``.
    moving_mask : ndarray or None, optional
        Boolean mask for ``moving_image``. The mask should evaluate to ``True``
        (or 1) on valid pixels. ``moving_mask`` should have the same shape
        as ``moving_image``. If ``None``, ``reference_mask`` will be used.
    overlap_ratio : float, optional
        Minimum allowed overlap ratio between images. The correlation for
        translations corresponding with an overlap ratio lower than this
        threshold will be ignored. A lower `overlap_ratio` leads to smaller
        maximum translation, while a higher `overlap_ratio` leads to greater
        robustness against spurious matches due to small overlap between
        masked images.

    Returns
    -------
    shifts : ndarray
        Shift vector (in pixels) required to register ``moving_image``
        with ``reference_image``. Axis ordering is consistent with
        numpy (e.g. Z, Y, X)

    References
    ----------
    .. [1] Dirk Padfield. Masked Object Registration in the Fourier Domain.
           IEEE Transactions on Image Processing, vol. 21(5),
           pp. 2706-2718 (2012). :DOI:`10.1109/TIP.2011.2181402`
    .. [2] D. Padfield. "Masked FFT registration". In Proc. Computer Vision and
           Pattern Recognition, pp. 2918-2925 (2010).
           :DOI:`10.1109/CVPR.2010.5540032`

    """
    if moving_mask is None:
        if reference_image.shape != moving_image.shape:
            raise ValueError(
                "Input images have different shapes, moving_mask must "
                "be explicitely set."
            )
        moving_mask = reference_mask.astype(bool)

    # We need masks to be of the same size as their respective images
    for im, mask in [(reference_image, reference_mask), (moving_image, moving_mask)]:
        if im.shape != mask.shape:
            raise ValueError("Image sizes must match their respective mask sizes.")

    xcorr = cross_correlate_masked(
        moving_image,
        reference_image,
        moving_mask,
        reference_mask,
        axes=tuple(range(moving_image.ndim)),
        mode="full",
        overlap_ratio=overlap_ratio,
    )

    # Generalize to the average of multiple equal maxima
    maxima = np.stack(np.nonzero(xcorr == xcorr.max()), axis=1)
    center = np.mean(maxima, axis=0)
    shifts = center - np.array(reference_image.shape) + 1

    # The mismatch in size will impact the center location of the
    # cross-correlation
    size_mismatch = np.array(moving_image.shape) - np.array(reference_image.shape)

    return -shifts + (size_mismatch / 2), xcorr.max()


def cross_correlate_tiles(
    positions,
    tiles,
    masks,
    overlaps,
    pixel_size,
    max_correction=0.3,
    generate_plot=False,
    cross_corr_param_file=None,
):
    """
    Perform cross-correlation-based alignment of image tiles and adjust their positions.

    This function aligns overlapping image tiles by calculating the relative shifts between them
    using masked phase cross-correlation. The shifts are then used to adjust the global positions
    of the tiles through least squares minimization. The function supports visualizing the tile
    positions and shift vectors if desired.

    Args:
        positions (ndarray): Nx2 array of the initial x, y coordinates of the tiles in micrometers.
        tiles (list of ndarrays): List of 2D arrays representing the image tiles.
        masks (list of ndarrays): List of binary masks representing the valid regions of the tiles.
        overlaps (list of tuples): Pairs of indices representing which tiles overlap and should be aligned.
        pixel_size (float): The pixel size in microns.
        max_correction (float, optional): Maximum allowed correction (in microns) for shifts between tiles. Defaults to 0.1.
        generate_plot (bool or string, optional): Whether to generate a plot visualizing the tile positions and shift vectors. Defaults to False.
                                                  If a string this will be the filename that the plot will be saved as.

    Returns:
        ndarray: Updated Nx2 array of the adjusted x, y coordinates of the tiles.

    Notes:
        - The alignment is solved as a least squares problem (Ax = b) to minimize the relative shifts between overlapping tiles.
        - If some tiles are not connected to others through reliable shift determinations, their positions are adjusted
        using the initial relative positions inferred from the microscope.

    References:
        - Masked phase cross-correlation: https://scikit-image.org/docs/stable/auto_examples/registration/plot_masked_register_translation.html
        - Dirk Padfield, "Masked object registration in the Fourier domain", IEEE Transactions on Image Processing, 2011.
    """
    pixels = tiles[0].shape
    cmap = plt.get_cmap("viridis")

    # Convert maximum allowed shift in microns (max_correction) to pixels (max_shift)
    max_shift = max_correction / pixel_size * 1e4

    # We will solve global alignment of tiles by least squares Ax = b
    # matrix problem (https://en.wikipedia.org/wiki/Linear_least_squares).
    # N is the number of tiles times two (for x and y coordinates)
    N = 2 * len(positions)
    # Initialize lists which we will append rows for the A and b matrices
    A = []
    b = []

    genplot = False
    if type(generate_plot) is bool:
        genplot = generate_plot
        show = True
        savefig = False
    elif type(generate_plot) is str:
        genplot = True
        show = False
        savefig = True

    if genplot:
        xcorfig, xcorax = plt.subplots(figsize=(8, 8))
        xcorax.plot(*positions.T, "ko", label="Initial tile positions")
        for i, pos in enumerate(positions):
            xcorax.annotate(str(i), pos)

    G = nx.Graph()
    # Add nodes to graph
    G.add_nodes_from(list(range(len(positions))))

    weights = []
    xcorr = []
    deltas = []
    shifts = []

    # TODO make this threaded
    if cross_corr_param_file is not None:
        xcorr, deltas = [
            load_array_from_hdf5(cross_corr_param_file, x)
            for x in ("cross_correlations", "relative_shifts")
        ]
    for ind, (i, j) in enumerate(tqdm(overlaps, desc="Cross-correlation alignment")):
        # Retrieve image shifts for the overlapping tiles
        x1 = positions[i]
        x2 = positions[j]

        # Calculate relative shift in pixels
        dx = [int(x) for x in (x2 - x1) / pixel_size * 1e4][::-1]
        if cross_corr_param_file is None:
            # Align tiles by masked phase cross correlation, see:
            # https://scikit-image.org/docs/stable/auto_examples/registration/plot_masked_register_translation.html
            # and Padfield, Dirk. "Masked object registration in the Fourier domain." IEEE Transactions on image processing 21.5 (2011): 2706-2718.

            # Get the indices for array overlap
            i1, i2 = array_overlap(dx[0], pixels[0])
            j1, j2 = array_overlap(dx[1], pixels[1])

            # Make masks for (estimated) overlapping areas of each array
            reference_mask = np.zeros_like(masks[i], dtype=bool)
            reference_mask[i1[0] : i1[1], j1[0] : j1[1]] = masks[i][
                i1[0] : i1[1], j1[0] : j1[1]
            ]
            moving_mask = np.zeros_like(masks[j], dtype=bool)
            moving_mask[i2[0] : i2[1], j2[0] : j2[1]] = masks[j][
                i2[0] : i2[1], j2[0] : j2[1]
            ]

            # Calculate shift by masked cross correlation with ordering (Y,X)
            detected_shift, xcorrmax = _masked_phase_cross_correlation(
                tiles[i],
                tiles[j],
                reference_mask=reference_mask,
                moving_mask=moving_mask,
            )
            # delta is measured shift in microns
            delta = np.asarray(detected_shift) * pixel_size * 1e-4
            xcorr.append(xcorrmax)
            deltas.append(delta)
        else:
            delta = deltas[ind]
            xcorrmax = xcorr[ind]
            detected_shift = delta / pixel_size * 1e-4

        # shifttoolarge=True
        if generate_plot:
            if ind == 0:
                label = "Shift vector from cross-correlation"
            else:
                label = None

        shifttoolarge = (
            np.linalg.norm(np.asarray(dx) * pixel_size * 1e-4 - delta) > max_correction
        )
        # detected_shift is position j (x2) - position i (x1)
        if not shifttoolarge:
            Arow = np.zeros((2, N))
            Arow[0, 2 * j] = 1
            Arow[0, 2 * i] = -1
            Arow[1, 2 * j + 1] = 1
            Arow[1, 2 * i + 1] = -1
            A.append(xcorrmax * Arow)
            b[len(b) :] = (xcorrmax * delta).tolist()
            # Add valid connection to graph
            G.add_edge(i, j)
        # Weight by the maximum value of cross-correlation
        weights.append(xcorrmax)

    # Join rows of A into numpy matrix
    if len(A) > 0:
        A = np.concatenate(A, axis=0)
    else:
        A = np.asarray(A)
    # Normalize by mean weight
    meanweight = np.mean(weights)
    A /= meanweight
    b = (np.asarray(b) / meanweight).tolist()

    # Function to find the closest points using KDTree
    def find_closest_points_with_kdtree(list1, list2):
        # Create KDTree for the second list
        tree = KDTree(list2)

        # Query the tree with all points in the first list
        distances, indices = tree.query(list1)

        # Find the closest pair
        min_distance_index = distances.argmin()
        closest_point_list1 = list1[min_distance_index]
        closest_point_list2 = list2[indices[min_distance_index]]

        return min_distance_index, indices[min_distance_index]

    # For some tiles there will be no reliable shift determinations
    # from cross-correlation linking them to the other tiles,
    # in this case default to using the original
    # shifts implied by the microscope image shifts if groups of tiles
    
    if not nx.is_connected(G):
        # if False:
        extraA = []
        islands = sorted(nx.connected_components(G), key=len)
        largest_island = islands.pop()
        largest_island_points = [positions[x] for x in largest_island]
        # anchor = largest_island.pop()
        # for i in np.where(np.all(A == 0, axis=0))[0]:
        for island in islands:
            island_points = [positions[i] for i in island]


            i, j = find_closest_points_with_kdtree(island_points, largest_island_points)

            i = list(island)[i]
            j = list(largest_island)[j]
            # i = x.pop()
            x1 = positions[j]
            x2 = positions[i]
            delta = x1 - x2

            # Loop over x and y components
            for jj in range(2):
                # Make new row to add to A matrix
                Arow = np.zeros((N))
                Arow[2 * j + jj] = 1
                Arow[2 * i + jj] = -1
                extraA.append(Arow)

            b[len(b) :] = delta[::-1].tolist()
        A = np.stack(A.tolist() + extraA, axis=0)

    def objective_function(x, A, b, positions, lam):
        obj = np.linalg.norm(A @ x - b)
        obj += lam * np.linalg.norm(positions - x)
        return obj

    from scipy.optimize import minimize

    xvec = positions[:, ::-1].ravel()
    # res = minimize(objective_function,xvec,args = (A,b,copy.deepcopy(xvec),0.0))
    # x = res.x
    # newpositions[]

    x, residuals, rank, s = np.linalg.lstsq(A, np.asarray(b), rcond=-1)
    # matax[1].plot(s)
    newy = x[::2]
    newx = x[1::2]

    # The absolute position is unconstrained in least squares minimization
    # Pin the position closest to the origin to its original absolute position
    i = np.argmin(np.linalg.norm(positions, axis=1))
    newx -= newx[i] - positions[i][0]
    newy -= newy[i] - positions[i][1]

    newpositions = np.zeros_like(positions)
    newpositions[:, 0] = newx
    newpositions[:, 1] = newy
    # newpositions = res.x.reshape(positions.shape)[:,::-1]
    if show:
        plt.show(block=True)
    return newpositions, xcorr, deltas
```
