## Supplementary material for "Whole cryo-FIB lamella acquisition at high magnification with rectangular beam montage cryo-electron tomography": Source code: visualize_irls.py

```
"""Animate the IRLS consensus fit that stitch.py's montage() runs internally.

Replays irls_positions() on the real edge measurements saved by a previous
stitch run (the "*_refined_positions.h5" sidecar), capturing every iteration's
weights and fitted positions. Each animation frame is one iteration: a
histogram of edge residuals (|measured - fitted displacement|) at that
iteration, with the Tukey/Huber weight function w(u) superimposed (shared,
normalized 0-1 axis: bar heights are density-normalized so no second y-scale
is needed).

Usage:
  python3 visualize_irls.py <refined_positions.h5> [options]
"""
import argparse
import logging
import os
import sys

import matplotlib

matplotlib.use("Agg")
import matplotlib.animation as animation
import matplotlib.pyplot as plt
import numpy as np

import processing_scripts.stitch as st

logging.basicConfig(level=logging.INFO, format="%(asctime)s %(levelname)s %(message)s")
logger = logging.getLogger("visualize_irls")

# dataviz palette (references/palette.md): blue = magnitude/histograms,
# orange = the one other identity on screen (the weight curve).
COLOR_HIST = "#2a78d6"
COLOR_WEIGHT = "#eb6834"
COLOR_THRESHOLD = "#52514e"


def load_edges(h5_path, binning):
    """`binning` must match the -b value the production stitch.py run used.

    The HDF5's "binned_pixel_size" dataset is a misnomer (see
    refine_montage_projmatch.py's _CanvasGeom docstring): stitch.py stores
    unbinned_pixel_size / binning there, not the canvas (binned) pixel size,
    which is unbinned_pixel_size * binning -- a factor of binning**2 apart.
    Un-mangle it here so the logged pixel size is the real bin-N value.
    """
    nominal = np.asarray(st.load_array_from_hdf5(h5_path, "Original_positions"), dtype=float)
    edge_ij = np.asarray(st.load_array_from_hdf5(h5_path, "overlaps"), dtype=int).reshape(-1, 2)
    edge_d = np.asarray(st.load_array_from_hdf5(h5_path, "relative_shifts"), dtype=float).reshape(-1, 2)
    misnamed = float(np.asarray(st.load_array_from_hdf5(h5_path, "binned_pixel_size")))
    pixel_size = misnamed * binning**2
    return nominal, edge_ij, edge_d, pixel_size


def irls_history(nominal_positions, edge_ij, edge_d, loss="tukey", threshold=None,
                  n_iterations=25, floor=0.0, tol=1e-9):
    """Re-run stitch.irls_positions(), keeping every iterate instead of just the last."""
    positions = nominal_positions.copy()
    if threshold is None:
        threshold = st._auto_threshold(st._edge_residuals(positions, edge_ij, edge_d), floor=floor)
    threshold = float(max(threshold, 1e-12))

    history = []
    for it in range(int(n_iterations)):
        residuals = st._edge_residuals(positions, edge_ij, edge_d)
        u = residuals / threshold
        if loss == "tukey":
            weights = np.where(u < 1.0, (1.0 - u**2) ** 2, 0.0)
        elif loss == "huber":
            weights = np.where(u <= 1.0, 1.0, 1.0 / np.maximum(u, 1e-12))
        else:
            raise ValueError(f"unknown loss {loss!r}; expected 'tukey' or 'huber'")
        new_positions, _n_bridges = st._least_squares_positions(
            nominal_positions, edge_ij, edge_d, weights
        )
        moved = float(np.max(np.linalg.norm(new_positions - positions, axis=1))) if len(positions) else 0.0
        history.append(dict(
            iteration=it, residuals=residuals, weights=weights,
            threshold=threshold, positions=new_positions.copy(), moved=moved,
        ))
        positions = new_positions
        if moved < tol:
            break
    return history


def weight_curve(u, loss):
    if loss == "tukey":
        return np.where(u < 1.0, (1.0 - u**2) ** 2, 0.0)
    return np.where(u <= 1.0, 1.0, 1.0 / np.maximum(u, 1e-12))


def build_animation(history, loss, bins, fps, hold_last, out_path):
    n_frames = len(history)

    # Stable axis ranges across all frames so bars don't jump around.
    all_resid_nm = np.concatenate([h["residuals"] for h in history]) * 1e3
    resid_xmax = max(
        3.0 * history[0]["threshold"] * 1e3,
        float(np.percentile(all_resid_nm, 95)),
    )

    fig, ax_l = plt.subplots(1, 1, figsize=(6.5, 5.5))
    fig.subplots_adjust(top=0.86, bottom=0.13)

    # Repeat the last (converged) frame a few times so the animation pauses there.
    frame_indices = list(range(n_frames)) + [n_frames - 1] * max(0, hold_last)

    def draw(idx):
        ax_l.clear()
        h = history[idx]
        resid_nm = h["residuals"] * 1e3
        threshold_nm = h["threshold"] * 1e3
        n_edges = len(resid_nm)
        n_active = int(np.sum(h["weights"] > 0))

        counts, edges = np.histogram(resid_nm, bins=bins, range=(0, resid_xmax))
        density = counts / counts.max() if counts.max() > 0 else counts.astype(float)
        ax_l.bar(
            edges[:-1], density, width=np.diff(edges), align="edge",
            color=COLOR_HIST, edgecolor="white", linewidth=0.3,
            label="residual histogram (normalized)",
        )
        u = np.linspace(0, resid_xmax, 300) / threshold_nm
        ax_l.plot(u * threshold_nm, weight_curve(u, loss), color=COLOR_WEIGHT, linewidth=2.2,
                   label=f"{loss} weight $w_{{ij}}$")
        ax_l.axvline(threshold_nm, color=COLOR_THRESHOLD, linestyle="--", linewidth=1.2,
                      label=f"tolerance ({threshold_nm:.0f} nm)")
        ax_l.set_xlim(0, resid_xmax)
        ax_l.set_ylim(0, 1.08)
        ax_l.set_xlabel("residual  |measured − fitted displacement|  (nm)")
        ax_l.set_ylabel("normalized count / weight")
        ax_l.set_title(f"residuals & weights  ({n_active}/{n_edges} edges active)")
        ax_l.legend(loc="upper right", fontsize=8, framealpha=0.9)

        fig.suptitle(
            f"IRLS consensus fit — iteration {idx + 1}/{n_frames}   "
            f"(max tile step {h['moved']*1e3:.1f} nm)",
            fontsize=12,
        )
        return []

    anim = animation.FuncAnimation(fig, draw, frames=frame_indices, blit=False)
    writer = animation.FFMpegWriter(fps=fps)
    logger.info("Writing %s (%d frames)...", out_path, len(frame_indices))
    anim.save(out_path, writer=writer)
    plt.close(fig)


def main():
    p = argparse.ArgumentParser(description=__doc__, formatter_class=argparse.RawDescriptionHelpFormatter)
    p.add_argument("h5_path", help="*_refined_positions.h5 sidecar from a stitch.py run")
    p.add_argument("--binning", type=int, default=4, help="Must match the production -b value")
    p.add_argument("--loss", choices=["tukey", "huber"], default="tukey")
    p.add_argument("--n-iterations", type=int, default=25)
    p.add_argument("--tol", type=float, default=1e-9)
    p.add_argument("--floor", type=float, default=0.0)
    p.add_argument("--threshold", type=float, default=None, help="Override the auto-derived tolerance (microns)")
    p.add_argument("--bins", type=int, default=50)
    p.add_argument("--fps", type=float, default=1.5)
    p.add_argument("--hold-last", type=int, default=6, help="Extra repeats of the final converged frame")
    p.add_argument("--out", default=None, help="Output .mp4 path (default: alongside the h5 file)")
    args = p.parse_args()

    nominal, edge_ij, edge_d, pixel_size = load_edges(args.h5_path, args.binning)
    logger.info("%d tiles, %d overlap edges, pixel size %.4f A/px (binned)", len(nominal), len(edge_ij), pixel_size)

    history = irls_history(
        nominal, edge_ij, edge_d, loss=args.loss, threshold=args.threshold,
        n_iterations=args.n_iterations, floor=args.floor, tol=args.tol,
    )
    logger.info(
        "IRLS ran %d iteration(s), converged=%s, final max step %.2f nm",
        len(history), history[-1]["moved"] < args.tol, history[-1]["moved"] * 1e3,
    )

    saved_refined = np.asarray(st.load_array_from_hdf5(args.h5_path, "Refined_positions"), dtype=float)
    replay_gap = np.max(np.linalg.norm(history[-1]["positions"] - saved_refined, axis=1)) * 1e3
    logger.info("Replay vs. saved Refined_positions: max discrepancy %.2f nm", replay_gap)

    out_path = args.out or os.path.splitext(args.h5_path)[0] + "_irls_animation.mp4"
    build_animation(
        history, args.loss, args.bins, args.fps, args.hold_last, out_path,
    )
    logger.info("Wrote %s", out_path)


if __name__ == "__main__":
    sys.exit(main())
```
