## Supplementary material for "Whole cryo-FIB lamella acquisition at high magnification with rectangular beam montage cryo-electron tomography": Source code: projection_matching_plan.md

# Per-tile montage refinement by projection matching

Design notes for `processing_scripts/refine_montage_projmatch.py`.

## Goal

AreTomo refines a tilt series by reprojecting the tomogram and shifting each
*whole image* to match. For a square-beam montage each tilt is 20 tiles stitched
together, and the residual error is largely *per tile* (beam-shift / stage
error, and tile-dependent specimen height). This script measures and corrects a
2D shift per (tilt, tile) instead of per tilt.

Output is an updated `Refined_positions` in a copy of the per-tilt HDF5, so
`stitch.py --positionfile` can re-stitch and the whole thing iterates.

All intensive steps run as slurm jobs. Nothing heavier than reading a header
runs on the login node.

## Stage status at a glance

| Stage | State | Evidence |
|---|---|---|
| **v0** geometry / conventions | **DONE, verified** | `rot-1 shift-1` @ +0.9105, unanimous over 3 tilts (job 28131022) |
| **tilt sign** | **DONE, derived, re-audited** | read off AreTomo's kernels + the `clip rotx` depth flip; matches a literal transcription at +0.96..+0.99. Citations re-read and the rotx flip re-run independently — see "Independent audit" |
| **v1** static reference | **DONE, re-measured on real data** | median residual 9.93 px = 272 A over the 20 un-clipped measurements (14.57 px over all 36), tile rejection halved to 36% (job 28219393). Two defects found in the process: whole-pixel quantisation, and no handling of ROI-clipped tiles |
| **v2** leave-one-out recon | not started — **now the open action** | required for correctness, not speed — see v1 caveat. Fold in the ROI fraction and the stable tile identity here |
| **tile identity across tilts** | **BROKEN, must fix before v3** | `Refined_positions` is renumbered per tilt; 19 of 20 tiles change index between tilt 0 and +3 |
| **v3** global solve + outer loop | not started | — |

Scripts: `04_AreTomo_nopatch.sh` (alignment + aligned stack) ->
`06_AreTomo_recon_bin4.sh` (reconstruction) -> `05_projmatch_v0.sh` (conventions)
-> `07_projmatch_v1.sh` (per-tile shifts).

## STATUS — where to pick this up

Last updated 2026-07-28.

Committed to git as of 2026-07-28 (`878b06f`, the script and this document
only), local commits, not pushed.

**Newest first:** v1 has been re-run through the fixed reprojection
(job 28219393) and its numbers are now real. The reprojection fixes bought
exactly what they should have: tile rejection halved and the residuals are
physically sized and structured. See "v1 re-run" immediately below.

### The single next action

**v2 — leave-one-out reconstruction.** Everything before it is settled and
measured. v1's magnitudes still cannot be read as residuals because its
reference contains the tile being aligned; v2 is what makes the numbers mean
something.

Four items found in the re-run should be folded into v2 rather than patched
into v1, in rough order of how much damage they do if left:

1. **Stable tile identity across tilts.** Currently broken and fatal to v3.
2. **ROI-clipped tiles**, which are neither detected nor down-weighted and are
   inflating the residuals by ~2.3x.
3. **Whole-pixel quantisation** — `--upsample` does nothing on the masked path.
4. The residual 36% rejection, much of which is item 2.

### v1 re-run (2026-07-28, job 28219393, 4m23s) — the fixes hold up

Run without `--settle-tilt-sign`, which is now only a wrong-sign control and
whose median-|shift| verdict rewards degenerate zeros. Clean exit, no warnings.

| | before (28131893, broken) | after (28219393) |
|---|---|---|
| tiles rejected | 39/56 = **70%** | 20/56 = **36%** |
| measurements | 17 | 36 |
| median \|shift\| | 1.00 px (stale) | **14.57 px = 399 A** |
| median weight | 0.416 | 0.448 |

Per check tilt: -3 deg 7.07 px (11 tiles), 0 deg 17.37 px (12), +3 deg 21.93 px
(13). Rejections 6/17, 8/20, 6/19.

**The rejection rate was mostly the broken reprojection, as suspected — but not
entirely.** Halving it confirms the diagnosis. Much of the remaining 36% is ROI
clipping — see the next section, which was written after this table and
supersedes the reading of these numbers.

**The residuals are anisotropic.** Perpendicular to the tilt axis the median is
7.0 px against 2.0 px along it, a 3.5x asymmetry on the 20 measurements from
un-clipped tiles. That is the signature the height term in the global solve is
built for: a tile at height `z_t` displaces by `z_t sin(alpha)` perpendicular to
the tilt axis and not at all along it. It is *suggestive only* at three check
tilts spanning 9-15 deg of effective angle. **The test is cheap and worth
doing:** re-run with `--n-check-tilts` large or `--check-tilt-strategy high` and
confirm the perpendicular component scales as `sin(alpha)` while the parallel
one does not. If it does, the height model is real geometry and not a fitted
convenience.

**But treat 14.57 px as an upper bound, not the result.** Restricted to tiles
wholly inside the ROI it is 9.93 px = 272 A; the ROI-clipped tiles median
23.17 px and drag the headline up. The 9.93 px figure is the one to quote.

### ROI clipping is not handled, and it is distorting the v1 numbers

`--ROI -5500 12000 -6000 12000` in `02_stitch.sh` crops the canvas below the
natural extent of the montage, so edge tiles lose real area off the sides. This
was already noted under "Correlation target: two modes" as an aside about mode
B. It is worse than an aside: **16 of the 36 v1 measurements come from tiles
that are not wholly inside the ROI**, and they behave measurably differently.

Per-tile footprints computed from `Refined_positions` against the 9000 x 8750
canvas, for the three check tilts:

| | n | median \|shift\| | median weight |
|---|---|---|---|
| wholly inside the ROI | 20 | **9.93 px** | 0.478 |
| clipped by the ROI | 16 | **23.17 px** | 0.399 |

Clipped tiles measure 2.3x larger shifts at lower confidence. The mechanism is
plain: masked NCC with a truncated mask has no content on the clipped side, so
the peak slides toward the surviving content, biasing the shift away from the
cut edge. Five of the eight largest shifts in the whole run are clipped tiles
with `w < 0.5`. Clipping runs to 24.9% of a tile still inside at tilt 0.

The correlation with the *fraction* inside is weak (Pearson -0.22 against
|shift|, +0.27 against weight) because the effect is closer to a threshold than
a gradient, and because one genuine large shift — tilt 0 tile 3, 36.2 px at
`w = 0.802`, the highest weight in the run — is a wholly-inside tile. So
clipping is not the whole story, but it is a large part of it.

It also accounts for much of the residual rejection: **six of the seven tiles
below 50% inside the ROI produced no measurement at all**, either skipped or
over `--max-shift`.

**Does the code check for this? Not really.** `measure_tile_shifts` has two
absolute size floors — `min_px = 5000` canvas px, then
`n_px < min_px // recon_bin**2` or `n_px < 200` in the aligned frame. Against a
full on-canvas tile of 5,892,480 px, 5000 px is 0.08%, so they only catch a tile
that has essentially vanished. Three deeper problems:

- **The ROI is never read.** Everything is derived from `index_map`, which is
  *already* both ROI-cropped and Voronoi-partitioned, so a tile truncated by the
  ROI and a tile carved down by `clip_masks_to_overlaps` are indistinguishable —
  both are simply a smaller mask. The two need separating: the Voronoi sliver is
  still valid data, the truncated footprint is a biased measurement.
- **The skips are `logger.debug`,** so tiles vanish from the INFO log without
  explanation and the counts do not reconcile — 18/20/19 tiles in, 17/20/19
  candidates out, with nothing said about the missing one.
- **Nothing down-weights a truncated tile.** It is measured as if whole.

**What to do,** in v2 where the measurement is rebuilt anyway: compute each
tile's full footprint from `Refined_positions` and the ROI (as above, it is
arithmetic on a 20 x 2 array — no need to touch `index_map`), and carry the
in-ROI fraction as a per-tile quantity. Then either fold it into the quality
weight, or reject below a threshold — and log the reason at INFO either way.
Mode B has the same issue and no `index_map` to hide it in.

### Tile indices are not stable across tilts — this breaks the v3 global solve

Found while checking the above, and more serious than the ROI itself.

`Refined_positions` has a **different length at different tilts**: 18 at -3 deg,
20 at 0 deg, 19 at +3 deg. The montage translates as it tilts, tiles fall
outside the ROI, and the survivors are written out renumbered `0..N-1`. Matching
tilt 0 to tilt +3 by position after removing the median montage offset
(row -67, col +455 px; tile pitch is ~1815 px, so the match is unambiguous at
~310 px residual):

    tilt 0 index  3  4  5  6  7 ... 19
    tilt+3 index  1  2  3  4  5 ... 17

**19 of 20 tiles carry a different index at the two tilts.** Tile `t` is simply
not the same physical tile from one tilt to the next.

For v1 this is harmless — it reports per tilt, and each HDF5's indices are
internally consistent, so writing `Refined_positions` back is also safe. For
**v3 it is fatal**. The global solve is written over `D(i, t)` with `t` a
physical tile: the height model fits `(a_t, z_t)` per tile across all tilts, the
seam term uses `overlaps` from one tilt's HDF5, and the gauge constraint sums
over `t` at fixed `i`. Every one of those silently mixes different physical
tiles. It would not crash; it would return a plausible, wrong answer, and the
"check fitted `z_t` varies smoothly across the montage" validation would fail
for a reason nobody would guess.

**The fix is available and cheap.** `Montage_imageshifts_9-A.txt` holds the 20
nominal beam-shift positions (header `41 / 0.0 / 20`, then `x y z` in unbinned
px) and is the canonical tile identity. Match each tilt's `Refined_positions` to
it by nearest neighbour after removing the per-tilt median offset, and key
everything on the nominal index. Do this in v2, before any cross-tilt
aggregation exists to be wrong.

### Found in the re-run: the `--upsample` flag is a no-op

Every one of the 36 reported shifts is an exact integer. `measure_tile_shifts`
passes `upsample_factor=upsample` (default 10) to
`skimage.registration.phase_cross_correlation`, but **skimage discards it
whenever a mask is supplied** — verified in the installed 0.24.0, where the
masked branch calls `_masked_phase_cross_correlation(...)` with neither
`upsample_factor` nor `space` and returns immediately. Padfield masked NCC has
no subpixel path in skimage.

So the measurement is quantised to one recon pixel = **27.4 A** at
`--recon-bin 4`. Against a 399 A median that is ~7%, tolerable for v1's
report-only role, but it is a hard floor on the whole method and the plan's
Algorithm section promises a "subpixel peak" that is not being computed. Fix
before v3, either by fitting a parabola to the masked NCC surface around the
integer peak, or in the torch port, where `GMnccXcf.cu` gives the full
correlation surface and a 3-point fit is a few lines. Until then, drop the
misleading `--upsample` flag or make it raise.

### Disk quota is no longer a constraint

`punim1452` is at 6479/8002 GB (80%), ~1.5 TB free, against "100%, 46 GB free"
when the bin-2 reconstruction failed. The 78.8 GB bin-2 volume is now
affordable, which would cut the quantisation above from 27.4 A to 13.7 A. Not
urgent — v2 rebuilds the reference anyway — but the constraint that forced
bin 4 is gone. Leftovers from that failure are still in `AreTomo_nopatch/`:
1024-byte stubs `recon_nopatch_bin2.mrc`, `toberotated_proj{XY,XZ}.mrc`.

### Independent audit of the tilt-sign work (2026-07-26, later session)

The derivation was re-checked from scratch against `~/gitprojects/AreTomo2`
rather than taken from these notes. It holds. Recorded so it does not get
re-litigated:

- Every source citation in `reproject_volume`'s comment block is accurate:
  `CTomoWbp.cpp:45-52` (cos/sin built straight from `pfTilts`, no negation),
  `CAlignFile.cpp:165-169` (that same `fTilt` written to the `TILT` column),
  `CProcessThread.cpp:505-515` (`GetFrame(iEndOldY - y)` reverses depth),
  `CSaveXF.cpp:60-70` (`A = R(-ROT)`, `d = A @ (-shift)`, i.e. `rot-1 shift-1`).
- `GForProj.cu:25-37` was re-derived independently and does collapse to
  `x' = xp/cos - tan·z'`. The shear is real, not a reinterpretation.
- The `clip rotx` depth flip was **re-run**, not trusted: a labelled 4x3x5
  volume in `(x, z, y)` order came back with output depth slices 0/1/2 holding
  internal `z` = 2/1/0. Confirms `z_disk = nz-1-z_internal`, and therefore the
  `+tan` in the shear matrix.
- Sign plumbing is consistent end to end: the rotx flip lives in the shear
  matrix, `run_measure` passes `.aln` `TILT` verbatim, and `--settle-tilt-sign`
  flips it to produce the wrong-sign control. No double negation.

### Fixed in that audit

- **`run_measure`'s docstring was stale** — it still described the abandoned
  procedure of reprojecting at both signs and keeping the smaller residual, the
  exact method the rest of the file documents as degenerate. Anyone reading that
  function first got the pre-change story. Rewritten.
- **`--measure` now requires `--canvas-stack`.** It previously crashed with a
  `TypeError` on `None` several minutes in, after the reprojection. v0 can fall
  back to the coverage mask; v1 has no fallback, so it fails fast instead.

### Repo / working-tree divergence — worth tidying

The slurm scripts this document names live only in the dataset's
`submission_scripts/` directory, not in the repo. `slurm_templates/` stops at
`04_AreTomo.sh`, so `04_AreTomo_nopatch.sh`, `05_projmatch_v0.sh`,
`06_AreTomo_recon_bin4.sh` and `07_projmatch_v1.sh` exist for job 28131022 et al.
but not for anyone reading the repo. `refine_montage_projmatch.py:155` and `:768`
point users at `04_AreTomo_nopatch.sh` by name.

**v0 is DONE and the convention is settled.** Job 28131022 (3m50s) returned

    WINNER: rot-1 shift-1   mean correlation +0.9105

unanimous across all three check tilts (+0.904, +0.916, +0.912), margin 0.095
over the runner-up, no warnings. Final warped-canvas vs aligned-stack checks
were +0.9045 / +0.9158 / +0.9121.

`rot-1 shift-1` is the **canonical** form taken from AreTomo's own `.xf` writer
(`AreTomo2/ImodUtil/CSaveXF.cpp:60-70`) — `A = R(-ROT)`, `d = A @ (-[TX,TY])`.
It is now the hard-coded default in `Convention`. There is **no 180 deg
rotation, no reflection and no TY flip**; the coordinate chain was right from
the beginning.

### What actually went wrong, and the lesson

Three successive searches returned pure noise (8 variants at high tilt, then 32
at high tilt, then 32 at low tilt; best correlations +0.0031, +0.0065). The
cause was **not** geometry. `_check_one_tilt` was correlating `canvas_proxy`, a
*binary tile-coverage mask* built from `index_map`, rather than image content.

Measured: 99.92% of the pixels the correlation scores are exactly 1.0. The
vector is constant, std 0.015 against the aligned slice's 2.68, so Pearson r is
structurally ~0 for **every** convention regardless of the geometry. The only
variance came from anti-aliased edge pixels after interpolation — which is
precisely the +/-0.01 band that kept appearing.

Swapping in the inpainted montage stack, the real input AreTomo consumed, moved
the winner from +0.0065 to +0.9105 with no other change.

**The lesson, for the next time this workflow reports something inconclusive:**
a flat, near-zero spread across *all* hypotheses is not a weak signal, it is a
broken measurement. Check what is being compared before enlarging the space of
how it is transformed. Two rounds were spent extending the search — the tell was
there in the first result.

### Fixed along the way

- **`--canvas-stack`** (new, and the fix that mattered). Supplies the montage
  stack AreTomo consumed, `stitched_motioncorr/Montage_9-A_inpainted.mrc`, one
  slice per tilt on the canvas indexed by **SEC**. Without it the script falls
  back to the coverage mask and now warns loudly that it cannot discriminate.
- **`select_check_tilts(strategy=...)`**, defaulting to **`low`** — tilts nearest
  zero, `(-3, 0, +3)`. The high-tilt frames on 9-A really are too dark to use
  (+54 and +60 overlay PNGs are 392/457 KB against 1.0 MB for -57), so this
  change stands on its own merits even though it was not the bug.
  `--check-tilt-strategy high` restores the old behaviour.
- **`Convention.rot_offset`** (degrees added to `ROT`; 180 flips the tilt axis)
  and **`Convention.flip_rows`** (mirror the row axis about the canvas centre).
  These closed a real gap — all 16 pre-existing variants had `det = +0.25`, i.e.
  they were *all* proper rotations, so no reflection was reachable by any
  combination of `rot_sign`, `rot_offset`, `shift_sign` and `transpose_xy`.
- **`Convention.flip_ty`** — negates the TY shift component alone. An
  MRC(y-up) vs numpy(row-down) *relabelling* conjugates the matrix to
  `R(-theta)` (already reachable via `rot_sign`) but sends `b -> F@b`, flipping
  TY and leaving TX. `shift_sign` flips both components and `transpose_xy` swaps
  them, so neither could express it.
- Transform **dedup** in `search_conventions`. All 64 flag combinations turn out
  to be distinct on the current axes, so it is a guard rather than a filter.

The flip axes all proved unnecessary here — `flipTY` variants surface as
runner-ups around +0.81 — but they were worth adding to rule the hypothesis out
rather than argue about it, and they cost one line each in the search.

### v0's remaining loose end — now closed

The **sign of the reprojection angle** is the one convention v0 structurally
cannot resolve, because the tilt angle never enters the 2D alignment transform —
only the reprojection. It was handed to v1, which failed to settle it
empirically. It has now been settled by **reading it off AreTomo's source**
instead of searching for it; see the next section.

## Tilt sign convention, taken from AreTomo (2026-07-26) — SETTLED

Not measured, derived, and then checked against a literal transcription of the
kernel. Two independent facts combine.

**1. AreTomo does not negate the tilt angle anywhere.**
`Recon/GBackProj.cu:31-33` backprojects by sampling the sinogram at

    xp = x*cos(TILT) + z*sin(TILT)

with `x`, `z` centred on the volume. `TILT` comes straight from `CAlignParam`
(`Recon/CTomoWbp.cpp:47-52`), which is the same array `CAlignFile` writes to the
`.aln` `TILT` column (`MrcUtil/CAlignFile.cpp:165-169`). `Recon/GForProj.cu:25-37`
inverts the same relation. So the angle to use is the `.aln` `TILT` as written,
unmodified — consistent with gotcha 3, where the `-TiltCor` offset is already
baked into that column.

**2. `clip rotx` reverses the depth axis.** Verified directly against IMOD's
`clip` on a labelled 4x3x5 test volume: output depth index `k` maps to input
internal `z = nz-1-k`. Re-run in a later session and reproduced exactly. That is
exactly what AreTomo's own `-FlipVol` does —
`CProcessThread.cpp:505-515` writes `pfDstFrm = GetFrame(iEndOldY - y)`. X and Y
directions are preserved. So the two routes to an xyz-view volume agree, and
both flip z relative to the reconstruction's internal frame.

Substituting `z_disk = -z_internal` into AreTomo's ray gives the same geometry
evaluated at **minus** the tilt angle. Hence, for a rotx'd volume:

    reproject at -TILT, where TILT is the .aln column verbatim

The negation is the rotx flip, not a free parameter. `--settle-tilt-sign` is
demoted to a control: it now reprojects at the deliberately wrong sign, and a
"win" for that sign means the *measurement* is broken, not the convention.

### Found while verifying: the reprojection was broken at high tilt

Checking the sign against a transcription of `Recon/GForProj.cu` exposed a
second and much larger error in `reproject_volume`, unrelated to signs.

The old implementation rotated the volume in the (z, x) plane inside its own
`nz`-tall array and summed over z — the obvious reading of "rotate then sum".
AreTomo does not do that. `GForProj.cu:28` starts each ray at
`fZStartp = -fXp*fSin/fCos - 0.5*iRayLength`, which **re-centres the sampled
span per output column** onto where that column's ray actually crosses the slab,
and runs it for `iRayLength = VolZ/cos` rather than `VolZ`. Expanding that
offset collapses the whole operation to a **shear**, not a rotation:

    x = xp/cos(TILT) - tan(TILT)*z        z = the slab's own z planes

Rotating instead discards every ray whose crossing point leaves `|z| < nz/2`.
For this canvas (`nz = 150`, `nx = 2186` at bin 4) that is nearly the entire
field: measured recovered intensity for a point source at increasing `|x|`,

| tilt | 0 px | 50 | 100 | 150 | 200 | 250 |
|---|---|---|---|---|---|---|
| 0 deg | 1.00 | 1.00 | 1.00 | 1.00 | 1.00 | 1.00 |
| 30 deg | 1.02 | 0.91 | 1.05 | 1.02 | **0.00** | **0.00** |
| 45 deg | 0.93 | 1.00 | 1.15 | **0.00** | **0.00** | **0.00** |
| 60 deg | 0.98 | 1.04 | **0.00** | **0.00** | **0.00** | **0.00** |

At 60 deg only `|x| < ~87 px` of a 1093 px half-width survived, so the reference
was essentially blank away from the tilt axis. This is very likely the real
cause of the ~70% tile rejection rate noted below, and it also explains why
`--settle-tilt-sign` was degenerate: with a mostly-empty reference, neither sign
has anything to correlate against.

`reproject_volume` now implements the shear, samples one slab plane per output
row (so the depth axis is not interpolated at all), and normalises by the per-ray
in-volume sample count to match `GForProj.cu:47-49`. Against a literal
nearest-neighbour transcription of the kernel, on a smooth random volume:

| tilt | new, derived sign | new, wrong sign | old rotate-and-sum |
|---|---|---|---|
| +20 | **+0.967** | +0.230 | +0.661 |
| +45 | **+0.957** | +0.100 | +0.425 |
| +60 | **+0.990** | +0.345 | +0.360 |
| -60 | **+0.993** | +0.455 | +0.332 |

The residual gap from 1.0 is nearest- versus linear-interpolation, and the
non-zero column coverage now matches the kernel exactly. Note the old code
scored only +0.33..+0.66 *even with the correct sign* — which is why the sign
could not be settled empirically before this was fixed.

**A half-pixel discrepancy that is not one.** `reproject_volume` centres both
input and output on `(n-1)/2`, where `GForProj.cu:26-27` measures the output
column from `(nx-1)/2` but adds it to an input centre of `nx*0.5` — apparently a
half-pixel inconsistency, and at `--recon-bin 4` a uniform 0.5 px bias would be
13.7 A, comparable to the residuals being measured. It cancels: the kernel
samples with `gfVol[... + (int)fX]`, and truncation puts the effective sample at
`fX - 0.5` in cell-centre coordinates. The two agree exactly. Do not "fix" this
by moving to `n/2` — that would introduce the bias rather than remove it.

**Consequence: v1's numbers were stale.** Job 28131893's residuals were measured
through the broken reprojection and should be discarded, not merely distrusted.
Superseded by job 28219393 — see "v1 re-run" in STATUS.

### v1 status (2026-07-26) — the superseded run, kept for the record

Reconstruction: `06_AreTomo_recon_bin4.sh` (job 28131468, 65 s) wrote
`recon_nopatch_bin4.mrc`, 2.95 GB, shape (150, 2250, 2186), thin axis first so
`clip rotx` worked. The bin-2 attempt in `04` had failed with *Disk quota
exceeded*, leaving a 1024-byte header — /data/gpfs is at 100% (46 GB free) and a
bin-2 volume of this canvas is 78.8 GB, so it could never have been written.

v1 itself (`07_projmatch_v1.sh`, job 28131893, 6m46s) ran to completion. Two
findings, one good and one not:

**Fixed: the masked NCC search was unbounded.** The first run returned median
residuals of **1101 px = 3.0 um** against a tile that is only 1.97 x 1.40 um —
shifts larger than the tile itself. Padfield masked NCC normalises by overlap
area, so as displacement grows and overlap shrinks the score is *inflated*; an
unbounded search is therefore biased towards large, small-overlap matches.
Cropping the correlation to the tile footprint plus `--max-shift` (default 200
canvas px) dropped the median to **1.00 px = 27.4 A**, a physically sensible
per-tile error.

**Not fixed: `--settle-tilt-sign` returned a degenerate answer.** It reported
`TILT SIGN: - wins (0.00 vs 1.00 px, ratio 1e9)`. That is not a measurement:

- the `-1` sign produced *fewer* surviving measurements (9 vs 17), because more
  tiles were rejected, and its median collapsed to exactly 0.00 px;
- comparing median |shift| therefore **rewards degenerate zeros** — whichever
  sign fails on more tiles looks better;
- the 1e9 ratio is just the `max(min, 1e-9)` guard dividing by zero;
- median *weight* actually favours the opposite sign (+1 at 0.416 vs -1 at
  0.384), i.e. the two criteria disagree.

**Superseded.** The sign is no longer v1's job — it has been derived from
AreTomo's source, see the tilt sign section above. The degeneracy is now
explained: the reprojection was blank away from the tilt axis, so neither sign
had anything to correlate against.

**Checked on the re-run:**

1. Roughly 70% of tiles were rejected (11/17, 14/20, 14/19). The prime suspect
   was the broken reprojection rather than `--max-shift 200` being too tight or
   mode A's Voronoi slivers being too small. **Confirmed, partially:** fixing
   the reprojection took it to 36% (6/17, 8/20, 6/19). The remainder is still
   unexplained and the other two suspects survive.
2. If a sign comparison is ever wanted again, score by **correlation weight over
   a common set of tiles**, not median |shift|, which rewards degenerate zeros.
   `07_projmatch_v1.sh` no longer passes `--settle-tilt-sign` at all.

**A caveat that may make this moot.** The near-zero residuals are exactly what
the Algorithm section warns about: *"Reprojecting a volume that contains the
tile being aligned is self-reinforcing — it will measure ~zero shift and report
success."* v1's static reference contains every tile, so ~0 is the built-in
answer and v1's magnitudes cannot be trusted as residuals. That was always
understood — v1 is "report only" for this reason — but it also means the tilt
sign may simply not be resolvable from a self-inclusive reference, and the
question may have to carry over to v2's leave-one-out reconstruction.

**Already verified** (no job needed — these ran interactively):

- Canvas rebuilt from `02_stitch.sh` arguments reproduces `index_map`
  (9000, 8750) *exactly*, at 6.852 A/px with tile-on-canvas (2880, 2046).
- `.aln` parses to 37 sections + 4 dark = 41.
- The SEC join is exact: `-60 + 3*SEC == TILT - 12` for all 37 sections,
  and the +12 deg `-TiltCor` offset is inferred automatically with zero spread.
- Mode A footprint loss measured: 56.1% mean, 13.3% worst tile.

**Formerly open, now closed:** the *sign* of the reprojection angle relative to
the reconstruction. Neither v0 nor v1 could resolve it empirically, so it was
read off AreTomo's source instead — see the tilt sign section above. The answer
is `-TILT` for a rotx'd volume, and the reason the empirical attempts failed
turned out to be a broken reprojection rather than a lack of leverage.

## Inputs (dataset 9-A, resolved)

| Input | Path | Contents used |
|---|---|---|
| Per-tilt positions | `stitched_motioncorr/Montage_9-A_9-A_<angle>_refined_positions.h5` | `Refined_positions` (20,2) µm, `index_map` (9000,8750) uint16, `overlaps` (44,2) |
| Raw tiles | `motion_corrected/Montage_9-A_9-A_<angle>.mrc` | 5760 x 4092 x 20, apix 3.426 A. 41 files, one per tilt |
| Initial shifts | `20260713_YeastLamella/Montage_imageshifts_9-A.txt` | header `41 / 0.0 / 20`, then 20 rows of `x y z` in unbinned px |
| AreTomo alignment | `AreTomo_nopatch/Montage_9-A_inpainted.aln` | `ROT GMAG TX TY TILT` per section + `# DarkFrame` list |
| Aligned stack | `AreTomo_nopatch/aligned_tiltseries_bin2.mrc` | AreTomo's own aligned montage — v0 ground truth |
| Reference volume | `AreTomo_nopatch/recon_nopatch_bin4.mrc` | v1 static reprojection source. **bin 4, not 2** — (150, 2250, 2186), 2.95 GB, from `06_AreTomo_recon_bin4.sh`. The bin-2 volume `04` tried to write is 78.8 GB and failed on quota |
| Stitch parameters | `submission_scripts/02_stitch.sh` | `PIXEL_SIZE=3.426`, `BINNING=2`, `--rotate 90`, `--ROI -5500 12000 -6000 12000`, `-f 80`, `-mt 0.5`, `-T reference.mrc` |

## Coordinate chain

Positions are in µm, ordered `(x=column, y=row)` — opposite to numpy. From
`stitch.py:354-460`, tile `t` lands at canvas pixel offset

    col0 = (pos[t,0]*1e4 - origin[0]) / pixel_size / binning
    row0 = (pos[t,1]*1e4 - origin[1]) / pixel_size / binning

with, from `stitch.py:1952-1955`, ROI given in *unbinned tile pixels*:

    origin = (roi[0], roi[2]) * pixel_size          = (-5500, -6000) * 3.426 A
    width  = (roi[1]-roi[0], roi[3]-roi[2]) * pixel_size
    canvas = width / pixel_size / binning           = 8750 x 9000  (checks out)

`origin` and `width` are shared across tilts, so canvas coordinates are directly
comparable between tilts.

AreTomo then applies, per section: rotation by `ROT` about the image centre
(bringing the tilt axis onto y), scale `GMAG`, translation `(TX, TY)`. Call this
`A_i`. A residual `m` measured in the aligned frame maps back to a position
update as

    dpos_µm = R(-ROT_i) @ m * pixel_size * binning / 1e4

with the convention pinned down empirically in v0 as `rot-1 shift-1`.

In code, `measure_tile_shifts` does **not** rebuild `R(-ROT)` by hand — it
inverts the very same `alignment_matrix()` used for the forward warp. That makes
the round trip self-consistent by construction and removes any opportunity for a
fresh sign error on the way back. Note the matrix must be built with
`out_bin = recon_bin`, so the inverse lands directly in canvas pixels.

**Patch/local alignment is deliberately not modelled.** The reference is
produced by `04_AreTomo_nopatch.sh` with `-Patch` omitted. Per-tile shifts *are*
the local alignment, parameterised physically by tile rather than by an
arbitrary patch grid; modelling both would double-correct.

## Pixel size ladder

The workflow bins repeatedly, so four different pixel sizes are in play.
Verified with `header` on 2026-07-25, extended 2026-07-26 for the bin-4 volume.

| Stage | File | apix (A) | Header trustworthy? |
|---|---|---|---|
| Motion-corrected tiles | `motion_corrected/Montage_9-A_9-A_<a>.mrc` | 3.426 | yes |
| Montage canvas (stitch `BINNING=2`) | `stitched_motioncorr/Montage_9-A.mrc` | 6.852 | yes |
| Montage, inpainted | `stitched_motioncorr/Montage_9-A_inpainted.mrc` | **1.000** | **NO — stripped** |
| AreTomo aligned stack (`-OutBin 2`) | `aligned_tiltseries_bin2.mrc` | 13.704 | labelled 13.74 |
| AreTomo reconstruction (`-OutBin 4`) | `recon_nopatch_bin4.mrc` | 27.408 | not checked |

Two traps here:

- The **inpainted** stack — the one AreTomo actually consumes — has had its
  pixel size reset to 1.0 by `inpaint_apply.py`. The pipeline only works
  because `-PixSize` is passed explicitly on the AreTomo command line. Worth
  fixing in `inpaint_apply.py` separately.
- **The aligned stack and the reconstruction are at *different* binnings**, 2
  and 4, and the code takes both separately as `--out-bin` and `--recon-bin`.
  Conflating them puts a factor of 2 into every measured shift, which would look
  like a plausible-but-wrong residual rather than an obvious failure. The
  reconstruction is coarser on purpose: at bin 2 this canvas gives a 78.8 GB
  volume, against 46 GB free.
- `03_AreTomo_motioncorr.sh` passes `-PixSize 6.87` where the true value is
  6.852, a 0.26% scale error baked into every downstream header.
  `04_AreTomo_nopatch.sh` uses 6.852. The difference is cosmetic for this work
  because alignment is pixel-based, but it is a good reason to
  **work in pixels throughout and never round-trip through Angstroms**.

Critically, `.aln` `TX`/`TY` are in **input-stack pixels**, i.e. montage canvas
pixels at 6.852 A — confirmed by `# RawSize = 8750 9000 41` matching the canvas.
The reference volume and aligned stack are `-OutBin` coarser, so a shift
measured against them scales by `out_bin` to reach canvas pixels. This is the
`out_bin` argument threaded through `alignment_matrix()`.

## Gotchas found while reading the existing pipeline

These are the things most likely to burn a day each.

1. **`binned_pixel_size` in the HDF5 is a misnomer.** `stitch.py:581` stores
   `pixel_size / binning` = 1.713 A, which is neither the tile pixel size nor
   the canvas pixel size — the canvas is `pixel_size * binning` = 6.852 A, a
   factor of 4 out. Do not use this field. `refine_montage_projmatch.py`
   ignores it and derives the canvas from the `02_stitch.sh` arguments instead,
   cross-checking against `index_map.shape`.
   `Refine_montage_from_aretomo.py:139` reads it as `pixel_size`, which looks
   like a latent bug in that script worth checking separately.

2. **`--rotate 90` is in the stitch command.** Every tile is `np.rot90(k=1)`'d
   before placement (`stitch.py:834`), and tile dimensions are swapped
   accordingly (`stitch.py:714`). Mode B must reproduce this exactly: the raw
   tile is (4092, 5760) in numpy order and becomes (5760, 4092) on the canvas.

3. **Join `.aln` sections to tilts by `SEC`, never by tilt angle.** This is the
   nastiest one, and an earlier draft of this plan had it exactly backwards.

   `-TiltCor 1 12` supplies a fixed 12 deg offset (lamella pre-tilt), and
   AreTomo writes the *corrected* angles into the `.aln`. So the `TILT` column
   runs **-45..+72** while the actual data runs **-60..+60** — a uniform +12
   deg shift. Matching on angle therefore pairs each tilt with the section
   12/3 = **4 positions away**, and every measurement comes out wrong while
   looking entirely plausible.

   The reliable key is `SEC`, the 0-based raw frame index, because `stitch.py`
   writes z-slices in ascending tilt order (`stitch.py:1982`):

       SEC == rank of the tilt angle among all 41 tilts, sorted ascending

   Verified exactly — `-60 + 3*SEC == TILT - 12` for all 37 sections, zero
   mismatches. `map_sections_to_tilts()` does this join and *infers* the offset
   rather than trusting a flag, so a different `-TiltCor` on another dataset is
   detected automatically. It errors if the inferred offsets are inconsistent.

   Consequence: `--tilt-offset` defaults to **0**, because the offset is
   already in `TILT`. Reprojection uses `TILT` as written.

4. **Dark frames: trust the indices, not the angles.** Four frames are dark,
   `# DarkFrame` indices `{0, 34, 37, 39}` — exactly the `SEC` values missing
   from the table (41 - 4 = 37). The *angles* on those lines are inconsistent:
   index 0 reports -48 (corrected) while index 34 reports 42 (nominal). Use the
   indices only.

   Note the aligned tilt series has 37 slices, not 41, so its z-index is the
   position among surviving sections in `SEC` order — a third distinct
   indexing.

6. **Tile indices are per-tilt, not physical.** `Refined_positions` holds only
   the tiles that survived stitching at that tilt, renumbered from zero, so its
   length varies (18/20/19 across the three check tilts on 9-A) and tile `t` is
   a different physical tile at different tilts. Safe within one tilt, fatal for
   anything that aggregates across tilts. Key on the nominal beam-shift index
   from `Montage_imageshifts_*.txt` instead. See the STATUS section.

7. **The ROI truncates tiles, and `index_map` hides it.** By the time a tile
   reaches `index_map` it has been both ROI-cropped and Voronoi-partitioned, and
   the two are indistinguishable. A truncated footprint biases masked NCC toward
   the surviving content. Compute the in-ROI fraction from `Refined_positions`
   and the ROI arguments, not from the mask.

5. **AreTomo volume output is (x, z, y)** and the pipeline runs `clip rotx` to
   get standard order. Use the rotx'd file and assert the axis order.

## Algorithm

Per tilt `i`, walking outward from the zero-tilt section (AreTomo's ordering in
`CProjAlignMain::mDoAll`), skipping dark frames:

1. **Leave-one-out reconstruction.** Backproject every *other* aligned tilt.
   Because the tilt axis is along y after global alignment, each row y is an
   independent x-z problem — see `AreTomo2/ProjAlign/GReproj.cu:mGBackProj`.
   Restrict z to a slab of `VolZ`.
2. **Reproject** at `TILT_i + tilt_offset` -> reference `R_i`, aligned frame.
3. **Tile footprints.** Forward-transform each tile's mask through the stitch
   placement and `A_i` into the aligned frame.
4. **Correlate** per tile: masked NCC (Padfield) against `R_i` over the tile
   mask, subpixel peak -> residual `m(i,t)` + quality weight `w(i,t)`.
5. **Map back** to `dpos_µm` per the chain above.

Leave-one-out is not optional. Reprojecting a volume that contains the tile
being aligned is self-reinforcing — it will measure ~zero shift and report
success.

### Correlation target: two modes

- **Mode A (iterations 1..n-1)** — crop the stitched montage where
  `index_map == t`. Fast, no resampling, reuses exactly the pixels AreTomo saw.
- **Mode B (final iteration)** — reload tile `t` from
  `motion_corrected/Montage_9-A_9-A_<angle>.mrc`, `np.rot90` it, mask with
  `make_mask` (`Utilities.py:1349`) using the stitch settings `shrinkn`/`-f 80`
  fringe, `-mt 0.5` threshold, `reference.mrc` template, then transform into the
  aligned frame independently.

Mode B matters more than it first appears, and this is now measured rather than
argued. `clip_masks_to_overlaps` plus the overwrite at `indxmap[...][msk] = i`
makes `index_map` a *partition* of the canvas — interior tiles are carved down
to a Voronoi-like sliver, discarding precisely the overlap regions where SNR is
best. On tilt 0 of dataset 9-A, against a full on-canvas tile of
2880 x 2046 = 5,892,480 px:

    mean index_map region  = 3,305,373 px = 56.1% of the full tile
    worst tile             =   781,005 px = 13.3%
    uncovered canvas       = 16.1%

Separately, the ROI crops the canvas below the natural tile extent: footprints
span rows -1879..10027 and cols -1537..8884 against a 9000 x 8750 canvas, so
edge tiles lose real area off the sides. **This turned out to matter far more
than "separately" suggests** — it affects 16 of v1's 36 measurements and doubles
their apparent residual. See "ROI clipping is not handled" in STATUS. Mode B
does not escape it either: reloading the raw tile recovers the pixels, but they
still have no counterpart in the reference, which is only as wide as the canvas.

So mode A correlates on roughly half a tile and, for the worst tile, an eighth.
Mode B recovers the full tile for the final measurement. Expect the largest
gains on the interior tiles that mode A carves up hardest.

### Global solve

Rather than applying `m(i,t)` directly, solve for updates `D(i,t)` in R^2:

    minimise  sum_{i,t} w(i,t) ||D(i,t) - m(i,t)||^2                data
            + lambda sum_{(t,u) in overlaps} ||D(i,t) - D(i,u)||^2  seam consistency
            + mu sum_{i,t} ||D(i,t) - (a_t + z_t sin(alpha_i))||^2  height model
    s.t.      sum_t D(i,t) = 0   for every tilt i                   gauge fix

- **`t` must be a physical tile, not an HDF5 row index.** `Refined_positions`
  renumbers per tilt, so every sum over `t` here — the gauge constraint, the
  height model, the seam term — is over a different set at each `i` unless the
  indices are mapped to the nominal beam-shift grid first. See gotcha 6.
- **The gauge constraint is essential.** The mean shift per tilt is exactly
  degenerate with AreTomo's `TX/TY`. Without it the refinement fights the global
  alignment and the outer loop will not converge. Refine only the differential
  part. It also makes the result insensitive to the global `-TiltCor` offset.
- **The height term is nearly free and strongly stabilising.** A laterally
  mispositioned tile gives a tilt-independent offset; a tile at height `z_t`
  gives an offset proportional to `sin(alpha)`. Fitting `(a_t, z_t)` collapses
  ~37 noisy 2D measurements per tile onto 4 parameters.
- **The seam term** reuses `overlaps` from the HDF5 to stop independently
  refined neighbours from opening new seams.

Linear least squares, sparse, negligible cost.

### Outer loop

    refine -> write h5 -> stitch.py --positionfile -> AreTomo (no -Patch) -> repeat

2-3 rounds, mode A throughout, mode B on the last.

## Implementation: PyTorch, not C

The hot loop is `grid_sample` (backprojection and reprojection) and `rfft2`
(masked NCC), both cuFFT/cuDNN-backed. Hand-written CUDA buys perhaps 1.5-2x,
which does not justify the cost. No C required.

The row-independence trick makes backprojection a single batched `grid_sample`
per tilt with `ny` folded into the channel dimension: sample a `(1, ny, 1, nx)`
tensor with a `(1, nz, nx, 2)` grid, accumulate over ~37 tilts. At stitch
binning 2 plus a correlation binning of 4, `nx ~ 1100`, `nz ~ 250`; chunked over
y this sits comfortably on an A100.

`AreTomo2/ProjAlign/GMnccXcf.cu` in this fork already implements masked
normalised cross-correlation and is a direct port target;
`Utilities.py:1533 _masked_phase_cross_correlation` is the numpy equivalent.

Environment: `~/.conda/envs/miss-alignment` has torch 2.8+cu128, h5py, mrcfile,
scipy. (The `Stitch` env used by `02_stitch.sh` has no torch.)

## Stages

- **v0 — geometry only. DONE, verified.**
  `processing_scripts/refine_montage_projmatch.py`. Parses HDF5 + `.aln`,
  rebuilds the canvas, forward-transforms tile footprints into the aligned
  frame, overlays them on `aligned_tiltseries_bin2.mrc`. No correlation, no
  reconstruction.

  Because AreTomo emits its own aligned stack (`-VolZ 0`), v0 has *ground
  truth*, so the sign conventions are searched rather than guessed:
  `--search-conventions` scores all **64** rot/shift/swap/rot180/flip variants by
  correlating the warped canvas against AreTomo's aligned frame. The canonical
  form is taken from AreTomo's own `.xf` writer
  (`AreTomo2/ImodUtil/CSaveXF.cpp:60-70`): `A = R(-ROT)`, `d = A @ (-[TX,TY])`,
  giving `aligned = R(-ROT) @ (p - centre - [TX,TY]) + centre`. That fixes the
  maths but not how it lands in numpy's (row, col) ordering, hence the search.
  The `rot_offset`, `flip_rows` and `flip_ty` axes were added on 2026-07-26 to
  close a genuine gap — the original eight variants were all proper rotations,
  so no reflection was reachable. In the event none was needed: the canonical
  convention won once real image content was correlated instead of a coverage
  mask. They remain in the search as cheap insurance.

  *Verified:* canvas rebuilt from `02_stitch.sh` arguments reproduces
  `index_map` (9000, 8750) exactly; `.aln` parses to 37 sections + 4 dark = 41;
  tile-on-canvas shape (2880, 2046) after the `--rotate 90` axis swap.
  *Settled (job 28131022):* `rot-1 shift-1` at +0.9105, unanimous across
  three tilts. Hard-coded as the `Convention` default. See STATUS.

- **v1 — static reference. DONE, report only.**
  `--measure` reprojects `recon_nopatch_bin4.mrc` with `reproject_volume()` and
  measures per-tile shifts in mode A via `measure_tile_shifts()`, report only.
  Run by `07_projmatch_v1.sh`.

  Reprojection exploits the same row-independence AreTomo does: after global
  alignment the tilt axis is along y, so the whole operation acts in the (z, x)
  plane with y untouched.

  Reprojection is a **shear**, not a rotation — AreTomo re-centres each output
  column's ray on where it crosses the slab (`GForProj.cu:28`). Because y is
  untouched, chunking over y is *exact* rather than approximate — re-verified
  bit-identical at chunk sizes 7 and 1000 after the shear change — which keeps
  peak memory to one chunk instead of a second copy of a multi-GB volume. Depth
  is not interpolated at all: one slab plane per output row.

  Requires both `--recon` and `--canvas-stack`; it exits early without either.

  *Fixed 2026-07-26:* the tilt sign (`-TILT`) and the rotate-vs-shear error,
  both taken from AreTomo's source and since re-audited against it.
  *Re-measured 2026-07-28* (job 28219393): median residual 14.57 px = 399 A over
  36 measurements, rejection down from 70% to 36%, residuals anisotropic in a
  way consistent with the height model. Shifts are whole-pixel — skimage's
  masked NCC ignores `upsample_factor`. See STATUS.

- **v2 — leave-one-out reconstruction** in torch, replacing v1's static volume.
  Not an optimisation of v1 but a correctness requirement: a reference
  containing the tile being aligned measures ~zero shift and reports success,
  which is what v1 currently does. The tilt-axis sign is settled and no longer
  carries over.
- **v3 — global solve + outer iteration + mode B final pass.**

## Validation

- Inject known synthetic per-tile shifts into a copy of the HDF5, re-stitch, and
  confirm recovery to sub-pixel.
- Check fitted `z_t` against tile position — should vary smoothly across the
  montage if it is real specimen geometry rather than fitted noise. Note this
  check *would* have caught the tile-identity bug, but only as an unexplained
  failure; assert the identity mapping directly as well.
- Track seam residuals in overlap regions before/after.
- v1 shifts should be small and structured; large random shifts mean a
  convention bug upstream in v0.
