## Supplementary material for "Whole cryo-FIB lamella acquisition at high magnification with rectangular beam montage cryo-electron tomography": Source code: pyproject.toml

```
[build-system]
requires = ["setuptools>=61", "wheel"]
build-backend = "setuptools.build_meta"

[tool.setuptools]
packages = ["processing_scripts"]

[project]
name = "SquareBeamMontaging"
version = "1.0.0"
description = "A tool for stitching square beam montage tomography data."
authors = [{name = "HamishGBrown"}]
requires-python = ">=3.6"
license = {text = "MIT"}
classifiers = [
    "Programming Language :: Python :: 3",
    "License :: OSI Approved :: MIT License",
    "Operating System :: OS Independent",
]
dependencies = [
    "numpy>=1.20,<2.1",  # 2.0.x has a memmap int32 overflow bug; 2.1+ requires Python>=3.10
    "scipy",
    "networkx",
    "tqdm",
    "scikit-image",
    "pypng",
    "Pillow",
    "matplotlib",
    "mrcfile",
    "h5py>=3.11",
]

[project.scripts]
stitch_square_beam              = "processing_scripts.stitch:main"
stitch_by_adjacent_tilt         = "processing_scripts.stitchbyadjacenttilt:main"
choose_fringe_size              = "processing_scripts.Choose_fringe_size:main"
determine_overlap_fraction      = "processing_scripts.Determine_overlap_fraction:main"
make_gainref                    = "processing_scripts.make_gainref:main"
refine_montage_from_aretomo     = "processing_scripts.Refine_montage_from_aretomo:main"
refine_montage_from_imod        = "processing_scripts.Refine_montage_from_imod_fiducials:main"
generate_image_shifts           = "processing_scripts.Generate_Image_shifts_for_Montage:main"
join_to_mrc                     = "processing_scripts.JointoMRC:main"
masked_motioncorr               = "processing_scripts.masked_motioncorr:main"
mask_and_inpaint                = "processing_scripts.mask_and_inpaint:main"
inpaint_apply                   = "processing_scripts.inpaint_apply:main"
beam_mask_motioncorr            = "processing_scripts.beam_mask_motioncorr:main"
montage_provenance              = "processing_scripts.sidecar:main"
montage_backfill                = "processing_scripts.backfill_sidecars:main"
montage_aretomo_record          = "processing_scripts.aretomo_record:main"
mrc_to_gif                      = "processing_scripts.mrc_to_gif:main"
montage_to_relion               = "processing_scripts.montage_to_relion:main"
overlay_picks_on_montage        = "processing_scripts.overlay_picks_on_montage:main"
picks_to_imod_model             = "processing_scripts.picks_to_imod_model:main"
score_subtomo_sum               = "processing_scripts.score_subtomo_sum:main"
roi_editor                      = "processing_scripts.roi_editor:main"
resize_reference_beam           = "processing_scripts.resize_reference_beam:main"
```
