## Supplementary material for "Whole cryo-FIB lamella acquisition at high magnification with rectangular beam montage cryo-electron tomography": Source code: README.md

# StitchTomo

![StitchTomo logo](StitchTomo_logo.svg)


Code for setting up, executing (in SerialEM) and pre-processing cryo-TEM montages with round, square and rectangular beams. Draws substantial inspiration from https://github.com/apeck12/montage

# Install

```
git clone https://github.com/HamishGBrown/StitchTomo.git
cd StitchTomo
pip install -e .
```

# Overview

The full pipeline for montage tomography is:

1. **Acquisition** — collect montage tilt series in SerialEM
2. **Motion correction** — align raw frames per tile (`beam_mask_motioncorr`)
3. **Stitching** — assemble tiles into full montage tilt series (`stitch_square_beam`)
4. **Masking & inpainting** — [optional and only if necessary] isolate lamella signal and fill background (`mask_and_inpaint`)
5. **Reconstruction** — tomogram reconstruction in IMOD or AreTomo2/3

# Step 1: Collect in SerialEM

Full details are in `Montage tomography SOP.docx`. In brief:

1. In low-dose mode, acquire Navigator maps of your lamellae using the View preset. Avoid low mag due to image shifts between magnifications.
2. On an adjacent vacuum area, run the DeterminOverlapFraction.txt to work out optimal beam overlap (aim for max 10% in each direction), copy the values for beam rotation and overlap into the setupPolygonMontage.txt script.
2. Draw navigator polygons around each of the items to acquire.
3. Using the SerialEM "Acquire at items" feature, acquire viewmag images of each of the items of interest, tick the box to make navigator maps and set  the "SetupPolygonMontage.txt" script to run after each acquisition. Check that 'Imageshift*.txt" files have been generated for each navigator item.
3. Set each of the new view image maps to acquire and run `acquire_montage.txt` at those points using the SerialEM "Acquire at items" feature.
4. SerialEM collects one multi-frame TIFF per tile per tilt angle, saving them alongside a `.mdoc` metadata file.

A single tilt's acquisition looks like this:

![image](SingleMontage.gif)

---

# Step 2: Motion correction

Raw frames from each tile are motion-corrected using `beam_mask_motioncorr`, which auto-detects the square beam boundary to restrict motion estimation to the illuminated area, then applies the derived shifts to the full detector frame so that downstream stitching is unaffected.

```
beam_mask_motioncorr \
    --input "/data/frames/*.tif" \
    --output-dir ./motioncorr \
    --pixel-size 1.56 \
    --gpu 0
```

Key arguments:
- `--input` — glob pattern or list of multi-frame TIFF files
- `--output-dir` — where to write the motion-corrected MRCs
- `--pixel-size` — pixel size in Å
- `--mask-threshold` / `--mask-shrink` — tune beam detection if the default fails (use `choose_mask_params` to find values interactively)
- `--save-diagnostic` — save a PNG showing the detected beam mask and crop region for each tile
- `--split-frames` — also write odd/even frame sums to `output-dir/odd/` and `output-dir/even/` for cryoCARE denoising

If you have access to an HPC, for large datasets, generate and submit a SLURM array job:

```
beam_mask_motioncorr --input "/data/frames/*.tif" --output-dir ./motioncorr \
    --pixel-size 1.56 --print-slurm > mc2_job.sh
sbatch mc2_job.sh
```

---

# Step 3: Stitch montage tilt series

Assemble the per-tile motion-corrected MRCs into a single montage tilt series per tilt angle, then join all tilts into one MRC stack:

```
stitch_square_beam \
    -i ./motioncorr \
    -I Imageshifts.txt \
    -o ./stitched \
    --mark-uncovered
```

Key arguments:
- `-i` — directory of per-tile MRCs from `beam_mask_motioncorr`, or a glob of per-tilt MRC stacks
- `-I` — image shifts file from `generate_image_shifts`
- `-o` — output directory (for individual tiff output) or single mrc file for direct stack output
- `-g` — gain reference (omit to skip gain correction, e.g. if already applied during motion correction)
- `-s` — skip cross-correlation refinement of tile positions and use image shifts directly
- `--mark-uncovered` — mark regions with no tile coverage with pixel value −1 rather than inpainting them; **recommended** when the output will be processed by `mask_and_inpaint`, which detects this sentinel automatically
- `-nt` — number of threads for parallel stitching
- `--correct-beam-edges` — correct plasmon-scattering darkening at beam edges (requires `--referencebeam`)

(If necessary), join all the per-tilt stitched images into a single MRC tilt series for reconstruction:

```
crop_to_smallest_common_size -i ./stitched -o ./stitched_stack
```

This crops all tilt images to the smallest common size (accounting for changes in montage field of view with tilt angle) and writes a single `.mrc` stack.

---

# Step 4: Mask and inpaint

The stitched montage contains background vacuum and thick regions outside the lamella, which degrade alignment and reconstruction. `mask_and_inpaint` provides an interactive GUI to define the lamella intensity window and inpaint everything outside it.

```
mask_and_inpaint Montage_stitched.mrc -o Montage_inpainted.mrc
```

In the GUI:
1. **Click on the histogram peak** corresponding to the lamella (the main bright peak in the electron image). The tool fits a Gaussian to that peak and propagates the fit across all tilts, tracking the tilt-angle-dependent intensity drift automatically.
2. Use the **N sigma** slider to widen or narrow the kept intensity window.
3. Toggle **Smooth µ** to stabilise the peak fits at high tilts where the signal weakens.
4. Use **Manual** mode to override the threshold for individual tilts where the automatic fit fails.
5. Click **Generate Output** to inpaint all masked regions using smooth interpolation and write the output MRC.

For large tilt series, use **Save SLURM Job** to write a batch script that applies the fitted parameters in parallel, then submit it:

```
sbatch Montage_stitched_inpaint_slurm.sh
```

---

# Step 5: Reconstruction

The inpainted tilt series `Montage_inpainted.mrc` is ready for standard tomographic reconstruction.

**IMOD:** Import into etomo or use `tilt` directly after alignment with `tiltalign`. The montage tilt series is large so consider binning during reconstruction.

**AreTomo2/3:** Consider using memory efficient implementation of [`AreTomo2`](https://github.com/HamishGBrown/MaskedMemoryEfficientAreTomo), this streams reconstructions to disk rather than storing the whole array in memory.

SLURM submission script templates for each step of the pipeline are provided in [`slurm_templates/`](slurm_templates/). Edit the `EDIT_ME` placeholders at the top of each script for your paths and parameters:

| Script | Purpose |
|---|---|
| `01_motion_correction.sh` | Array job: motion-correct one TIFF per task |
| `02_stitch.sh` | Stitch tiles into a tilt series MRC stack |
| `03_inpaint_apply.sh` | Apply inpainting params (generated by `mask_and_inpaint` GUI) |
| `04_AreTomo.sh` | AreTomo2 reconstruction |
