## Supplementary material for "Whole cryo-FIB lamella acquisition at high magnification with rectangular beam montage cryo-electron tomography": Source code: DetermineOverlapFraction.txt

#!Python
#ScriptName	DetermineOverlapFraction
import os
import copy
import json
import numpy as np
import matplotlib
matplotlib.use('TkAgg')  # X11-forwarding compatible backend
from matplotlib import pyplot as plt
from matplotlib.widgets import Slider
import argparse
from collections import deque
from matplotlib.patches import Rectangle
import serialem as sem 
from matplotlib.widgets import Button

# ---------------------------------------------------------------------------
# Beam shape for this microscope/beam setup. Set once when configuring the
# scripts for a given instrument:
#   "square" - rectangular beam, interactive x/y overlap-factor sliders.
#   "round"  - circular beam tiled hexagonally, interactive hexagon side
#              length slider.
#
# This script uses matplotlib for its interactive sliders. If you'd rather
# not install matplotlib on this PC (e.g. it has no internet access to pip
# install with), use DetermineOverlapFraction_Tkinter.txt instead -- same
# geometry and BeamGeometry.json output, built on stdlib tkinter only.
# ---------------------------------------------------------------------------
BEAM_SHAPE = "round"

def fourier_crop(ain, shapeout):
    """
    Crop or pad a Fourier-transformed array to match the desired output shape.

    Parameters:
    -----------
    ain : numpy.ndarray
        Input array to be cropped or padded in Fourier space.
    shapeout : tuple
        Desired shape of the output array.

    Returns:
    --------
    numpy.ndarray : Cropped or padded array in Fourier space.
    """

    def crop1d(array, s, d):
        # Number of dimensions of array
        N = len(array.shape)
        # Size of array that will be transferred to new grid
        s_ = min(array.shape[d], s)
        # Indices of grid region to transfer to new grid
        ind1 = (
            (np.s_[:],) * ((N + d) % N)
            + (np.s_[: s_ // 2 + s_ % 2],)
            + (np.s_[:],) * (N - (N + d) % N - 1)
        )
        ind2 = (
            (np.s_[:],) * ((N + d) % N)
            + (np.s_[-s_ // 2 + s_ % 2 :],)
            + (np.s_[:],) * (N - (N + d) % N - 1)
        )
        if s > array.shape[d]:
            xtra = list(array.shape)
            xtra[d] = s - array.shape[d]
            return np.concatenate(
                [array[ind1], np.zeros(xtra, dtype=array.dtype), array[ind2]], axis=d
            )
        else:
            return np.concatenate([array[ind1], array[ind2]], axis=d)

    array = copy.deepcopy(ain)
    for i, s in enumerate(shapeout):
        array = crop1d(array, s, i - len(shapeout))
    return array


def fourier_interpolate(
    ain, shapeout, norm="conserve_val", N=None, qspace_in=False, qspace_out=False
):
    """
    Perfom fourier interpolation on array ain so that its shape matches shapeout.

    Arguments
    ---------
    ain : (...,Ni,..,Ny,Nx) array_like
        Input numpy array, interpolation will be applied to the n trailing
        dimensions where n is the length of shapeout.
    shapeout : int (n,) , array_like
        Desired shape of output array
    norm : str, optional  {'conserve_val','conserve_norm','conserve_L1'}
        Normalization of output. If 'conserve_val' then array values are preserved
        if 'conserve_norm' L2 norm is conserved under interpolation and if
        'conserve_L1' L1 norm is conserved under interpolation
    N : int, optional
        Number of (trailing) dimensions to Fourier interpolate. By default take
        the length of shapeout
    qspace_in : bool, optional
        Set to True if the input array is in reciprocal space, False if not (default).
        Be careful with setting this to True for a non-complex array.
    qspace_out : bool, optional
        Set to True for reciprocal space output, False for real-space output (default).
    """
    # Import required FFT functions
    from numpy.fft import fftn

    if N is None:
        N = len(shapeout)

    # Make input complex
    aout = np.zeros(np.shape(ain)[:-N] + tuple(shapeout), dtype=complex)

    # Get input dimensions
    shapein = np.shape(ain)[-N:]

    # axes to Fourier transform
    axes = np.arange(-N, 0)

    if qspace_in:
        a = np.asarray(ain, dtype=complex)
    else:
        a = fftn(np.asarray(ain, dtype=complex), axes=axes)

    aout = fourier_crop(a, shapeout)
    # aout = np.fft.fftshift(crop(np.fft.fftshift(a,axes=axes),shapeout),axes=axes)

    # Fourier transform result with appropriate normalization
    if norm == "conserve_val":
        aout *= np.prod(shapeout) / np.prod(np.shape(ain)[-N:])
    elif norm == "conserve_norm":
        aout *= np.sqrt(np.prod(shapeout) / np.prod(np.shape(ain)[-N:]))

    if not qspace_out:
        aout = np.fft.ifftn(aout, axes=axes)

    # Return correct array data type
    if not np.iscomplexobj(ain):
        return np.real(aout)
    else:
        return aout

def convolve(array1, array2, axes=None):
    """
    Fourier convolution of two arrays over specified axes.

    array2 is broadcast to match array1 so axes refers to the dimensions of
    array1
    """
    # input and output shape
    s = array1.shape
    # Broadcast array2 to match array1
    a2 = np.broadcast_to(array2, s)
    # Axes of transformation
    a = axes
    if a is not None:
        s = [s[i] for i in a]
    if np.iscomplexobj(array1) or np.iscomplexobj(a2):
        return np.fft.ifftn(np.fft.fftn(array1, s, a) * np.fft.fftn(a2, s, a), s, a)
    else:
        return np.fft.irfftn(np.fft.rfftn(array1, s, a) * np.fft.rfftn(a2, s, a), s, a)

def Gaussian(sigma, gridshape):
    r"""
    Calculate a 2D Gaussian function.

    Notes
    -----
    Functional form
    .. math:: 1 / \sqrt { 2 \pi \sigma }  e^{ - ( x^2 + y^2 ) / 2 / \sigma^2 }

    Parameters
    ----------
    sigma : float or (2,) array_like
        The standard deviation of the Gaussian function in pixels
    gridshape : (2,) array_like
        Number of pixels in the grid.
    """
    ysqr = np.fft.fftfreq(gridshape[0], d=1 / gridshape[0]) ** 2
    xsqr = np.fft.fftfreq(gridshape[1], d=1 / gridshape[1]) ** 2

    gaussian = np.exp(-(ysqr[:, None] + xsqr[None, :]) / sigma**2 / 2)
    return gaussian / np.sum(gaussian)


def circular_mask(size, radius=None, center=None):
    """
    Create a binary circular mask in a 2D array.

    Parameters:
    - size (int, int): Size of the array (rows, cols).
    - radius (int, optional): Radius of the circle. Defaults to min(size)/2.
    - center (int, int, optional): Center of the circle (row, col). Defaults to the center of the array.

    Returns:
    - mask (2D numpy array): Binary mask with a circle.
    """
    rows, cols = size

    if center is None:
        center = (rows // 2, cols // 2)
    if radius is None:
        radius = min(rows, cols) // 2

    Y, X = np.ogrid[:rows, :cols]
    dist_from_center = np.sqrt((X - center[1]) ** 2 + (Y - center[0]) ** 2)

    return dist_from_center <= radius


def binary_erosion_fft(mask, structure):
    """
    Erode a binary mask by a flat structuring element, using the script's
    own FFT-based `convolve()` instead of scipy.ndimage.binary_erosion.

    A pixel survives erosion iff every element of `structure` centered on
    it falls on foreground -- equivalent to: convolve the mask with the
    structuring element and keep pixels where the convolved value equals
    the structuring element's full weight.

    `convolve()` performs a *circular* convolution and expects its second
    argument already in the zero-centered, wraparound layout (peak at index
    (0,0), like `Gaussian()` produces) rather than spatially centered in the
    middle of the array (like `circular_mask()` produces). `structure` is
    re-centered with `np.roll` to that layout before convolving. The mask is
    also zero-padded by half the structuring element's size on each side
    first, so real mask content near the border doesn't wrap around the
    image edges and get treated as if it were foreground there -- matching
    scipy's default `border_value=0`.
    """
    sy, sx = structure.shape
    pad_y, pad_x = sy // 2 + 1, sx // 2 + 1
    padded = np.pad(mask.astype(float), ((pad_y, pad_y), (pad_x, pad_x)), mode="constant")

    kernel = np.zeros_like(padded)
    kernel[:sy, :sx] = structure.astype(float)
    kernel = np.roll(kernel, (-(sy // 2), -(sx // 2)), axis=(0, 1))

    conv = convolve(padded, kernel)
    conv = conv[pad_y:pad_y + mask.shape[0], pad_x:pad_x + mask.shape[1]]
    return conv >= structure.sum() - 0.5


def binary_fill_holes_bfs(mask):
    """
    Fill enclosed holes in a binary mask without scipy.ndimage.binary_fill_holes.

    Flood-fills the background (the inverse of `mask`) starting from the
    image border using 4-connectivity; any background pixel *not* reached
    is an enclosed hole and gets set to foreground.
    """
    h, w = mask.shape
    reachable = np.zeros_like(mask, dtype=bool)
    q = deque()

    def visit(y, x):
        if not mask[y, x] and not reachable[y, x]:
            reachable[y, x] = True
            q.append((y, x))

    for x in range(w):
        visit(0, x)
        visit(h - 1, x)
    for y in range(h):
        visit(y, 0)
        visit(y, w - 1)

    while q:
        y, x = q.popleft()
        for ny, nx in ((y - 1, x), (y + 1, x), (y, x - 1), (y, x + 1)):
            if 0 <= ny < h and 0 <= nx < w:
                visit(ny, nx)

    return mask | ~reachable


def rotate_nearest(mask, angle_degrees):
    """
    Nearest-neighbor rotation about the array center, for the square-beam
    angle-preview plot only -- replaces scipy.ndimage.rotate(reshape=False).
    Nearest-neighbor (vs scipy's default cubic spline) is fine since the
    input is already binary and this only feeds a preview image.
    """
    h, w = mask.shape
    cy, cx = h / 2, w / 2
    theta = np.deg2rad(angle_degrees)
    cos_t, sin_t = np.cos(theta), np.sin(theta)
    yy, xx = np.mgrid[0:h, 0:w]
    yy_c, xx_c = yy - cy, xx - cx
    src_y = cos_t * yy_c + sin_t * xx_c + cy
    src_x = -sin_t * yy_c + cos_t * xx_c + cx
    src_y = np.round(src_y).astype(int)
    src_x = np.round(src_x).astype(int)
    valid = (src_y >= 0) & (src_y < h) & (src_x >= 0) & (src_x < w)
    out = np.zeros_like(mask)
    out[valid] = mask[src_y[valid], src_x[valid]]
    return out


def make_mask(im, shrinkn=20, smoothing_kernel=3):
    """
    Generate a binary mask from an image by applying a Gaussian filter, filling holes,
    and then shrinking the mask with morphological erosion.

    Parameters:
    -----------
    im : numpy.ndarray
        Input 2D image array.
    shrinkn : int, optional
        Factor determining the size of the structuring element for shrinking the mask.
        Default is 20.
    smoothing_kernel : float,optional
        Sigma of the gaussian smoothing kernel before applying the mask (3 by default)

    Returns:
    --------
    numpy.ndarray : Binary mask of the same shape as the input image.
    """
    # Step 1: Apply a Gaussian filter to smooth the image
    smoothed_im = convolve(im, Gaussian(smoothing_kernel, im.shape))

    # Step 2: Create an initial binary mask based on a threshold (0.4 * median of the original image)
    mask = smoothed_im > 0.4 * np.median(im)

    # Step 3: Fill any holes in the initial mask
    mask = binary_fill_holes_bfs(mask)

    # Step 4: Create a circular structuring element to use for binary erosion
    struct_elem = circular_mask([shrinkn * 2, shrinkn * 2], radius=shrinkn / 2)

    # Step 5: Apply binary erosion to shrink the mask
    mask = binary_erosion_fft(mask, structure=struct_elem)

    return mask

def parse_commandline():
    """
    Parse commandline input.
    """
    parser = argparse.ArgumentParser(
        description="Determine actual overlap fraction for square beam data."
    )
    parser.add_argument(
        "-i", "--input", help="*.mrc file for raw data", required=True, type=str
    )
    parser.add_argument(
        "-o", "--overlapfactor", help="Overlap factor in x and y", required=True, type=int, nargs=2,metavar=("oy","ox")
    )
    return vars(parser.parse_args())

def roll_no_periodic(arr, shift, fill_value=0, angle=0):
    """
    Rolls an array along the given shift vector but without periodic boundary conditions.
    Vacated positions will be filled with fill_value. The shift can be rotated by a given angle.

    Parameters:
    - arr: numpy array to be shifted.
    - shift: tuple of ints, amount to shift in (y, x) directions. Positive values shift right/down, negative values shift left/up.
    - fill_value: value to place in the vacated positions.
    - angle: float, angle in degrees to rotate the shift vector.

    Returns:
    - A new numpy array with the same shape as arr, but shifted.
    """

    # Rotate the shift vector by the given angle
    angle_rad = np.deg2rad(angle)
    shift_rotated = (
        int(shift[0] * np.cos(angle_rad) - shift[1] * np.sin(angle_rad)),
        int(shift[0] * np.sin(angle_rad) + shift[1] * np.cos(angle_rad))
    )

    result = np.full_like(arr, fill_value)  # Create an array filled with the fill_value

    src_slice = [slice(None)] * arr.ndim
    dst_slice = [slice(None)] * arr.ndim

    for ax, s in enumerate(shift_rotated):
        if s == 0:
            continue  # No shift for this axis

        # Determine the slices that will remain after the shift
        if s > 0:
            src_slice[ax] = slice(0, -s)
            dst_slice[ax] = slice(s, None)
        elif s < 0:
            src_slice[ax] = slice(-s, None)
            dst_slice[ax] = slice(0, s)

    result[tuple(dst_slice)] = arr[tuple(src_slice)]

    return result

def rotation_matrix(angle_degrees):
    """
    Generates a 2D rotation matrix for a given angle in degrees.

    Parameters:
    - angle_degrees: The rotation angle in degrees.

    Returns:
    - A 2x2 NumPy array representing the rotation matrix.
    """
    angle_radians = np.deg2rad(angle_degrees)
    cos_theta = np.cos(angle_radians)
    sin_theta = np.sin(angle_radians)
    return np.array([[cos_theta, -sin_theta], [sin_theta, cos_theta]])


def setup_square_beam_controls(fig, ax, mask, angle):
    ax_overlapy = plt.axes([0.33, 0.1, 0.25, 0.03])
    ax_overlapx = plt.axes([0.66, 0.1, 0.25, 0.03])
    oy,ox = [10,10]
    overlapy_slider = Slider(ax_overlapy, "Y overlap", 2, 20, valinit=oy, valstep=0.1,valfmt='%0.1f')
    overlapx_slider = Slider(ax_overlapx, "X overlap", 2, 20, valinit=ox, valstep=0.1,valfmt='%0.1f')

    def makeoverlapmaps(oy,ox,mask,angle=0):
        overlapmapy = copy.deepcopy(mask).astype(int)
        s = mask.shape[0]
        overlapmapy += roll_no_periodic(mask, [mask.shape[0]-s//oy,0], angle=angle)
        overlapmapy += roll_no_periodic(mask, [s//oy-mask.shape[0],0], angle=angle)
        overlapmapx = copy.deepcopy(mask)
        overlapmapx += roll_no_periodic(mask, [0,s//ox-mask.shape[1]], angle=angle)
        overlapmapx += roll_no_periodic(mask, [0,mask.shape[1]-s//ox], angle=angle)
        return overlapmapy,overlapmapx

    def make_combined_rgb(overlapmapy, overlapmapx):
        # Red = y-neighbour overlap, green = x-neighbour overlap (each 0..2,
        # scaled to 0..1). Yellow marks a point confirmed doubly-covered in
        # both directions; solid red-only or green-only flags a direction
        # with no confirmed neighbour overlap there.
        rgb = np.zeros(overlapmapy.shape + (3,))
        rgb[..., 0] = overlapmapy / 2.0
        rgb[..., 1] = overlapmapx / 2.0
        return rgb

    overlapmapy,overlapmapx = makeoverlapmaps(oy,ox,mask,angle=angle)
    shifty = np.asarray((mask.shape[0] - mask.shape[0]//oy,0)) @ rotation_matrix(angle).T
    shiftx = np.asarray((0,mask.shape[1] - mask.shape[1]//ox))@ rotation_matrix(angle).T
    im1 = ax[1].imshow(make_combined_rgb(overlapmapy, overlapmapx))
    ax[1].axis("off")
    plusrecty = Rectangle(shifty[::-1],mask.shape[1],mask.shape[0],fill=False,edgecolor='red')
    minusrecty = Rectangle(-shifty[::-1],mask.shape[1],mask.shape[0],fill=False,edgecolor='red')
    ax[1].add_patch(plusrecty)
    ax[1].add_patch(minusrecty)
    plusrectx = Rectangle(shiftx[::-1],mask.shape[1],mask.shape[0],fill=False,edgecolor='lime')
    minusrectx = Rectangle(-shiftx[::-1],mask.shape[1],mask.shape[0],fill=False,edgecolor='lime')
    ax[1].add_patch(plusrectx)
    ax[1].add_patch(minusrectx)

    def update(val):
        oy = overlapy_slider.val
        ox = overlapx_slider.val
        shifty = np.asarray((mask.shape[0] - mask.shape[0]//oy,0)) @ rotation_matrix(angle).T
        shiftx = np.asarray((0,mask.shape[1] - mask.shape[1]//ox))@ rotation_matrix(angle).T
        plusrecty.set_xy(shifty[::-1])
        minusrecty.set_xy(-shifty[::-1])
        plusrectx.set_xy(shiftx[::-1])
        minusrectx.set_xy(-shiftx[::-1])
        overlapmapy,overlapmapx = makeoverlapmaps(oy,ox,mask,angle=angle)
        overlapyfraction = np.sum(overlapmapy == 2) / np.prod(overlapmapy.shape)
        overlapxfraction = np.sum(overlapmapx == 2) / np.prod(overlapmapx.shape)
        im1.set_data(make_combined_rgb(overlapmapy, overlapmapx))
        ax[1].set_title(
            'Overlapy fraction (red): {0:.2f}   |   Overlapx fraction (green): {1:.2f}'.format(
                overlapyfraction, overlapxfraction
            )
        )
        fig.canvas.draw_idle()

    overlapx_slider.on_changed(update)
    overlapy_slider.on_changed(update)
    update(None)

    # Widgets must stay externally referenced or Python's garbage collector
    # can reap them (and their on_changed callback) with no error -- the
    # window keeps working, the slider just goes silently dead. Return them
    # so the caller can keep them alive at least until plt.show() returns.
    return overlapy_slider, overlapx_slider


def setup_round_beam_controls(fig, ax, mask, angle, binning=1.0):
    """
    Interactive hexagon-side-length control for a round beam tiled on a
    hexagonal (triangular-lattice) grid. The hexagon side length equals the
    centre-to-centre spacing of neighbouring beam positions (a regular
    hexagon's side length equals its circumradius), so the six neighbour
    directions are the same base shift vector rotated by 0/60/.../300
    degrees (plus the calibrated beam angle) about the beam centre.

    Parameters
    ----------
    binning : float, optional
        Ratio of the raw (unbinned) record image width to the downsampled
        preview `mask` width (512 px). The slider itself operates in preview
        pixels -- the geometry in make_hex_overlap_map needs that -- but the
        reported side length is converted to unbinned pixels, since that is
        the unit SetupPolygonMontage.txt expects.
    """
    max_side = min(mask.shape)
    hexside_init = min(max(2, min(mask.shape) / 2), max_side)

    ax_hexside = plt.axes([0.375, 0.1, 0.35, 0.03])
    hexside_slider = Slider(
        ax_hexside, "Hexagon side length (display px)", 2, max_side,
        valinit=hexside_init, valstep=1, valfmt='%0.0f'
    )

    def make_hex_overlap_map(side_length, angle=0):
        overlap_map = mask.astype(int).copy()
        for i in range(6):
            overlap_map = overlap_map + roll_no_periodic(mask, [side_length, 0], angle=angle + i * 60)
        return overlap_map

    overlap_map = make_hex_overlap_map(hexside_init, angle=angle)
    im1 = ax[1].imshow(overlap_map, vmin=0, vmax=3)
    ax[1].axis("off")

    def update(val):
        side_length = hexside_slider.val
        overlap_map = make_hex_overlap_map(side_length, angle=angle)
        overlap_fraction = np.sum(overlap_map >= 2) / np.prod(overlap_map.shape)
        im1.set_data(overlap_map)
        ax[1].set_title(
            'Hex overlap fraction: {0:.2f}  |  side length = {1:.0f} px (display) '
            '= {2:.0f} px (unbinned)'.format(overlap_fraction, side_length, side_length * binning)
        )
        fig.canvas.draw_idle()

    hexside_slider.on_changed(update)
    update(hexside_init)

    # See the matching comment in setup_square_beam_controls: without an
    # external reference, the slider (and its on_changed callback) can be
    # silently garbage-collected before it's ever interacted with.
    return (hexside_slider,)


def main():
    # Take record image, save it as reference and read from serial EM buffer
    a = sem.R()

    sem.OpenNewFile('ReferenceBeam.mrc')
    sem.Save('A')
    sem.CloseFile()
    print('ReferenceBeam.mrc saved in ' + sem.ReportDirectory() + ', for use in stitching')

    image = np.asarray(sem.bufferImage('A'))

    # Preview is downsampled to a fixed 512 px width to keep the sliders
    # responsive; record it so reported pixel values can be converted back
    # to unbinned (raw record image) pixels.
    binning = image.shape[1] / 512

    image = fourier_interpolate(image, (int(image.shape[0]/image.shape[1]*512),512))
    oy,ox=[10,10]
    mask = np.where(make_mask(image, 20,3),1,0)  # Initial blur with small sigma

    # For rectangular beams, work out rotation of rectangle with respect to camera
    if BEAM_SHAPE=='square':
        fig,axes  = plt.subplots(1,2,figsize=(12,6))
        ax_angle = plt.axes([0.05, 0.05, 0.5, 0.03])

        maskax,rotated_mask_ax = axes
        maskax.imshow(mask)
        angle_slider = Slider(ax_angle, "Angle", -180, 180, valinit=0, valstep=1, valfmt='%0.0f')
       
        rotated_mask_ax.set_title("Rotated Mask")
        rotated_mask_ax.axis("off")
        rotated_mask_im = rotated_mask_ax.imshow(mask, cmap='gray')

        def rotate_mask(mask, angle):
            return rotate_nearest(mask, -angle)

        def update_rotation(val):
            angle = angle_slider.val
            rotated_mask = rotate_mask(mask, angle)
            rotated_mask_im.set_data(rotated_mask)
            fig.canvas.draw_idle()

        angle_slider.on_changed(update_rotation)
        plt.show()
        vmin = np.percentile(image, 10)
        vmax = np.percentile(image, 90)

        angle = angle_slider.val
    else:
        #For round beam the angle not relevant, set to zero
        angle = 0.0
   

    # Plot the original image and create space for the eroded image
    fig, ax = plt.subplots(1, 3, figsize=(18, 6))
    plt.subplots_adjust(bottom=0.25)

    ax[0].set_title("Original Image")
    ax[0].axis("off")
    ax[0].imshow(np.where(mask,1,0))
    ax[2].axis("off")

    if BEAM_SHAPE == "round":
        widgets = setup_round_beam_controls(fig, ax, mask, angle, binning=binning)
    else:
        widgets = setup_square_beam_controls(fig, ax, mask, angle)

    # Keep a live reference to the slider(s) for as long as the window is
    # open -- see the comment in each setup_*_beam_controls function.
    fig._widgets = widgets

    # Show the interactive plot
    plt.show()
    wdir = sem.ReportDirectory()
    fig.savefig(os.path.join(wdir,'overlap.pdf'))

    # Final slider value(s), read after the window is closed. overlap here
    # matches the "overlap" argument SetupPolygonMontage.txt expects: [oy,ox]
    # for a square beam, or a single hexagon side length in *unbinned* camera
    # pixels (hence * binning) for a round beam -- see setup_round_beam_controls.
    if BEAM_SHAPE == "round":
        (hexside_slider,) = widgets
        overlap = hexside_slider.val * binning
    else:
        overlapy_slider, overlapx_slider = widgets
        overlap = [overlapy_slider.val, overlapx_slider.val]

    geometry_path = os.path.join(wdir, 'BeamGeometry.json')
    with open(geometry_path, 'w') as f:
        json.dump(
            {"beam_shape": BEAM_SHAPE, "rotation_deg": angle, "overlap": overlap},
            f, indent=2,
        )
    print('Wrote beam geometry to ' + geometry_path + ' for SetupPolygonMontage.txt to read')

if __name__=='__main__':
    main()
