## Supplementary material for "Whole cryo-FIB lamella acquisition at high magnification with rectangular beam montage cryo-electron tomography": Source code: DetermineOverlapFraction_Tkinter.txt

#!Python
#ScriptName	DetermineOverlapFraction_Tkinter
import os
import copy
import json
import numpy as np
import tkinter as tk
import argparse
from collections import deque
import serialem as sem

# ---------------------------------------------------------------------------
# Beam shape for this microscope/beam setup. Set once when configuring the
# scripts for a given instrument:
#   "square" - rectangular beam, interactive x/y overlap-factor sliders.
#   "round"  - circular beam tiled hexagonally, interactive hexagon side
#              length slider.
#
# Tkinter-only variant of DetermineOverlapFraction.txt: same geometry, same
# BeamGeometry.json output, but the GUI uses only Python's stdlib tkinter
# (ships with every standard Windows Python install) instead of matplotlib,
# for microscope control PCs that are offline and can't easily pip install
# matplotlib. Prefer DetermineOverlapFraction.txt when matplotlib is already
# available there -- its plotting code is simpler to read and extend.
# ---------------------------------------------------------------------------
BEAM_SHAPE = "round"

def fourier_crop(ain, shapeout):
    """
    Crop or pad a Fourier-transformed array to match the desired output shape.

    Parameters:
    -----------
    ain : numpy.ndarray
        Input array to be cropped or padded in Fourier space.
    shapeout : tuple
        Desired shape of the output array.

    Returns:
    --------
    numpy.ndarray : Cropped or padded array in Fourier space.
    """

    def crop1d(array, s, d):
        # Number of dimensions of array
        N = len(array.shape)
        # Size of array that will be transferred to new grid
        s_ = min(array.shape[d], s)
        # Indices of grid region to transfer to new grid
        ind1 = (
            (np.s_[:],) * ((N + d) % N)
            + (np.s_[: s_ // 2 + s_ % 2],)
            + (np.s_[:],) * (N - (N + d) % N - 1)
        )
        ind2 = (
            (np.s_[:],) * ((N + d) % N)
            + (np.s_[-s_ // 2 + s_ % 2 :],)
            + (np.s_[:],) * (N - (N + d) % N - 1)
        )
        if s > array.shape[d]:
            xtra = list(array.shape)
            xtra[d] = s - array.shape[d]
            return np.concatenate(
                [array[ind1], np.zeros(xtra, dtype=array.dtype), array[ind2]], axis=d
            )
        else:
            return np.concatenate([array[ind1], array[ind2]], axis=d)

    array = copy.deepcopy(ain)
    for i, s in enumerate(shapeout):
        array = crop1d(array, s, i - len(shapeout))
    return array


def fourier_interpolate(
    ain, shapeout, norm="conserve_val", N=None, qspace_in=False, qspace_out=False
):
    """
    Perfom fourier interpolation on array ain so that its shape matches shapeout.

    Arguments
    ---------
    ain : (...,Ni,..,Ny,Nx) array_like
        Input numpy array, interpolation will be applied to the n trailing
        dimensions where n is the length of shapeout.
    shapeout : int (n,) , array_like
        Desired shape of output array
    norm : str, optional  {'conserve_val','conserve_norm','conserve_L1'}
        Normalization of output. If 'conserve_val' then array values are preserved
        if 'conserve_norm' L2 norm is conserved under interpolation and if
        'conserve_L1' L1 norm is conserved under interpolation
    N : int, optional
        Number of (trailing) dimensions to Fourier interpolate. By default take
        the length of shapeout
    qspace_in : bool, optional
        Set to True if the input array is in reciprocal space, False if not (default).
        Be careful with setting this to True for a non-complex array.
    qspace_out : bool, optional
        Set to True for reciprocal space output, False for real-space output (default).
    """
    # Import required FFT functions
    from numpy.fft import fftn

    if N is None:
        N = len(shapeout)

    # Make input complex
    aout = np.zeros(np.shape(ain)[:-N] + tuple(shapeout), dtype=complex)

    # Get input dimensions
    shapein = np.shape(ain)[-N:]

    # axes to Fourier transform
    axes = np.arange(-N, 0)

    if qspace_in:
        a = np.asarray(ain, dtype=complex)
    else:
        a = fftn(np.asarray(ain, dtype=complex), axes=axes)

    aout = fourier_crop(a, shapeout)
    # aout = np.fft.fftshift(crop(np.fft.fftshift(a,axes=axes),shapeout),axes=axes)

    # Fourier transform result with appropriate normalization
    if norm == "conserve_val":
        aout *= np.prod(shapeout) / np.prod(np.shape(ain)[-N:])
    elif norm == "conserve_norm":
        aout *= np.sqrt(np.prod(shapeout) / np.prod(np.shape(ain)[-N:]))

    if not qspace_out:
        aout = np.fft.ifftn(aout, axes=axes)

    # Return correct array data type
    if not np.iscomplexobj(ain):
        return np.real(aout)
    else:
        return aout

def convolve(array1, array2, axes=None):
    """
    Fourier convolution of two arrays over specified axes.

    array2 is broadcast to match array1 so axes refers to the dimensions of
    array1
    """
    # input and output shape
    s = array1.shape
    # Broadcast array2 to match array1
    a2 = np.broadcast_to(array2, s)
    # Axes of transformation
    a = axes
    if a is not None:
        s = [s[i] for i in a]
    if np.iscomplexobj(array1) or np.iscomplexobj(a2):
        return np.fft.ifftn(np.fft.fftn(array1, s, a) * np.fft.fftn(a2, s, a), s, a)
    else:
        return np.fft.irfftn(np.fft.rfftn(array1, s, a) * np.fft.rfftn(a2, s, a), s, a)

def Gaussian(sigma, gridshape):
    r"""
    Calculate a 2D Gaussian function.

    Notes
    -----
    Functional form
    .. math:: 1 / \sqrt { 2 \pi \sigma }  e^{ - ( x^2 + y^2 ) / 2 / \sigma^2 }

    Parameters
    ----------
    sigma : float or (2,) array_like
        The standard deviation of the Gaussian function in pixels
    gridshape : (2,) array_like
        Number of pixels in the grid.
    """
    ysqr = np.fft.fftfreq(gridshape[0], d=1 / gridshape[0]) ** 2
    xsqr = np.fft.fftfreq(gridshape[1], d=1 / gridshape[1]) ** 2

    gaussian = np.exp(-(ysqr[:, None] + xsqr[None, :]) / sigma**2 / 2)
    return gaussian / np.sum(gaussian)


def circular_mask(size, radius=None, center=None):
    """
    Create a binary circular mask in a 2D array.

    Parameters:
    - size (int, int): Size of the array (rows, cols).
    - radius (int, optional): Radius of the circle. Defaults to min(size)/2.
    - center (int, int, optional): Center of the circle (row, col). Defaults to the center of the array.

    Returns:
    - mask (2D numpy array): Binary mask with a circle.
    """
    rows, cols = size

    if center is None:
        center = (rows // 2, cols // 2)
    if radius is None:
        radius = min(rows, cols) // 2

    Y, X = np.ogrid[:rows, :cols]
    dist_from_center = np.sqrt((X - center[1]) ** 2 + (Y - center[0]) ** 2)

    return dist_from_center <= radius


def binary_erosion_fft(mask, structure):
    """
    Erode a binary mask by a flat structuring element, using the script's
    own FFT-based `convolve()` instead of scipy.ndimage.binary_erosion.

    A pixel survives erosion iff every element of `structure` centered on
    it falls on foreground -- equivalent to: convolve the mask with the
    structuring element and keep pixels where the convolved value equals
    the structuring element's full weight.

    `convolve()` performs a *circular* convolution and expects its second
    argument already in the zero-centered, wraparound layout (peak at index
    (0,0), like `Gaussian()` produces) rather than spatially centered in the
    middle of the array (like `circular_mask()` produces). `structure` is
    re-centered with `np.roll` to that layout before convolving. The mask is
    also zero-padded by half the structuring element's size on each side
    first, so real mask content near the border doesn't wrap around the
    image edges and get treated as if it were foreground there -- matching
    scipy's default `border_value=0`.
    """
    sy, sx = structure.shape
    pad_y, pad_x = sy // 2 + 1, sx // 2 + 1
    padded = np.pad(mask.astype(float), ((pad_y, pad_y), (pad_x, pad_x)), mode="constant")

    kernel = np.zeros_like(padded)
    kernel[:sy, :sx] = structure.astype(float)
    kernel = np.roll(kernel, (-(sy // 2), -(sx // 2)), axis=(0, 1))

    conv = convolve(padded, kernel)
    conv = conv[pad_y:pad_y + mask.shape[0], pad_x:pad_x + mask.shape[1]]
    return conv >= structure.sum() - 0.5


def binary_fill_holes_bfs(mask):
    """
    Fill enclosed holes in a binary mask without scipy.ndimage.binary_fill_holes.

    Flood-fills the background (the inverse of `mask`) starting from the
    image border using 4-connectivity; any background pixel *not* reached
    is an enclosed hole and gets set to foreground.
    """
    h, w = mask.shape
    reachable = np.zeros_like(mask, dtype=bool)
    q = deque()

    def visit(y, x):
        if not mask[y, x] and not reachable[y, x]:
            reachable[y, x] = True
            q.append((y, x))

    for x in range(w):
        visit(0, x)
        visit(h - 1, x)
    for y in range(h):
        visit(y, 0)
        visit(y, w - 1)

    while q:
        y, x = q.popleft()
        for ny, nx in ((y - 1, x), (y + 1, x), (y, x - 1), (y, x + 1)):
            if 0 <= ny < h and 0 <= nx < w:
                visit(ny, nx)

    return mask | ~reachable


def rotate_nearest(mask, angle_degrees):
    """
    Nearest-neighbor rotation about the array center, for the square-beam
    angle-preview plot only -- replaces scipy.ndimage.rotate(reshape=False).
    Nearest-neighbor (vs scipy's default cubic spline) is fine since the
    input is already binary and this only feeds a preview image.
    """
    h, w = mask.shape
    cy, cx = h / 2, w / 2
    theta = np.deg2rad(angle_degrees)
    cos_t, sin_t = np.cos(theta), np.sin(theta)
    yy, xx = np.mgrid[0:h, 0:w]
    yy_c, xx_c = yy - cy, xx - cx
    src_y = cos_t * yy_c + sin_t * xx_c + cy
    src_x = -sin_t * yy_c + cos_t * xx_c + cx
    src_y = np.round(src_y).astype(int)
    src_x = np.round(src_x).astype(int)
    valid = (src_y >= 0) & (src_y < h) & (src_x >= 0) & (src_x < w)
    out = np.zeros_like(mask)
    out[valid] = mask[src_y[valid], src_x[valid]]
    return out


def make_mask(im, shrinkn=20, smoothing_kernel=3):
    """
    Generate a binary mask from an image by applying a Gaussian filter, filling holes,
    and then shrinking the mask with morphological erosion.

    Parameters:
    -----------
    im : numpy.ndarray
        Input 2D image array.
    shrinkn : int, optional
        Factor determining the size of the structuring element for shrinking the mask.
        Default is 20.
    smoothing_kernel : float,optional
        Sigma of the gaussian smoothing kernel before applying the mask (3 by default)

    Returns:
    --------
    numpy.ndarray : Binary mask of the same shape as the input image.
    """
    # Step 1: Apply a Gaussian filter to smooth the image
    smoothed_im = convolve(im, Gaussian(smoothing_kernel, im.shape))

    # Step 2: Create an initial binary mask based on a threshold (0.4 * median of the original image)
    mask = smoothed_im > 0.4 * np.median(im)

    # Step 3: Fill any holes in the initial mask
    mask = binary_fill_holes_bfs(mask)

    # Step 4: Create a circular structuring element to use for binary erosion
    struct_elem = circular_mask([shrinkn * 2, shrinkn * 2], radius=shrinkn / 2)

    # Step 5: Apply binary erosion to shrink the mask
    mask = binary_erosion_fft(mask, structure=struct_elem)

    return mask

def parse_commandline():
    """
    Parse commandline input.
    """
    parser = argparse.ArgumentParser(
        description="Determine actual overlap fraction for square beam data."
    )
    parser.add_argument(
        "-i", "--input", help="*.mrc file for raw data", required=True, type=str
    )
    parser.add_argument(
        "-o", "--overlapfactor", help="Overlap factor in x and y", required=True, type=int, nargs=2,metavar=("oy","ox")
    )
    return vars(parser.parse_args())

def roll_no_periodic(arr, shift, fill_value=0, angle=0):
    """
    Rolls an array along the given shift vector but without periodic boundary conditions.
    Vacated positions will be filled with fill_value. The shift can be rotated by a given angle.

    Parameters:
    - arr: numpy array to be shifted.
    - shift: tuple of ints, amount to shift in (y, x) directions. Positive values shift right/down, negative values shift left/up.
    - fill_value: value to place in the vacated positions.
    - angle: float, angle in degrees to rotate the shift vector.

    Returns:
    - A new numpy array with the same shape as arr, but shifted.
    """

    # Rotate the shift vector by the given angle
    angle_rad = np.deg2rad(angle)
    shift_rotated = (
        int(shift[0] * np.cos(angle_rad) - shift[1] * np.sin(angle_rad)),
        int(shift[0] * np.sin(angle_rad) + shift[1] * np.cos(angle_rad))
    )

    result = np.full_like(arr, fill_value)  # Create an array filled with the fill_value

    src_slice = [slice(None)] * arr.ndim
    dst_slice = [slice(None)] * arr.ndim

    for ax, s in enumerate(shift_rotated):
        if s == 0:
            continue  # No shift for this axis

        # Determine the slices that will remain after the shift
        if s > 0:
            src_slice[ax] = slice(0, -s)
            dst_slice[ax] = slice(s, None)
        elif s < 0:
            src_slice[ax] = slice(-s, None)
            dst_slice[ax] = slice(0, s)

    result[tuple(dst_slice)] = arr[tuple(src_slice)]

    return result

def rotation_matrix(angle_degrees):
    """
    Generates a 2D rotation matrix for a given angle in degrees.

    Parameters:
    - angle_degrees: The rotation angle in degrees.

    Returns:
    - A 2x2 NumPy array representing the rotation matrix.
    """
    angle_radians = np.deg2rad(angle_degrees)
    cos_theta = np.cos(angle_radians)
    sin_theta = np.sin(angle_radians)
    return np.array([[cos_theta, -sin_theta], [sin_theta, cos_theta]])


# ---------------------------------------------------------------------------
# Tkinter <-> numpy image glue (no Pillow, no matplotlib)
# ---------------------------------------------------------------------------

def array_to_photoimage(rgb):
    """Wrap an (H, W, 3) uint8 array as a Tkinter PhotoImage.

    Builds a binary PPM (P6) buffer in memory and hands it to PhotoImage's
    `data` argument -- Tk's photo image reads PPM natively, so this needs
    neither Pillow nor matplotlib to get a numpy array on screen.
    """
    h, w = rgb.shape[:2]
    header = "P6 {0} {1} 255 ".format(w, h).encode("ascii")
    body = np.ascontiguousarray(rgb, dtype=np.uint8).tobytes()
    return tk.PhotoImage(data=header + body)


def mask_to_rgb(mask):
    """Binary mask -> white-on-black RGB preview image."""
    gray = np.where(mask, 255, 0).astype(np.uint8)
    return np.stack([gray, gray, gray], axis=-1)


# Small hand-rolled colormap for the round-beam overlap-count preview (0..3
# copies of the mask landing on a pixel), standing in for matplotlib's
# default colormap since there's no matplotlib here to supply one.
_COUNT_LUT = np.array([
    [20, 20, 20],    # 0: not covered
    [40, 90, 200],   # 1: seen once
    [60, 170, 90],   # 2: confirmed double coverage
    [250, 220, 40],  # 3+: triple coverage or more
], dtype=np.uint8)


def counts_to_rgb(counts, vmax=3):
    """Map small integer overlap counts to an RGB preview via `_COUNT_LUT`."""
    idx = np.clip(counts, 0, vmax).astype(int)
    return _COUNT_LUT[idx]


def save_array_as_gif(rgb, path):
    """Write an (H, W, 3) uint8 array to disk as a GIF.

    Uses Tkinter's own PhotoImage encoder instead of matplotlib's
    `fig.savefig` -- a short-lived, hidden Tk root is created purely to host
    the image and torn down immediately after, nothing is shown on screen.
    """
    helper_root = tk.Tk()
    helper_root.withdraw()
    try:
        photo = array_to_photoimage(rgb)
        photo.write(path, format="gif")
    finally:
        helper_root.destroy()


# ---------------------------------------------------------------------------
# Interactive windows (one Tk() root per stage; each blocks in mainloop()
# until the user closes it, then the caller reads the final slider value(s)
# out of the Tk variables below)
# ---------------------------------------------------------------------------

def choose_beam_angle(mask):
    """Angle slider + live-rotated mask preview (square beam only).

    Blocks until the window is closed; returns the final angle in degrees.
    """
    root = tk.Tk()
    root.title("Beam angle")
    h, w = mask.shape

    frame = tk.Frame(root)
    frame.pack()
    orig_canvas = tk.Canvas(frame, width=w, height=h)
    orig_canvas.grid(row=0, column=0, padx=4, pady=4)
    rot_canvas = tk.Canvas(frame, width=w, height=h)
    rot_canvas.grid(row=0, column=1, padx=4, pady=4)
    tk.Label(frame, text="Original mask").grid(row=1, column=0)
    tk.Label(frame, text="Rotated mask").grid(row=1, column=1)

    # Static original-mask preview never changes, so it needs no update logic
    orig_photo = array_to_photoimage(mask_to_rgb(mask))
    orig_canvas.create_image(0, 0, anchor="nw", image=orig_photo)

    # Tkinter photo images must stay referenced by Python or they vanish --
    # the canvas item goes blank with no error -- so keep this live in a
    # dict that update() below can freely reassign.
    photo_holder = {"rotated": array_to_photoimage(mask_to_rgb(mask))}
    rot_image_id = rot_canvas.create_image(0, 0, anchor="nw", image=photo_holder["rotated"])

    def update(_value=None):
        rotated = rotate_nearest(mask, -angle_var.get())
        photo_holder["rotated"] = array_to_photoimage(mask_to_rgb(rotated))
        rot_canvas.itemconfig(rot_image_id, image=photo_holder["rotated"])

    angle_var = tk.DoubleVar(value=0.0)
    tk.Scale(
        root, from_=-180, to=180, resolution=1, orient="horizontal", length=400,
        label="Angle", variable=angle_var, command=update,
    ).pack(fill="x", padx=8, pady=4)
    tk.Button(root, text="Done", command=root.destroy).pack(pady=4)

    update()
    root.mainloop()
    return angle_var.get()


def choose_square_overlap(mask, angle):
    """Interactive Y/X overlap-fraction sliders for a rectangular beam.

    Mirrors `setup_square_beam_controls` in DetermineOverlapFraction.txt --
    same overlap-map geometry, same red/green combined preview. Blocks until
    the window is closed; returns (oy, ox, final_rgb_preview).
    """
    root = tk.Tk()
    root.title("Square beam overlap")
    h, w = mask.shape

    canvas = tk.Canvas(root, width=w, height=h)
    canvas.pack(padx=4, pady=4)
    title_var = tk.StringVar()
    tk.Label(root, textvariable=title_var).pack()

    def make_overlap_maps(oy, ox):
        overlapmapy = mask.astype(int).copy()
        s = mask.shape[0]
        overlapmapy = overlapmapy + roll_no_periodic(mask, [mask.shape[0] - s // oy, 0], angle=angle)
        overlapmapy = overlapmapy + roll_no_periodic(mask, [s // oy - mask.shape[0], 0], angle=angle)
        overlapmapx = mask.astype(int).copy()
        overlapmapx = overlapmapx + roll_no_periodic(mask, [0, s // ox - mask.shape[1]], angle=angle)
        overlapmapx = overlapmapx + roll_no_periodic(mask, [0, mask.shape[1] - s // ox], angle=angle)
        return overlapmapy, overlapmapx

    def combined_rgb(overlapmapy, overlapmapx):
        # Red = y-neighbour overlap, green = x-neighbour overlap (each 0..2,
        # scaled to 0..255). Yellow marks a point confirmed doubly-covered in
        # both directions; solid red-only or green-only flags a direction
        # with no confirmed neighbour overlap there.
        rgb = np.zeros(overlapmapy.shape + (3,), dtype=np.uint8)
        rgb[..., 0] = (overlapmapy / 2.0 * 255).astype(np.uint8)
        rgb[..., 1] = (overlapmapx / 2.0 * 255).astype(np.uint8)
        return rgb

    def rect_coords(shift):
        # shift is (row, col); Rectangle/Canvas coordinates want (x, y) =
        # (col, row), matching the matplotlib version's `shift[::-1]`
        x, y = float(shift[1]), float(shift[0])
        return x, y, x + w, y + h

    oy0, ox0 = 10.0, 10.0
    overlapmapy0, overlapmapx0 = make_overlap_maps(oy0, ox0)
    photo_holder = {"photo": array_to_photoimage(combined_rgb(overlapmapy0, overlapmapx0))}
    image_id = canvas.create_image(0, 0, anchor="nw", image=photo_holder["photo"])

    shifty0 = np.asarray((h - h // oy0, 0)) @ rotation_matrix(angle).T
    shiftx0 = np.asarray((0, w - w // ox0)) @ rotation_matrix(angle).T
    plus_recty = canvas.create_rectangle(*rect_coords(shifty0), outline="red")
    minus_recty = canvas.create_rectangle(*rect_coords(-shifty0), outline="red")
    plus_rectx = canvas.create_rectangle(*rect_coords(shiftx0), outline="lime")
    minus_rectx = canvas.create_rectangle(*rect_coords(-shiftx0), outline="lime")

    def update(_value=None):
        oy, ox = oy_var.get(), ox_var.get()
        shifty = np.asarray((h - h // oy, 0)) @ rotation_matrix(angle).T
        shiftx = np.asarray((0, w - w // ox)) @ rotation_matrix(angle).T
        canvas.coords(plus_recty, *rect_coords(shifty))
        canvas.coords(minus_recty, *rect_coords(-shifty))
        canvas.coords(plus_rectx, *rect_coords(shiftx))
        canvas.coords(minus_rectx, *rect_coords(-shiftx))

        overlapmapy, overlapmapx = make_overlap_maps(oy, ox)
        overlapyfraction = np.sum(overlapmapy == 2) / np.prod(overlapmapy.shape)
        overlapxfraction = np.sum(overlapmapx == 2) / np.prod(overlapmapx.shape)
        photo_holder["photo"] = array_to_photoimage(combined_rgb(overlapmapy, overlapmapx))
        canvas.itemconfig(image_id, image=photo_holder["photo"])
        title_var.set(
            "Overlapy fraction (red): {0:.2f}   |   Overlapx fraction (green): {1:.2f}".format(
                overlapyfraction, overlapxfraction
            )
        )

    oy_var = tk.DoubleVar(value=oy0)
    ox_var = tk.DoubleVar(value=ox0)
    tk.Scale(
        root, from_=2, to=20, resolution=0.1, orient="horizontal", length=400,
        label="Y overlap", variable=oy_var, command=update,
    ).pack(fill="x", padx=8)
    tk.Scale(
        root, from_=2, to=20, resolution=0.1, orient="horizontal", length=400,
        label="X overlap", variable=ox_var, command=update,
    ).pack(fill="x", padx=8)
    tk.Button(root, text="Done", command=root.destroy).pack(pady=4)

    update()
    root.mainloop()

    final_oy, final_ox = oy_var.get(), ox_var.get()
    final_map_y, final_map_x = make_overlap_maps(final_oy, final_ox)
    return final_oy, final_ox, combined_rgb(final_map_y, final_map_x)


def choose_hex_overlap(mask, angle, binning=1.0):
    """Interactive hexagon-side-length control for a round beam tiled on a
    hexagonal (triangular-lattice) grid.

    Mirrors `setup_round_beam_controls` in DetermineOverlapFraction.txt: the
    hexagon side length equals the centre-to-centre spacing of neighbouring
    beam positions (a regular hexagon's side length equals its
    circumradius), so the six neighbour directions are the same base shift
    vector rotated by 0/60/.../300 degrees (plus the calibrated beam angle)
    about the beam centre.

    Parameters
    ----------
    binning : float, optional
        Ratio of the raw (unbinned) record image width to the downsampled
        preview `mask` width (512 px). The slider itself operates in preview
        pixels -- the geometry in make_hex_overlap_map needs that -- but the
        returned side length is converted to unbinned pixels, since that is
        the unit SetupPolygonMontage.txt expects.

    Blocks until the window is closed; returns
    (side_length_unbinned_px, final_rgb_preview).
    """
    root = tk.Tk()
    root.title("Round beam overlap")
    h, w = mask.shape
    max_side = min(mask.shape)
    side_init = min(max(2, min(mask.shape) / 2), max_side)

    canvas = tk.Canvas(root, width=w, height=h)
    canvas.pack(padx=4, pady=4)
    title_var = tk.StringVar()
    tk.Label(root, textvariable=title_var).pack()

    def make_hex_overlap_map(side_length):
        overlap_map = mask.astype(int).copy()
        for i in range(6):
            overlap_map = overlap_map + roll_no_periodic(mask, [side_length, 0], angle=angle + i * 60)
        return overlap_map

    overlap_map0 = make_hex_overlap_map(side_init)
    photo_holder = {"photo": array_to_photoimage(counts_to_rgb(overlap_map0))}
    image_id = canvas.create_image(0, 0, anchor="nw", image=photo_holder["photo"])

    def update(_value=None):
        side_length = side_var.get()
        overlap_map = make_hex_overlap_map(side_length)
        overlap_fraction = np.sum(overlap_map >= 2) / np.prod(overlap_map.shape)
        photo_holder["photo"] = array_to_photoimage(counts_to_rgb(overlap_map))
        canvas.itemconfig(image_id, image=photo_holder["photo"])
        title_var.set(
            "Hex overlap fraction: {0:.2f}  |  side length = {1:.0f} px (display) "
            "= {2:.0f} px (unbinned)".format(overlap_fraction, side_length, side_length * binning)
        )

    side_var = tk.DoubleVar(value=side_init)
    tk.Scale(
        root, from_=2, to=max_side, resolution=1, orient="horizontal", length=400,
        label="Hexagon side length (display px)", variable=side_var, command=update,
    ).pack(fill="x", padx=8)
    tk.Button(root, text="Done", command=root.destroy).pack(pady=4)

    update()
    root.mainloop()

    final_side = side_var.get()
    final_map = make_hex_overlap_map(final_side)
    return final_side * binning, counts_to_rgb(final_map)


def main():
    # Take record image, save it as reference and read from SerialEM buffer
    a = sem.R()

    sem.OpenNewFile('ReferenceBeam.mrc')
    sem.Save('A')
    sem.CloseFile()
    print('ReferenceBeam.mrc saved in ' + sem.ReportDirectory() + ', for use in stitching')

    image = np.asarray(sem.bufferImage('A'))

    # Preview is downsampled to a fixed 512 px width to keep the sliders
    # responsive; record it so reported pixel values can be converted back
    # to unbinned (raw record image) pixels.
    binning = image.shape[1] / 512

    image = fourier_interpolate(image, (int(image.shape[0] / image.shape[1] * 512), 512))
    mask = np.where(make_mask(image, 20, 3), 1, 0)  # Initial blur with small sigma

    # For rectangular beams, work out rotation of rectangle with respect to camera
    if BEAM_SHAPE == 'square':
        angle = choose_beam_angle(mask)
    else:
        # For round beam the angle is not relevant, set to zero
        angle = 0.0

    # Final slider value(s), read after each window is closed. overlap here
    # matches the "overlap" argument SetupPolygonMontage.txt expects: [oy,ox]
    # for a square beam, or a single hexagon side length in *unbinned* camera
    # pixels (hence * binning) for a round beam -- see choose_hex_overlap.
    if BEAM_SHAPE == "round":
        overlap, final_rgb = choose_hex_overlap(mask, angle, binning=binning)
    else:
        oy, ox, final_rgb = choose_square_overlap(mask, angle)
        overlap = [oy, ox]

    wdir = sem.ReportDirectory()
    save_array_as_gif(final_rgb, os.path.join(wdir, 'overlap.gif'))

    geometry_path = os.path.join(wdir, 'BeamGeometry.json')
    with open(geometry_path, 'w') as f:
        json.dump(
            {"beam_shape": BEAM_SHAPE, "rotation_deg": angle, "overlap": overlap},
            f, indent=2,
        )
    print('Wrote beam geometry to ' + geometry_path + ' for SetupPolygonMontage.txt to read')

if __name__=='__main__':
    main()
