## Supplementary material for "Whole cryo-FIB lamella acquisition at high magnification with rectangular beam montage cryo-electron tomography": Source code: MontageTiltSeries.txt

MacroName MontageTiltSeries

##Realign to center image up to 3 times when the shift is more than 1um, end with nonzero image shift
##Center reference image taken at view mode and saved in buffer T
LmtRATimes = 3
LmtISShift = 1
ISDelay = 0.5
Beamcentering = 0

## Set to anything but 0 for coma free image shift
## Set to 0 to not balance coma in image shift
ComaFree = 1

Require arrays
ReportNavItem

## Get data directory, image prefix, tile coordinate file, number of tilts in series 
## and montage tiles per tilt from commandline
Echo Outputting to Directory:
ReportDirectory
FramesDirectory = $ReportedValue1\Frames
makeDirectory $FramesDirectory
SetFolderForFrames $FramesDirectory
OFPrefix = Montage_$navLabel
SetFrameBaseName 0 1 -1 $OFPrefix

## Set filename format for frames
SetFrameNameFormat 1 -1 0x468


ParFilename = Montage_imageshifts_$navLabel.txt

## Initialize arrays for storing stage positions
NewArray PlusStagePos -1 2

NewArray MinusStagePos -1 2

## Opens text file $ParFilename as read-only (r) and with ID MTSPar
OpenTextFile MTSPar r 0 $ParFilename
## Read line from image shift text file (id MTSPar) to array TSNumber (number of tilt series)
NewArray TSNumber -1 1

# Get number of tilts in series
ReadLineToArray MTSPar TSNumber 
echo $TSNumber

## Initialize Angle, Montage image shift (MontIS_D) and last Montage Image shifts (LastMontIS_D)
## arrays
NewArray Angle -1 1
NewArray MontIS_D -1 3
NewArray LastMontIS_D -1 3 
echo lastmontISD $LastMontIS_D

# Get matrix to convert from camera values for image shift (which is what is inputted)
# to specimen coordinates (in microns)
GoToLowDoseArea R
BinningForCameraMatrix 1
CameraToSpecimenMatrix 0
echo  $ReportedValue1 $ReportedValue2 $ReportedValue3  $ReportedValue4

# 2 * is to account for the fact that serial-EM takes the "un-binned" pixel size of the K3 to be in super-res mode
XperX =  $ReportedValue1
XperY =  $ReportedValue2
YperX = $ReportedValue3
YperY =  $ReportedValue4

## Loop over tilts
Loop $TSNumber

   ## Reset image shift
   SetImageShift 0 0

   ## Read line from image shift text file (id MTSPar) to array Angle
   ReadLineToArray MTSPar Angle


   ## Write tilt angle to log window
   Echo Tilt to $Angle
   
   if $Angle > 0
      Angleminusone = $Angle - 1
      TiltTo $Angleminusone
   elseif $Angle < 0
      Angleminusone = $Angle + 1
      TiltTo $Angleminusone
   Endif

   TiltTo $Angle
   ResetImageShift

   If ABS $Angle > 0
        # Move stage to previously aligned x,y position
        If $Angle > 0
            MoveStageTo $PlusStagePos[1] $PlusStagePos[2]
        Else
            MoveStageTo $MinusStagePos[1] $MinusStagePos[2]
        EndIf

        ## Loop alignment procedure at view preset up to
        ## $LmtRATimes (3 by default)
        Loop $LmtRATimes
            ## Take view image
            View

            ## Align to image stored in buffer R (view image taken at previous tilt)
            If $Angle > 0
                AlignTo R
            Else
                Alignto S
            Endif
            ReportSpecimenShift 

            ## Magnitude of shift by Pythagoras theorem
            IS = $reportedValue1 * $reportedValue1 + $reportedValue2 * $reportedValue2

            ## If measured shift greater than threshhold reset image shift and stage shift
            ## to position, then align again. Otherwise break loop
            If $IS > $LmtISShift
                ResetImageShift
            Else
                Break 
            Endif 
        EndLoop
   Else
      # Realign to nav item for 0 degree tilt
      RealignToNavItem  1 1 $LmtISShift $LmtRATimes  3
   Endif

   View
   ## Copy View image to buffers R and S, R will be overwritten at each
   ## positive tilt and S will be overwritten at each negative tilt
   If $Angle > -1
      Copy A R
   Endif
   If $Angle < 1
      Copy A S
   Endif

   ## Get and store stage positions, positive and negative (tilt) stage
   ## positions will be stored seperately
   ReportStageXYZ
   If $Angle > -1
      PlusStagePos[1] = $ReportedValue1
      PlusStagePos[2] = $ReportedValue2
   Endif
   If $Angle < 1
      MinusStagePos[1] = $ReportedValue1
      MinusStagePos[2] = $ReportedValue2
   Endif

   
   ## 1 means beam centering using centroid - beam should be fully in image
   If $Beamcentering == 1
      Trial 
      CenterBeamFromImage
   Endif

   AutoFocus 

   #Read number of tiles to variable TileNumber
   ReadLineToArray MTSPar TileNumber
   Echo $TileNumber tiles for this tilt   

   ## Open output file
   output_file = $OFPrefix_$Angle.mrc
   OpenNewFile $output_file 

   LastMontIS_D[1] = 0
   LastMontIS_D[2] = 0
   LastMontIS_D[3] = 0

   # Loop over times
   Echo $TileNumber tiles for this tilt
   Loop $TileNumber indx

      ## Read image shift from file
      ReadLineToArray MTSPar MontIS_D

      ## Convert image shift from camera pixel units to specimen units
      ## by multiplying by appropriate matrix
      Xout = $MontIS_D[1] * $XperX + $MontIS_D[2] * $XperY
      Yout = $MontIS_D[1] * $YperX + $MontIS_D[2] * $YperY 

      ## Difference in image shift from last image shift vector
      IS_X = $Xout - $LastMontIS_D[1]
      IS_Y = $Yout - $LastMontIS_D[2]

      ## Defocus change
      DelDef = $MontIS_D[3] - $LastMontIS_D[3]

      ## Apply image shift
      ImageShiftByMicrons $IS_X $IS_Y 1 $ComaFree

      ## Apply defocus offset if necessary
      if $DelDef != 0
         ChangeFocus $DelDef
         echo change focus by $DelDef
      Endif

      ReportSpecimenShift

      ## Delay after image shift and then record and save image
      Delay $ISDelay
      Record
      Save 

      # Center beam
      If $Beamcentering == 1
         CenterBeamFromImage 1
      Endif

      # Store current image shift as last image shift for next loop
      # iteration
      LastMontIS_D[1] = $Xout
      LastMontIS_D[2] = $Yout
      LastMontIS_D[3] = $MontIS_D[3]
   
   # End loop over montage tiles
   EndLoop

   CloseFile
# End loop over tilts
EndLoop 

TiltTo 0
SetImageShift 0 0
