## Supplementary material for "Whole cryo-FIB lamella acquisition at high magnification with rectangular beam montage cryo-electron tomography": Source code: SetupPolygonMontage.txt

#!Python
#ScriptName	SetupPolygonMontage

import numpy as np
import os
import re
import json
import matplotlib.pyplot as plt
import serialem as sem


# ---------------------------------------------------------------------------
# beam_shape/rotation/overlap are no longer hand-copied here -- they are
# loaded at the bottom of this file from BeamGeometry.json, which
# DetermineOverlapFraction.txt writes next to ReferenceBeam.mrc after its
# sliders are closed (see load_beam_geometry and the __main__ block).
#   "square" - rectangular beam, tiled on a rectangular grid. `overlap` is
#              [overlapy, overlapx], the two reciprocal-style factors this
#              file has always used (spacing fraction = 1 - 1/overlap).
#   "round"  - circular beam, tiled on a hexagonal (triangular-lattice) grid.
#              `overlap` is a single number: the hexagon side length in
#              unbinned camera pixels, i.e. the centre-to-centre spacing
#              between beam positions.
# ---------------------------------------------------------------------------


def polygon_area(points):
    """
    Area of a closed simple polygon via the shoelace formula.

    Parameters:
    - points: An (N, 2) array of polygon vertices (ring not repeated).

    Returns:
    - The (unsigned) area, so the vertex winding direction does not matter.
    """
    x, y = np.asarray(points, dtype=float).T
    return 0.5 * np.abs(np.dot(x, np.roll(y, -1)) - np.dot(y, np.roll(x, -1)))


def clip_polygon_to_box(points, xmin, ymin, xmax, ymax):
    """
    Clip a polygon against an axis-aligned rectangle (Sutherland-Hodgman).

    The clip region is convex so the algorithm is exact. For a concave subject
    polygon the output can contain zero-width connecting edges running along the
    rectangle boundary, but those contribute no area to the shoelace formula, so
    the area of the result is still correct.

    Parameters:
    - points: An (N, 2) array of polygon vertices (ring not repeated).
    - xmin, ymin, xmax, ymax: The bounds of the clipping rectangle.

    Returns:
    - An (M, 2) array of clipped vertices, empty if there is no overlap.
    """
    poly = np.asarray(points, dtype=float)

    # Clip in turn against each of the four edges. keep_greater says which side
    # of the clip line is inside the rectangle.
    for axis, bound, keep_greater in (
        (0, xmin, True),
        (0, xmax, False),
        (1, ymin, True),
        (1, ymax, False),
    ):
        if len(poly) == 0:
            return np.empty((0, 2))

        # Signed distance to the clip line, positive on the inside
        d = (poly[:, axis] - bound) * (1.0 if keep_greater else -1.0)

        out = []
        for p, q, dp, dq in zip(poly, np.roll(poly, -1, axis=0), d, np.roll(d, -1)):
            # Keep vertices that are inside
            if dp >= 0:
                out.append(p)
            # Edge crosses the clip line, so add the crossing point
            if (dp > 0) != (dq > 0):
                out.append(p + (q - p) * (dp / (dp - dq)))
        poly = np.asarray(out) if out else np.empty((0, 2))

    return poly


def rect_polygon_overlap_area(polygon_points, xmin, ymin, xmax, ymax):
    """
    Area of intersection of a simple polygon with an axis-aligned rectangle.

    Replaces shapely's Polygon(...).intersection(box(...)).area using numpy only.
    """
    clipped = clip_polygon_to_box(polygon_points, xmin, ymin, xmax, ymax)
    if len(clipped) < 3:
        return 0.0
    return polygon_area(clipped)


def rectangles_with_overlap(
    polygon_points, rect_points, rect_width, rect_height, overlap_criterion
):
    """
    Function to find rectangles that overlap with a polygon by more than a given fractional area criterion.

    Parameters:
    - polygon_points: A list of points describing the polygon (list of tuples).
    - rect_points: A list of bottom-left corner points for each rectangle (list of tuples).
    - rect_width: The width of each rectangle.
    - rect_height: The height of each rectangle.
    - overlap_criterion: The fractional overlap criterion (between 0 and 1).

    Returns:
    - A list of bottom-left corner points of rectangles that have a fractional overlap greater than the criterion.
    """
    # List to hold rectangles that satisfy the overlap criterion
    overlapping_rectangles = []
    from matplotlib.patches import Rectangle

    #fig,ax = plt.subplots()
    #ax.plot(*np.asarray(polygon_points).T,'k--')
    # Loop through each rectangle
    for rect_point in rect_points:
        # Calculate the intersection area between the polygon and the rectangle
        # given by the bottom-left corner point and the given dimensions
        overlap_area = rect_polygon_overlap_area(
            polygon_points,
            rect_point[0],
            rect_point[1],
            rect_point[0] + rect_width,
            rect_point[1] + rect_height,
        )

        # Calculate the area of the rectangle
        rect_area = rect_width * rect_height

        # Calculate the fractional overlap
        fractional_overlap = overlap_area / rect_area
        # ax.add_patch(Rectangle(rect_point,rect_width,rect_height,ec='b',fc='none'))
        #ax.plot((rect_point[0],rect_point[0],rect_point[0] + rect_width,rect_point[0] + rect_width),(rect_point[1], rect_point[1] + rect_height, rect_point[1] + rect_height,rect_point[1]),'r-')

        # If the fractional overlap is greater than the overlap criterion, add the rectangle to the result
        if fractional_overlap > overlap_criterion:
            overlapping_rectangles.append(rect_point)
            # ax.plot((rect_point[0],rect_point[0],rect_point[0] + rect_width,rect_point[0] + rect_width),(rect_point[1], rect_point[1] + rect_height, rect_point[1] + rect_height,rect_point[1]),'b-')
    #plt.show(block=True)
    if len(overlapping_rectangles) < 1:
        raise ValueError(f"No acquisitions fit inside given polygon.")
    return np.asarray(overlapping_rectangles)


def extract_points(file_path, item_index):
    """
    Extracts PtsX and PtsY for a given Item index from a SerialEM nav file using string find method.

    Parameters:
    - file_path: path to the file
    - item_index: the index of the Item to extract

    Returns:
    - A numpy array of shape (N, 2) where N is the number of points, containing the PtsX and PtsY values.

    Raises:
    - ValueError: If PtsX or PtsY are missing for the specified item.
    """
    with open(file_path, "r") as file:
        content = file.read()

    # Locate the start of the [Item = item_index] section
    item_header = f"[Item = {item_index}]"
    item_start = content.find(item_header)

    if item_start == -1:
        raise ValueError(f"Item {item_index} not found in the file.")

    # Find the end of this item section (next empty line or new item)
    item_end = content.find("\n\n", item_start)
    if item_end == -1:
        item_end = len(content)

    # Extract the content of the item section
    item_content = content[item_start:item_end]

    # Find and extract the PtsX and PtsY values
    ptsx_start = item_content.find("PtsX = ")
    ptsy_start = item_content.find("PtsY = ")

    if ptsx_start == -1:
        raise ValueError(f"PtsX not found for Item {item_index}.")
    if ptsy_start == -1:
        raise ValueError(f"PtsY not found for Item {item_index}.")

    # Extract the lines with PtsX and PtsY values
    ptsx_start += len("PtsX = ")
    ptsy_start += len("PtsY = ")

    ptsx_end = item_content.find("|\n", ptsx_start)
    ptsy_end = item_content.find("|\n", ptsy_start)

    ptsx_str = item_content[ptsx_start:ptsx_end].strip()
    ptsy_str = item_content[ptsy_start:ptsy_end].strip()

    # Convert the space-separated values to lists of floats
    ptsx = list(map(float, ptsx_str.split()))
    ptsy = list(map(float, ptsy_str.split()))

    if len(ptsx) != len(ptsy):
        raise ValueError("Mismatch between the number of PtsX and PtsY values.")

    # Combine PtsX and PtsY into a NumPy array
    points_array = np.column_stack((ptsx, ptsy))

    return points_array


def load_beam_geometry(path):
    """
    Load beam_shape/rotation_deg/overlap from the BeamGeometry.json written
    by DetermineOverlapFraction.txt, so those values don't have to be
    hand-copied out of that script's slider titles into this one.

    Returns
    -------
    (beam_shape, rotation_deg, overlap) : tuple
    """
    if not os.path.exists(path):
        raise FileNotFoundError(
            "No BeamGeometry.json found at {0}. Run DetermineOverlapFraction.txt "
            "in this directory first -- it writes this file once its sliders "
            "are closed.".format(path)
        )
    with open(path) as f:
        geometry = json.load(f)
    return geometry["beam_shape"], geometry["rotation_deg"], geometry["overlap"]


def regular_polygon_points(radius, n_sides=28):
    """
    Corner points (relative to a center at the origin, closed loop) of a
    regular polygon approximating a circle of the given radius.

    SerialEM's navigator scripting has no native circle item -- only
    AddStagePointsAsPolygon, which draws a closed polygon from a point list
    -- so a many-sided regular polygon is the standard way to draw a
    round-beam footprint there; at typical navigator zoom levels it's
    visually indistinguishable from a true circle.
    """
    theta = np.linspace(0, 2 * np.pi, n_sides, endpoint=False)
    theta = np.append(theta, theta[0])  # repeat first point to close the loop
    return radius * np.cos(theta), radius * np.sin(theta)


def plot_points_in_serial_EM_navigator(imageshifts, M, nx, ny, x, y, beam_shape="square"):
    import serialem as sem
    # Get a unique group number to add squares to
    gid = int(sem.GetUniqueNavID())

    # Plot zero-tilt image shifts on navigator
    ids = []
    for imshift in imageshifts:
        xx = imshift[0]
        yy = imshift[1]

        if beam_shape == "round":
            # Largest circle inscribed in the nx x ny record footprint --
            # the same footprint already used elsewhere as the round beam's
            # effective per-tile area (see the overlap-filtering comment in
            # main_serialem/main_sandbox).
            relcornersx, relcornersy = regular_polygon_points(min(nx, ny) / 2)
        else:
            # Generate corners in pixel coordinates
            relcornersx = np.asarray([nx / 2 * (1 - 2 * ((i % 4) // 2)) for i in range(5)])
            relcornersy = np.asarray([ny / 2 * (1 - 2 * ((i + 1) % 4 // 2)) for i in range(5)])

        cornersx = xx + relcornersx
        cornersy = yy + relcornersy

        # Use matrix to convert to specimen coordinates
        cornersx, cornersy = M @ np.stack((cornersx, cornersy), axis=0)

        # Add "origin" (nav point coordinates) to
        cornersx += x
        cornersy += y

        # Write arrays back to serial-EM land
        sem.SetVariable("cornersx", "\n".join([str(a) for a in cornersx]))
        sem.SetVariable("cornersy", "\n".join([str(a) for a in cornersy]))

        # Add image shift acquisitions as polygons in display and group them
        ids.append(int(sem.AddStagePointsAsPolygon("cornersx", "cornersy", 0)))
    for id_ in ids:
        sem.ChangeItemGroupID(id_, gid)
    return ids, gid


def dose_symmetric_tilts(maxtilt, step, nreflect):
    """
    Generates a sequence of symmetric positive and negative tilt angles
    based on the given maximum tilt, step size, and reflection group size.

    The function first creates arrays of positive and negative tilts, then
    iteratively adds a set number (`nreflect`) of positive and negative tilts
    in an alternating pattern until all possible tilts are used.

    Parameters
    ----------
    maxtilt : float
        The maximum absolute tilt angle to generate (positive and negative).
    step : float
        The step size between consecutive tilt angles.
    nreflect : int
        The number of tilt angles to alternate between positive and negative
        in each reflection group.

    Returns
    -------
    tilts : list of float
        A list of tilt angles arranged in alternating positive and negative
        groups, symmetrically distributed.

    Example
    -------
    >>> dose_symmetric_tilts(10, 2, 3)
    [0, 2, 4, -2, -4, 6, 8, -6, -8]
    """

    # Create an array of positive tilt values ranging from 0 to (maxtilt).
    # step/10 max np.arange inclusive
    postilts = np.arange(0, maxtilt + step / 10, step)

    # Create an array of negative tilt values ranging from -step to -(maxtilt - step).
    negtilts = np.arange(-step, -maxtilt - step / 10, -step)

    # Determine the number of positive and negative tilt values generated.
    npos = postilts.shape[0]
    nneg = negtilts.shape[0]

    # Initialize an empty list to store the final tilt sequence.
    tilts = []

    # Loop through the tilt arrays, alternating between positive and negative groups.
    # The range ensures we loop enough times to cover all the tilts, based on nreflect.
    for i in range(int(np.ceil(npos / nreflect))):

        # Add a block of 'nreflect' positive tilt values to the tilt list.
        # The minimum ensures we don't exceed the number of positive tilts available.
        tilts += list(postilts[i * nreflect : min((i + 1) * nreflect, npos)])

        # Add a block of 'nreflect' negative tilt values to the tilt list.
        # The minimum ensures we don't exceed the number of negative tilts available.
        tilts += list(negtilts[i * nreflect : min((i + 1) * nreflect, nneg)])

    # Return the final list of alternating positive and negative tilt values.
    return tilts


def overlap_factor_to_sep(overlap_factor, detector_pixels, beam_shape="square"):
    """
    Convert the user-facing overlap parameter into fractional row/column
    tile-to-tile spacing (each as a fraction of one tile's y/x extent).

    For beam_shape="square", overlap_factor is [overlapy, overlapx], the
    reciprocal-style factors this file has always used: spacing fraction =
    1 - 1/overlap_factor.

    For beam_shape="round", overlap_factor is a single number: the hexagon
    side length in unbinned camera pixels (as reported by
    DetermineOverlapFraction.txt's round-beam control) -- the centre-to-centre
    tile spacing directly, rather than a reciprocal factor. Row (y) spacing
    is compressed by sqrt(3)/2 for triangular-lattice circle packing; the
    half-column row offset generate_montage_shifts already applies gives the
    required lattice offset in x.
    """
    if beam_shape == "round":
        hex_side_px = float(overlap_factor)
        return np.asarray([
            hex_side_px / detector_pixels[1] * np.sqrt(3) / 2,
            hex_side_px / detector_pixels[0],
        ])

    o = 1 / np.asarray(overlap_factor)
    return 1 - o


def generate_montage_shifts(
    overlap_factor, tiles, detector_pixels, shift=[0.0, 0.0], beam_shape="square"
):
    """
    Generates a list of montage shift coordinates to cover a large area using a grid
    of overlapping detector tiles. The generated coordinates represent shifts in pixel
    space required to position the montage tiles, accounting for the overlap between tiles
    and shifting the grid each tilt to avoid excessive dose on any particular region.

    Parameters
    ----------
    overlap_factor : list/tuple of float, or a single float
        For beam_shape="square": [overlapy, overlapx], the fractional overlap
        factors between adjacent tiles (see overlap_factor_to_sep). For
        beam_shape="round": a single number, the hexagon side length in
        unbinned camera pixels.
    tiles : tuple of int
        The number of tiles in the x (width) and y (height) directions (n, m).
    detector_pixels : list or tuple of int
        The dimensions of the detector in pixels (nx, ny).
    shift : list or tuple of float, optional
        An optional initial shift to apply to the montage coordinates in the x and y
        directions. Default is [0.0, 0.0].
    beam_shape : str, optional
        "square" (default) for a rectangular beam, or "round" for a circular
        beam. Both patterns already offset alternate rows by half the column
        spacing (see Notes); "round" additionally compresses the row-to-row
        spacing by sqrt(3)/2, turning that brick offset into the tighter
        triangular-lattice packing that covers a plane with circles using
        the least overlap. See load_beam_geometry and the __main__ block
        below for how this is selected.

    Returns
    -------
    coords : ndarray of shape (n_tiles, 2)
        An array of coordinates in pixel space, where each row corresponds to the shift
        needed for a particular tile in the montage. The x and y shifts are provided
        relative to the detector center.

    Notes
    -----
    - The function calculates the montage shifts such that the center of the grid is
      positioned at (0, 0), with each tile shifted by an amount determined by the overlap
      factor.
    - Alternate rows are offset by half the column spacing (a brick pattern) regardless
      of beam_shape; for beam_shape="round" this is exactly the triangular-lattice
      offset once the row spacing is also compressed below.
    - The x coordinates reverse direction on each row (serpentine or zigzag pattern) to
      minimize the magnitude of successive shifts.
    - The y coordinates are calculated to ensure coverage based on the overlap and tile
      dimensions.

    Example
    -------
    >>> generate_montage_shifts([0.1, 0.1], (3, 3), [1024, 1024])
    array([[ -512.,  -512.],
           [  512.,  -512.],
           [ 1536.,  -512.],
           [ 1536.,     0.],
           [  512.,     0.],
           [ -512.,     0.],
           [ -512.,   512.],
           [  512.,   512.],
           [ 1536.,   512.]])
    """

    # Get number of tiles in x and y direction
    n, m = tiles

    # Fractional seperation of individual images (see overlap_factor_to_sep
    # for how this differs between beam_shape="square" and "round")
    sep = overlap_factor_to_sep(overlap_factor, detector_pixels, beam_shape=beam_shape)

    # Equivalent fractional overlap, used below to size the tail of each row/column
    o = 1 - sep

    # Generate a list of fractional y coordinates for the montage centers
    # centered on 0. (shift+0.5)%1.0)+0.5 ensures that we always start within
    # half a field of view of the desired area, m+0.5 ensures that we always
    # cover the desired area. -m/2 centers the coordinates on zero, the step size
    # is 1 - the overlap fraction
    M = (
        np.arange(
            sep[0] / 2 - ((shift[1] + sep[0] / 2) % sep[0]), m + sep[0] / 2, sep[0]
        )
        - m / 2
    )  # + 0.5*((m+1)%2)

    # N +=
    coords = []
    for iim, im in enumerate(M):
        # Generate a list of fractional x coordinates for the montage centers
        # centered on 0. (shift+0.5)%1.0)+0.5 ensures that we always start within
        # half a field of view of the desired area
        halfshift = sep[1] * (iim % 2)/2
        
        start =  ((shift[1] + halfshift  + sep[1]) % sep[1]) - 0.5
        stop = n + 0.5 - o[1]
        N = (
            np.arange(start,stop,sep[1]
            )
            - n / 2
        )
        # Reverse direction of x coordinate every row to minimize magnitude of
        # successive image shifts
        for iin in N[:: 1 - 2 * int(iim % 2)]:
            # Each row must be shifted either left or right by 1/4 of the x
            # dimension
            coords += [[iin, im]]
    
    # import sys;sys.exit()
    # Return the result in units of detector pixels
    return np.asarray(coords) * np.asarray(detector_pixels)


def estimate_montage_time(
    imageshifts, record_time, delaytime=0.5, measured_overhead=1.7, newtilt_overhead=60.0
):
    """
    ESTIMATE ONLY of total montage acquisition time, via

        T = sum_tilts n_tilt * (Rtime + delaytime + measured_overhead) + newtilt_overhead

    where n_tilt is the number of Record acquisitions at that tilt (taken
    from len(imageshifts[i])).

    Parameters
    ----------
    imageshifts : list of ndarray
        Per-tilt image shift arrays, as built in main_serialem/main_sandbox;
        n_tilt for each tilt is len(imageshifts[i]).
    record_time : float
        The Record preset exposure time in seconds (e.g.
        sem.ReportExposure('R')[0]).
    delaytime : float, optional
        Extra per-shot delay set by the user elsewhere. Default 0.5 s.
    measured_overhead : float, optional
        Fixed per-shot overhead measured on this microscope; will vary
        microscope to microscope. Default 1.7 s.
    newtilt_overhead : float, optional
        Overhead per tilt for going to a new tilt angle, tracking/aligning
        and autofocusing. Default 60 s.

    Returns
    -------
    total_seconds : float
        The estimated total acquisition time in seconds.
    """
    total_seconds = 0.0
    for ishifts in imageshifts:
        n_tilt = len(ishifts)
        total_seconds += n_tilt * (record_time + delaytime + measured_overhead) + newtilt_overhead

    hours, remainder = divmod(total_seconds, 3600)
    minutes, seconds = divmod(remainder, 60)
    print(
        "ESTIMATE ONLY (actual run time will vary): predicted montage "
        "acquisition time is approximately {0:d}h {1:02d}m {2:02d}s "
        "({3:.0f} s total)".format(int(hours), int(minutes), int(seconds), total_seconds)
    )
    return total_seconds


def write_tilts_and_image_shifts_to_file(filename, tilts, imageshifts, zs):

    f = open(filename, "w", encoding="utf-8")
    f.write("{0:d}|\n".format(len(tilts)))

    for tilt, ishifts, z in zip(tilts, imageshifts, zs):
        f.write("{0}|\n{1}|\n".format(tilt, len(ishifts)))
        np.savetxt(
            f,
            np.concatenate((ishifts, z.reshape((z.shape[0], 1))), axis=-1),
            fmt="%5.3f",
            newline="|\n",
        )

    f.close()


def calculate_defocii(imageshifts, angles, M):

    # Generate tilt axis vector
    zs = []

    for alpha, imshift in zip(angles, imageshifts):
        # Use camera to specimen transformation matrix to calculate
        # component of image shift

        zs.append(np.tan(np.deg2rad(alpha)) * (M @ imshift.T).T[:, 1])
    return zs

def plot_tilts_and_image_shifts(polygonpoints,tilts,imageshifts,M,ny,nx):
    fig = plt.figure()  
    camerapoints = np.asarray([[0,0],[nx,0],[nx,ny],[0,ny],[0,0]]) @ M.T
    ax = fig.add_subplot(projection='3d')
    
    threedpolypoints = np.concatenate((polygonpoints,polygonpoints[0:1,:]),axis=0)
    Z = np.zeros(threedpolypoints.shape[0])
    threedpolypoints = np.concatenate((threedpolypoints,Z.reshape((Z.shape[0],1))),axis=1)
    ax.plot3D(*(threedpolypoints.T), c='b',alpha=0.5)
    for i, (tilt, imagexy) in enumerate(zip(tilts, imageshifts)):
        if tilt<0:
            continue
        points = imagexy @ M.T
        print((points.reshape((points.shape[0], 1,points.shape[1])) + camerapoints).shape)

        for shift in points.reshape((points.shape[0], 1,points.shape[1])) + camerapoints:
            Z = shift[...,1]*np.tan(np.deg2rad(tilt))
            ax.plot3D(*(np.concatenate([shift,Z.reshape((Z.shape[0],1))],axis=1)).T, 'r-')
        # plt.show()
        # ax.plot(imshift[:, 0], imshift[:, 1], "o", label=f"Tilt: {tilt}")
        # for j, (x, y) in enumerate(imshift):
        #     ax.text(x, y, f"{j}", fontsize=8)
        # for j, (x, y) in enumerate(imshift):    
    plt.show()

def main_serialem(step,maxtilt,ntiltgroup,overlap,outdir,polyoverlapfrac,rotation=0,beam_shape="square",delaytime=0.5):
    import serialem as sem

    # Get coordinates from polygon and label from last acquired item
    # this script is intended by run after acquisition using "Acquire at items"
    # in serial-EM so the nav item will be the polygon
    item_index, x, y , z, label = sem.ReportNavItem()

    label = sem.GetVariable("navLabel").strip()

    # No way to directly get polygon points in serialEM so the following hack will
    # have to surfice...
    tempnavfile = os.path.join(outdir, "Temp.nav")
    sem.SaveNavigator(tempnavfile)
    polygonpoints = extract_points(tempnavfile, label)
    os.remove(tempnavfile)

    # Get the stage position and label from the last acquired image which should be the view map.
    _________, vx, vy , vz,  label = sem.ReportOtherItem(-1)
    label = sem.GetVariable("navLabel").strip()

    # Generate output filename using navigation item albel
    fnamout = "Montage_imageshifts_{0}.txt".format(label)
    fnamout = os.path.join(outdir, fnamout)

    # Get information about record preset (pixels and pixel size)
    sem.GoToLowDoseArea("R")
    nx, ny, rotflip, p, up, camnum = sem.CameraProperties()

    # Get camera to specimen matrix to convert between camera pixel
    # measurements and coordinates on the specimen
    # 2 is to adjust for the fact that "binning 1" for the K3 is really superres
    # see https://groups.google.com/a/colorado.edu/g/serialem/c/uoEtxqA4xRA
    # delete if not using a K2 or K3
    sem.BinningForCameraMatrix(1)
    M = np.asarray(sem.CameraToStageMatrix(0)).reshape((2,2))
    print(M)
    # M = np.asarray(sem.CameraToStageMatrix(0)).reshape((2, 2))
    Minv = np.linalg.inv(M)
    # Convert pixel size to micron
    p = p * 1e-4

    # Allows the user to specify a rotation of the montage to match a rotated
    # beam
    Mrot = rotation_matrix(rotation)
    Mrotinv = np.linalg.inv(Mrot)

    # Generate tilt series
    tilts = dose_symmetric_tilts(maxtilt, step, ntiltgroup)
    print(
        "{0} tilts between {1} and {2} in steps of {3}".format(
            len(tilts), min(tilts), max(tilts), step
        )
    )

    # Get size (range) and origin of polygon to fit in initial guess of montage
    # tiles
    transformed_poly_points = (Minv @ polygonpoints.T).T
    extent = np.ptp((Mrotinv @ transformed_poly_points.T).T, axis=0)
    origin = (
        np.max(transformed_poly_points, axis=0)
        + np.min(transformed_poly_points, axis=0)
    ) / 2

    ntilts = len(tilts)

    # Fractional row/column tile spacing, used below to give each tilt a
    # different sub-tile phase so the same ground isn't dosed on every tilt
    # (see overlap_factor_to_sep for the square vs. round difference).
    sep = overlap_factor_to_sep(overlap, [nx, ny], beam_shape=beam_shape)

    imageshifts = []
    for itilts in range(ntilts):

        # Generate montage shifts for this tilt, shift by overlap
        # output is in pixels
        imagexy = generate_montage_shifts(
            overlap,
            [extent[0] / nx, extent[1] / ny],
            [nx, ny],
            shift=[(itilts * (1 - sep[0])) % 1.0, (itilts * (1 - sep[1])) % 1.0],
            beam_shape=beam_shape,
        )

        # Shift image shifts by 1/2 a Record field of view
        fov = np.asarray((nx, ny))
        imagexy -= fov / 2

        # Rotate image shifts
        imagexy = imagexy @ Mrot.T

        # Dilate the polygon by the cosine of the tilt angle to account for
        # the fact that the field of view is larger at higher tilts
        dilated_poly_points = polygonpoints.copy()
        dilated_poly_points = (dilated_poly_points[:,1] - vy)*np.cos(np.deg2rad(tilts[itilts])) + vy
        transformed_poly_points = (Minv @ polygonpoints.T).T
        # dilated_poly_points[:,1] = (transformed_poly_points[:,1] - vy)*np.cos(np.deg2rad(tilts[itilts])) + vy

        # Remove images with insufficient overlap with the polygon. For
        # beam_shape="round" this treats each tile's footprint as the nx x ny
        # record rectangle rather than the true circular beam outline -- an
        # approximation, but nx x ny is already the useful (inscribed-square)
        # data extracted per tile downstream, so it is the right footprint
        # to test against.
        imagexy = (
            rectangles_with_overlap(
                transformed_poly_points, imagexy + origin, nx, ny, polyoverlapfrac
            )
            - origin + fov/2
        )

        imageshifts.append(imagexy)

    ids, gid = plot_points_in_serial_EM_navigator(imageshifts[0], M, nx, ny, vx, vy, beam_shape=beam_shape)

    # Estimate total acquisition time (see estimate_montage_time docstring
    # for the formula and caveats -- this is an estimate only)
    record_time = sem.ReportExposure("R")[0]
    estimate_montage_time(imageshifts, record_time, delaytime=delaytime)

    # Calculate defocii to compensate for distance from tilt axis
    zs = calculate_defocii(imageshifts,tilts,M)

    # Save image shifts for Serial-EM in units of pixels in the camera
    # basis (x and y aligned with camera axes). Imageshifts will be 
    # converted to basis aligned with tilt axis inside SerialEM
    write_tilts_and_image_shifts_to_file(fnamout,tilts,imageshifts,zs)
    print('Save {0}'.format(fnamout))

def rotation_matrix(angle_degrees):
    """
    Generates a 2D rotation matrix for a given angle in degrees.

    Parameters:
    - angle_degrees: The rotation angle in degrees.

    Returns:
    - A 2x2 NumPy array representing the rotation matrix.
    """
    angle_radians = np.deg2rad(angle_degrees)
    cos_theta = np.cos(angle_radians)
    sin_theta = np.sin(angle_radians)
    return np.array([[cos_theta, -sin_theta], [sin_theta, cos_theta]])

def main_sandbox(step,maxtilt,ntiltgroup,overlap,outdir,polyoverlapfrac,polygonpoints=None,nx=4092,ny=4092,p=0.65,rotation=0,beam_shape="square"):

    # Get coordinates from polygon and label from last acquired item
    # this script is intended by run after acquisition using "Acquire at items"
    # in serial-EM so the nav item will be the polygon
    if polygonpoints is None:
        polygonpoints = np.asarray([[-1.0,0.0],[-1.0,1.0],[-0.75,1.0],[-0.75,0.75],[0.75,0.75],[0.75,1.0],[1.0,1.0],[1.0,0.0]])
    
    vx,vy = np.ptp(polygonpoints,axis=0)
    

    # Generate output filename using navigation item albel
    fnamout = "Montage_imageshifts.txt"
    fnamout = os.path.join(outdir, fnamout)

    # Convert pixel size to micron
    pmicron = p * 1e-4

    # Generate tilt series
    tilts = dose_symmetric_tilts(maxtilt, step, ntiltgroup)
    print(
        "{0} tilts between {1} and {2} in steps of {3}".format(
            len(tilts), min(tilts), max(tilts), step
        )
    )
    Mrot = rotation_matrix(rotation)
    Mrotinv = np.linalg.inv(Mrot)
    M = np.eye(2)*pmicron
    Minv = np.linalg.inv(M)

    # Get size (range) and origin of polygon to fit in initial guess of montage
    # tiles
    transformed_poly_points = ( Minv @ polygonpoints.T).T
    extent = np.ptp((Mrotinv @ transformed_poly_points.T).T, axis=0)
    origin = (
        np.max(transformed_poly_points, axis=0)
        + np.min(transformed_poly_points, axis=0)
    ) / 2

    # Generate corners in pixel coordinates
    cornersx = np.asarray([nx / 2 * (1 - 2 * ((i % 4) // 2)) for i in range(5)])
    cornersy = np.asarray([ny / 2 * (1 - 2 * ((i + 1) % 4 // 2)) for i in range(5)])
    corners = np.stack((cornersx, cornersy), axis=1)

    beamcorners = (Mrot @ corners.T).T 
    print(beamcorners,np.ptp(beamcorners,axis=0)/np.asarray((nx,ny)))
    # sys.exit()
    beamcorners /= np.amax(np.ptp(beamcorners,axis=0))/np.asarray((nx,ny))


        
    ntilts = len(tilts)

    # Fractional row/column tile spacing, used below to give each tilt a
    # different sub-tile phase so the same ground isn't dosed on every tilt
    # (see overlap_factor_to_sep for the square vs. round difference).
    sep = overlap_factor_to_sep(overlap, [nx, ny], beam_shape=beam_shape)

    imageshifts = []
    for itilts in range(ntilts):

        # Generate montage shifts for this tilt, shift by overlap
        # output is in pixels
        imagexy = generate_montage_shifts(
            overlap,
            [extent[0] / nx, extent[1] / ny],
            [nx, ny],
            shift=[(itilts * (1 - sep[0])) % 1.0, (itilts * (1 - sep[1])) % 1.0],
            beam_shape=beam_shape,
        )

        # Shift image shifts by 1/2 a Record field of view
        fov = np.asarray((nx, ny))
        imagexy -= fov / 2

        # Rotate image shifts
        imagexy = imagexy @ Mrot.T

        # Dilate the polygon by the cosine of the tilt angle to account for
        # the fact that the field of view is larger at higher tilts
        dilated_transformed_points = transformed_poly_points.copy()
        dilated_transformed_points[:,1] = (transformed_poly_points[:,1] - vy)*np.cos(np.deg2rad(tilts[itilts])) + vy
        
        
        # Remove images with insufficient overlap with the polygon
        imagexy = (
            rectangles_with_overlap(
                dilated_transformed_points, imagexy + origin, nx, ny, polyoverlapfrac
            )
            - origin + fov/2
        )

       
        imageshifts.append(imagexy)
    plot_tilts_and_image_shifts(polygonpoints,tilts,imageshifts,M,ny,nx)


    # Calculate defocii to compensate for distance from tilt axis
    zs = calculate_defocii(imageshifts,tilts,M)

    # Save image shifts for Serial-EM in units of pixels in the camera
    # basis (x and y aligned with camera axes). Imageshifts will be 
    # converted to basis aligned with tilt axis inside SerialEM
    write_tilts_and_image_shifts_to_file(fnamout,tilts,imageshifts,zs)
    
    print('Save {0}'.format(fnamout))

if __name__ == "__main__":

    # Parse commandline arguments
    step = 3
    maxtilt = 60
    ntiltgroup = 3

    outdir = sem.ReportDirectory()
    polyoverlapfrac = 0.2
    delaytime = 0.5  # extra per-shot delay set elsewhere; keep in sync with that script

    # beam_shape, rotation and overlap come from DetermineOverlapFraction.txt's
    # BeamGeometry.json, written next to ReferenceBeam.mrc in this same
    # directory -- see load_beam_geometry.
    BEAM_SHAPE, rotation, overlap = load_beam_geometry(os.path.join(outdir, "BeamGeometry.json"))

    main_serialem(step,maxtilt,ntiltgroup,overlap,outdir,polyoverlapfrac,rotation=rotation,beam_shape=BEAM_SHAPE,delaytime=delaytime)
    #main_sandbox(step,maxtilt,ntiltgroup,overlap,outdir,polyoverlapfrac,rotation=rotation,beam_shape=BEAM_SHAPE)
    
