## Supplementary material for "Whole cryo-FIB lamella acquisition at high magnification with rectangular beam montage cryo-electron tomography": Source code: 01_motion_correction.sh

```
#!/bin/bash
#SBATCH --job-name=mc2_montage
#SBATCH --output=logs/mc2_%A_%a.out
#SBATCH --error=logs/mc2_%A_%a.err
#SBATCH -p gpu-l40s
#SBATCH --mem=32G
#SBATCH --time=00:30:00
#SBATCH --gres=gpu:1
#SBATCH --array=0-EDIT_ME    # set upper bound to (number of TIFFs - 1)
#
# Motion-correct one multi-frame TIFF per array task.
# Run beam_mask_motioncorr --print-slurm to auto-generate this script
# with the array size and paths filled in for your dataset.
#
# ── edit these ────────────────────────────────────────────────────────────────
FRAMES_DIR="EDIT_ME"          # directory containing raw multi-frame TIFFs
TIFF_GLOB="*.tif"             # Edit to target only tifs following a certain pattern
BASE_DIR="EDIT_ME"            # Base directory for all processing outputs
OUTPUT_DIR="${BASE_DIR}/motion_corrected" # where to write motion-corrected MRCs
PIXEL_SIZE=EDIT_ME            # pixel size in Å
VOLTAGE=EDIT_ME                 # acceleration voltage in kV, for CTF estimation/phase-flipping
CS=EDIT_ME                      # spherical aberration in mm, for CTF estimation/phase-flipping
AMP_CONTRAST=0.07           # amplitude contrast fraction, for CTF estimation/phase-flipping
ROTATION=0                 # rotation of the input frames in degrees, so that it matches
                           # Serial-EM's preferred camera orientation
SPLIT_FRAMES=0              # 1 = also write odd/even half-sums (for cryoCARE etc), 0 = skip
BATCH_SIZE=1                 # TIFFs per array task. 1 matches the #SBATCH --array bound above
                             # (one task per TIFF); raise it to cover more TIFFs with fewer
                             # array tasks -- e.g. 280 for 4475 TIFFs over --array=0-15.
START_OFFSET=0               # first TIFF index this array covers; 0 unless resuming a partial run

# Beam-detection threshold for the mask used to find the crop fed to
# MotionCor2, as a fraction of each tile's own image median -- matching
# choose_mask_params' live "x median" readout exactly, so a value confirmed
# there transfers here per-tile unchanged, regardless of dose/detector mode.
# NOT the same mask/tool as Choose_fringe_size.py (that one tunes stitch.py's
# LATER fringe-removal mask, a different step). Tune with choose_mask_params
# on a real frame from FRAMES_DIR before submitting. Prefer this fraction
# over the older --mask-threshold (absolute counts): a fixed absolute count
# from one dataset does not transfer to another shot at a different dose --
# e.g. single-electron counting frames can have a median under 1 count, where
# an absolute threshold tuned for an integrating-mode dataset leaves the mask
# empty and beam_mask_motioncorr fails with "largest inscribed square is only
# 0px".
MASK_THRESHOLD_FRACTION=EDIT_ME
MASK_SHRINK=20
OVERWRITE=1                  # 0 = --no-overwrite, skip tiles whose .stack_markers
                             # marker already exists -- set to 0 and resubmit
                             # (chained with --dependency=afterany:<jobid> on the
                             # timed-out job) to pick up where an array that hit
                             # its --time limit left off, without redoing tiles
                             # that already finished.

MOTIONCOR2=/programs/x86_64-linux/motioncor2/1.6.4/MotionCor2_1.6.4_Cuda121_Mar312023
CTFFIND4=/programs/x86_64-linux/ctffind4/4.1.14/ctffind4/bin/ctffind4
# ── end of edit section ───────────────────────────────────────────────────────

source /programs/sbgrid.shrc
module load CUDA/12.2.0

set -eo pipefail
mkdir -p "${OUTPUT_DIR}/logs"

# Each task writes a provenance sidecar per output tile as it goes -- they are
# on by default, so there is no path to configure and nothing to remember. That
# replaces the manifest, which needed a shared file named consistently across
# every step plus a dependent 01b job to fold the array's spool into it. Both
# of those failed silently on Montage_13A_new: the name had a typo and the
# fold never ran. Pass --no-sidecar to skip them.

# Map tiff files to all different slurm jobs
mapfile -t TIFFS < <(ls "${FRAMES_DIR}"/${TIFF_GLOB} | sort)
BATCH_START=$(( START_OFFSET + SLURM_ARRAY_TASK_ID * BATCH_SIZE ))
INPUTS=("${TIFFS[@]:$BATCH_START:$BATCH_SIZE}")

# Pass the split-frames argument if asked for
SPLIT_FRAMES_ARGS=()
[ "${SPLIT_FRAMES}" = "1" ] && SPLIT_FRAMES_ARGS+=(--split-frames)

OVERWRITE_ARGS=()
[ "${OVERWRITE}" = "0" ] && OVERWRITE_ARGS+=(--no-overwrite)

# Run beam_mask_motioncorr on the files
beam_mask_motioncorr \
    --input         "${INPUTS[@]}" \
    --output-dir    "${OUTPUT_DIR}" \
    --pixel-size    ${PIXEL_SIZE} \
    --motioncor2    ${MOTIONCOR2} \
    --ctffind4      ${CTFFIND4} \
    --voltage       ${VOLTAGE} \
    --cs            ${CS} \
    --amp-contrast  ${AMP_CONTRAST} \
    --rotate ${ROTATION} \
    --mask-threshold-fraction ${MASK_THRESHOLD_FRACTION} \
    --mask-shrink ${MASK_SHRINK} \
    "${SPLIT_FRAMES_ARGS[@]}" \
    "${OVERWRITE_ARGS[@]}" \
    --stack-output \
    --save-diagnostic diagnostics \
    --gpu 0


my-job-stats -a -n -s
```
