## Supplementary material for "Whole cryo-FIB lamella acquisition at high magnification with rectangular beam montage cryo-electron tomography": Source code: 02_stitch.sh

```
#!/bin/bash
#SBATCH --job-name=stitch_montage
#SBATCH --output=/home/%u/logs/%x_%A_%a.out
#SBATCH --error=/home/%u/logs/%x_%A_%a.out
#SBATCH -p cascade,sapphire
#SBATCH --array=0-19
#SBATCH --mem=128G
#SBATCH --time=04:00:00
#SBATCH --cpus-per-task=16
#
# Stitch per-tile motion-corrected MRCs into a single tilt series MRC stack.
# stitch_square_beam writes every discovered tilt straight into its own
# z-slice of STITCHED_FILE (one shared, flock-guarded MRC stack) since all
# tilts share one canvas size, so no separate crop/join step is needed.
#
# ── edit these ────────────────────────────────────────────────────────────────
BASE_DIR="EDIT_ME"            # Base directory for all processing outputs
INPUT_DIR="${BASE_DIR}/motion_corrected" # directory of per-tile MRCs either from motion_correction
                              # or fresh off the TEM (ie. summed)
IMAGE_SHIFTS="EDIT_ME"        # Imageshifts.txt from Serial-EM
STITCHED_FILE="${BASE_DIR}/EDIT_ME.mrc"  # output .mrc file for per-tilt stitched images
N_THREADS=8                   # match --cpus-per-task above, the program tries
                              # to use all available cpus otherwise
BINNING=2                     # binning of the input tiles
INPUT_FRAME=motion-corrected  # what MOTIONCORR_DIR holds. Set to 'raw' when
                              # stitching uncorrected tiles summed tiles raw
                              # from the microscope
MARK_UNCOVERED=true           # true if manual masking/inpainting (03_inpaint_apply.sh)
                              # follows; false to skip marking gap sentinels
STITCH_ODD_EVEN=0             # 1 = also stitch odd/even motion-corrected half-sums
                              #     (requires SPLIT_FRAMES=1 in 01_motion_correction.sh)
ODD_INPUT_DIR="${INPUT_DIR}/odd"    # odd half-sums written by motion correction
EVEN_INPUT_DIR="${INPUT_DIR}/even"  # even half-sums written by motion correction
STITCHED_ODD_FILE="${STITCHED_FILE%.mrc}_odd.mrc"
STITCHED_EVEN_FILE="${STITCHED_FILE%.mrc}_even.mrc"
# ── end of edit section ───────────────────────────────────────────────────────

set -eo pipefail

# Ensure STITCHED_FILE's destination exists before stitch_square_beam runs.

STITCH_TARGET="${STITCHED_FILE}"
if [[ "${STITCH_TARGET%:*}" == *.mrc && "${STITCH_TARGET##*:}" =~ ^-?[0-9]+$ ]]; then
    STITCH_TARGET="${STITCH_TARGET%:*}"  # drop the ':N' slice-index suffix
fi
case "${STITCH_TARGET,,}" in
    *.mrc|*.tif|*.tiff)
        mkdir -p "$(dirname "${STITCH_TARGET}")"
        ;;
    *)
        mkdir -p "${STITCH_TARGET}"
        ;;
esac

MARK_UNCOVERED_FLAG=()
if [ "${MARK_UNCOVERED}" = true ]; then
    MARK_UNCOVERED_FLAG=(--mark-uncovered)
fi

# Arguments shared by all three stitch_square_beam calls -- only -i, -o and
# -s differ between them. 
common_args=(
    -I "${IMAGE_SHIFTS}"
    --input-frame ${INPUT_FRAME}
    "${MARK_UNCOVERED_FLAG[@]}"
    -nt ${N_THREADS}
    -b ${BINNING}
    --correct_beam_edges
    --tilt-index "$SLURM_ARRAY_TASK_ID"
    --consensus lstsq
    -mt ${MASK_THRESHOLD_FRACTION}
    -f ${MASK_SHRINK}
    --min_mean_intensity 0.1
)

# Run stitching
stitch_square_beam \
    -i "${INPUT_DIR}" \
    -o "${STITCHED_FILE}" \
    "${common_args[@]}"


# Re-stitch the dose-split motion-corrected half-sums with the alignment found
# above. -s reuses each tilt's saved positions, tile selection and beam masks,
# so odd/even stacks differ from the full-dose stack only in dose-split noise.
# The half-set variants write their own sidecars beside their own output
# stacks, so they neither need nor can collide with the full-dose ones.
if [ "${STITCH_ODD_EVEN}" = "1" ]; then
    stitch_square_beam \
        -i "${ODD_INPUT_DIR}" \
        -o "${STITCHED_ODD_FILE}" \
        "${common_args[@]}" \
        -s

    stitch_square_beam \
        -i "${EVEN_INPUT_DIR}" \
        -o "${STITCHED_EVEN_FILE}" \
        "${common_args[@]}" \
        -s
fi

my-job-stats -a -n -s
```
