## Supplementary material for "Whole cryo-FIB lamella acquisition at high magnification with rectangular beam montage cryo-electron tomography": Source code: 03_inpaint_apply.sh

```
#!/bin/bash
#SBATCH --job-name=inpaint
#SBATCH --output=logs/inpaint_%j.out
#SBATCH --error=logs/inpaint_%j.err
#SBATCH -p cascade,sapphire
#SBATCH --mem=64G
#SBATCH --time=02:00:00
#SBATCH --cpus-per-task=16
#
# Apply pre-fitted inpainting parameters to a montage tilt series MRC.
# The params JSON is generated interactively by running:
#
#   mask_and_inpaint <input.mrc>
#
# then clicking "Save SLURM Job" in the GUI, which writes both the params JSON
# and a ready-to-submit SLURM script automatically.  This template is provided
# as a reference; the GUI-generated script is usually easier to use.
#
# ── edit these ────────────────────────────────────────────────────────────────
INPUT="EDIT_ME"               # stitched MRC tilt series (output of 02_stitch)
OUTPUT="EDIT_ME"              # output path for inpainted MRC
PARAMS="EDIT_ME"              # per-tilt (lo, hi) JSON written by mask_and_inpaint GUI
N_TOTAL=16                    # total thread budget (match --cpus-per-task above)
NPERTILT=4                    # DCT threads per tilt; N_TOTAL // NPERTILT tilts run concurrently
BIN=2                         # downsample factor for the smoothn fit (1 disables binning)
# ── end of edit section ───────────────────────────────────────────────────────

source /programs/sbgrid.shrc

mkdir -p logs

inpaint_apply \
    --input    "${INPUT}" \
    --output   "${OUTPUT}" \
    --params   "${PARAMS}" \
    --ntotal   ${N_TOTAL} \
    --npertilt ${NPERTILT} \
    --bin      ${BIN}

my-job-stats -a -n -s
```
