## Supplementary material for "Whole cryo-FIB lamella acquisition at high magnification with rectangular beam montage cryo-electron tomography": Source code: 04_AreTomo.sh

```
#!/bin/bash
#SBATCH --job-name=AreTomo
#SBATCH --output=logs/aretomo_%j.out
#SBATCH --error=logs/aretomo_%j.out
#SBATCH -p gpu-a100-short
#SBATCH --mem=100G
#SBATCH --time=2:00:00
#SBATCH --gres=gpu:1
#
# Reconstruct full-dose (and, if configured, odd/even half-dose) tomograms
# with AreTomo.
#
# AreTomo's raw output has axis order (x, z, y). IMOD's `clip rotx`
# (single-threaded, memory-bound, no GPU) reorders it to (x, y, z) -- it is
# offloaded to a CPU-only job, 04b_clip_rotx.sh, queued automatically below,
# so it does not sit on this job's GPU allocation. Recording into the
# provenance (via montage_aretomo_record) also happens there, once each volume
# is in its final rotated form: recording it here, before rotation, would
# either record the wrong axis order/z_sign or need a second, duplicate
# record once the rotation lands -- see aretomo_record.py's docstring on
# `--output`/output_pending.
#
# ── edit these ────────────────────────────────────────────────────────────────
BASE_DIR="EDIT_ME"                              # Base directory for all processing outputs
OUTPUT_DIR="${BASE_DIR}/AreTomo"
FULL_TILT_SERIES="${BASE_DIR}/stitched/EDIT_ME" # full-dose stitched tilt series
EVEN_TILT_SERIES="${BASE_DIR}/stitched/EDIT_ME" # even half-dose, if denoising -- leave as EDIT_ME to skip
ODD_TILT_SERIES="${BASE_DIR}/stitched/EDIT_ME"  # odd half-dose, if denoising -- leave as EDIT_ME to skip
MASK_FILE="${BASE_DIR}/stitched/EDIT_ME"
ANGLE_FILE="${BASE_DIR}/stitched/EDIT_ME"       # tiltAngles.txt written by stitching
ARETOMO="/home/hgbrown/gitprojects/AreTomo2/AreTomo2" # path to AreTomo executable
PATCHX=5        # x patches for AreTomo
PATCHY=5        # y patches for AreTomo
VOLZ=2000       # Z volume (unbinned pixels)
OUTBIN=1        # AreTomo's own -OutBin
PIXEL_SIZE="EDIT_ME"  # pixel size of the stitched tilt series -- account for stitch_square_beam's --binning
AUTO_ROTX=1     # queue 04b_clip_rotx.sh to follow this job. Set to 0 to
                # submit it by hand instead (see the note near the bottom).
# ── end of edit section ───────────────────────────────────────────────────────

module load foss/2022a CUDA/11.8.0 UCX-CUDA/1.13.1-CUDA-11.8.0 cuDNN/8.7.0.84-CUDA-11.8.0
module load GCC/11.3.0  OpenMPI/4.1.4

mkdir -p "${OUTPUT_DIR}" logs

# AreTomo names the .aln after -InMrC but writes it into the -OutMrC
# directory. Derive it rather than typing it: the two do not have to match,
# and when they do not, the file the script looks for is never written.
ALN="${OUTPUT_DIR}/$(basename "${FULL_TILT_SERIES%.mrc}").aln"

# Common arguments in an array, so montage_aretomo_record (in 04b, after
# rotation) is given the *same* command line AreTomo ran rather than an
# approximation of it. AreTomo itself is not wrapped: it runs exactly as
# before and keeps its own exit code.
common=(-Patch ${PATCHX} ${PATCHY} -MaskFile "${MASK_FILE}" -VolZ ${VOLZ}
        -TiltCor 1 -AngFile "${ANGLE_FILE}" -OutBin ${OUTBIN}
        -PixSize ${PIXEL_SIZE})

# Each variant's *unrotated* AreTomo output and its final (post-`clip rotx`)
# home get appended to ROTX_SPEC as an add_variant call. 04b sources this
# file with its own add_variant defined -- one that actually rotates and
# records, rather than just collecting -- so the two scripts hand off
# structured per-variant data without needing a third format. Job-ID-suffixed
# so a later re-run cannot collide with one still queued.
ROTX_SPEC="${OUTPUT_DIR}/.rotx_pending_${SLURM_JOB_ID}.sh"
: > "${ROTX_SPEC}"
{
    printf 'ALN=%q\n' "${ALN}"
    printf 'ARETOMO_BINARY=%q\n' "${ARETOMO}"
} >> "${ROTX_SPEC}"

# add_variant NAME UNROTATED FINAL -- ARGV...
add_variant() {
    local name="$1" unrotated="$2" final="$3"; shift 3
    [ "$1" = "--" ] && shift
    {
        printf 'add_variant %q %q %q --' "${name}" "${unrotated}" "${final}"
        printf ' %q' "$@"
        printf '\n'
    } >> "${ROTX_SPEC}"
}

# 1. Full dose. This run solves the alignment and writes $ALN; the frame the
#    tomogram lives in is declared from its sidecar, written in 04b.
full_unrotated="${OUTPUT_DIR}/recon_patch_${PATCHX}_${PATCHY}_mask.mrc"
full_final="${OUTPUT_DIR}/recon__${PATCHX}_${PATCHY}__rotx.mrc"
full_args=(-InMrC "${FULL_TILT_SERIES}" -OutMrC "${full_unrotated}" "${common[@]}")
${ARETOMO} "${full_args[@]}"
add_variant full "${full_unrotated}" "${full_final}" -- "${full_args[@]}"

# 2-3. Half sets, reusing that alignment with -AlnFile so all three volumes
#      sit in the same frame. montage_aretomo_record (in 04b) marks these
#      'supplied' and checks the file against the one run 1 wrote: a
#      re-solved alignment paired with a stale half set is otherwise
#      invisible until the halves disagree. Skipped if left as EDIT_ME.
if [[ "${EVEN_TILT_SERIES}" != *EDIT_ME* && "${ODD_TILT_SERIES}" != *EDIT_ME* ]]; then
    even_unrotated="${OUTPUT_DIR}/recon_patch_${PATCHX}_${PATCHY}_mask_even.mrc"
    even_final="${OUTPUT_DIR}/recon_patch_${PATCHX}_${PATCHY}_mask_even_rotx.mrc"
    even_args=(-InMrC "${EVEN_TILT_SERIES}" -OutMrC "${even_unrotated}"
               "${common[@]}" -AlnFile "${ALN}")
    ${ARETOMO} "${even_args[@]}"
    add_variant even "${even_unrotated}" "${even_final}" -- "${even_args[@]}"

    odd_unrotated="${OUTPUT_DIR}/recon_patch_${PATCHX}_${PATCHY}_mask_odd.mrc"
    odd_final="${OUTPUT_DIR}/recon_patch_${PATCHX}_${PATCHY}_mask_odd_rotx.mrc"
    odd_args=(-InMrC "${ODD_TILT_SERIES}" -OutMrC "${odd_unrotated}"
              "${common[@]}" -AlnFile "${ALN}")
    ${ARETOMO} "${odd_args[@]}"
    add_variant odd "${odd_unrotated}" "${odd_final}" -- "${odd_args[@]}"
else
    echo "EVEN_TILT_SERIES/ODD_TILT_SERIES left as EDIT_ME -- skipping half-set reconstruction."
fi

# ── queue clip rotx + sidecar recording ─────────────────────────────────────
# afterany, not afterok: if a later variant's AreTomo run fails, the earlier
# ones that did complete are still worth rotating and recording -- same
# rationale as any array whose later tasks can fail independently.
echo "ROTX_SPEC: ${ROTX_SPEC}"
if [[ "${AUTO_ROTX}" == "1" ]]; then
    if command -v sbatch >/dev/null 2>&1; then
        ROTX_JOB=$(sbatch --parsable \
            --dependency=afterany:"${SLURM_JOB_ID}" \
            --chdir="${SLURM_SUBMIT_DIR}" \
            --export=ALL,ROTX_SPEC="${ROTX_SPEC}" \
            "${SLURM_SUBMIT_DIR}/04b_clip_rotx.sh")
        echo "Queued 04b_clip_rotx.sh as job ${ROTX_JOB}, to run after ${SLURM_JOB_ID}."
    else
        echo "WARNING: no sbatch on this node; run 04b_clip_rotx.sh by hand, e.g.:" >&2
        echo "  ROTX_SPEC=${ROTX_SPEC} sbatch 04b_clip_rotx.sh" >&2
    fi
fi

# With AUTO_ROTX=0 this is the manual two-step submission:
#
#   ARETOMO_JOB=$(sbatch --parsable 04_AreTomo.sh)
#   sbatch --dependency=afterany:${ARETOMO_JOB} \
#       --export=ALL,ROTX_SPEC=<the path this job echoed above> 04b_clip_rotx.sh
#
# ROTX_SPEC's path is deterministic (OUTPUT_DIR/.rotx_pending_<job ID>.sh),
# so it can be computed ahead of submission if scripting this.

my-job-stats -a -n -s
```
