## Supplementary material for "Whole cryo-FIB lamella acquisition at high magnification with rectangular beam montage cryo-electron tomography": Source code: 11_montage_to_relion.sh

```
#!/bin/bash
#SBATCH --job-name=montage2relion
#SBATCH --output=/home/hgbrown/logs/montage2relion_%A.out
#SBATCH --error=/home/hgbrown/logs/montage2relion_%A.out
#SBATCH -p cascade,sapphire
#SBATCH --ntasks=1
#SBATCH --cpus-per-task=4
#SBATCH --mem=64G
#SBATCH --time=02:00:00
# set -x
#
# Turn 3D picks made in a montage tomogram into a RELION-5 tomogram +
# particle set, one "tilt image" per (tilt, tile).
#
# STAGE 1 validates the coordinate chain and STAGE 2 exports it. Run stage 1
# first and LOOK AT THE PNGs. The exporter will happily write a well-formed
# star file full of noise if a sign convention is wrong.
#
# ── edit these ────────────────────────────────────────────────────────────────
BASE=/home/hgbrown/20240917_Montage_stitching/20260713Yeastattempt2

# The picks, as the IMOD model straight out of 3dmod. It is converted to the
# text form every stage reads by model2point below, so the model stays the
# master copy and there is no hand-made .txt to go stale against it. A .txt
# here still works and is passed through untouched.
PICKS="${BASE}/AreTomo_motioncorr/Ribosomes.mod"
PICKS="${BASE}/AreTomo_motioncorr/Points_for_reprojection_check.mod"

# TOMOGRAM AND ALN MUST BE THE SAME RUN.
#
# The picks are 3D coordinates in one specific volume, and the .aln is the only
# description of where that volume sits. Pairing a volume with a different run's
# alignment is not a small error: the local (patch) field differs between runs,
# so the reprojection is wrong by the whole field, position-dependently, which
# is precisely what cost the 191 A of job 28886886.
#
# Two consistent pairs exist here. The -Patch 5 3 one is the volume these check
# points were picked in (2026-08-07), so it is the pair to verify against; the
# archived .aln is the alignment that run solved, kept when 03_AreTomo_motioncorr
# installed the newer one.
#   -Patch 5 3   recon_patch53mask_bin2.mrc    Montage_9-A_inpainted_20260805-1627.aln
#   -Patch 10 6  recon_patch10x6mask_bin2.mrc  Montage_9-A_inpainted.aln
# Switching to the 10x6 volume means re-picking in it first.
TOMOGRAM="${BASE}/AreTomo_motioncorr/recon_patch53mask_bin2.mrc"
ALN="${BASE}/AreTomo_motioncorr/Montage_9-A_inpainted_20260805-1627.aln"

POSITIONS_DIR="${BASE}/stitched_motioncorr"
TILE_DIR="${BASE}/motion_corrected"
CTF_RESULTS="${TILE_DIR}/ctf_results.txt"
CTF_BASE="Montage_9-A_9-A"

# Geometry now comes from the provenance sidecars written beside each output,
# read automatically by walking back from TOMOGRAM: its own sidecar names the
# stack it was reconstructed from, that stack's sidecar carries the ROI, tile
# pixel size, stitch binning and -- the one that used to be guessed -- the
# rotation stitch actually applied. Nothing downstream restates 02_stitch.sh
# from memory and there is no path to configure.
#
# That anchoring is also what enforces the warning above: TOMOGRAM's sidecar
# names ITS aln and ITS input, so a -Patch 5 3 volume can no longer be paired
# with a -Patch 10 6 run's geometry by editing one variable and not the other.
#
# For a series processed before sidecars existed, reconstruct them once:
#   montage_backfill ${BASE}
# and pass --set for anything only the submission scripts know. Add
# --no-sidecars below to ignore them and use the hard-coded defaults instead.

ALIGNED_STACK="${BASE}/AreTomo_motioncorr/recon_patch53mask_bin2tiltseries.mrc"
CANVAS_STACK="${POSITIONS_DIR}/Montage_9-A.mrc"

OVERLAY_DIR="${BASE}/relion/overlays"
OUT_DIR="${BASE}/relion/tomo"

BOX=256          # extraction box, unbinned tile pixels (256 x 3.426 A = 877 A)
DOSE_PER_TILT=0  # e-/A^2 per exposure. 0 disables dose weighting -- set it!
# 1 = overlay, 2 = star files, 3 = imod models, 4 = measure reprojection residual.
# Edit the default here; `STAGE=4 sbatch ...` overrides it for one run.
STAGE="${STAGE:-3}"
echo "STAGE=${STAGE}"

# Which tilts get a per-tile model (stage 3). Empty = the section nearest 0 deg,
# which is the one tilt where an error perpendicular to the tilt axis is
# multiplied by sin(theta) = 0 and therefore invisible -- so ask for the
# extremes too. "--tilt all" writes one model per tilt.
TILT_ARGS="--tilt -57 --tilt -24 --tilt 0 --tilt 24 --tilt 60"
# STAGE="${STAGE:-1}"
# ── end of edit section ───────────────────────────────────────────────────────

# Passed to every stage. Empty by default: the sidecars supply the geometry,
# and anything set here OVERRIDES them, which is occasionally what you want
# for a test and never what you want by accident.
GEOM_ARGS=""

# NOTE: the `Stitch` conda env is currently broken -- ~/.conda/envs/Stitch has
# only conda-meta/ and etc/, no bin/, so `conda activate Stitch` yields no
# python. That breaks 02_stitch.sh and 03_inpaint_apply.sh too and wants
# rebuilding. Until then run out of the repo with the base Anaconda3, which has
# numpy/scipy/h5py/mrcfile/matplotlib and is all these two tools need.
module load Anaconda3
REPO=/home/hgbrown/gitprojects/Square_beam_montaging
# Take the interpreter from the module root, not from PATH. `python` on PATH is
# whatever env happens to be first -- the IsoNet2 build env here, which has no
# h5py -- and stage 3 sources sbgrid for point2model, which puts python 2.7.2
# in front of everything and turns any annotated signature into a SyntaxError.
PY="${EBROOTANACONDA3}/bin/python"
cd "${REPO}"

set -eo pipefail
export MPLBACKEND=Agg
unset DISPLAY

mkdir -p "${OVERLAY_DIR}" "${OUT_DIR}"

# Picks: .mod -> .txt, once, before any stage runs.
#
# -float because load_picks() takes fractional coordinates and the default
# integer rounding would throw away up to half a voxel (~7 A at bin 2). No
# -object/-contour (the loader wants three columns; it would take the last
# three anyway) and no -zcoord, which shifts Z by -0.5. That combination was
# checked to reproduce the hand-made Ribosomes.txt exactly. IMOD 4.11 prints
# "the -float entry is no longer needed" -- it is float by default there; the
# flag is kept for whichever IMOD sbgrid supplies.
#
# The output is named for the tool that made it so it cannot be confused with,
# or overwrite, a .txt written by hand from an older version of the model.
if [ "${PICKS##*.}" = "mod" ]; then
    PICKS_MOD="${PICKS}"
    PICKS="${PICKS_MOD%.mod}_model2point.txt"
    if ! command -v model2point >/dev/null 2>&1; then
        # IMOD is on PATH interactively (~/Software/IMOD). A batch job that
        # inherited a thinner environment gets it from sbgrid instead; see the
        # stage 3 note for why the source is wrapped in `set +e`. PY is already
        # an absolute path, so sbgrid's python 2.7.2 landing in front of
        # everything does not matter here.
        set +e
        source /programs/sbgrid.shrc
        set -e
    fi
    if ! command -v model2point >/dev/null 2>&1; then
        echo "model2point not on PATH; cannot convert ${PICKS_MOD}" >&2
        exit 1
    fi
    model2point -float "${PICKS_MOD}" "${PICKS}"
    echo "picks: ${PICKS_MOD} -> ${PICKS}  ($(wc -l < "${PICKS}") points)"
fi

if [ "${STAGE}" = "1" ]; then
    # Stage 1: does the chain land the picks on density?
    #
    # Three views, each isolating one link: `aligned` tests only the
    # tomogram->aligned shear, `canvas` adds the inverse alignment (and settles
    # the 180 deg that footprint-based rotation detection cannot), `patches`
    # tests the whole thing by cutting boxes out of the raw tiles.
    #
    # The number to read off is the mean patch's |peak|/rms. Below ~2 the
    # geometry is wrong (or the picks are not real) and stage 2 is pointless.
    # If it reports a residual offset, feed that back as --extra-shift.
    ${PY} -m processing_scripts.overlay_picks_on_montage \
        --picks          "${PICKS}" \
        --tomogram       "${TOMOGRAM}" \
        --aln            "${ALN}" \
        --positions-dir  "${POSITIONS_DIR}" \
        --tile-dir       "${TILE_DIR}" \
        --aligned-stack  "${ALIGNED_STACK}" \
        --canvas-stack   "${CANVAS_STACK}" \
        --mode all \
        --box  ${BOX} \
        ${GEOM_ARGS} \
        -o     "${OVERLAY_DIR}"
elif [ "${STAGE}" = "3" ]; then
    # Stage 3: IMOD models, to check the chain against the images in 3dmod.
    #
    #   picks_tiltseries_canvas.mod  one contour per particle, one point per
    #       tilt, on the stitched montage. Select a contour and step through Z:
    #       the point should stay on the same feature. Drift that grows with
    #       |tilt| is the Z/shear term, a constant offset is TX/TY.
    #   picks_tiles_<tilt>.mod       the raw per-tilt stack, where each Z
    #       section is a TILE. Green = the tile each particle was assigned to,
    #       red = boxes rejected for crossing the square-beam edge (they should
    #       form a band around each tile's illuminated square).
    #
    # Model Z is 0-based, matching the image index; 3dmod's slider counts from 1.
    # sbgrid.shrc ends with `let SB_RUNTIME=endtime-starttime`, and `let`
    # returns status 1 when the expression evaluates to 0. Sourcing it in under
    # a second therefore makes `source` return 1, and under `set -e` the job
    # dies right there with an empty log. It is a race against the clock, so it
    # fails intermittently -- job 28842562 passed, 28843292 did not, same script.
    set +e
    source /programs/sbgrid.shrc   # for point2model
    set -e
    ${PY} -m processing_scripts.picks_to_imod_model \
        --picks             "${PICKS}" \
        --tomogram          "${TOMOGRAM}" \
        --aln               "${ALN}" \
        --positions-dir     "${POSITIONS_DIR}" \
        --tile-dir          "${TILE_DIR}" \
        --frame             canvas \
        --tiltseries-image  "${CANVAS_STACK}" \
        --box               ${BOX} \
        --mode both \
        ${TILT_ARGS} \
        ${GEOM_ARGS} \
        -o "${OVERLAY_DIR}"
elif [ "${STAGE}" = "4" ]; then
    # Stage 4: measure the reprojection residual and fit it.
    #
    # For when stage 3 shows the picks drifting perpendicular to the tilt axis.
    # Cross-correlates a patch cut at each pick's PREDICTED position against the
    # same pick at the reference tilt, then fits the residual against the terms
    # that could cause it -- a constant, sin(theta) (z centre), cos(theta)-1
    # (x centre) and dx*sin(theta) (tilt angle). The fit names the culprit
    # instead of leaving it to be guessed.
    #
    # Use picks on obvious high-contrast features; ribosomes are too small and
    # too crowded to cross-correlate one at a time.
    # Run it BOTH ways, in one job, so the comparison is like-for-like.
    #
    # `nolocal` reproduces the old behaviour: global ROT/TX/TY only, ignoring
    # the .aln's patch table. That is the configuration that measured 14.0
    # voxels rms = 191 A perpendicular in job 28886886, so it is the control.
    # `local` applies AreTomo's patch field the way the reconstruction did.
    # If the diagnosis is right the perpendicular residual falls in `local`
    # and `nolocal` reproduces ~191 A; if both are unchanged, the patch field
    # was not the cause and the sin(theta) z-centre term is next.
    for mode in nolocal local; do
        extra=""
        [ "${mode}" = "nolocal" ] && extra="--no-local"
        echo
        echo "──────── reprojection residual: ${mode} ────────"
        mkdir -p "${OVERLAY_DIR}/residual_${mode}"
        ${PY} -m processing_scripts.measure_reprojection_residual \
            --picks          "${PICKS}" \
            --tomogram       "${TOMOGRAM}" \
            --aln            "${ALN}" \
            --positions-dir  "${POSITIONS_DIR}" \
            --tile-dir       "${TILE_DIR}" \
            --canvas-stack   "${CANVAS_STACK}" \
            ${GEOM_ARGS} ${extra} \
            -o "${OVERLAY_DIR}/residual_${mode}"
    done
else
    # Stage 2: write the star files.
    #
    # --extra-shift comes from stage 1. --handedness flips the sign of the
    # along-beam (defocus) axis. The CTF fits CANNOT settle it here: the
    # acquisition script already flattens the defocus across each montage
    # (ChangeFocus per tile), so there is no geometric ramp in the data whose
    # sign could be read. Refine both ways and keep the better.
    ${PY} -m processing_scripts.montage_to_relion \
        --picks          "${PICKS}" \
        --tomogram       "${TOMOGRAM}" \
        --aln            "${ALN}" \
        --positions-dir  "${POSITIONS_DIR}" \
        --tile-dir       "${TILE_DIR}" \
        --ctf-results    "${CTF_RESULTS}" \
        --ctf-base       "${CTF_BASE}" \
        --box            ${BOX} \
        --dose-per-tilt  ${DOSE_PER_TILT} \
        --extra-shift    0 0 \
        --handedness     1 \
        ${GEOM_ARGS} \
        -o "${OUT_DIR}"

    echo
    echo "Then, with RELION 5 on PATH (source /programs/sbgrid.shrc):"
    echo "  relion_tomo_subtomo --i ${OUT_DIR}/optimisation_set.star \\"
    echo "      --b ${BOX} --crop $((BOX/2)) --bin 2 --j 8 --o Subtomograms/"
fi

my-job-stats -a -n -s
```
