## Supplementary material for "Whole cryo-FIB lamella acquisition at high magnification with rectangular beam montage cryo-electron tomography": Source code: 12_subtomo_validate.sh

```
#!/bin/bash
#SBATCH --job-name=subtomo_val
#SBATCH --output=/home/hgbrown/logs/subtomo_val_%A.out
#SBATCH --error=/home/hgbrown/logs/subtomo_val_%A.out
#SBATCH -p cascade,sapphire
#SBATCH --ntasks=1
#SBATCH --cpus-per-task=8
# relion_tomo_subtomo loads the WHOLE tilt series into memory before extracting,
# and one image per (tilt, tile) makes that 642 x 4092 x 5760 x 4 B = 60.5 GB of
# image data alone. 64 G was OOM-killed at 67 GB resident (job 28884970).
#SBATCH --mem=256G
#SBATCH --time=04:00:00
#
# Is the pseudo-subtomogram extraction actually pulling particles out?
#
# The test is NOT "does the sum look like a blob". With unknown orientations the
# sum of 128 randomly-oriented ribosomes is a featureless ball either way, and
# crowded cytoplasm averages to a ball too. So the same extraction is run twice:
# once on the picks, once on DECOY positions displaced 750 A sideways within the
# same tiles -- same dose, same defocus, same tiles, differing in one respect
# only, that they are not on a particle. The real sum has to beat the decoy.
#
# Read off: peak/rms of the two sums, and the radial profile. A 300 A ribosome
# at 6.85 A/px is ~44 px across; anything much bigger is not a particle.
#
# ── edit these ────────────────────────────────────────────────────────────────
export BASE=/home/hgbrown/20240917_Montage_stitching/20260713Yeastattempt2
TOMO_DIR="${BASE}/relion/tomo"            # from 11_montage_to_relion.sh STAGE=2
OUT="${BASE}/relion/validate"

# Box arithmetic, and it matters. `--box` in 11_montage_to_relion.sh is in
# UNBINNED tile pixels and decides where rlnTomoVisibleFrames is set. RELION's
# `--b` is in BINNED pixels, so it reads `b * bin` unbinned pixels per particle.
# Keep b*bin <= box or RELION reads past the region we declared visible -- off
# the tile, or across the square-beam edge, which is inpainted.
#   BOX 256 unbinned (877 A) -> --b 128 --bin 2 reads exactly 256.
BIN=2
B=128
# Do NOT crop tight. At 7 um the CTF delocalises signal by lambda*df*s = 69 A at
# 20 A and 138 A at 10 A, so a 300 A ribosome needs 300 + 2*delocalisation of
# box to keep its own signal. 128 * 6.852 = 877 A; 96 (658 A) is already marginal.
CROP=128
JITTER=750       # decoy displacement, A, in the specimen plane
# ── end of edit section ───────────────────────────────────────────────────────

module load Anaconda3
REPO=/home/hgbrown/gitprojects/Square_beam_montaging
PY="${EBROOTANACONDA3}/bin/python"
cd "${REPO}"

set -eo pipefail
mkdir -p "${OUT}"

# sbgrid.shrc ends with `let SB_RUNTIME=end-start`, and `let` returns 1 when the
# result is 0, so sourcing it in under a second kills the job under `set -e`.
set +e
source /programs/sbgrid.shrc
set -e

for SET in real decoy; do
    if [ "${SET}" = "real" ]; then
        SRC="${TOMO_DIR}"
    else
        SRC="${OUT}/decoy_tomo"
        # Rebuild the whole export from displaced picks rather than editing the
        # star file: the decoys must go through the same tile assignment and
        # visibility test, or the comparison is not like for like.
        ${PY} - "${JITTER}" <<'EOF'
import sys, numpy as np, os
base = os.environ["BASE"]
picks = np.loadtxt(f"{base}/AreTomo_motioncorr/Ribosomes.txt")
rng = np.random.default_rng(0)
d = float(sys.argv[1]) / (3.426 * 4)          # A -> reconstruction voxels
ang = rng.uniform(0, 2 * np.pi, len(picks))
picks[:, 0] += d * np.cos(ang)
picks[:, 1] += d * np.sin(ang)
os.makedirs(f"{base}/relion/validate", exist_ok=True)
np.savetxt(f"{base}/relion/validate/Ribosomes_decoy.txt", picks, fmt="%12.2f")
EOF
        ${PY} -m processing_scripts.montage_to_relion \
            --picks          "${OUT}/Ribosomes_decoy.txt" \
            --tomogram       "${BASE}/AreTomo_motioncorr/recon_patch53mask_bin2.mrc" \
            --aln            "${BASE}/AreTomo_motioncorr/Montage_9-A_inpainted.aln" \
            --positions-dir  "${BASE}/stitched_motioncorr" \
            --tile-dir       "${BASE}/motion_corrected" \
            --ctf-results    "${BASE}/motion_corrected/ctf_results.txt" \
            --ctf-base       Montage_9-A_9-A \
            --box 256 --dose-per-tilt 0 --extra-shift 0 0 --handedness 1 \
            --tomo-name recon_patch53mask_bin2 \
            -o "${SRC}"
    fi

    echo "=============== ${SET} ==============="
    mkdir -p "${OUT}/${SET}"
    # RELION resolves rlnTomoTomogramsFile / rlnTomoTiltSeriesStarFile against
    # the CURRENT DIRECTORY, not against the star file that names them, so it
    # has to be run from the export directory. Everything else is absolute.
    #
    # Expect "has relion-4 definition of projection matrices; converting them
    # now". That is RELION spotting that our matrices take a corner-origin
    # coordinate (which projection_matrix builds deliberately) and shifting them
    # to the RELION-5 centred convention. It should be the right conversion, but
    # it is asserted rather than verified -- the decoy comparison below is what
    # would catch it if it is not.
    ( cd "${SRC}" && relion_tomo_subtomo \
        --i    optimisation_set.star \
        --b    ${B} --crop ${CROP} --bin ${BIN} \
        --j    ${SLURM_CPUS_PER_TASK} \
        --sum  --no_writing \
        --o    "${OUT}/${SET}/" )
done

# Score both sums the same way and print the comparison.
${PY} -m processing_scripts.score_subtomo_sum \
    --real  "${OUT}/real"  --decoy "${OUT}/decoy" \
    --pixel-size $(${PY} -c "print(3.426*${BIN})") \
    -o "${OUT}"

my-job-stats -a -n -s
```
