## Supplementary material for "Whole cryo-FIB lamella acquisition at high magnification with rectangular beam montage cryo-electron tomography": Source code: test_stitch.py

```
import unittest
import numpy as np
from stitch import (
    make_gain_ref,
    stitch,
    generate_montage_shifts,
    dose_symmetric_tilts,
    parse_image_shifts,
    parse_commandline,
    main,
)

class TestStitch(unittest.TestCase):

    def test_make_gain_ref(self):
        # Test the make_gain_ref function
        images = [np.random.rand(100, 100) for _ in range(5)]
        gain_ref = make_gain_ref(images, binning=2)
        self.assertEqual(gain_ref.shape, (50, 50))

    def test_stitch(self):
        # Test the stitch function
        ims = [np.random.rand(100, 100) for _ in range(4)]
        positions = np.array([[0, 0], [0, 100], [100, 0], [100, 100]])
        msks = [np.ones((100, 100), dtype=bool) for _ in range(4)]
        pixel_size = 1.0
        result = stitch(ims, positions, msks, pixel_size)
        self.assertEqual(result.shape, (200, 200))

    def test_generate_montage_shifts(self):
        # Test the generate_montage_shifts function
        overlap_factor = [0.1, 0.1]
        tiles = (3, 3)
        detector_pixels = [1024, 1024]
        shifts = generate_montage_shifts(overlap_factor, tiles, detector_pixels)
        self.assertEqual(shifts.shape, (9, 2))

    def test_dose_symmetric_tilts(self):
        # Test the dose_symmetric_tilts function
        tilts = dose_symmetric_tilts(10, 2, 3)
        self.assertEqual(len(tilts), 9)

    def test_parse_image_shifts(self):
        # Test the parse_image_shifts function
        image_shifts = parse_image_shifts("test_image_shifts.txt")
        self.assertEqual(len(image_shifts), 3)

    def test_parse_commandline(self):
        # Test the parse_commandline function
        args = parse_commandline()
        self.assertIn("input", args)
        self.assertIn("output", args)

if __name__ == "__main__":
    unittest.main()
```
